# The Developmental Transcriptome for *Lytechinus variegatus* Exhibits Temporally Punctuated Gene Expression Changes

**DOI:** 10.1101/572388

**Authors:** John D. Hogan, Jessica L. Keenan, Lingqi Luo, Dakota Y. Hawkins, Jonas Ibn-Salem, Arjun Lamba, Daphne Schatzberg, Michael L. Piacentino, Daniel T. Zuch, Amanda B. Core, Carolyn Blumberg, Bernd Timmermann, José Horacio Grau, Emily Speranza, Miguel A. Andrade-Narravo, Naoki Irie, Albert J. Poustka, Cynthia A. Bradham

**Author notes:** co-first authors (John D. Hogan, Jessica L. Keenan, Lingqi Luo).

## Abstract

Embryonic development is arguably the most complex process an organism undergoes during its lifetime, and understanding this complexity is best approached with a systems-level perspective. The sea urchin has become a highly valuable model organism for understanding developmental specification, morphogenesis, and evolution. As a non-chordate deuterostome, the sea urchin occupies an important evolutionary niche between protostomes and vertebrates. *Lytechinus variegatus* (Lv) is an Atlantic species that has been well studied, and which has provided important insights into signal transduction, patterning, and morphogenetic changes during embryonic and larval development. The Pacific species, *Strongylocentrotus purpuratus* (Sp), is another well-studied sea urchin, particularly for gene regulatory networks (GRNs) and *cis*-regulatory analyses. A well-annotated genome and transcriptome for Sp are available, but similar resources have not been developed for Lv. Here, we provide an analysis of the Lv transcriptome at 11 timepoints during embryonic and larval development. The data indicate that the gene regulatory networks that underlie specification are well-conserved among sea urchin species. We show that the major transitions in variation of embryonic transcription divide the developmental time series into four distinct, temporally sequential phases. Our work shows that sea urchin development occurs via sequential intervals of relatively stable gene expression states that are punctuated by abrupt transitions.

## Introduction

Although only about half of the gene complement of sea urchins is expressed by the developing embryo, approximately 90% of the signaling ligands, kinases, small GTPases, and transcription factors are expressed during development (Beane et al., 2006b; Bradham et al., 2006; Croce et al., 2006b; Howard-Ashby et al., 2006a; Howard-Ashby et al., 2006b; Lapraz et al., 2006; Materna et al., 2006; Samanta et al., 2006; Sodergren et al., 2006; Walton et al., 2006). The embryonic utilization of the vast majority of its signaling and transcriptional regulatory gene repertoire highlights the intrinsic complexity at the basis of development, which in turn underscores the value and importance of systems-level perspectives for understanding developmental mechanisms.

There is a rich history of investigation in sea urchin embryos that provides a wealth of knowledge regarding the anatomical and cellular changes that accompany sea urchin embryogenesis (Driesch, 1892; Horstadius, 1935, 1939; Gustafson and Wolpert, 1961a, b; Wolpert and Gustafson, 1961; Gustafson and Wolpert, 1967; Horstadius, 1973). Work in more recent decades has uncovered many of the important signals and transcription factors that drive specification and development in these embryos (Logan et al., 1999; Sherwood and McClay, 1999; Angerer et al., 2000; Angerer et al., 2001; Sweet et al., 2002; Oliveri et al., 2003; Bradham et al., 2004; Duboc et al., 2004; Rottinger et al., 2004; Wikramanayake et al., 2004; Duboc et al., 2005; Bradham and McClay, 2006; Oliveri et al., 2006; Duloquin et al., 2007; Duboc et al., 2008; Rottinger et al., 2008; Yaguchi et al., 2008; Bradham et al., 2009; Lapraz et al., 2009; Sethi et al., 2009; Walton et al., 2009; Wei et al., 2009; Yaguchi et al., 2010; Luo and Su, 2012; Sethi et al., 2012; Materna et al., 2013b; McIntyre et al., 2013; Range et al., 2013; Krupke and Burke, 2014; Khadka et al., 2018). This work has culminated in gene regulatory network (GRN) models that describe the specification of the endomesoderm and the ectoderm (Davidson et al., 2002; Su et al., 2009; Saudemont et al., 2010; Peter and Davidson, 2011; Rafiq et al., 2012; Materna et al., 2013a; Li et al., 2014), and more recently, efforts have been made to connect the specification networks to morphogenesis (Annunziata and Arnone, 2014; Saunders and McClay, 2014; Martik and McClay, 2015). Sea urchins are nonchordate deuterostomes, and as such, they occupy an important evolutionary niche between protostomes and vertebrates. The availability of GRN models and global sequence resources has enabled evolutionary, comparative, and population studies at the molecular and network level (Hinman et al., 2003; Gao and Davidson, 2008; Garfield et al., 2013; Wygoda et al., 2014; Erkenbrack et al., 2018).

With the advent of systems biology, the sea urchin has emerged as an important developmental model for global analyses. The sea urchin larva is relatively simple morphologically, since it possesses relatively few cell types and lacks complex organs and structures (Angerer and Angerer, 2003). *Ex vivo* fertilization allows for the routine collection of synchronously developing, large cultures of embryos, providing sample sizes appropriate for systems-level measurements. The sea urchin genome has not undergone a duplication, and lacks the extensive redundancy found in vertebrates (Bradham et al., 2006; Howard-Ashby et al., 2006a; Lapraz et al., 2006; Materna et al., 2006; Sodergren et al., 2006), although the small GTPases and Wnt genes are present at comparable numbers in the sea urchin and human genomes (Beane et al., 2006b; Croce et al., 2006b). However, among signaling proteins and transcription factor genes in general, sea urchins possess the diversity of vertebrate genomes without the complexity engendered by genetic redundancy (Bradham et al., 2006; Howard-Ashby et al., 2006b; Lapraz et al., 2006; Sodergren et al., 2006). For example, although sea urchins possess approximately 30% fewer kinase genes than humans, sea urchins lack only four of the 186 kinase subfamilies found in humans (Bradham et al., 2006; Sodergren et al., 2006).

*Lytechinus variegatus* (Lv) is a well-studied Atlantic sea urchin species, and many important insights in signal transduction, patterning, and morphogenesis have been obtained from Lv (Armstrong et al., 1993; Ettensohn and Malinda, 1993; Ruffins and Ettensohn, 1996; Guss and Ettensohn, 1997; Logan et al., 1999; Sherwood and McClay, 1999; Sweet et al., 2002; Beane et al., 2006a; Bradham and McClay, 2006; Croce et al., 2006a; Ettensohn et al., 2007; Wu and McClay, 2007; Bradham et al., 2009; Walton et al., 2009; McIntyre et al., 2013; Saunders and McClay, 2014; Martik and McClay, 2015; Piacentino et al., 2015; Schatzberg et al., 2015; Piacentino et al., 2016a; Piacentino et al., 2016b). Investigations in *Strongylocentrotus purpuratus* (Sp), a well-studied Pacific sea urchin species, have been particularly important for GRN and *cis*-regulatory analyses; the latter in particular depend on interspecies comparisons, which have often been made between Sp and Lv (Wei et al., 1995; Xu et al., 1996; Yuh et al., 2001; Yuh et al., 2002; Oliveri et al., 2003; Revilla-i-Domingo et al., 2004; Yuh et al., 2004; Minokawa et al., 2005; Ransick and Davidson, 2006; Lee et al., 2007; Livi and Davidson, 2007; Nam et al., 2007; Ochiai et al., 2008; Sethi et al., 2009; Su et al., 2009; Ben-Tabou de-Leon and Davidson, 2010; Damle and Davidson, 2011; Li et al., 2013; Materna et al., 2013a; Li et al., 2014; Erkenbrack et al., 2018). While a well-annotated genome and transcriptome for Sp are available (Sodergren et al., 2006; Tu et al., 2012; Tu et al., 2014), similar resources have been lacking for Lv.

This study presents the developmental transcriptome for the sea urchin *L. variegatus* at 11 timepoints during embryonic and larval development, and provides an online database of the sequences along with annotation, Gene Ontology (GO), Pfam, and BLAST information, which we anticipate will be an important resource for the sea urchin community and a foundation for subsequent systems-level efforts, such as tissue-specific sequencing and proteomics. Unbiased analyses partition the developmental time course into four phases with relatively little internal variation that are separated by large transitions in gene expression, demonstrating that developmental gene expression is temporally punctuated rather than smooth and thus underlining the modularity of development in this organism.

## Results

### Sequence Collection, Annotation, and Validation

To determine the profile of gene expression during the development of *Lytechinus variegatus* (Lv), we sequenced the whole embryo transcriptome at 11 stages of embryonic and larval development (Fig. 1A). These stages were chosen because the intervals between them correspond to important transitions in the development of this species. Between 2-cell and 60-cell stages, the process of Wnt8/ß-catenin-dependent anterior-posterior (AP) specification initiates, while Nodal-dependent dorsal-ventral (DV) specification initiates during the transition from early blastula (EB) to hatched blastula stage (HB) in Lv (Hardin et al., 1992; Davidson et al., 1998; Wikramanayake et al., 1998; Logan et al., 1999; Wikramanayake et al., 2004; Bradham and McClay, 2006). Vegetal/posterior cells become elongated at thickened vegetal plate stage (TVP), prior to the ingression of the skeletogenic primary mesenchyme cells (PMCs, Fig. 1A red) at mesenchyme blastula (MB) stage (Miller and McClay, 1997; Wu et al., 2007). The remaining vegetal plate buckles inward, invaginates, and undergoes convergent extension at early, mid and late gastrula stages (EG, MG, LG), respectively (Hardin, 1996; Beane et al., 2006a). At LG, the secondary mesenchyme cells (SMCs; Fig. 1A orange) delaminate from the tip of the gut, and give rise to pharyngeal muscle cells, pigment cells, coelomic pouch cells, and blastocoelar cells (Ruffins and Ettensohn, 1996; Logan and McClay, 1997; Sherwood and McClay, 1999). The PMCs secrete skeletal triradiates that are visible at LG (Fig. 1A, yellow), and undergo substantial growth between LG and early pluteus (EP) stages (Wolpert and Gustafson, 1961). After LG, the larval mouth forms (Fig. 1A, “M”) (Hardin and McClay, 1990). Finally, neuronal development becomes detectable between EP and late pluteus (LP) in Lv (Bradham et al., 2009).

**Figure 1.**
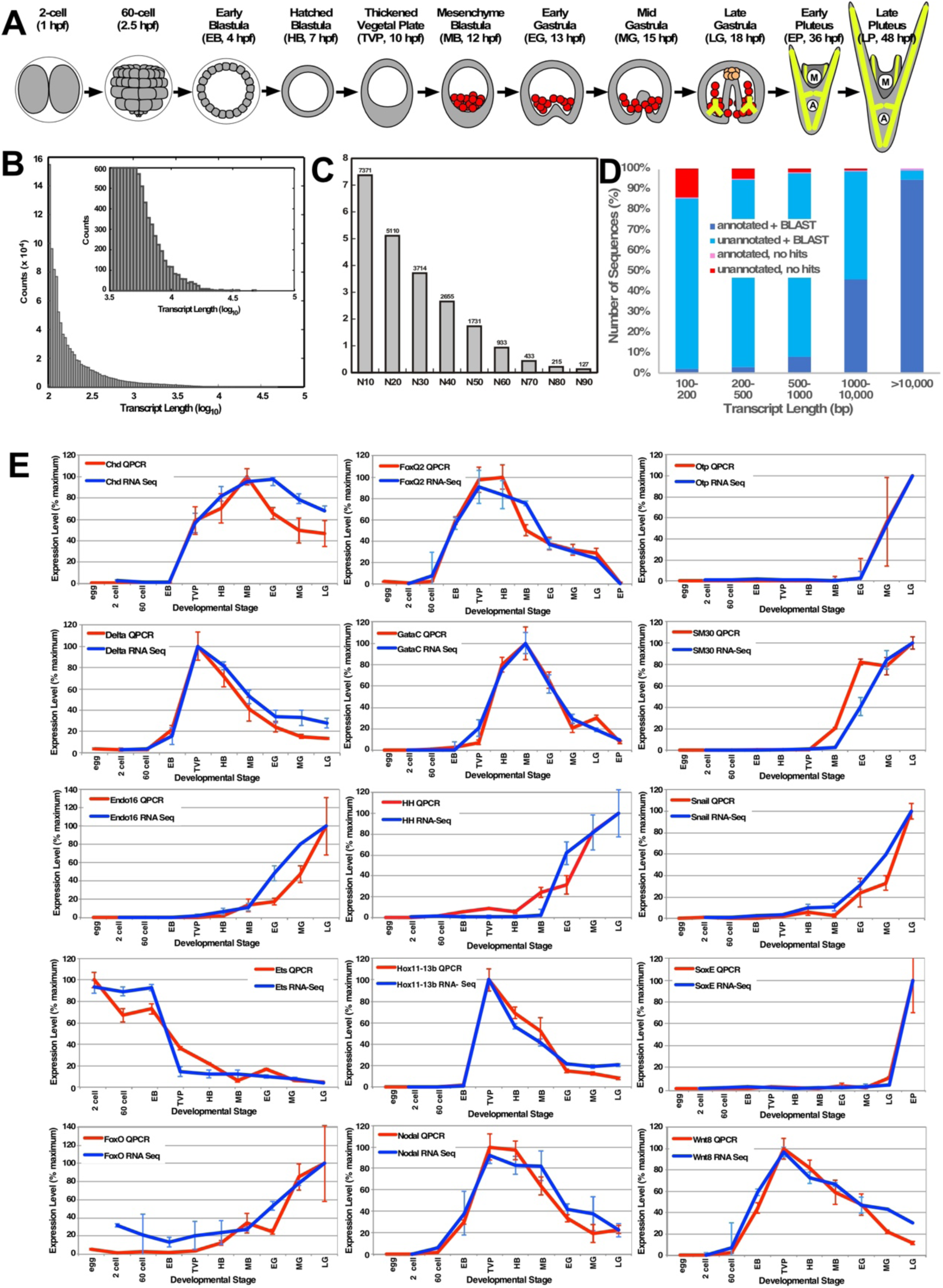
Transcriptome assembly analysis. **A.** A schematic indicating the developmental stages sequenced in this study. The earliest stages are enclosed in a fertilization envelope from which the embryo hatches at HB. The skeletogenic primary mesenchyme cells (PMCs) are indicated in red, while the skeleton is depicted in yellow. The secondary mesenchyme cells (SMCs) are depicted in orange. The PMCs ingress from the vegetal plate at MB, while the SMCs delaminate from the tip of the archenteron at LG. The growth of the skeleton supports the final shape of the larva. The mouth (M) and anus (A) are indicated. **B.** A histogram of transcript lengths (kb) for the Lv assembly is shown, with the upper range enlarged in the inset. **C.** A plot of N10-N90 values for the scaffolds in the Lv assembly is shown. See also Fig. S1. **D.** A plot of genomic alignments for annotated and unannotated genes is shown as a function of the transcript length. **E.** RNA-seq (blue) and QPCR (red) quantitations are compared for the indicated well-known genes. QPCRs were normalized to Lv-Setmar (Fig. S2). In each case, the results were scaled from 0 to 100, then plotted as average ± SEM for three biological replicates. See Table S1 for qPCR primer sequences.

We utilized the Illumina GAII, HiSeq, and HiSeq4000 platforms to collect the sequence data from three biological replicates and SOAPdenovo-Trans to assemble the resulting reads. The combined RNA-seq data generated 956,587 scaffolds. The largest scaffold was 49,229 base pairs (bp), and the N50 was 1731 bp (Fig. 1B, C). The distribution of transcript lengths (Fig. 1B) and N(x) values (Fig. 1C) illustrates that the size distribution for this assembly is regular and smooth, indicating that the assembly contains a well-distributed range of scaffold sizes. We initially used DESeq to normalize the expression values across the samples; however, DESeq did not yield good agreement between samples for the overall range of expression values (Fig. S1A). This is probably due to the sequencing platform differences. Based on previous comparative analyses (Bradnam et al., 2013), we instead employed quantile normalization (Hansen and Irizarry, 2012), which produced substantially better agreement (Fig. S1B).

We annotated the scaffolds and contigs by comparing them via BLASTx (Altschul et al., 1997) to known *S. purpuratus* (Sp) genes, which are well annotated and highly similar to Lv genes, particularly at the protein-coding level. We identified 57,500 predicted Lv transcripts that correspond to predicted Sp genes, of which 87.3% are annotated. These resolved to 18825 unique matches to Sp genes, which is comparable to the number of genes identified in transcriptome analysis in Sp (Tu et al., 2012), and reflects 62.9% of the currently predicted 29,949 Sp genes from genomic analyses (www.echinobase.org). We identified GO terms for the annotated sequences using BLAST2GO, and Pfam identifiers using HMMer; of the annotated transcripts, 42.4% have both GO and Pfam identifiers, 3.2% have only GO terms, 37.4% have only Pfam identifiers, and 17.0% have neither.

We aligned the scaffolds to the Lv genome sequenced at Baylor University (www.echinobase.org) using BLAST analysis (Altschul et al., 1997), and found that 88.9% of the total scaffolds aligned to the genome at e = 10^−6^ or less, with 95.2% of the annotated scaffolds and 88.5% of the unannotated scaffolds aligned. Among the aligned sequences, an average of 94.2% of the length of the transcriptome sequences aligned to the genome. We note that as the scaffold sequence length increases, the fraction of both annotated sequences and sequences that align to the genome increases (Fig. 1D). However, since many of our annotated transcriptome scaffolds were longer than the genome scaffolds, we were not able to use the Baylor genome to improve our assembly. We further evaluated the assembly quality by searching for the presence of the 248 most conserved eukaryotic genes (CEGs) using CEGMA, which employs hidden Markov models for orthologous genes to identify sequences matching the defined set of CEGs (Parra et al., 2007). We found 240 of the 248 genes (96.8%), providing an estimate of the completeness and accuracy of the assembly.

To validate the quantitation of the expression data, we used qPCR analysis of three independent biological replicates distinct from those used for RNA-seq analysis, then compared the results with the quantile-normalized RNA-seq expression data for 15 well-known genes. These genes were chosen to reflect a range of expression profiles, with maximal expression for each gene occurring across the range of the sequenced stages (Fig. 1E). Sea urchin qPCR analysis typically employs ubiquitin as a normalization gene, despite the order of magnitude change in ubiquitin expression level during development (Fig. S2A). We therefore sought a less dynamic gene for use as a normalizer in these analyses, and identified Lv-Setmar as a gene with very consistent expression across this developmental time course (Fig. S2). We thus used Lv-Setmar to normalize qPCR results in this study. Expression trends were generally in good agreement between RNA-seq and qPCR quantitations. The Pearson correlation for the RNA-seq and qPCR measures for these 15 genes was 0.962, indicating that the quantitation of the RNA-seq results reliably matches independent empirical measurements of gene expression. We cloned about twenty genes based on the sequences predicted by the RNA-seq assemblies, and in all cases, the empirical clone sequences match the predicted gene models very well (~99%), providing another indicator that the assembly is reliable, and that the inclusion of misassembled artifacts among known genes is minimal.

### Expression Analysis: GRN Network Circuits

We evaluated the timing of expression for groups of genes that function in well-studied GRN circuits within five sea urchin tissues, in keeping with the analyses performed by Gildor et al (Gildor and Ben-Tabou de-Leon, 2015). The PMC lineage, which gives rise to the skeletogenic mesoderm, arises at the 16-cell stage and is a crucial source of inductive signals for endomesoderm specification (Activin B) and subsequent mesoderm segregation (Delta) (Sherwood and McClay, 1999; Sweet et al., 2002; Revilla-i-Domingo et al., 2004; Sethi et al., 2009). In *S. purpuratus* (Sp), PMC specification depends on a coherent feed-forward circuit that is driven by the maternal transcription factor Ets1, which activates Alx1, then Hex. Alx1 and Ets1 together drive Dri expression; all four of these factors are required for the expression of SM50, a skeletogenetic differentiation gene (Revilla-i-Domingo et al., 2004; Oliveri et al., 2008; Damle and Davidson, 2011) (Fig. 2A1). Comparisons with Sp and *P. lividus* (Pl), a Mediterranean species, show that Ets1 is maternally expressed in all three species, while Alx1 is maternal in Lv and Pl, but not Sp. The timing of Hex, Dri, and SM50 expression varies, in that SM50 and Dri are coincident and precede Hex in Sp, while Dri and Hex are coincident and precede SM50 in Lv (Fig. 3B1, C1) and in Pl (Gildor and Ben-Tabou de-Leon, 2015). These results indicate that some wiring differences exist in the regulation of Hex in particular, shifting it to slightly later expression in Sp, and show a lack of requirement for Hex to drive SM50 in Sp, suggesting a closer relationship between Pl and Lv relative to Sp regarding this circuit.

**Figure 2.**
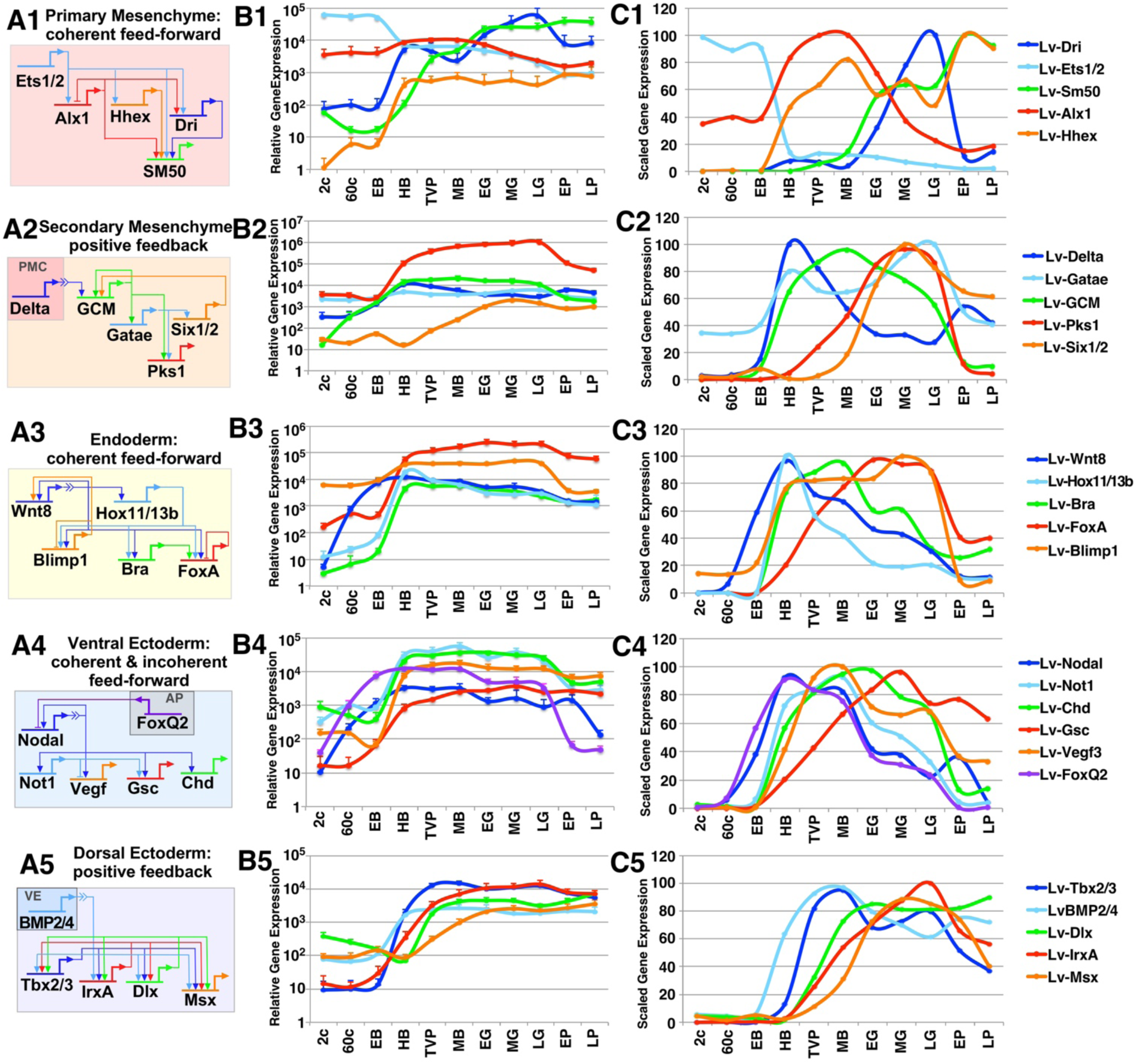
A comparison of the relative timing of gene expression onsets in five known GRN circuits that operate in distinct territories. **A.** Schematics illustrating five well-known and conserved network motifs, each composed of five or six genes that encode transcription factors or signals (i.e. Delta, Wnt8, Nodal, Vegf, and BMP2/4) in a range of territories, including the PMCs (1), the SMCs (2), the endoderm (3), the ventral ectoderm (4) and the dorsal ectoderm (5). AP, apical plate; VE, vegetal ectoderm. **B.** Unscaled and **C.** scaled temporal gene expression profiles are shown for the genes depicted in A. The unscaled plots reveal the relative gene expression levels, while the scaled plots more clearly show the temporal onset relationships.

**Figure 3.**
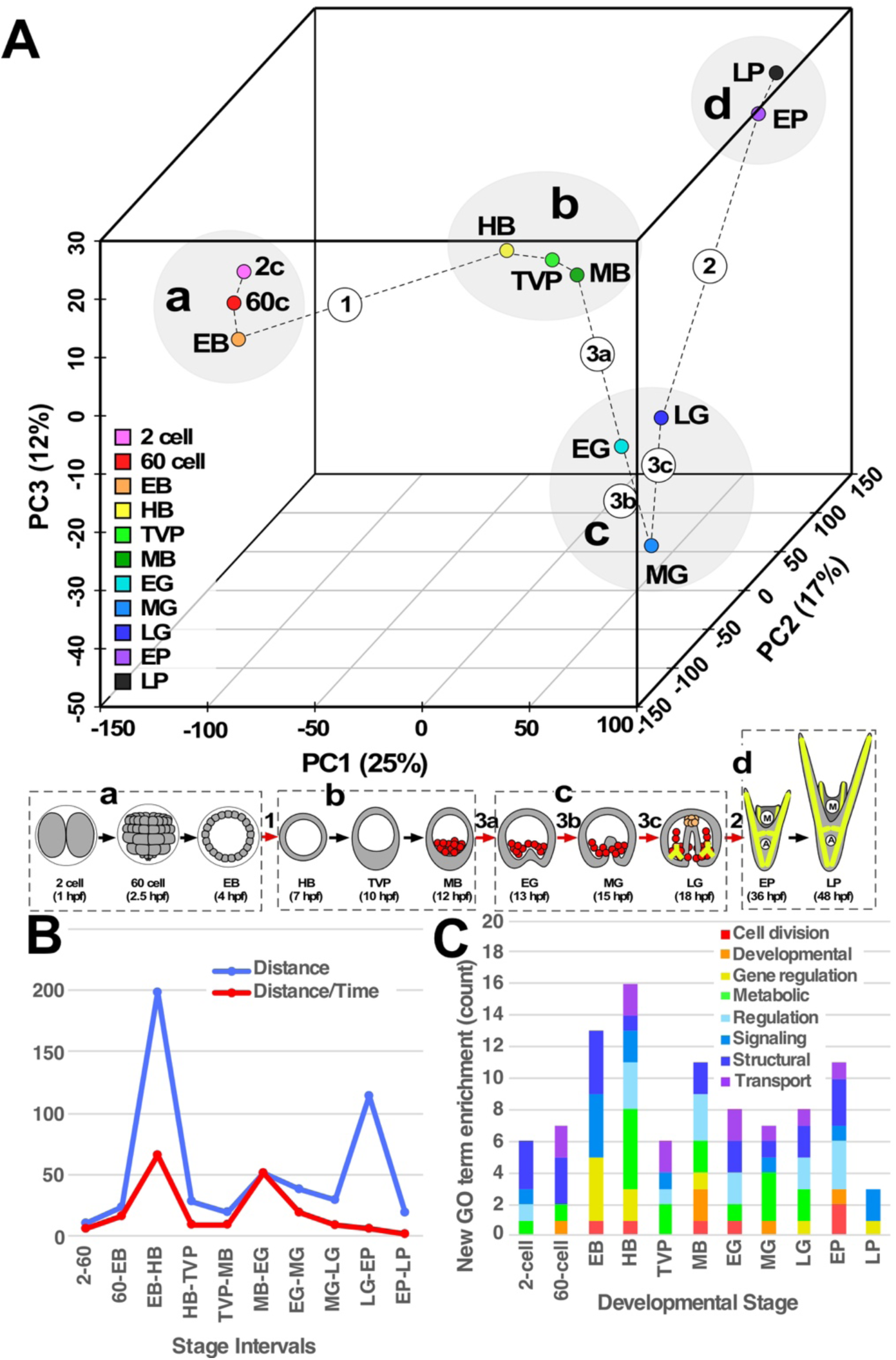
Principal component analysis (PCA) indicates that the transition from early blastula to hatched blastula stage accounts for the largest variation in the Lv transcriptome, and separates the developmental stages into four phases of gene expression. **A.** PCs were calculated using the complete set of transcripts from each sequenced developmental stage, and the results are plotted along the first three principal components (PC1, PC2, and PC3). The temporal relationships between the stages are indicated with dashed lines. The largest transitions along each PC are indicated by circled numbers; for the third PC, three transitions are indicated (3a, 3b, and 3c). This series of transitions divides the developmental stages into 4 phases, designated a-d. The major transitions (red arrows) and phases (dotted boxes) are noted in the schematic below the plot. See also Table S2 for PCA values. **B.** Euclidean distances in phase space (blue) and the rate of change (red) are shown between each developmental stage. **C.** The number of newly enriched GO terms within each stage is plotted. See also Fig. S4 and Table S3.

SMCs are segregated from endoderm via reception of a Delta signal from the adjacent PMCs (Sherwood and McClay, 1999; Sweet et al., 2002). Delta-Notch signaling is mediated by a positive feedback circuit in which the Notch intracellular domain co-activates the transcription factor Gcm, which activates GataE and itself; GataE in turn activates Six1/2 which feeds back to Gcm; GataE and Gcm together activate the differentiation gene Pks1 (Lee and Davidson, 2004; Ransick and Davidson, 2006; Lee et al., 2007; Croce and McClay, 2010; Ransick and Davidson, 2012; Materna et al., 2013a) (Fig. 2A2). In Lv, Delta exhibits early non-zero expression, with an increase at early blastula stage (EB), while the onset of Gcm occurs between the 2- and 60-cell stages, prior to the increase in Delta levels at early blastula stage (EB) (Fig. 2B2). GataE and Six1/2 also increase expression at EB, although Six1/2 exhibits only a small transient increase, followed by a much larger increase beginning at mesenchyme blastula stage (MB). Pks1 expression occurs last, with onset at hatched blastula stage (HB) (Fig. 2B2, C2), similar to what is observed in Pl and Sp (Gildor and Ben-Tabou de-Leon, 2015). The late peak of Six1/2 corresponds to prolonged expression of Gcm as well as a second peak of GataE, consistent with indirect Six1/2 input, via Gcm, into GataE regulation in Lv.

Endoderm specification relies on Wnt8 signaling (Wikramanayake et al., 2004), which drives a coherent feed-forward circuit in which Wnt8 inputs (via ß-catenin) drive Hox11/13b, Blimp1, and Brachyury (Bra) expression, with Hox11/13b also driving Blimp1 and Bra expression; Wnt8, Hox11/13b and Bra each feed into FoxA expression, which is autorepressive (Wikramanayake et al., 2004; Minokawa et al., 2005; Smith et al., 2007; Smith and Davidson, 2008; Smith et al., 2008; Ben-Tabou de-Leon and Davidson, 2010; Peter and Davidson, 2010, 2011) (Fig. 2A3). In Lv, Wnt8 expression occurs first between the 2- and 60-cell stages; all four of the other genes in the circuit onset between EB and HB, with FoxA reaching its maximum value the most slowly, consistent with autorepressive wiring (Fig. 2B3, C3). This is largely similar to Sp and Pl (Gildor and Ben-Tabou de-Leon, 2015), although the slow peak of FoxA expression is distinct, implying stronger autorepression and/or weaker activation effects for FoxA in Lv.

The ventral region of the ectoderm is specified by Nodal signaling, which activates Not1 and Gsc expression, both of which are required for ventral specification, and Chordin (Chd), which inhibits BMP signaling in the ventral region (Duboc et al., 2004; Bradham and McClay, 2006; Bradham et al., 2009; Lapraz et al., 2009; Su et al., 2009; Saudemont et al., 2010; Li et al., 2012, 2013; Li et al., 2014). FoxQ2 directs neural specification, and Nodal expression is suppressed by apical (animal plate) FoxQ2 expression; this antagonism is thought to promote the boundary between the ventral region and the adjacent apical neural region (Yaguchi et al., 2008). A pair of posterior lateral subdomains within the ventral ectoderm express VEGF, which signals to promote PMC positioning and biomineralization adjacent to those subdomains, and to drive posterior secondary skeletal patterning (Duloquin et al., 2007; Adomako-Ankomah and Ettensohn, 2013; Piacentino et al., 2016b). VEGF is induced by Nodal (probably indirectly) and repressed by Not1; the ventral-centric Not1-mediated repression is thought to participate in the spatial restriction of VEGF expression to the posteriolateral subdomains (Li et al., 2012). For Lv embryos, we observe that the onset of expression of Nodal and FoxQ2 is temporally coincident, between 2- and 60-cell stages (Fig. 2B4). The other genes in this circuit each onset between EB and HB, except Gsc, which onsets between 60-cell and EB, and shows the slowest activation. These dynamics appear more similar to Pl with respect to VEGF, which exhibits a maternal phase of expression and is similarly preceded by Gsc; however, the late-peaking expression of Gsc is more similar to Sp (Gildor and Ben-Tabou de-Leon, 2015).

Dorsal specification requires BMP2/4 signaling. Interestingly, BMP2/4 is expressed in the ventral territory, but signals only in the dorsal region; this spatial disconnection is likely due to the expression of the BMP inhibitor Chd in the ventral region (Angerer et al., 2000; Duboc et al., 2004; Bradham et al., 2009; Lapraz et al., 2009; van Heijster et al., 2014). BMP2/4 signaling activates the expression of transcription factors Tbx2/3, IrxA, Dlx, and Msx (Lapraz et al., 2009; Su et al., 2009; Saudemont et al., 2010); loss of function analyses suggest that positive feedback circuitry interconnects these genes (Saudemont et al., 2010; Ben-Tabou de-Leon et al., 2013). In Lv, we observe the activation of BMP2/4 expression between EB and HB, then activation of Tbx2/3 between HB and TVP, and finally the other genes activate between TVP and MB (Fig. 2B5, 2C5). This is dissimilar to both Sp and Pl: in Sp, Tbx2/3 onset is coincident with the other downstream genes rather than preceding them, while in Pl, Tbx2/3 expression onset is coincident with BMP2/4 (Gildor and Ben-Tabou de-Leon, 2015). Aside from this, the dynamics in Lv appear more similar to Pl, in which the onset of Msx is delayed compared to the remaining genes as it is in Lv (Fig. 2B5, C5). Overall, these results indicate that these five circuits are generally well-conserved across a range of sea urchin species, with only minor variations in gene expression timing between Sp, Pl, and Lv. This in turn suggests that the overall architecture of the sea urchin specification GRN models are well-conserved.

### Expression Analysis: PCA reveals Distinct Phases of Developmental Gene Expression

We used principal component analysis (PCA) (Sclens, 2005) to evaluate the overall variation in the expression data over the developmental time course captured by our sample range. We included all expressed transcripts in this analysis, using averaged expression values from our three biological replicates. The results show that the first three principal components (PCs) account for 54% of the variation in the data. PC1 is the most recognizable of the axes in that it corresponds well with time, since all the stages occur in temporal order along this component, with 2-cell and LG stages being minor exceptions (Table S2). The largest transition is between EB and HB along PC1 (Fig. 3A, Table S2), which corresponds with the onset of DV specification and the elaboration of AP specification. The second largest transition is between LG and EP along PC2, which corresponds with a significant degree of morphological change, including formation of the mouth, development of the skeleton and ciliary band, and overall morphogenesis. There are three sizeable transitions along PC3, which occur sequentially along the progression from MB to LG. Interestingly, transitions 3b and 3c produce nearly opposite effects, such that EG and LG are positioned similarly in phase space. This group of developmental stages comprises gastrulation, during which significant migration of the endoderm and mesoderm occurs. This set of transitions effectively separates the embryonic stages into four distinct modules or phases with relatively little internal variation, which we designated a-d (Fig. 3). These results show that developmental gene expression dynamics in Lv proceed in an abrupt, punctuated manner, rather than smoothly over time.

We compared the Euclidean distances and the rate of change between consecutive stages (Fig. 3B). These results show that transitions 1 and 3 are distinct and rapid, while transition 2, between LG and EP, is comparatively slow and not distinct from the prior and subsequent interstage rates (Fig. 3B, red). This is unsurprising since the transition 2 corresponds to the largest temporal gap in our dataset. This shows that the transcriptional changes that occur at HB and EG are quantitatively distinct and represent transcriptional bursts, unlike the transcriptional change at EP.

To better understand the nature of the gene expression changes that occur as development proceeds, we evaluated gene ontology (GO) term enrichment in each of our sequenced stages relative to temporally adjacent stages. We identified approximately 100 enriched GO terms, which we grouped into eight categories. We found that more new GO terms were enriched at HB (hatched blastula stage) compared to other stages (Fig. 3C), consistent with the PCA results (see also S.Fig. 4 and S.Table 3). The results show that transcriptional burst at HB stage is reflected by enrichment of GO terms across a range of categories, and includes ciliary motility, matching the onset of cilia-mediated swimming at this stage, and redox homeostasis, in keeping with known roles for redox signaling in mediating DV specification at this time (Coffman and Davidson, 2001; Coffman et al., 2004; Coffman et al., 2009; Modell and Bradham, 2011; Coffman et al., 2014; Chang et al., 2017) (Fig. S4; Table S3). However, while EP and EG both show a large number of newly enriched GO terms, they do not rank as the second and third stages in this regard as expected from the PCA results. Instead, EB (early blastula) has the second largest set of newly enriched GO terms, and MB (mesenchyme blastula) has more than EG (early gastrula). Thus, GO term enrichment does not consistently reflect the overall variance detected by PCA. We also evaluated GO enrichment among those transcripts expressed only within a single phase (Fig. S5 and Table S4). These results reveal interesting functional correlates, such as cell division during blastula stages in phase a, gene expression, signal transduction, and epidermis development during phase b when a significant degree of signaling and specification occurs, and neural-specific GO terms in phase d, when neural differentiation becomes apparent.

**Figure 4.**
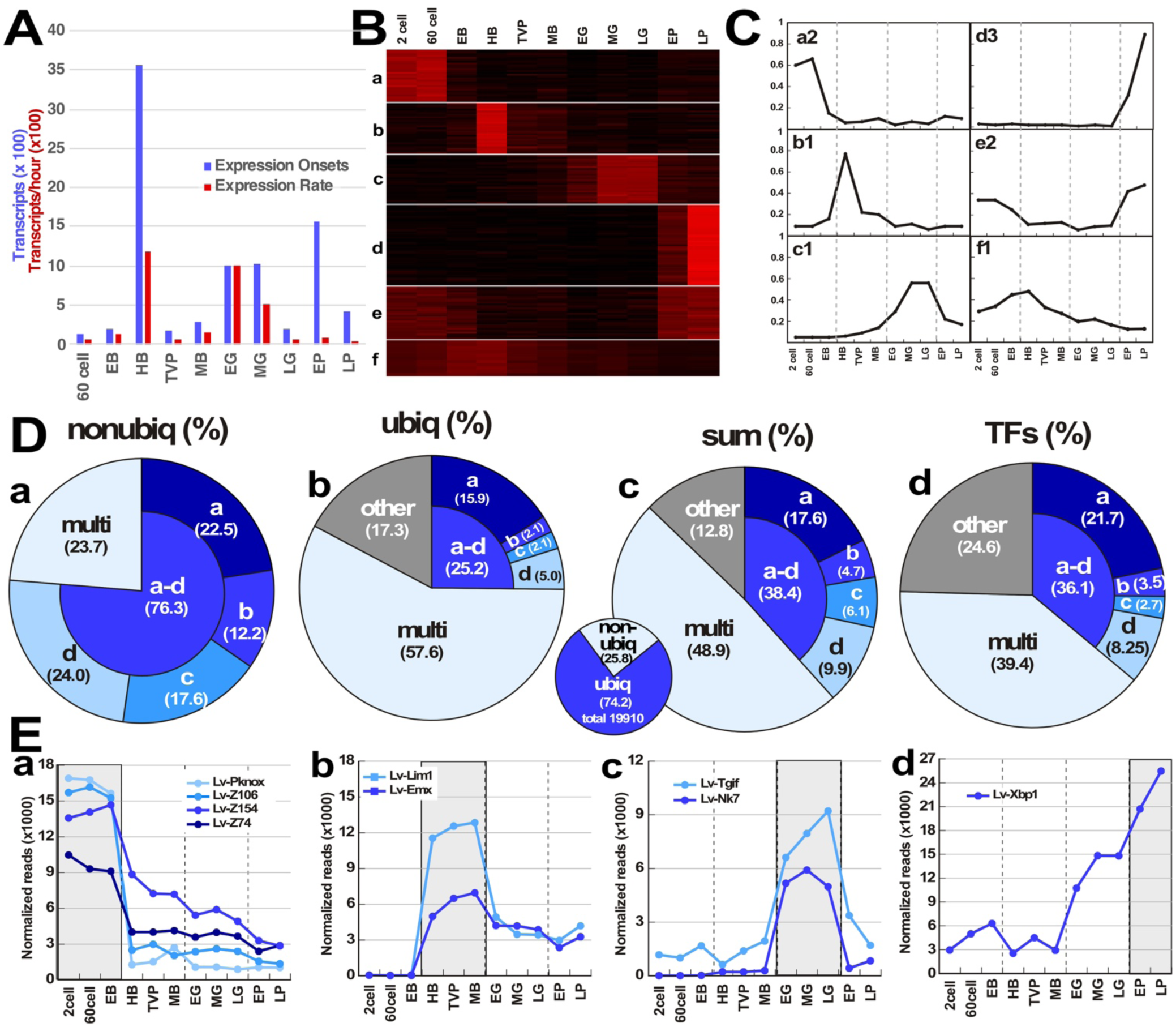
*K*-means clustering corroborates the PCA results. *K*-means clustering was performed on the annotated transcripts, divided into two groups: the non-ubiquitously expressed and ubiquitously expressed transcripts. Transcripts encoding transcription factors were also analyzed separately. **A.** The number of newly expressed transcripts (blue) and the rate of their expression (red) at each stage was estimated from the *k*-means clusters. For these calculations, each cluster profile was assigned to a stage of initial expression, then the number of transcripts per stage was summed. **B.** Exemplars of heat maps are shown that depict the major categories of gene expression profiles: confined to a single expression phase (a-d), confined to multiple phases (e, multi), and non-adherent to the expression phases (f, other). **C.** Average normalized expression profile plots for the heat maps shown in B, with the phase boundaries indicated by vertical dotted lines. See also Fig. S6-8 for the complete *k*-means heat maps and averaged expression plots for each phase, which include these exemplars, and Tables S5-7 for gene lists. **D.** The distribution of annotated transcripts in each of the six categories illustrated in B and C is shown for the non-ubiquitous transcripts (a), the ubiquitous transcripts (b), the sum of non-ubiquitous and ubiquitous transcripts (c), and the transcription factors (TFs, d). The inset pie chart shows the proportions of the ubiquitous and non-ubiquitous transcripts. **E.** Expression profiles for TFs whose expression is confined to a specific expression phase and which exhibited both the lowest expression level variance across the phase and the highest expression level within that phase (a-d).

### Expression Analysis: Gene Clustering reproduces Distinct Phases of Gene Expression

We used *k*-means clustering (Steinley, 2006; Do and Choi, 2008) to define the temporal expression patterns within the Lv transcriptome. We limited this analysis to the transcripts with Sp homologs (“annotated transcripts”, although a fraction of these remain unnamed), then partitioned these sequences into ubiquitous transcripts (expression > 1% of the maximum average expression level per transcript at all timepoints), or non-ubiquitous transcripts (expression ≤ 1% of the maximum for that transcript for at least one timepoint). We reasoned that non-ubiquitous transcripts are subject to regulatory control that is more complex than the regulation of ubiquitous transcripts, and thus the expression profiles for the non-ubiquitous genes are likely to be distinct from those for the ubiquitous genes. For the set of 5,136 non-ubiquitous sequences, we identified 19 clusters of gene expression profiles, with an average of 270 transcripts/cluster (Fig. S6, Table S5). For the set of 14,774 ubiquitous transcripts, we identified 37 clusters of gene expression profiles, with an average of 400 genes/cluster (Fig. S7, Table S6). In comparison to the non-ubiquitous set, the number of clusters is nearly double, while the number of genes is almost triple for the ubiquitous set; the averages show that the ubiquitous set is less complex than the non-ubiquitous set, as anticipated. Since sea urchin developmental specification relies on the hierarchical deployment of transcription factor (TF) networks, we also performed *k*-means cluster analysis on the 521 TF transcripts within the Lv transcriptome, which identified 23 clusters of gene expression, with an average of 23 transcripts/cluster (Fig. S8, Table S7). Unsurprisingly, the complexity of expression patterns is considerably increased among the TFs.

We estimated the number of newly expressed genes and the rate of new gene onsets in each stage from these analyses, and found that, excluding genes with maternal/2-cell stage onset, HB exhibits the largest number of newly expressed transcripts, followed by EP, then EG and MG (Fig. 4A, blue). These results are in very good agreement with the PCA findings (Fig. 3A). However, as with the distance measures from the PCA (Fig. 3B), HB and EG exhibit the largest rates of gene expression onset (Fig. 4A, red), consistent with the expected bursts of gene expression at these two stages. In contrast, the large number of new gene expression at EP are not expressed at a high rate and thus do not correspond to a transcriptional burst.

From the perspective of the four expression phases defined by the PCA (Fig. 3A), we noted striking agreement between these phases and the gene expression profiles within most of the *k*-means clusters. Accordingly, we sorted the clusters into six sets, with examples shown in Fig. 4B and C: those whose expression is strictly confined to one of the four phases (a-d), those whose expression occurs in more than one phase but changes at the phase boundary, “respecting” that boundary (e, “multi”), and finally those whose expression does not change at the phase boundaries (f, “other”) (Fig. S5-7).

For the non-ubiquitous set of transcripts, the majority (76.3%) are confined to a single phase, and a minority (23.7%) are expressed in more than one phase (Fig. 4Da, Fig. S6, Table S5). There are two notable points: first, a comparatively large number of the non-ubiquitous transcripts exhibit expression that is confined to a single developmental stage (Fig. S6); second, the cluster averages all exhibit expression profiles that conform to the phase boundaries, with no examples of the “other” type that is not restricted by the phase boundaries among the non-ubiquitous transcripts.

In contrast, the ubiquitous set of sequences exhibits only a minority of phase-specific transcripts (a-d, 25.2%), while the majority of transcripts were expressed in more than one phase (e, multi, 57.6%); finally, 17.3% of the transcripts exhibited temporal profiles that were not restricted by phase boundaries (f) (Fig. 4Db, Fig. S7, Table S6). Since there is measurable expression for every sequenced developmental stage among the ubiquitous set, we considered expression to be positive above a threshold of 15% of the maximum for each transcript in this analysis, to group the profiles. Only a very small fraction of transcript profiles in this set exhibit single stage-specific expression even with this high threshold (Fig. S7); instead, most of the ubiquitously expressed transcripts are in the “multi” category. Together, the non-ubiquitous and ubiquitous transcripts exhibit an intermediate distribution, with 12.8% of the annotated transcripts in the “other” category (Fig. 4Dc).

The TFs contain a relatively large fraction of “other” genes (24.6%) (Fig. 4Dd). Very few TF clusters have expression in only one developmental stage, while the majority of TFs are expressed in three or more sequential stages, with 75.4% of profiles conforming to the expression phases (Fig. S8, Table S7). This interesting result suggests that in general, TF expression is both more temporally continuous than general gene expression, and also bridges the expression phases more often than general gene expression.

Overall, the *k*-means cluster analyses suggested that global chromatin changes might underlie the PCA transitions. In mammals, particular TFs have been identified as “pioneer factors” that bind chromatin prior to other TFs, and function to promote an open chromatin conformation that permits the subsequent binding of other TFs (Zaret and Carroll, 2011). In this way, pioneer factors control which parts of the genome are available for transcription by initializing specific global chromatin states (Mullen et al., 2011; Zaret and Carroll, 2011). Since most (87.2%, Fig. 4Dc) of the developmental transcripts exhibit expression profiles that conform to the phases defined by PCA (Fig. 3A), we reasoned that a group of transcription factors, functionally similar to pioneer factors, might maintain, rather than initialize, specific global chromatin states; such factors could be responsible for and underlie the major expression phases within the Lv transcriptome.

To identify any such putative “chromatin-state maintenance” candidates, we sought TFs whose expression meets three criteria: first, expression is confined to a single expression phase, consistent with maintaining that corresponding state; second, expression exhibits minimal variation across the relevant phase, consistent with a primary role in regulating chromatin status; and third, expression is at a high level within the phase, consistent with binding at a relatively large number of genomic locations. For the final criterion, we used an order of magnitude cut-off in expression level for each set of least-variably expressed transcription factor transcripts.

Surprisingly, this analysis identified only a few factors for each expression phase, and nine factors in total (Fig. 4Ea-d). Four TFs were found for phase a, and two for phase c (Fig. 4Ea, 4Ec). For phase d, only one TF is strongly expressed specifically in the relevant time period, although its expression is relatively variable within that period, and is also elevated earlier (Fig. 4Ed). Similarly, for phase b, the expression profiles for only two TFs meet our criteria, and each exhibits fairly high expression levels within the subsequent phases as well (Fig. 4Eb). Thus, in both of these cases, the expression profiles for the identified TFs are not strictly confined to a single phase, but nonetheless are most strongly elevated in the phase of interest. This speculative analysis neglects more complex cases and makes the simplifying assumption of proportionate transcription and translation; however, it nonetheless provides a starting point for further analyses. It will be of interest to determine whether these factors influence chromatin state, and to determine their global genomic binding locations and how these locations overlap with those of other factors. It will also be of interest to determine whether global chromatin status is consistent across the expression phases, but variable between them, using approaches such as ATAC-seq (Buenrostro et al., 2013).

The *k*-means clustering results for the annotated subset of transcripts reinforce the global PCA findings, and indicate first that overall onset of new transcripts is maximal at predicted phase transitions, and second, that the expression phases are reflected by 87.2% of all the annotated transcripts. These data also show that the non-ubiquitous set of transcripts has a strikingly different, temporally restricted set of expression profiles compared with the other analyzed sets, and is completely adherent to the expression phases, whereas the other sets are 75% or more adherent (Fig. 4C). The overall and TF-specific results each have implications for the temporal behavior of the TF networks that drive developmental specification, suggesting that network composition is relatively stable within each phase, and is punctuated by comparatively abrupt changes at specific intervals that correspond to the major phase transitions. Phase c, with two internal transitions, appears to be the exception, and instead exhibits steady change over time rather than a stable state (Fig. 4A), in keeping with comparatively large PC transitions within that module (Fig. 3A, B). Overall, these results demonstrate abrupt changes in gene expression at the intervals corresponding to the major phase transitions, corroborating those findings.

### Expression Analysis: The Specification Network Corroborates Expression Phases

To determine whether the specification GRNs conform with the expression phases, we evaluated the temporal expression of the Lv genes that correspond to genes within known specification network models for the sea urchin species Sp and Pl. These genes include transcription factors, a small number of signals (Wnts, Delta, Univin, Nodal, BMPs, and VEGF) and signal inhibitors (Chordin and Lefty), and a few differentiation genes (Pks1, Endo16, Msp130, SM30, and SM50). We determined the onset of expression for each gene in this set of known GRN genes (Fig. 5) to learn whether the phase transitions are evident, as well as to determine whether the overall hierarchy of onset in Lv agrees with the logic of the known networks, in an extension of the circuit analysis described above (Fig. 2). We calculated the onset of gene expression using the Sigmoid function in Python as described (Gildor and Ben-Tabou de-Leon, 2015) for ~ 70% of these genes; the expression profiles for the remainder were not amenable to this analysis because of bimodality or other irregularities; in these cases, expression onset was interpolated by comparison with expression profiles with calculable onsets. This analysis is based on the assumption that the same or very similar networks operate in Lv, as has been indicated for the PMCs (Saunders and McClay, 2014) and as suggested by our circuit analysis herein (Fig. 2).

**Figure 5.**
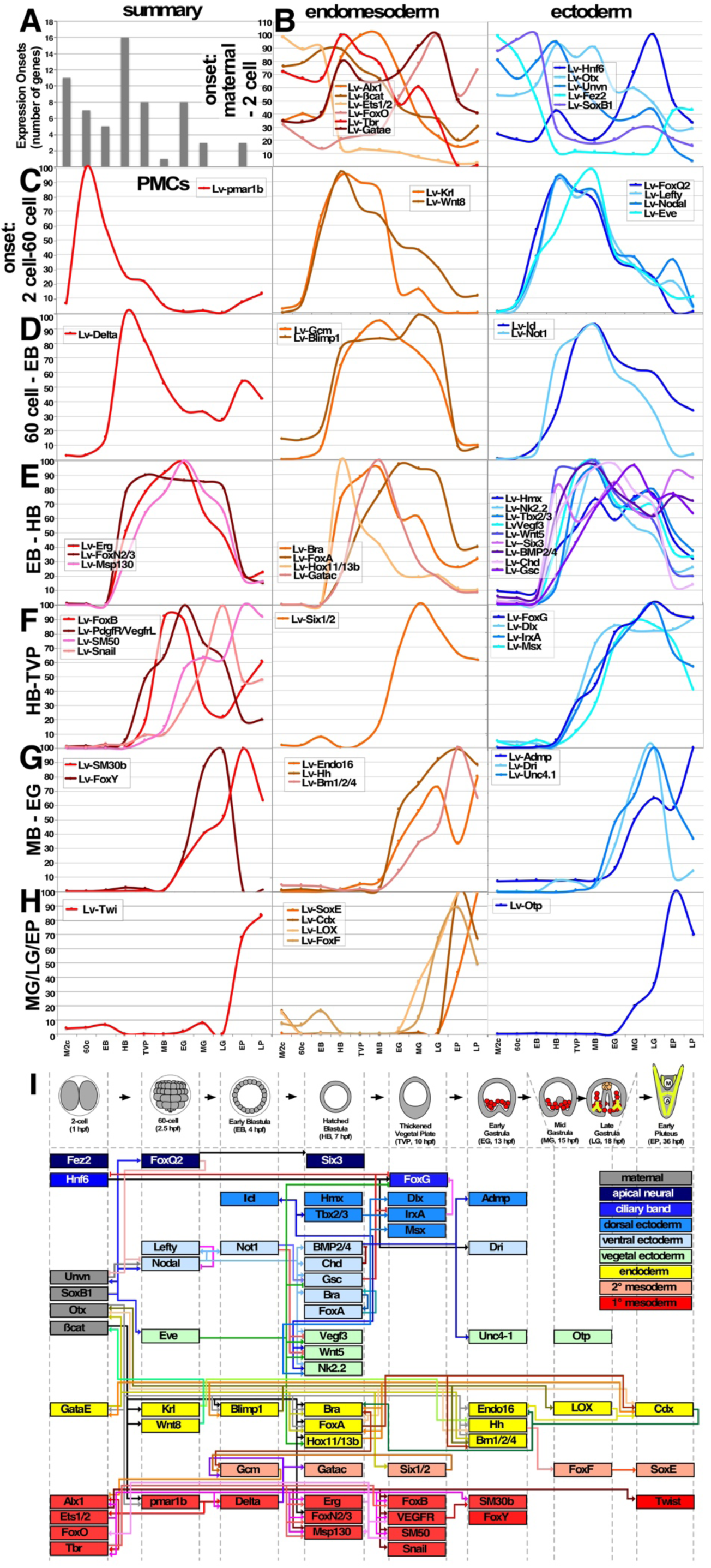
Among the major GRN genes, the maximal number of onsets occurs between EB and HB stages, consistent with the PCA results. **A.** A summary of the onset analysis for 68 genes in the known endomesodermal or ectodermal gene regulatory network (GRN) models, presented as the number of GRN gene expression onsets at each developmental stage. **B.** Genes with maternal onsets, separated into endomesodermal and ectodermal genes. **C.-H.** Genes with zygotic onsets are separated by stage of onset and tissue (primary mesenchyme cells (PMC), endomesoderm, or ectoderm). Onsets at midgastrula (MG) or later are combined into single plots (H). **I.** The network relationships among the GRN genes, sorted by time of onset in Lv and tissue. The expression onset for the indicated genes is depicted by their appearance along the horizontal axis, representing time, with the associated stages indicated schematically along the top. The ectodermal territories are ventral (light blue), dorsal (medium blue), the ciliary band (royal blue), which is positioned as a stripe at the boundary between the dorsal and ventral ectodermal territories, the animal plate, which is neurogenic (deep blue), and the posterior-most ectodermal belt which is derived from vegetal blastomeres (light green). The endomesodermal territories are the primary mesenchyme cells (PMCs, red), the secondary mesenchyme cells (orange), and the endoderm (yellow). Maternally expressed genes are also indicated (grey). Midgastrula and late gastrula onsets are combined into a single column. Colored edges are drawn based on relationships identified in Sp and/or Pl. Some genes are expressed in more than one tissue. In some cases, the gene is shown in both tissues (e.g. Bra, FoxA). In other cases, the gene is shown in the earliest tissue (e.g. ß-catenin).

We sorted the results by tissue, separating PMCs, endomesoderm, and ectoderm. We then grouped the results by time of onset (Fig. 5B-H). The predicted wiring diagram among these genes in Lv is shown in Fig. 5I. The results show that, for the most part, the relationships between the onsets of expression for GRN genes do not violate the network logic established for Sp and Pl (Davidson et al., 2002; Su et al., 2009; Saudemont et al., 2010; Peter and Davidson, 2011; Materna et al., 2013a; Li et al., 2014).

In the stages that correspond to phase a (2-cell to EB), key regulators for each major tissue are expressed (Fig. 5B-D). These include Pmar (mesoderm), Wnt8 (endoderm), Nodal (ectoderm), and FoxQ2 (neural ectoderm) (Oliveri et al., 2002; Duboc et al., 2004; Wikramanayake et al., 2004; Yaguchi et al., 2008), whose expression in Lv is first detected at the 60-cell stage. Other key regulators SoxB1, Otx, and ß-catenin are maternally expressed, as is Univin, which is important for ectoderm specification and skeletal patterning (Range et al., 2007; Li et al., 2014; Piacentino et al., 2015). Delta is expressed by the micromeres (PMC precursors) at EB, and signals for SMC specification, which separates the SMCs from the endoderm (Sherwood and McClay, 1999; Sweet et al., 2002). Lefty is also expressed at EB by the ectoderm, downstream from Nodal; here, Lefty functions to restrain Nodal signaling to the ventral side (Duboc et al., 2008).

At hatched blastula (HB) stage, the largest number of GRN genes initiate expression, consistent with the PCA findings (Fig. 5A, 5E). These genes include BMP2/4, which is the key signal for dorsal specification and is expressed ventrally downstream from Nodal (Angerer et al., 2000; Duboc et al., 2004; Bradham et al., 2009), and Six3, which is upstream from many neural genes in the anterior plate (Wei et al., 2009). Many additional genes, including signals and TFs, are initially expressed at HB. The transition from EB to HB corresponds to the largest PCA transition, and this network characterization further corroborates those findings and agrees with the results from *k*-means clustering of abrupt changes during the phase transitions. Since the extant GRN models are focused on early specification, it is difficult to similarly analyze the later PCA transitions because the network becomes too sparse at later stages.

A few genes in this analysis do not agree with predictions from Sp and Pl. Among them, the PMC gene Alx1 is expressed earlier than the Sp and Pl GRNs predict, since it is present before Pmar is expressed, yet is modeled as becoming expressed downstream of the double-negative gate regulated by Pmar (Oliveri et al., 2002; Oliveri et al., 2003; Rafiq et al., 2012). However, others have similarly found that Alx1 expression precedes the double-negative gate in both Sp and Lv (Sharma and Ettensohn, 2010). Other PMC genes are also expressed earlier than predicted and prior to Pmar, including Ets1/2, FoxO, and Tbr; these results were corroborated by qPCR analysis (Fig. 1 and not shown). However, the expression of their targets is delayed until after the double-negative gate has operated, and the time of onset for the targets agrees with previous results in Lv (Saunders and McClay, 2014), suggesting that additional temporal regulatory control is involved in regulating the PMC genes expressed at HB and later stages (Fig. 5I, red). Taken together, these results are consistent with largely similar GRNs driving specification in Lv, Sp, and Pl, with the exception of the earliest timepoints in the PMC network. Further, the results indicate that the specification network, rather than smoothly changing over developmental time, is instead relatively stable at most stages reflected in Figure 5, and is punctuated by an abrupt transition at HB stage, consistent with the PCA and *k*-means clustering results.

### Expression Analysis: The Metabolic Network Corroborates Expression Phases

The other major network operant during development is the metabolic network, which might be predicted to be more stable and, unlike the GRN, to not conform with the expression phases and transitions. To evaluate this, we used iPath 2.0 (Letunic et al., 2008; Yamada et al., 2011) to visualize expression changes in the metabolic enzyme network members during Lv development (Fig. 6). We determined the number of expression changes for 372 metabolic enzymes in the network between each pair of sequential stages in our initial assembly, and we found that the metabolic network is surprisingly dynamic. Since it is unclear what degree of change in expression is meaningful for metabolic genes, we considered cut-offs of both two-fold and four-fold. The results show that the transition from EB to HB has the maximum number of changes at both cut-offs, while the transition from LG to EP has the second largest number of changes at both cut-offs (Fig. 6A). These maxima correspond to the first two major transitions identified in the PCA, further corroborating it. The third major PC transition, between MB and EG, had a correspondingly large number of changes in the metabolic network at a two-fold, but not four-fold, cut-off. More than half of the total set of genes exhibited a two-fold change at HB, suggesting that this degree of change may be inconsequential. We visualized the network changes that occur during the first two major transitions using the Sp filter in iPath 2.0 (Fig. S9A), then mapped the edges that changed four-fold or more in the first two major PC transitions (Fig. 6B, C). We observed expression changes across the network, without an apparent concentration of changes in any particular network region. We also mapped expression levels across the network at each developmental timepoint (Fig. S9B-L). These metabolic maps show the temporal dynamics within the network during development, which are similarly diffuse throughout the network. It is difficult to interpret the precise repercussions of these changes on the behavior of the network as a whole, since many metabolic proteins are subject to post-translational modification, increasing the difficulties associated with network modeling (Fendt et al., 2010; Zelezniak et al., 2014); moreover, to our knowledge, metabolic flux level information and flux balance models are not currently available for developing Lv sea urchins. These results show that the major phase transitions can also be observed within the metabolic networks, and establish the dynamics of expression of the metabolic network in normally developing Lv embryos, providing a foundation for further studies. Metabolism is a largely untapped area in sea urchin development, and metabolic modeling is an interesting problem in this context, given the parallels in metabolism between early mammalian embryos and tumors (Smith and Sturmey, 2013), and because unlike the mammal, the sea urchin embryo is a nutritionally closed system.

**Figure 6.**
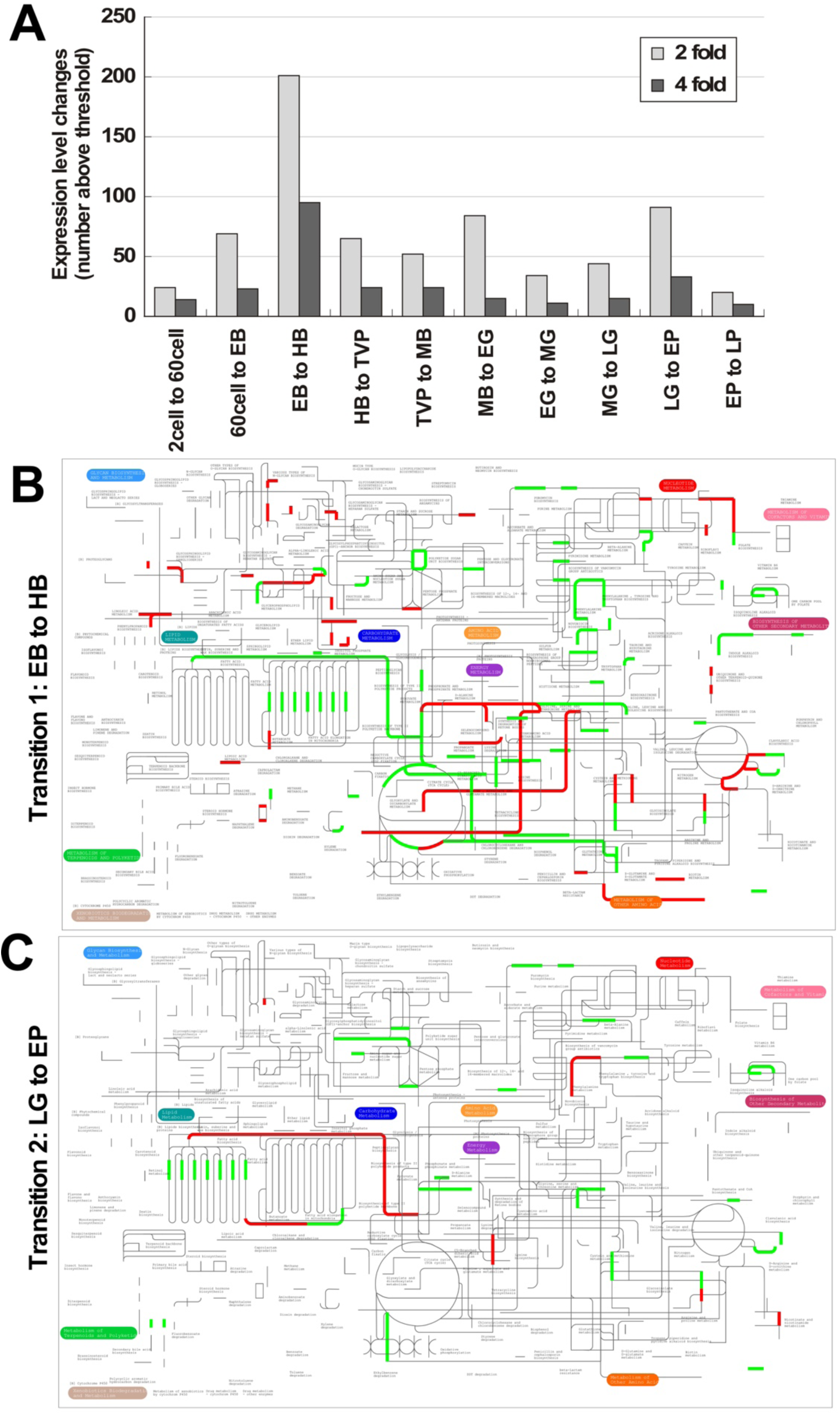
Metabolic gene expression changes coincide with expression phase transitions. Metabolic genes were analyzed using iPath, and expression changes between stages were determined for each metabolic gene. **A.** The number of changes in the expression levels of 372 genes encoding metabolic enzymes is shown at the indicated thresholds for each sequential transition in the dataset. **B., C.** The sea urchin metabolic network, with increases (green) and decreases (red) above a 4-fold threshold mapped onto the network for the first two phase transitions, EB to HB (B) and LG to EP (C). See also Fig. S9.

## Discussion

In this study, we compared the embryonic transcriptome at 11 distinct timepoints corresponding with major developmental events. We find that the timing of expression of known genes is compatible with established network models from other sea urchin species, both for particular network motifs, and for the known GRN genes in general. These results indicate that the GRN architecture is likely well-conserved among sea urchin species.

We used principal component analysis (PCA) to evaluate the gene expression variation during Lv development, and observed unexpectedly sharp transitions between specific developmental stages, with comparatively smaller variation between most others. These sharp transitions divide the sampled time course into four phases, and this result was corroborated by *k*-means clustering, gene regulatory network analysis, and metabolic network analysis, especially for the first major transition between early and hatched blastula. This transition in particular is reflected by a burst of rapid transcription, and likely reflects a transition between developmental milestones (Levin et al., 2012), although interspecies comparisons will be required to confirm that. Unfortunately, the currently available developmental transcriptome for Sp is sparsely sampled at early timepoints (Tu et al., 2012; Tu et al., 2014), precluding a direct comparison at this time.

The temporal non-smoothness of developmental gene expression has been observed in other model embryos, including zebrafish, mice, and pigs (Tang et al., 2011; Yang et al., 2013; Zhong et al., 2018). In the mouse embryo transcriptome, PCA analysis revealed one major transition that corresponds to the switch from maternal to zygotic gene expression, while the porcine transcriptome exhibits transitions in PCA space between 2-cell and 4-cell stages, and between 8-cell and morula stages (Tang et al., 2011; Zhong et al., 2018). In other embryonic transcriptomes, developmental gene expression dynamics analyzed with PCA exhibit smooth changes, including Arabidopsis, Maize, and *Drosophila* embryos (Cherbas et al., 2011; Chen et al., 2014; Hofmann et al., 2019). However, these studies did not consider transcriptional rates, and such rate differences may be present but not obvious from PCA results, as is the case for *C. elegans* and other nematode worms, whose developmental gene expression profiles exhibit smooth PCA trajectories, yet are punctuated by temporal gene expression rate changes (Levin et al., 2012). It will be of interest to learn whether the strongly punctuated changes in gene expression observed in Lv sea urchin embryos, and/or punctuated changes in transcriptional rates are a general feature of developing animal and plant embryos.

It is interesting that the non-ubiquitously expressed genes, whose overall transcriptional pattern is of moderate complexity based on *k*-means analyses, exhibit a collectively distinct expression profile, with a large number of stage-specific transcripts, a majority of phase-specific transcripts, and no transcripts that violate the expression phases. In contrast, the relatively low complexity ubiquitously expressed genes and the high complexity transcription factor subset each exhibit many fewer stage-specific and phase-specific transcripts, temporally broader gene expression in general, and a significant fraction of transcripts whose expression violates the expression phase boundaries. This fraction is largest for the TFs, suggesting that these genes in particular mediate integration across the expression phases.

The phase transitions from early blastula (EB) to hatched blastula (HB) and from late gastrula (LG) to early pluteus (EP) exhibit the largest variation, and this is reflected in the *k*-means clusters and the metabolic and gene regulatory networks. The first and largest transition, from EB to HB, ranks as the transition with the largest number of newly enriched GO terms, newly expressed transcripts, GRN gene onsets, and metabolic network gene expression changes. The second largest transition, from LG to EP, similarly ranks second for each of these metrics except new GO term enrichments and GRN gene onsets. In the latter case, this is likely because the GRN is too sparse at later stages to evaluate this transition (Fig. 5). Although the transition between LG and EP does not reflect a transcriptional burst, it nonetheless reflects a significant amount of gene expression change as evidenced by the large PC distance, *k*-means clustering and metabolic analyses. The transition from MB to EG is the third largest of the phase transitions, while EG to MG and MG to LG are also each relatively large; the stages in phase c exhibit more internal variation than within the other phases. Each of the transitions in phase c exhibits a large number of newly enriched GO terms, and the first two exhibit equivalent numbers of transcript onsets, while MB to EG specifically exhibits a comparatively large number of GRN gene onsets and metabolic network gene expression changes (with a 2-fold threshold). MB to EG is the second transcriptional burst (Fig. 3B), implying that the transition to EG is from a second developmental milestone in sea urchin embryos.

Hatching is a significant event in the embryonic life cycle that is mediated by the secretion of hatching enzyme that proteolytically degrades the fertilization envelope (Roe and Lennarz, 1990). Hatching is a critical event in the lifecycle of embryos in general, with both costs and benefits (Warkentin, 2011b, a). The timing of hatching is often plastic in response to the environment, and in many instances, embryos that delay hatching itself otherwise develop on schedule (Warkentin, 2011b, a); this is the case for sand dollar embryos (Armstrong et al., 2013), and probably for echinoderm embryos in general, implying an uncoupling between the expression of hatching enzyme and the deployment of the specification GRNs.

In sea urchins, hatching marks the expression of numerous genes, including Pmar target genes within the PMCs, the targets of the Otx/GataE/Blimp lock-down loop in the endoderm, GCM targets in the SMCs, Eve targets in the posteriolateral ectoderm (which are likely involved in instructive signaling to PMCs), the targets of Nodal signals that comprise the initial ventral ectoderm specification TFs, the onset of dorsal ectoderm specification signaling via BMP2/4 and Chd expression, and neural Six3 expression (Fig. 5). Together, these changes correspond to increasingly well-defined specification states across all the major tissues in the embryo. There is a strong increase in the rate of gene expression for this transition reflecting a burst of transcription.

In contrast, the second transcriptional burst, between MB and EG, probably primarily reflects the morphogenetic changes that drive gastrulation, with specification state changes providing comparatively smaller contributions; for example, the DV axis is committed by EG stage in sea urchins (Hardin et al., 1992; Hardin and Armstrong, 1997; Bradham and McClay, 2006; Piacentino et al., 2015). However, left-right specification remains incomplete at EG, as does regional specification of the gut (Annunziata and Arnone, 2014; Piacentino et al., 2016a). Intriguingly, very few new transcripts are expressed at MB, when the PMCs undergo an epithelial-mesenchymal transition (EMT) and ingress into the blastocoel. These results suggest that the EMT that produces the PMCs is primarily driven by post-transcriptional regulation rather than by new gene expression, while the invagination of the gut involves comparatively more transcriptional regulation.

Together, these results demonstrate that sea urchin developmental gene expression changes are comparatively small between most contiguous stages, with exceptions corresponding to major transcriptional bursts between early and hatched blastula, and mesenchyme blastula and early gastrula. These findings underscore the modularity of development, which is especially pronounced in this echinoderm model system. It will be of interest to extend these studies to other echinoderm embryos, and to determine whether chromatin state changes accompany and underlie these major transitions.

## Methods

### RNA-seq, *de novo* assembly, and analysis

*L. variegatus* total RNA was prepared from 1 × 10^6^ control embryos per timepoint using TRIzol (Invitrogen) and DNase treatment, along with an additional six samples from embryos treated with the perturbants nickel chloride (Sigma) or SB203580 (Calbiochem) and collected at EG, MG, and LG. Data associated with the latter samples were described previously (Piacentino et al., 2016b) and are excluded from further analysis here; however, those transcripts were included in the assembly pipeline herein. RNA quantitation and integrity were determined using a Qubit^®^ 2.0 Fluorometer (Life Technologies) and a 2100 Bioanalyzer (Agilent Technologies). Total RNA was subjected to three iterations of polyA selection using Dynabeads (Life Technologies) prior to cDNA synthesis. Sequencing libraries were prepared from 1 µg of size-selected cDNA for standard Illumina paired-end sequencing. The average insert size of the gastrula stage libraries was ~180 bp, and was ~280 bp for all others. These smaller insert sizes aid in preventing chimeric assembly products (Xie et al., 2014). EG, MG, and LG samples (both control and treated) were initially sequenced on an Illumina GAII platform (Morozova et al., 2009); the remaining eight control samples were sequenced using the HiSeq Illumina platform at a later time. In both cases, 101 bp paired-end reads were obtained. Biological replicates were sequenced using HiSeq4000 with barcoding; the average insert size was 200 bp, and 101 bp paired-ends reads were generated and adaptor- and quality-trimmed (BGI, Inc.). Prior to *de novo* sequence assembly, a custom Python script was used to trim raw Illumina reads of adapter sequences (on average 1-3%) and low quality reads (Phred score below 11). Reads containing Ns were excluded. An average of 10% of the sequences were excluded by this procedure. Overlapping reads were joined into longer reads and PCR duplicates were excluded. Approximately 1.5 billion reads were used for *de novo* assemblies simultaneously. The final assembly was generated using SOAPdenovo-Trans, which avoids chimeric assembly artifacts by requiring a minimum of three read pairs to define the distance and order between adjacent contigs (Xie et al., 2014). Settings (other than default) used were K31, M3, F and G200. Per default, up to five transcripts per locus were allowed. Assembled reads shorter than 100 bp were excluded. Reads were mapped to the assembly with Bowtie2 using the argument k=20 and otherwise default parameter settings. Count values were generated using a custom Python script, and were initially normalized using DESeq (Anders and Huber, 2010), then with the quantile normalization package in R (Hansen and Irizarry, 2012). Quantile-normalized values were used for all subsequent analyses. Annotation was performed using BLASTx (Altschul et al., 1990) against the *S. purpuratus* predicted protein database, using a cut-off of e = 1 × 10^−7^; these annotations were supplemented by BLASTx against the nr database on NCBI. GO terms and Enzyme Commission (EC) numbers (for iPath2 analyses) were assigned using BLAST2GO (Conesa et al., 2005), and Pfam domains assigned using HMMer (Eddy, 1998; Sammut et al., 2008). GO enrichment analysis was performed using iPAGE (Goodarzi et al., 2009). Metabolic analysis was performed using iPath (Yamada et al., 2011).The results from the sequencing and assembly can be accessed at NCBI (BioProject accession number PRJNA241187) and at https://lvedge.bu.edu.

### qPCR analysis

qPCRs were performed as described (Bradham and McClay, 2006), except that gene expression measurements were normalized to Lv-Setmar. All qPCR analyses were performed on three independent biological replicates, in triplicate. qPCR primer sequences are provided in Table S1.

### PCA and *k*-means clustering

Principal components were identified using the prcomp function from the R package stats (R Core Team, 2014), and plotted with the R package scatterplot3d (Ligges and Mächler, 2003). *K*-means cluster analyses were performed using Cluster 3.0 (http://bonsai.hgc.jp/~mdehoon/software/cluster/software.htm#ctv) across a range of *k* values, and optimal *k* values were selected based on manual inspection for cluster uniqueness and uniformity. Final clusters were the optimal solution from 5,000 trials. Heat maps were generated using JavaTreeView (Saldanha, 2004) and manually organized in Canvas (ACD Systems).

### Database

An online database that provides access to the RNA-Seq data herein was generated using a Python-based interface with a MySQL database, and was named LvEDGEdb (Fig. S3). It is accessible at https://lvedge.bu.edu. The database is searchable and provides gene expression results graphically or numerically, as well as GO terms and Pfam domains, and sequences as fasta files. The database is also searchable by BLAST, using a ViroBLAST interface (Deng et al., 2007). Registered database users can contribute new or revised annotations.

## Acknowledgements

We thank Adrian Reich (Brown University) and Smadar Ben Tabou de Leon (University of Haifa) for helpful discussions, Keith Bradnam (UC Davis) for the CEGMA analysis, Shile Zhang (Boston University) for suggesting the PCA, and Gary Benson (Boston University) for organizing the challenge project and database courses at Boston University, whose students (J.D.H., J.L.K., L.L.) contributed to the bioinformatics analyses and database construction herein. This study was supported by start-up funds from Boston University (CAB) and by NSF OIS 1257825 and 1656752 (CAB); CB was supported by the RISE program at Boston University.

## Author Contributions

The study was conceived by CAB

The sequencing samples were collected by ABC and DS, then sequenced, assembled and annotated by BT, AJP, JDH, JLK, ES, JHG. JI-S, and NI

Bioinformatics analyses and database construction were performed by JDH, JLK, LL, DYH, AL, CB, JHG, JI-S, ES, MA and CAB

Biological analyses were performed by MLP, DS, and DTZ The manuscript was written by CAB and edited by

## Supplemental Figure Legends

**Figure S1.** Box-and-whisker plots depicting the range of expression values per developmental stage, with DESeq normalization (**A**) or quantile normalization (**B**). See also Figure 1.

**Figure S2. Lv-Setmar expression has low variation over developmental time. A.** A plot of Lv-setmar versus Lv-ubiquitin expression over time demonstrates that Lv-setmar exhibits less temporal variation than Lv-ubiquitin. **B.** A representative gel showing Lv-setmar qPCR products amplified from cDNAs representing each sequenced stage in this study, demonstrating comparable product levels and an absence of spurious amplification products. See also Figure 1E.

**Figure S3. LvEDGE database.** Screen shots showing the home page (**A**), the search window (**B**), an example search with a temporal expression plot (**C**), and the numerical data reflected in the plot (**D**) for the LvEDGE public database, which hosts the data described herein.

**Figure S4. GO term enrichment is shown at each sequenced developmental stage.** 96 GO terms identified as enriched in one or more of the sequenced stages were sorted into nine categories (various colors), and are shown here for each instance of enrichment relative to the adjacent timepoints. See also Fig. 1D and Table S3.

**Figure S5. GO term enrichment among expression phases.** Only transcripts whose expression is limited to a specific expression phase were considered for this analysis. The results are shown as an enrichment heat map. See also Table S4.

**Figure S6. *k*-means cluster analysis of the non-ubiquitous transcriptome reveals complete adherence to the expression phases.** *K*-means clustering (*k*=19) was performed on annotated sequences whose expression is ≤ 1% of the maximum expression level per gene for at least one timepoint (n = 4792 sequences). Left: A heat map depicts the clusters; bright red is maximal expression. Right: Plots of the average normalized expression values for each cluster. Expression phase boundaries are indicated by vertical dashed lines. Clusters a2, b1, c1, d3 and e2 are also shown in Figure 4. See also Table S5.

**Figure S7. Cluster analysis of the ubiquitous transcriptome reveals clusters that do not adhere to the expression phases.** *K*-means clustering (*k*=37) was performed on annotated sequences whose expression is > 1% of the maximum expression level for all timepoints (n = 14,773 sequences). Left: A heat map depicts the clusters; bright red is maximal expression. Right: Plots of the average normalized expression values for each cluster. Expression phase boundaries are indicated by vertical dashed lines. Cluster f1 is also shown in Figure 4. See also Table S6.

**Figure S8. Cluster analysis of the transcription factors reveals that most are expressed in three or more sequential stages.** *K*-means clustering (*k* = 23) was performed on all the identified transcription factors in the Lv transcriptome (n = 521). Left: A heat map depicts the clusters; bright red is maximum expression. Right: Plots of the average normalized expression value for each cluster. Phase boundaries are indicated by vertical dashed lines. See also Figure 4 and Table S7.

**Figure S9. Temporal metabolic network analysis. A.** The *S. purpuratus*-specific network, produced by the built-in filter in iPath. **B.-L.** The metabolic network expressed in each indicated developmental stage. For these maps, zero expression is depicted in black; expression from 0 – 100 is in light blue, 100 – 1000 is in medium blue, and >1000 is in dark blue. Thin gray lines indicate edges expected to be present based on the filter, but for which expression was not detected in any developmental stage. See also Figure 6.

