## Supplemental Figures and Tables for "The Developmental Transcriptome for *Lytechinus variegatus* Exhibits Temporally Punctuated Gene Expression Changes"

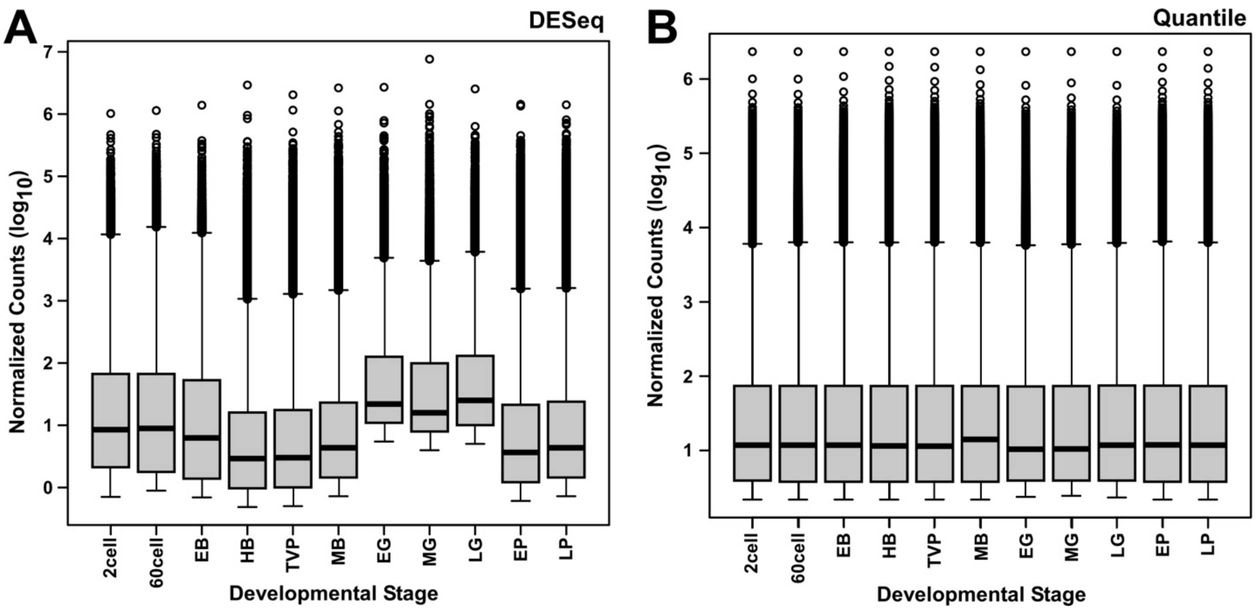

**Figure S1.** Box-and-whisker plots depicting the range of expression values per developmental stage, with DESeq normalization (**A**) or quantile normalization (**B**). See also Figure 1.

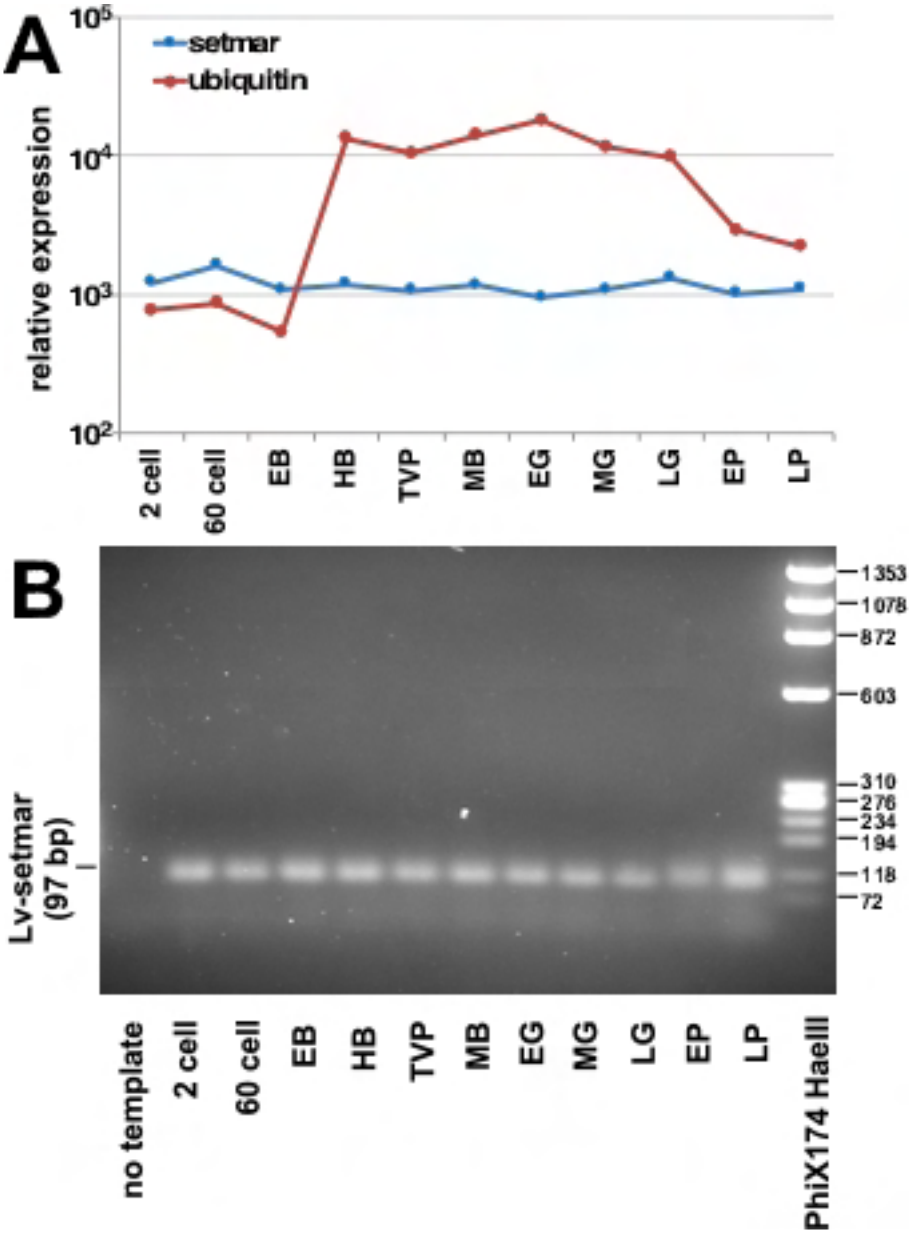

**Figure S2. Lv-Setmar expression has low variation over developmental time. A.** A plot of Lv-setmar versus Lv-ubiquitin expression over time demonstrates that Lv-setmar exhibits less temporal variation than Lv-ubiquitin. **B.** A representative gel showing Lv-setmar qPCR products amplified from cDNAs representing each sequenced stage in this study, demonstrating comparable product levels and an absence of spurious amplification products. See also Figure 1E.

**A**

Home Search BLAST Add Annotations Admin Change Password Help Create Account Login Logged in as **guest**.

**Welcome to LvEDGEdb!**

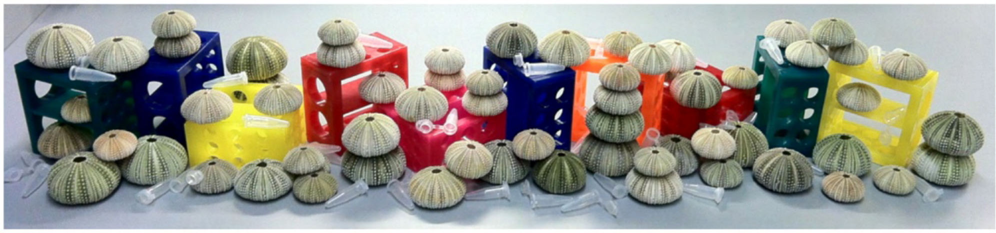

**B**

Home Search BLAST Add Annotations Admin Change Password Help Create Account Login Logged in as **guest**.

**Search by (results will match any fields entered):**

| | | |
|---|---|---|
| Gene Name | | (e.g. LvNodal) |
| Keyword in Description | | (e.g. ATPase) |
| Contig or Scaffold Number (Nov. 2012 assembly) | | (e.g. C1710637, scaffold18) |
| SPU ID | | (e.g. SPU_003582) |
| GO ID | | (e.g. GO:0000030) |
| Pfam Domain | | (e.g. HCR) |

Submit

**C**

**Displaying search results for:**
Gene Name: BMP2/4

| Gene Name | Description | Annotated By | Sequence | Graph | Embryonic Development Data (Help!) | Screen Data (Help!) | SPU ID# | Best BLAST Hit GenBank ID | Pfam Domain | GO Ids |
|---|---|---|---|---|---|---|---|---|---|---|
| LvBmp2/4 | none | albert | fasta | Show Graph | Link | Link | SPU_000669 | AAD28038 | TGFb_propeptide, | GO:0005576, GO:0008083, GO:0040007, |

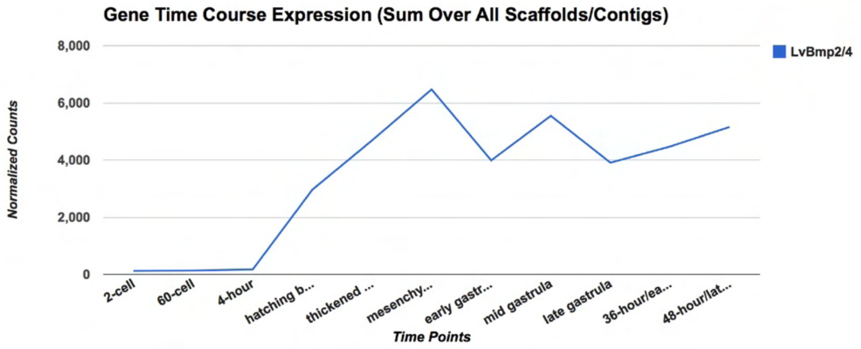

**D**

| singleton | 2-cell | 60-cell | 4-hour | hatching blastula | thickened vegetal plate | mesenchyme blastula | early gastrula | mid gastrula | late gastrula | 36-hour/early pluteus | 48-hour/late pluteus |
|---|---|---|---|---|---|---|---|---|---|---|---|
| scaffold8980 | 119.562 | 134.514 | 170.867 | 2961.72 | 4682.66 | 6467.62 | 3985.68 | 5542.98 | 3906.14 | 4469.73 | 5154.73 |

Click here for raw text file

**Figure S3. LvEDGE database.** Screen shots showing the home page (**A**), the search window (**B**), an example search with a temporal expression plot (**C**), and the numerical data reflected in the plot (**D**) for the LvEDGE public database, which hosts the data described herein.

| | | stage | | | | | | | | | | |
|---|---|---|---|---|---|---|---|---|---|---|---|---|
| | | 1 | 2 | 3 | 4 | 5 | 6 | 7 | 8 | 9 | 10 | 11 |
| Category | Subcategory | 2-cell | 60-cell | EB | HB | TVP | MB | EG | MG | LG | EP | LP |
| Cell Division | | | | Cytokinesis | Mitosis | | checkpoint | cell division | | | meiotic recombination | |
| | | | | | | | | | | | cell cycle | |
| Development | | | stem cell maintenance | | | | left-right asymmetry | | cell morphogenesis | | left-right asymmetry | |
| | | | | | | | regulation of multicellular organismal process | | | | cell soma | cell soma |
| Gene Expression Control | Chromatin | | | chromatin modification | | | | | | chromatin binding | | SWI/SNF complex |
| | | | | methylated histone Binding | | | | | | | | |
| | DNA binding | | | sequence-specific DNA binding | sequence-specific DNA binding | sequence-specific DNA binding | negative regulation of transcription factor activity | | sequence-specific DNA binding | sequence-specific DNA binding | | |
| | Processing | | | nuclear mRNA splicing, via spliceosome | nuclear mRNA splicing, via spliceosome | | | nuclear mRNA splicing, via spliceosome | nuclear mRNA splicing, via spliceosome | nuclear mRNA splicing, via spliceosome | nuclear mRNA splicing, via spliceosome | |
| | | | | | mRNA processing | | | | | | | |
| | Translation | | | | translation initiation factor activity | translation initiation factor activity | | | | translation initiation factor activity | translation initiation factor activity | |
| Metabolic | | fatty acid metabolic process | NADH dehydrogenase (ubiquinone) activity | | cell redox homeostasis | cell redox homeostasis | primary metabolic process | mitochondrial inner membrane | mitochondrial matrix | glycosaminoglycan biosynthetic process | ATP synthesis coupled proton transport | |
| | | | | | oxidoreductase activity, acting on paired donors, with incorporation or reduction of molecular oxygen | monooxygenase activity | monooxygenase activity | | oxidoreductase activity | positive regulation of cellular metabolic process | oxidoreductase activity | |
| | | | | | sulfuric ester hydrolase activity | ATP catabolic process | transferase activity, transferring hexosyl groups | | oxidoreductase activity, acting on the aldehyde or oxo group of donors, NAD or NADP as acceptor | | | |
| | | | | | ATP synthesis coupled proton transport | | | | | | | |
| | | | | | DNA Metabolic Process | | | | | | | |
| Regulation | cytoskeleton | | | | ciliary or flagellar motility | | microtubule motor activity | | | | actin binding | |
| | protease | metallo-endopeptidase activity | metallo-endopeptidase activity | | serine-type endopeptidase activity | serine-type endopeptidase activity | threonine endopeptidase activity | threonine endopeptidase activity | metallo-endopeptidase activity | metallo-endopeptidase activity | serine-type peptidase activity | |
| | | | | | peptidase activity | metallopeptidase activity | metallopeptidase activity | | | | | |
| | | | | | | | platelet activation | platelet activation | | | | |
| | other | | | | | | | identical protein binding | | protein ubiquitination | protein polyubiquitination | |
| | | | | | | | | regulation of cellular process | | positive regulation of cellular metabolic process | | |
| Signaling | | G-protein coupled receptor activity | G-protein coupled receptor activity | G-protein coupled receptor activity | G-protein coupled receptor activity | G-protein coupled receptor activity | G-protein coupled receptor activity | G-protein coupled receptor activity | G-protein coupled receptor activity | G-protein coupled receptor activity | G-protein coupled receptor activity | G-protein coupled receptor activity |
| | | | | phosphoinositide phospholipase C activity | neuropeptide Y receptor activity | neuropeptide Y receptor activity | neuropeptide Y receptor activity | | cGMP biosynthetic process | | | regulation of apoptosis |
| | | | | non-membrane spanning protein tyrosine kinase activity | scavenger receptor activity | scavenger receptor activity | | | | | receptor binding | MAPKKK cascade |
| | | | | growth factor activity | | small GTPase mediated signal transduction | | | | | small GTPase mediated signal transduction | small GTPase mediated signal transduction |
| | | | | positive regulation of GTPase activity | | | | | | | | |
| Structural | cytoskeleton | axonemal dynein complex | microtubule-based process | dendritic shaft | | | | | | dendritic spine | microtubule | |
| | | cytoskeletal part | | | | | | | | | | |
| | | sarcomere organization | | | | | | | | | | |
| | organelle | | nucleosome assembly | nucleosome | nucleosome | | | | nucleosome | nucleosome | nuclear envelope | |
| | | | | integral to Golgi membrane | | | | intracellular organelle part | intracellular organelle | | | |
| | | | structural constituent of ribosome | structural constituent of ribosome | structural constituent of ribosome | structural constituent of ribosome | structural constituent of ribosome | structural constituent of ribosome | structural constituent of ribosome | ribosome | structural constituent of ribosome | ribosome |
| | other | | | cytoplasmic part | protein polymerization | | Proteinaceous extracellular matrix | intracellular part | intracellular part | | Extracellular region part | |
| | | | | | | | Extracellular region | | | | | |

**Figure S4. GO term enrichment is shown at each sequenced developmental stage.** 96 GO terms identified as enriched in one or more of the sequenced stages were sorted into nine categories (various colors), and are shown here for each instance of enrichment relative to the adjacent timepoints. See also Fig. 1D and Table S3.

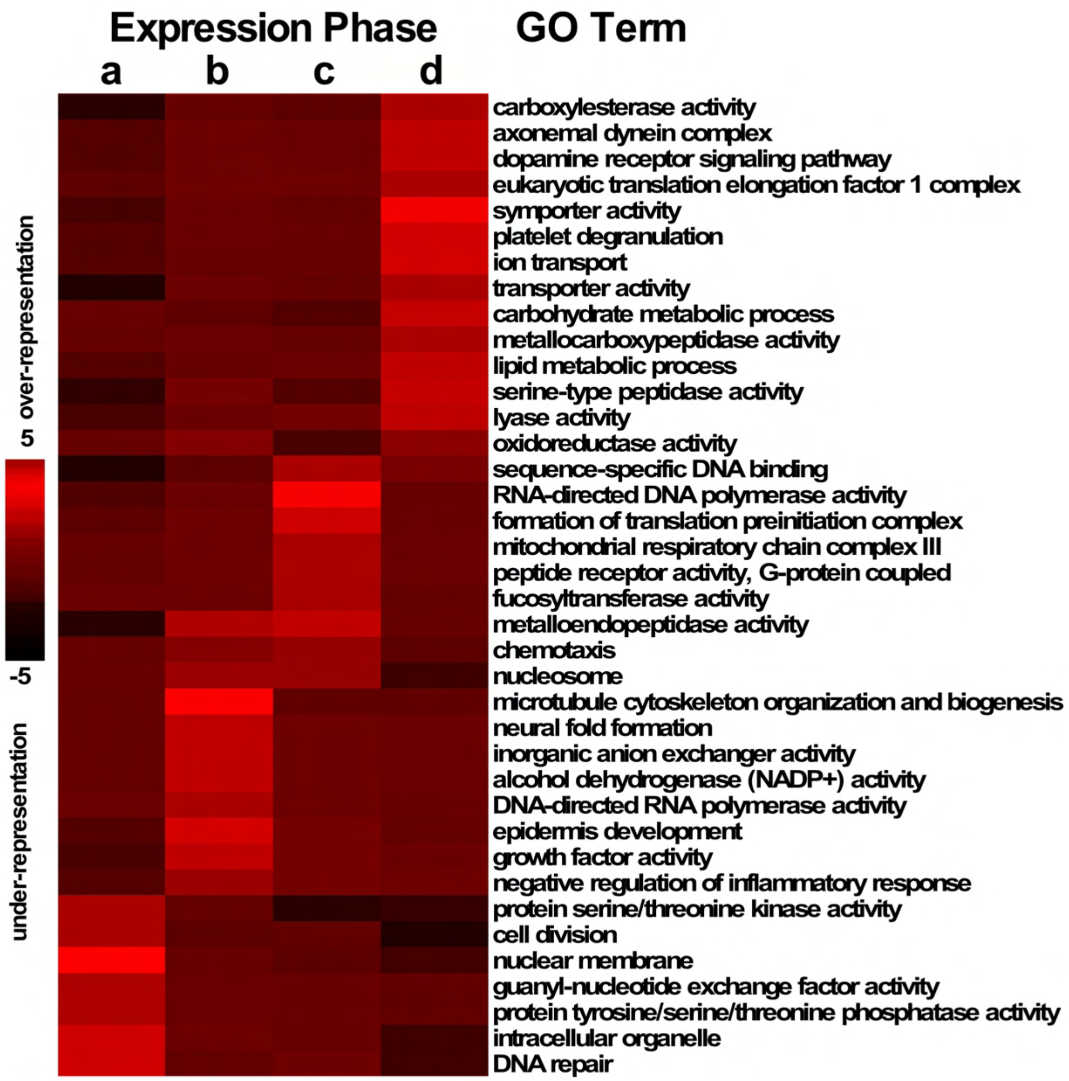

**Figure S5. GO term enrichment among expression phases.** Only transcripts whose expression is limited to a specific expression phase were considered for this analysis. The results are shown as an enrichment heat map. See also Table S4.

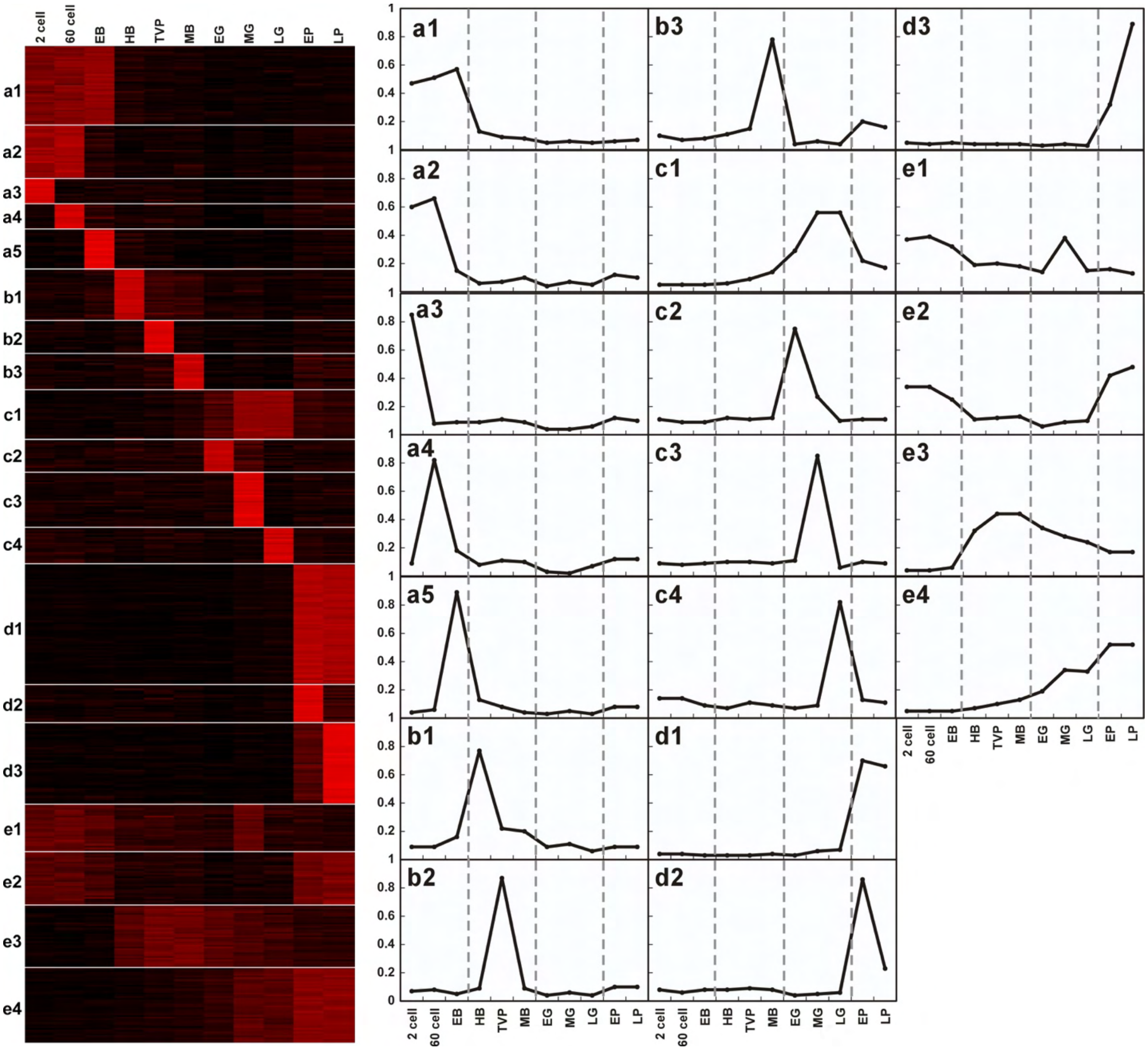

**Figure S6. *k*-means cluster analysis of the non-ubiquitous transcriptome reveals complete adherence to the expression phases.** *K*-means clustering (*k*=19) was performed on annotated sequences whose expression is ≤ 1% of the maximum expression level per gene for at least one timepoint (n = 4792 sequences). Left: A heat map depicts the clusters; bright red is maximal expression. Right: Plots of the average normalized expression values for each cluster. Expression phase boundaries are indicated by vertical dashed lines. Clusters a2, b1, c1, d3 and e2 are also shown in Figure 4. See also Table S5.

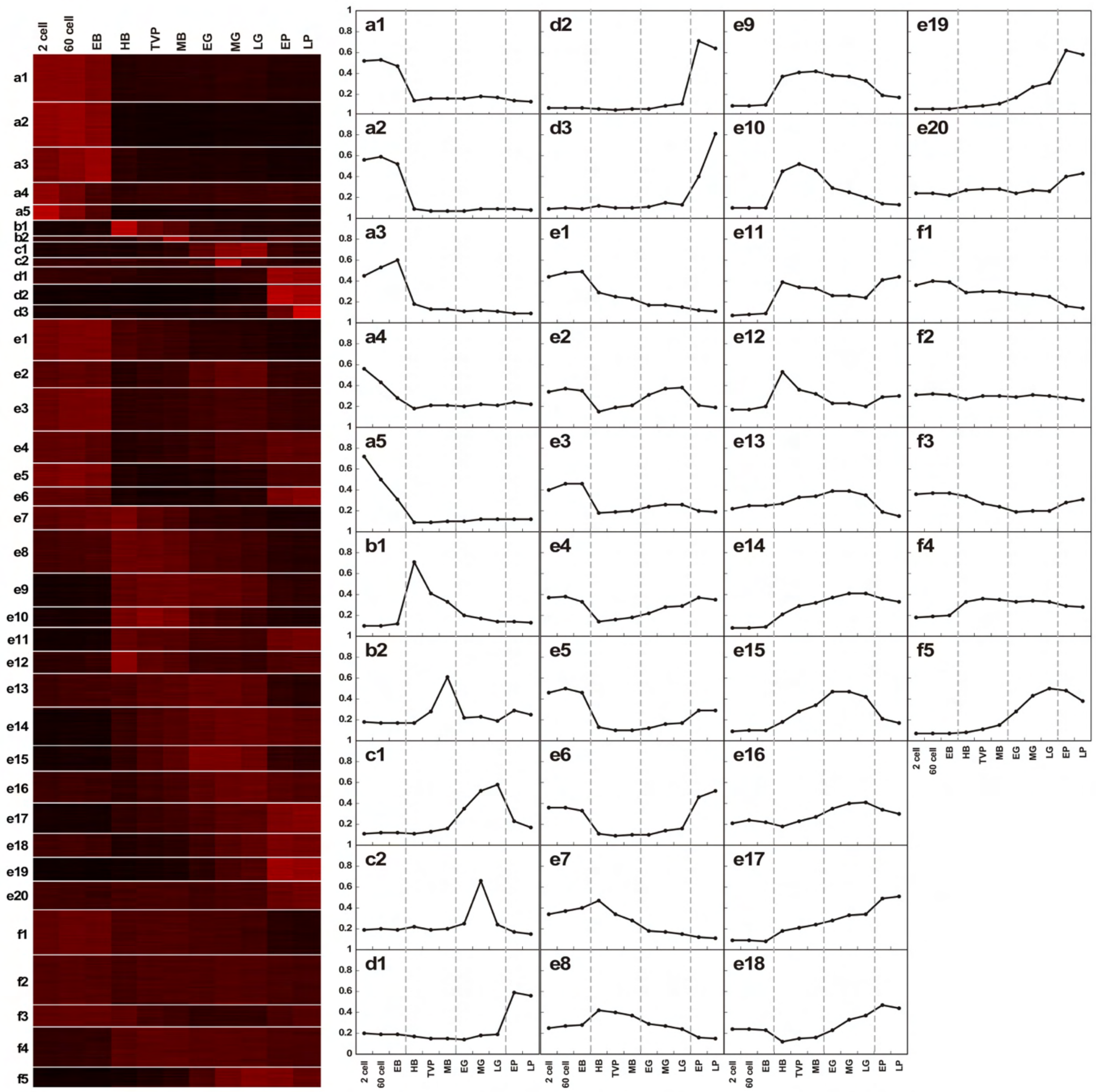

**Figure S7. Cluster analysis of the ubiquitous transcriptome reveals clusters that do not adhere to the expression phases.** *K*-means clustering (*k*=37) was performed on annotated sequences whose expression is > 1% of the maximum expression level for all timepoints (n = 14,773 sequences). Left: A heat map depicts the clusters; bright red is maximal expression. Right: Plots of the average normalized expression values for each cluster. Expression phase boundaries are indicated by vertical dashed lines. Cluster f1 is also shown in Figure 4. See also Table S6.

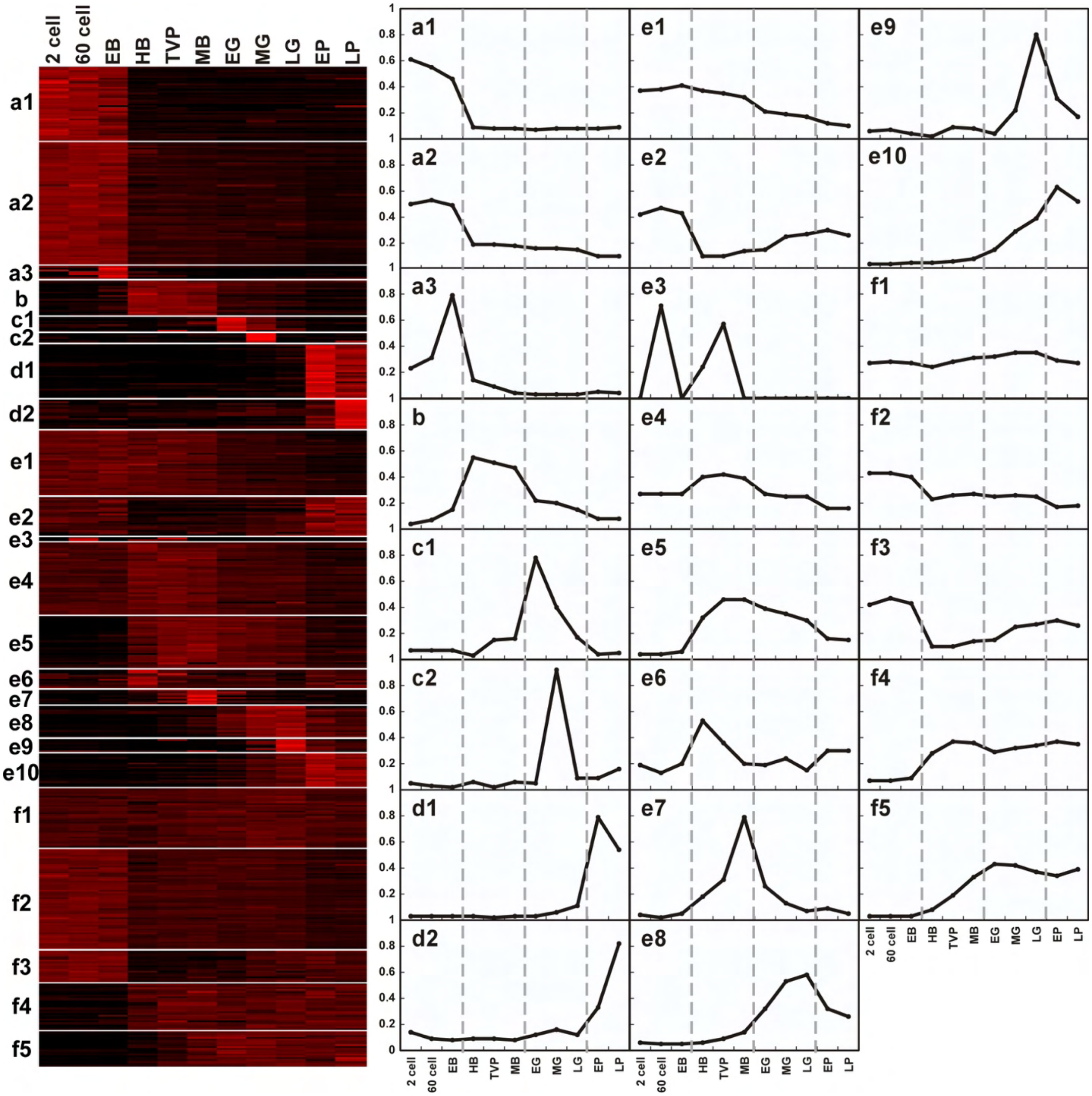

**Figure S8. Cluster analysis of the transcription factors reveals that most are expressed in three or more sequential stages.** *K*-means clustering ($k$ = 23) was performed on all the identified transcription factors in the Lv transcriptome (n = 521). Left: A heat map depicts the clusters; bright red is maximum expression. Right: Plots of the average normalized expression value for each cluster. Phase boundaries are indicated by vertical dashed lines. See also Figure 4 and Table S7.

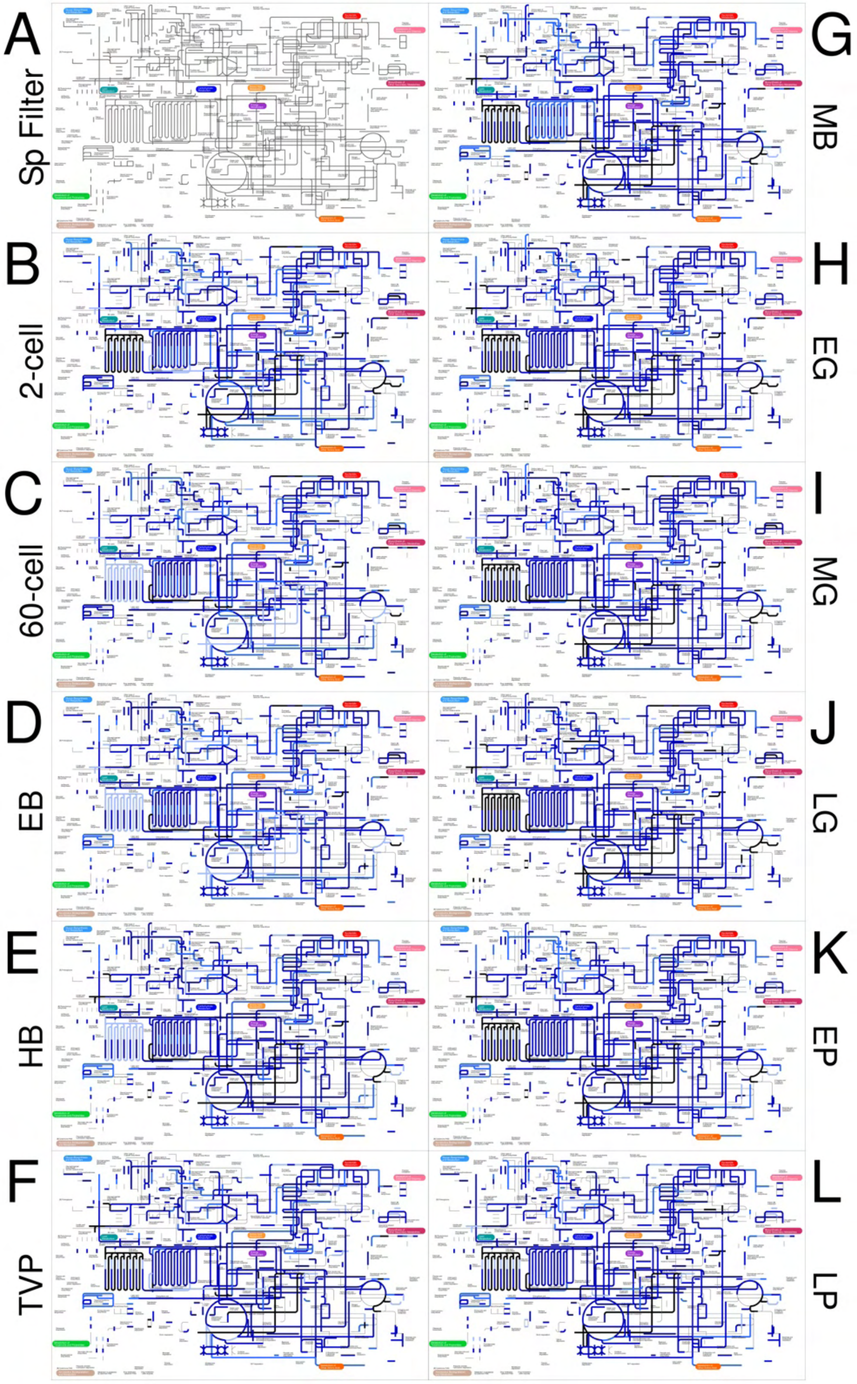

**Figure S9. Temporal metabolic network analysis. A.** The *S. purpuratus*-specific network, produced by the built-in filter in iPath. **B.-L.** The metabolic network expressed in each indicated developmental stage. For these maps, zero expression is depicted in black; expression from 0 – 100 is in light blue, 100 – 1000 is in medium blue, and >1000 is in dark blue. Thin gray lines indicate edges expected to be present based on the filter, but for which expression was not detected in any developmental stage. See also Figure 6.

### Table S1. qPCR Primer Sequences

| Gene | 5' Primer (5' -> 3') | 3' Primer (5' -> 3') |
|---|---|---|
| LvAlx1 | CCGGTACCAGTACCACCAAC | AGGATGGGAGTGTTGACTGC |
| LvBlimp | GACGATGAGGACGAGTTGGT | CGGTTTCCATTGCATCTTCT |
| LvBMP2/4 | CCATCAGGAGCTATCATCACG | ATCACTGTGTGTCTGTGGTG |
| LvBMP5-8 | TACCCTCCACAACGATACACCC | GCCTTTAGCAACGGACCATACA |
| LvBrachyury | CAGGCTAGAGGACGTGGAAC | CCACTCCCCATTGACGATCT |
| LvChordin | CCCACTCAAGGCAGAGATAC | GAAGGTACACCCTGGAACAC |
| LvDelta | CTGCCAAAACAACATCAACG | AAACCTCCCATCACATCTCG |
| LvEndo16 | CGGATCGGAATCAGAGTCAT | AGCAGCATCCAACTCCAAAT |
| LvErg | GGACACTTGCTAACCCGGTA | TTCTCCCCATCTTCTGGCTA |
| LvEts1 | CGGAGGCATTGAGAATCAAG | TAGGCATGCGTTCATCAGAG |
| LvEve | CATCCCCTATCACGATCACC | TGTGGAGGCCAAACTCTAGG |
| LvFoxG | GAGCGATACAGAGCCGGAGA | CTGAAGGGAGGTTTTTCACC |
| LvFoxO | AGCAATAGTTCAGCGGGTTG | ACTCTCCCTCGCTTCCTCTC |
| LvFoxQ2 | GAGAAGCGTCTATTGCTGTGTG | ACGAATACTGTTCCTCCAGCTC |
| LvGataC | GGGCCGAGAATGTGTAAACT | AAGCGGTCTGTTCTGTCCAT |
| LvGataE | TAAACCTCAGCGCCAATACC | CATTCCGATTCTGCTTTGCT |
| LvGcm | TCCAAAGACACCTGTCTACCAG | GGAGAAGCACTGGAACAAAGAC |
| LvGoosecoid | TCACCGAAGAACAACTGGAA | TAAGCCTCACATGCCCTCTT |
| LvHh | GGGAATCAAACTTCGACCAA | AGATCCTTGCCTGGTCACTG |
| LvHnf6 | GGGAGGAGATCAACACGAAA | CTGCCGCTCTTGAGCTTACT |
| LvHox11/13b | ACCTCATCCACCACCTCATC | GAAGTTGCGATTGATGACCA |
| LvIrxA | CCAGGAATCGATGCGGTGATGG | CATCTCGATGATCACGGTGGTGG |
| LvNodal | CCATGTTCATGGTGGAATTG | GAATGTCGGGAAGGAGGTCT |
| LvOtp | CCCAACTCAACGAACTGGAG | GAGGCCGTGAGATGGTAGAA |
| LvOtx | GCCAAACGTTGAACTCATGC | AGCTGAGCTCGGGTAAAAGTC |
| LvPtc | AGGTTGTCGTGGAAACCAGT | GTGGGGGTACTGATGGTGAG |
| LvSetmar[1] | GCCATCATGTCCTTGTCTCA | CACATGAAGCTTGATCAGGTAGTC |
| LvSix1/2 | CCCACAACCCTTACCCTTCT | TTCCCATTCATAGGCAGGTC |
| LvSix3 | GGCTCAAGCCACAGGACTAA | GGGTTGGGACTGAGTAAGCA |
| LvSM30 | CCACAGCTTCTGCAACAACA | AGGCTGGATGGCGTAGTATG |
| LvSmo | GATGCTCCTGGTATGGTTGG | GTGCTCAATCCCGCTATCAC |
| LvSnail | CCGAGGTCTTTTCTCGTCAA | ACTTGCCACTGACACAACACC |
| LvSoxE | TCAGTATCCTCAGCTCCACAAC | GATGAAGGGCTGCTTCTCCT |
| LvTbrain | CTCTCTCACCAGACTGCTCACT | GTGGTGGAATGTGGGGAATC |
| LvTbx2/3 | CCCATGGAAACACAGAGC | TGACCATTTCCGTTCCTC |
| LvWnt5a | CGAGCTCTTCATCCTTGGTACA | TGGTCCTGGTAGAGCTGACATA |
| LvWnt8 | AACATGCACAACAACGAAGC | GACGCGTACTGCCTTGAAGT |

[1]normalization gene

**Table S2. Principal components corresponding to Fig. 3.**
Averaged PCA values, with the major new transitions for each PC indicated with boxes.

| stage | PC1 | PC2 | PC3 |
|---|---|---|---|
| 2c | -141.2 | 24.82 | 1.47 |
| 60c | -142.71 | 15.77 | -2.85 |
| EB | -133.55 | -6.06 | -6.04 |
| HB | 28.06 | -116.22 | 24.24 |
| TVP | 41.13 | -91.52 | 19.03 |
| MB | 46.62 | -73.54 | 14.27 |
| EG | 54.91 | -36.05 | -20.23 |
| MG | 58.30 | -4.18 | -41.77 |
| LG | 55.75 | 17.22 | -22.61 |
| EP | 64.54 | 124.57 | 15.04 |
| LP | 67.14 | 143.11 | 19.49 |

**Table S3. GO term enrichment by stage**

| stage of enrichment | Category | Subcategory | GO term |
|---|---|---|---|
| 1 | metabolic | | GO:0006631 fatty acid metabolic process |
| 1 | regulation | protease | GO:0004222 metalloendopeptidase activity |
| 1 | signaling | | GO:0004930 G-protein coupled receptor activity |
| 1 | structural | cytoskeleton | GO:0005858 axonemal dynein complex |
| 1 | structural | cytoskeleton | GO:0044430 cytoskeletal part |
| 1 | structural | cytoskeleton | GO:0045214 sarcomere organization |
| 2 | developmental | | GO:0019827 stem cell maintenance |
| 2 | metabolic | | GO:0008137 NADH dehydrogenase (ubiquinone) activity |
| 2 | structural | cytoskeleton | GO:0007017 microtubule-based process |
| 2 | structural | organelle | GO:0003735 structural constituent of ribosome |
| 2 | structural | organelle | GO:0006334 nucleosome assembly |
| 2 | transport | protein | GO:0006886 intracellular protein transport |
| 2 | transport | protein | GO:0015031 protein transport |
| 3 | cell division | | GO:0000910 cytokinesis |
| 3 | gene expression | chromatin | GO:0016568 chromatin modification |
| 3 | gene expression | chromatin | GO:0035064 methylated histone residue binding |
| 3 | gene expression | DNA binding | GO:0043565 sequence-specific DNA binding |
| 3 | gene expression | processing | GO:0000398 nuclear mRNA splicing, via spliceosome |
| 3 | signaling | | GO:0004435 phosphoinositide phospholipase C activity |
| 3 | signaling | | GO:0004715 non-membrane spanning protein tyrosine kina[illegible] |
| 3 | signaling | | GO:0008083 growth factor activity |
| 3 | signaling | | GO:0043547 positive regulation of GTPase activity |
| 3 | structural | cytoskeleton | GO:0043198 dendritic shaft |
| 3 | structural | cytosol | GO:0044444 cytoplasmic part |
| 3 | structural | organelle | GO:0000786 nucleosome |
| 3 | structural | organelle | GO:0030173 integral to Golgi membrane |
| 4 | cell division | | GO:0007067 mitosis |
| 4 | gene expression | processing | GO:0006397 mRNA processing |
| 4 | gene expression | splicing | GO:0000398 nuclear mRNA splicing, via spliceosome |
| 4 | gene expression | translation | GO:0003743 translation initiation factor activity |
| 4 | metabolic | | GO:0006259 DNA metabolic process |
| 4 | metabolic | | GO:0008484 sulfuric ester hydrolase activity |
| 4 | metabolic | | GO:0015986 ATP synthesis coupled proton transport |
| 4 | metabolic | | GO:0016705 oxidoreductase activity, acting on paired donor[illegible] oxygen |

| | | | |
|---|---|---|---|
| 4 | metabolic | | GO:0045454 cell redox homeostasis |
| 4 | regulation | cytoskeleton | GO:0001539 ciliary or flagellar motility |
| 4 | regulation | protease | GO:0004252 serine-type endopeptidase activity |
| 4 | regulation | protease | GO:0008233 peptidase activity |
| 4 | signaling | | GO:0004983 neuropeptide Y receptor activity |
| 4 | signaling | | GO:0005044 scavenger receptor activity |
| 4 | structural | | GO:0051258 protein polymerization |
| 4 | transport | channels&pumps | GO:0005215 transporter activity |
| 4 | transport | channels&pumps | GO:0015171 amino acid transporter activity |
| 5 | metabolic | | GO:0004497 monooxygenase activity |
| 5 | metabolic | | GO:0006200 ATP catabolic process |
| 5 | regulation | protease | GO:0008237 metallopeptidase activity |
| 5 | signaling | | GO:0007264 small GTPase mediated signal transduction |
| 5 | transport | channels&pumps | GO:0008508 bile acid:sodium symporter activity |
| 5 | transport | protein | GO:0006886 intracellular protein transport |
| 6 | cell division | | GO:0000075 cell cycle checkpoint |
| 6 | developmental | | GO:0007368 determination of left/right symmetry |
| 6 | developmental | | GO:0051239 regulation of multicellular organismal process |
| 6 | gene expression | DNA binding | GO:0043433 negative regulation of transcription factor activ |
| 6 | metabolic | | GO:0016758 transferase activity, transferring hexosyl group |
| 6 | metabolic | | GO:0044238 primary metabolic process |
| 6 | regulation | cytoskeleton | GO:0003777 microtubule motor activity |
| 6 | regulation | protease | GO:0004298 threonine endopeptidase activity |
| 6 | regulation | protease | GO:0030168 platelet activation |
| 6 | structural | | GO:0005576 extracellular region |
| 6 | structural | | GO:0005578 proteinaceous extracellular matrix |
| 7 | cell division | | GO:0051301 cell division |
| 7 | gene expression | processing | GO:0000398 nuclear mRNA splicing, via spliceosome |
| 7 | metabolic | | GO:0005743 mitochondrial inner membrane |
| 7 | regulation | binding | GO:0042802 identical protein binding |
| 7 | regulation | | GO:0050794 regulation of cellular process |
| 7 | structural | cytosol | GO:0044424 intracellular part |
| 7 | structural | organelle | GO:0044446 intracellular organelle part |
| 7 | transport | channels&pumps | GO:0005507 copper ion binding |
| 7 | transport | Golgi | GO:0016192 vesicle-mediated transport |
| 8 | developmental | | GO:0000902 cell morphogenesis |
| 8 | gene expression | DNA binding | GO:0043565 sequence-specific DNA binding |
| 8 | metabolic | - | GO:0005759 mitochondrial matrix |
| 8 | metabolic | | GO:0016491 oxidoreductase activity |

| | | | |
|---|---|---|---|
| 8 | metabolic | | GO:0016620 oxidoreductase activity, acting on the aldehyde |
| 8 | regulation | protease | GO:0004222 metalloendopeptidase activity |
| 8 | signaling | | GO:0006182 cGMP biosynthetic process |
| 8 | structural | organelle | GO:0000786 nucleosome |
| 8 | structural | organelle | GO:0043229 intracellular organelle |
| 8 | transport | protein | GO:0006891 intra-Golgi vesicle-mediated transport |
| 9 | gene expression | DNA binding | GO:0003682 chromatin binding |
| 9 | gene expression | translation | GO:0003743 translation initiation factor activity |
| 9 | metabolic | | GO:0006024 glycosaminoglycan biosynthetic process |
| 9 | metabolic | | GO:0031325 positive regulation of cellular metabolic proces |
| 9 | regulation | ubiq | GO:0016567 protein ubiquitination |
| 9 | structural | cytoskeleton | GO:0043197 dendritic spine |
| 9 | structural | organelle | GO:0005840 ribosome |
| 9 | transport | channels&pumps | GO:0005262 calcium channel activity |
| 10 | cell division | | GO:0007049 cell cycle |
| 10 | cell division | | GO:0007131 meiotic recombination |
| 10 | development | | GO:0007368 determination of left/right symmetry |
| 10 | development | | GO:0043025 cell soma |
| *10* | *gene expression* | *polymerase* | *GO:0006261 DNA-dependent DNA replication* |
| 10 | metabolic | | GO:0004091 carboxylesterase activity |
| 10 | metabolic | | GO:0016491 oxidoreductase activity |
| 10 | regulation | cytoskeleton | GO:0003779 actin binding |
| 10 | regulation | protease | GO:0008236 serine-type peptidase activity |
| 10 | regulation | ubiq | GO:0000209 protein polyubiquitination |
| 10 | signaling | | GO:0005102 receptor binding |
| 10 | signaling | | GO:0007264 small GTPase mediated signal transduction |
| 10 | structural | cytoskeleton | GO:0005874 microtubule |
| *10* | *structural* | *organelle* | *GO:0003735 structural constituent of ribosome* |
| 10 | structural | organelle | GO:0005635 nuclear envelope |
| 10 | structural | | GO:0044421 extracellular region part |
| 10 | transport | channels&pumps | GO:0005215 transporter activity |
| 10 | transport | channels&pumps | GO:0015293 symporter activity |
| 11 | gene expression | chromatin | GO:0016514 SWI/SNF complex |
| 11 | signaling | | GO:0000165 MAPKKK cascade |
| 11 | signaling | | GO:0042981 regulation of apoptosis |
| 11 | structural | organelle | GO:0005840 ribosome |

| Table S4. GO terms enrichment for genes expressed only within a single expression phase | | |
|---|---|---|
| ENRICHED PHASE | GO | NAME |
| a | GO:0004674 | protein serine/threonine kinase activity |
| a | GO:0005085 | guanyl-nucleotide exchange factor activity |
| a | GO:0006281 | DNA repair |
| a | GO:0008138 | protein tyrosine/serine/threonine phosphatase activity |
| a | GO:0031965 | nuclear membrane |
| a | GO:0043229 | intracellular organelle |
| a | GO:0051301 | cell division |
| b | GO:0000226 | microtubule cytoskeleton organization and biogenesis |
| b | GO:0001842 | neural fold formation |
| b | GO:0003899 | DNA-directed RNA polymerase activity |
| b | GO:0005452 | inorganic anion exchanger activity |
| b | GO:0008083 | growth factor activity |
| b | GO:0008106 | alcohol dehydrogenase (NADP+) activity |
| b | GO:0008544 | epidermis development |
| b | GO:0050728 | negative regulation of inflammatory response |
| b, c | GO:0000786 | nucleosome |
| b, c | GO:0004222 | metalloendopeptidase activity |
| c | GO:0001731 | formation of translation preinitiation complex |
| c | GO:0003964 | RNA-directed DNA polymerase activity |
| c | GO:0005750 | mitochondrial respiratory chain complex III |
| c | GO:0006935 | chemotaxis |
| c | GO:0008417 | fucosyltransferase activity |
| c | GO:0008528 | peptide receptor activity, G-protein coupled |
| c | GO:0043565 | sequence-specific DNA binding |
| d | GO:0002576 | platelet degranulation |
| d | GO:0004091 | carboxylesterase activity |
| d | GO:0004181 | metallocarboxypeptidase activity |
| d | GO:0005215 | transporter activity |
| d | GO:0005853 | eukaryotic translation elongation factor 1 complex |
| d | GO:0005858 | axonemal dynein complex |
| d | GO:0005975 | carbohydrate metabolic process |
| d | GO:0006629 | lipid metabolic process |
| d | GO:0006811 | ion transport |
| d | GO:0007212 | dopamine receptor signaling pathway |
| d | GO:0008236 | serine-type peptidase activity |

| | | |
|---|---|---|
| d | GO:0015293 | symporter activity |
| d | GO:0016705 | oxidoreductase activity, acting on paired donors, with incorporation or reduction [illegible] |

| **Table S5. k-means clusters for non-ubiquitously expressed transcripts** | | |
|---|---|---|
| **CLUSTER** | **SPU ID** | **NAME** |
| a1 | SPU_000081 | Lv-PionL |
| a1 | SPU_000139 | NA |
| a1 | SPU_000143 | Lv-Hypp_296 |
| a1 | SPU_000162 | NA |
| a1 | SPU_000218 | NA |
| a1 | SPU_000350 | NA |
| a1 | SPU_000369 | NA |
| a1 | SPU_000436 | Lv-Cyp3L2 |
| a1 | SPU_000533 | Lv-Igf_21 |
| a1 | SPU_000576 | Lv-Gpr151L |
| a1 | SPU_000584 | NA |
| a1 | SPU_000599 | Lv-Hypp_310 |
| a1 | SPU_000633 | Lv-Plasmgnr1 |
| a1 | SPU_000983 | NA |
| a1 | SPU_001219 | Lv-Hypp_1314 |
| a1 | SPU_001227 | NA |
| a1 | SPU_001290 | none |
| a1 | SPU_001312 | Lv-ClvhH2b |
| a1 | SPU_001379 | NA |
| a1 | SPU_001610 | Lv-Fn3_15 |
| a1 | SPU_001671 | NA |
| a1 | SPU_001850 | Lv-Vstc/GnrhLr |
| a1 | SPU_001964 | NA |
| a1 | SPU_002045 | Lv-Cdt1L |
| a1 | SPU_002076 | NA |
| a1 | SPU_002142 | Lv-Ece1 |
| a1 | SPU_002154 | Lv-Hnmt |
| a1 | SPU_002171 | Lv-Chst11L |
| a1 | SPU_002221 | Lv-Hypp_1378 |
| a1 | SPU_002288 | NA |
| a1 | SPU_002357 | NA |
| a1 | SPU_002359 | NA |
| a1 | SPU_002400 | NA |
| a1 | SPU_002422 | Lv-AplnrL |

| | | |
|---|---|---|
| a1 | SPU_002426 | Lv-Hln |
| a1 | SPU_002670 | none |
| a1 | SPU_002752 | none |
| a1 | SPU_002845 | Lv-Hypp_1416 |
| a1 | SPU_002922 | none |
| a1 | SPU_002948 | none |
| a1 | SPU_003012 | NA |
| a1 | SPU_003031 | none |
| a1 | SPU_003183 | Lv-Ttll11_1 |
| a1 | SPU_003252 | NA |
| a1 | SPU_003349 | none |
| a1 | SPU_003377 | NA |
| a1 | SPU_003527 | NA |
| a1 | SPU_003544 | NA |
| a1 | SPU_003605 | Lv-Svep1L_4 |
| a1 | SPU_003659 | Lv-Fn3/Egf_12 |
| a1 | SPU_003703 | Lv-Mmp11L |
| a1 | SPU_003707 | Lv-EgfiL |
| a1 | SPU_003790 | NA |
| a1 | SPU_003813 | NA |
| a1 | SPU_004019 | Lv-Seccltcp1 |
| a1 | SPU_004054 | Lv-Hypp_1496 |
| a1 | SPU_004086 | NA |
| a1 | SPU_004094 | NA |
| a1 | SPU_004170 | Lv-SacSl_4 |
| a1 | SPU_004176 | none |
| a1 | SPU_004200 | none |
| a1 | SPU_004393 | NA |
| a1 | SPU_004409 | none |
| a1 | SPU_004469 | Lv-Hypp_1526 |
| a1 | SPU_004508 | none |
| a1 | SPU_004586 | Lv-Ty/Ldla |
| a1 | SPU_004686 | Lv-Echn-21 |
| a1 | SPU_004716 | Lv-Hypp_426 |
| a1 | SPU_004793 | Lv-Fut4L_2 |
| a1 | SPU_004817 | none |
| a1 | SPU_004907 | Lv-Kcnq2 |
| a1 | SPU_004987 | Lv-Hypp_435 |

| | | |
|---|---|---|
| a1 | SPU_005002 | NA |
| a1 | SPU_005078 | none |
| a1 | SPU_005189 | none |
| a1 | SPU_005235 | Lv-Unc44L_42 |
| a1 | SPU_005351 | none |
| a1 | SPU_005404 | Lv-AlkL |
| a1 | SPU_005410 | Lv-Nlr184 |
| a1 | SPU_005510 | Lv-Clca4L |
| a1 | SPU_005533 | none |
| a1 | SPU_005603 | Lv-Hypp_456 |
| a1 | SPU_005705 | Lv-Vps13aL |
| a1 | SPU_005735 | NA |
| a1 | SPU_005762 | Lv-Gem |
| a1 | SPU_005774 | NA |
| a1 | SPU_005829 | none |
| a1 | SPU_005984 | NA |
| a1 | SPU_006083 | NA |
| a1 | SPU_006140 | Lv-GagpL |
| a1 | SPU_006195 | NA |
| a1 | SPU_006346 | Lv-YP30/FascL_5 |
| a1 | SPU_006614 | Lv-RoboL |
| a1 | SPU_006616 | Lv-Negr1L |
| a1 | SPU_006632 | Lv-Neh2 |
| a1 | SPU_006643 | Lv-Trim3L_3 |
| a1 | SPU_006647 | none |
| a1 | SPU_006653 | Lv-Hypp_1672 |
| a1 | SPU_006774 | Lv-Hypp_1679 |
| a1 | SPU_006981 | none |
| a1 | SPU_007026 | NA |
| a1 | SPU_007099 | Lv-Tmem26 |
| a1 | SPU_007183 | none |
| a1 | SPU_007203 | Lv-Igf1p |
| a1 | SPU_007216 | Lv-Cvp18 |
| a1 | SPU_007262 | none |
| a1 | SPU_007263 | none |
| a1 | SPU_007289 | Lv-EfgiaL |
| a1 | SPU_007383 | NA |
| a1 | SPU_007398 | none |

| | | |
|---|---|---|
| a1 | SPU_007465 | Lv-Adrb2L_2 |
| a1 | SPU_007470 | Lv-Clca_2 |
| a1 | SPU_007496 | NA |
| a1 | SPU_007558 | Lv-Cyp4L5 |
| a1 | SPU_007608 | Lv-Hypp_1734 |
| a1 | SPU_007637 | NA |
| a1 | SPU_007660 | NA |
| a1 | SPU_007662 | Lv-Ccdc28b |
| a1 | SPU_007714 | NA |
| a1 | SPU_007789 | Lv-Fn3f_10 |
| a1 | SPU_007824 | NA |
| a1 | SPU_007920 | Lv-Ids_1 |
| a1 | SPU_007948 | Lv-Syt6-1_1 |
| a1 | SPU_008003 | Lv-NeurmdnBr_1 |
| a1 | SPU_008145 | Lv-Echn25 |
| a1 | SPU_008194 | Lv-Boule |
| a1 | SPU_008233 | NA |
| a1 | SPU_008351 | NA |
| a1 | SPU_008404 | NA |
| a1 | SPU_008420 | none |
| a1 | SPU_008615 | none |
| a1 | SPU_008756 | Lv-Adam/Ts7/12L |
| a1 | SPU_008808 | none |
| a1 | SPU_008850 | NA |
| a1 | SPU_008921 | Lv-Drd2aL |
| a1 | SPU_009186 | Lv-Mib1L_1 |
| a1 | SPU_009208 | NA |
| a1 | SPU_009290 | none |
| a1 | SPU_009392 | Lv-Ace |
| a1 | SPU_009474 | Lv-Z364 |
| a1 | SPU_009476 | NA |
| a1 | SPU_009490 | none |
| a1 | SPU_009585 | Lv-Hypp_1860 |
| a1 | SPU_009586 | none |
| a1 | SPU_009607 | Lv-Oit3L |
| a1 | SPU_009608 | Lv-Hypp_52 |
| a1 | SPU_009610 | NA |
| a1 | SPU_009899 | Lv-Popdc3 |

| | | |
|---|---|---|
| a1 | SPU_010066 | Lv-Ph4_3 |
| a1 | SPU_010195 | Lv-Hypp_1903 |
| a1 | SPU_010232 | Lv-Srcr86 |
| a1 | SPU_010271 | none |
| a1 | SPU_010289 | Lv-Plaa |
| a1 | SPU_010429 | Lv-Fn3_38 |
| a1 | SPU_010471 | Lv-Cara8LA |
| a1 | SPU_010473 | Lv-Hypp_3065 |
| a1 | SPU_010483 | NA |
| a1 | SPU_010499 | NA |
| a1 | SPU_010589 | Lv-Hypp_1925 |
| a1 | SPU_010606 | Lv-Endtr |
| a1 | SPU_010661 | Lv-Mib2-2 |
| a1 | SPU_010662 | none |
| a1 | SPU_010942 | NA |
| a1 | SPU_011191 | Lv-CycA_1 |
| a1 | SPU_011228 | NA |
| a1 | SPU_011392 | Lv-Knck16L |
| a1 | SPU_011406 | none |
| a1 | SPU_011452 | NA |
| a1 | SPU_011470 | Lv-Mapkkk9 |
| a1 | SPU_011526 | Lv-Tnpo3_1 |
| a1 | SPU_011558 | Lv-Hypp_1988 |
| a1 | SPU_011732 | Lv-Gage4 |
| a1 | SPU_011761 | Lv-Z295 |
| a1 | SPU_011763 | Lv-Hs3st4L |
| a1 | SPU_011926 | Lv-Cuzd1L |
| a1 | SPU_012104 | none |
| a1 | SPU_012144 | NA |
| a1 | SPU_012162 | none |
| a1 | SPU_012432 | Lv-Lrp4_1 |
| a1 | SPU_012513 | none |
| a1 | SPU_012535 | NA |
| a1 | SPU_012538 | Lv-Hypp_3061 |
| a1 | SPU_012554 | none |
| a1 | SPU_012582 | Lv-Hypp_3062 |
| a1 | SPU_012593 | Lv-Clca6 |
| a1 | SPU_012842 | Lv-Hypp_2051 |

| | | |
|---|---|---|
| a1 | SPU_012876 | none |
| a1 | SPU_012900 | Lv-Nup98_2 |
| a1 | SPU_012945 | Lv-cacc1 |
| a1 | SPU_013000 | Lv-Kiaa1432 |
| a1 | SPU_013030 | none |
| a1 | SPU_013110 | Lv-Hypp_2068 |
| a1 | SPU_013276 | Lv-Zmiz1 |
| a1 | SPU_013535 | NA |
| a1 | SPU_013555 | NA |
| a1 | SPU_013910 | NA |
| a1 | SPU_013996 | NA |
| a1 | SPU_014000 | NA |
| a1 | SPU_014077 | Lv-Hypp_2126 |
| a1 | SPU_014151 | Lv-Clca_7 |
| a1 | SPU_014220 | Lv-Clect/Tm_14 |
| a1 | SPU_014257 | Lv-ProtglyL |
| a1 | SPU_014354 | NA |
| a1 | SPU_014388 | Lv-Kctd21L_4 |
| a1 | SPU_014396 | NA |
| a1 | SPU_014429 | Lv-Snx14_1 |
| a1 | SPU_014438 | Lv-Usp47_1 |
| a1 | SPU_014667 | Lv-Cub/So/Gpcr |
| a1 | SPU_014692 | Lv-AmnL |
| a1 | SPU_014721 | NA |
| a1 | SPU_014753 | NA |
| a1 | SPU_014813 | NA |
| a1 | SPU_014905 | none |
| a1 | SPU_014979 | none |
| a1 | SPU_015095 | Lv-Siat8f |
| a1 | SPU_015240 | Lv-Heatr4_2 |
| a1 | SPU_015246 | Lv-Radr_1 |
| a1 | SPU_015259 | Lv-Orxn/np/qrfpLr2 |
| a1 | SPU_015268 | NA |
| a1 | SPU_015421 | Lv-Trx2 |
| a1 | SPU_015457 | Lv-Hypp_2213 |
| a1 | SPU_015469 | Lv-Hypp_2214 |
| a1 | SPU_015552 | NA |
| a1 | SPU_015699 | Lv-C1qL_7 |

| | | |
|---|---|---|
| a1 | SPU_015756 | Lv-Hypp_2228 |
| a1 | SPU_015761 | Lv-Adrb2_1 |
| a1 | SPU_015792 | Lv-Notchh1_11 |
| a1 | SPU_015802 | Lv-Clca_8 |
| a1 | SPU_015821 | Lv-Hypp_2233 |
| a1 | SPU_015847 | Lv-Hypp_2234 |
| a1 | SPU_015884 | Lv-Hypp_3057 |
| a1 | SPU_015896 | none |
| a1 | SPU_015917 | Lv-Crhr2L |
| a1 | SPU_015933 | Lv-Hypp_2240 |
| a1 | SPU_015945 | Lv-Elft |
| a1 | SPU_015947 | NA |
| a1 | SPU_016079 | Lv-Alk4-5-7_1 |
| a1 | SPU_016206 | Lv-Rxfp2L_4 |
| a1 | SPU_016222 | Lv-Vwdf_2 |
| a1 | SPU_016317 | NA |
| a1 | SPU_016406 | Lv-Dsp3 |
| a1 | SPU_016433 | NA |
| a1 | SPU_016541 | NA |
| a1 | SPU_016647 | none |
| a1 | SPU_016866 | Lv-Yp303 |
| a1 | SPU_016885 | Lv-Sned1_4 |
| a1 | SPU_016920 | Lv-Casr_1 |
| a1 | SPU_016943 | NA |
| a1 | SPU_017002 | NA |
| a1 | SPU_017003 | Lv-Hypp_854 |
| a1 | SPU_017141 | Lv-Prpf8 |
| a1 | SPU_017182 | Lv-Hypp_2308 |
| a1 | SPU_017269 | Lv-Adra2aL |
| a1 | SPU_017380 | Lv-Six4 |
| a1 | SPU_017459 | Lv-Hypp_2327 |
| a1 | SPU_017496 | none |
| a1 | SPU_017514 | Lv-Magi2 |
| a1 | SPU_017536 | Lv-Ttrspn_15 |
| a1 | SPU_017637 | Lv-EglL |
| a1 | SPU_017735 | Lv-Tlr198 |
| a1 | SPU_017790 | Lv-NekA |
| a1 | SPU_017919 | Lv-Adra1b_1 |

| | | |
|---|---|---|
| a1 | SPU_017986 | NA |
| a1 | SPU_018035 | Lv-Syt9 |
| a1 | SPU_018049 | Lv-Mib2-3 |
| a1 | SPU_018160 | Lv-Kiaa1884L |
| a1 | SPU_018182 | NA |
| a1 | SPU_018292 | NA |
| a1 | SPU_018332 | Lv-Xpo4_1 |
| a1 | SPU_018396 | NA |
| a1 | SPU_018449 | NA |
| a1 | SPU_018729 | none |
| a1 | SPU_018779 | NA |
| a1 | SPU_018812 | NA |
| a1 | SPU_018859 | NA |
| a1 | SPU_018909 | Lv-Z39 |
| a1 | SPU_018917 | NA |
| a1 | SPU_018940 | Lv-Clect/Tm_15 |
| a1 | SPU_018998 | Lv-Rho_1 |
| a1 | SPU_019008 | Lv-Dycl |
| a1 | SPU_019129 | Lv-Eyg |
| a1 | SPU_019269 | NA |
| a1 | SPU_019469 | Lv-Cnp_2 |
| a1 | SPU_019612 | Lv-Ataxin7 |
| a1 | SPU_019614 | Lv-Jph3 |
| a1 | SPU_019617 | Lv-RigIL13 |
| a1 | SPU_019681 | Lv-P4htmL |
| a1 | SPU_019687 | Lv-Cpn2L_1 |
| a1 | SPU_019787 | Lv-Cacna1gL |
| a1 | SPU_019798 | Lv-Mib2L_1 |
| a1 | SPU_019811 | Lv-Dgkb2 |
| a1 | SPU_019870 | none |
| a1 | SPU_019879 | NA |
| a1 | SPU_019891 | NA |
| a1 | SPU_019967 | Lv-Echn1 |
| a1 | SPU_020007 | NA |
| a1 | SPU_020049 | Lv-Gpr21L_14 |
| a1 | SPU_020210 | none |
| a1 | SPU_020305 | Lv-Trpa1L_29 |
| a1 | SPU_020357 | NA |

| | | |
|---|---|---|
| a1 | SPU_020377 | Lv-Clect/Pan_6 |
| a1 | SPU_020690 | NA |
| a1 | SPU_020770 | Lv-Gen1L |
| a1 | SPU_020838 | Lv-Cts13 |
| a1 | SPU_020885 | Lv-P160 |
| a1 | SPU_020972 | NA |
| a1 | SPU_021163 | Lv-Syne1L_2 |
| a1 | SPU_021205 | Lv-Z357 |
| a1 | SPU_021254 | Lv-Egf/Lnb/Tm7/Gpcr_2 |
| a1 | SPU_021257 | none |
| a1 | SPU_021325 | NA |
| a1 | SPU_021381 | none |
| a1 | SPU_021392 | NA |
| a1 | SPU_021396 | NA |
| a1 | SPU_021502 | Lv-Tlr212 |
| a1 | SPU_021567 | NA |
| a1 | SPU_021600 | Lv-Notch2_6 |
| a1 | SPU_021613 | Lv-Adra1aL_9 |
| a1 | SPU_021805 | none |
| a1 | SPU_021922 | Lv-Hypp_176 |
| a1 | SPU_021949 | NA |
| a1 | SPU_022075 | Lv-ClvhH2a |
| a1 | SPU_022280 | Lv-Diras |
| a1 | SPU_022336 | NA |
| a1 | SPU_022345 | Lv-PppL_133 |
| a1 | SPU_022376 | Lv-KpLr2 |
| a1 | SPU_022685 | none |
| a1 | SPU_022741 | Lv-Hypp_1056 |
| a1 | SPU_022801 | none |
| a1 | SPU_022897 | Lv-Tpx2L2 |
| a1 | SPU_022994 | NA |
| a1 | SPU_023167 | NA |
| a1 | SPU_023335 | Lv-Piwil2 |
| a1 | SPU_023472 | NA |
| a1 | SPU_023473 | none |
| a1 | SPU_023490 | NA |
| a1 | SPU_023824 | NA |
| a1 | SPU_023825 | Lv-Hypp_2689 |

| | | |
|---|---|---|
| a1 | SPU_024106 | NA |
| a1 | SPU_024156 | Lv-Lrrc15L |
| a1 | SPU_024211 | Lv-Hypp_1109 |
| a1 | SPU_024293 | Lv-Hypp_1112 |
| a1 | SPU_024403 | none |
| a1 | SPU_024482 | NA |
| a1 | SPU_024519 | NA |
| a1 | SPU_024631 | none |
| a1 | SPU_024708 | NA |
| a1 | SPU_024744 | NA |
| a1 | SPU_024839 | Lv-Mgat4c_1 |
| a1 | SPU_024908 | Lv-Chrna7_7 |
| a1 | SPU_024925 | Lv-Sfe1 |
| a1 | SPU_025009 | Lv-D2hgdhL |
| a1 | SPU_025021 | NA |
| a1 | SPU_025034 | Lv-Chrm1L_10 |
| a1 | SPU_025083 | Lv-Hypp_2768 |
| a1 | SPU_025143 | NA |
| a1 | SPU_025229 | none |
| a1 | SPU_025233 | Lv-Hypp_2782 |
| a1 | SPU_025277 | Lv-SelpL_2 |
| a1 | SPU_025382 | none |
| a1 | SPU_025539 | NA |
| a1 | SPU_025842 | Lv-HyalinL_2 |
| a1 | SPU_026122 | Lv-Znfx1-21 |
| a1 | SPU_026288 | Lv-Hcn1 |
| a1 | SPU_026432 | Lv-Drd2L |
| a1 | SPU_026514 | Lv-PlxL_1 |
| a1 | SPU_026534 | none |
| a1 | SPU_026582 | Lv-Rptp_mu/Fn3_1 |
| a1 | SPU_026682 | Lv-Gabbr1_8 |
| a1 | SPU_026800 | Lv-Amy_9 |
| a1 | SPU_026843 | Lv-Echn43 |
| a1 | SPU_026903 | NA |
| a1 | SPU_026991 | NA |
| a1 | SPU_026997 | none |
| a1 | SPU_027043 | NA |
| a1 | SPU_027081 | Lv-Hypp_2885 |

| | | |
|---|---|---|
| a1 | SPU_027204 | NA |
| a1 | SPU_027278 | Lv-Mos |
| a1 | SPU_027315 | none |
| a1 | SPU_027330 | Lv-Bahcc1 |
| a1 | SPU_027379 | Lv-Srcr220 |
| a1 | SPU_027396 | Lv-EchnL_11 |
| a1 | SPU_027416 | Lv-Hypp_2900 |
| a1 | SPU_027504 | NA |
| a1 | SPU_027568 | Lv-Hox3 |
| a1 | SPU_027749 | none |
| a1 | SPU_027759 | Lv-Plxnb2L |
| a1 | SPU_027765 | Lv-Unk_51 |
| a1 | SPU_027868 | Lv-Slc2a5_1 |
| a1 | SPU_027952 | Lv-Fn3/Igf_27 |
| a1 | SPU_028038 | Lv-Rasgrf2 |
| a1 | SPU_028062 | Lv-EhH3_57 |
| a1 | SPU_028075 | none |
| a1 | SPU_028084 | Lv-Kcnq5L |
| a1 | SPU_028088 | NA |
| a1 | SPU_028321 | NA |
| a1 | SPU_028484 | NA |
| a1 | SPU_028502 | NA |
| a1 | SPU_028619 | Lv-Fbn2_1 |
| a1 | SPU_028718 | NA |
| a1 | SPU_028801 | NA |
| a1 | SPU_028839 | NA |
| a1 | SPU_028863 | Lv-Prss7L_1 |
| a1 | SPU_028923 | NA |
| a1 | SPU_028942 | Lv-PppL_168 |
| a1 | SPU_030026 | NA |
| a1 | SPU_030034 | Lv-Acy1l2L_6 |
| a2 | SPU_000033 | Lv-Perld1L |
| a2 | SPU_000100 | Lv-Hypp_1243 |
| a2 | SPU_000269 | Lv-Plxna2 |
| a2 | SPU_000370 | Lv-Hypp_1255 |
| a2 | SPU_000699 | Lv-Hypp_1277 |
| a2 | SPU_000700 | Lv-Hypp_1278 |
| a2 | SPU_000701 | Lv-Hypp_1279 |

| | | |
|---|---|---|
| a2 | SPU_001057 | NA |
| a2 | SPU_001066 | NA |
| a2 | SPU_001177 | Lv-Srcr12 |
| a2 | SPU_001229 | Lv-Srcr13 |
| a2 | SPU_001254 | Lv-Hypp_3111 |
| a2 | SPU_001385 | Lv-Unc44L_12 |
| a2 | SPU_001410 | none |
| a2 | SPU_001675 | Lv-Galr2L_3 |
| a2 | SPU_001676 | Lv-Igfbp |
| a2 | SPU_001735 | none |
| a2 | SPU_002413 | Lv-Hypp_3103 |
| a2 | SPU_002418 | Lv-Mnk2 |
| a2 | SPU_002667 | NA |
| a2 | SPU_002701 | Lv-Hypp_367 |
| a2 | SPU_002814 | Lv-Hypp_1493 |
| a2 | SPU_002868 | Lv-Nlr27 |
| a2 | SPU_002872 | Lv-Clca |
| a2 | SPU_002940 | NA |
| a2 | SPU_002979 | none |
| a2 | SPU_003016 | Lv-Htr1aL_2 |
| a2 | SPU_003578 | Lv-Tlr016 |
| a2 | SPU_003664 | Lv-Clca_17 |
| a2 | SPU_003737 | Lv-Hypp_1477 |
| a2 | SPU_003808 | Lv-Gpr22 |
| a2 | SPU_003854 | NA |
| a2 | SPU_003935 | Lv-Unc44L_34 |
| a2 | SPU_003964 | Lv-Srcr30 |
| a2 | SPU_004440 | Lv-Hypp_417 |
| a2 | SPU_004507 | Lv-Hypp_418 |
| a2 | SPU_004537 | none |
| a2 | SPU_004725 | Lv-Clca1 |
| a2 | SPU_004971 | Lv-Notch2-8 |
| a2 | SPU_004979 | Lv-Hypp_1554 |
| a2 | SPU_005016 | Lv-Hypp_1557 |
| a2 | SPU_005024 | Lv-Unc44L_38 |
| a2 | SPU_005025 | Lv-Hypp_1558 |
| a2 | SPU_005027 | Lv-Unc44L_39 |
| a2 | SPU_005043 | Lv-Hypp_1559 |

| | | |
|---|---|---|
| a2 | SPU_005076 | Lv-Unc44_276 |
| a2 | SPU_005421 | Lv-Trpa1-3 |
| a2 | SPU_005985 | Lv-Adrb1L_2 |
| a2 | SPU_006159 | Lv-Pitx3_1 |
| a2 | SPU_006237 | Lv-Krp95s_1 |
| a2 | SPU_006345 | Lv-YP30/FascL_4 |
| a2 | SPU_006516 | Lv-PppL_248 |
| a2 | SPU_006591 | NA |
| a2 | SPU_006600 | NA |
| a2 | SPU_006648 | Lv-Clect_67 |
| a2 | SPU_006671 | NA |
| a2 | SPU_006866 | Lv-CspeN6a |
| a2 | SPU_007016 | Lv-Hypp_1693 |
| a2 | SPU_007040 | Lv-Clect_16 |
| a2 | SPU_007046 | Lv-Z162 |
| a2 | SPU_007079 | none |
| a2 | SPU_007134 | none |
| a2 | SPU_007256 | Lv-Hypp_1708 |
| a2 | SPU_007355 | NA |
| a2 | SPU_007370 | Lv-Srcr17 |
| a2 | SPU_007466 | none |
| a2 | SPU_007791 | Lv-Fn3/Egff_19 |
| a2 | SPU_007807 | Lv-HtrlaL |
| a2 | SPU_007858 | NA |
| a2 | SPU_007916 | Lv-Cub/Egf/Lnb/7Tm/Gpcr |
| a2 | SPU_008037 | none |
| a2 | SPU_008156 | Lv-Echn26 |
| a2 | SPU_008236 | Lv-Hypp_540 |
| a2 | SPU_008383 | Lv-Egfia_3 |
| a2 | SPU_008387 | Lv-Fn3/Igf_33 |
| a2 | SPU_008435 | Lv-Ig/Fn3f |
| a2 | SPU_008509 | Lv-Galr2L_12 |
| a2 | SPU_008630 | NA |
| a2 | SPU_008750 | none |
| a2 | SPU_008884 | Lv-Surreal/Gpcr51 |
| a2 | SPU_008948 | Lv-Fer2 |
| a2 | SPU_009062 | Lv-Hypp_1822 |
| a2 | SPU_009064 | NA |

| | | |
|---|---|---|
| a2 | SPU_009188 | Lv-Amt2 |
| a2 | SPU_009403 | NA |
| a2 | SPU_009498 | Lv-Chrna10L |
| a2 | SPU_009612 | none |
| a2 | SPU_009763 | Lv-Tie1L_5 |
| a2 | SPU_009777 | NA |
| a2 | SPU_009852 | Lv-Unk_81 |
| a2 | SPU_010020 | Lv-Gtf2ir2b |
| a2 | SPU_010034 | NA |
| a2 | SPU_010177 | Lv-Gpr64L_4 |
| a2 | SPU_010484 | NA |
| a2 | SPU_010809 | none |
| a2 | SPU_010949 | Lv-Hypp_1948 |
| a2 | SPU_011070 | Lv-Ilkap |
| a2 | SPU_011076 | Lv-JamL1 |
| a2 | SPU_011327 | none |
| a2 | SPU_011411 | Lv-Drd2L_15 |
| a2 | SPU_011422 | NA |
| a2 | SPU_011448 | Lv-Tmem151bL |
| a2 | SPU_011553 | Lv-Hypp_1987 |
| a2 | SPU_011632 | Lv-Clect_74 |
| a2 | SPU_011741 | Lv-Stk4L_1 |
| a2 | SPU_011866 | NA |
| a2 | SPU_011887 | none |
| a2 | SPU_011953 | Lv-Rxfp1_1 |
| a2 | SPU_012054 | Lv-Cd97L |
| a2 | SPU_012121 | Lv-Clect/Egf/Tm_3 |
| a2 | SPU_012141 | Lv-Etub3 |
| a2 | SPU_012156 | Lv-EthrL |
| a2 | SPU_012159 | Lv-Srcr105 |
| a2 | SPU_012284 | Lv-Sgsm1_1 |
| a2 | SPU_012314 | Lv-Clect_26 |
| a2 | SPU_012397 | NA |
| a2 | SPU_012483 | none |
| a2 | SPU_012534 | none |
| a2 | SPU_012537 | Lv-Capn5L |
| a2 | SPU_012749 | none |
| a2 | SPU_012952 | Lv-Hypp_2058 |

| | | |
|---|---|---|
| a2 | SPU_012974 | Lv-Galr2L_16 |
| a2 | SPU_012991 | none |
| a2 | SPU_013072 | NA |
| a2 | SPU_013109 | none |
| a2 | SPU_013227 | NA |
| a2 | SPU_013291 | Lv-PolypL_73 |
| a2 | SPU_013453 | Lv-Hypp_716 |
| a2 | SPU_013476 | Lv-CadherinL5 |
| a2 | SPU_013490 | NA |
| a2 | SPU_013865 | Lv-Hypp_2111 |
| a2 | SPU_013979 | Lv-Ltb4dh_1 |
| a2 | SPU_014057 | Lv-Birc2b |
| a2 | SPU_014309 | Lv-Hypp_2139 |
| a2 | SPU_014464 | Lv-Adamts7_1 |
| a2 | SPU_014530 | Lv-Hypp_102 |
| a2 | SPU_014537 | none |
| a2 | SPU_014560 | Lv-Hypp_2158 |
| a2 | SPU_014602 | NA |
| a2 | SPU_014624 | none |
| a2 | SPU_014734 | Lv-Znfx1_9 |
| a2 | SPU_014845 | Lv-TlrL_6 |
| a2 | SPU_015033 | Lv-Nlr195 |
| a2 | SPU_015050 | none |
| a2 | SPU_015070 | Lv-Hypp_2186 |
| a2 | SPU_015087 | Lv-Tsr1 |
| a2 | SPU_015156 | Lv-Cyp3L14 |
| a2 | SPU_015234 | Lv-Galr2 |
| a2 | SPU_015242 | none |
| a2 | SPU_015337 | none |
| a2 | SPU_015383 | none |
| a2 | SPU_015557 | Lv-RtL_39 |
| a2 | SPU_015801 | none |
| a2 | SPU_015862 | none |
| a2 | SPU_015863 | Lv-PlxL |
| a2 | SPU_015910 | Lv-Hypp_2239 |
| a2 | SPU_015957 | Lv-NeurmdnULr2 |
| a2 | SPU_015991 | NA |
| a2 | SPU_015992 | Lv-Hypp_807 |

| | | |
|---|---|---|
| a2 | SPU_016202 | Lv-Hypp_2252 |
| a2 | SPU_016242 | Lv-Plxna4L_4 |
| a2 | SPU_016366 | Lv-Gal3st1L_1 |
| a2 | SPU_016694 | none |
| a2 | SPU_016716 | Lv-Ret |
| a2 | SPU_016721 | Lv-Slc16a12L_11 |
| a2 | SPU_016755 | none |
| a2 | SPU_016902 | NA |
| a2 | SPU_016911 | Lv-Hypp_2287 |
| a2 | SPU_016915 | Lv-Hypp_2288 |
| a2 | SPU_016924 | Lv-Transaa |
| a2 | SPU_016986 | Lv-Unc44_121 |
| a2 | SPU_017022 | Lv-Fn3/Igf_14 |
| a2 | SPU_017065 | Lv-Usp52_2 |
| a2 | SPU_017140 | NA |
| a2 | SPU_017205 | Lv-Hypp_3047 |
| a2 | SPU_017352 | Lv-Hox1 |
| a2 | SPU_017591 | Lv-Hypp_884 |
| a2 | SPU_017611 | none |
| a2 | SPU_017670 | NA |
| a2 | SPU_017693 | Lv-AlkL_2 |
| a2 | SPU_017712 | none |
| a2 | SPU_017776 | Lv-Unken3-5dL_2 |
| a2 | SPU_017868 | none |
| a2 | SPU_017979 | Lv-Fn3f_28 |
| a2 | SPU_018140 | none |
| a2 | SPU_018392 | Lv-Tbx20_1 |
| a2 | SPU_018523 | none |
| a2 | SPU_018663 | none |
| a2 | SPU_018669 | Lv-Hypp_919 |
| a2 | SPU_018690 | none |
| a2 | SPU_019182 | none |
| a2 | SPU_019336 | NA |
| a2 | SPU_019353 | Lv-Hypp2434 |
| a2 | SPU_019432 | none |
| a2 | SPU_019595 | Lv-Hypp_2453 |
| a2 | SPU_019670 | Lv-Fbs_3 |
| a2 | SPU_019918 | none |

| | | |
|---|---|---|
| a2 | SPU_020132 | Lv-Znfx1-12 |
| a2 | SPU_020244 | Lv-Hyalin114 |
| a2 | SPU_020620 | Lv-Rej2 |
| a2 | SPU_020630 | Lv-Mib2L_4 |
| a2 | SPU_020837 | Lv-Cts12 |
| a2 | SPU_020940 | Lv-Rgs22 |
| a2 | SPU_021054 | NA |
| a2 | SPU_021113 | none |
| a2 | SPU_021153 | Lv-Dyrk1A |
| a2 | SPU_021258 | Lv-Hypp_1020 |
| a2 | SPU_021418 | none |
| a2 | SPU_021508 | none |
| a2 | SPU_021748 | Lv-Hypp_2575 |
| a2 | SPU_021799 | none |
| a2 | SPU_021831 | Lv-Slc16a14L |
| a2 | SPU_021866 | Lv-Notch1-18 |
| a2 | SPU_021890 | Lv-Srcr171 |
| a2 | SPU_022041 | Lv-Gpr128L |
| a2 | SPU_022166 | none |
| a2 | SPU_022299 | none |
| a2 | SPU_022348 | none |
| a2 | SPU_022822 | Lv-PolypL_118 |
| a2 | SPU_022851 | Lv-Opsin4 |
| a2 | SPU_022941 | Lv-CspeN2e |
| a2 | SPU_022960 | none |
| a2 | SPU_023088 | none |
| a2 | SPU_023254 | none |
| a2 | SPU_023387 | none |
| a2 | SPU_023458 | none |
| a2 | SPU_023459 | NA |
| a2 | SPU_023477 | Lv-Lrp4L |
| a2 | SPU_024027 | Lv-Dll1 |
| a2 | SPU_024242 | Lv-Unc44_169 |
| a2 | SPU_024269 | Lv-Ubr3 |
| a2 | SPU_024390 | Lv-Srcr203 |
| a2 | SPU_024738 | Lv-Hypp_2749 |
| a2 | SPU_024837 | Lv-B4galt6_7 |
| a2 | SPU_024842 | none |

| | | |
|---|---|---|
| a2 | SPU_024891 | Lv-Lipf |
| a2 | SPU_025142 | Lv-Hypp_1141 |
| a2 | SPU_025209 | Lv-Dhx16_2 |
| a2 | SPU_025372 | Lv-Harbi1L_1 |
| a2 | SPU_025474 | none |
| a2 | SPU_025538 | Lv-Hypp_1153 |
| a2 | SPU_025598 | Lv-TgoL_1 |
| a2 | SPU_025662 | Lv-Slc22a13_4 |
| a2 | SPU_025663 | Lv-Echn42 |
| a2 | SPU_025811 | Lv-Gpr153L |
| a2 | SPU_025827 | Lv-Mphosph9L_1 |
| a2 | SPU_025885 | Lv-RigIL5 |
| a2 | SPU_025958 | Lv-Fmo2_4 |
| a2 | SPU_026060 | Lv-Ldla/Egf/Gpcr |
| a2 | SPU_026290 | Lv-Hypp_2834 |
| a2 | SPU_026439 | Lv-Ugt2b33_5 |
| a2 | SPU_026479 | Lv-Traf1 |
| a2 | SPU_026588 | Lv-Echn13 |
| a2 | SPU_026721 | NA |
| a2 | SPU_026871 | NA |
| a2 | SPU_026876 | Lv-Chrna7_8 |
| a2 | SPU_026998 | Lv-Csmd1L |
| a2 | SPU_027003 | NA |
| a2 | SPU_027025 | Lv-Znfx1-17 |
| a2 | SPU_027035 | Lv-Nlr194 |
| a2 | SPU_027154 | Lv-Slc16a12L_20 |
| a2 | SPU_027350 | none |
| a2 | SPU_027372 | none |
| a2 | SPU_027409 | Lv-Tie1L_8 |
| a2 | SPU_027424 | Lv-B3galnt1 |
| a2 | SPU_027578 | NA |
| a2 | SPU_027686 | Lv-Tie1L |
| a2 | SPU_027741 | Lv-Egf/Sea |
| a2 | SPU_027771 | Lv-Endrvt118 |
| a2 | SPU_027865 | Lv-Hypp_1216 |
| a2 | SPU_028154 | Lv-Hypp_2937 |
| a2 | SPU_028237 | Lv-Inpp1_2 |
| a2 | SPU_028527 | none |

| | | |
|---|---|---|
| a2 | SPU_028911 | Lv-Vps13cL |
| a2 | SPU_030150 | Lv-Atl7L |
| a3 | SPU_000038 | Lv-Slc16a9L |
| a3 | SPU_000481 | Lv-Hypp_1258 |
| a3 | SPU_000499 | Lv-Unc44L_4 |
| a3 | SPU_000713 | NA |
| a3 | SPU_000804 | Lv-Tax1bp1-2 |
| a3 | SPU_000917 | Lv-Lrp4L_4 |
| a3 | SPU_001004 | Lv-Srcr9 |
| a3 | SPU_001319 | none |
| a3 | SPU_001357 | Lv-Hsp20.1 |
| a3 | SPU_001363 | Lv-Gcs1_1 |
| a3 | SPU_001485 | Lv-Drd2L_4 |
| a3 | SPU_001620 | none |
| a3 | SPU_002212 | Lv-Hypp_349 |
| a3 | SPU_003356 | NA |
| a3 | SPU_003462 | Lv-TrafA |
| a3 | SPU_003568 | Lv-Hypp_389 |
| a3 | SPU_003704 | Lv-Lef1 |
| a3 | SPU_003756 | Lv-Hypp_1481 |
| a3 | SPU_003980 | Lv-Hypp_1492 |
| a3 | SPU_004389 | NA |
| a3 | SPU_004483 | NA |
| a3 | SPU_004565 | Lv-Pir-2 |
| a3 | SPU_004603 | NA |
| a3 | SPU_004714 | none |
| a3 | SPU_004754 | Lv-Hypp_1541 |
| a3 | SPU_005203 | Lv-Oxct1 |
| a3 | SPU_005723 | Lv-Mmp18/19L6 |
| a3 | SPU_005831 | NA |
| a3 | SPU_005837 | none |
| a3 | SPU_006178 | NA |
| a3 | SPU_006292 | Lv-Ccdc150L |
| a3 | SPU_006468 | Lv-Znfx1_7 |
| a3 | SPU_006644 | none |
| a3 | SPU_006655 | Lv-SacSl_8 |
| a3 | SPU_006731 | Lv-Srcr53 |
| a3 | SPU_006946 | none |

| | | |
|---|---|---|
| a3 | SPU_007118 | Lv-Clect/Tm_2 |
| a3 | SPU_007199 | Lv-Mtnr1aL |
| a3 | SPU_007299 | Lv-Unc44_210 |
| a3 | SPU_007425 | NA |
| a3 | SPU_007709 | none |
| a3 | SPU_007888 | Lv-Hypp_528 |
| a3 | SPU_007980 | Lv-Mm24 |
| a3 | SPU_008498 | Lv-Nlr134 |
| a3 | SPU_008602 | NA |
| a3 | SPU_008711 | Lv-Htr2bL |
| a3 | SPU_008748 | Lv-Gabbr1_6 |
| a3 | SPU_008801 | Lv-Dynlc2-3c |
| a3 | SPU_008802 | Lv-AnL11 |
| a3 | SPU_009510 | NA |
| a3 | SPU_009570 | Lv-Hypp_1858 |
| a3 | SPU_009823 | Lv-Unc44L_64 |
| a3 | SPU_009827 | NA |
| a3 | SPU_009978 | Lv-Hypp_599 |
| a3 | SPU_010100 | Lv-Clect_21 |
| a3 | SPU_010212 | NA |
| a3 | SPU_010485 | NA |
| a3 | SPU_010514 | NA |
| a3 | SPU_010793 | Lv-Slc14_1 |
| a3 | SPU_010810 | Lv-Hypp_1940 |
| a3 | SPU_011432 | none |
| a3 | SPU_011684 | Lv-Dynlc2-4d |
| a3 | SPU_011749 | Lv-Unc44_298 |
| a3 | SPU_011752 | Lv-Srcr102 |
| a3 | SPU_012500 | Lv-Cyp2L25 |
| a3 | SPU_012882 | Lv-Endrvt81 |
| a3 | SPU_012973 | NA |
| a3 | SPU_013210 | Lv-Unc44L_87 |
| a3 | SPU_013339 | Lv-Hypp_2992 |
| a3 | SPU_013445 | Lv-Plxna4L_1 |
| a3 | SPU_013585 | none |
| a3 | SPU_013636 | none |
| a3 | SPU_013762 | Lv-Clca2 |
| a3 | SPU_013824 | Lv-Tlr074 |

| | | |
|---|---|---|
| a3 | SPU_014253 | Lv-Nkf1 |
| a3 | SPU_014499 | Lv-Gpr52L_7 |
| a3 | SPU_014586 | none |
| a3 | SPU_015569 | Lv-Hypp_787 |
| a3 | SPU_016335 | Lv-Mtif2L1 |
| a3 | SPU_016529 | Lv-Clect_79 |
| a3 | SPU_016833 | Lv-Hypp_2281 |
| a3 | SPU_017397 | Lv-Mtrf1l_1 |
| a3 | SPU_017713 | Lv-Unc44_127 |
| a3 | SPU_017933 | Lv-Srcr140 |
| a3 | SPU_018056 | none |
| a3 | SPU_018448 | NA |
| a3 | SPU_018484 | Lv-Wdr18 |
| a3 | SPU_018578 | Lv-Ankrd44 |
| a3 | SPU_018722 | none |
| a3 | SPU_018752 | NA |
| a3 | SPU_019594 | none |
| a3 | SPU_019861 | none |
| a3 | SPU_020350 | NA |
| a3 | SPU_020394 | Lv-Mib1L |
| a3 | SPU_020531 | none |
| a3 | SPU_020935 | Lv-Gpr1L_9 |
| a3 | SPU_021353 | Lv-Ace_2 |
| a3 | SPU_021509 | Lv-Srcr166 |
| a3 | SPU_021985 | none |
| a3 | SPU_022064 | none |
| a3 | SPU_022404 | NA |
| a3 | SPU_022568 | Lv-Srcr195 |
| a3 | SPU_022572 | NA |
| a3 | SPU_022582 | Lv-Hypp_188 |
| a3 | SPU_022597 | NA |
| a3 | SPU_023033 | NA |
| a3 | SPU_023041 | none |
| a3 | SPU_023415 | none |
| a3 | SPU_023636 | none |
| a3 | SPU_023702 | Lv-Slc5a6_3 |
| a3 | SPU_023724 | NA |
| a3 | SPU_023797 | Lv-Clect_44 |

| | | |
|---|---|---|
| a3 | SPU_023821 | Lv-Hypp_2688 |
| a3 | SPU_023862 | Lv-Abcc5_1 |
| a3 | SPU_024099 | NA |
| a3 | SPU_024147 | Lv-Hypp_1108 |
| a3 | SPU_024682 | Lv-Hypp_1120 |
| a3 | SPU_025114 | Lv-Unc44_345 |
| a3 | SPU_025389 | NA |
| a3 | SPU_025667 | Lv-Unc44_175 |
| a3 | SPU_026144 | none |
| a3 | SPU_026165 | Lv-Reeler/Ccp |
| a3 | SPU_026370 | Lv-Taar7eL_5 |
| a3 | SPU_026615 | NA |
| a3 | SPU_026654 | Lv-Samhd1_7 |
| a3 | SPU_026802 | Lv-Hypp_3121 |
| a3 | SPU_027026 | none |
| a3 | SPU_027243 | Lv-Gpr126L_3 |
| a3 | SPU_028387 | Lv-Nlr69 |
| a3 | SPU_028518 | none |
| a3 | SPU_028603 | Lv-Fucoltf |
| a3 | SPU_028684 | Lv-Apob_2 |
| a3 | SPU_028820 | Lv-Nlr138 |
| a3 | SPU_028927 | Lv-Clca_16 |
| a4 | SPU_000292 | NA |
| a4 | SPU_000470 | NA |
| a4 | SPU_000516 | NA |
| a4 | SPU_000532 | Lv-Igf_13 |
| a4 | SPU_000643 | Lv-Hypp_3012 |
| a4 | SPU_000901 | NA |
| a4 | SPU_001118 | none |
| a4 | SPU_001267 | none |
| a4 | SPU_001301 | none |
| a4 | SPU_001419 | Lv-EchnL_2 |
| a4 | SPU_001465 | Lv-AnL10b_1 |
| a4 | SPU_001575 | Lv-NotchL_7 |
| a4 | SPU_001611 | Lv-Unc44L_16 |
| a4 | SPU_002168 | NA |
| a4 | SPU_002612 | NA |
| a4 | SPU_002720 | NA |

| | | |
|---|---|---|
| a4 | SPU_002731 | NA |
| a4 | SPU_003246 | NA |
| a4 | SPU_003343 | Lv-DhddsL |
| a4 | SPU_003876 | Lv-Rcbtb1_1 |
| a4 | SPU_004099 | Lv-Hypp_1500 |
| a4 | SPU_004707 | Lv-Hypp_1538 |
| a4 | SPU_004849 | Lv-Slc14 |
| a4 | SPU_004997 | NA |
| a4 | SPU_005039 | Lv-Ldla/Egf/Lnb/7TM/Gpcr |
| a4 | SPU_005084 | none |
| a4 | SPU_005091 | none |
| a4 | SPU_005260 | Lv-Abca10a |
| a4 | SPU_005492 | NA |
| a4 | SPU_005506 | NA |
| a4 | SPU_005745 | none |
| a4 | SPU_005995 | Lv-Gabbr1_4 |
| a4 | SPU_006102 | Lv-Slc1a2 |
| a4 | SPU_006146 | Lv-Hypp_1636 |
| a4 | SPU_006551 | none |
| a4 | SPU_007217 | Lv-Slc7a11_7 |
| a4 | SPU_007230 | none |
| a4 | SPU_007323 | Lv-Fn3/Igf_32 |
| a4 | SPU_007371 | NA |
| a4 | SPU_007421 | none |
| a4 | SPU_007570 | Lv-Slc22a5L |
| a4 | SPU_007707 | NA |
| a4 | SPU_007739 | Lv-Npr3 |
| a4 | SPU_008150 | NA |
| a4 | SPU_008228 | Lv-Tlr043 |
| a4 | SPU_008442 | Lv-Hypp_34 |
| a4 | SPU_008494 | Lv-Hypp_1787 |
| a4 | SPU_008583 | NA |
| a4 | SPU_009144 | NA |
| a4 | SPU_009222 | Lv-Clect/Egf_1 |
| a4 | SPU_009489 | Lv-Hypp_578 |
| a4 | SPU_009699 | Lv-Hypp_585 |
| a4 | SPU_009805 | Lv-Hypp_56 |
| a4 | SPU_009875 | Lv-AgrnL_1 |

| | | |
|---|---|---|
| a4 | SPU_010192 | NA |
| a4 | SPU_010242 | none |
| a4 | SPU_010284 | Lv-Cdk4 |
| a4 | SPU_010557 | Lv-Mocs1_1 |
| a4 | SPU_010584 | NA |
| a4 | SPU_010987 | Lv-Ercc6L |
| a4 | SPU_011012 | none |
| a4 | SPU_011201 | Lv-Sec7-containing |
| a4 | SPU_011678 | Lv-Hypp_654 |
| a4 | SPU_011995 | Lv-Egfib_5 |
| a4 | SPU_011996 | none |
| a4 | SPU_012688 | none |
| a4 | SPU_013790 | NA |
| a4 | SPU_014333 | NA |
| a4 | SPU_014722 | Lv-Pmar1 |
| a4 | SPU_014834 | Lv-Hypp_2171 |
| a4 | SPU_014856 | Lv-Gpi_2 |
| a4 | SPU_015157 | none |
| a4 | SPU_015291 | NA |
| a4 | SPU_015314 | Lv-Slc5a12_4 |
| a4 | SPU_015387 | Lv-Srcr125 |
| a4 | SPU_015476 | Lv-Atp13a3_3 |
| a4 | SPU_015585 | NA |
| a4 | SPU_015641 | none |
| a4 | SPU_015780 | Lv-Adra1bL_9 |
| a4 | SPU_015828 | Lv-Pmar1c |
| a4 | SPU_015829 | Lv-Pmar1d |
| a4 | SPU_016218 | none |
| a4 | SPU_016551 | NA |
| a4 | SPU_016624 | Lv-Unc44_116 |
| a4 | SPU_016947 | Lv-Slc17a5_6 |
| a4 | SPU_017001 | Lv-Hypp_2297 |
| a4 | SPU_017287 | Lv-Hand |
| a4 | SPU_017499 | NA |
| a4 | SPU_017520 | NA |
| a4 | SPU_017678 | Lv-Fbn3 |
| a4 | SPU_018047 | none |
| a4 | SPU_018754 | Lv-Unc44_136 |

| | | |
|---|---|---|
| a4 | SPU_018942 | Lv-Unc44_240 |
| a4 | SPU_019255 | none |
| a4 | SPU_019362 | Lv-Chrm2L_6 |
| a4 | SPU_019698 | Lv-Hypp_2462 |
| a4 | SPU_019728 | none |
| a4 | SPU_019789 | Lv-Hypp_946 |
| a4 | SPU_020279 | none |
| a4 | SPU_020301 | Lv-Acsf2_3 |
| a4 | SPU_020580 | Lv-Samhd1_4 |
| a4 | SPU_020648 | Lv-Birc2c |
| a4 | SPU_020822 | Lv-Srcr161 |
| a4 | SPU_020982 | Lv-Tll2L_2 |
| a4 | SPU_021220 | none |
| a4 | SPU_021346 | Lv-Egfia_9 |
| a4 | SPU_021414 | NA |
| a4 | SPU_021770 | none |
| a4 | SPU_021793 | Lv-Slc8a2 |
| a4 | SPU_022226 | none |
| a4 | SPU_022421 | Lv-Znfx1 |
| a4 | SPU_022491 | Lv-Hypp_2612 |
| a4 | SPU_022584 | Lv-Acot9_1 |
| a4 | SPU_022661 | Lv-Echn38 |
| a4 | SPU_022911 | Lv-Tlr117 |
| a4 | SPU_023055 | NA |
| a4 | SPU_023923 | Lv-Fn3/Egf_4 |
| a4 | SPU_024018 | NA |
| a4 | SPU_024032 | Lv-Apn5 |
| a4 | SPU_025004 | none |
| a4 | SPU_025061 | Lv-Unk_23 |
| a4 | SPU_025509 | none |
| a4 | SPU_025546 | NA |
| a4 | SPU_025597 | Lv-TgoL |
| a4 | SPU_025604 | none |
| a4 | SPU_026259 | Lv-PtprB |
| a4 | SPU_026779 | Lv-Rock1 |
| a4 | SPU_027031 | Lv-Galr2_1 |
| a4 | SPU_027456 | Lv-Unk_27 |
| a4 | SPU_027612 | Lv-GagpolL_13 |

| | | |
|---|---|---|
| a4 | SPU_027746 | Lv-Unc44_261 |
| a4 | SPU_028127 | Lv-Hypp_2934 |
| a4 | SPU_028645 | Lv-HtrL_1 |
| a4 | SPU_028697 | Lv-Slc7a1 |
| a4 | SPU_030101 | Lv-TlrP46 |
| a5 | SPU_000413 | Lv-Chrm5L |
| a5 | SPU_000507 | none |
| a5 | SPU_000567 | Lv-He2 |
| a5 | SPU_000596 | none |
| a5 | SPU_000976 | Lv-Psmd7Lp26SnATPase7 |
| a5 | SPU_001013 | none |
| a5 | SPU_001274 | Lv-Clect_4 |
| a5 | SPU_001285 | Lv-Slc7a6_4 |
| a5 | SPU_001400 | Lv-Cpa3L_1 |
| a5 | SPU_001407 | Lv-PppL_739 |
| a5 | SPU_001474 | none |
| a5 | SPU_001607 | none |
| a5 | SPU_001737 | NA |
| a5 | SPU_001941 | Lv-Hypp_1356 |
| a5 | SPU_001957 | Lv-Fn3f_1 |
| a5 | SPU_002019 | none |
| a5 | SPU_002152 | none |
| a5 | SPU_002496 | NA |
| a5 | SPU_002576 | none |
| a5 | SPU_002708 | Lv-Z382 |
| a5 | SPU_002746 | Lv-Fbln2L_2 |
| a5 | SPU_002771 | none |
| a5 | SPU_002823 | none |
| a5 | SPU_002831 | Lv-Cyp2L7 |
| a5 | SPU_002981 | Lv-Sned1 |
| a5 | SPU_003000 | none |
| a5 | SPU_003096 | NA |
| a5 | SPU_003186 | Lv-Nlr72 |
| a5 | SPU_003337 | Lv-Hfm1-2 |
| a5 | SPU_003666 | none |
| a5 | SPU_003714 | Lv-Crat_1 |
| a5 | SPU_003950 | Lv-Kidins220 |
| a5 | SPU_004058 | Lv-Adrb2L |

| | | |
|---|---|---|
| a5 | SPU_004123 | Lv-Tmprss3 |
| a5 | SPU_004232 | none |
| a5 | SPU_004305 | none |
| a5 | SPU_004370 | Lv-Hypp_1518 |
| a5 | SPU_004448 | none |
| a5 | SPU_004581 | Lv-Adora2aL_3 |
| a5 | SPU_004935 | Lv-Ndufa8_1 |
| a5 | SPU_005146 | none |
| a5 | SPU_005620 | NA |
| a5 | SPU_005754 | Lv-ctbs |
| a5 | SPU_005811 | Lv-Uqcrfs1 |
| a5 | SPU_005882 | Lv-Tssk1 |
| a5 | SPU_005972 | Lv-Unc44_208 |
| a5 | SPU_005992 | Lv-Adndatrnas |
| a5 | SPU_006425 | Lv-Hypp_481 |
| a5 | SPU_006533 | Lv-SphH1 |
| a5 | SPU_006708 | none |
| a5 | SPU_006714 | none |
| a5 | SPU_006729 | Lv-Lvn1.4 |
| a5 | SPU_006822 | Lv-Surreal/Gpcr26 |
| a5 | SPU_006833 | Lv-Surreal/Gpcr81 |
| a5 | SPU_007330 | NA |
| a5 | SPU_007432 | Lv-Adrb2L_1 |
| a5 | SPU_007690 | none |
| a5 | SPU_008016 | none |
| a5 | SPU_008047 | Lv-Asp_10 |
| a5 | SPU_008098 | Lv-Hypp_533 |
| a5 | SPU_008226 | Lv-Galr2L_11 |
| a5 | SPU_008276 | none |
| a5 | SPU_008310 | none |
| a5 | SPU_008397 | Lv-Slc29a4 |
| a5 | SPU_008752 | Lv-Nfe2_1 |
| a5 | SPU_008800 | NA |
| a5 | SPU_009106 | none |
| a5 | SPU_009386 | NA |
| a5 | SPU_009395 | Lv-MucL_2 |
| a5 | SPU_009546 | Lv-Copz1_1 |
| a5 | SPU_009826 | Lv-RasD1/2 |

| | | |
|---|---|---|
| a5 | SPU_010045 | Lv-Rbp4L |
| a5 | SPU_010416 | Lv-Speedy/Ringo |
| a5 | SPU_010531 | Lv-Atub12 |
| a5 | SPU_010600 | Lv-Fbxo47 |
| a5 | SPU_010817 | Lv-Ankrp_1 |
| a5 | SPU_010888 | none |
| a5 | SPU_011003 | none |
| a5 | SPU_011036 | Lv-Nr1 |
| a5 | SPU_011230 | Lv-Ssr4_1 |
| a5 | SPU_011329 | Lv-Sdha |
| a5 | SPU_011447 | none |
| a5 | SPU_011552 | NA |
| a5 | SPU_011731 | Lv-Asp_13 |
| a5 | SPU_011826 | Lv-EchnL_6 |
| a5 | SPU_011899 | Lv-Hypp_2006 |
| a5 | SPU_011947 | Lv-Hrh3_2 |
| a5 | SPU_012035 | NA |
| a5 | SPU_012334 | Lv-Hypp_682 |
| a5 | SPU_012509 | none |
| a5 | SPU_012566 | none |
| a5 | SPU_012619 | Lv-Sastk |
| a5 | SPU_012624 | none |
| a5 | SPU_012673 | none |
| a5 | SPU_013034 | none |
| a5 | SPU_013220 | Lv-Fn3/Egff_10 |
| a5 | SPU_013221 | Lv-Unc44L_89 |
| a5 | SPU_013260 | none |
| a5 | SPU_013495 | none |
| a5 | SPU_013613 | Lv-B3galt1_10 |
| a5 | SPU_013657 | Lv-Adrb2L_4 |
| a5 | SPU_013820 | Lv-Eno3 |
| a5 | SPU_013839 | Lv-Pdap1 |
| a5 | SPU_014090 | Lv-Hypp_3019 |
| a5 | SPU_014184 | Lv-Clect_31 |
| a5 | SPU_014335 | Lv-Hypp_98 |
| a5 | SPU_014500 | Lv-Hypp_2155 |
| a5 | SPU_014508 | Lv-Hypp_756 |
| a5 | SPU_014593 | none |

| | | |
|---|---|---|
| a5 | SPU_014730 | Lv-PlasmgnL2 |
| a5 | SPU_014745 | Lv-Adrb2L_7 |
| a5 | SPU_014857 | none |
| a5 | SPU_014865 | NA |
| a5 | SPU_015007 | Lv-Fn3/Egf_10 |
| a5 | SPU_015279 | none |
| a5 | SPU_015293 | Lv-FaMeT |
| a5 | SPU_015492 | Lv-Grik3 |
| a5 | SPU_015512 | Lv-Gpr21L_11 |
| a5 | SPU_015513 | none |
| a5 | SPU_015519 | none |
| a5 | SPU_015592 | Lv-Cct8_1 |
| a5 | SPU_015631 | Lv-Hypp_2222 |
| a5 | SPU_016094 | none |
| a5 | SPU_016191 | none |
| a5 | SPU_016295 | none |
| a5 | SPU_016360 | none |
| a5 | SPU_016565 | Lv-SphH2b2 |
| a5 | SPU_016704 | Lv-Hypp_2269 |
| a5 | SPU_016711 | Lv-Sf3a3_1 |
| a5 | SPU_016785 | Lv-Trdn |
| a5 | SPU_016875 | Lv-Trappc10L |
| a5 | SPU_016880 | Lv-Srcr136 |
| a5 | SPU_016942 | Lv-Hrh3L_12 |
| a5 | SPU_017170 | Lv-Treh_2 |
| a5 | SPU_017195 | Lv-Hypp_2309 |
| a5 | SPU_017214 | Lv-Hypp_866 |
| a5 | SPU_017371 | Lv-Hypp_2322 |
| a5 | SPU_017475 | NA |
| a5 | SPU_017588 | Lv-Hypp_136 |
| a5 | SPU_017730 | none |
| a5 | SPU_017759 | Lv-Ppp1cb_1 |
| a5 | SPU_017796 | Lv-Ank1 |
| a5 | SPU_017913 | Lv-Hypp_901 |
| a5 | SPU_018305 | none |
| a5 | SPU_018547 | Lv-Tie1L_1 |
| a5 | SPU_018603 | Lv-Gpr21L_13 |
| a5 | SPU_018633 | none |

| | | |
|---|---|---|
| a5 | SPU_018708 | NA |
| a5 | SPU_018930 | none |
| a5 | SPU_019077 | none |
| a5 | SPU_019260 | Lv-Imp3_1 |
| a5 | SPU_019575 | Lv-Clect_39 |
| a5 | SPU_019642 | Lv-Unc44_242 |
| a5 | SPU_020170 | NA |
| a5 | SPU_020375 | none |
| a5 | SPU_020590 | Lv-Fn3f_6 |
| a5 | SPU_020621 | none |
| a5 | SPU_020842 | none |
| a5 | SPU_021405 | Lv-Hypp_3032 |
| a5 | SPU_021571 | Lv-Ndufv2_1 |
| a5 | SPU_021594 | none |
| a5 | SPU_021906 | none |
| a5 | SPU_021925 | Lv-Wdr23 |
| a5 | SPU_023185 | Lv-Lnb/7Tm/GpcrC |
| a5 | SPU_023345 | Lv-Wdfy3 |
| a5 | SPU_023360 | none |
| a5 | SPU_023606 | Lv-Hypp_1092 |
| a5 | SPU_023746 | Lv-Znfx1-14 |
| a5 | SPU_024263 | Lv-SerpL9 |
| a5 | SPU_024614 | Lv-F2 |
| a5 | SPU_024712 | none |
| a5 | SPU_024769 | Lv-Noggin |
| a5 | SPU_024861 | none |
| a5 | SPU_024872 | Lv-NifuL |
| a5 | SPU_024888 | Lv-Galr2L_30 |
| a5 | SPU_024923 | NA |
| a5 | SPU_025013 | none |
| a5 | SPU_025134 | Lv-Hypp_2774 |
| a5 | SPU_025248 | Lv-Clect_49 |
| a5 | SPU_025403 | Lv-Slc5a3-3 |
| a5 | SPU_025468 | none |
| a5 | SPU_025494 | Lv-Notch2_7 |
| a5 | SPU_025576 | none |
| a5 | SPU_025861 | none |
| a5 | SPU_025957 | NA |

| | | |
|---|---|---|
| a5 | SPU_026059 | Lv-Surreal/Gpcr53 |
| a5 | SPU_026152 | Lv-Znfx1-16 |
| a5 | SPU_026718 | Lv-Ogdh_1 |
| a5 | SPU_026745 | Lv-P120L |
| a5 | SPU_027034 | none |
| a5 | SPU_027336 | Lv-Ras/Dva |
| a5 | SPU_027465 | Lv-Unc44_182 |
| a5 | SPU_027473 | Lv-Hypp_1204 |
| a5 | SPU_027775 | Lv-Rdh3 |
| a5 | SPU_027863 | Lv-C10orf112L_4 |
| a5 | SPU_027866 | Lv-Gpr54L_6 |
| a5 | SPU_028077 | Lv-Cndpdp2L |
| a5 | SPU_028086 | none |
| a5 | SPU_028288 | Lv-F2L_2 |
| a5 | SPU_028344 | Lv-Akrf_4 |
| a5 | SPU_028499 | Lv-B3gnt7_1 |
| a5 | SPU_028581 | Lv-Lrp2L |
| a5 | SPU_028590 | none |
| a5 | SPU_028922 | Lv-Cyp2L66 |
| b1 | SPU_000004 | Lv-Hypp_1240 |
| b1 | SPU_000027 | Lv-Rtl |
| b1 | SPU_000428 | Lv-Tlr004 |
| b1 | SPU_000459 | Lv-Smyd3L |
| b1 | SPU_000566 | Lv-He1 |
| b1 | SPU_000629 | Lv-C2orf42L |
| b1 | SPU_001096 | NA |
| b1 | SPU_001133 | Lv-Unc44_2641 |
| b1 | SPU_001201 | Lv-Ph4 |
| b1 | SPU_001220 | Lv-Hypp_1315 |
| b1 | SPU_001238 | none |
| b1 | SPU_001766 | NA |
| b1 | SPU_001784 | Lv-Hypp_339 |
| b1 | SPU_001904 | none |
| b1 | SPU_001993 | Lv-Tlr162 |
| b1 | SPU_002240 | Lv-Gpr21L_4 |
| b1 | SPU_002349 | NA |
| b1 | SPU_002597 | NA |
| b1 | SPU_002631 | Lv-Hox11/13b |

| | | |
|---|---|---|
| b1 | SPU_002705 | Lv-Znfx1_4 |
| b1 | SPU_002844 | NA |
| b1 | SPU_003036 | Lv-Hypp_1426 |
| b1 | SPU_003268 | none |
| b1 | SPU_003424 | Lv-Tssk4 |
| b1 | SPU_003477 | Lv-Sf3B2_1 |
| b1 | SPU_003563 | NA |
| b1 | SPU_003594 | none |
| b1 | SPU_003604 | none |
| b1 | SPU_004113 | Lv-An |
| b1 | SPU_004139 | Lv-Tlr019 |
| b1 | SPU_004148 | Lv-Krl |
| b1 | SPU_004460 | NA |
| b1 | SPU_004497 | Lv-Hypp_1531 |
| b1 | SPU_004502 | Lv-Gpr103L |
| b1 | SPU_004610 | NA |
| b1 | SPU_004624 | Lv-Hypp_421 |
| b1 | SPU_004650 | none |
| b1 | SPU_004675 | Lv-Rfesd |
| b1 | SPU_004711 | none |
| b1 | SPU_004792 | Lv-Tlr167 |
| b1 | SPU_004871 | Lv-Echn-22 |
| b1 | SPU_004953 | Lv-Hypp_3087 |
| b1 | SPU_005113 | NA |
| b1 | SPU_005342 | NA |
| b1 | SPU_005437 | none |
| b1 | SPU_005477 | Lv-ItgaL |
| b1 | SPU_005674 | NA |
| b1 | SPU_005728 | NA |
| b1 | SPU_006207 | Lv-Hypp_3077 |
| b1 | SPU_006267 | NA |
| b1 | SPU_006486 | Lv-Slc10a2_1 |
| b1 | SPU_006488 | none |
| b1 | SPU_006489 | none |
| b1 | SPU_006508 | Lv-Clect_15 |
| b1 | SPU_006784 | Lv-Pi3k |
| b1 | SPU_007030 | none |
| b1 | SPU_007042 | Lv-Hypp_3014 |

| | | |
|---|---|---|
| b1 | SPU_007270 | Lv-KctdL_1 |
| b1 | SPU_007316 | Lv-Hypp_1713 |
| b1 | SPU_007453 | Lv-Hypp_1724 |
| b1 | SPU_007615 | Lv-Hypp_1735 |
| b1 | SPU_007774 | NA |
| b1 | SPU_007805 | none |
| b1 | SPU_008084 | NA |
| b1 | SPU_008220 | Lv-Endrvt65 |
| b1 | SPU_008275 | none |
| b1 | SPU_008336 | NA |
| b1 | SPU_008337 | NA |
| b1 | SPU_008375 | Lv-Endrvt67 |
| b1 | SPU_008514 | Lv-Srcr69 |
| b1 | SPU_008676 | none |
| b1 | SPU_008841 | NA |
| b1 | SPU_008937 | Lv-Ugt2b1_3 |
| b1 | SPU_008961 | Lv-Hypp_1817 |
| b1 | SPU_009562 | Lv-Srcr77 |
| b1 | SPU_009564 | Lv-Glt1 |
| b1 | SPU_009604 | NA |
| b1 | SPU_009947 | NA |
| b1 | SPU_010129 | NA |
| b1 | SPU_010318 | Lv-Surreal/Gpcr33 |
| b1 | SPU_010400 | Lv-Hypp_1915 |
| b1 | SPU_010465 | Lv-Unc44_290 |
| b1 | SPU_010615 | Lv-Clect/Pan_13 |
| b1 | SPU_010768 | Lv-RtL_24 |
| b1 | SPU_011000 | none |
| b1 | SPU_011518 | Lv-Chst11L_2 |
| b1 | SPU_011627 | Lv-EhH2b_38 |
| b1 | SPU_011751 | none |
| b1 | SPU_011780 | Lv-Hypp_661 |
| b1 | SPU_011827 | NA |
| b1 | SPU_011831 | Lv-Gpr151L_3 |
| b1 | SPU_011834 | none |
| b1 | SPU_012060 | Lv-Unc44L_81 |
| b1 | SPU_012168 | NA |
| b1 | SPU_012267 | Lv-Unc44L_83 |

| | | |
|---|---|---|
| b1 | SPU_012298 | none |
| b1 | SPU_012308 | NA |
| b1 | SPU_012327 | NA |
| b1 | SPU_012388 | Lv-Igvc1 |
| b1 | SPU_012400 | Lv-NmbrL_1 |
| b1 | SPU_012410 | Lv-Srcr107 |
| b1 | SPU_012420 | Lv-PppL_195 |
| b1 | SPU_012421 | Lv-PolppL_93 |
| b1 | SPU_012448 | Lv-Ovo |
| b1 | SPU_012588 | NA |
| b1 | SPU_012595 | Lv-PolppL_95 |
| b1 | SPU_012625 | Lv-SlspL |
| b1 | SPU_012631 | Lv-Notchh2_2 |
| b1 | SPU_012867 | NA |
| b1 | SPU_013162 | Lv-Tlr186 |
| b1 | SPU_013213 | Lv-Unc44L_88 |
| b1 | SPU_013390 | Lv-Endrvt84 |
| b1 | SPU_013787 | Lv-RtL_35 |
| b1 | SPU_013899 | Lv-Htr1L_3 |
| b1 | SPU_013958 | Lv-Srcr111 |
| b1 | SPU_014152 | NA |
| b1 | SPU_014266 | NA |
| b1 | SPU_014287 | Lv-St8sia5 |
| b1 | SPU_014555 | Lv-Adra1aL_5 |
| b1 | SPU_014737 | Lv-Fam55cL_1 |
| b1 | SPU_014927 | NA |
| b1 | SPU_014932 | NA |
| b1 | SPU_014941 | Lv-Hypp_768 |
| b1 | SPU_014992 | Lv-Srcr120 |
| b1 | SPU_015010 | NA |
| b1 | SPU_015072 | Lv-Galr2L_18 |
| b1 | SPU_015140 | Lv-Tcknr2 |
| b1 | SPU_015185 | Lv-Tlr191 |
| b1 | SPU_015203 | NA |
| b1 | SPU_015309 | none |
| b1 | SPU_015333 | NA |
| b1 | SPU_015444 | Lv-Hypp_785 |
| b1 | SPU_015570 | Lv-Hypp_788 |

| | | |
|---|---|---|
| b1 | SPU_015601 | Lv-Ank10 |
| b1 | SPU_015632 | Lv-Chrm3L_9 |
| b1 | SPU_015963 | Lv-Inpp5a_1 |
| b1 | SPU_015983 | Lv-MyoD3 |
| b1 | SPU_016162 | none |
| b1 | SPU_016188 | Lv-ReelerL_4 |
| b1 | SPU_016199 | Lv-Hypp_817 |
| b1 | SPU_016207 | none |
| b1 | SPU_016257 | Lv-Nlr142 |
| b1 | SPU_016332 | NA |
| b1 | SPU_016501 | Lv-Tlr086 |
| b1 | SPU_016607 | none |
| b1 | SPU_016703 | Lv-Dhyg_3 |
| b1 | SPU_017171 | Lv-Fbg/Egf/Clect |
| b1 | SPU_017196 | Lv-Nlr132 |
| b1 | SPU_017336 | Lv-Gluld1_15 |
| b1 | SPU_017464 | Lv-Hypp_2328 |
| b1 | SPU_017505 | Lv-Nlr131 |
| b1 | SPU_017506 | Lv-Nlr131_1 |
| b1 | SPU_017523 | Lv-CspeN3a |
| b1 | SPU_017639 | Lv-Fmo |
| b1 | SPU_017718 | NA |
| b1 | SPU_017866 | Lv-Gpr1L_6 |
| b1 | SPU_017907 | NA |
| b1 | SPU_017931 | Lv-PppL_118 |
| b1 | SPU_017984 | Lv-Hypp_904 |
| b1 | SPU_018001 | Lv-CckarL_1 |
| b1 | SPU_018014 | none |
| b1 | SPU_018015 | none |
| b1 | SPU_018104 | Lv-Unc44_319 |
| b1 | SPU_018105 | Lv-Unc44_320 |
| b1 | SPU_018138 | Lv-Hypp_140 |
| b1 | SPU_018227 | Lv-Oprk1L_7 |
| b1 | SPU_018230 | none |
| b1 | SPU_018266 | Lv-Hypp_2378 |
| b1 | SPU_018279 | none |
| b1 | SPU_018360 | NA |
| b1 | SPU_018431 | Lv-Slc2a5 |

| | | |
|---|---|---|
| b1 | SPU_018472 | Lv-Mmp1/3L1 |
| b1 | SPU_018726 | NA |
| b1 | SPU_018869 | none |
| b1 | SPU_018954 | Lv-Awh |
| b1 | SPU_018985 | Lv-Srcr147 |
| b1 | SPU_019204 | NA |
| b1 | SPU_019322 | Lv-ShprhL |
| b1 | SPU_019822 | none |
| b1 | SPU_019869 | none |
| b1 | SPU_019878 | NA |
| b1 | SPU_019965 | Lv-Slc16a4L_4 |
| b1 | SPU_020103 | NA |
| b1 | SPU_020110 | Lv-Z474_1 |
| b1 | SPU_020253 | none |
| b1 | SPU_020271 | NA |
| b1 | SPU_020400 | Lv-Hypp_973 |
| b1 | SPU_020594 | Lv-Chrm3L_15 |
| b1 | SPU_020666 | Lv-TlrP22 |
| b1 | SPU_020941 | Lv-Polyp_4 |
| b1 | SPU_021006 | none |
| b1 | SPU_021015 | Lv-Mtnr1aL_2 |
| b1 | SPU_021052 | NA |
| b1 | SPU_021240 | none |
| b1 | SPU_021389 | Lv-Unc44_249 |
| b1 | SPU_021436 | NA |
| b1 | SPU_021553 | NA |
| b1 | SPU_021772 | Lv-Ddc_1 |
| b1 | SPU_021776 | Lv-Adra2bL |
| b1 | SPU_022000 | Lv-Srcr174 |
| b1 | SPU_022085 | Lv-Srcr175 |
| b1 | SPU_022124 | Lv-B3galt2_1 |
| b1 | SPU_022287 | Lv-Srcr187 |
| b1 | SPU_022390 | Lv-Gpr21L_15 |
| b1 | SPU_022402 | Lv-Hrh3L_17 |
| b1 | SPU_022738 | Lv-Hypp_2627 |
| b1 | SPU_022909 | Lv-Tlr116 |
| b1 | SPU_023193 | Lv-TlrP29 |
| b1 | SPU_023230 | Lv-Hypp_1078 |

| | | |
|---|---|---|
| b1 | SPU_023238 | Lv-Fut4_4 |
| b1 | SPU_023579 | NA |
| b1 | SPU_023598 | Lv-Fam166b |
| b1 | SPU_023786 | NA |
| b1 | SPU_023858 | none |
| b1 | SPU_023912 | none |
| b1 | SPU_023979 | Lv-Aqp_3 |
| b1 | SPU_024078 | NA |
| b1 | SPU_024148 | none |
| b1 | SPU_024653 | Lv-C1qL_4 |
| b1 | SPU_024714 | Lv-B3galt5_3 |
| b1 | SPU_024871 | none |
| b1 | SPU_024924 | Lv-Orxn/np/qrfpLr1 |
| b1 | SPU_024992 | Lv-Chrm3_1 |
| b1 | SPU_025006 | none |
| b1 | SPU_025168 | Lv-Rergl4 |
| b1 | SPU_025322 | NA |
| b1 | SPU_025333 | NA |
| b1 | SPU_025461 | NA |
| b1 | SPU_026047 | none |
| b1 | SPU_026208 | NA |
| b1 | SPU_026274 | NA |
| b1 | SPU_026299 | Lv-Lrr2 |
| b1 | SPU_026323 | Lv-Slc10a2L_2 |
| b1 | SPU_026713 | Lv-Hypp_2863 |
| b1 | SPU_026818 | Lv-C12orf10L |
| b1 | SPU_026929 | NA |
| b1 | SPU_026941 | Lv-Tmprss8L |
| b1 | SPU_027178 | NA |
| b1 | SPU_027351 | none |
| b1 | SPU_027546 | NA |
| b1 | SPU_027864 | none |
| b1 | SPU_027941 | none |
| b1 | SPU_027951 | Lv-Fn3/Igf_6 |
| b1 | SPU_027955 | Lv-Hypp_1219 |
| b1 | SPU_028031 | Lv-Furin_2 |
| b1 | SPU_028206 | Lv-Gpr101L_7 |
| b1 | SPU_028285 | Lv-TpaseL_21 |

| | | |
|---|---|---|
| b1 | SPU_028331 | NA |
| b1 | SPU_028358 | Lv-Akrf_2 |
| b1 | SPU_028382 | NA |
| b1 | SPU_028500 | NA |
| b1 | SPU_028513 | none |
| b1 | SPU_028583 | Lv-Zic |
| b1 | SPU_028709 | Lv-Pst |
| b1 | SPU_028742 | Lv-Astacin2 |
| b1 | SPU_028815 | Lv-EhH2b_94 |
| b1 | SPU_028928 | Lv-Clca2L_1 |
| b1 | SPU_030011 | Lv-Mmp2L_1 |
| b1 | SPU_030116 | Lv-TlrP60 |
| b1 | SPU_030130 | Lv-TlrP74 |
| b1 | SPU_030131 | NA |
| b2 | SPU_000266 | none |
| b2 | SPU_000378 | Lv-Samd9L |
| b2 | SPU_000615 | NA |
| b2 | SPU_000728 | Lv-Hypp_1286 |
| b2 | SPU_000998 | Lv-Hypp_2978 |
| b2 | SPU_001157 | Lv-Hypp_326 |
| b2 | SPU_001348 | Lv-Hypp_331 |
| b2 | SPU_001494 | Lv-Tmem67 |
| b2 | SPU_001519 | Lv-Barx |
| b2 | SPU_001589 | none |
| b2 | SPU_001666 | Lv-Kcnk5L |
| b2 | SPU_001707 | Lv-Ank5 |
| b2 | SPU_001781 | Lv-Nlr20 |
| b2 | SPU_002144 | Lv-Clect_5 |
| b2 | SPU_002365 | Lv-Ints8-2 |
| b2 | SPU_002641 | Lv-Nlr12 |
| b2 | SPU_002682 | NA |
| b2 | SPU_002919 | none |
| b2 | SPU_002943 | Lv-Unc44L_29 |
| b2 | SPU_002995 | none |
| b2 | SPU_003024 | NA |
| b2 | SPU_003126 | Lv-Srcrr1 |
| b2 | SPU_003215 | Lv-GagpolL_91111 |
| b2 | SPU_004263 | Lv-Ugt_2 |

| | | |
|---|---|---|
| b2 | SPU_004264 | Lv-PppL_242 |
| b2 | SPU_004321 | Lv-Kiaa1012L |
| b2 | SPU_004903 | none |
| b2 | SPU_004956 | none |
| b2 | SPU_005089 | Lv-TlrP02 |
| b2 | SPU_005094 | Lv-Unc44L_41 |
| b2 | SPU_005147 | none |
| b2 | SPU_005184 | Lv-Hypp_1570 |
| b2 | SPU_005294 | Lv-Hypp_1576 |
| b2 | SPU_005413 | Lv-PolppL_48 |
| b2 | SPU_005695 | NA |
| b2 | SPU_005716 | Lv-Hypp_461 |
| b2 | SPU_005928 | Lv-Unc44L_46 |
| b2 | SPU_005976 | none |
| b2 | SPU_006006 | Lv-Rab40_1 |
| b2 | SPU_006315 | none |
| b2 | SPU_006538 | Lv-Srcr51 |
| b2 | SPU_006541 | none |
| b2 | SPU_006611 | Lv-TlrP03 |
| b2 | SPU_006780 | Lv-Hypp_1681 |
| b2 | SPU_007047 | Lv-EhH1_12 |
| b2 | SPU_007177 | Lv-Hypp_1702 |
| b2 | SPU_007476 | Lv-Lrr/Igr_17 |
| b2 | SPU_007499 | Lv-Acaa1b |
| b2 | SPU_007509 | NA |
| b2 | SPU_007648 | Lv-CubnL_9 |
| b2 | SPU_007759 | Lv-Abcc5A1 |
| b2 | SPU_007839 | none |
| b2 | SPU_007925 | none |
| b2 | SPU_008074 | Lv-Hypp_30 |
| b2 | SPU_008109 | none |
| b2 | SPU_008190 | Lv-Hrh3L_6 |
| b2 | SPU_008323 | Lv-Hypp_1777 |
| b2 | SPU_008359 | none |
| b2 | SPU_008421 | NA |
| b2 | SPU_008510 | Lv-Hypp_551 |
| b2 | SPU_008620 | NA |
| b2 | SPU_008829 | NA |

| | | |
|---|---|---|
| b2 | SPU_009459 | Lv-Tlr173 |
| b2 | SPU_009541 | none |
| b2 | SPU_009578 | Lv-Hypp_1859 |
| b2 | SPU_009581 | NA |
| b2 | SPU_009842 | Lv-Tk3 |
| b2 | SPU_010261 | none |
| b2 | SPU_010385 | Lv-Eea1 |
| b2 | SPU_010524 | Lv-Casp2L |
| b2 | SPU_010619 | Lv-Tlr057 |
| b2 | SPU_010711 | Lv-Hebp1 |
| b2 | SPU_011192 | Lv-Clect/Egf_14 |
| b2 | SPU_011215 | Lv-Manba_2 |
| b2 | SPU_011221 | Lv-Endrvt78 |
| b2 | SPU_011773 | Lv-Atl2 |
| b2 | SPU_011794 | none |
| b2 | SPU_011804 | Lv-Fn3/Egf_7 |
| b2 | SPU_011820 | Lv-Hypp_2001 |
| b2 | SPU_011924 | Lv-Gpr54L |
| b2 | SPU_011950 | Lv-Unc44L_78 |
| b2 | SPU_012218 | none |
| b2 | SPU_012463 | Lv-Clect/Fn3_16 |
| b2 | SPU_012763 | Lv-Chst1_1 |
| b2 | SPU_012812 | Lv-Dnah9_2 |
| b2 | SPU_013559 | Lv-Trrap_4 |
| b2 | SPU_013586 | Lv-Slc25a27 |
| b2 | SPU_013653 | none |
| b2 | SPU_013780 | Lv-Hypp_2107 |
| b2 | SPU_013806 | Lv-Unk_60 |
| b2 | SPU_013848 | Lv-PolppL_103 |
| b2 | SPU_014110 | NA |
| b2 | SPU_014545 | Lv-Hypp_757 |
| b2 | SPU_014809 | Lv-FoxL2 |
| b2 | SPU_014867 | none |
| b2 | SPU_014993 | Lv-Srcr121 |
| b2 | SPU_015096 | Lv-Cyp2L28 |
| b2 | SPU_015196 | NA |
| b2 | SPU_015340 | Lv-Nlr16 |
| b2 | SPU_015431 | Lv-Unc44L_108 |

| | | |
|---|---|---|
| b2 | SPU_015531 | none |
| b2 | SPU_015711 | Lv-Slc16a4L_3 |
| b2 | SPU_015728 | Lv-Efgib_2 |
| b2 | SPU_015803 | Lv-Clca_9 |
| b2 | SPU_015993 | Lv-Hypp_2242 |
| b2 | SPU_016097 | NA |
| b2 | SPU_016502 | none |
| b2 | SPU_016812 | Lv-Sstr4 |
| b2 | SPU_016996 | Lv-AkrL_2 |
| b2 | SPU_017330 | Lv-Hypp_2320 |
| b2 | SPU_017566 | Lv-Unc44_239 |
| b2 | SPU_017668 | NA |
| b2 | SPU_017739 | none |
| b2 | SPU_017970 | Lv-Hypp_903 |
| b2 | SPU_018006 | Lv-Gpr151L_11 |
| b2 | SPU_018509 | Lv-Ahctf1L |
| b2 | SPU_019159 | NA |
| b2 | SPU_019222 | Lv-Hypp_150 |
| b2 | SPU_019273 | Lv-MyID/G |
| b2 | SPU_019865 | NA |
| b2 | SPU_020314 | Lv-Hrh3_10 |
| b2 | SPU_020369 | Lv-Taar1L_3 |
| b2 | SPU_020545 | Lv-Adam15L2 |
| b2 | SPU_020700 | none |
| b2 | SPU_020957 | none |
| b2 | SPU_020976 | Lv-Unc44_149 |
| b2 | SPU_021415 | Lv-Tlr111 |
| b2 | SPU_021750 | Lv-Nox5_3 |
| b2 | SPU_021957 | none |
| b2 | SPU_022079 | Lv-Mystat1 |
| b2 | SPU_022106 | Lv-Gemin5-2 |
| b2 | SPU_022294 | Lv-Nlr39 |
| b2 | SPU_022497 | Lv-Ank8 |
| b2 | SPU_022618 | Lv-Unc44_162 |
| b2 | SPU_022720 | Lv-Hypp_2625 |
| b2 | SPU_022814 | Lv-Srcr197 |
| b2 | SPU_022931 | Lv-Mib1L_6 |
| b2 | SPU_022977 | none |

| | | |
|---|---|---|
| b2 | SPU_023365 | Lv-Tmtc3 |
| b2 | SPU_023513 | Lv-Ppk2L |
| b2 | SPU_023745 | Lv-NotchL_95 |
| b2 | SPU_023816 | NA |
| b2 | SPU_023822 | none |
| b2 | SPU_023840 | Lv-Srcr200 |
| b2 | SPU_024054 | Lv-Hypp_1103 |
| b2 | SPU_024166 | none |
| b2 | SPU_024395 | Lv-Camk4b |
| b2 | SPU_024410 | Lv-Htr1aL_7 |
| b2 | SPU_024566 | Lv-Cub/Gpcr |
| b2 | SPU_024889 | Lv-Taar7eL_4 |
| b2 | SPU_025180 | Lv-Mrc1L_12 |
| b2 | SPU_025214 | NA |
| b2 | SPU_025273 | none |
| b2 | SPU_025438 | NA |
| b2 | SPU_025473 | Lv-Hypp_2790 |
| b2 | SPU_025549 | none |
| b2 | SPU_025551 | none |
| b2 | SPU_025600 | Lv-Nlr147 |
| b2 | SPU_025805 | Lv-Hypp_1157 |
| b2 | SPU_026232 | Lv-Hypp_2829 |
| b2 | SPU_026334 | Lv-Csmd2L_2 |
| b2 | SPU_026532 | Lv-Tbc1d16_1 |
| b2 | SPU_026854 | Lv-Polr2b_1 |
| b2 | SPU_026939 | Lv-Endrvt10 |
| b2 | SPU_027011 | Lv-Hypp_2879 |
| b2 | SPU_027074 | Lv-Hypp_1196 |
| b2 | SPU_027094 | Lv-Rxfp1_4 |
| b2 | SPU_027205 | Lv-Gor |
| b2 | SPU_027938 | Lv-Dynlc2-6b |
| b2 | SPU_028335 | Lv-Hrh1L_8 |
| b2 | SPU_028797 | Lv-Abcc9b |
| b2 | SPU_028798 | Lv-Abcc9_1 |
| b2 | SPU_028864 | Lv-Prss7L_2 |
| b2 | SPU_028907 | none |
| b2 | SPU_030046 | Lv-CbapsL_1 |
| b3 | SPU_000254 | Lv-Trim3l |

| | | |
|---|---|---|
| b3 | SPU_000697 | Lv-Unc44L_5 |
| b3 | SPU_000869 | none |
| b3 | SPU_001099 | none |
| b3 | SPU_001326 | none |
| b3 | SPU_001463 | NA |
| b3 | SPU_001509 | Lv-EhH4_2 |
| b3 | SPU_001537 | Lv-Gnrhr |
| b3 | SPU_001627 | Lv-Gpr101L |
| b3 | SPU_001820 | Lv-Unc44L_18 |
| b3 | SPU_002047 | none |
| b3 | SPU_002261 | Lv-Hypp_1385 |
| b3 | SPU_002310 | none |
| b3 | SPU_002416 | NA |
| b3 | SPU_002423 | Lv-Nlr56 |
| b3 | SPU_002624 | none |
| b3 | SPU_002676 | NA |
| b3 | SPU_002706 | Lv-Znfx1_5 |
| b3 | SPU_002778 | Lv-Unc44L_26 |
| b3 | SPU_003393 | Lv-Gpr52L |
| b3 | SPU_003526 | Lv-Srcr26 |
| b3 | SPU_003557 | none |
| b3 | SPU_003965 | Lv-Srcr31 |
| b3 | SPU_004150 | Lv-Tlr020 |
| b3 | SPU_004201 | Lv-Hypp_1505 |
| b3 | SPU_004244 | none |
| b3 | SPU_004359 | none |
| b3 | SPU_004387 | Lv-Hypp_1520 |
| b3 | SPU_004539 | Lv-Hypp_1533 |
| b3 | SPU_004636 | Lv-Hypp_1536 |
| b3 | SPU_004708 | Lv-Gpr51_3 |
| b3 | SPU_004726 | Lv-Sr/Lrr/Gpcr_2 |
| b3 | SPU_004800 | none |
| b3 | SPU_004931 | Lv-Hypp_1553 |
| b3 | SPU_005060 | Lv-Fn3_21 |
| b3 | SPU_005075 | Lv-Sacs |
| b3 | SPU_005154 | Lv-Srcr41 |
| b3 | SPU_005521 | Lv-Cyp3L13 |
| b3 | SPU_005755 | Lv-Znfx1-18 |

| | | |
|---|---|---|
| b3 | SPU_005853 | Lv-Znfx1_6 |
| b3 | SPU_006097 | none |
| b3 | SPU_006219 | Lv-Nlr198 |
| b3 | SPU_006408 | Lv-Hypp_1660 |
| b3 | SPU_006477 | none |
| b3 | SPU_006540 | Lv-Lrr/Gpcr_15 |
| b3 | SPU_006970 | Lv-KctdL |
| b3 | SPU_007123 | Lv-Unc44L_56 |
| b3 | SPU_007430 | Lv-Tlr038 |
| b3 | SPU_007467 | Lv-Taar6L |
| b3 | SPU_007636 | Lv-Z306 |
| b3 | SPU_007781 | Lv-Srcr58 |
| b3 | SPU_007896 | Lv-Srcr63 |
| b3 | SPU_007901 | Lv-Hypp_531 |
| b3 | SPU_007968 | none |
| b3 | SPU_007988 | NA |
| b3 | SPU_007996 | Lv-RtL_16 |
| b3 | SPU_008021 | none |
| b3 | SPU_008036 | none |
| b3 | SPU_008060 | Lv-Kctd21L |
| b3 | SPU_008124 | Lv-Hypp_1766 |
| b3 | SPU_008263 | Lv-Tstsk1_1 |
| b3 | SPU_008482 | NA |
| b3 | SPU_008522 | Lv-SacSl_2 |
| b3 | SPU_008524 | Lv-Hypp_1790 |
| b3 | SPU_008578 | NA |
| b3 | SPU_008585 | Lv-Man1a2 |
| b3 | SPU_008814 | Lv-Hypp_1808 |
| b3 | SPU_009314 | Lv-Hypp_575 |
| b3 | SPU_009602 | NA |
| b3 | SPU_009764 | Lv-Krp95s_2 |
| b3 | SPU_009927 | Lv-AnL10 |
| b3 | SPU_009988 | Lv-Ig/Srcr1 |
| b3 | SPU_010053 | Lv-Nlr102 |
| b3 | SPU_010512 | Lv-Hypp_1922 |
| b3 | SPU_010603 | none |
| b3 | SPU_010880 | Lv-Gpr84L_3 |
| b3 | SPU_010925 | none |

| | | |
|---|---|---|
| b3 | SPU_010953 | Lv-Srcr94 |
| b3 | SPU_011004 | Lv-Unc44L_68 |
| b3 | SPU_011225 | NA |
| b3 | SPU_011275 | Lv-Alk_1 |
| b3 | SPU_011465 | Lv-Ccdc57 |
| b3 | SPU_011482 | Lv-Hypp_1981 |
| b3 | SPU_011540 | Lv-Tlr064 |
| b3 | SPU_011843 | Lv-Serrp2 |
| b3 | SPU_012059 | Lv-Unc44L_80 |
| b3 | SPU_012274 | Lv-Chrm4L_1 |
| b3 | SPU_012378 | Lv-TaarL |
| b3 | SPU_013120 | Lv-Hypp_2070 |
| b3 | SPU_013218 | Lv-C3orf32L |
| b3 | SPU_013313 | Lv-Polr1a |
| b3 | SPU_013462 | none |
| b3 | SPU_013532 | Lv-B3gnt5_3 |
| b3 | SPU_013766 | none |
| b3 | SPU_013847 | Lv-PolppL_102 |
| b3 | SPU_013943 | Lv-Slc7a6_7 |
| b3 | SPU_013978 | Lv-Igf_18 |
| b3 | SPU_013986 | Lv-Chst2L_5 |
| b3 | SPU_014023 | Lv-Slc16a14L_4 |
| b3 | SPU_014039 | Lv-TlrL_5 |
| b3 | SPU_014292 | Lv-Unc44_232 |
| b3 | SPU_014338 | NA |
| b3 | SPU_014599 | NA |
| b3 | SPU_014661 | Lv-Unc44L_99 |
| b3 | SPU_014814 | none |
| b3 | SPU_015030 | none |
| b3 | SPU_015499 | none |
| b3 | SPU_015616 | Lv-Hypp_3021 |
| b3 | SPU_015868 | NA |
| b3 | SPU_015886 | Lv-Adrb2L_10 |
| b3 | SPU_016036 | Lv-Hypp_2995 |
| b3 | SPU_016313 | none |
| b3 | SPU_016347 | Lv-Kctd4_1 |
| b3 | SPU_016445 | Lv-Myor4 |
| b3 | SPU_016672 | none |

| | | |
|---|---|---|
| b3 | SPU_016701 | Lv-Rdh8_10 |
| b3 | SPU_016724 | Lv-Slc16a12L_12 |
| b3 | SPU_016870 | Lv-Hypp_2284 |
| b3 | SPU_017180 | Lv-Tlr088 |
| b3 | SPU_017183 | Lv-Hypp_864 |
| b3 | SPU_017332 | Lv-Hypp_874 |
| b3 | SPU_017517 | Lv-Fbxo39 |
| b3 | SPU_017674 | none |
| b3 | SPU_017706 | Lv-Hypp_891 |
| b3 | SPU_017763 | Lv-Taar7bL |
| b3 | SPU_017828 | Lv-SacSl |
| b3 | SPU_017865 | Lv-Gpr1L_5 |
| b3 | SPU_017928 | Lv-Unc44_132 |
| b3 | SPU_017994 | Lv-Hypp_2364 |
| b3 | SPU_018055 | Lv-Tlr090 |
| b3 | SPU_018839 | Lv-Hypp_923 |
| b3 | SPU_019186 | NA |
| b3 | SPU_019268 | Lv-Shox |
| b3 | SPU_019313 | Lv-Hypp_934 |
| b3 | SPU_019382 | Lv-Tmem26_4 |
| b3 | SPU_019564 | Lv-Hypp_156 |
| b3 | SPU_019565 | Lv-Hypp_939 |
| b3 | SPU_019924 | Lv-AnkyL |
| b3 | SPU_020092 | none |
| b3 | SPU_020277 | none |
| b3 | SPU_020355 | Lv-Hypp_2502 |
| b3 | SPU_020372 | Lv-PppL_272 |
| b3 | SPU_020650 | Lv-Srcr160 |
| b3 | SPU_020936 | NA |
| b3 | SPU_020984 | none |
| b3 | SPU_020997 | Lv-Tlr107 |
| b3 | SPU_021053 | Lv-B3galt1L_4 |
| b3 | SPU_021152 | Lv-Unc44_152 |
| b3 | SPU_021224 | none |
| b3 | SPU_021249 | NA |
| b3 | SPU_021617 | NA |
| b3 | SPU_021745 | Lv-Fn3/Igf_40 |
| b3 | SPU_021994 | none |

| | | |
|---|---|---|
| b3 | SPU_022001 | Lv-Nlr137 |
| b3 | SPU_022278 | none |
| b3 | SPU_022288 | Lv-Srcr188 |
| b3 | SPU_022602 | Lv-Clca_15 |
| b3 | SPU_022844 | NA |
| b3 | SPU_023031 | NA |
| b3 | SPU_023520 | Lv-Cd2bp2_1 |
| b3 | SPU_023747 | none |
| b3 | SPU_024062 | Lv-Tlr122 |
| b3 | SPU_024118 | Lv-Hypp_2711 |
| b3 | SPU_024290 | NA |
| b3 | SPU_024359 | Lv-Herc1-3 |
| b3 | SPU_024759 | Lv-Hypp_1125 |
| b3 | SPU_024858 | none |
| b3 | SPU_024937 | none |
| b3 | SPU_025135 | NA |
| b3 | SPU_025360 | Lv-SacSl_1 |
| b3 | SPU_025440 | none |
| b3 | SPU_025445 | Lv-Slc5a9L_1 |
| b3 | SPU_025594 | Lv-Rnh1L |
| b3 | SPU_025918 | none |
| b3 | SPU_026120 | Lv-Hypp_1168 |
| b3 | SPU_026321 | Lv-Fn3_41 |
| b3 | SPU_026367 | none |
| b3 | SPU_026507 | Lv-Egf/Lnb/7Tm/Gpcr_2 |
| b3 | SPU_026614 | Lv-My8 |
| b3 | SPU_026702 | Lv-Prrdh_4 |
| b3 | SPU_026720 | NA |
| b3 | SPU_027095 | Lv-Dmbt1L |
| b3 | SPU_027177 | Lv-Hrh2L_9 |
| b3 | SPU_027249 | none |
| b3 | SPU_027300 | Lv-Nlr170 |
| b3 | SPU_027762 | Lv-Hypp_274 |
| b3 | SPU_028043 | none |
| b3 | SPU_028380 | Lv-TlrP36 |
| b3 | SPU_028879 | Lv-Hypp_3026 |
| b3 | SPU_030071 | Lv-Vc1_2 |
| c1 | SPU_000192 | Lv-Clect_61 |

| | | |
|---|---|---|
| c1 | SPU_000239 | Lv-Rrm |
| c1 | SPU_000325 | Lv-Lgr6L_2 |
| c1 | SPU_000394 | none |
| c1 | SPU_000438 | Lv-p58-b |
| c1 | SPU_000439 | Lv-p58-a |
| c1 | SPU_000537 | Lv-Dlk1L |
| c1 | SPU_000634 | Lv-Hecw1L |
| c1 | SPU_000662 | Lv-Hypp_1271 |
| c1 | SPU_000678 | NA |
| c1 | SPU_000710 | Lv-B3galt2 |
| c1 | SPU_000947 | none |
| c1 | SPU_000954 | none |
| c1 | SPU_001059 | Lv-Mab21l2 |
| c1 | SPU_001113 | Lv-Sna |
| c1 | SPU_001152 | Lv-Mif7 |
| c1 | SPU_001210 | NA |
| c1 | SPU_001343 | none |
| c1 | SPU_001574 | none |
| c1 | SPU_001601 | Lv-Srcr16 |
| c1 | SPU_001739 | Lv-Unc4-1 |
| c1 | SPU_001802 | Lv-IgfalsL |
| c1 | SPU_002272 | Lv-Nlr105 |
| c1 | SPU_002677 | Lv-Ptf1a |
| c1 | SPU_002881 | Lv-Etfdh |
| c1 | SPU_002895 | Lv-Pks1 |
| c1 | SPU_003010 | Lv-Soat |
| c1 | SPU_003105 | Lv-Hypp_1433 |
| c1 | SPU_003143 | Lv-Hypp_1435 |
| c1 | SPU_003224 | none |
| c1 | SPU_003306 | NA |
| c1 | SPU_003511 | Lv-Cryz |
| c1 | SPU_003597 | NA |
| c1 | SPU_003640 | Lv-Nlr116 |
| c1 | SPU_003785 | Lv-Odc1 |
| c1 | SPU_003819 | Lv-Tmem195 |
| c1 | SPU_003885 | Lv-Hypp_1486 |
| c1 | SPU_003900 | Lv-MyV_2 |
| c1 | SPU_003941 | Lv-Slc15a4 |

| | | |
|---|---|---|
| c1 | SPU_004137 | Lv-Ephx2_1 |
| c1 | SPU_004155 | none |
| c1 | SPU_004158 | none |
| c1 | SPU_004159 | none |
| c1 | SPU_004499 | Lv-Acacb |
| c1 | SPU_004731 | none |
| c1 | SPU_004746 | Lv-Fn3/Igf_29 |
| c1 | SPU_004810 | none |
| c1 | SPU_005157 | Lv-Fpgs_6 |
| c1 | SPU_005170 | Lv-Hox7_1 |
| c1 | SPU_005365 | Lv-Hypp_444 |
| c1 | SPU_005398 | none |
| c1 | SPU_005569 | Lv-Opsin1 |
| c1 | SPU_005683 | Lv-Hypp_1597 |
| c1 | SPU_005846 | Lv-Hrh4L |
| c1 | SPU_006052 | NA |
| c1 | SPU_006187 | Lv-Sult1c_2 |
| c1 | SPU_006218 | Lv-Tlr032 |
| c1 | SPU_006247 | Lv-Hypp_1642 |
| c1 | SPU_006347 | Lv-Slc16a7_2 |
| c1 | SPU_006431 | Lv-Galr2L_6 |
| c1 | SPU_006470 | Lv-Kcnc2 |
| c1 | SPU_006545 | Lv-Cryz_1 |
| c1 | SPU_006559 | NA |
| c1 | SPU_006676 | Lv-FoxA |
| c1 | SPU_006814 | Lv-Hairy2/4 |
| c1 | SPU_007088 | Lv-Sarmr14 |
| c1 | SPU_007110 | Lv-Srcr54 |
| c1 | SPU_007113 | Lv-Nlr98 |
| c1 | SPU_007226 | Lv-Eif3e |
| c1 | SPU_007265 | Lv-Wdr36 |
| c1 | SPU_007484 | Lv-Cycpln |
| c1 | SPU_007765 | Lv-Chst1 |
| c1 | SPU_008094 | Lv-St8sia4_1 |
| c1 | SPU_008204 | none |
| c1 | SPU_008658 | Lv-Cara7LB |
| c1 | SPU_008784 | Lv-Hypp_1803 |
| c1 | SPU_008887 | Lv-Unc44L_61 |

| | | |
|---|---|---|
| c1 | SPU_008985 | NA |
| c1 | SPU_009043 | NA |
| c1 | SPU_009048 | Lv-Pax2 |
| c1 | SPU_009100 | Lv-Erg1 |
| c1 | SPU_009201 | none |
| c1 | SPU_009320 | Lv-blaZ |
| c1 | SPU_009352 | none |
| c1 | SPU_009609 | Lv-Slc7a11 |
| c1 | SPU_009616 | NA |
| c1 | SPU_009719 | Lv-Oprd1L_1 |
| c1 | SPU_010051 | NA |
| c1 | SPU_010207 | Lv-Sstr4L_1 |
| c1 | SPU_010305 | Lv-Hnf1aL |
| c1 | SPU_010330 | NA |
| c1 | SPU_010351 | Lv-IrxA |
| c1 | SPU_010403 | Lv-FoxY |
| c1 | SPU_010560 | Lv-Lgr5L |
| c1 | SPU_010561 | Lv-Lrr/Gpcr_8 |
| c1 | SPU_010575 | Lv-Tlr056 |
| c1 | SPU_010591 | Lv-Mdn1_4 |
| c1 | SPU_010650 | Lv-Cycs |
| c1 | SPU_010651 | Lv-Cycs_1 |
| c1 | SPU_011381 | none |
| c1 | SPU_011616 | Lv-Prkrir |
| c1 | SPU_011798 | none |
| c1 | SPU_011951 | NA |
| c1 | SPU_012063 | Lv-Hypp_667 |
| c1 | SPU_012154 | NA |
| c1 | SPU_012190 | NA |
| c1 | SPU_012313 | none |
| c1 | SPU_012348 | Lv-Fmo5_1 |
| c1 | SPU_012484 | Lv-Hypp_2034 |
| c1 | SPU_012491 | Lv-Nk1 |
| c1 | SPU_012615 | Lv-Slc16a12_1 |
| c1 | SPU_012929 | Lv-Hypp_3018 |
| c1 | SPU_013031 | none |
| c1 | SPU_013228 | NA |
| c1 | SPU_013294 | none |

| | | |
|---|---|---|
| c1 | SPU_013473 | Lv-Acaca_1 |
| c1 | SPU_013595 | Lv-Gpr84L_4 |
| c1 | SPU_013641 | NA |
| c1 | SPU_013783 | Lv-Mc5rL |
| c1 | SPU_013801 | NA |
| c1 | SPU_013821 | Lv-Msp130_1 |
| c1 | SPU_013823 | Lv-Msp130r3 |
| c1 | SPU_013933 | none |
| c1 | SPU_014123 | NA |
| c1 | SPU_014219 | Lv-Clect/Tm_4 |
| c1 | SPU_014270 | none |
| c1 | SPU_014296 | none |
| c1 | SPU_014539 | Lv-Pax2-5-8 |
| c1 | SPU_014679 | Lv-Unc44L_100 |
| c1 | SPU_014709 | Lv-LsampL |
| c1 | SPU_014720 | NA |
| c1 | SPU_014822 | none |
| c1 | SPU_014828 | Lv-Astacin4 |
| c1 | SPU_014947 | Lv-Fmo2_2 |
| c1 | SPU_015076 | Lv-Sstr4L_4 |
| c1 | SPU_015078 | Lv-Galr2L_20 |
| c1 | SPU_015289 | none |
| c1 | SPU_015362 | Lv-Z167 |
| c1 | SPU_015371 | Lv-Acel1 |
| c1 | SPU_015640 | Lv-ScratchX |
| c1 | SPU_015978 | NA |
| c1 | SPU_015982 | SpGsc |
| c1 | SPU_015990 | none |
| c1 | SPU_016235 | Lv-Kifcb |
| c1 | SPU_016248 | NA |
| c1 | SPU_016348 | Lv-Kctd21L_5 |
| c1 | SPU_016582 | none |
| c1 | SPU_016587 | Lv-Fpgs_3 |
| c1 | SPU_016638 | Lv-Fut8L_1 |
| c1 | SPU_016984 | none |
| c1 | SPU_017181 | none |
| c1 | SPU_017328 | NA |
| c1 | SPU_017379 | Lv-Six1/2 |

| | | |
|---|---|---|
| c1 | SPU_017442 | none |
| c1 | SPU_017480 | Lv-Hypp_880 |
| c1 | SPU_017625 | Lv-Hypp_2347 |
| c1 | SPU_017862 | Lv-Gpr1L_2 |
| c1 | SPU_017992 | NA |
| c1 | SPU_018352 | Lv-Unk_63 |
| c1 | SPU_018403 | none |
| c1 | SPU_018406 | NA |
| c1 | SPU_018408 | Lv-P16 |
| c1 | SPU_018436 | none |
| c1 | SPU_018441 | Lv-Snd1_1 |
| c1 | SPU_018443 | NA |
| c1 | SPU_018499 | none |
| c1 | SPU_018516 | Lv-Prss12L_2 |
| c1 | SPU_018542 | Lv-Sdr |
| c1 | SPU_018824 | NA |
| c1 | SPU_019053 | Lv-Ank2_1 |
| c1 | SPU_019109 | none |
| c1 | SPU_019417 | Lv-Acly_2 |
| c1 | SPU_019761 | NA |
| c1 | SPU_019843 | NA |
| c1 | SPU_020035 | Lv-Mif4 |
| c1 | SPU_020036 | Lv-Mif5 |
| c1 | SPU_020188 | none |
| c1 | SPU_020705 | NA |
| c1 | SPU_020734 | Lv-Hypp_2522 |
| c1 | SPU_020928 | Lv-B3galt6 |
| c1 | SPU_021056 | Lv-Nme1 |
| c1 | SPU_021189 | Lv-Ddesat |
| c1 | SPU_021192 | Lv-Hnf4 |
| c1 | SPU_021252 | Lv-Fn3_6 |
| c1 | SPU_021284 | Lv-Unc44_153 |
| c1 | SPU_021297 | none |
| c1 | SPU_021314 | Lv-Mfsd6l_1 |
| c1 | SPU_021457 | Lv-Srcr165 |
| c1 | SPU_021659 | Lv-Hypp_1028 |
| c1 | SPU_021841 | Lv-Sarmr6 |
| c1 | SPU_022067 | none |

| | | |
|---|---|---|
| c1 | SPU_022203 | Lv-Slc10a2L_3 |
| c1 | SPU_022276 | Lv-Rrp12_1 |
| c1 | SPU_022337 | Lv-Rxfp2_4 |
| c1 | SPU_022386 | Lv-Eif3m |
| c1 | SPU_022391 | Lv-A1cfL |
| c1 | SPU_022420 | Lv-Znfx1-19 |
| c1 | SPU_022694 | Lv-Naalad2_1 |
| c1 | SPU_022719 | Lv-Gal/lec12 |
| c1 | SPU_022903 | NA |
| c1 | SPU_023050 | Lv-Hypp_2645 |
| c1 | SPU_023098 | Lv-Abcc9c |
| c1 | SPU_023212 | Lv-Hypp_1077 |
| c1 | SPU_023220 | Lv-Ft1a |
| c1 | SPU_023340 | Lv-Hypp_2668 |
| c1 | SPU_023430 | Lv-Alox5_9 |
| c1 | SPU_023544 | Lv-Tlr121 |
| c1 | SPU_023612 | none |
| c1 | SPU_023678 | Lv-Enpp7 |
| c1 | SPU_023738 | Lv-Fn3f_49 |
| c1 | SPU_024031 | Lv-Znfx1-20 |
| c1 | SPU_024043 | Lv-Hs3st1L_2 |
| c1 | SPU_024363 | Lv-Hypp_1113 |
| c1 | SPU_024375 | NA |
| c1 | SPU_024655 | Lv-Gpr54L_3 |
| c1 | SPU_024662 | none |
| c1 | SPU_025121 | NA |
| c1 | SPU_025145 | Lv-B4galt5_3 |
| c1 | SPU_025235 | Lv-Npnt |
| c1 | SPU_025314 | Lv-Npffr2L_4 |
| c1 | SPU_025384 | Lv-PppL_228 |
| c1 | SPU_025517 | Lv-Cd157 |
| c1 | SPU_025601 | Lv-Dr_1 |
| c1 | SPU_025813 | Lv-Lsm2 |
| c1 | SPU_025835 | NA |
| c1 | SPU_025912 | NA |
| c1 | SPU_025934 | NA |
| c1 | SPU_025974 | none |
| c1 | SPU_026167 | Lv-Galr2L_32 |

| | | |
|---|---|---|
| c1 | SPU_026283 | Lv-Hypp_2833 |
| c1 | SPU_026298 | Lv-Lrr1 |
| c1 | SPU_026506 | NA |
| c1 | SPU_026566 | Lv-Hypp_2852 |
| c1 | SPU_026617 | Lv-Naalad2L |
| c1 | SPU_026831 | NA |
| c1 | SPU_026877 | Lv-Irf4 |
| c1 | SPU_026948 | none |
| c1 | SPU_027441 | NA |
| c1 | SPU_027584 | Lv-Surreal/Gpcr93 |
| c1 | SPU_027757 | Lv-Endrvt53 |
| c1 | SPU_027760 | none |
| c1 | SPU_027772 | Lv-Unc44_184 |
| c1 | SPU_027819 | Lv-Pofut1 |
| c1 | SPU_027923 | Lv-B4galt5_4 |
| c1 | SPU_028057 | NA |
| c1 | SPU_028389 | Lv-Uqcrb_1 |
| c1 | SPU_028669 | Lv-Srcr226 |
| c1 | SPU_028705 | NA |
| c1 | SPU_028788 | Lv-Hypp_290 |
| c1 | SPU_028804 | Lv-Srcr228 |
| c1 | SPU_030076 | Lv-Tlr224 |
| c1 | SPU_030193 | NA |
| c2 | SPU_000161 | NA |
| c2 | SPU_000173 | Lv-Slc30a3 |
| c2 | SPU_000407 | none |
| c2 | SPU_000580 | Lv-Srcr3 |
| c2 | SPU_001086 | none |
| c2 | SPU_001230 | none |
| c2 | SPU_001248 | Lv-Hypp_1319 |
| c2 | SPU_001461 | Lv-Ptprol |
| c2 | SPU_001523 | none |
| c2 | SPU_001549 | Lv-Nlr169 |
| c2 | SPU_001618 | none |
| c2 | SPU_001866 | none |
| c2 | SPU_002506 | none |
| c2 | SPU_003539 | Lv-Nlr3 |
| c2 | SPU_003681 | Lv-Atol2 |

| | | |
|---|---|---|
| c2 | SPU_003904 | Lv-Slc10a2_13 |
| c2 | SPU_004114 | Lv-AnL1 |
| c2 | SPU_004115 | Lv-AnL2 |
| c2 | SPU_004116 | none |
| c2 | SPU_004117 | Lv-AnL4 |
| c2 | SPU_004481 | Lv-Ugtla9-2 |
| c2 | SPU_004560 | Lv-Hypp_1534 |
| c2 | SPU_004593 | Lv-Slc16a4_1 |
| c2 | SPU_004759 | Lv-Rergl7 |
| c2 | SPU_004832 | Lv-Gluld1_16 |
| c2 | SPU_005596 | Lv-Clect/Egf_7 |
| c2 | SPU_005900 | Lv-Fn3/Igf_30 |
| c2 | SPU_006046 | none |
| c2 | SPU_006053 | NA |
| c2 | SPU_006100 | Lv-TtnL |
| c2 | SPU_006232 | Lv-MyoD2 |
| c2 | SPU_006359 | none |
| c2 | SPU_006496 | none |
| c2 | SPU_006507 | Lv-Chrm1L_1 |
| c2 | SPU_006832 | Lv-Unc44L_55 |
| c2 | SPU_006885 | Lv-Hypp_1686 |
| c2 | SPU_006914 | Lv-Tssk2 |
| c2 | SPU_006982 | none |
| c2 | SPU_007001 | NA |
| c2 | SPU_007298 | Lv-Unc44L_57 |
| c2 | SPU_007600 | none |
| c2 | SPU_007717 | Lv-Hypp_526 |
| c2 | SPU_007900 | Lv-Srcr66 |
| c2 | SPU_007907 | Lv-Slco10a2 |
| c2 | SPU_008030 | NA |
| c2 | SPU_008463 | none |
| c2 | SPU_008478 | Lv-Tmem26_1 |
| c2 | SPU_008690 | Lv-Fram1 |
| c2 | SPU_008707 | Lv-Nlr149 |
| c2 | SPU_008933 | Lv-Kyl |
| c2 | SPU_009479 | none |
| c2 | SPU_009649 | Lv-Hypp_584 |
| c2 | SPU_009776 | Lv-Hmcn1L |

| | | |
|---|---|---|
| c2 | SPU_009789 | none |
| c2 | SPU_009987 | NA |
| c2 | SPU_009997 | Lv-Unc44L_65 |
| c2 | SPU_010213 | Lv-St6galnac5_3 |
| c2 | SPU_010366 | NA |
| c2 | SPU_010582 | Lv-Tif1 |
| c2 | SPU_010681 | Lv-Hypp_624 |
| c2 | SPU_010715 | none |
| c2 | SPU_010832 | NA |
| c2 | SPU_010878 | NA |
| c2 | SPU_010989 | Lv-Hypp_1952 |
| c2 | SPU_010999 | NA |
| c2 | SPU_011149 | NA |
| c2 | SPU_011344 | Lv-Fn3/Egff_21 |
| c2 | SPU_011439 | NA |
| c2 | SPU_011492 | Lv-CubnL_3 |
| c2 | SPU_011501 | none |
| c2 | SPU_011718 | Lv-Aars_3 |
| c2 | SPU_011742 | Lv-Stk4L |
| c2 | SPU_012153 | Lv-Unc44L_82 |
| c2 | SPU_012305 | NA |
| c2 | SPU_012458 | NA |
| c2 | SPU_012586 | Lv-Fax1 |
| c2 | SPU_012735 | NA |
| c2 | SPU_012738 | Lv-Hypp_2048 |
| c2 | SPU_013234 | none |
| c2 | SPU_013894 | NA |
| c2 | SPU_013927 | NA |
| c2 | SPU_014177 | Lv-Lbx |
| c2 | SPU_014224 | NA |
| c2 | SPU_014451 | Lv-KifC1L |
| c2 | SPU_014600 | Lv-Unc44_233 |
| c2 | SPU_014647 | Lv-C20orf152L |
| c2 | SPU_014708 | none |
| c2 | SPU_015172 | Lv-C7orf63_1 |
| c2 | SPU_015269 | Lv-Ank6 |
| c2 | SPU_015439 | Lv-Hypp_784 |
| c2 | SPU_015511 | Lv-Tlr184 |

| | | |
|---|---|---|
| c2 | SPU_015539 | Lv-Srcr126 |
| c2 | SPU_015604 | Lv-ReelerL_3 |
| c2 | SPU_015974 | Lv-Tdc1 |
| c2 | SPU_016468 | Lv-Tlr085 |
| c2 | SPU_016488 | NA |
| c2 | SPU_016819 | NA |
| c2 | SPU_017524 | Lv-FUT |
| c2 | SPU_017549 | Lv-Hypp_2340 |
| c2 | SPU_017810 | Lv-Clect_36 |
| c2 | SPU_018164 | Lv-Mvp_2 |
| c2 | SPU_018241 | Lv-Slc16a14L_5 |
| c2 | SPU_018416 | Lv-Prss23L |
| c2 | SPU_018519 | Lv-Tlr098 |
| c2 | SPU_018539 | Lv-Hypp_916 |
| c2 | SPU_018651 | Lv-Unc44_134 |
| c2 | SPU_018951 | Lv-Prx1 |
| c2 | SPU_019375 | Lv-AplnrL_2 |
| c2 | SPU_019471 | Lv-Ugt1L_7 |
| c2 | SPU_019852 | Lv-Ptprlec1 |
| c2 | SPU_020183 | Lv-Unc44_143 |
| c2 | SPU_020556 | none |
| c2 | SPU_020652 | Lv-Tlr206 |
| c2 | SPU_020712 | Lv-Clca_14 |
| c2 | SPU_020819 | NA |
| c2 | SPU_020821 | NA |
| c2 | SPU_021002 | Lv-Slsp_14 |
| c2 | SPU_021245 | Lv-GagpolL_8 |
| c2 | SPU_021293 | Lv-Iqgapph |
| c2 | SPU_021528 | Lv-Galr2L_27 |
| c2 | SPU_021785 | Lv-Nlgn4xL |
| c2 | SPU_021818 | Lv-Exoc8_1 |
| c2 | SPU_021863 | NA |
| c2 | SPU_021996 | Lv-Lgsn |
| c2 | SPU_021999 | Lv-Hypp_1035 |
| c2 | SPU_022246 | NA |
| c2 | SPU_022277 | NA |
| c2 | SPU_022387 | none |
| c2 | SPU_022405 | NA |

| | | |
|---|---|---|
| c2 | SPU_022573 | Lv-Nk7 |
| c2 | SPU_022758 | Lv-Hypp_2631 |
| c2 | SPU_022769 | Lv-Poln |
| c2 | SPU_022816 | Lv-Alx4 |
| c2 | SPU_023116 | NA |
| c2 | SPU_023277 | Lv-Phf10_1 |
| c2 | SPU_023349 | Lv-Slc7_1 |
| c2 | SPU_023552 | Lv-Endrvt106 |
| c2 | SPU_023561 | NA |
| c2 | SPU_023721 | Lv-TpaseL_16 |
| c2 | SPU_023943 | NA |
| c2 | SPU_024121 | none |
| c2 | SPU_024386 | Lv-Tlr128 |
| c2 | SPU_024843 | none |
| c2 | SPU_024882 | Lv-Fn3/Igf_26 |
| c2 | SPU_025060 | none |
| c2 | SPU_025178 | Lv-Ly/Egf/Sea_1 |
| c2 | SPU_025204 | Lv-Nlr90 |
| c2 | SPU_025351 | Lv-Hypp_1144 |
| c2 | SPU_025446 | Lv-Adra1L_4 |
| c2 | SPU_025731 | Lv-Lrr/Igr_13 |
| c2 | SPU_025902 | Lv-Itih_3 |
| c2 | SPU_025966 | Lv-Lrr15L |
| c2 | SPU_026044 | Lv-Unc44_255 |
| c2 | SPU_026373 | Lv-Cyp2L62 |
| c2 | SPU_026849 | Lv-Srcr216 |
| c2 | SPU_027259 | Lv-PppL_153 |
| c2 | SPU_027333 | Lv-Unc44_181 |
| c2 | SPU_027571 | none |
| c2 | SPU_027668 | Lv-Unc44_183 |
| c2 | SPU_027836 | none |
| c2 | SPU_027906 | Lv-C-lectin/PMC1 |
| c2 | SPU_028056 | Lv-Galr2L_36 |
| c2 | SPU_028279 | none |
| c2 | SPU_028418 | NA |
| c2 | SPU_028751 | none |
| c2 | SPU_028752 | none |
| c2 | SPU_028755 | none |

| | | |
|---|---|---|
| c2 | SPU_030013 | Lv-MeprinAaL |
| c2 | SPU_030198 | Lv-Il17-20 |
| c3 | SPU_000030 | Lv-Gpr21_1 |
| c3 | SPU_000160 | none |
| c3 | SPU_000271 | Lv-Clect/Egf_4 |
| c3 | SPU_000368 | Lv-Hypp_1253 |
| c3 | SPU_000653 | Lv-Echn16 |
| c3 | SPU_000715 | Lv-PolppL-2 |
| c3 | SPU_000746 | Lv-Cyp2L1p |
| c3 | SPU_000951 | Lv-St8sia4 |
| c3 | SPU_000990 | Lv-Atol1 |
| c3 | SPU_001215 | none |
| c3 | SPU_001386 | Lv-Hypp_1324 |
| c3 | SPU_001458 | Lv-Tlr159 |
| c3 | SPU_001674 | Lv-Htr7L |
| c3 | SPU_001763 | Lv-Srcr19 |
| c3 | SPU_001951 | Lv-Tacr2L_2 |
| c3 | SPU_001981 | Lv-Ccdc151L |
| c3 | SPU_001988 | Lv-Unc44L_20 |
| c3 | SPU_002005 | Lv-Gal/lec |
| c3 | SPU_002007 | Lv-C20orf152 |
| c3 | SPU_002037 | Lv-Ube4A_1 |
| c3 | SPU_002084 | none |
| c3 | SPU_002224 | Lv-Tlr013 |
| c3 | SPU_002255 | Lv-Hypp_1383 |
| c3 | SPU_002312 | Lv-Pip5k3L |
| c3 | SPU_002550 | Lv-MacpfB.1 |
| c3 | SPU_002739 | Lv-Tmem26-2 |
| c3 | SPU_002756 | Lv-St6galnac2 |
| c3 | SPU_002762 | Lv-Ankrd27-2 |
| c3 | SPU_002763 | Lv-Tie2L |
| c3 | SPU_002797 | none |
| c3 | SPU_002957 | Lv-Hypp_372 |
| c3 | SPU_002985 | Lv-Kif3a |
| c3 | SPU_003384 | Lv-Srcr25 |
| c3 | SPU_003542 | none |
| c3 | SPU_003620 | none |
| c3 | SPU_003667 | Lv-Slc5a2 |

| | | |
|---|---|---|
| c3 | SPU_003695 | Lv-Hypp_1475 |
| c3 | SPU_003878 | none |
| c3 | SPU_004053 | Lv-Nlr13 |
| c3 | SPU_004342 | none |
| c3 | SPU_004479 | Lv-ALv-6 |
| c3 | SPU_004511 | Lv-Aqp9 |
| c3 | SPU_004644 | none |
| c3 | SPU_005046 | Lv-Hypp_1560 |
| c3 | SPU_005081 | Lv-AspL |
| c3 | SPU_005088 | Lv-Tlr025 |
| c3 | SPU_005270 | Lv-Ccp/Zp |
| c3 | SPU_005330 | Lv-Slco2blL |
| c3 | SPU_005331 | Lv-Slco5a1_1 |
| c3 | SPU_005383 | Lv-Nlr140 |
| c3 | SPU_005461 | NA |
| c3 | SPU_005530 | Lv-Lphn2L |
| c3 | SPU_005619 | Lv-Unc44L_44 |
| c3 | SPU_005642 | Lv-LrpL_2 |
| c3 | SPU_005859 | none |
| c3 | SPU_005921 | Lv-Hypp_1615 |
| c3 | SPU_006045 | Lv-Srcr46 |
| c3 | SPU_006273 | Lv-Hypp_1649 |
| c3 | SPU_006277 | Lv-GagpolL_1 |
| c3 | SPU_006279 | Lv-Srp9 |
| c3 | SPU_006341 | none |
| c3 | SPU_006512 | Lv-Hypp_1666 |
| c3 | SPU_006703 | Lv-PolypL_935 |
| c3 | SPU_006715 | none |
| c3 | SPU_006737 | Lv-Opsin5 |
| c3 | SPU_006799 | Lv-Hypp_1684 |
| c3 | SPU_006908 | Lv-Stk22A |
| c3 | SPU_006978 | Lv-Pdcd11L |
| c3 | SPU_007090 | Lv-Galr2L_1 |
| c3 | SPU_007122 | Lv-Bhmt_3 |
| c3 | SPU_007444 | Lv-Mtnr1b |
| c3 | SPU_007792 | Lv-Mfag4L |
| c3 | SPU_007852 | Lv-Runt2 |
| c3 | SPU_007889 | Lv-Hypp_529 |

| | | |
|---|---|---|
| c3 | SPU_007989 | Lv-Hypp_1757 |
| c3 | SPU_007993 | none |
| c3 | SPU_008057 | Lv-Kctd4 |
| c3 | SPU_008278 | Lv-Tlr045 |
| c3 | SPU_008363 | Lv-Prg4 |
| c3 | SPU_008392 | Lv-Hypp_1783 |
| c3 | SPU_008885 | NA |
| c3 | SPU_008947 | Lv-Htr4L_4 |
| c3 | SPU_008957 | Lv-DeltaL |
| c3 | SPU_008973 | Lv-Hrh3L_7 |
| c3 | SPU_009047 | none |
| c3 | SPU_009176 | NA |
| c3 | SPU_009179 | Lv-Rxfp2_1 |
| c3 | SPU_009187 | Lv-Hypp_1831 |
| c3 | SPU_009401 | Lv-C1qc |
| c3 | SPU_009580 | Lv-Surreal/Gpcr30 |
| c3 | SPU_009594 | Lv-Spon2Lf |
| c3 | SPU_009659 | Lv-Nlr127 |
| c3 | SPU_009788 | Lv-Api_1 |
| c3 | SPU_010091 | Lv-Nlr176 |
| c3 | SPU_010144 | Lv-Fn3_26 |
| c3 | SPU_010264 | Lv-Hyr |
| c3 | SPU_010325 | none |
| c3 | SPU_010357 | Lv-Gpr52L_1 |
| c3 | SPU_010365 | none |
| c3 | SPU_010372 | none |
| c3 | SPU_010467 | none |
| c3 | SPU_010980 | Lv-Alox5_6 |
| c3 | SPU_011023 | none |
| c3 | SPU_011096 | Lv-Hypp_1958 |
| c3 | SPU_011164 | Lv-Hypp_642 |
| c3 | SPU_011219 | Lv-Klhl2L |
| c3 | SPU_011233 | none |
| c3 | SPU_011269 | none |
| c3 | SPU_011431 | none |
| c3 | SPU_011457 | Lv-Taut_1 |
| c3 | SPU_011607 | NA |
| c3 | SPU_011608 | Lv-Chrna6L_2 |

| | | |
|---|---|---|
| c3 | SPU_011630 | Lv-Hypp_1994 |
| c3 | SPU_011639 | none |
| c3 | SPU_011825 | none |
| c3 | SPU_011971 | Lv-Clect/Ldl_6a |
| c3 | SPU_011979 | Lv-Hypp_666 |
| c3 | SPU_012001 | Lv-Egf/Ty |
| c3 | SPU_012176 | Lv-RtL_31 |
| c3 | SPU_012207 | Lv-Sat2_6 |
| c3 | SPU_012266 | Lv-Hypp_2025 |
| c3 | SPU_012375 | Lv-CckbrL_2 |
| c3 | SPU_012752 | Lv-Lrp4L_7 |
| c3 | SPU_012978 | Lv-PlexA4_4 |
| c3 | SPU_013190 | Lv-Tie1L_2 |
| c3 | SPU_013270 | Lv-Adrb2L_3 |
| c3 | SPU_013289 | Lv-Hsp701F |
| c3 | SPU_013547 | none |
| c3 | SPU_013759 | Lv-Asp_15 |
| c3 | SPU_013789 | Lv-Gpr52L_6 |
| c3 | SPU_013837 | Lv-Lhh2ah |
| c3 | SPU_013897 | NA |
| c3 | SPU_013916 | none |
| c3 | SPU_013962 | Lv-Sim_1 |
| c3 | SPU_014192 | Lv-Echn33 |
| c3 | SPU_014278 | Lv-Elovl6_5 |
| c3 | SPU_014327 | Lv-Hypp_2140 |
| c3 | SPU_014349 | Lv-Hypp_2142 |
| c3 | SPU_014350 | Lv-Iap1 |
| c3 | SPU_014353 | Lv-Hypp_750 |
| c3 | SPU_014385 | Lv-Dhx40 |
| c3 | SPU_014519 | Lv-Scnn1a |
| c3 | SPU_014677 | Lv-MacpfC3 |
| c3 | SPU_014701 | NA |
| c3 | SPU_014703 | Lv-Veb4 |
| c3 | SPU_014848 | Lv-Hypp_2173 |
| c3 | SPU_014855 | Lv-Rhbdf1-2 |
| c3 | SPU_015015 | none |
| c3 | SPU_015073 | Lv-Galr2L_19 |
| c3 | SPU_015251 | NA |

| | | |
|---|---|---|
| c3 | SPU_015744 | none |
| c3 | SPU_015787 | Lv-Hypp_2230 |
| c3 | SPU_015791 | Lv-Hypp_2231 |
| c3 | SPU_015846 | Lv-Fn3/Igf_10 |
| c3 | SPU_015897 | Lv-Hypp_2237 |
| c3 | SPU_016160 | Lv-Hypp_2249 |
| c3 | SPU_016590 | Lv-Ift52 |
| c3 | SPU_016690 | Lv-Hypp_2267 |
| c3 | SPU_016732 | none |
| c3 | SPU_016759 | Lv-Hypp_841 |
| c3 | SPU_016787 | Lv-Adrrcpta1B |
| c3 | SPU_016939 | Lv-Htr1bL_4 |
| c3 | SPU_016974 | Lv-TlrP17 |
| c3 | SPU_017080 | Lv-Hypp_2303 |
| c3 | SPU_017489 | Lv-Hypp_2329 |
| c3 | SPU_017513 | Lv-Sfi1 |
| c3 | SPU_017537 | none |
| c3 | SPU_017740 | NA |
| c3 | SPU_017889 | Lv-Fn3/Igf_39 |
| c3 | SPU_017908 | Lv-Gpr101L_4 |
| c3 | SPU_017991 | Lv-Gfm2L |
| c3 | SPU_018090 | Lv-Wdr4_1 |
| c3 | SPU_018226 | none |
| c3 | SPU_018248 | Lv-Bmp9 |
| c3 | SPU_018339 | Lv-QrfprL_4 |
| c3 | SPU_018629 | none |
| c3 | SPU_018668 | Lv-Znfx1-11 |
| c3 | SPU_018743 | Lv-Rptp_mu/pfn3 |
| c3 | SPU_018928 | Lv-Tlr101 |
| c3 | SPU_019047 | Lv-Pipox_3 |
| c3 | SPU_019242 | none |
| c3 | SPU_019419 | none |
| c3 | SPU_019501 | none |
| c3 | SPU_019507 | none |
| c3 | SPU_019721 | Lv-EchnL_7 |
| c3 | SPU_020106 | Lv-Hypp_2487 |
| c3 | SPU_020353 | Lv-Hypp_2501 |
| c3 | SPU_020428 | Lv-Tlr204 |

| | | |
|---|---|---|
| c3 | SPU_020557 | Lv-Hypp_2512 |
| c3 | SPU_020626 | none |
| c3 | SPU_020660 | NA |
| c3 | SPU_020785 | none |
| c3 | SPU_020903 | none |
| c3 | SPU_020975 | Lv-Engrailed |
| c3 | SPU_021102 | none |
| c3 | SPU_021134 | Lv-EthrL_1 |
| c3 | SPU_021151 | Lv-Adra2aL_1 |
| c3 | SPU_021178 | Lv-TlrL_2 |
| c3 | SPU_021213 | none |
| c3 | SPU_021219 | Lv-Gpr54 |
| c3 | SPU_021260 | none |
| c3 | SPU_021309 | Lv-Hox8 |
| c3 | SPU_021361 | Lv-Hypp_2547 |
| c3 | SPU_021526 | Lv-Ams |
| c3 | SPU_021535 | Lv-Prss23_2 |
| c3 | SPU_021984 | Lv-Pdha1_1 |
| c3 | SPU_022411 | Lv-Ttrspn_17 |
| c3 | SPU_022442 | Lv-Hypp_1046 |
| c3 | SPU_022460 | none |
| c3 | SPU_022461 | none |
| c3 | SPU_022481 | Lv-Sstr5L_1 |
| c3 | SPU_022554 | Lv-Ac/Sc3 |
| c3 | SPU_022744 | none |
| c3 | SPU_022885 | Lv-Hypp_1060 |
| c3 | SPU_022920 | Lv-Pappa_1 |
| c3 | SPU_022934 | Lv-PlasmgnL1 |
| c3 | SPU_022949 | NA |
| c3 | SPU_022962 | Lv-Galr2L_28 |
| c3 | SPU_023026 | Lv-Vps29h |
| c3 | SPU_023082 | none |
| c3 | SPU_023105 | Lv-Hypp_1072 |
| c3 | SPU_023133 | Lv-Slc17a5_9 |
| c3 | SPU_023152 | Lv-Cradd1 |
| c3 | SPU_023154 | none |
| c3 | SPU_023165 | Lv-Fn3/Egff_22 |
| c3 | SPU_023239 | none |

| | | |
|---|---|---|
| c3 | SPU_023251 | Lv-Hypp_2661 |
| c3 | SPU_023411 | Lv-Drd2L_23 |
| c3 | SPU_023479 | Lv-Hypp_2672 |
| c3 | SPU_023719 | Lv-Hypp_2684 |
| c3 | SPU_023749 | Lv-Unc44_166 |
| c3 | SPU_023793 | Lv-Hypp_2686 |
| c3 | SPU_023890 | Lv-Cryz_3 |
| c3 | SPU_024116 | Lv-Hypp_1106 |
| c3 | SPU_024272 | Lv-Unc44_341 |
| c3 | SPU_024280 | Lv-Hypp_2722 |
| c3 | SPU_024300 | Lv-Sstr5L_2 |
| c3 | SPU_024454 | Lv-Zpdcp |
| c3 | SPU_024699 | Lv-Slc7a11_10 |
| c3 | SPU_024724 | Lv-Tesk2L_3 |
| c3 | SPU_024952 | none |
| c3 | SPU_025037 | Lv-Htr1L_6 |
| c3 | SPU_025120 | Lv-Tnxb |
| c3 | SPU_025133 | Lv-Hypp_1140 |
| c3 | SPU_025177 | Lv-Hypp_2776 |
| c3 | SPU_025219 | Lv-Mekk5 |
| c3 | SPU_025383 | Lv-PppL_141 |
| c3 | SPU_025400 | none |
| c3 | SPU_025439 | Lv-Rdh8_3 |
| c3 | SPU_025680 | Lv-Nlr7 |
| c3 | SPU_025871 | Lv-Hypp_2806 |
| c3 | SPU_025911 | none |
| c3 | SPU_025914 | Lv-Nlr15 |
| c3 | SPU_025964 | Lv-Ttc22 |
| c3 | SPU_026512 | Lv-Cep63L |
| c3 | SPU_026533 | Lv-Ia1/Ic140s |
| c3 | SPU_026738 | Lv-Hypp_1186 |
| c3 | SPU_026808 | Lv-Endrvt115 |
| c3 | SPU_026926 | none |
| c3 | SPU_026927 | Lv-Hypp_1192 |
| c3 | SPU_027056 | none |
| c3 | SPU_027175 | Lv-PolypL_86 |
| c3 | SPU_027241 | Lv-Hypp_2897 |
| c3 | SPU_027348 | Lv-CelL |

| | | |
|---|---|---|
| c3 | SPU_027459 | Lv-EctdL |
| c3 | SPU_027475 | none |
| c3 | SPU_027724 | NA |
| c3 | SPU_027787 | Lv-Hypp_1214 |
| c3 | SPU_027815 | Lv-Tlr152 |
| c3 | SPU_027861 | Lv-Mamdc2_3 |
| c3 | SPU_027884 | Lv-Hrh3L_20 |
| c3 | SPU_027908 | Lv-Galr2L_35 |
| c3 | SPU_027942 | none |
| c3 | SPU_027944 | Lv-Adra1aL_12 |
| c3 | SPU_027948 | Lv-PlxnaL_2 |
| c3 | SPU_028013 | Lv-Ddc_3 |
| c3 | SPU_028101 | Lv-St6galnac5_4 |
| c3 | SPU_028160 | none |
| c3 | SPU_028212 | Lv-ZanL_1 |
| c3 | SPU_028244 | Lv-Harbi1_4 |
| c3 | SPU_028276 | Lv-Cub/Zp_1 |
| c3 | SPU_028295 | Lv-Hypp_1225 |
| c3 | SPU_028453 | Lv-PppL_236 |
| c3 | SPU_028498 | none |
| c3 | SPU_028595 | Lv-Nlr115 |
| c3 | SPU_028859 | Lv-Unk_66 |
| c3 | SPU_028933 | none |
| c3 | SPU_030185 | Lv-Il17-10 |
| c3 | SPU_030200 | Lv-Il17-22 |
| c4 | SPU_000179 | Lv-Fut8 |
| c4 | SPU_000359 | Lv-Gnmt_1 |
| c4 | SPU_000519 | Lv-Bche-5 |
| c4 | SPU_000524 | none |
| c4 | SPU_001143 | Lv-Ank3 |
| c4 | SPU_001151 | NA |
| c4 | SPU_001163 | Lv-Unc44L_8 |
| c4 | SPU_001176 | Lv-Znf862-2 |
| c4 | SPU_001223 | Lv-PppL_737 |
| c4 | SPU_001440 | none |
| c4 | SPU_001659 | Lv-Ugt_3 |
| c4 | SPU_001877 | Lv-Tlr010 |
| c4 | SPU_001903 | Lv-LacL_1 |

| | | |
|---|---|---|
| c4 | SPU_001919 | none |
| c4 | SPU_002031 | Lv-PapL2 |
| c4 | SPU_002243 | Lv-Tll2L_29 |
| c4 | SPU_002462 | none |
| c4 | SPU_002541 | none |
| c4 | SPU_002704 | none |
| c4 | SPU_003163 | Lv-Hypp_7375 |
| c4 | SPU_003369 | Lv-Nfxl1l |
| c4 | SPU_003675 | Lv-Hypp_393 |
| c4 | SPU_003722 | none |
| c4 | SPU_003920 | Lv-Barx_1 |
| c4 | SPU_004258 | none |
| c4 | SPU_004384 | none |
| c4 | SPU_004491 | Lv-Hypp_1530 |
| c4 | SPU_004768 | Lv-Hypp_1542 |
| c4 | SPU_005098 | Lv-Hypp_1564 |
| c4 | SPU_005444 | NA |
| c4 | SPU_005495 | none |
| c4 | SPU_005741 | Lv-Kif2CL |
| c4 | SPU_005820 | NA |
| c4 | SPU_005877 | none |
| c4 | SPU_005945 | none |
| c4 | SPU_006088 | Lv-PppL_185 |
| c4 | SPU_006089 | none |
| c4 | SPU_006356 | Lv-Sstr2L |
| c4 | SPU_006450 | Lv-Hrh3L_1 |
| c4 | SPU_006639 | NA |
| c4 | SPU_006927 | Lv-Orxn/np/qrfpLr3 |
| c4 | SPU_007585 | none |
| c4 | SPU_007740 | none |
| c4 | SPU_007967 | none |
| c4 | SPU_008100 | Lv-Mrc2L_1 |
| c4 | SPU_008167 | Lv-Hypp_1770 |
| c4 | SPU_008431 | Lv-Nlr61 |
| c4 | SPU_008449 | Lv-Copb1_1 |
| c4 | SPU_008659 | none |
| c4 | SPU_008837 | Lv-Hypp_1809 |
| c4 | SPU_008873 | Lv-Pi16L |

| | | |
|---|---|---|
| c4 | SPU_009030 | none |
| c4 | SPU_009081 | Lv-Hypp_41 |
| c4 | SPU_009157 | Lv-Hypp_1828 |
| c4 | SPU_009238 | none |
| c4 | SPU_009284 | NA |
| c4 | SPU_009496 | Lv-Srcr76 |
| c4 | SPU_009573 | Lv-PolppL_83 |
| c4 | SPU_009673 | Lv-Mfnl1 |
| c4 | SPU_009754 | NA |
| c4 | SPU_009864 | none |
| c4 | SPU_010194 | Lv-Hypp_1902 |
| c4 | SPU_010262 | none |
| c4 | SPU_010389 | none |
| c4 | SPU_011008 | Lv-Trpa1L_2 |
| c4 | SPU_011478 | Lv-Chrna9L_1 |
| c4 | SPU_011525 | Lv-Mamdc4-6 |
| c4 | SPU_011670 | Lv-Tmprss9L_1 |
| c4 | SPU_011682 | Lv-Dynlc2-4a |
| c4 | SPU_011845 | Lv-Ona13 |
| c4 | SPU_011875 | Lv-Ap1cfi2 |
| c4 | SPU_011989 | none |
| c4 | SPU_012101 | Lv-Hypp_668 |
| c4 | SPU_012295 | Lv-Apn7_1 |
| c4 | SPU_012635 | Lv-Hypp_689 |
| c4 | SPU_012795 | Lv-Hypp_2050 |
| c4 | SPU_012954 | Lv-Hypp_698 |
| c4 | SPU_012982 | none |
| c4 | SPU_013153 | Lv-Hrh3_5 |
| c4 | SPU_013165 | none |
| c4 | SPU_013550 | none |
| c4 | SPU_013591 | Lv-Chst10 |
| c4 | SPU_013827 | Lv-Hypp_730 |
| c4 | SPU_014122 | NA |
| c4 | SPU_014124 | NA |
| c4 | SPU_014212 | Lv-Nmur2L_1 |
| c4 | SPU_014461 | Lv-Pitx1 |
| c4 | SPU_014570 | none |
| c4 | SPU_014595 | Lv-St8sia2_2 |

| | | |
|---|---|---|
| c4 | SPU_014652 | Lv-SacsL_12 |
| c4 | SPU_014858 | Lv-Tie1L_1 |
| c4 | SPU_014894 | Lv-Oplah_1 |
| c4 | SPU_014980 | none |
| c4 | SPU_015047 | Lv-Endrvt90 |
| c4 | SPU_015183 | none |
| c4 | SPU_015498 | Lv-Unc44L_110 |
| c4 | SPU_015502 | none |
| c4 | SPU_015808 | Lv-Hypp_116 |
| c4 | SPU_015844 | Lv-Fn3/Igf_9 |
| c4 | SPU_016043 | Lv-Ttrspn_14 |
| c4 | SPU_016071 | NA |
| c4 | SPU_016187 | Lv-Znfx1_10 |
| c4 | SPU_016209 | none |
| c4 | SPU_016210 | Lv-Hypp_818 |
| c4 | SPU_016306 | Lv-C9orf102L |
| c4 | SPU_016584 | Lv-HyalinL_1 |
| c4 | SPU_016615 | none |
| c4 | SPU_017060 | Lv-Hypp_2301 |
| c4 | SPU_017178 | Lv-TrimL |
| c4 | SPU_017223 | Lv-Hypp_867 |
| c4 | SPU_017238 | Lv-Cavl |
| c4 | SPU_017392 | Lv-PppL_208 |
| c4 | SPU_017400 | Lv-Unc44_238 |
| c4 | SPU_017741 | Lv-Rasgef1b |
| c4 | SPU_017873 | NA |
| c4 | SPU_018223 | Lv-Hypp_911 |
| c4 | SPU_018228 | Lv-R/Ptp/10D |
| c4 | SPU_018501 | Lv-Clect_83 |
| c4 | SPU_018550 | Lv-Gal/lec14 |
| c4 | SPU_018836 | Lv-Rprd1b_1 |
| c4 | SPU_018867 | NA |
| c4 | SPU_019335 | Lv-Hypp_2432 |
| c4 | SPU_019351 | Lv-Il17-6 |
| c4 | SPU_019547 | Lv-Hypp_2447 |
| c4 | SPU_019874 | NA |
| c4 | SPU_019935 | Lv-Hypp_2473 |
| c4 | SPU_019978 | Lv-Hypp_955 |

| | | |
|---|---|---|
| c4 | SPU_020139 | Lv-Cola2L_2 |
| c4 | SPU_020510 | none |
| c4 | SPU_021119 | Lv-MyoD |
| c4 | SPU_021318 | Lv-Unc44_154 |
| c4 | SPU_021356 | NA |
| c4 | SPU_021408 | NA |
| c4 | SPU_021420 | NA |
| c4 | SPU_021708 | none |
| c4 | SPU_021786 | none |
| c4 | SPU_021943 | Lv-Mon2-2 |
| c4 | SPU_021980 | Lv-Hypp_1034 |
| c4 | SPU_022043 | none |
| c4 | SPU_022443 | Lv-Mamdc4_8 |
| c4 | SPU_022611 | Lv-Unk_65 |
| c4 | SPU_022859 | none |
| c4 | SPU_022886 | Lv-Echn39 |
| c4 | SPU_022913 | Lv-Hypp_1061 |
| c4 | SPU_022915 | none |
| c4 | SPU_022967 | none |
| c4 | SPU_023087 | Lv-Unc44_164 |
| c4 | SPU_023550 | Lv-Nlr67 |
| c4 | SPU_023551 | none |
| c4 | SPU_023871 | Lv-Hypp_2691 |
| c4 | SPU_024000 | Lv-Hypp_1102 |
| c4 | SPU_024088 | none |
| c4 | SPU_024225 | Lv-Nme5_1 |
| c4 | SPU_024230 | Lv-Chst11_3 |
| c4 | SPU_024232 | Lv-Slc5a11 |
| c4 | SPU_024385 | Lv-Tlr127 |
| c4 | SPU_024512 | Lv-Hypp_2739 |
| c4 | SPU_024571 | none |
| c4 | SPU_024915 | none |
| c4 | SPU_025102 | Lv-Vdp_1 |
| c4 | SPU_025167 | Lv-Nlr156 |
| c4 | SPU_025316 | Lv-Clect_116 |
| c4 | SPU_025455 | none |
| c4 | SPU_025571 | Lv-Klhl10 |
| c4 | SPU_025769 | Lv-Unc44_176 |

| | | |
|---|---|---|
| c4 | SPU_025965 | Lv-Ttc22L |
| c4 | SPU_026161 | NA |
| c4 | SPU_026200 | Lv-Tlr142 |
| c4 | SPU_026234 | Lv-Srcr211 |
| c4 | SPU_026296 | Lv-TlrL_17 |
| c4 | SPU_026452 | Lv-Slc10a2_12 |
| c4 | SPU_026520 | none |
| c4 | SPU_026524 | NA |
| c4 | SPU_026563 | none |
| c4 | SPU_027030 | none |
| c4 | SPU_027138 | none |
| c4 | SPU_027319 | NA |
| c4 | SPU_027449 | NA |
| c4 | SPU_027563 | Lv-Aldoaa_1 |
| c4 | SPU_027577 | Lv-Hypp_2911 |
| c4 | SPU_027590 | Lv-Hypp_2914 |
| c4 | SPU_027722 | Lv-Hypp_2916 |
| c4 | SPU_027753 | Lv-Z347_1 |
| c4 | SPU_027776 | none |
| c4 | SPU_028035 | Lv-Sdr_2 |
| c4 | SPU_028372 | none |
| c4 | SPU_028400 | Lv-Sned1L_2 |
| c4 | SPU_028578 | Lv-Hypp_2959 |
| c4 | SPU_028597 | none |
| c4 | SPU_028723 | Lv-Ttrspn_24 |
| c4 | SPU_028793 | Lv-PolpL_89 |
| c4 | SPU_028825 | NA |
| c4 | SPU_030095 | Lv-Tlr183 |
| d1 | SPU_000001 | Lv-Sox |
| d1 | SPU_000076 | Lv-Gat_2 |
| d1 | SPU_000080 | NA |
| d1 | SPU_000090 | NA |
| d1 | SPU_000250 | Lv-Hypp_1246 |
| d1 | SPU_000276 | Lv-PaxC |
| d1 | SPU_000309 | Lv-AbcL |
| d1 | SPU_000372 | Lv-Hypp_1256 |
| d1 | SPU_000381 | Lv-FbsL_2 |
| d1 | SPU_000426 | Lv-Arsj |

| | | |
|---|---|---|
| d1 | SPU_000441 | Lv-Igcam/Igcam |
| d1 | SPU_000442 | Lv-Mlckb |
| d1 | SPU_000518 | Lv-Bche-2 |
| d1 | SPU_000521 | none |
| d1 | SPU_000594 | none |
| d1 | SPU_000616 | Lv-Pde10a-2 |
| d1 | SPU_000625 | Lv-March8L_2 |
| d1 | SPU_000682 | Lv-Hs3st3a1L |
| d1 | SPU_000725 | Lv-Mamdc4-3 |
| d1 | SPU_000772 | Lv-B3galt1_1 |
| d1 | SPU_000775 | Lv-Spg11L |
| d1 | SPU_000781 | Lv-B3galt5_1 |
| d1 | SPU_000836 | Lv-Th |
| d1 | SPU_000863 | Lv-Nlr152 |
| d1 | SPU_000930 | Lv-Hypp_1295 |
| d1 | SPU_000953 | none |
| d1 | SPU_000970 | Lv-Bbox1 |
| d1 | SPU_001045 | Lv-Btub2_1 |
| d1 | SPU_001076 | Lv-Chst11-4 |
| d1 | SPU_001109 | none |
| d1 | SPU_001110 | none |
| d1 | SPU_001139 | Lv-Kiaa1161 |
| d1 | SPU_001140 | Lv-Kiaa1161L_2 |
| d1 | SPU_001155 | none |
| d1 | SPU_001270 | NA |
| d1 | SPU_001383 | Lv-Unc44L_11 |
| d1 | SPU_001399 | Lv-Cpa1L |
| d1 | SPU_001447 | NA |
| d1 | SPU_001453 | Lv-Ugt2b4 |
| d1 | SPU_001456 | Lv-Mbd2L |
| d1 | SPU_001531 | Lv-Cpp |
| d1 | SPU_001532 | Lv-FibA |
| d1 | SPU_001563 | Lv-Slc20a1_1 |
| d1 | SPU_001639 | Lv-Lct-8 |
| d1 | SPU_001697 | NA |
| d1 | SPU_001899 | none |
| d1 | SPU_001912 | NA |
| d1 | SPU_002036 | NA |

| | | |
|---|---|---|
| d1 | SPU_002120 | none |
| d1 | SPU_002192 | none |
| d1 | SPU_002270 | Lv-Trpc6 |
| d1 | SPU_002280 | Lv-Cspe3/7L |
| d1 | SPU_002350 | Lv-Srcr23 |
| d1 | SPU_002382 | NA |
| d1 | SPU_002406 | none |
| d1 | SPU_002455 | Lv-Loxl2 |
| d1 | SPU_002481 | Lv-Slc5a9 |
| d1 | SPU_002514 | Lv-SmpdL |
| d1 | SPU_002587 | Lv-Ef1bg |
| d1 | SPU_002605 | Lv-AchE_5 |
| d1 | SPU_002606 | Lv-Rpl14 |
| d1 | SPU_002611 | none |
| d1 | SPU_002665 | Lv-Hypp_363 |
| d1 | SPU_002784 | none |
| d1 | SPU_002787 | Lv-Glur3 |
| d1 | SPU_002816 | Lv-Hb9 |
| d1 | SPU_002947 | NA |
| d1 | SPU_002967 | Lv-Oprk1L_4 |
| d1 | SPU_002970 | none |
| d1 | SPU_002971 | Lv-Ache |
| d1 | SPU_002978 | none |
| d1 | SPU_003053 | Lv-Hypp_1427 |
| d1 | SPU_003242 | Lv-DlcL1 |
| d1 | SPU_003277 | Lv-Vwd/Mucl |
| d1 | SPU_003293 | Lv-Fn3/Egff_8 |
| d1 | SPU_003298 | NA |
| d1 | SPU_003354 | Lv-Hypp_1446 |
| d1 | SPU_003437 | NA |
| d1 | SPU_003531 | Lv-Slc43a2 |
| d1 | SPU_003566 | NA |
| d1 | SPU_003606 | Lv-Cyp2L9 |
| d1 | SPU_003610 | Lv-Clect/Tm_8 |
| d1 | SPU_003711 | Lv-Gat_3 |
| d1 | SPU_003745 | Lv-Hs3st3b1L |
| d1 | SPU_003823 | Lv-Ugt2b6 |
| d1 | SPU_003842 | Lv-Mrlc2 |

| | | |
|---|---|---|
| d1 | SPU_003880 | Lv-Rt22 |
| d1 | SPU_003952 | Lv-Hypp_1489 |
| d1 | SPU_003972 | Lv-Kcnk16 |
| d1 | SPU_003991 | Lv-Susd2-3 |
| d1 | SPU_004033 | NA |
| d1 | SPU_004039 | none |
| d1 | SPU_004074 | NA |
| d1 | SPU_004096 | Lv-Rrp22_1 |
| d1 | SPU_004131 | Lv-Cox6b1 |
| d1 | SPU_004157 | Lv-Cpa3 |
| d1 | SPU_004174 | Lv-Wdr59-2 |
| d1 | SPU_004215 | Lv-Hypp_405 |
| d1 | SPU_004265 | none |
| d1 | SPU_004338 | NA |
| d1 | SPU_004345 | none |
| d1 | SPU_004347 | Lv-Fn3/Egf_5 |
| d1 | SPU_004431 | Lv-Pcs_2 |
| d1 | SPU_004503 | Lv-Slc7a6 |
| d1 | SPU_004599 | Lv-Pitx2 |
| d1 | SPU_004609 | Lv-Rpl31 |
| d1 | SPU_004626 | Lv-Pdpi(cyclophilin)L4 |
| d1 | SPU_004637 | Lv-Dhrs |
| d1 | SPU_004677 | Lv-Thred |
| d1 | SPU_004775 | Lv-Hypp_1544 |
| d1 | SPU_004845 | Lv-Egfrp |
| d1 | SPU_004885 | Lv-Apod |
| d1 | SPU_004961 | NA |
| d1 | SPU_005015 | Lv-Fn3_40 |
| d1 | SPU_005045 | Lv-Fbxl2 |
| d1 | SPU_005199 | Lv-Hypp_1571 |
| d1 | SPU_005250 | none |
| d1 | SPU_005334 | NA |
| d1 | SPU_005353 | none |
| d1 | SPU_005392 | Lv-Taar7eL_2 |
| d1 | SPU_005465 | Lv-Unc44L_43 |
| d1 | SPU_005500 | Lv-C1qL |
| d1 | SPU_005636 | Lv-Tmem20 |
| d1 | SPU_005669 | Lv-Cav1 |

| | | |
|---|---|---|
| d1 | SPU_005773 | none |
| d1 | SPU_005819 | Lv-GhdcL_6 |
| d1 | SPU_005833 | Lv-Slco3a1_1 |
| d1 | SPU_005893 | Lv-Hypp_10 |
| d1 | SPU_005894 | Lv-Hypp_11 |
| d1 | SPU_005912 | none |
| d1 | SPU_005917 | Lv-C10orf112L |
| d1 | SPU_005989 | Lv-Clect_13 |
| d1 | SPU_006002 | Lv-Slc16a14_1 |
| d1 | SPU_006003 | none |
| d1 | SPU_006033 | Lv-Sirt3L |
| d1 | SPU_006051 | Lv-Unc44L_50 |
| d1 | SPU_006123 | Lv-AchE |
| d1 | SPU_006155 | NA |
| d1 | SPU_006161 | Lv-Vacht |
| d1 | SPU_006360 | Lv-Hrh4L_1 |
| d1 | SPU_006494 | none |
| d1 | SPU_006495 | none |
| d1 | SPU_006524 | Lv-Slc22a3_2 |
| d1 | SPU_006537 | Lv-Slc22a3_9 |
| d1 | SPU_006610 | Lv-Nlr4 |
| d1 | SPU_006628 | Lv-Astacin6 |
| d1 | SPU_006646 | Lv-Fam151bL |
| d1 | SPU_006649 | Lv-Clect_102 |
| d1 | SPU_006712 | Lv-Cyth1L |
| d1 | SPU_006800 | Lv-Ugt2a1 |
| d1 | SPU_006891 | none |
| d1 | SPU_006919 | Lv-Hrh1L |
| d1 | SPU_006929 | NA |
| d1 | SPU_006934 | Lv-CfhL |
| d1 | SPU_006942 | Lv-Hypp_502 |
| d1 | SPU_007010 | Lv-Hypp_1692 |
| d1 | SPU_007073 | none |
| d1 | SPU_007093 | none |
| d1 | SPU_007094 | Lv-Galr2L_7 |
| d1 | SPU_007186 | Lv-Mglur3_3 |
| d1 | SPU_007308 | Lv-Megf6 |
| d1 | SPU_007328 | none |

| | | |
|---|---|---|
| d1 | SPU_007404 | Lv-Cyp1L6 |
| d1 | SPU_007420 | NA |
| d1 | SPU_007553 | Lv-Go |
| d1 | SPU_007599 | Lv-Glass2 |
| d1 | SPU_007602 | NA |
| d1 | SPU_007603 | Lv-Slco |
| d1 | SPU_007796 | Lv-Surreal/Gpcr85 |
| d1 | SPU_007797 | Lv-Asp_8 |
| d1 | SPU_008164 | none |
| d1 | SPU_008332 | Lv-Traf4 |
| d1 | SPU_008356 | Lv-Cthrc1 |
| d1 | SPU_008566 | none |
| d1 | SPU_008631 | Lv-Lct_2 |
| d1 | SPU_008633 | Lv-Lct_3 |
| d1 | SPU_008769 | NA |
| d1 | SPU_008788 | Lv-Chst3L_1 |
| d1 | SPU_008817 | Lv-Disp1_1 |
| d1 | SPU_008862 | Lv-Timp1b |
| d1 | SPU_008935 | Lv-Unc44_215 |
| d1 | SPU_008968 | Lv-Hypp_1818 |
| d1 | SPU_009142 | Lv-Slc16a9_4 |
| d1 | SPU_009145 | Lv-Srcr74 |
| d1 | SPU_009191 | NA |
| d1 | SPU_009231 | Lv-Ncsl |
| d1 | SPU_009232 | none |
| d1 | SPU_009307 | none |
| d1 | SPU_009341 | Lv-Hypp_1843 |
| d1 | SPU_009365 | Lv-Lact2L |
| d1 | SPU_009432 | Lv-SpnprA |
| d1 | SPU_009466 | Lv-Ebr1L_6 |
| d1 | SPU_009482 | Lv-CskaIIa |
| d1 | SPU_009656 | none |
| d1 | SPU_009701 | NA |
| d1 | SPU_009740 | none |
| d1 | SPU_009816 | NA |
| d1 | SPU_009825 | Lv-Cyp2L20 |
| d1 | SPU_009854 | NA |
| d1 | SPU_009879 | none |

| | | |
|---|---|---|
| d1 | SPU_010006 | Lv-Ttn |
| d1 | SPU_010007 | Lv-Conn/Unc89L |
| d1 | SPU_010137 | Lv-Slc20a1_3 |
| d1 | SPU_010143 | Lv-Cyp2L21 |
| d1 | SPU_010167 | Lv-Thytrprhr |
| d1 | SPU_010179 | Lv-Tacr2L_1 |
| d1 | SPU_010206 | Lv-Tmprss5L |
| d1 | SPU_010409 | NA |
| d1 | SPU_010425 | Lv-Odadc3L1 |
| d1 | SPU_010431 | Lv-Adora2aL_1 |
| d1 | SPU_010518 | NA |
| d1 | SPU_010622 | NA |
| d1 | SPU_010751 | Lv-Hypp_1939 |
| d1 | SPU_010792 | NA |
| d1 | SPU_010831 | Lv-Glycos |
| d1 | SPU_010839 | none |
| d1 | SPU_010845 | Lv-Csk3 |
| d1 | SPU_010884 | Lv-Htr4L_7 |
| d1 | SPU_010934 | none |
| d1 | SPU_011050 | Lv-Slc16a12L_5 |
| d1 | SPU_011054 | none |
| d1 | SPU_011057 | none |
| d1 | SPU_011062 | NA |
| d1 | SPU_011103 | none |
| d1 | SPU_011109 | Lv-Hypp_1960 |
| d1 | SPU_011133 | Lv-Fut4_2 |
| d1 | SPU_011137 | Lv-Slc16a10L |
| d1 | SPU_011147 | NA |
| d1 | SPU_011155 | NA |
| d1 | SPU_011162 | NA |
| d1 | SPU_011280 | Lv-Sp8 |
| d1 | SPU_011320 | Lv-Drd1 |
| d1 | SPU_011331 | NA |
| d1 | SPU_011366 | Lv-Hypp_1975 |
| d1 | SPU_011383 | none |
| d1 | SPU_011389 | none |
| d1 | SPU_011497 | Lv-Dur3_1 |
| d1 | SPU_011541 | Lv-Tlrp40 |

| | | |
|---|---|---|
| d1 | SPU_011730 | none |
| d1 | SPU_011736 | Lv-Colf_3 |
| d1 | SPU_011737 | none |
| d1 | SPU_011759 | Lv-Lrr/Igr_12 |
| d1 | SPU_011768 | Lv-Slc37a3 |
| d1 | SPU_011795 | Lv-Hypp_2000 |
| d1 | SPU_011809 | Lv-Arrdc4 |
| d1 | SPU_011818 | NA |
| d1 | SPU_011892 | Lv-Glipr2L |
| d1 | SPU_011928 | Lv-Slc16a7 |
| d1 | SPU_011933 | Lv-Atp13a3 |
| d1 | SPU_011938 | NA |
| d1 | SPU_011978 | Lv-Enpep |
| d1 | SPU_012007 | Lv-Mfhas1 |
| d1 | SPU_012008 | Lv-Myor2 |
| d1 | SPU_012092 | NA |
| d1 | SPU_012095 | Lv-Aqp8 |
| d1 | SPU_012146 | none |
| d1 | SPU_012185 | Lv-Ache_21 |
| d1 | SPU_012187 | none |
| d1 | SPU_012247 | Lv-Tsp1_1 |
| d1 | SPU_012279 | Lv-Slc5a9_2 |
| d1 | SPU_012289 | Lv-Fut5 |
| d1 | SPU_012322 | Lv-Echn3 |
| d1 | SPU_012323 | Lv-Cntnap4L |
| d1 | SPU_012358 | Lv-Kcnk1 |
| d1 | SPU_012371 | none |
| d1 | SPU_012381 | none |
| d1 | SPU_012476 | Lv-Mmr |
| d1 | SPU_012502 | Lv-Hypp_2036 |
| d1 | SPU_012518 | Lv-Cara7LA |
| d1 | SPU_012541 | Lv-Syt7L_1 |
| d1 | SPU_012562 | Lv-Htr6L |
| d1 | SPU_012597 | NA |
| d1 | SPU_012683 | Lv-Zizimin2 |
| d1 | SPU_012692 | Lv-Hypp_79 |
| d1 | SPU_012699 | Lv-Nkx6-1 |
| d1 | SPU_012710 | NA |

| | | |
|---|---|---|
| d1 | SPU_012746 | Lv-Kcnk4 |
| d1 | SPU_012806 | Lv-Slc45a2 |
| d1 | SPU_012855 | Lv-Clect/Fn3_18 |
| d1 | SPU_012935 | none |
| d1 | SPU_013040 | Lv-Hypp_700 |
| d1 | SPU_013122 | Lv-Plxna2L_1 |
| d1 | SPU_013205 | none |
| d1 | SPU_013284 | Lv-Chst9_2 |
| d1 | SPU_013361 | Lv-Slc7a9_1 |
| d1 | SPU_013368 | Lv-Prdx6 |
| d1 | SPU_013410 | none |
| d1 | SPU_013429 | Lv-Elovl6_3 |
| d1 | SPU_013465 | Lv-Nlr89 |
| d1 | SPU_013498 | Lv-Gpr133_2 |
| d1 | SPU_013504 | NA |
| d1 | SPU_013616 | Lv-Hypp_724 |
| d1 | SPU_013621 | none |
| d1 | SPU_013626 | Lv-Clca_6 |
| d1 | SPU_013717 | none |
| d1 | SPU_013736 | none |
| d1 | SPU_013825 | Lv-Clect_76 |
| d1 | SPU_013872 | Lv-Clect_106 |
| d1 | SPU_013884 | NA |
| d1 | SPU_013923 | none |
| d1 | SPU_013960 | Lv-Allgif1 |
| d1 | SPU_014010 | Lv-Chst3L_2 |
| d1 | SPU_014024 | Lv-Slco4a1_5 |
| d1 | SPU_014025 | Lv-Slco2a1_1 |
| d1 | SPU_014043 | Lv-Slc10a2_14 |
| d1 | SPU_014084 | Lv-Susd2 |
| d1 | SPU_014109 | none |
| d1 | SPU_014136 | Lv-Slc10a2_8 |
| d1 | SPU_014170 | Lv-SoxF |
| d1 | SPU_014274 | NA |
| d1 | SPU_014340 | Lv-CfhL-1 |
| d1 | SPU_014379 | Lv-Hypp_2144 |
| d1 | SPU_014395 | Lv-DisLA2 |
| d1 | SPU_014487 | Lv-Hypp_2153 |

| | | |
|---|---|---|
| d1 | SPU_014488 | NA |
| d1 | SPU_014496 | Lv-Msp130r4 |
| d1 | SPU_014556 | Lv-Hypp_2157 |
| d1 | SPU_014606 | Lv-CadherinL6 |
| d1 | SPU_014614 | Lv-Krn1 |
| d1 | SPU_014664 | Lv-Glur6_1 |
| d1 | SPU_014680 | none |
| d1 | SPU_014689 | Lv-Cndrep |
| d1 | SPU_014761 | Lv-Nlr111 |
| d1 | SPU_014779 | Lv-Unc44L_101 |
| d1 | SPU_014827 | Lv-Hypp_765 |
| d1 | SPU_014844 | Lv-Srcr117 |
| d1 | SPU_014877 | Lv-Chrna9_3 |
| d1 | SPU_015027 | Lv-Acel_2 |
| d1 | SPU_015161 | NA |
| d1 | SPU_015218 | Lv-Hypp_2199 |
| d1 | SPU_015227 | none |
| d1 | SPU_015228 | NA |
| d1 | SPU_015286 | Lv-Hypp_111 |
| d1 | SPU_015344 | none |
| d1 | SPU_015485 | Lv-Gluld1_7 |
| d1 | SPU_015655 | none |
| d1 | SPU_015681 | NA |
| d1 | SPU_015719 | Lv-FoxL1 |
| d1 | SPU_015795 | Lv-Slc10a2_5 |
| d1 | SPU_015837 | NA |
| d1 | SPU_015866 | none |
| d1 | SPU_015875 | Lv-Unc44L_112 |
| d1 | SPU_015922 | none |
| d1 | SPU_015938 | Lv-Srcr129 |
| d1 | SPU_016020 | Lv-Enpep_2 |
| d1 | SPU_016033 | Lv-Lrr/Gpcr_2 |
| d1 | SPU_016037 | Lv-Hypp_808 |
| d1 | SPU_016184 | Lv-Hypp_2250 |
| d1 | SPU_016196 | none |
| d1 | SPU_016260 | Lv-Nlgn4yL |
| d1 | SPU_016283 | Lv-Nos_2 |
| d1 | SPU_016349 | none |

| | | |
|---|---|---|
| d1 | SPU_016379 | Lv-Rej9 |
| d1 | SPU_016399 | NA |
| d1 | SPU_016402 | NA |
| d1 | SPU_016432 | Lv-Alpl_1 |
| d1 | SPU_016482 | Lv-C3orf16L |
| d1 | SPU_016519 | none |
| d1 | SPU_016583 | Lv-PolypL_75 |
| d1 | SPU_016595 | Lv-Soat_1 |
| d1 | SPU_016612 | Lv-Scdfr2L_2 |
| d1 | SPU_016629 | Lv-Clect/Egfl/Tm_5 |
| d1 | SPU_016685 | Lv-Ap2 |
| d1 | SPU_016778 | Lv-Unc44_315 |
| d1 | SPU_016786 | Lv-Rhox3 |
| d1 | SPU_016792 | Lv-Tob |
| d1 | SPU_016820 | none |
| d1 | SPU_016823 | Lv-Rbp1_1 |
| d1 | SPU_016988 | NA |
| d1 | SPU_017054 | Lv-Nlr17 |
| d1 | SPU_017176 | Lv-Hypp_2307 |
| d1 | SPU_017194 | Lv-Srcr138 |
| d1 | SPU_017234 | Lv-PtprA |
| d1 | SPU_017267 | Lv-Muc19L |
| d1 | SPU_017304 | none |
| d1 | SPU_017305 | none |
| d1 | SPU_017340 | none |
| d1 | SPU_017341 | NA |
| d1 | SPU_017345 | NA |
| d1 | SPU_017361 | Lv-Vwd/Egf |
| d1 | SPU_017373 | Lv-Gst_pi |
| d1 | SPU_017375 | Lv-Nr2e6 |
| d1 | SPU_017460 | NA |
| d1 | SPU_017468 | Lv-Sat2L |
| d1 | SPU_017516 | Lv-Abcc10_1 |
| d1 | SPU_017622 | NA |
| d1 | SPU_017623 | none |
| d1 | SPU_017624 | Lv-Prokr1 |
| d1 | SPU_017659 | none |
| d1 | SPU_017661 | Lv-Mgam |

| | | |
|---|---|---|
| d1 | SPU_017662 | Lv-Mgam_2 |
| d1 | SPU_017673 | Lv-Dom3zL_1 |
| d1 | SPU_017793 | Lv-Slc39a12L |
| d1 | SPU_017814 | Lv-Odadc3L3 |
| d1 | SPU_017875 | Lv-Efcab8 |
| d1 | SPU_017918 | Lv-5ht1F |
| d1 | SPU_018088 | none |
| d1 | SPU_018091 | Lv-Htrla |
| d1 | SPU_018294 | Lv-Lrr/Gpcr_7 |
| d1 | SPU_018402 | Lv-Hypp_2385 |
| d1 | SPU_018474 | none |
| d1 | SPU_018541 | Lv-Gpr1L_8 |
| d1 | SPU_018709 | NA |
| d1 | SPU_018735 | Lv-Itih_1 |
| d1 | SPU_018738 | Lv-Slc16aL_1 |
| d1 | SPU_018768 | Lv-ApdpL_1 |
| d1 | SPU_018827 | NA |
| d1 | SPU_018912 | none |
| d1 | SPU_018919 | Lv-Hypp_2410 |
| d1 | SPU_018945 | Lv-Hypp_2412 |
| d1 | SPU_018974 | Lv-C4mso |
| d1 | SPU_019143 | none |
| d1 | SPU_019170 | none |
| d1 | SPU_019201 | Lv-Hypp_931-2 |
| d1 | SPU_019212 | Lv-Slc7a4_5 |
| d1 | SPU_019220 | Lv-Unc44_321 |
| d1 | SPU_019290 | Lv-Otp |
| d1 | SPU_019345 | Lv-FoxQ1 |
| d1 | SPU_019489 | Lv-P2rx4 |
| d1 | SPU_019549 | Lv-TrypsinL_4 |
| d1 | SPU_019624 | Lv-Smpdl3b |
| d1 | SPU_019709 | Lv-Nacha2_2 |
| d1 | SPU_019710 | Lv-Nacha4 |
| d1 | SPU_019774 | Lv-Osta |
| d1 | SPU_019877 | Lv-Dnajc7L |
| d1 | SPU_019970 | Lv-Nos1 |
| d1 | SPU_019979 | Lv-Slc24a6_1 |
| d1 | SPU_019981 | Lv-Hypp_956 |

| | | |
|---|---|---|
| d1 | SPU_019990 | NA |
| d1 | SPU_020032 | none |
| d1 | SPU_020033 | none |
| d1 | SPU_020041 | NA |
| d1 | SPU_020052 | Lv-UbcL |
| d1 | SPU_020114 | Lv-Hypp_2489 |
| d1 | SPU_020142 | Lv-Mtnr1b_1 |
| d1 | SPU_020234 | Lv-Cyp2L42 |
| d1 | SPU_020235 | none |
| d1 | SPU_020330 | Lv-Grm |
| d1 | SPU_020343 | Lv-Hypp_2500 |
| d1 | SPU_020346 | Lv-Tbx6 |
| d1 | SPU_020376 | Lv-PrpLr |
| d1 | SPU_020432 | NA |
| d1 | SPU_020457 | Lv-B7L3 |
| d1 | SPU_020467 | Lv-Slc5a6_2 |
| d1 | SPU_020474 | none |
| d1 | SPU_020519 | NA |
| d1 | SPU_020686 | NA |
| d1 | SPU_020759 | Lv-Kcnk3_1 |
| d1 | SPU_020970 | Lv-Siat6_1 |
| d1 | SPU_020979 | Lv-Gpr158 |
| d1 | SPU_021031 | Lv-Hmcn1L_1 |
| d1 | SPU_021156 | Lv-Arrdc3_3 |
| d1 | SPU_021157 | Lv-Chrna9_4 |
| d1 | SPU_021158 | Lv-Nacha10 |
| d1 | SPU_021159 | Lv-Nacha6 |
| d1 | SPU_021185 | Lv-Cyp17r3 |
| d1 | SPU_021241 | none |
| d1 | SPU_021355 | Lv-Ace_1 |
| d1 | SPU_021372 | none |
| d1 | SPU_021495 | Lv-FibD |
| d1 | SPU_021500 | Lv-Ugt2b11 |
| d1 | SPU_021584 | Lv-Syt9L_1 |
| d1 | SPU_021675 | Lv-Gal/lec10 |
| d1 | SPU_021698 | none |
| d1 | SPU_021873 | Lv-Rdh8_8 |
| d1 | SPU_021876 | Lv-Muc5b |

| | | |
|---|---|---|
| d1 | SPU_021897 | none |
| d1 | SPU_021899 | none |
| d1 | SPU_022021 | Lv-Igf_12 |
| d1 | SPU_022096 | Lv-Hypp_1037 |
| d1 | SPU_022121 | none |
| d1 | SPU_022125 | none |
| d1 | SPU_022217 | Lv-Hypp_2605 |
| d1 | SPU_022229 | Lv-RtL_56 |
| d1 | SPU_022312 | Lv-Sult1b1_1 |
| d1 | SPU_022318 | Lv-MacpfD.2 |
| d1 | SPU_022319 | none |
| d1 | SPU_022382 | Lv-Sert |
| d1 | SPU_022471 | Lv-Hypp_1047 |
| d1 | SPU_022506 | Lv-Slc6a2 |
| d1 | SPU_022542 | Lv-Hypp_2614 |
| d1 | SPU_022586 | Lv-Fn3/Egf/Igf |
| d1 | SPU_022592 | none |
| d1 | SPU_022651 | Lv-Agpat1_1 |
| d1 | SPU_022730 | Lv-Gpr54L_2 |
| d1 | SPU_022753 | Lv-Sea/Ccp_1 |
| d1 | SPU_022763 | Lv-AchE_7 |
| d1 | SPU_022846 | Lv-FoxABL |
| d1 | SPU_022882 | Lv-Colf_7 |
| d1 | SPU_022943 | NA |
| d1 | SPU_023069 | Lv-TrafB |
| d1 | SPU_023072 | Lv-Unc44_163 |
| d1 | SPU_023077 | Lv-Hypp_1069 |
| d1 | SPU_023078 | Lv-Hypp_2648 |
| d1 | SPU_023097 | NA |
| d1 | SPU_023130 | NA |
| d1 | SPU_023170 | Lv-Hypp_2652 |
| d1 | SPU_023223 | Lv-Orxn/HpcLr2 |
| d1 | SPU_023291 | none |
| d1 | SPU_023292 | Lv-Hypp_2665 |
| d1 | SPU_023467 | Lv-Nmral1_2 |
| d1 | SPU_023522 | Lv-Z66 |
| d1 | SPU_023543 | Lv-Fstl5 |
| d1 | SPU_023584 | Lv-Apob |

| | | |
|---|---|---|
| d1 | SPU_023642 | Lv-Nlr153 |
| d1 | SPU_023711 | NA |
| d1 | SPU_023713 | Lv-Unc44_340 |
| d1 | SPU_023734 | Lv-Glra1 |
| d1 | SPU_023850 | none |
| d1 | SPU_023874 | Lv-Hypp_2692 |
| d1 | SPU_023894 | Lv-FoxI |
| d1 | SPU_023902 | Lv-Hypp_2695 |
| d1 | SPU_023944 | none |
| d1 | SPU_023984 | Lv-Naalad2L_13 |
| d1 | SPU_024003 | Lv-Mmp1/3L2 |
| d1 | SPU_024007 | none |
| d1 | SPU_024057 | Lv-Hypp_2706 |
| d1 | SPU_024167 | Lv-Pla2g12 |
| d1 | SPU_024176 | Lv-Hypp_211 |
| d1 | SPU_024188 | Lv-Anxa7_1 |
| d1 | SPU_024193 | none |
| d1 | SPU_024195 | NA |
| d1 | SPU_024200 | Lv-Scnng |
| d1 | SPU_024249 | Lv-Chat |
| d1 | SPU_024320 | Lv-Cyp2L57 |
| d1 | SPU_024558 | Lv-Hypp_218 |
| d1 | SPU_024602 | none |
| d1 | SPU_024747 | Lv-Mchr1L |
| d1 | SPU_024972 | NA |
| d1 | SPU_024978 | Lv-Glur2_1 |
| d1 | SPU_024993 | Lv-Cav3 |
| d1 | SPU_025055 | Lv-Slc3a1_1 |
| d1 | SPU_025092 | Lv-Lct_1 |
| d1 | SPU_025146 | Lv-Slc46a1L_8 |
| d1 | SPU_025158 | none |
| d1 | SPU_025203 | Lv-Hypp_2778 |
| d1 | SPU_025228 | Lv-Nt5e_3 |
| d1 | SPU_025244 | Lv-Mrlc |
| d1 | SPU_025266 | Lv-GabrA |
| d1 | SPU_025323 | none |
| d1 | SPU_025392 | Lv-Slc27a2-5 |
| d1 | SPU_025396 | none |

| | | |
|---|---|---|
| d1 | SPU_025407 | NA |
| d1 | SPU_025477 | NA |
| d1 | SPU_025632 | Lv-Pou4f2 |
| d1 | SPU_025728 | Lv-Scnn1a_1 |
| d1 | SPU_025729 | none |
| d1 | SPU_025916 | Lv-Ei24_1 |
| d1 | SPU_025935 | Lv-Slc34a2_2 |
| d1 | SPU_025947 | Lv-Slc32a1 |
| d1 | SPU_026008 | Lv-FcolI/II/IIIf |
| d1 | SPU_026009 | Lv-A2col |
| d1 | SPU_026021 | Lv-Slc43a2L |
| d1 | SPU_026056 | Lv-Rspo3 |
| d1 | SPU_026127 | Lv-Ugt2b1_2 |
| d1 | SPU_026203 | Lv-Rpesp |
| d1 | SPU_026221 | Lv-SLc5a9 |
| d1 | SPU_026241 | Lv-Srcr212 |
| d1 | SPU_026414 | Lv-OprL |
| d1 | SPU_026451 | Lv-Hypp_2844 |
| d1 | SPU_026466 | Lv-Abcc5g |
| d1 | SPU_026607 | Lv-Mamdc4-14 |
| d1 | SPU_026694 | Lv-Slc7a9_2 |
| d1 | SPU_026696 | Lv-Slc7_2 |
| d1 | SPU_026751 | Lv-Acel_6 |
| d1 | SPU_026873 | NA |
| d1 | SPU_026910 | Lv-Acel |
| d1 | SPU_026973 | none |
| d1 | SPU_026989 | none |
| d1 | SPU_027076 | none |
| d1 | SPU_027091 | Lv-Ghdc_1 |
| d1 | SPU_027098 | none |
| d1 | SPU_027121 | Lv-Vwd/Egf/Fn3 |
| d1 | SPU_027139 | Lv-Hypp_2890 |
| d1 | SPU_027170 | Lv-Hypp_2891 |
| d1 | SPU_027295 | Lv-Chdh-4 |
| d1 | SPU_027308 | Lv-St3gal |
| d1 | SPU_027422 | Lv-Slc16a4L_6 |
| d1 | SPU_027500 | NA |
| d1 | SPU_027513 | Lv-Nlr47 |

| | | |
|---|---|---|
| d1 | SPU_027537 | none |
| d1 | SPU_027572 | none |
| d1 | SPU_027648 | Lv-FoxD_1 |
| d1 | SPU_027729 | Lv-Sea/Egf_1 |
| d1 | SPU_027732 | Lv-Hypp_2917 |
| d1 | SPU_027747 | Lv-Ttrspn_22 |
| d1 | SPU_027794 | none |
| d1 | SPU_027808 | Lv-Nlr154 |
| d1 | SPU_027831 | Lv-BcatL_1 |
| d1 | SPU_027870 | Lv-Acsl5_5 |
| d1 | SPU_027892 | Lv-Rpl35al |
| d1 | SPU_027899 | Lv-Glur1 |
| d1 | SPU_027927 | Lv-Nlr43 |
| d1 | SPU_027928 | Lv-Cpa |
| d1 | SPU_028004 | Lv-Slc7a6_3 |
| d1 | SPU_028040 | Lv-GucyL |
| d1 | SPU_028085 | Lv-Hypp_2931 |
| d1 | SPU_028200 | none |
| d1 | SPU_028203 | Lv-ApobL |
| d1 | SPU_028207 | Lv-Ccap/VasLr |
| d1 | SPU_028316 | Lv-Vav_2 |
| d1 | SPU_028379 | none |
| d1 | SPU_028392 | Lv-Opn4L_1 |
| d1 | SPU_028403 | Lv-Susd2_2 |
| d1 | SPU_028466 | Lv-Slc43a2L_1 |
| d1 | SPU_028497 | Lv-Tmprss3_1 |
| d1 | SPU_028511 | Lv-DspchdLB |
| d1 | SPU_028615 | none |
| d1 | SPU_028618 | Lv-RabO2 |
| d1 | SPU_028635 | Lv-C20orf3_2 |
| d1 | SPU_028686 | none |
| d1 | SPU_028720 | Lv-Pde9a_1 |
| d1 | SPU_028727 | Lv-Hypp_1236 |
| d1 | SPU_028749 | Lv-Mt5/MmpL2 |
| d1 | SPU_028802 | Lv-Rdh8_4 |
| d1 | SPU_028836 | NA |
| d1 | SPU_028844 | Lv-Ccp/Zp_3 |
| d1 | SPU_028862 | none |

| | | |
|---|---|---|
| d1 | SPU_030059 | Lv-Twi |
| d1 | SPU_030149 | Lv-Ash3L |
| d2 | SPU_000157 | Lv-DpL |
| d2 | SPU_000158 | NA |
| d2 | SPU_000353 | Lv-Hypp_299 |
| d2 | SPU_000427 | none |
| d2 | SPU_000482 | none |
| d2 | SPU_000657 | NA |
| d2 | SPU_000740 | NA |
| d2 | SPU_000837 | Lv-Clect_99 |
| d2 | SPU_000980 | Lv-Gal3st1-2 |
| d2 | SPU_001036 | Lv-Treh_1 |
| d2 | SPU_001405 | none |
| d2 | SPU_001750 | NA |
| d2 | SPU_002033 | Lv-Cuzd1 |
| d2 | SPU_002265 | Lv-PolppL_14 |
| d2 | SPU_002275 | Lv-Unc44_266 |
| d2 | SPU_002331 | Lv-Gal3st1L |
| d2 | SPU_002396 | Lv-Sacs-3 |
| d2 | SPU_002513 | Lv-SpdecL |
| d2 | SPU_002595 | Lv-PolppL_17 |
| d2 | SPU_002601 | none |
| d2 | SPU_002685 | Lv-Hypp_3105 |
| d2 | SPU_002921 | Lv-IceL1a |
| d2 | SPU_003103 | Lv-Slc10a1 |
| d2 | SPU_003266 | Lv-Anpep |
| d2 | SPU_003315 | none |
| d2 | SPU_003316 | none |
| d2 | SPU_003778 | NA |
| d2 | SPU_003901 | Lv-My5 |
| d2 | SPU_004196 | none |
| d2 | SPU_004233 | none |
| d2 | SPU_004475 | NA |
| d2 | SPU_004542 | Lv-NotchL_917 |
| d2 | SPU_004556 | NA |
| d2 | SPU_004659 | Lv-DisLA3 |
| d2 | SPU_004744 | Lv-Hypp_427 |
| d2 | SPU_004752 | Lv-Adora2aL_4 |

| | | |
|---|---|---|
| d2 | SPU_004869 | Lv-Sm30F |
| d2 | SPU_004922 | NA |
| d2 | SPU_005030 | none |
| d2 | SPU_005108 | Lv-Hypp_1565 |
| d2 | SPU_005377 | none |
| d2 | SPU_005536 | Lv-Hypp_1587 |
| d2 | SPU_005758 | Lv-Ldla/Gpcr |
| d2 | SPU_005783 | NA |
| d2 | SPU_006327 | Lv-Unc44L_52 |
| d2 | SPU_006397 | Lv-Hypp_479 |
| d2 | SPU_006465 | Lv-Hypp_482 |
| d2 | SPU_006469 | none |
| d2 | SPU_006659 | Lv-Srcr/Lrr1 |
| d2 | SPU_006941 | NA |
| d2 | SPU_007009 | Lv-Slc16a6_1 |
| d2 | SPU_007147 | Lv-Ngn |
| d2 | SPU_007423 | NA |
| d2 | SPU_007692 | none |
| d2 | SPU_007826 | Lv-Hypp_1748 |
| d2 | SPU_007985 | Lv-Slc7 |
| d2 | SPU_007992 | NA |
| d2 | SPU_008127 | Lv-Gpr21L_1 |
| d2 | SPU_008490 | Lv-C1orf222L |
| d2 | SPU_008512 | Lv-Naalad2L_7 |
| d2 | SPU_008598 | NA |
| d2 | SPU_008661 | Lv-Asp_11 |
| d2 | SPU_008813 | Lv-Mgam_6 |
| d2 | SPU_008836 | Lv-Srcr72 |
| d2 | SPU_008964 | NA |
| d2 | SPU_009626 | NA |
| d2 | SPU_009979 | NA |
| d2 | SPU_010069 | none |
| d2 | SPU_010124 | NA |
| d2 | SPU_010321 | Lv-Ank9 |
| d2 | SPU_010408 | NA |
| d2 | SPU_010640 | none |
| d2 | SPU_010657 | Lv-Chrm4L |
| d2 | SPU_010868 | NA |

| | | |
|---|---|---|
| d2 | SPU_011157 | Lv-Rab14_1 |
| d2 | SPU_011209 | Lv-Unc44_292 |
| d2 | SPU_011318 | Lv-Surreal/Gpcr11 |
| d2 | SPU_011579 | none |
| d2 | SPU_012077 | NA |
| d2 | SPU_012188 | Lv-Hypp_674 |
| d2 | SPU_012249 | Lv-Hypp_2022 |
| d2 | SPU_012326 | Lv-Hypp_681 |
| d2 | SPU_012351 | NA |
| d2 | SPU_012416 | Lv-Dnahc2 |
| d2 | SPU_012567 | none |
| d2 | SPU_012594 | NA |
| d2 | SPU_012783 | NA |
| d2 | SPU_012975 | NA |
| d2 | SPU_013121 | none |
| d2 | SPU_013464 | Lv-Phox2 |
| d2 | SPU_013530 | Lv-B3gnt5_2 |
| d2 | SPU_013587 | Lv-Slc25a35_1 |
| d2 | SPU_013622 | none |
| d2 | SPU_013684 | NA |
| d2 | SPU_013709 | Lv-Vc1_1 |
| d2 | SPU_013977 | NA |
| d2 | SPU_014031 | Lv-Fn3f_64 |
| d2 | SPU_014215 | none |
| d2 | SPU_014401 | Lv-NtL |
| d2 | SPU_014982 | NA |
| d2 | SPU_015181 | Lv-Hypp_2195 |
| d2 | SPU_015326 | Lv-Hypp_781 |
| d2 | SPU_015338 | Lv-Serrp12 |
| d2 | SPU_015434 | Lv-Slc7a6_5 |
| d2 | SPU_015488 | Lv-Gluld1_13 |
| d2 | SPU_015541 | Lv-Tnni3k |
| d2 | SPU_015702 | none |
| d2 | SPU_016164 | NA |
| d2 | SPU_016182 | none |
| d2 | SPU_016301 | Lv-Cuzd1L_1 |
| d2 | SPU_016324 | NA |
| d2 | SPU_016428 | Lv-Cpa2 |

| | | |
|---|---|---|
| d2 | SPU_016545 | NA |
| d2 | SPU_016566 | NA |
| d2 | SPU_016610 | Lv-Aadc_1 |
| d2 | SPU_017096 | Lv-Surreal/Gpcr88 |
| d2 | SPU_017497 | none |
| d2 | SPU_017526 | NA |
| d2 | SPU_017529 | Lv-Tlr089 |
| d2 | SPU_017660 | none |
| d2 | SPU_017704 | NA |
| d2 | SPU_018301 | none |
| d2 | SPU_018393 | Lv-Scratch |
| d2 | SPU_018650 | NA |
| d2 | SPU_019029 | Lv-Endrvt99 |
| d2 | SPU_019151 | none |
| d2 | SPU_019249 | Lv-Hypp_2429 |
| d2 | SPU_019281 | Lv-Chrm2L_5 |
| d2 | SPU_019618 | none |
| d2 | SPU_019711 | NA |
| d2 | SPU_019991 | Lv-Unc44_141 |
| d2 | SPU_020423 | none |
| d2 | SPU_020424 | NA |
| d2 | SPU_020665 | none |
| d2 | SPU_020895 | none |
| d2 | SPU_020919 | none |
| d2 | SPU_020934 | Lv-Drd2L_22 |
| d2 | SPU_020962 | Lv-Samhd1_5 |
| d2 | SPU_021111 | none |
| d2 | SPU_021242 | Lv-Ap |
| d2 | SPU_021244 | Lv-Fn3f_12 |
| d2 | SPU_021380 | NA |
| d2 | SPU_021468 | none |
| d2 | SPU_021637 | Lv-Hypp_2567 |
| d2 | SPU_021681 | Lv-Unc44_156 |
| d2 | SPU_021898 | none |
| d2 | SPU_021908 | Lv-Tlr214 |
| d2 | SPU_021993 | Lv-Fbgp_1 |
| d2 | SPU_021998 | Lv-Adrb2L_15 |
| d2 | SPU_022523 | NA |

| | | |
|---|---|---|
| d2 | SPU_022577 | Lv-Hypp_1049 |
| d2 | SPU_022655 | NA |
| d2 | SPU_022811 | Lv-Hypp_2633 |
| d2 | SPU_022978 | none |
| d2 | SPU_023474 | Lv-Hypp_1086 |
| d2 | SPU_023629 | NA |
| d2 | SPU_023767 | NA |
| d2 | SPU_023846 | Lv-Slc13a2_2 |
| d2 | SPU_023893 | Lv-Tlx3 |
| d2 | SPU_023913 | none |
| d2 | SPU_024123 | NA |
| d2 | SPU_024365 | Lv-Unc44_342 |
| d2 | SPU_024377 | none |
| d2 | SPU_024404 | Lv-Tlr129 |
| d2 | SPU_024715 | Lv-Cdx |
| d2 | SPU_025048 | NA |
| d2 | SPU_025070 | none |
| d2 | SPU_025227 | Lv-Nt5e_2 |
| d2 | SPU_025391 | none |
| d2 | SPU_025500 | Lv-Clca4L_1 |
| d2 | SPU_025722 | Lv-Cara12LB |
| d2 | SPU_026069 | Lv-Unc44_177 |
| d2 | SPU_026245 | Lv-EchnL_9 |
| d2 | SPU_026311 | NA |
| d2 | SPU_026316 | Lv-Drd2L_24 |
| d2 | SPU_026655 | Lv-Unc44_256 |
| d2 | SPU_026742 | Lv-Sbf2 |
| d2 | SPU_026868 | Lv-Vps13d_3 |
| d2 | SPU_026931 | NA |
| d2 | SPU_027114 | Lv-AnL7_1 |
| d2 | SPU_027128 | none |
| d2 | SPU_027129 | Lv-Tmem86aL |
| d2 | SPU_027263 | Lv-PppL_155 |
| d2 | SPU_027347 | Lv-Fibcd1_3 |
| d2 | SPU_027497 | none |
| d2 | SPU_027623 | Lv-Mist |
| d2 | SPU_027924 | Lv-UxyltL |
| d2 | SPU_028119 | NA |

| | | |
|---|---|---|
| d2 | SPU_028159 | Lv-Hrmn |
| d2 | SPU_028314 | Lv-Hypp_2949 |
| d2 | SPU_028549 | none |
| d2 | SPU_028552 | NA |
| d2 | SPU_028584 | none |
| d2 | SPU_028610 | none |
| d2 | SPU_028767 | Lv-PppL_290 |
| d2 | SPU_030081 | NA |
| d2 | SPU_030107 | Lv-TlrP51 |
| d2 | SPU_030109 | Lv-TlrP53 |
| d2 | SPU_030112 | NA |
| d3 | SPU_000120 | NA |
| d3 | SPU_000140 | Lv-Sst5L |
| d3 | SPU_000150 | none |
| d3 | SPU_000153 | Lv-AchE_8 |
| d3 | SPU_000314 | Lv-Cnga3_2 |
| d3 | SPU_000527 | Lv-CubnL_8 |
| d3 | SPU_000528 | Lv-Serp |
| d3 | SPU_000529 | Lv-Hypp_305 |
| d3 | SPU_000530 | Lv-Taar7eL |
| d3 | SPU_000691 | Lv-Clect/Tsp1 |
| d3 | SPU_000720 | Lv-Oprk1L_2 |
| d3 | SPU_000928 | Lv-Hypp_1293 |
| d3 | SPU_000929 | Lv-Hypp_1294 |
| d3 | SPU_000942 | Lv-Slc27a6-2 |
| d3 | SPU_001180 | NA |
| d3 | SPU_001342 | none |
| d3 | SPU_001398 | Lv-Cpa2L_2 |
| d3 | SPU_001443 | none |
| d3 | SPU_001516 | Lv-Mamdc4-16 |
| d3 | SPU_001625 | Lv-Chrm3L_2 |
| d3 | SPU_001663 | none |
| d3 | SPU_001821 | Lv-EchL3 |
| d3 | SPU_001868 | Lv-ALv-2 |
| d3 | SPU_001869 | NA |
| d3 | SPU_001871 | Lv-Unc44L_19 |
| d3 | SPU_001922 | Lv-Unc44_201 |
| d3 | SPU_001956 | Lv-Cnga3 |

| | | |
|---|---|---|
| d3 | SPU_001999 | Lv-Hypp_1363 |
| d3 | SPU_002048 | Lv-Samhd1L |
| d3 | SPU_002098 | Lv-Hypp_1370 |
| d3 | SPU_002237 | Lv-Hypp_1379 |
| d3 | SPU_002248 | Lv-Adrb3L |
| d3 | SPU_002283 | Lv-Hypp_1386 |
| d3 | SPU_002367 | Lv-ReelerL_1 |
| d3 | SPU_002453 | NA |
| d3 | SPU_002526 | Lv-Hypp_1400 |
| d3 | SPU_002630 | Lv-Hox8_1 |
| d3 | SPU_002832 | Lv-Cyp4VL3 |
| d3 | SPU_002841 | Lv-Hypp_1415 |
| d3 | SPU_002955 | Lv-Serrp11 |
| d3 | SPU_003098 | Lv-Hypp_1432 |
| d3 | SPU_003101 | Lv-Surreal/Gpcr6 |
| d3 | SPU_003125 | Lv-Sstr1L |
| d3 | SPU_003170 | Lv-Adam/TsL6 |
| d3 | SPU_003395 | NA |
| d3 | SPU_003423 | Lv-Hypp_1451 |
| d3 | SPU_003451 | Lv-Opsin2 |
| d3 | SPU_003571 | Lv-Hypp_1463 |
| d3 | SPU_003676 | Lv-YP30/FascL |
| d3 | SPU_003741 | none |
| d3 | SPU_003859 | Lv-Mgam_1 |
| d3 | SPU_003931 | NA |
| d3 | SPU_004087 | Lv-Rps21_1 |
| d3 | SPU_004135 | Lv-Cah10L |
| d3 | SPU_004182 | NA |
| d3 | SPU_004185 | Lv-Ache-10 |
| d3 | SPU_004186 | Lv-Bche |
| d3 | SPU_004421 | Lv-Lucf_2 |
| d3 | SPU_004803 | NA |
| d3 | SPU_004804 | Lv-Serp-3 |
| d3 | SPU_004831 | Lv-Gal/lec2 |
| d3 | SPU_004913 | Lv-Naalad2L_1 |
| d3 | SPU_005012 | Lv-Hypp_1555 |
| d3 | SPU_005013 | Lv-Clect/Tm_9 |
| d3 | SPU_005320 | NA |

| | | |
|---|---|---|
| d3 | SPU_005399 | Lv-Pi16 |
| d3 | SPU_005426 | Lv-Clect/Tsp2 |
| d3 | SPU_005578 | Lv-Adra1aL_2 |
| d3 | SPU_005595 | Lv-Sstr4L |
| d3 | SPU_005635 | Lv-Tmem20_7 |
| d3 | SPU_005658 | none |
| d3 | SPU_005909 | NA |
| d3 | SPU_005951 | Lv-Gal3st3L |
| d3 | SPU_005952 | Lv-Slco4a1 |
| d3 | SPU_006010 | NA |
| d3 | SPU_006032 | Lv-Hypp_1626 |
| d3 | SPU_006049 | Lv-Hypp_1627 |
| d3 | SPU_006075 | Lv-AadacL_1 |
| d3 | SPU_006150 | Lv-Tbx1 |
| d3 | SPU_006330 | Lv-Amy |
| d3 | SPU_006417 | none |
| d3 | SPU_006418 | Lv-Hypp_480 |
| d3 | SPU_006561 | Lv-Gat |
| d3 | SPU_006631 | Lv-Dlg1 |
| d3 | SPU_007037 | NA |
| d3 | SPU_007043 | Lv-Slc32a1_1 |
| d3 | SPU_007329 | Lv-Pnlip_1 |
| d3 | SPU_007356 | Lv-Hypp_1716 |
| d3 | SPU_007492 | Lv-Aldh5a1 |
| d3 | SPU_007538 | Lv-Cd38 |
| d3 | SPU_007616 | none |
| d3 | SPU_007737 | none |
| d3 | SPU_007793 | Lv-Asp_7 |
| d3 | SPU_007811 | Lv-Slc5 |
| d3 | SPU_007814 | Lv-Slc46a1L |
| d3 | SPU_007816 | none |
| d3 | SPU_007929 | Lv-Dnase1 |
| d3 | SPU_007930 | Lv-Dnaseg |
| d3 | SPU_008056 | NA |
| d3 | SPU_008159 | Lv-Kcnk13 |
| d3 | SPU_008212 | Lv-Agtr1 |
| d3 | SPU_008352 | none |
| d3 | SPU_008370 | Lv-Dbh_1 |

| | | |
|---|---|---|
| d3 | SPU_008455 | Lv-Iars_1 |
| d3 | SPU_008812 | Lv-Si2 |
| d3 | SPU_008821 | NA |
| d3 | SPU_008854 | none |
| d3 | SPU_008864 | Lv-Timp2d |
| d3 | SPU_008950 | Lv-Gpcr |
| d3 | SPU_009034 | Lv-Unc44L_62 |
| d3 | SPU_009041 | Lv-Ache_16 |
| d3 | SPU_009091 | NA |
| d3 | SPU_009192 | Lv-Rergl3 |
| d3 | SPU_009344 | NA |
| d3 | SPU_009353 | NA |
| d3 | SPU_009368 | Lv-Cts3 |
| d3 | SPU_009371 | Lv-Slc22a5 |
| d3 | SPU_009416 | Lv-Ugt2b34_2 |
| d3 | SPU_009438 | none |
| d3 | SPU_009458 | Lv-Hao1_15 |
| d3 | SPU_009491 | Lv-Mocs1 |
| d3 | SPU_009519 | Lv-Hypp_579 |
| d3 | SPU_009790 | Lv-Hypp_1871 |
| d3 | SPU_009806 | Lv-Hypp_1873 |
| d3 | SPU_009905 | none |
| d3 | SPU_009989 | Lv-Srcr81 |
| d3 | SPU_010001 | NA |
| d3 | SPU_010077 | none |
| d3 | SPU_010369 | Lv-Slc22a4 |
| d3 | SPU_010370 | NA |
| d3 | SPU_010371 | Lv-Slc22a3_3 |
| d3 | SPU_010394 | none |
| d3 | SPU_010450 | none |
| d3 | SPU_010930 | none |
| d3 | SPU_011041 | Lv-Slc7a11_4 |
| d3 | SPU_011097 | Lv-Nlr196 |
| d3 | SPU_011131 | Lv-Hypp_639 |
| d3 | SPU_011163 | Lv-Clect_25 |
| d3 | SPU_011185 | Lv-Wscd2_1 |
| d3 | SPU_011338 | none |
| d3 | SPU_011342 | NA |

| | | |
|---|---|---|
| d3 | SPU_011463 | Lv-Hypp_648 |
| d3 | SPU_011522 | none |
| d3 | SPU_011586 | Lv-AT_V_1 |
| d3 | SPU_011600 | NA |
| d3 | SPU_011637 | Lv-Lrr/Igr_14 |
| d3 | SPU_011650 | Lv-Peci_1 |
| d3 | SPU_011819 | Lv-EmapL1 |
| d3 | SPU_011865 | Lv-Kiaa1161L_1 |
| d3 | SPU_011869 | Lv-Ttrhde |
| d3 | SPU_012012 | none |
| d3 | SPU_012020 | Lv-Cpa3L_2 |
| d3 | SPU_012021 | Lv-Cpa1L_1 |
| d3 | SPU_012145 | none |
| d3 | SPU_012246 | Lv-Acp5 |
| d3 | SPU_012255 | Lv-Sec14L2_1 |
| d3 | SPU_012256 | none |
| d3 | SPU_012273 | Lv-Hrh3L |
| d3 | SPU_012341 | Lv-Dnase1_1 |
| d3 | SPU_012342 | Lv-Sult1c_3 |
| d3 | SPU_012355 | Lv-Igf_16 |
| d3 | SPU_012418 | Lv-Dnahc2_1 |
| d3 | SPU_012474 | Lv-Amy2a-2 |
| d3 | SPU_012539 | NA |
| d3 | SPU_012646 | Lv-Orct |
| d3 | SPU_012742 | NA |
| d3 | SPU_012771 | NA |
| d3 | SPU_012907 | none |
| d3 | SPU_012910 | Lv-Secrtag |
| d3 | SPU_013033 | Lv-Hypp_2065 |
| d3 | SPU_013080 | Lv-Slc16a12L_8 |
| d3 | SPU_013142 | Lv-SommL |
| d3 | SPU_013233 | Lv-Hypp_708 |
| d3 | SPU_013292 | none |
| d3 | SPU_013359 | Lv-Hrh2_2 |
| d3 | SPU_013506 | Lv-Asp_14 |
| d3 | SPU_013635 | none |
| d3 | SPU_013649 | Lv-Clect_28 |
| d3 | SPU_013685 | none |

| | | |
|---|---|---|
| d3 | SPU_013726 | Lv-Kctd15 |
| d3 | SPU_013770 | Lv-Ahcy |
| d3 | SPU_013781 | Lv-Chrm4L_2 |
| d3 | SPU_013782 | Lv-Hrh1L_7 |
| d3 | SPU_013937 | Lv-PpcL |
| d3 | SPU_013938 | Lv-Hypp_3123 |
| d3 | SPU_014040 | Lv-Gpr112_4 |
| d3 | SPU_014139 | Lv-Slc46a3L_2 |
| d3 | SPU_014202 | Lv-Slc22a3_5 |
| d3 | SPU_014218 | Lv-Gal/lec/Egf/Tm |
| d3 | SPU_014229 | Lv-MacpfF.1 |
| d3 | SPU_014264 | none |
| d3 | SPU_014265 | Lv-Asp_16 |
| d3 | SPU_014268 | none |
| d3 | SPU_014303 | Lv-Crabp1 |
| d3 | SPU_014409 | none |
| d3 | SPU_014423 | none |
| d3 | SPU_014447 | Lv-Hypp_2150 |
| d3 | SPU_014551 | Lv-So/Gpcr |
| d3 | SPU_014640 | Lv-Pam_2 |
| d3 | SPU_014671 | Lv-Amy_3 |
| d3 | SPU_014846 | Lv-Mapkbp1L |
| d3 | SPU_014849 | Lv-Hypp_2174 |
| d3 | SPU_014859 | Lv-Srcr118 |
| d3 | SPU_014952 | none |
| d3 | SPU_014953 | Lv-Unc44L_104 |
| d3 | SPU_014990 | none |
| d3 | SPU_015000 | Lv-Bhmt_7 |
| d3 | SPU_015099 | Lv-PlbL |
| d3 | SPU_015100 | Lv-Phlpb |
| d3 | SPU_015113 | Lv-Pnlip_8 |
| d3 | SPU_015155 | Lv-Prrdh |
| d3 | SPU_015178 | Lv-Cpa2L |
| d3 | SPU_015179 | Lv-Cpa2L_1 |
| d3 | SPU_015186 | Lv-Drd2L_19 |
| d3 | SPU_015257 | Lv-Vdac2 |
| d3 | SPU_015258 | Lv-Hypp_2203 |
| d3 | SPU_015278 | NA |

| | | |
|---|---|---|
| d3 | SPU_015510 | NA |
| d3 | SPU_015670 | Lv-YP30/FascL_6 |
| d3 | SPU_015798 | none |
| d3 | SPU_015892 | Lv-Scnn1aL |
| d3 | SPU_015893 | Lv-Scnn1aL_1 |
| d3 | SPU_015943 | none |
| d3 | SPU_016147 | Lv-Wdr86 |
| d3 | SPU_016178 | Lv-Amds_2 |
| d3 | SPU_016307 | none |
| d3 | SPU_016308 | none |
| d3 | SPU_016389 | Lv-Lrrtm1L |
| d3 | SPU_016419 | Lv-Tbc1d4 |
| d3 | SPU_016427 | NA |
| d3 | SPU_016479 | Lv-Unc44L_114 |
| d3 | SPU_016681 | Lv-Clect/Egf_3 |
| d3 | SPU_016693 | Lv-C1galt1 |
| d3 | SPU_016700 | Lv-Gdnfa1 |
| d3 | SPU_016779 | none |
| d3 | SPU_016814 | Lv-Gpr151L_10 |
| d3 | SPU_016881 | Lv-SoxE |
| d3 | SPU_016927 | Lv-Unc44_119 |
| d3 | SPU_016941 | Lv-Htr1dL |
| d3 | SPU_017007 | Lv-Clect_111 |
| d3 | SPU_017024 | Lv-Kremen1 |
| d3 | SPU_017031 | NA |
| d3 | SPU_017111 | NA |
| d3 | SPU_017129 | NA |
| d3 | SPU_017146 | NA |
| d3 | SPU_017239 | Lv-064_3 |
| d3 | SPU_017250 | NA |
| d3 | SPU_017257 | NA |
| d3 | SPU_017291 | Lv-Chst15L_1 |
| d3 | SPU_017294 | Lv-Hypp_872 |
| d3 | SPU_017386 | NA |
| d3 | SPU_017387 | NA |
| d3 | SPU_017416 | none |
| d3 | SPU_017461 | NA |
| d3 | SPU_017564 | NA |

| | | |
|---|---|---|
| d3 | SPU_017647 | Lv-Bmp11b |
| d3 | SPU_017737 | none |
| d3 | SPU_017789 | NA |
| d3 | SPU_017791 | Lv-Fcolf_1 |
| d3 | SPU_017879 | NA |
| d3 | SPU_017888 | Lv-Clecsf13 |
| d3 | SPU_018003 | Lv-Aifm2 |
| d3 | SPU_018177 | Lv-Bbox1_6 |
| d3 | SPU_018252 | Lv-Srcr141 |
| d3 | SPU_018286 | none |
| d3 | SPU_018465 | Lv-2Apt |
| d3 | SPU_018658 | Lv-SommL_2 |
| d3 | SPU_018734 | Lv-Hypp_2403 |
| d3 | SPU_018781 | Lv-Hypp_3041 |
| d3 | SPU_018826 | Lv-5ht1a |
| d3 | SPU_018984 | none |
| d3 | SPU_019038 | none |
| d3 | SPU_019086 | NA |
| d3 | SPU_019105 | NA |
| d3 | SPU_019108 | Lv-Adrb2L_12 |
| d3 | SPU_019167 | NA |
| d3 | SPU_019240 | Lv-Rxfp2 |
| d3 | SPU_019284 | Lv-Rad18L |
| d3 | SPU_019293 | NA |
| d3 | SPU_019437 | Lv-Clect/Tsp3 |
| d3 | SPU_019464 | Lv-Cyp3L10 |
| d3 | SPU_019480 | NA |
| d3 | SPU_019581 | Lv-Clca_11 |
| d3 | SPU_019605 | none |
| d3 | SPU_019849 | Lv-Sec14L2_5 |
| d3 | SPU_019928 | none |
| d3 | SPU_019944 | Lv-Ccp/Egf/Tsp1/HyalinL3 |
| d3 | SPU_019946 | none |
| d3 | SPU_019956 | Lv-Hypp_2475 |
| d3 | SPU_019983 | Lv-Dan |
| d3 | SPU_020043 | NA |
| d3 | SPU_020107 | Lv-AchE_4 |
| d3 | SPU_020113 | Lv-Slsp_12 |

| | | |
|---|---|---|
| d3 | SPU_020208 | Lv-Hypp_2496 |
| d3 | SPU_020237 | NA |
| d3 | SPU_020300 | none |
| d3 | SPU_020318 | none |
| d3 | SPU_020354 | NA |
| d3 | SPU_020516 | Lv-Gyar |
| d3 | SPU_020551 | none |
| d3 | SPU_020633 | Lv-Pcalr |
| d3 | SPU_020876 | Lv-Cyp26L |
| d3 | SPU_020883 | Lv-Hypp_2524 |
| d3 | SPU_020921 | Lv-Hypp_1005 |
| d3 | SPU_021080 | Lv-Ankrd34c |
| d3 | SPU_021228 | Lv-Cf5/8L1 |
| d3 | SPU_021363 | Lv-Oprm1L |
| d3 | SPU_021423 | Lv-Sec14LX |
| d3 | SPU_021545 | NA |
| d3 | SPU_021555 | Lv-Salmfap |
| d3 | SPU_021752 | Lv-Gba_3 |
| d3 | SPU_021975 | NA |
| d3 | SPU_022047 | Lv-Pla2g1b_8 |
| d3 | SPU_022050 | Lv-Sc4mol_2 |
| d3 | SPU_022077 | Lv-Hypp_2596 |
| d3 | SPU_022102 | Lv-PcbpdB1L |
| d3 | SPU_022123 | none |
| d3 | SPU_022149 | Lv-Srcr180 |
| d3 | SPU_022152 | Lv-Clect_41 |
| d3 | SPU_022409 | none |
| d3 | SPU_022505 | Lv-Naglu_2 |
| d3 | SPU_022507 | none |
| d3 | SPU_022530 | none |
| d3 | SPU_022545 | Lv-Hypp_2616 |
| d3 | SPU_022551 | Lv-Hypp_1048 |
| d3 | SPU_022600 | NA |
| d3 | SPU_022613 | Lv-Hypp_3004 |
| d3 | SPU_022773 | Lv-Mtnr1aL_3 |
| d3 | SPU_022951 | NA |
| d3 | SPU_022954 | none |
| d3 | SPU_023043 | Lv-Hypp_1066 |

| | | |
|---|---|---|
| d3 | SPU_023234 | NA |
| d3 | SPU_023242 | Lv-Hypp_2659 |
| d3 | SPU_023364 | Lv-Slco4a1_15 |
| d3 | SPU_023385 | none |
| d3 | SPU_023441 | Lv-Tbx4 |
| d3 | SPU_023485 | Lv-Pnlip_10 |
| d3 | SPU_023512 | Lv-Tsp/Igf |
| d3 | SPU_023540 | none |
| d3 | SPU_023548 | Lv-Fic |
| d3 | SPU_023675 | Lv-Chrna2 |
| d3 | SPU_023884 | none |
| d3 | SPU_023911 | none |
| d3 | SPU_024012 | NA |
| d3 | SPU_024055 | Lv-Myls4 |
| d3 | SPU_024058 | none |
| d3 | SPU_024246 | Lv-Tsc2 |
| d3 | SPU_024422 | NA |
| d3 | SPU_024486 | Lv-Dsf |
| d3 | SPU_024539 | Lv-Prprp |
| d3 | SPU_024630 | NA |
| d3 | SPU_024713 | Lv-Hypp_2748 |
| d3 | SPU_024730 | Lv-Fth1_1 |
| d3 | SPU_024749 | Lv-Galr2L_29 |
| d3 | SPU_024836 | Lv-Cyp27L |
| d3 | SPU_024907 | Lv-Slsp_22 |
| d3 | SPU_024942 | NA |
| d3 | SPU_024977 | none |
| d3 | SPU_024994 | Lv-Follistatin |
| d3 | SPU_025379 | Lv-Hypp_2788 |
| d3 | SPU_025419 | none |
| d3 | SPU_025565 | Lv-ScavrB2L2 |
| d3 | SPU_025605 | NA |
| d3 | SPU_025652 | none |
| d3 | SPU_025666 | Lv-Notch2_12 |
| d3 | SPU_025752 | none |
| d3 | SPU_025853 | NA |
| d3 | SPU_025855 | Lv-Hypp_2805 |
| d3 | SPU_025925 | Lv-Hypp_1160 |

| | | |
|---|---|---|
| d3 | SPU_025969 | Lv-Cyp1L8 |
| d3 | SPU_026003 | Lv-Kcna3 |
| d3 | SPU_026119 | Lv-MacpfD.3 |
| d3 | SPU_026128 | NA |
| d3 | SPU_026325 | none |
| d3 | SPU_026329 | Lv-Mgst1 |
| d3 | SPU_026360 | Lv-Cyp2L61 |
| d3 | SPU_026443 | Lv-Pm20d2 |
| d3 | SPU_026586 | none |
| d3 | SPU_026604 | NA |
| d3 | SPU_026942 | NA |
| d3 | SPU_027077 | none |
| d3 | SPU_027230 | NA |
| d3 | SPU_027274 | Lv-Rpia_1 |
| d3 | SPU_027398 | none |
| d3 | SPU_027543 | NA |
| d3 | SPU_027549 | Lv-Adora2aL_7 |
| d3 | SPU_027565 | Lv-CpBL |
| d3 | SPU_027769 | Lv-Rpl14_1 |
| d3 | SPU_027797 | Lv-Slc7a11_11 |
| d3 | SPU_027835 | Lv-Hypp_2921 |
| d3 | SPU_027838 | NA |
| d3 | SPU_027859 | Lv-Stom_6 |
| d3 | SPU_027894 | none |
| d3 | SPU_027907 | Lv-Hypp_1218 |
| d3 | SPU_027929 | NA |
| d3 | SPU_028021 | Lv-Acel_7 |
| d3 | SPU_028037 | Lv-Oprl1L |
| d3 | SPU_028044 | NA |
| d3 | SPU_028116 | Lv-Angptl1_1 |
| d3 | SPU_028187 | Lv-Fb2 |
| d3 | SPU_028278 | none |
| d3 | SPU_028370 | Lv-Dnah1 |
| d3 | SPU_028420 | none |
| d3 | SPU_028519 | Lv-Ebr1L_10 |
| d3 | SPU_028569 | Lv-Pla2g1b_4 |
| d3 | SPU_028683 | Lv-Vtgn3 |
| d3 | SPU_028714 | NA |

| | | |
|---|---|---|
| d3 | SPU_028756 | Lv-MacpfE.2 |
| d3 | SPU_028766 | NA |
| d3 | SPU_028792 | Lv-Hypp_2970 |
| d3 | SPU_028799 | none |
| d3 | SPU_028821 | NA |
| d3 | SPU_028841 | Lv-Mgam_4 |
| d3 | SPU_028853 | none |
| d3 | SPU_028937 | Lv-Naalad2L_11 |
| e1 | SPU_000330 | Lv-HgfL |
| e1 | SPU_000389 | Lv-TekL_2 |
| e1 | SPU_000409 | NA |
| e1 | SPU_000621 | Lv-Lrp1L_1 |
| e1 | SPU_000791 | Lv-Ids_5 |
| e1 | SPU_000921 | NA |
| e1 | SPU_001015 | none |
| e1 | SPU_001126 | none |
| e1 | SPU_001275 | none |
| e1 | SPU_001600 | Lv-Mpped2 |
| e1 | SPU_001683 | NA |
| e1 | SPU_001708 | Lv-Hspg2 |
| e1 | SPU_001965 | Lv-Adra1bL_2 |
| e1 | SPU_002001 | Lv-Btbd17L |
| e1 | SPU_002026 | none |
| e1 | SPU_002542 | Lv-Hypp_1402 |
| e1 | SPU_002636 | Lv-Bbox1_1 |
| e1 | SPU_002795 | Lv-Myostatin |
| e1 | SPU_002848 | Lv-Unc44L_27 |
| e1 | SPU_002964 | NA |
| e1 | SPU_003003 | Lv-Adra1bL_3 |
| e1 | SPU_003090 | Lv-Chrna6L_3 |
| e1 | SPU_003214 | Lv-Col9a1 |
| e1 | SPU_003264 | Lv-Apn4 |
| e1 | SPU_003303 | Lv-Nlr175 |
| e1 | SPU_003334 | none |
| e1 | SPU_003587 | Lv-Chst3 |
| e1 | SPU_003809 | Lv-Unc44L_32 |
| e1 | SPU_003946 | Lv-Hypp_401 |
| e1 | SPU_003988 | none |

| | | |
|---|---|---|
| e1 | SPU_004111 | none |
| e1 | SPU_004211 | Lv-Hypp_3099 |
| e1 | SPU_004346 | NA |
| e1 | SPU_004425 | NA |
| e1 | SPU_004501 | Lv-Fn3/Igf_28 |
| e1 | SPU_004629 | Lv-Drd2 |
| e1 | SPU_004791 | Lv-Tlr023 |
| e1 | SPU_004851 | Lv-Hypp_1548 |
| e1 | SPU_004995 | NA |
| e1 | SPU_005058 | Lv-Rsl1d1-2 |
| e1 | SPU_005064 | NA |
| e1 | SPU_005247 | NA |
| e1 | SPU_005323 | Lv-Samd9L_4 |
| e1 | SPU_005460 | NA |
| e1 | SPU_005488 | NA |
| e1 | SPU_005549 | none |
| e1 | SPU_005725 | NA |
| e1 | SPU_005926 | NA |
| e1 | SPU_005927 | none |
| e1 | SPU_005980 | none |
| e1 | SPU_006067 | NA |
| e1 | SPU_006070 | Lv-Ciapin1_1 |
| e1 | SPU_006185 | none |
| e1 | SPU_006208 | Lv-Gpr112L_10 |
| e1 | SPU_006245 | none |
| e1 | SPU_006376 | Lv-Hypp_1658 |
| e1 | SPU_006615 | Lv-Unc44L_54 |
| e1 | SPU_006728 | Lv-Igsf10L |
| e1 | SPU_006741 | NA |
| e1 | SPU_007269 | Lv-Hypp_1710 |
| e1 | SPU_007300 | Lv-Mrgbp_1 |
| e1 | SPU_007372 | Lv-Srcr15 |
| e1 | SPU_007440 | Lv-Z282 |
| e1 | SPU_007446 | NA |
| e1 | SPU_007535 | NA |
| e1 | SPU_007584 | Lv-March4L |
| e1 | SPU_007647 | Lv-Hypp_1740 |
| e1 | SPU_007761 | Lv-Galr2L_10 |

| | | |
|---|---|---|
| e1 | SPU_007861 | NA |
| e1 | SPU_008269 | Lv-Hypp_1774 |
| e1 | SPU_008419 | Lv-Apba1 |
| e1 | SPU_008675 | Lv-Hypp_556 |
| e1 | SPU_008782 | none |
| e1 | SPU_008861 | Lv-Timp2b |
| e1 | SPU_009244 | Lv-Egfib_1 |
| e1 | SPU_009294 | NA |
| e1 | SPU_009391 | Lv-Hypp_1848 |
| e1 | SPU_009486 | NA |
| e1 | SPU_009576 | Lv-Igf |
| e1 | SPU_009582 | NA |
| e1 | SPU_009793 | Lv-Rdh8_7 |
| e1 | SPU_010104 | Lv-Clca_5 |
| e1 | SPU_010222 | NA |
| e1 | SPU_010286 | NA |
| e1 | SPU_010320 | Lv-Tlr176 |
| e1 | SPU_010350 | Lv-Wdr90 |
| e1 | SPU_010454 | NA |
| e1 | SPU_010527 | Lv-Traf |
| e1 | SPU_010834 | none |
| e1 | SPU_010893 | Lv-Alox5_1 |
| e1 | SPU_011150 | Lv-Hypp_640 |
| e1 | SPU_011236 | Lv-Ache_19 |
| e1 | SPU_011519 | Lv-Mib1 |
| e1 | SPU_011793 | Lv-Gpr126L_1 |
| e1 | SPU_011803 | none |
| e1 | SPU_011881 | Lv-Lonp1 |
| e1 | SPU_011883 | Lv-Tll2L_1 |
| e1 | SPU_011987 | none |
| e1 | SPU_012123 | none |
| e1 | SPU_012234 | NA |
| e1 | SPU_012365 | NA |
| e1 | SPU_012379 | Lv-Loxhd1_3 |
| e1 | SPU_012435 | Lv-Hrh2L_6 |
| e1 | SPU_012510 | Lv-Hypp_2037 |
| e1 | SPU_012645 | Lv-Z166 |
| e1 | SPU_013141 | Lv-Gpr151L_4 |

| | | |
|---|---|---|
| e1 | SPU_013334 | Lv-MucL_4 |
| e1 | SPU_013436 | Lv-Gsx |
| e1 | SPU_013537 | Lv-Hypp_719 |
| e1 | SPU_013589 | none |
| e1 | SPU_013628 | Lv-Ccdc108_1 |
| e1 | SPU_013654 | Lv-A2mgL |
| e1 | SPU_013830 | NA |
| e1 | SPU_014047 | Lv-Utp20_3 |
| e1 | SPU_014277 | none |
| e1 | SPU_014485 | none |
| e1 | SPU_014511 | NA |
| e1 | SPU_014636 | none |
| e1 | SPU_014741 | Lv-Galr2L_17 |
| e1 | SPU_014768 | Lv-Cts8 |
| e1 | SPU_014785 | Lv-Tck |
| e1 | SPU_014838 | Lv-Hypp_2172 |
| e1 | SPU_014965 | NA |
| e1 | SPU_015065 | Lv-Clect/Pan_10 |
| e1 | SPU_015097 | Lv-Guan/Cyc |
| e1 | SPU_015664 | Lv-Hypp_796 |
| e1 | SPU_015777 | none |
| e1 | SPU_015810 | Lv-Hdac4L |
| e1 | SPU_015964 | NA |
| e1 | SPU_016109 | none |
| e1 | SPU_016195 | Lv-Srcr132 |
| e1 | SPU_016329 | none |
| e1 | SPU_016373 | Lv-Srcr133 |
| e1 | SPU_016535 | Lv-Echn35 |
| e1 | SPU_016623 | none |
| e1 | SPU_016826 | Lv-Pecr_1 |
| e1 | SPU_016863 | none |
| e1 | SPU_016900 | Lv-Ugt2b7 |
| e1 | SPU_016926 | Lv-Nlr143 |
| e1 | SPU_017127 | Lv-Srcr137 |
| e1 | SPU_017261 | Lv-Nbn1 |
| e1 | SPU_017337 | Lv-Slc16a12L_13 |
| e1 | SPU_017353 | Lv-Hypp_876 |
| e1 | SPU_017449 | NA |

| | | |
|---|---|---|
| e1 | SPU_017766 | Lv-Fn3f_58 |
| e1 | SPU_017783 | none |
| e1 | SPU_017839 | Lv-Ank2 |
| e1 | SPU_017864 | Lv-Gpr1L_4 |
| e1 | SPU_017887 | Lv-Clect/PanApple/Scop |
| e1 | SPU_018116 | Lv-Hypp_3044 |
| e1 | SPU_018384 | Lv-Nlr120 |
| e1 | SPU_018476 | Lv-Fn3f_56 |
| e1 | SPU_018531 | Lv-Slc17a5_4 |
| e1 | SPU_018579 | Lv-Llda/Egf/Ig/Lnb/Tm7/Gpcr |
| e1 | SPU_018661 | NA |
| e1 | SPU_018711 | NA |
| e1 | SPU_018828 | NA |
| e1 | SPU_018868 | Lv-Hypp_2408 |
| e1 | SPU_018959 | NA |
| e1 | SPU_018989 | Lv-Hypp_2415 |
| e1 | SPU_019244 | NA |
| e1 | SPU_019279 | Lv-ColpL |
| e1 | SPU_019346 | none |
| e1 | SPU_019364 | none |
| e1 | SPU_019398 | Lv-Ddc |
| e1 | SPU_019467 | none |
| e1 | SPU_019576 | Lv-Clect/Egf_11 |
| e1 | SPU_019667 | Lv-Cecr1 |
| e1 | SPU_019680 | none |
| e1 | SPU_019835 | Lv-TlrP20 |
| e1 | SPU_019895 | none |
| e1 | SPU_020313 | NA |
| e1 | SPU_020452 | Lv-Grp94 |
| e1 | SPU_020561 | Lv-Hypp_168 |
| e1 | SPU_020818 | Lv-Echn10 |
| e1 | SPU_020824 | Lv-CubnL |
| e1 | SPU_020846 | Lv-Z233 |
| e1 | SPU_021083 | Lv-Fn3f_59 |
| e1 | SPU_021357 | Lv-Acel_5 |
| e1 | SPU_021373 | Lv-Chrm1L |
| e1 | SPU_021521 | Lv-PppL_273 |
| e1 | SPU_021754 | Lv-Mtd |

| | | |
|---|---|---|
| e1 | SPU_021815 | NA |
| e1 | SPU_021849 | NA |
| e1 | SPU_022341 | Lv-Unc44_159 |
| e1 | SPU_022400 | Lv-Hypp_2607 |
| e1 | SPU_022499 | Lv-Ptpn13L |
| e1 | SPU_022522 | none |
| e1 | SPU_022535 | NA |
| e1 | SPU_022566 | Lv-Srcrr2 |
| e1 | SPU_022617 | Lv-Hypp_3005 |
| e1 | SPU_022838 | Lv-Il17-7 |
| e1 | SPU_022990 | Lv-Slc35e2 |
| e1 | SPU_023178 | none |
| e1 | SPU_023181 | NA |
| e1 | SPU_023248 | Lv-Smarcc2_1 |
| e1 | SPU_023405 | Lv-Adrb2L_18 |
| e1 | SPU_023545 | NA |
| e1 | SPU_023826 | Lv-Oit3L_1 |
| e1 | SPU_023987 | NA |
| e1 | SPU_024113 | Lv-Cyp2L56 |
| e1 | SPU_024474 | NA |
| e1 | SPU_024651 | Lv-Tmem145 |
| e1 | SPU_024673 | Lv-Sema/Psi_2 |
| e1 | SPU_024906 | NA |
| e1 | SPU_025020 | Lv-Hypp_1135 |
| e1 | SPU_025035 | Lv-HtrL |
| e1 | SPU_025109 | Lv-EhH2b_83 |
| e1 | SPU_025216 | NA |
| e1 | SPU_025231 | Lv-Hypp_2780 |
| e1 | SPU_025433 | Lv-Fn3/Igf_44 |
| e1 | SPU_025448 | Lv-Thsd7b_1 |
| e1 | SPU_025584 | Lv-Tbr |
| e1 | SPU_025673 | NA |
| e1 | SPU_025837 | Lv-LrpL_7 |
| e1 | SPU_026320 | Lv-Nlr176_1 |
| e1 | SPU_026651 | NA |
| e1 | SPU_026652 | Lv-Slc16a5L |
| e1 | SPU_026858 | none |
| e1 | SPU_026904 | Lv-Npr1_2 |

| | | |
|---|---|---|
| e1 | SPU_026976 | Lv-Unc44_180 |
| e1 | SPU_026992 | Lv-Ralgps1L |
| e1 | SPU_027014 | Lv-Ptprfn2(novel) |
| e1 | SPU_027059 | Lv-Hao1_13 |
| e1 | SPU_027118 | Lv-Vwd/Vwcp |
| e1 | SPU_027442 | Lv-PppL_284 |
| e1 | SPU_027466 | NA |
| e1 | SPU_027610 | Lv-Nlr135 |
| e1 | SPU_027619 | Lv-Srcr222 |
| e1 | SPU_027742 | Lv-Hypp_1211 |
| e1 | SPU_027971 | Lv-Cacng5L |
| e1 | SPU_028048 | Lv-Nrp2L_2 |
| e1 | SPU_028171 | Lv-SphH2b1 |
| e1 | SPU_028281 | Lv-Hypp_2946 |
| e1 | SPU_028292 | Lv-Unc44_186 |
| e1 | SPU_028342 | Lv-Tbrg1 |
| e1 | SPU_028364 | NA |
| e1 | SPU_028510 | Lv-B7L5 |
| e1 | SPU_028758 | Lv-Aadac |
| e1 | SPU_028806 | NA |
| e1 | SPU_028902 | Lv-IgfbpL7 |
| e1 | SPU_030125 | NA |
| e1 | SPU_030196 | Lv-Il17-18 |
| e2 | SPU_000543 | Lv-Clect/Egf/Tm |
| e2 | SPU_000642 | NA |
| e2 | SPU_000651 | NA |
| e2 | SPU_000716 | Lv-PppL_171 |
| e2 | SPU_000798 | Lv-Gabbr1 |
| e2 | SPU_000808 | Lv-Hypp_1288 |
| e2 | SPU_000828 | Lv-Sm30D |
| e2 | SPU_000888 | none |
| e2 | SPU_000907 | NA |
| e2 | SPU_000978 | Lv-Plxna4 |
| e2 | SPU_001023 | Lv-Trpc5_5 |
| e2 | SPU_001027 | Lv-Clect/Egf/Tm_1 |
| e2 | SPU_001056 | Lv-Gal/Lect/Lnb/7Tm/Gpcr |
| e2 | SPU_001115 | Lv-Hypp_1307 |
| e2 | SPU_001162 | Lv-Galr2L |

| | | |
|---|---|---|
| e2 | SPU_001380 | NA |
| e2 | SPU_001702 | Lv-Syt17 |
| e2 | SPU_001809 | none |
| e2 | SPU_002077 | Lv-Mib2-4 |
| e2 | SPU_002369 | NA |
| e2 | SPU_002470 | Lv-Fn3f_19 |
| e2 | SPU_002527 | Lv-Hypp_1401 |
| e2 | SPU_002661 | Lv-Slc17a5_1 |
| e2 | SPU_002963 | Lv-Fmo2 |
| e2 | SPU_003021 | Lv-Nt5e |
| e2 | SPU_003141 | NA |
| e2 | SPU_003342 | Lv-Hypp_1445 |
| e2 | SPU_003452 | Lv-Hypp_384 |
| e2 | SPU_003492 | Lv-Rxfp2L_3 |
| e2 | SPU_003639 | none |
| e2 | SPU_003688 | Lv-Hypp_1473 |
| e2 | SPU_003908 | NA |
| e2 | SPU_004101 | Lv-Srcr36 |
| e2 | SPU_004184 | NA |
| e2 | SPU_004364 | Lv-Hypp_1516 |
| e2 | SPU_004366 | Lv-Hbtf |
| e2 | SPU_004616 | none |
| e2 | SPU_004632 | Lv-Ccnb1ip1 |
| e2 | SPU_004670 | NA |
| e2 | SPU_004720 | Lv-Srr_1 |
| e2 | SPU_004891 | none |
| e2 | SPU_005120 | Lv-Hypp_438 |
| e2 | SPU_005126 | none |
| e2 | SPU_005345 | Lv-Ugt_4 |
| e2 | SPU_005366 | none |
| e2 | SPU_005472 | Lv-Nkx2.5 |
| e2 | SPU_005631 | Lv-Tmem20_3 |
| e2 | SPU_005639 | none |
| e2 | SPU_005781 | Lv-Ty/Wap/Vwc |
| e2 | SPU_005870 | NA |
| e2 | SPU_005967 | Lv-Htr1b |
| e2 | SPU_006014 | Lv-Mmp13L |
| e2 | SPU_006217 | Lv-Accn2 |

| | | |
|---|---|---|
| e2 | SPU_006297 | Lv-Gdpd1 |
| e2 | SPU_006298 | Lv-Suox2 |
| e2 | SPU_006409 | Lv-Hypp_1661 |
| e2 | SPU_006532 | Lv-TropT |
| e2 | SPU_006594 | Lv-Htr4L_3 |
| e2 | SPU_006719 | none |
| e2 | SPU_006996 | NA |
| e2 | SPU_007076 | Lv-Hypp_1695 |
| e2 | SPU_007102 | none |
| e2 | SPU_007104 | none |
| e2 | SPU_007131 | none |
| e2 | SPU_007290 | NA |
| e2 | SPU_007320 | NA |
| e2 | SPU_007534 | Lv-Ltbp4L_1 |
| e2 | SPU_007713 | none |
| e2 | SPU_007806 | Lv-Htr1L_1 |
| e2 | SPU_007856 | none |
| e2 | SPU_008096 | Lv-Siatf_1 |
| e2 | SPU_008303 | NA |
| e2 | SPU_008357 | none |
| e2 | SPU_008467 | Lv-Cngb1 |
| e2 | SPU_008542 | Lv-Slc5a3 |
| e2 | SPU_008806 | NA |
| e2 | SPU_008859 | Lv-Timp2 |
| e2 | SPU_008972 | NA |
| e2 | SPU_009038 | Lv-Galr2L_14 |
| e2 | SPU_009040 | none |
| e2 | SPU_009098 | Lv-Xkr6L |
| e2 | SPU_009149 | NA |
| e2 | SPU_009316 | none |
| e2 | SPU_009512 | Lv-Cyp2L18 |
| e2 | SPU_009597 | Lv-CckbrL_1 |
| e2 | SPU_009617 | Lv-Fmo2_1 |
| e2 | SPU_009755 | NA |
| e2 | SPU_009935 | Lv-Hypp_1882 |
| e2 | SPU_010018 | Lv-Hypp_602 |
| e2 | SPU_010093 | Lv-Ldlaf |
| e2 | SPU_010122 | Lv-Hypp_1899 |

| | | |
|---|---|---|
| e2 | SPU_010127 | NA |
| e2 | SPU_010133 | none |
| e2 | SPU_010154 | NA |
| e2 | SPU_010205 | Lv-Tmprss4 |
| e2 | SPU_010250 | none |
| e2 | SPU_010265 | Lv-Accn2_1 |
| e2 | SPU_010291 | Lv-Fn3/Igf_36 |
| e2 | SPU_010413 | NA |
| e2 | SPU_010432 | none |
| e2 | SPU_010721 | Lv-Cyp1L5 |
| e2 | SPU_010870 | Lv-Gpr64L_12 |
| e2 | SPU_010907 | NA |
| e2 | SPU_010968 | Lv-Egf |
| e2 | SPU_011026 | Lv-C10orf112L_1 |
| e2 | SPU_011132 | Lv-Hypp_1962 |
| e2 | SPU_011360 | Lv-NeurmdnBr_2 |
| e2 | SPU_011393 | Lv-Hrh3L_8 |
| e2 | SPU_011402 | Lv-Gpr133L_1 |
| e2 | SPU_011589 | Lv-Npr1_1 |
| e2 | SPU_011672 | Lv-Adra1bL_1 |
| e2 | SPU_011813 | NA |
| e2 | SPU_011898 | Lv-Unc44L_77 |
| e2 | SPU_012011 | Lv-Echn31 |
| e2 | SPU_012143 | NA |
| e2 | SPU_012268 | Lv-Hypp_678 |
| e2 | SPU_012285 | Lv-Hypp_680 |
| e2 | SPU_012352 | Lv-Igf_6 |
| e2 | SPU_012689 | Lv-Adra1aL_1 |
| e2 | SPU_012794 | none |
| e2 | SPU_012944 | none |
| e2 | SPU_012976 | Lv-NatL |
| e2 | SPU_013246 | Lv-Hypp_3058 |
| e2 | SPU_013308 | Lv-Chdh-2 |
| e2 | SPU_013631 | Lv-Hypp_725 |
| e2 | SPU_013964 | NA |
| e2 | SPU_014178 | none |
| e2 | SPU_014240 | Lv-Lrr/Igr_16 |
| e2 | SPU_014380 | Lv-Tacr2L_3 |

| | | |
|---|---|---|
| e2 | SPU_014389 | Lv-Hypp_754 |
| e2 | SPU_014457 | none |
| e2 | SPU_014514 | Lv-Adra1aL_4 |
| e2 | SPU_014528 | Lv-Giant |
| e2 | SPU_014764 | Lv-Gpr21L_10 |
| e2 | SPU_014853 | Lv-FcgbpL_1 |
| e2 | SPU_014864 | Lv-Hsp703A |
| e2 | SPU_014874 | Lv-Cyp3L9 |
| e2 | SPU_014890 | NA |
| e2 | SPU_014935 | Lv-Cpa2L_3 |
| e2 | SPU_014939 | NA |
| e2 | SPU_015036 | Lv-Sacs-2 |
| e2 | SPU_015139 | Lv-Tcknr1 |
| e2 | SPU_015233 | Lv-Cntnap2L |
| e2 | SPU_015273 | Lv-Hypp_777 |
| e2 | SPU_015470 | Lv-Catsper3L |
| e2 | SPU_015620 | none |
| e2 | SPU_015639 | Lv-Chrna6_1 |
| e2 | SPU_015667 | Lv-NmbrL_2 |
| e2 | SPU_015672 | none |
| e2 | SPU_015790 | none |
| e2 | SPU_015796 | Lv-Hypp_2232 |
| e2 | SPU_016076 | Lv-Accn1_2 |
| e2 | SPU_016185 | Lv-Slc22a5_1 |
| e2 | SPU_016270 | NA |
| e2 | SPU_016354 | Lv-Gluld1_17 |
| e2 | SPU_016455 | Lv-Tlk1c |
| e2 | SPU_016497 | Lv-Hypp_827 |
| e2 | SPU_016542 | Lv-Drel |
| e2 | SPU_016543 | Lv-Orct_2 |
| e2 | SPU_016679 | Lv-Slc13a5 |
| e2 | SPU_016698 | Lv-Syt15-1_1 |
| e2 | SPU_016923 | Lv-Hypp_2289 |
| e2 | SPU_017097 | NA |
| e2 | SPU_017189 | none |
| e2 | SPU_017324 | NA |
| e2 | SPU_017351 | none |
| e2 | SPU_017580 | Lv-Unc44_126 |

| | | |
|---|---|---|
| e2 | SPU_017663 | none |
| e2 | SPU_017689 | Lv-Npy1rL |
| e2 | SPU_017762 | Lv-Fn3_34 |
| e2 | SPU_017822 | Lv-Unc44_128 |
| e2 | SPU_017929 | Lv-Fn3f_41 |
| e2 | SPU_018013 | none |
| e2 | SPU_018141 | Lv-Amt5 |
| e2 | SPU_018263 | Lv-Hypp_2377 |
| e2 | SPU_018314 | Lv-Cctn/Cctngrpr |
| e2 | SPU_018737 | NA |
| e2 | SPU_018804 | Lv-Klhl12 |
| e2 | SPU_018870 | NA |
| e2 | SPU_018963 | Lv-NmbrL_4 |
| e2 | SPU_019094 | Lv-Hao1_18 |
| e2 | SPU_019189 | Lv-Triol1 |
| e2 | SPU_019190 | NA |
| e2 | SPU_019252 | NA |
| e2 | SPU_019264 | none |
| e2 | SPU_019455 | Lv-Hsf5L |
| e2 | SPU_019716 | Lv-Hypp_943 |
| e2 | SPU_019760 | NA |
| e2 | SPU_019795 | NA |
| e2 | SPU_019796 | Lv-Hypp_2465 |
| e2 | SPU_019909 | none |
| e2 | SPU_019995 | Lv-Syt5 |
| e2 | SPU_020436 | NA |
| e2 | SPU_020447 | NA |
| e2 | SPU_020461 | NA |
| e2 | SPU_020549 | Lv-Ctbs |
| e2 | SPU_020763 | Lv-Hypp_989 |
| e2 | SPU_020828 | Lv-Znfx1-13 |
| e2 | SPU_021063 | Lv-Slc1 |
| e2 | SPU_021124 | Lv-Srcr163 |
| e2 | SPU_021136 | none |
| e2 | SPU_021291 | Lv-Ccap/NpSLr |
| e2 | SPU_021329 | NA |
| e2 | SPU_021455 | Lv-Slc5a9_6 |
| e2 | SPU_021579 | Lv-Adrb2L_14 |

| | | |
|---|---|---|
| e2 | SPU_021657 | Lv-Bhmt_6 |
| e2 | SPU_021788 | Lv-Slc5a12_3 |
| e2 | SPU_021905 | Lv-Hypp_2587 |
| e2 | SPU_021919 | Lv-Sea/Ccp |
| e2 | SPU_021921 | NA |
| e2 | SPU_022093 | Lv-Ldla/Egf/Ig/Lnb/7Tm/Gpcr |
| e2 | SPU_022209 | Lv-Syt15-2a |
| e2 | SPU_022252 | Lv-Slc17a6 |
| e2 | SPU_022320 | Lv-Ldla/Egf/Ig/Gpcr_1 |
| e2 | SPU_022396 | Lv-Clect_42 |
| e2 | SPU_022408 | Lv-Ache-11 |
| e2 | SPU_022716 | Lv-Naalad2L_2 |
| e2 | SPU_022772 | Lv-Adra1aL_10 |
| e2 | SPU_022795 | none |
| e2 | SPU_022862 | Lv-Hypp_2635 |
| e2 | SPU_022933 | Lv-Hypp_1062 |
| e2 | SPU_022981 | Lv-Hypp_1063 |
| e2 | SPU_022989 | Lv-Gabbr1_7 |
| e2 | SPU_023257 | none |
| e2 | SPU_023299 | NA |
| e2 | SPU_023578 | none |
| e2 | SPU_023615 | NA |
| e2 | SPU_023706 | Lv-GramarL1 |
| e2 | SPU_023813 | Lv-Nec2 |
| e2 | SPU_024119 | none |
| e2 | SPU_024218 | Lv-Clect_92 |
| e2 | SPU_024304 | Lv-Scai |
| e2 | SPU_024318 | Lv-Hypp_2724 |
| e2 | SPU_024460 | Lv-Pnpla4L |
| e2 | SPU_024779 | Lv-Hypp_1127 |
| e2 | SPU_024812 | Lv-Surreal/Gpcr135 |
| e2 | SPU_024817 | Lv-Surreal/Gpcr142 |
| e2 | SPU_024824 | NA |
| e2 | SPU_024883 | Lv-Cret_1 |
| e2 | SPU_024905 | Lv-Slsp_20 |
| e2 | SPU_024918 | Lv-NeuroD1 |
| e2 | SPU_024935 | Lv-Hypp_2758 |
| e2 | SPU_024971 | Lv-Hypp_2761 |

| | | |
|---|---|---|
| e2 | SPU_024988 | Lv-Kiaa0840L |
| e2 | SPU_025073 | none |
| e2 | SPU_025268 | Lv-PppL_224 |
| e2 | SPU_025280 | NA |
| e2 | SPU_025341 | none |
| e2 | SPU_025932 | Lv-Npr1L |
| e2 | SPU_026002 | NA |
| e2 | SPU_026016 | none |
| e2 | SPU_026042 | NA |
| e2 | SPU_026391 | Lv-Hypp_2836 |
| e2 | SPU_026417 | Lv-Cubnr_3 |
| e2 | SPU_026543 | Lv-Notch1-20 |
| e2 | SPU_026658 | NA |
| e2 | SPU_026736 | none |
| e2 | SPU_027010 | NA |
| e2 | SPU_027290 | Lv-Ptpase |
| e2 | SPU_027605 | Lv-Hypp_271 |
| e2 | SPU_027700 | Lv-Gpr61 |
| e2 | SPU_027807 | Lv-PppL_235 |
| e2 | SPU_028012 | Lv-Mib2_10 |
| e2 | SPU_028097 | Lv-Slc5a9_10 |
| e2 | SPU_028132 | NA |
| e2 | SPU_028158 | Lv-Aldh9aLf |
| e2 | SPU_028188 | Lv-Fb |
| e2 | SPU_028198 | none |
| e2 | SPU_028290 | Lv-Slc22a9_1 |
| e2 | SPU_028298 | Lv-Clect_114 |
| e2 | SPU_028634 | NA |
| e2 | SPU_028694 | NA |
| e2 | SPU_028762 | Lv-TpaseL_22 |
| e2 | SPU_028895 | Lv-Loxhd1_2 |
| e3 | SPU_000056 | Lv-C10orf93L |
| e3 | SPU_000064 | Lv-Mus_achlrM3 |
| e3 | SPU_000148 | Lv-Epdmnrp2 |
| e3 | SPU_000152 | none |
| e3 | SPU_000310 | Lv-Pdgfr/vegfrL |
| e3 | SPU_000465 | NA |
| e3 | SPU_000635 | Lv-GagpolL_3 |

| | | |
|---|---|---|
| e3 | SPU_000756 | Lv-Nk2-2 |
| e3 | SPU_000757 | Lv-Nkx2.1 |
| e3 | SPU_000834 | NA |
| e3 | SPU_000896 | NA |
| e3 | SPU_000910 | Lv-Slc16_1 |
| e3 | SPU_000918 | Lv-Egfi2 |
| e3 | SPU_001153 | none |
| e3 | SPU_001218 | none |
| e3 | SPU_001470 | NA |
| e3 | SPU_001556 | NA |
| e3 | SPU_001615 | NA |
| e3 | SPU_002081 | NA |
| e3 | SPU_002088 | Lv-Msp130 |
| e3 | SPU_002129 | Lv-Not |
| e3 | SPU_002169 | Lv-Sstr5 |
| e3 | SPU_002245 | Lv-Hypp_1381 |
| e3 | SPU_002298 | none |
| e3 | SPU_002324 | Lv-ElavL |
| e3 | SPU_002351 | Lv-Chrm1L_2 |
| e3 | SPU_002592 | Lv-Emx |
| e3 | SPU_002734 | none |
| e3 | SPU_002852 | Lv-Atf4L |
| e3 | SPU_002958 | none |
| e3 | SPU_003231 | Lv-Cyp17r1 |
| e3 | SPU_003345 | Lv-Dkk1 |
| e3 | SPU_003407 | Lv-Ascc3l1 |
| e3 | SPU_003419 | Lv-Tlr164 |
| e3 | SPU_003551 | none |
| e3 | SPU_003768 | Lv-3Apcol |
| e3 | SPU_003799 | NA |
| e3 | SPU_003814 | NA |
| e3 | SPU_003853 | none |
| e3 | SPU_004018 | Lv-Fn3/Egff_16 |
| e3 | SPU_004091 | none |
| e3 | SPU_004100 | NA |
| e3 | SPU_004136 | Lv-P19 |
| e3 | SPU_004292 | Lv-Rbl1 |
| e3 | SPU_004551 | Lv-FoxB |

| | | |
|---|---|---|
| e3 | SPU_004905 | NA |
| e3 | SPU_005072 | Lv-Hypp_1563 |
| e3 | SPU_005148 | Lv-Tlr169 |
| e3 | SPU_005590 | Lv-Slc23a1_1 |
| e3 | SPU_005605 | Lv-C12orf63L |
| e3 | SPU_005676 | NA |
| e3 | SPU_005702 | Lv-TransglutL |
| e3 | SPU_005816 | Lv-Hypp_1606 |
| e3 | SPU_005944 | Lv-Nola1 |
| e3 | SPU_005982 | Lv-PolppL_53 |
| e3 | SPU_006233 | Lv-Flii |
| e3 | SPU_006242 | Lv-Fgf9/16/20 |
| e3 | SPU_006248 | Lv-PppL_246 |
| e3 | SPU_006272 | NA |
| e3 | SPU_006340 | none |
| e3 | SPU_006421 | none |
| e3 | SPU_006462 | Lv-Gcm |
| e3 | SPU_006500 | Lv-Hypp_1664 |
| e3 | SPU_006527 | none |
| e3 | SPU_006586 | Lv-Hypp_485 |
| e3 | SPU_006702 | Lv-Kiaa0415L_1 |
| e3 | SPU_006753 | Lv-E2f3 |
| e3 | SPU_006860 | Lv-Rdh12_2 |
| e3 | SPU_006932 | Lv-Drb1 |
| e3 | SPU_006991 | Lv-Lim1 |
| e3 | SPU_007008 | Lv-Flilh |
| e3 | SPU_007028 | Lv-Trim2-3 |
| e3 | SPU_007165 | Lv-Abcc5E |
| e3 | SPU_007332 | Lv-Unc44L_58 |
| e3 | SPU_007434 | Lv-Disp1-3 |
| e3 | SPU_007638 | NA |
| e3 | SPU_007776 | Lv-Chrna7_1 |
| e3 | SPU_008112 | Lv-Phb1 |
| e3 | SPU_008297 | none |
| e3 | SPU_008342 | Lv-Hypp_1779 |
| e3 | SPU_008408 | Lv-Cecr5L |
| e3 | SPU_008632 | Lv-Cyp2U |
| e3 | SPU_008664 | Lv-Nav2 |

| | | |
|---|---|---|
| e3 | SPU_008666 | Lv-Pear1L |
| e3 | SPU_009011 | NA |
| e3 | SPU_009017 | Lv-Nlr10 |
| e3 | SPU_009052 | NA |
| e3 | SPU_009071 | Lv-Spef2 |
| e3 | SPU_009315 | Lv-Fam55c_1 |
| e3 | SPU_009550 | Lv-Hypp_581 |
| e3 | SPU_009606 | none |
| e3 | SPU_009668 | none |
| e3 | SPU_009676 | Lv-Srcr78 |
| e3 | SPU_009730 | none |
| e3 | SPU_009751 | none |
| e3 | SPU_009860 | NA |
| e3 | SPU_009911 | Lv-Lefty |
| e3 | SPU_009956 | none |
| e3 | SPU_009958 | Lv-Rt32 |
| e3 | SPU_010027 | Lv-Hypp_1893 |
| e3 | SPU_010041 | none |
| e3 | SPU_010064 | none |
| e3 | SPU_010234 | NA |
| e3 | SPU_010401 | Lv-Hypp_1916 |
| e3 | SPU_010594 | Lv-Hypp_1926 |
| e3 | SPU_010645 | Lv-B3gt1 |
| e3 | SPU_010705 | Lv-Ugt2c1 |
| e3 | SPU_010717 | Lv-Hypp_1936 |
| e3 | SPU_010764 | Lv-Ak7 |
| e3 | SPU_011064 | Lv-Nodal |
| e3 | SPU_011073 | Lv-Ttrspn_7 |
| e3 | SPU_011156 | Lv-TlrP07 |
| e3 | SPU_011252 | none |
| e3 | SPU_011339 | NA |
| e3 | SPU_011343 | none |
| e3 | SPU_011419 | NA |
| e3 | SPU_011420 | none |
| e3 | SPU_011555 | NA |
| e3 | SPU_011561 | none |
| e3 | SPU_011644 | none |
| e3 | SPU_011756 | Lv-Wnt1 |

| | | |
|---|---|---|
| e3 | SPU_011841 | NA |
| e3 | SPU_011876 | NA |
| e3 | SPU_012069 | Lv-Surreal/Gpcr107 |
| e3 | SPU_012253 | Lv-Eve |
| e3 | SPU_012377 | Lv-Mrpl35 |
| e3 | SPU_012403 | NA |
| e3 | SPU_012408 | NA |
| e3 | SPU_012426 | Lv-Ache_22 |
| e3 | SPU_013015 | Lv-Bra |
| e3 | SPU_013047 | Lv-Nkx3-2 |
| e3 | SPU_013086 | Lv-Gpc35_2 |
| e3 | SPU_013102 | Lv-Cyp3L6 |
| e3 | SPU_013202 | Lv-Ift172_1 |
| e3 | SPU_013355 | NA |
| e3 | SPU_013519 | NA |
| e3 | SPU_013552 | Lv-Hebp2L |
| e3 | SPU_013553 | Lv-Hypp_721 |
| e3 | SPU_013554 | Lv-Taar7dL |
| e3 | SPU_013623 | Lv-Adra1aL_3 |
| e3 | SPU_013666 | Lv-Hypp_727 |
| e3 | SPU_013670 | Lv-Mt1-4/MmpL6 |
| e3 | SPU_013758 | Lv-Hypp_2103 |
| e3 | SPU_013822 | Lv-Msp130r1 |
| e3 | SPU_014006 | NA |
| e3 | SPU_014289 | Lv-Rx |
| e3 | SPU_014367 | NA |
| e3 | SPU_014770 | NA |
| e3 | SPU_014919 | NA |
| e3 | SPU_014937 | Lv-Trpm3 |
| e3 | SPU_014994 | Lv-Srcr122 |
| e3 | SPU_015080 | Lv-Abcg12 |
| e3 | SPU_015146 | Lv-Hypp_107 |
| e3 | SPU_015175 | Lv-PppL_200 |
| e3 | SPU_015239 | Lv-Hypp_2201 |
| e3 | SPU_015243 | Lv-FoxN2/3 |
| e3 | SPU_015303 | Lv-Tlr082 |
| e3 | SPU_015374 | Lv-Id |
| e3 | SPU_015393 | Lv-Hypp_2210 |

| | | |
|---|---|---|
| e3 | SPU_015518 | NA |
| e3 | SPU_015612 | Lv-Lrr/Igr_7 |
| e3 | SPU_015659 | NA |
| e3 | SPU_015944 | Lv-Hypp_2241 |
| e3 | SPU_015958 | Lv-PppL_201 |
| e3 | SPU_016155 | NA |
| e3 | SPU_016174 | none |
| e3 | SPU_016213 | NA |
| e3 | SPU_016225 | NA |
| e3 | SPU_016371 | Lv-Clmdln |
| e3 | SPU_016397 | Lv-Hypp_2259 |
| e3 | SPU_016506 | Lv-Msp130r2 |
| e3 | SPU_016525 | Lv-Fibf |
| e3 | SPU_016556 | NA |
| e3 | SPU_016800 | NA |
| e3 | SPU_016952 | none |
| e3 | SPU_016970 | Lv-Rtp4L |
| e3 | SPU_017019 | Lv-Risl2 |
| e3 | SPU_017090 | Lv-Gulo_1 |
| e3 | SPU_017123 | NA |
| e3 | SPU_017130 | Lv-PppL_114 |
| e3 | SPU_017134 | none |
| e3 | SPU_017295 | Lv-Clect_35 |
| e3 | SPU_017511 | Lv-TgfbrtII |
| e3 | SPU_017518 | NA |
| e3 | SPU_017574 | Lv-Ajpx2 |
| e3 | SPU_017614 | NA |
| e3 | SPU_017636 | Lv-Hypp_886 |
| e3 | SPU_017655 | Lv-Gpr101L_2 |
| e3 | SPU_017697 | Lv-TpaseL_9 |
| e3 | SPU_017809 | NA |
| e3 | SPU_017812 | Lv-Hypp_2356 |
| e3 | SPU_017821 | Lv-RtL_70 |
| e3 | SPU_017848 | Lv-Z313_2 |
| e3 | SPU_017967 | Lv-Slc46a3_1 |
| e3 | SPU_017985 | none |
| e3 | SPU_018071 | NA |
| e3 | SPU_018096 | Lv-PppL_270 |

| | | |
|---|---|---|
| e3 | SPU_018127 | Lv-Slc10a2_4 |
| e3 | SPU_018147 | Lv-Slsp_5 |
| e3 | SPU_018179 | Lv-Abcc5l |
| e3 | SPU_018341 | NA |
| e3 | SPU_018452 | NA |
| e3 | SPU_018483 | Lv-Erg |
| e3 | SPU_018798 | Lv-NotchL_3 |
| e3 | SPU_019002 | Lv-FoxQ2_1 |
| e3 | SPU_019050 | none |
| e3 | SPU_019173 | Lv-Hypp_929 |
| e3 | SPU_019319 | Lv-Hypp_2431 |
| e3 | SPU_019344 | Lv-VwdeL_1 |
| e3 | SPU_019370 | Lv-Srcr152 |
| e3 | SPU_019420 | Lv-Pres_1 |
| e3 | SPU_019457 | Lv-Hypp_2443 |
| e3 | SPU_019536 | Lv-Ugt8_2 |
| e3 | SPU_020008 | Lv-Sarmr2 |
| e3 | SPU_020023 | Lv-Icdc |
| e3 | SPU_020071 | Lv-Hypp_2481 |
| e3 | SPU_020178 | Lv-Unc44_327 |
| e3 | SPU_020228 | NA |
| e3 | SPU_020311 | Lv-Klf2/4 |
| e3 | SPU_020371 | Lv-Wnt8 |
| e3 | SPU_020451 | Lv-Bra_1 |
| e3 | SPU_020471 | Lv-Olah |
| e3 | SPU_020612 | Lv-Megf10 |
| e3 | SPU_020613 | Lv-Hypp_2515 |
| e3 | SPU_020680 | Lv-FgfrL |
| e3 | SPU_020695 | Lv-Morn3 |
| e3 | SPU_020849 | Lv-Abcg11 |
| e3 | SPU_021058 | none |
| e3 | SPU_021147 | none |
| e3 | SPU_021218 | NA |
| e3 | SPU_021265 | NA |
| e3 | SPU_021268 | NA |
| e3 | SPU_021287 | NA |
| e3 | SPU_021299 | Lv-TlrP24 |
| e3 | SPU_021362 | NA |

| | | |
|---|---|---|
| e3 | SPU_021364 | Lv-Amy_7 |
| e3 | SPU_021385 | Lv-ApL |
| e3 | SPU_021433 | Lv-Rhogap5 |
| e3 | SPU_021626 | Lv-Hypp_1027 |
| e3 | SPU_021694 | NA |
| e3 | SPU_021947 | Lv-Z474 |
| e3 | SPU_021958 | Lv-Hypp_177 |
| e3 | SPU_021978 | NA |
| e3 | SPU_022024 | Lv-Z410 |
| e3 | SPU_022168 | NA |
| e3 | SPU_022182 | Lv-Efcab6-2 |
| e3 | SPU_022364 | Lv-Amy2-2 |
| e3 | SPU_022462 | none |
| e3 | SPU_022472 | none |
| e3 | SPU_022541 | Lv-Samd11 |
| e3 | SPU_022543 | Lv-Hypp_2615 |
| e3 | SPU_022567 | Lv-Srcr194 |
| e3 | SPU_022570 | Lv-Casc1 |
| e3 | SPU_022598 | Lv-FReD |
| e3 | SPU_022817 | Lv-Alx1_1 |
| e3 | SPU_022883 | NA |
| e3 | SPU_022995 | Lv-Hypp_2641 |
| e3 | SPU_023045 | Lv-Hypp_1067 |
| e3 | SPU_023111 | NA |
| e3 | SPU_023115 | Lv-Ptpriz |
| e3 | SPU_023275 | none |
| e3 | SPU_023294 | NA |
| e3 | SPU_023317 | NA |
| e3 | SPU_023386 | Lv-Tbx2-3 |
| e3 | SPU_023390 | NA |
| e3 | SPU_023462 | Lv-Rabrp3 |
| e3 | SPU_023503 | NA |
| e3 | SPU_023577 | Lv-Lnb/7Tm/GpcrD |
| e3 | SPU_023759 | Lv-Nlr188 |
| e3 | SPU_023910 | Lv-Hypp_1098 |
| e3 | SPU_023949 | none |
| e3 | SPU_023964 | Lv-Spsb3 |
| e3 | SPU_023993 | Lv-Unk_31 |

| | | |
|---|---|---|
| e3 | SPU_024189 | Lv-Sp5 |
| e3 | SPU_024285 | Lv-Hypp_213 |
| e3 | SPU_024370 | Lv-Hypp_2725 |
| e3 | SPU_024479 | NA |
| e3 | SPU_024487 | Lv-Srcr206 |
| e3 | SPU_024828 | none |
| e3 | SPU_024921 | Lv-GagpolL_10 |
| e3 | SPU_024961 | Lv-Hypp_2760 |
| e3 | SPU_024991 | none |
| e3 | SPU_025116 | Lv-Endrvt113 |
| e3 | SPU_025125 | Lv-Z147 |
| e3 | SPU_025195 | NA |
| e3 | SPU_025271 | Lv-GagpL_6 |
| e3 | SPU_025358 | none |
| e3 | SPU_025479 | NA |
| e3 | SPU_025638 | none |
| e3 | SPU_026075 | none |
| e3 | SPU_026076 | Lv-A2bp1 |
| e3 | SPU_026081 | NA |
| e3 | SPU_026162 | none |
| e3 | SPU_026423 | NA |
| e3 | SPU_026623 | Lv-Hypp_258 |
| e3 | SPU_026767 | Lv-Abcg10 |
| e3 | SPU_026847 | Lv-PppL_151 |
| e3 | SPU_026869 | NA |
| e3 | SPU_026922 | Lv-Chst11_14 |
| e3 | SPU_027015 | Lv-GataC |
| e3 | SPU_027021 | Lv-Hypp_2880 |
| e3 | SPU_027037 | Lv-Srcr217 |
| e3 | SPU_027104 | none |
| e3 | SPU_027284 | none |
| e3 | SPU_027349 | none |
| e3 | SPU_027696 | Lv-Hypp_273 |
| e3 | SPU_028093 | Lv-Scl |
| e3 | SPU_028113 | NA |
| e3 | SPU_028230 | NA |
| e3 | SPU_028299 | NA |
| e3 | SPU_028395 | Lv-Pks2 |

| | | |
|---|---|---|
| e3 | SPU_028428 | Lv-Sec14L_3 |
| e3 | SPU_028431 | Lv-Sec14L2_3 |
| e3 | SPU_028572 | NA |
| e3 | SPU_028869 | Lv-Gagpol10 |
| e3 | SPU_030006 | Lv-Mt4/MmpL |
| e3 | SPU_030148 | Lv-Vegf3 |
| e3 | SPU_030197 | Lv-Il17-19 |
| e4 | SPU_000007 | Lv-Wnt7 |
| e4 | SPU_000053 | Lv-Egf/HyalinL5 |
| e4 | SPU_000164 | Lv-Clect |
| e4 | SPU_000175 | NA |
| e4 | SPU_000279 | Lv-Cyp17a1 |
| e4 | SPU_000308 | Lv-Rhogap24l2 |
| e4 | SPU_000335 | Lv-Hypp_1249 |
| e4 | SPU_000379 | Lv-Rpl23a |
| e4 | SPU_000395 | Lv-Rpl6 |
| e4 | SPU_000399 | none |
| e4 | SPU_000593 | NA |
| e4 | SPU_000672 | Lv-Nlr41 |
| e4 | SPU_000693 | NA |
| e4 | SPU_000737 | Lv-Tmem |
| e4 | SPU_000789 | NA |
| e4 | SPU_000826 | Lv-Sm30b |
| e4 | SPU_000829 | NA |
| e4 | SPU_000975 | Lv-FoxF |
| e4 | SPU_001309 | Lv-Kpna2 |
| e4 | SPU_001414 | Lv-PolppL_12 |
| e4 | SPU_001438 | Lv-CckbrL |
| e4 | SPU_001521 | none |
| e4 | SPU_001628 | Lv-Wnt3 |
| e4 | SPU_001636 | NA |
| e4 | SPU_001657 | Lv-Cebpa |
| e4 | SPU_001673 | Lv-Dcdsh |
| e4 | SPU_001753 | Lv-Mamdc4-4 |
| e4 | SPU_001838 | Lv-MtoL1 |
| e4 | SPU_002092 | Lv-P35L |
| e4 | SPU_002157 | none |
| e4 | SPU_002202 | NA |

| | | |
|---|---|---|
| e4 | SPU_002232 | Lv-Nid2L |
| e4 | SPU_002460 | NA |
| e4 | SPU_002464 | Lv-Rpl22 |
| e4 | SPU_002633 | Lv-Hox9/10 |
| e4 | SPU_002743 | Lv-Gluld1_1 |
| e4 | SPU_002745 | Lv-Hypp_369 |
| e4 | SPU_002880 | Lv-Rpl21 |
| e4 | SPU_002893 | none |
| e4 | SPU_002896 | none |
| e4 | SPU_002928 | Lv-Tpte |
| e4 | SPU_002969 | Lv-Selenbp1_1 |
| e4 | SPU_003136 | Lv-Rerg |
| e4 | SPU_003309 | Lv-Lrrtm2L |
| e4 | SPU_003623 | NA |
| e4 | SPU_003698 | none |
| e4 | SPU_003725 | Lv-Tph |
| e4 | SPU_003843 | none |
| e4 | SPU_003886 | Lv-Trpc5_6 |
| e4 | SPU_003905 | Lv-Rpl9 |
| e4 | SPU_003934 | NA |
| e4 | SPU_003947 | Lv-Hypp_1487 |
| e4 | SPU_003985 | Lv-Gelsolin |
| e4 | SPU_004043 | Lv-Nlr30 |
| e4 | SPU_004059 | NA |
| e4 | SPU_004160 | Lv-Srcr37 |
| e4 | SPU_004282 | Lv-Fn3/Igf_3 |
| e4 | SPU_004377 | NA |
| e4 | SPU_004622 | NA |
| e4 | SPU_004681 | Lv-Gpr64-2 |
| e4 | SPU_004748 | NA |
| e4 | SPU_004815 | Lv-Slc43a3_1 |
| e4 | SPU_004876 | Lv-Fth1 |
| e4 | SPU_004952 | none |
| e4 | SPU_004996 | Lv-Rps15a |
| e4 | SPU_005152 | Lv-Sat2 |
| e4 | SPU_005233 | Lv-PppL_243 |
| e4 | SPU_005242 | NA |
| e4 | SPU_005305 | NA |

| | | |
|---|---|---|
| e4 | SPU_005498 | Lv-Surreal/Gpcr104 |
| e4 | SPU_005571 | Lv-Rpl29 |
| e4 | SPU_005699 | Lv-Rps17 |
| e4 | SPU_005700 | Lv-Rps17_1 |
| e4 | SPU_005750 | NA |
| e4 | SPU_005775 | Lv-Hypp_464 |
| e4 | SPU_005825 | Lv-Rps2_1 |
| e4 | SPU_005884 | Lv-Rack |
| e4 | SPU_005983 | Lv-Il17-1 |
| e4 | SPU_005990 | Lv-Sm29 |
| e4 | SPU_006034 | Lv-Rps27 |
| e4 | SPU_006077 | NA |
| e4 | SPU_006188 | NA |
| e4 | SPU_006213 | Lv-Rpl13a |
| e4 | SPU_006415 | NA |
| e4 | SPU_006458 | Lv-Tlr033 |
| e4 | SPU_006459 | Lv-Rps18 |
| e4 | SPU_006487 | Lv-Rps19 |
| e4 | SPU_006570 | none |
| e4 | SPU_006605 | Lv-Rps9_1 |
| e4 | SPU_006622 | Lv-Unk_69 |
| e4 | SPU_006652 | Lv-Asp_1 |
| e4 | SPU_006677 | none |
| e4 | SPU_006764 | none |
| e4 | SPU_006777 | Lv-Hypp_1680 |
| e4 | SPU_006898 | Lv-Hypp_1687 |
| e4 | SPU_006917 | Lv-Runt1 |
| e4 | SPU_006968 | Lv-Ars_2 |
| e4 | SPU_007101 | none |
| e4 | SPU_007115 | Lv-Hypp_20 |
| e4 | SPU_007239 | none |
| e4 | SPU_007246 | Lv-Kcnk5 |
| e4 | SPU_007373 | none |
| e4 | SPU_007506 | none |
| e4 | SPU_007521 | Lv-Armc4-2 |
| e4 | SPU_007593 | none |
| e4 | SPU_007752 | Lv-Unc44_283 |
| e4 | SPU_007822 | Lv-Bmp3b |

| | | |
|---|---|---|
| e4 | SPU_007882 | Lv-C-lectin |
| e4 | SPU_007936 | none |
| e4 | SPU_008108 | none |
| e4 | SPU_008211 | Lv-Rps6 |
| e4 | SPU_008394 | Lv-Rps9 |
| e4 | SPU_008456 | Lv-Tlr047 |
| e4 | SPU_008513 | Lv-Gpr133L |
| e4 | SPU_008558 | Lv-Rpl37a |
| e4 | SPU_008650 | Lv-Rps24 |
| e4 | SPU_008751 | none |
| e4 | SPU_008776 | Lv-Ndufb10 |
| e4 | SPU_008869 | Lv-Gabbr1_2 |
| e4 | SPU_008920 | Lv-Hypp_1813 |
| e4 | SPU_009076 | Lv-6Afcol |
| e4 | SPU_009183 | Lv-Hypp_1830 |
| e4 | SPU_009406 | Lv-Prokr2 |
| e4 | SPU_009587 | Lv-Taar4L |
| e4 | SPU_009724 | none |
| e4 | SPU_009771 | Lv-FoxG |
| e4 | SPU_009786 | NA |
| e4 | SPU_009925 | Lv-Mmp18/19L4 |
| e4 | SPU_010042 | Lv-Gpr112_1 |
| e4 | SPU_010062 | Lv-Srcr83 |
| e4 | SPU_010279 | none |
| e4 | SPU_010405 | Lv-Znf503 |
| e4 | SPU_010479 | Lv-Hibch_1 |
| e4 | SPU_010581 | Lv-Ctsr |
| e4 | SPU_010626 | Lv-Rpl30 |
| e4 | SPU_010692 | Lv-Rpl8 |
| e4 | SPU_010916 | Lv-Rpl7 |
| e4 | SPU_011005 | Lv-AnkrpL |
| e4 | SPU_011015 | none |
| e4 | SPU_011038 | Lv-Endo16 |
| e4 | SPU_011039 | Lv-Gage1 |
| e4 | SPU_011075 | Lv-Rpl7a |
| e4 | SPU_011111 | none |
| e4 | SPU_011121 | Lv-Tmem55b |
| e4 | SPU_011122 | Lv-Pipox_2 |

| | | |
|---|---|---|
| e4 | SPU_011180 | Lv-Fn3/Egff_1 |
| e4 | SPU_011188 | none |
| e4 | SPU_011248 | none |
| e4 | SPU_011297 | Lv-Mbx1 |
| e4 | SPU_011352 | Lv-Rps3A |
| e4 | SPU_011572 | Lv-Rps20 |
| e4 | SPU_011573 | none |
| e4 | SPU_011692 | Lv-CcLta1 |
| e4 | SPU_011698 | Lv-Gpr21 |
| e4 | SPU_011703 | none |
| e4 | SPU_011817 | NA |
| e4 | SPU_011873 | NA |
| e4 | SPU_012050 | Lv-Z11 |
| e4 | SPU_012157 | Lv-St8sia3_1 |
| e4 | SPU_012238 | Lv-Hh |
| e4 | SPU_012296 | Lv-Ahrl |
| e4 | SPU_012552 | Lv-Notch2_11 |
| e4 | SPU_012574 | NA |
| e4 | SPU_012598 | NA |
| e4 | SPU_012865 | none |
| e4 | SPU_013417 | Lv-Echn32 |
| e4 | SPU_013474 | none |
| e4 | SPU_013570 | Lv-Wnt6 |
| e4 | SPU_013662 | Lv-Rps3 |
| e4 | SPU_013760 | Lv-Sstr2L_1 |
| e4 | SPU_013774 | Lv-D2rlL |
| e4 | SPU_013834 | Lv-Rps28 |
| e4 | SPU_013854 | none |
| e4 | SPU_013994 | NA |
| e4 | SPU_014016 | Lv-Mamdc2-2 |
| e4 | SPU_014074 | Lv-Lrp2 |
| e4 | SPU_014157 | Lv-Lmx1 |
| e4 | SPU_014176 | Lv-Gluld1_5 |
| e4 | SPU_014298 | none |
| e4 | SPU_014425 | Lv-NrplaL |
| e4 | SPU_014454 | Lv-Rpl17 |
| e4 | SPU_014756 | none |
| e4 | SPU_014911 | Lv-Ache_23 |

| | | |
|---|---|---|
| e4 | SPU_015056 | none |
| e4 | SPU_015105 | Lv-Nlr63 |
| e4 | SPU_015170 | Lv-Hypp_773 |
| e4 | SPU_015284 | NA |
| e4 | SPU_015294 | Lv-Rpl36a |
| e4 | SPU_015475 | Lv-C20orf54L_1 |
| e4 | SPU_015596 | Lv-AgrnL_2 |
| e4 | SPU_015650 | Lv-Kidins220L_2 |
| e4 | SPU_015666 | Lv-Rpl4 |
| e4 | SPU_015712 | Lv-Hey4 |
| e4 | SPU_015726 | Lv-Clect_78 |
| e4 | SPU_015740 | Lv-Lrrc/Gpcrlr |
| e4 | SPU_015818 | Lv-Rpl17_1 |
| e4 | SPU_015831 | Lv-B4galt5_5 |
| e4 | SPU_015838 | none |
| e4 | SPU_015870 | Lv-Rpip1 |
| e4 | SPU_015908 | Lv-Nfat |
| e4 | SPU_015946 | Lv-Slc7a6_6 |
| e4 | SPU_016011 | Lv-Glyt_6 |
| e4 | SPU_016052 | Lv-Vtgn2 |
| e4 | SPU_016057 | Lv-Abtb1_1 |
| e4 | SPU_016135 | Lv-Gal3st1_9 |
| e4 | SPU_016148 | NA |
| e4 | SPU_016152 | Lv-Cyp3L15 |
| e4 | SPU_016200 | Lv-Rps15 |
| e4 | SPU_016303 | NA |
| e4 | SPU_016365 | NA |
| e4 | SPU_016384 | Lv-Lrrtm2 |
| e4 | SPU_016481 | NA |
| e4 | SPU_016558 | Lv-Ghdc1_6 |
| e4 | SPU_016758 | Lv-Nme2 |
| e4 | SPU_016771 | Lv-Clect_109 |
| e4 | SPU_016794 | Lv-Nlr114 |
| e4 | SPU_016966 | Lv-Hypp_2294 |
| e4 | SPU_017026 | none |
| e4 | SPU_017131 | Lv-PppL_206 |
| e4 | SPU_017147 | NA |
| e4 | SPU_017296 | none |

| | | |
|---|---|---|
| e4 | SPU_017312 | NA |
| e4 | SPU_017519 | NA |
| e4 | SPU_017686 | Lv-Rpl10a |
| e4 | SPU_017688 | Lv-Galr2L_24 |
| e4 | SPU_017799 | Lv-Fut4_1 |
| e4 | SPU_017849 | Lv-Z313_3 |
| e4 | SPU_017850 | NA |
| e4 | SPU_017856 | Lv-Taar7bL_1 |
| e4 | SPU_017917 | none |
| e4 | SPU_017924 | Lv-Hypp_902 |
| e4 | SPU_018052 | none |
| e4 | SPU_018084 | Lv-Gnmt_6 |
| e4 | SPU_018178 | Lv-Hypp_2374 |
| e4 | SPU_018204 | Lv-Nacha3 |
| e4 | SPU_018318 | Lv-Rpl13 |
| e4 | SPU_018374 | none |
| e4 | SPU_018385 | none |
| e4 | SPU_018405 | Lv-Hypp_2386 |
| e4 | SPU_018407 | Lv-Hypp_2999 |
| e4 | SPU_018811 | Lv-Sm50 |
| e4 | SPU_018813 | Lv-Sm37 |
| e4 | SPU_018819 | none |
| e4 | SPU_018866 | Lv-Rps8 |
| e4 | SPU_018874 | Lv-Sult1b1 |
| e4 | SPU_018877 | none |
| e4 | SPU_018884 | Lv-Rpl39 |
| e4 | SPU_018937 | Lv-Hypp_2411 |
| e4 | SPU_018966 | Lv-Hrh3L_14 |
| e4 | SPU_019048 | Lv-Hypp_9417 |
| e4 | SPU_019089 | Lv-Z133 |
| e4 | SPU_019134 | Lv-Phgdh_2 |
| e4 | SPU_019137 | none |
| e4 | SPU_019250 | Lv-Hypp_933 |
| e4 | SPU_019352 | Lv-Hypp_2433 |
| e4 | SPU_019366 | Lv-BsxL |
| e4 | SPU_019379 | none |
| e4 | SPU_019504 | none |
| e4 | SPU_019735 | Lv-C6st |

| | | |
|---|---|---|
| e4 | SPU_019788 | Lv-Rpl27 |
| e4 | SPU_020081 | NA |
| e4 | SPU_020194 | none |
| e4 | SPU_020308 | Lv-Rps3A_1 |
| e4 | SPU_020324 | Lv-AmidL_3 |
| e4 | SPU_020431 | Lv-Unc44_328 |
| e4 | SPU_020565 | Lv-Msxl |
| e4 | SPU_020683 | Lv-Oach_2 |
| e4 | SPU_020694 | none |
| e4 | SPU_020733 | NA |
| e4 | SPU_020743 | NA |
| e4 | SPU_020745 | Lv-Rps12 |
| e4 | SPU_020805 | NA |
| e4 | SPU_020809 | Lv-Unc44_147 |
| e4 | SPU_020933 | Lv-Uchl3 |
| e4 | SPU_020993 | none |
| e4 | SPU_021064 | Lv-Tmem20L_2 |
| e4 | SPU_021065 | Lv-Tmem45bL |
| e4 | SPU_021093 | Lv-Hypp_2535 |
| e4 | SPU_021216 | NA |
| e4 | SPU_021393 | Lv-HyalinL_7 |
| e4 | SPU_021447 | Lv-Nlr54 |
| e4 | SPU_021453 | none |
| e4 | SPU_021669 | Lv-Atub2 |
| e4 | SPU_021739 | Lv-Arp |
| e4 | SPU_021832 | Lv-Lnb/Tm7_2 |
| e4 | SPU_021901 | none |
| e4 | SPU_022062 | none |
| e4 | SPU_022161 | Lv-Rps13 |
| e4 | SPU_022314 | Lv-Fn3f_43 |
| e4 | SPU_022427 | Lv-Ogfod2_1 |
| e4 | SPU_022607 | Lv-Rps4 |
| e4 | SPU_022616 | NA |
| e4 | SPU_022677 | Lv-InvsL |
| e4 | SPU_022726 | none |
| e4 | SPU_022796 | Lv-Slc5a7_6 |
| e4 | SPU_022868 | Lv-Jag2 |
| e4 | SPU_022966 | Lv-Gpr84L_5 |

| | | |
|---|---|---|
| e4 | SPU_023059 | Lv-Slc10a2_11 |
| e4 | SPU_023067 | Lv-Cyp2L53 |
| e4 | SPU_023071 | none |
| e4 | SPU_023136 | Lv-Upk_3 |
| e4 | SPU_023206 | NA |
| e4 | SPU_023236 | Lv-Unk_78 |
| e4 | SPU_023252 | Lv-Drd2_1 |
| e4 | SPU_023320 | Lv-Trpc5_2 |
| e4 | SPU_023337 | Lv-Hypp_2667 |
| e4 | SPU_023353 | Lv-Ars_1 |
| e4 | SPU_023356 | Lv-Hypp_1081 |
| e4 | SPU_023413 | none |
| e4 | SPU_023517 | Lv-Rps25 |
| e4 | SPU_023566 | Lv-Nckap1_1 |
| e4 | SPU_023569 | none |
| e4 | SPU_023692 | Lv-Rpl18a |
| e4 | SPU_023730 | Lv-Isl |
| e4 | SPU_023842 | NA |
| e4 | SPU_023860 | Lv-Kiaa1095 |
| e4 | SPU_024139 | Lv-FoxC |
| e4 | SPU_024141 | none |
| e4 | SPU_024163 | NA |
| e4 | SPU_024266 | Lv-Hypp_2720 |
| e4 | SPU_024354 | none |
| e4 | SPU_024413 | none |
| e4 | SPU_024499 | Lv-Dynlc2-5c |
| e4 | SPU_024596 | Lv-Dnahc8L_1 |
| e4 | SPU_024654 | NA |
| e4 | SPU_024683 | Lv-Hypp_1121 |
| e4 | SPU_024705 | none |
| e4 | SPU_024786 | NA |
| e4 | SPU_024811 | Lv-Surreal/Gpcr139 |
| e4 | SPU_024813 | Lv-Htr4L_9 |
| e4 | SPU_024899 | Lv-Endrvt111 |
| e4 | SPU_024936 | none |
| e4 | SPU_025547 | Lv-Slc5a11_2 |
| e4 | SPU_025612 | Lv-Runt1_1 |
| e4 | SPU_025709 | Lv-Cnga3_1 |

| | | |
|---|---|---|
| e4 | SPU_025859 | Lv-Fbxl16 |
| e4 | SPU_025982 | none |
| e4 | SPU_026099 | Lv-Lox |
| e4 | SPU_026108 | none |
| e4 | SPU_026112 | NA |
| e4 | SPU_026132 | NA |
| e4 | SPU_026175 | Lv-Hypp_2826 |
| e4 | SPU_026304 | Lv-Nlr180 |
| e4 | SPU_026347 | Lv-Unk_88 |
| e4 | SPU_026354 | Lv-Hypp_1175 |
| e4 | SPU_026440 | NA |
| e4 | SPU_026701 | NA |
| e4 | SPU_026855 | Lv-Allc_1 |
| e4 | SPU_026891 | Lv-Hypp_2872 |
| e4 | SPU_026909 | Lv-Dcst2-2 |
| e4 | SPU_027123 | NA |
| e4 | SPU_027134 | Lv-Hypp_2888 |
| e4 | SPU_027145 | Lv-Eph |
| e4 | SPU_027148 | Lv-Hypp_263 |
| e4 | SPU_027166 | NA |
| e4 | SPU_027215 | Lv-Hhex |
| e4 | SPU_027252 | Lv-Aqp8_1 |
| e4 | SPU_027253 | none |
| e4 | SPU_027434 | Lv-Chrna10 |
| e4 | SPU_027463 | Lv-Slc22a13L |
| e4 | SPU_027550 | none |
| e4 | SPU_027633 | Lv-Opsin3.2 |
| e4 | SPU_027634 | Lv-Opsin3.1 |
| e4 | SPU_027773 | NA |
| e4 | SPU_027795 | NA |
| e4 | SPU_027956 | Lv-Igf_8 |
| e4 | SPU_028014 | Lv-KyL_2 |
| e4 | SPU_028107 | Lv-Unc44_263 |
| e4 | SPU_028120 | Lv-Dmrt |
| e4 | SPU_028125 | Lv-Scnn1gL |
| e4 | SPU_028155 | Lv-Rpl28 |
| e4 | SPU_028243 | Lv-Hypp_2942 |
| e4 | SPU_028247 | Lv-Rpl36 |

| | | |
|---|---|---|
| e4 | SPU_028271 | none |
| e4 | SPU_028349 | Lv-PppL_165 |
| e4 | SPU_028366 | Lv-Cask_1 |
| e4 | SPU_028485 | Lv-Nlr182 |
| e4 | SPU_028528 | Lv-Lmbr2_2 |
| e4 | SPU_028530 | none |
| e4 | SPU_028681 | Lv-Nlr8 |
| e4 | SPU_028688 | none |
| e4 | SPU_028823 | Lv-Drd2L_27 |
| e4 | SPU_030147 | Lv-Pm27 |

| Table S6. k-means clusters for ubiquitously expressed transcripts | | |
|---|---|---|
| **CLUSTER** | **SPU_ID** | **NAME** |
| a1 | SPU_000016 | Lv-Unk_48 |
| a1 | SPU_000023 | Lv-Mical3l |
| a1 | SPU_000037 | Lv-Veph1_1 |
| a1 | SPU_000052 | Lv-Aytl2 |
| a1 | SPU_000098 | Lv-Clasp2 |
| a1 | SPU_000197 | Lv-Figl1L |
| a1 | SPU_000208 | Lv-Dspchd |
| a1 | SPU_000376 | Lv-Lrrc7 |
| a1 | SPU_000497 | Lv-Ppp2r4 |
| a1 | SPU_000531 | Lv-Ubr4 |
| a1 | SPU_000558 | none |
| a1 | SPU_000605 | Lv-Hypp_1265 |
| a1 | SPU_000638 | Lv-Zfand2b |
| a1 | SPU_000683 | Lv-Hypp_1273 |
| a1 | SPU_000709 | Lv-Hao1_1 |
| a1 | SPU_000736 | Lv-Fn3_11 |
| a1 | SPU_000783 | Lv-C17orf59L |
| a1 | SPU_000815 | Lv-Mib2L |
| a1 | SPU_000824 | Lv-Tstsk2 |
| a1 | SPU_000882 | Lv-Cspe9L1a |
| a1 | SPU_000897 | Lv-Pak7_1 |
| a1 | SPU_000969 | none |
| a1 | SPU_001071 | Lv-Ccdc46 |
| a1 | SPU_001128 | Lv-Nit2 |
| a1 | SPU_001228 | Lv-Cypf |
| a1 | SPU_001276 | Lv-Gpr112L_2 |
| a1 | SPU_001316 | Lv-Hs3st2L |
| a1 | SPU_001334 | Lv-Tmem194 |
| a1 | SPU_001337 | Lv-Mtrf1L |
| a1 | SPU_001354 | none |
| a1 | SPU_001359 | none |
| a1 | SPU_001388 | Lv-Atrx |
| a1 | SPU_001445 | Lv-Fam161aL |
| a1 | SPU_001468 | Lv-Hypp_1331 |
| a1 | SPU_001481 | Lv-Slc41a1 |

| | | |
|---|---|---|
| a1 | SPU_001483 | Lv-Z59 |
| a1 | SPU_001500 | Lv-Ck1d |
| a1 | SPU_001528 | Lv-Gtf3c2L |
| a1 | SPU_001530 | Lv-Sec16a |
| a1 | SPU_001577 | Lv-Ndufaf2 |
| a1 | SPU_001593 | NA |
| a1 | SPU_001622 | Lv-Cyp2L2 |
| a1 | SPU_001623 | Lv-Ube2f |
| a1 | SPU_001682 | Lv-Fn3f_61 |
| a1 | SPU_001712 | Lv-CycE |
| a1 | SPU_001715 | Lv-01715 |
| a1 | SPU_001771 | Lv-Nsun6 |
| a1 | SPU_001825 | none |
| a1 | SPU_001856 | none |
| a1 | SPU_001859 | Lv-Pura |
| a1 | SPU_001892 | Lv-Sema6d |
| a1 | SPU_001921 | NA |
| a1 | SPU_001939 | Lv-Syde1 |
| a1 | SPU_001940 | NA |
| a1 | SPU_001953 | Lv-TorlaipL |
| a1 | SPU_002025 | Lv-Agrin/Nt |
| a1 | SPU_002055 | none |
| a1 | SPU_002078 | Lv-Muc4L_3 |
| a1 | SPU_002096 | Lv-Hypp_342 |
| a1 | SPU_002116 | Lv-Hypp_3008 |
| a1 | SPU_002122 | Lv-Skap2 |
| a1 | SPU_002140 | Lv-Z475 |
| a1 | SPU_002179 | Lv-Cdc91L1 |
| a1 | SPU_002182 | Lv-Pdpr |
| a1 | SPU_002190 | none |
| a1 | SPU_002206 | NA |
| a1 | SPU_002216 | Lv-C9orf93L |
| a1 | SPU_002287 | NA |
| a1 | SPU_002291 | Lv-Mtmr14 |
| a1 | SPU_002303 | NA |
| a1 | SPU_002313 | Lv-Pip5k3_L2 |
| a1 | SPU_002346 | NA |
| a1 | SPU_002469 | Lv-Fn3f_62 |

| | | |
|---|---|---|
| a1 | SPU_002476 | NA |
| a1 | SPU_002509 | Lv-Fbw2 |
| a1 | SPU_002512 | Lv-AftphL |
| a1 | SPU_002523 | Lv-Ccdc85a |
| a1 | SPU_002529 | none |
| a1 | SPU_002557 | Lv-C6orf192L_1 |
| a1 | SPU_002558 | Lv-Slc17a5 |
| a1 | SPU_002615 | Lv-Furin |
| a1 | SPU_002679 | Lv-Hypp_365 |
| a1 | SPU_002691 | Lv-Tmco4 |
| a1 | SPU_002697 | Lv-Clect_6 |
| a1 | SPU_002798 | none |
| a1 | SPU_002860 | none |
| a1 | SPU_002988 | Lv-Z49 |
| a1 | SPU_003129 | none |
| a1 | SPU_003172 | Lv-Mospd2 |
| a1 | SPU_003308 | Lv-Dclr1 |
| a1 | SPU_003325 | Lv-Nmnat3 |
| a1 | SPU_003335 | Lv-Psme4-2 |
| a1 | SPU_003378 | Lv-Z108 |
| a1 | SPU_003398 | none |
| a1 | SPU_003416 | Lv-AtrnL1 |
| a1 | SPU_003482 | Lv-Hypp_1454 |
| a1 | SPU_003525 | Lv-Gal3st1L_4 |
| a1 | SPU_003592 | Lv-Orai2L |
| a1 | SPU_003619 | Lv-Nlr201 |
| a1 | SPU_003726 | Lv-FryhL |
| a1 | SPU_003773 | Lv-Z392 |
| a1 | SPU_003779 | Lv-Lrrc59 |
| a1 | SPU_003824 | Lv-Bcas3 |
| a1 | SPU_003868 | Lv-Taf4 |
| a1 | SPU_003893 | Lv-Hypp_3097 |
| a1 | SPU_003967 | Lv-CycC |
| a1 | SPU_004006 | Lv-Ttk1 |
| a1 | SPU_004030 | none |
| a1 | SPU_004056 | Lv-Trafd1 |
| a1 | SPU_004110 | Lv-Unk_3 |
| a1 | SPU_004120 | none |

| | | |
|---|---|---|
| a1 | SPU_004253 | Lv-Rnf34L |
| a1 | SPU_004289 | Lv-Slc35b4 |
| a1 | SPU_004317 | Lv-Swi/Snf |
| a1 | SPU_004363 | Lv-Sirt4 |
| a1 | SPU_004394 | NA |
| a1 | SPU_004396 | Lv-PolppL_37 |
| a1 | SPU_004401 | Lv-Pomt2 |
| a1 | SPU_004528 | Lv-Btbd7 |
| a1 | SPU_004564 | Lv-Alg8h |
| a1 | SPU_004651 | none |
| a1 | SPU_004778 | Lv-Gnn |
| a1 | SPU_004783 | Lv-Shq1h |
| a1 | SPU_004788 | Lv-Gapvd1_2 |
| a1 | SPU_004822 | Lv-Hypp_431 |
| a1 | SPU_004833 | NA |
| a1 | SPU_004883 | Lv-Perld1 |
| a1 | SPU_004898 | Lv-Cdc42se2 |
| a1 | SPU_004899 | none |
| a1 | SPU_004954 | Lv-Pp2arsBi2 |
| a1 | SPU_005073 | Lv-Pi3kcIII_1 |
| a1 | SPU_005079 | NA |
| a1 | SPU_005105 | Lv-Chibbyp |
| a1 | SPU_005117 | Lv-Ipo7 |
| a1 | SPU_005177 | Lv-B3galt1_3 |
| a1 | SPU_005257 | Lv-Ube2s |
| a1 | SPU_005263 | Lv-Hypp_1573 |
| a1 | SPU_005288 | Lv-Pcyox1 |
| a1 | SPU_005291 | none |
| a1 | SPU_005420 | Lv-Srcr42 |
| a1 | SPU_005494 | Lv-Hypp_1585 |
| a1 | SPU_005523 | Lv-St6galnac4 |
| a1 | SPU_005692 | Lv-Ankra2L |
| a1 | SPU_005701 | Lv-Dnajhc21 |
| a1 | SPU_005738 | none |
| a1 | SPU_005771 | Lv-Wbp4 |
| a1 | SPU_005891 | Lv-C4orf14L |
| a1 | SPU_005961 | Lv-Hook3 |
| a1 | SPU_005999 | Lv-Fras1 |

| | | |
|---|---|---|
| a1 | SPU_006060 | Lv-Hif1an |
| a1 | SPU_006138 | none |
| a1 | SPU_006148 | none |
| a1 | SPU_006170 | Lv-Wdr81 |
| a1 | SPU_006256 | Lv-Ppp1r3bL |
| a1 | SPU_006394 | Lv-Lycat_2 |
| a1 | SPU_006442 | Lv-Neu1_2 |
| a1 | SPU_006463 | Lv-Arfrp1 |
| a1 | SPU_006481 | Lv-Gcl |
| a1 | SPU_006547 | Lv-Copiner |
| a1 | SPU_006579 | Lv-Tbpaf_rna_polIB |
| a1 | SPU_006603 | Lv-Lipe |
| a1 | SPU_006673 | Lv-Ccdc138 |
| a1 | SPU_006679 | Lv-Nat11 |
| a1 | SPU_006689 | Lv-Rnbn1L |
| a1 | SPU_006711 | Lv-Vnn1 |
| a1 | SPU_006735 | Lv-Cmip |
| a1 | SPU_006758 | none |
| a1 | SPU_006769 | Lv-Sec24b_1 |
| a1 | SPU_006783 | none |
| a1 | SPU_006807 | none |
| a1 | SPU_006808 | Lv-Af9 |
| a1 | SPU_006838 | Lv-C14orf100L |
| a1 | SPU_006854 | Lv-PrccL_1 |
| a1 | SPU_006865 | Lv-Unk_50 |
| a1 | SPU_006903 | Lv-Slc35c1 |
| a1 | SPU_006912 | Lv-Presnln |
| a1 | SPU_006999 | Lv-Hax1L |
| a1 | SPU_007121 | Lv-Fxn |
| a1 | SPU_007195 | Lv-Slc25a40 |
| a1 | SPU_007212 | Lv-Hypp_1706 |
| a1 | SPU_007240 | Lv-Ged |
| a1 | SPU_007294 | Lv-Ugt1L_1 |
| a1 | SPU_007304 | Lv-Lrch4 |
| a1 | SPU_007311 | NA |
| a1 | SPU_007324 | Lv-Dnz11 |
| a1 | SPU_007433 | Lv-Depdc5-3 |
| a1 | SPU_007443 | Lv-Pitpnm2 |

| | | |
|---|---|---|
| a1 | SPU_007475 | NA |
| a1 | SPU_007497 | Lv-Rld2L2 |
| a1 | SPU_007510 | Lv-Fbx15 |
| a1 | SPU_007517 | none |
| a1 | SPU_007530 | Lv-Chk1 |
| a1 | SPU_007590 | none |
| a1 | SPU_007685 | none |
| a1 | SPU_007721 | Lv-Grb2 |
| a1 | SPU_007939 | Lv-Acy1l2L_4 |
| a1 | SPU_007975 | Lv-Aggf1_1 |
| a1 | SPU_008024 | Lv-Haspin |
| a1 | SPU_008091 | none |
| a1 | SPU_008244 | Lv-Senp1 |
| a1 | SPU_008281 | NA |
| a1 | SPU_008288 | Lv-Phkb |
| a1 | SPU_008343 | Lv-Svop |
| a1 | SPU_008469 | Lv-PpiL2 |
| a1 | SPU_008518 | Lv-Gcc1 |
| a1 | SPU_008663 | Lv-RoboL_1 |
| a1 | SPU_008764 | Lv-Opa1 |
| a1 | SPU_008775 | Lv-Uvrag |
| a1 | SPU_008879 | Lv-Tk1 |
| a1 | SPU_008907 | Lv-Gipc1 |
| a1 | SPU_008923 | Lv-Scyl3 |
| a1 | SPU_008966 | Lv-Sgk496 |
| a1 | SPU_008984 | Lv-Fpgs_2 |
| a1 | SPU_009066 | Lv-Mtap |
| a1 | SPU_009070 | none |
| a1 | SPU_009096 | Lv-Xkr6 |
| a1 | SPU_009126 | Lv-Usp16 |
| a1 | SPU_009167 | Lv-Med25 |
| a1 | SPU_009177 | none |
| a1 | SPU_009332 | Lv-Wdr91 |
| a1 | SPU_009434 | Lv-Ire1 |
| a1 | SPU_009441 | Lv-Kiaa0052_1 |
| a1 | SPU_009472 | Lv-Tmem39a |
| a1 | SPU_009473 | Lv-Capn7 |
| a1 | SPU_009507 | Lv-Wac |

| | | |
|---|---|---|
| a1 | SPU_009522 | Lv-Pten |
| a1 | SPU_009577 | Lv-Trim2L |
| a1 | SPU_009600 | Lv-PlexA2_1 |
| a1 | SPU_009647 | Lv-Sacm1L |
| a1 | SPU_009648 | Lv-Ranbp9L |
| a1 | SPU_009651 | NA |
| a1 | SPU_009663 | Lv-C20orf12L-2 |
| a1 | SPU_009703 | Lv-Mesdc1 |
| a1 | SPU_009781 | Lv-Nwd1_1 |
| a1 | SPU_009808 | Lv-Hypp_57 |
| a1 | SPU_009834 | none |
| a1 | SPU_009868 | Lv-Selo |
| a1 | SPU_009904 | Lv-Pdss2 |
| a1 | SPU_009913 | Lv-Ttf |
| a1 | SPU_009965 | Lv-Hypp_1884 |
| a1 | SPU_010115 | Lv-Sumf1 |
| a1 | SPU_010141 | Lv-Pak7 |
| a1 | SPU_010189 | Lv-En_IV |
| a1 | SPU_010245 | Lv-Socs7 |
| a1 | SPU_010293 | Lv-Slc25a29 |
| a1 | SPU_010348 | none |
| a1 | SPU_010361 | Lv-Hypp_1912 |
| a1 | SPU_010417 | Lv-Tarbp1 |
| a1 | SPU_010448 | Lv-Gpr133 |
| a1 | SPU_010449 | Lv-Gtf3c1L-2 |
| a1 | SPU_010458 | Lv-Z443 |
| a1 | SPU_010491 | none |
| a1 | SPU_010507 | none |
| a1 | SPU_010625 | Lv-Ext1 |
| a1 | SPU_010673 | none |
| a1 | SPU_010724 | Lv-Fastkd5L |
| a1 | SPU_010780 | none |
| a1 | SPU_010857 | Lv-Ncoa4 |
| a1 | SPU_011165 | Lv-Kctd21L_3 |
| a1 | SPU_011240 | Lv-Cxorf58 |
| a1 | SPU_011304 | Lv-Klhl29 |
| a1 | SPU_011310 | Lv-Ncapd2-2 |
| a1 | SPU_011326 | Lv-Drd1L_3 |

| | | |
|---|---|---|
| a1 | SPU_011358 | Lv-Elp3h |
| a1 | SPU_011403 | Lv-Hrh3L_9 |
| a1 | SPU_011466 | Lv-Sirt7 |
| a1 | SPU_011502 | Lv-Z437 |
| a1 | SPU_011604 | Lv-Glo1_1 |
| a1 | SPU_011614 | Lv-Samd9L_7 |
| a1 | SPU_011685 | Lv-Rictor |
| a1 | SPU_011710 | Lv-Mtpap |
| a1 | SPU_011716 | Lv-Klhl8 |
| a1 | SPU_011821 | Lv-Fbxw7L_1 |
| a1 | SPU_011835 | Lv-Klhl20 |
| a1 | SPU_011838 | Lv-Mast1 |
| a1 | SPU_011918 | Lv-Citph |
| a1 | SPU_011930 | NA |
| a1 | SPU_012017 | none |
| a1 | SPU_012025 | Lv-Dus1ll |
| a1 | SPU_012048 | Lv-Irak1bp1L |
| a1 | SPU_012073 | Lv-Ptp9Q22 |
| a1 | SPU_012172 | Lv-Ttc4 |
| a1 | SPU_012217 | Lv-Pex13_1 |
| a1 | SPU_012307 | Lv-Z322 |
| a1 | SPU_012344 | Lv-Dhdh_1 |
| a1 | SPU_012444 | NA |
| a1 | SPU_012495 | Lv-Kcmf1 |
| a1 | SPU_012496 | NA |
| a1 | SPU_012503 | Lv-Ndrk |
| a1 | SPU_012651 | Lv-Hao1_7 |
| a1 | SPU_012669 | none |
| a1 | SPU_012818 | Lv-Pet112l |
| a1 | SPU_012843 | Lv-Hypp_80 |
| a1 | SPU_012981 | Lv-Hypp_85 |
| a1 | SPU_012994 | NA |
| a1 | SPU_013099 | Lv-ASRGL1_ |
| a1 | SPU_013124 | Lv-Kiaa0746L |
| a1 | SPU_013168 | Lv-Slc30a5 |
| a1 | SPU_013181 | Lv-Slc30a9 |
| a1 | SPU_013189 | Lv-MucL_3 |
| a1 | SPU_013283 | Lv-Chsy1 |

| | | |
|---|---|---|
| a1 | SPU_013285 | Lv-Lgals3bpL |
| a1 | SPU_013307 | Lv-Z124_1 |
| a1 | SPU_013328 | none |
| a1 | SPU_013394 | Lv-TropC |
| a1 | SPU_013433 | Lv-Scyl3 |
| a1 | SPU_013461 | Lv-Hypp_717 |
| a1 | SPU_013647 | Lv-Cacybp |
| a1 | SPU_013737 | Lv-Rbm34L |
| a1 | SPU_013739 | Lv-B9d2 |
| a1 | SPU_013763 | Lv-Fam20c_1 |
| a1 | SPU_013768 | NA |
| a1 | SPU_013771 | Lv-Cep76 |
| a1 | SPU_013773 | Lv-Dusp15/22A |
| a1 | SPU_013788 | Lv-Ahr |
| a1 | SPU_013862 | Lv-Epb41l4A |
| a1 | SPU_013881 | Lv-Hypp_2113 |
| a1 | SPU_013904 | Lv-Z484 |
| a1 | SPU_013920 | Lv-Brf2L |
| a1 | SPU_013982 | Lv-Hypp_96 |
| a1 | SPU_014053 | none |
| a1 | SPU_014089 | Lv-Wdr91_1 |
| a1 | SPU_014106 | Lv-Dhx34_3 |
| a1 | SPU_014181 | Lv-ActL |
| a1 | SPU_014200 | Lv-Hypp_742 |
| a1 | SPU_014208 | Lv-GagpolL_7 |
| a1 | SPU_014235 | Lv-Abi1L |
| a1 | SPU_014242 | Lv-Bscl2 |
| a1 | SPU_014246 | Lv-Trmt5 |
| a1 | SPU_014247 | Lv-Egl9h2 |
| a1 | SPU_014271 | Lv-Fbxo41 |
| a1 | SPU_014325 | none |
| a1 | SPU_014334 | Lv-CacTta1Hs |
| a1 | SPU_014390 | Lv-Mfsd7L |
| a1 | SPU_014406 | Lv-Kcnn1 |
| a1 | SPU_014419 | Lv-Bat2 |
| a1 | SPU_014437 | Lv-C14orf102L |
| a1 | SPU_014445 | none |
| a1 | SPU_014526 | none |

| | | |
|---|---|---|
| a1 | SPU_014685 | Lv-Z80 |
| a1 | SPU_014686 | Lv-Z478_1 |
| a1 | SPU_014777 | Lv-Ldla/Egf/Ig/Lnb7/7Tm/Gpcr |
| a1 | SPU_014819 | Lv-Dex1 |
| a1 | SPU_014887 | Lv-Ypel5_1 |
| a1 | SPU_014908 | Lv-Ect2 |
| a1 | SPU_014913 | Lv-Rab4 |
| a1 | SPU_015023 | Lv-Zmynd11 |
| a1 | SPU_015034 | Lv-Hypp_2183 |
| a1 | SPU_015083 | Lv-Arrdc3_1 |
| a1 | SPU_015104 | Lv-C9orf86L |
| a1 | SPU_015126 | Lv-Usp47 |
| a1 | SPU_015169 | Lv-St8sia1_1 |
| a1 | SPU_015208 | Lv-Arl6ip4L |
| a1 | SPU_015298 | Lv-Hypp_780 |
| a1 | SPU_015307 | Lv-Tmem209 |
| a1 | SPU_015324 | Lv-Suv420h1L |
| a1 | SPU_015381 | Lv-Fn3/Egf_8 |
| a1 | SPU_015395 | Lv-Cdr2 |
| a1 | SPU_015398 | NA |
| a1 | SPU_015422 | Lv-C12orf44L |
| a1 | SPU_015486 | Lv-Gluld1_8 |
| a1 | SPU_015556 | Lv-Ap4e1 |
| a1 | SPU_015593 | Lv-Gpr45 |
| a1 | SPU_015709 | Lv-Dcun1d1 |
| a1 | SPU_015723 | Lv-Tcf25 |
| a1 | SPU_015819 | Lv-Homer2 |
| a1 | SPU_015882 | Lv-Znf608L |
| a1 | SPU_015909 | Lv-Dync1li2 |
| a1 | SPU_016026 | Lv-Hypp_2994 |
| a1 | SPU_016123 | none |
| a1 | SPU_016143 | NA |
| a1 | SPU_016149 | Lv-Z265 |
| a1 | SPU_016167 | none |
| a1 | SPU_016263 | none |
| a1 | SPU_016285 | Lv-Hypp_126 |
| a1 | SPU_016302 | Lv-C1galt1_1 |
| a1 | SPU_016328 | none |

| | | |
|---|---|---|
| a1 | SPU_016351 | Lv-Sh3d19L |
| a1 | SPU_016358 | Lv-Crocc-3 |
| a1 | SPU_016370 | Lv-Pionh |
| a1 | SPU_016374 | Lv-Srcr134 |
| a1 | SPU_016451 | Lv-Snx17 |
| a1 | SPU_016459 | Lv-Ipo13 |
| a1 | SPU_016499 | none |
| a1 | SPU_016518 | Lv-Z376 |
| a1 | SPU_016548 | Lv-Unk_34 |
| a1 | SPU_016576 | Lv-Pdcd2L |
| a1 | SPU_016603 | none |
| a1 | SPU_016622 | Lv-Endrvt92 |
| a1 | SPU_016656 | NA |
| a1 | SPU_016661 | Lv-Usp1 |
| a1 | SPU_016858 | Lv-Med14L_1 |
| a1 | SPU_016878 | Lv-Slc39a11 |
| a1 | SPU_016897 | Lv-Slc5a6_1 |
| a1 | SPU_016913 | Lv-Alg9 |
| a1 | SPU_016961 | Lv-Tmem9l |
| a1 | SPU_017027 | Lv-Map4k4 |
| a1 | SPU_017078 | NA |
| a1 | SPU_017126 | Lv-Nup153 |
| a1 | SPU_017144 | Lv-Vps33B |
| a1 | SPU_017156 | Lv-Shank2 |
| a1 | SPU_017164 | Lv-Zdhhc7_1 |
| a1 | SPU_017167 | none |
| a1 | SPU_017410 | Lv-C4orf29L |
| a1 | SPU_017493 | NA |
| a1 | SPU_017551 | Lv-Hypp_883 |
| a1 | SPU_017562 | NA |
| a1 | SPU_017567 | Lv-Slc35e1 |
| a1 | SPU_017626 | Lv-Rnf38L |
| a1 | SPU_017691 | Lv-Fpgt |
| a1 | SPU_017701 | Lv-Nwd1 |
| a1 | SPU_017738 | Lv-Abhd2 |
| a1 | SPU_017779 | none |
| a1 | SPU_017804 | Lv-Tm7sf3 |
| a1 | SPU_017820 | Lv-Hypp_845 |

| | | |
|---|---|---|
| a1 | SPU_017838 | Lv-B4galt7 |
| a1 | SPU_017876 | Lv-Gphn |
| a1 | SPU_017894 | Lv-Hypp_900 |
| a1 | SPU_017976 | Lv-Slc12a9 |
| a1 | SPU_018008 | Lv-Rnf113a |
| a1 | SPU_018010 | Lv-Apc |
| a1 | SPU_018046 | Lv-Dhodh |
| a1 | SPU_018073 | Lv-Hisppd2a |
| a1 | SPU_018075 | Lv-Z160 |
| a1 | SPU_018080 | Lv-Lrr/Igr_9 |
| a1 | SPU_018192 | Lv-Ttc13 |
| a1 | SPU_018198 | Lv-Lystr |
| a1 | SPU_018201 | NA |
| a1 | SPU_018308 | Lv-MterfL |
| a1 | SPU_018323 | Lv-Rapgef1 |
| a1 | SPU_018328 | Lv-Znhit6 |
| a1 | SPU_018373 | Lv-Z308 |
| a1 | SPU_018457 | Lv-Psf3 |
| a1 | SPU_018478 | Lv-Grm8 |
| a1 | SPU_018543 | none |
| a1 | SPU_018555 | Lv-Cul4_2 |
| a1 | SPU_018644 | Lv-Btb |
| a1 | SPU_018680 | Lv-Z338 |
| a1 | SPU_018695 | Lv-Slc38a7 |
| a1 | SPU_018782 | Lv-Kcnk10_1 |
| a1 | SPU_018845 | Lv-Nr1m4 |
| a1 | SPU_018952 | Lv-Tbc1d13 |
| a1 | SPU_018957 | Lv-Hypp_3036 |
| a1 | SPU_019007 | Lv-B4galt7_1 |
| a1 | SPU_019024 | Lv-Cbp |
| a1 | SPU_019025 | Lv-Cbpp |
| a1 | SPU_019045 | Lv-Ppp2rsBa |
| a1 | SPU_019265 | Lv-PolypL_115 |
| a1 | SPU_019297 | Lv-Catsper1 |
| a1 | SPU_019359 | Lv-Hypp_155 |
| a1 | SPU_019360 | Lv-Tbc1d15_1 |
| a1 | SPU_019411 | Lv-Hypp_2439 |
| a1 | SPU_019434 | none |

| | | |
|---|---|---|
| a1 | SPU_019453 | Lv-C19orf29 |
| a1 | SPU_019495 | Lv-Fam38b |
| a1 | SPU_019538 | Lv-Hypp_938 |
| a1 | SPU_019568 | Lv-Cdc25 |
| a1 | SPU_019630 | Lv-C7orf73L |
| a1 | SPU_019652 | Lv-Z189 |
| a1 | SPU_019783 | Lv-Cadherin_4 |
| a1 | SPU_019800 | Lv-Hypp_2467 |
| a1 | SPU_019862 | Lv-Ubr4_5 |
| a1 | SPU_019872 | Lv-Cgsp1 |
| a1 | SPU_019875 | Lv-Rnf41 |
| a1 | SPU_019948 | NA |
| a1 | SPU_019998 | Lv-Vnn1_1 |
| a1 | SPU_020029 | Lv-Sec23a |
| a1 | SPU_020031 | Lv-Pam |
| a1 | SPU_020068 | Lv-Hypp_960 |
| a1 | SPU_020069 | Lv-Mrck |
| a1 | SPU_020079 | none |
| a1 | SPU_020164 | none |
| a1 | SPU_020190 | Lv-Stk10 |
| a1 | SPU_020356 | Lv-Rfwd3 |
| a1 | SPU_020378 | Lv-Prdm9L |
| a1 | SPU_020391 | Lv-Cox10 |
| a1 | SPU_020458 | none |
| a1 | SPU_020462 | none |
| a1 | SPU_020481 | Lv-Z310 |
| a1 | SPU_020497 | Lv-Atp6v0a1_3 |
| a1 | SPU_020525 | Lv-Fa58C/Hyr/Lnb/7Tm/Gpcr |
| a1 | SPU_020542 | Lv-Ptpry2 |
| a1 | SPU_020585 | Lv-Ggt1_9 |
| a1 | SPU_020592 | Lv-Dgkd |
| a1 | SPU_020610 | Lv-Zfand5b |
| a1 | SPU_020611 | Lv-Fray |
| a1 | SPU_020669 | Lv-Abcc4a |
| a1 | SPU_020701 | none |
| a1 | SPU_020797 | Lv-C18orf22 |
| a1 | SPU_020806 | none |
| a1 | SPU_020879 | Lv-Diaph |

| | | |
|---|---|---|
| a1 | SPU_020923 | Lv-Trim45 |
| a1 | SPU_020964 | Lv-Gfod2 |
| a1 | SPU_020966 | Lv-Ship |
| a1 | SPU_021020 | Lv-Slc30a3_2 |
| a1 | SPU_021036 | Lv-Kiaa1267L |
| a1 | SPU_021038 | Lv-Cep78 |
| a1 | SPU_021039 | Lv-Cep78L |
| a1 | SPU_021108 | Lv-Hypp_2537 |
| a1 | SPU_021211 | Lv-Nkiras |
| a1 | SPU_021315 | Lv-Ndor1 |
| a1 | SPU_021471 | none |
| a1 | SPU_021484 | Lv-March6 |
| a1 | SPU_021493 | Lv-Ctdp1 |
| a1 | SPU_021597 | none |
| a1 | SPU_021599 | Lv-Ptpr1 |
| a1 | SPU_021649 | Lv-Rasa3 |
| a1 | SPU_021714 | none |
| a1 | SPU_021771 | Lv-Ube3b |
| a1 | SPU_021783 | Lv-Glrx_1 |
| a1 | SPU_021797 | Lv-Pnpt1 |
| a1 | SPU_021812 | Lv-CycT |
| a1 | SPU_021845 | Lv-SmtnL_2 |
| a1 | SPU_021858 | Lv-Lrrkc |
| a1 | SPU_021896 | Lv-Ubp24 |
| a1 | SPU_021923 | Lv-Vti1ah |
| a1 | SPU_021935 | Lv-Hdac8 |
| a1 | SPU_021961 | Lv-Cnksr2L |
| a1 | SPU_022013 | Lv-Hypp_179 |
| a1 | SPU_022191 | Lv-Zyg11b |
| a1 | SPU_022200 | Lv-Wnk |
| a1 | SPU_022357 | Lv-Xpo7 |
| a1 | SPU_022450 | Lv-Blnk |
| a1 | SPU_022486 | Lv-Zdhhc23 |
| a1 | SPU_022500 | Lv-Hypp_185 |
| a1 | SPU_022590 | Lv-Cyp2L51 |
| a1 | SPU_022630 | Lv-Arih2 |
| a1 | SPU_022668 | Lv-Hypp_1053 |
| a1 | SPU_022669 | Lv-Mtmr6/7/8 |

| | | |
|---|---|---|
| a1 | SPU_022673 | none |
| a1 | SPU_022708 | Lv-Myd88L3 |
| a1 | SPU_022710 | Lv-Helz |
| a1 | SPU_022717 | Lv-Pi3kp110b |
| a1 | SPU_022820 | Lv-Soxb1 |
| a1 | SPU_022840 | Lv-Kif12 |
| a1 | SPU_022867 | Lv-CdkaL1 |
| a1 | SPU_022872 | Lv-Sans |
| a1 | SPU_022879 | Lv-Icmt |
| a1 | SPU_022950 | Lv-Wapal |
| a1 | SPU_022983 | Lv-C4orf27 |
| a1 | SPU_023070 | Lv-Usp32_2 |
| a1 | SPU_023269 | none |
| a1 | SPU_023389 | Lv-Ptr4 |
| a1 | SPU_023395 | Lv-UreF |
| a1 | SPU_023426 | Lv-Slc25a30 |
| a1 | SPU_023444 | NA |
| a1 | SPU_023515 | Lv-Phkb_1 |
| a1 | SPU_023546 | Lv-Slc11a2_1 |
| a1 | SPU_023563 | Lv-PolypL_83 |
| a1 | SPU_023592 | Lv-Setdb1_1 |
| a1 | SPU_023602 | Lv-Hypp_2678 |
| a1 | SPU_023648 | Lv-Mgat1 |
| a1 | SPU_023671 | Lv-Scavr |
| a1 | SPU_023685 | Lv-Taf7 |
| a1 | SPU_023686 | Lv-Eif3s5L |
| a1 | SPU_023791 | Lv-Tnni3k_1 |
| a1 | SPU_023800 | Lv-C5orf34L |
| a1 | SPU_023837 | Lv-Rab3gap2 |
| a1 | SPU_023856 | Lv-Afdn |
| a1 | SPU_023864 | none |
| a1 | SPU_023939 | Lv-Hypp_1100 |
| a1 | SPU_024035 | Lv-Aak1 |
| a1 | SPU_024048 | Lv-Oach_3 |
| a1 | SPU_024081 | Lv-Dyx1c1 |
| a1 | SPU_024092 | Lv-ACVR |
| a1 | SPU_024151 | Lv-St14L_3 |
| a1 | SPU_024224 | Lv-Coq4_1 |

| | | |
|---|---|---|
| a1 | SPU_024257 | Lv-LamG/Egff5 |
| a1 | SPU_024268 | Lv-Pnlcd1 |
| a1 | SPU_024277 | none |
| a1 | SPU_024287 | Lv-Cytsa-2 |
| a1 | SPU_024299 | Lv-Unk_52 |
| a1 | SPU_024316 | Lv-Mekkk5 |
| a1 | SPU_024329 | Lv-Cuedc1 |
| a1 | SPU_024350 | Lv-Tdrdcp_1 |
| a1 | SPU_024393 | NA |
| a1 | SPU_024405 | Lv-St6galnac2_1 |
| a1 | SPU_024443 | Lv-Abp34L |
| a1 | SPU_024480 | Lv-Chz10L |
| a1 | SPU_024534 | Lv-Faf1 |
| a1 | SPU_024637 | Lv-Ssx2ip |
| a1 | SPU_024656 | Lv-Dscr1l2_1 |
| a1 | SPU_024695 | Lv-JmyL |
| a1 | SPU_024794 | Lv-Pen2 |
| a1 | SPU_024838 | Lv-Madd |
| a1 | SPU_024875 | Lv-Rfp19a |
| a1 | SPU_024901 | NA |
| a1 | SPU_025011 | Lv-Hypp_1134 |
| a1 | SPU_025190 | Lv-Fbxw7_1 |
| a1 | SPU_025253 | NA |
| a1 | SPU_025330 | Lv-Hypp_1143 |
| a1 | SPU_025343 | Lv-Frmd1 |
| a1 | SPU_025512 | Lv-C19orf29L |
| a1 | SPU_025531 | none |
| a1 | SPU_025533 | Lv-Tdrd5 |
| a1 | SPU_025535 | Lv-Dolpp1 |
| a1 | SPU_025578 | Lv-Pro_ser_thr_ppip1 |
| a1 | SPU_025610 | Lv-PppL_144 |
| a1 | SPU_025624 | Lv-Taf1 |
| a1 | SPU_025628 | Lv-Nfxl1-2 |
| a1 | SPU_025631 | Lv-Rnf219 |
| a1 | SPU_025671 | Lv-Tbpaf_RnapIA |
| a1 | SPU_025682 | Lv-Rspry1L |
| a1 | SPU_025757 | Lv-Hypp_1156 |
| a1 | SPU_025759 | Lv-Rptpd_fn3r |

| | | |
|---|---|---|
| a1 | SPU_025764 | Lv-Dhx33 |
| a1 | SPU_025776 | Lv-Apaf1 |
| a1 | SPU_025778 | Lv-Kiaa1407L |
| a1 | SPU_025781 | Lv-Slc10a2L |
| a1 | SPU_025891 | Lv-Crot_2 |
| a1 | SPU_025990 | none |
| a1 | SPU_026013 | Lv-Atp/Gtpbp1 |
| a1 | SPU_026049 | Lv-Ggt1_7 |
| a1 | SPU_026177 | Lv-Ints6 |
| a1 | SPU_026187 | Lv-Ubr4_7 |
| a1 | SPU_026195 | none |
| a1 | SPU_026209 | Lv-Z445 |
| a1 | SPU_026211 | none |
| a1 | SPU_026270 | Lv-Ube2j2 |
| a1 | SPU_026350 | none |
| a1 | SPU_026366 | none |
| a1 | SPU_026385 | Lv-Z427 |
| a1 | SPU_026450 | Lv-Mcm7 |
| a1 | SPU_026503 | Lv-Krp170p |
| a1 | SPU_026509 | Lv-Insc |
| a1 | SPU_026527 | Lv-PcnxL |
| a1 | SPU_026683 | Lv-Mitd1 |
| a1 | SPU_026697 | Lv-Hypp_2860 |
| a1 | SPU_026717 | NA |
| a1 | SPU_026822 | Lv-MthfsdL |
| a1 | SPU_026828 | Lv-Msn1 |
| a1 | SPU_026834 | Lv-Fuca2_1 |
| a1 | SPU_026844 | Lv-Hypp_1188 |
| a1 | SPU_027072 | Lv-Tecpr1 |
| a1 | SPU_027099 | Lv-Hypp_2886 |
| a1 | SPU_027122 | Lv-Hypp_1198 |
| a1 | SPU_027136 | Lv-Ldla/Lrr/Gpcr |
| a1 | SPU_027161 | Lv-Tecpr1_1 |
| a1 | SPU_027171 | none |
| a1 | SPU_027231 | Lv-Gal3st1_14 |
| a1 | SPU_027280 | none |
| a1 | SPU_027411 | Lv-Tdrdcp_2 |
| a1 | SPU_027457 | Lv-Hsdl1 |

| | | |
|---|---|---|
| a1 | SPU_027522 | none |
| a1 | SPU_027544 | Lv-Hypp_269 |
| a1 | SPU_027567 | Lv-Skiv2l2 |
| a1 | SPU_027573 | NA |
| a1 | SPU_027580 | Lv-Arp8ph |
| a1 | SPU_027589 | Lv-Hypp_2913 |
| a1 | SPU_027662 | Lv-Neu3 |
| a1 | SPU_027709 | Lv-Z125 |
| a1 | SPU_027728 | Lv-Pla2g4b_1 |
| a1 | SPU_027743 | Lv-Fbxl20 |
| a1 | SPU_027825 | Lv-Hypp_1215 |
| a1 | SPU_027873 | none |
| a1 | SPU_027888 | Lv-Bace1 |
| a1 | SPU_027903 | Lv-Pibf1L |
| a1 | SPU_027912 | Lv-Z365 |
| a1 | SPU_027946 | Lv-Csad_1 |
| a1 | SPU_027973 | none |
| a1 | SPU_028017 | Lv-Usp40 |
| a1 | SPU_028053 | Lv-Rad1 |
| a1 | SPU_028074 | Lv-Zcchc17 |
| a1 | SPU_028080 | Lv-Selp |
| a1 | SPU_028103 | Lv-Prtelsn |
| a1 | SPU_028186 | Lv-Hypp_2941 |
| a1 | SPU_028197 | Lv-Auts2L |
| a1 | SPU_028222 | Lv-Z146 |
| a1 | SPU_028236 | Lv-Cdc2l5 |
| a1 | SPU_028256 | Lv-Hypp_281 |
| a1 | SPU_028393 | Lv-NotchL_96 |
| a1 | SPU_028438 | Lv-Cxxc1L |
| a1 | SPU_028448 | Lv-Ormdl2 |
| a1 | SPU_028469 | Lv-SerpL1 |
| a1 | SPU_028544 | none |
| a1 | SPU_028616 | Lv-Hypp_2961 |
| a1 | SPU_028877 | Lv-Fhdc1_2 |
| a1 | SPU_028898 | Lv-Traf6 |
| a1 | SPU_028921 | Lv-Hells |
| a1 | SPU_030065 | Lv-Triad_2 |
| a2 | SPU_000102 | Lv-Snrk |

| | | |
|---|---|---|
| a2 | SPU_000151 | Lv-Cdk6 |
| a2 | SPU_000169 | Lv-Ncoa7l |
| a2 | SPU_000215 | none |
| a2 | SPU_000237 | Lv-Jip1 |
| a2 | SPU_000257 | Lv-Rab3fap1 |
| a2 | SPU_000280 | Lv-Zc2p1L |
| a2 | SPU_000355 | none |
| a2 | SPU_000487 | Lv-Dak |
| a2 | SPU_000542 | Lv-Clect/Pan_7 |
| a2 | SPU_000559 | Lv-Sec24b |
| a2 | SPU_000617 | none |
| a2 | SPU_000698 | Lv-Hypp_1276 |
| a2 | SPU_000703 | Lv-Hypp_1280 |
| a2 | SPU_000810 | Lv-Efcab7 |
| a2 | SPU_000922 | Lv-Ext(multiple)L3 |
| a2 | SPU_001069 | Lv-Fbf1 |
| a2 | SPU_001070 | none |
| a2 | SPU_001097 | none |
| a2 | SPU_001167 | none |
| a2 | SPU_001284 | Lv-Kif1BL3 |
| a2 | SPU_001313 | Lv-Hypp_1321 |
| a2 | SPU_001317 | Lv-Eme1h |
| a2 | SPU_001396 | Lv-Jip3_2 |
| a2 | SPU_001426 | Lv-Syne1L_3 |
| a2 | SPU_001496 | Lv-Dnajhc1 |
| a2 | SPU_001536 | Lv-GnRHLr |
| a2 | SPU_001546 | Lv-Fam20b |
| a2 | SPU_001612 | Lv-Tmem169 |
| a2 | SPU_001792 | NA |
| a2 | SPU_001834 | Lv-Mpdz-2 |
| a2 | SPU_001894 | NA |
| a2 | SPU_001896 | Lv-Cdan1L |
| a2 | SPU_001969 | none |
| a2 | SPU_002046 | Lv-Cdt1 |
| a2 | SPU_002069 | Lv-Sgcz |
| a2 | SPU_002094 | Lv-CckarL |
| a2 | SPU_002149 | none |
| a2 | SPU_002163 | Lv-Znfx1_2 |

| | | |
|---|---|---|
| a2 | SPU_002215 | Lv-Agsd |
| a2 | SPU_002223 | Lv-My11hp_smph2 |
| a2 | SPU_002229 | Lv-Slc29a3L |
| a2 | SPU_002269 | Lv-C6orf192L |
| a2 | SPU_002333 | Lv-Cnpy2h |
| a2 | SPU_002379 | Lv-Tyki |
| a2 | SPU_002427 | Lv-Slc7a2 |
| a2 | SPU_002516 | Lv-Gpr51 |
| a2 | SPU_002519 | Lv-Ap4m1 |
| a2 | SPU_002555 | Lv-Scrib |
| a2 | SPU_002656 | Lv-Cyp2L5 |
| a2 | SPU_002663 | Lv-Etub1 |
| a2 | SPU_002666 | Lv-Plekha7-2 |
| a2 | SPU_002736 | Lv-Slc12a4 |
| a2 | SPU_002773 | NA |
| a2 | SPU_002843 | Lv-C2orf18L |
| a2 | SPU_002866 | Lv-Deltex4 |
| a2 | SPU_002874 | Lv-Ets1/2 |
| a2 | SPU_002907 | Lv-Gpt2_1 |
| a2 | SPU_002990 | Lv-St8sia4_3 |
| a2 | SPU_003007 | none |
| a2 | SPU_003032 | Lv-G11 |
| a2 | SPU_003099 | Lv-Gpatch2 |
| a2 | SPU_003121 | Lv-Hypp_1434 |
| a2 | SPU_003228 | Lv-Rabl2 |
| a2 | SPU_003229 | Lv-Slc40a1 |
| a2 | SPU_003236 | Lv-Tnpo3 |
| a2 | SPU_003250 | Lv-Hypp_3100 |
| a2 | SPU_003328 | Lv-Kiaa0859L |
| a2 | SPU_003355 | Lv-C2h2/swim1l |
| a2 | SPU_003386 | Lv-Hypp_1450 |
| a2 | SPU_003445 | none |
| a2 | SPU_003486 | Lv-ScavrBIIIL |
| a2 | SPU_003490 | Lv-Z131 |
| a2 | SPU_003498 | Lv-Hypp_1458 |
| a2 | SPU_003528 | Lv-CycA |
| a2 | SPU_003680 | Lv-Hypp_3092 |
| a2 | SPU_003747 | Lv-Mov10L_2 |

| | | |
|---|---|---|
| a2 | SPU_003802 | Lv-Dullardh |
| a2 | SPU_003883 | Lv-Galnt13-2 |
| a2 | SPU_003898 | Lv-Gq |
| a2 | SPU_003911 | Lv-Il1ap |
| a2 | SPU_003930 | Lv-Srcr28 |
| a2 | SPU_004222 | Lv-Rilpl1L |
| a2 | SPU_004230 | Lv-Plcd |
| a2 | SPU_004243 | Lv-Npr1_5 |
| a2 | SPU_004259 | Lv-Trubh2 |
| a2 | SPU_004260 | Lv-Hs2St |
| a2 | SPU_004357 | Lv-Udg |
| a2 | SPU_004382 | Lv-Whsc1 |
| a2 | SPU_004422 | Lv-Mpdz-3 |
| a2 | SPU_004428 | Lv-Hypp_1522 |
| a2 | SPU_004473 | Lv-Sec14L2 |
| a2 | SPU_004557 | Lv-Sarmr11 |
| a2 | SPU_004567 | Lv-Gabbr1_3 |
| a2 | SPU_004589 | Lv-Hypp_420 |
| a2 | SPU_004606 | none |
| a2 | SPU_004679 | none |
| a2 | SPU_004825 | Lv-IunhL |
| a2 | SPU_004864 | Lv-Mpp5-2 |
| a2 | SPU_004888 | Lv-Mbr1 |
| a2 | SPU_004925 | Lv-Htr1L |
| a2 | SPU_005023 | Lv-Unc44L_37 |
| a2 | SPU_005055 | Lv-Tk2_1 |
| a2 | SPU_005099 | Lv-Mtmr5/13 |
| a2 | SPU_005104 | Lv-Trubh1 |
| a2 | SPU_005198 | Lv-YP30/FascL_3 |
| a2 | SPU_005255 | Lv-Hypp_5 |
| a2 | SPU_005349 | NA |
| a2 | SPU_005363 | Lv-Mib2L_2 |
| a2 | SPU_005529 | Lv-Kiaa0317L |
| a2 | SPU_005573 | none |
| a2 | SPU_005580 | Lv-Kiaa1239 |
| a2 | SPU_005587 | Lv-PolypL |
| a2 | SPU_005589 | Lv-Rt25 |
| a2 | SPU_005626 | Lv-Tesk |

| | | |
|---|---|---|
| a2 | SPU_005857 | Lv-Ergic2 |
| a2 | SPU_005860 | Lv-Srcr/Ptpc1 |
| a2 | SPU_005871 | Lv-Il1r1 |
| a2 | SPU_005947 | none |
| a2 | SPU_005954 | Lv-Npr1 |
| a2 | SPU_005970 | Lv-Unc44L_47 |
| a2 | SPU_005994 | Lv-Hypp_468 |
| a2 | SPU_006005 | Lv-Ust_2 |
| a2 | SPU_006021 | Lv-Gpr180 |
| a2 | SPU_006029 | Lv-KnkL |
| a2 | SPU_006065 | NA |
| a2 | SPU_006124 | Lv-Bokl |
| a2 | SPU_006132 | Lv-Ppm1l |
| a2 | SPU_006133 | Lv-Hypp_1634 |
| a2 | SPU_006141 | none |
| a2 | SPU_006216 | Lv-Flj20433L |
| a2 | SPU_006255 | Lv-Ppp1r3aL |
| a2 | SPU_006262 | Lv-Tmco7 |
| a2 | SPU_006299 | none |
| a2 | SPU_006314 | Lv-Hypp_1656 |
| a2 | SPU_006319 | Lv-Traf7 |
| a2 | SPU_006379 | Lv-Gle1l |
| a2 | SPU_006497 | Lv-Z136 |
| a2 | SPU_006501 | Lv-Hypp_1665 |
| a2 | SPU_006574 | Lv-Cyp2L14 |
| a2 | SPU_006602 | Lv-Kif13A2 |
| a2 | SPU_006640 | none |
| a2 | SPU_006657 | Lv-Mpdz |
| a2 | SPU_006658 | NA |
| a2 | SPU_006663 | Lv-Xylt1 |
| a2 | SPU_006674 | Lv-Clspn2 |
| a2 | SPU_006725 | Lv-Psd3L |
| a2 | SPU_006815 | NA |
| a2 | SPU_006843 | Lv-Slc16a12_3 |
| a2 | SPU_007000 | Lv-Rasa1_2 |
| a2 | SPU_007221 | Lv-Cp_1 |
| a2 | SPU_007234 | Lv-Uhrf1L |
| a2 | SPU_007295 | Lv-Snap47 |

| | | |
|---|---|---|
| a2 | SPU_007478 | Lv-Depdc7 |
| a2 | SPU_007526 | Lv-Relr2 |
| a2 | SPU_007532 | Lv-KirreL3 |
| a2 | SPU_007587 | Lv-Mkk3/6 |
| a2 | SPU_007597 | Lv-Hypp_522 |
| a2 | SPU_007645 | none |
| a2 | SPU_007654 | Lv-Hypp_2987 |
| a2 | SPU_007747 | Lv-Grm3 |
| a2 | SPU_007756 | Lv-Cyb561d1 |
| a2 | SPU_007877 | Lv-Ankrd49 |
| a2 | SPU_007923 | Lv-Hypp_532 |
| a2 | SPU_007950 | Lv-Asphd2 |
| a2 | SPU_008116 | Lv-Rpap3 |
| a2 | SPU_008160 | Lv-Trpm_3 |
| a2 | SPU_008179 | Lv-Tanc2 |
| a2 | SPU_008237 | Lv-Hypp_31 |
| a2 | SPU_008251 | Lv-Txndc15 |
| a2 | SPU_008259 | Lv-Lrr/Gpcr_6 |
| a2 | SPU_008277 | Lv-Cadps |
| a2 | SPU_008355 | Lv-Hypp_1781 |
| a2 | SPU_008453 | Lv-Smarcal1 |
| a2 | SPU_008475 | Lv-Hypp_549 |
| a2 | SPU_008502 | Lv-Pp4r1 |
| a2 | SPU_008511 | none |
| a2 | SPU_008523 | Lv-Hypp_1789 |
| a2 | SPU_008715 | Lv-Slc39a10_1 |
| a2 | SPU_008787 | none |
| a2 | SPU_008848 | none |
| a2 | SPU_008855 | Lv-Syn3 |
| a2 | SPU_008876 | Lv-Bfar |
| a2 | SPU_008882 | NA |
| a2 | SPU_008926 | Lv-Ldla/Ig/Gpcr |
| a2 | SPU_008940 | none |
| a2 | SPU_008949 | Lv-CDNA1 |
| a2 | SPU_009025 | Lv-Kctd21 |
| a2 | SPU_009132 | none |
| a2 | SPU_009198 | Lv-Z479 |
| a2 | SPU_009215 | Lv-Fmi2 |

| | | |
|---|---|---|
| a2 | SPU_009325 | Lv-Pth2rL |
| a2 | SPU_009362 | none |
| a2 | SPU_009388 | Lv-Php |
| a2 | SPU_009409 | Lv-Tmem179L |
| a2 | SPU_009412 | Lv-Trmt61a |
| a2 | SPU_009593 | Lv-Slbp |
| a2 | SPU_009715 | Lv-Rad30 |
| a2 | SPU_009758 | Lv-Phospho2 |
| a2 | SPU_009762 | Lv-Strabs |
| a2 | SPU_009766 | Lv-Hrh2 |
| a2 | SPU_009774 | Lv-C10orf97L |
| a2 | SPU_009807 | Lv-Opa1L(N-term) |
| a2 | SPU_009812 | Lv-Calpain_1 |
| a2 | SPU_009867 | Lv-Sentrin |
| a2 | SPU_009877 | Lv-Dhx30 |
| a2 | SPU_009900 | Lv-Bves |
| a2 | SPU_009929 | Lv-Plcb3 |
| a2 | SPU_009942 | Lv-Ddx58_2 |
| a2 | SPU_009963 | none |
| a2 | SPU_010017 | Lv-Ptpmt1 |
| a2 | SPU_010032 | Lv-C6st_1 |
| a2 | SPU_010120 | Lv-Acy1 |
| a2 | SPU_010168 | Lv-Fog |
| a2 | SPU_010200 | Lv-Hypp_64 |
| a2 | SPU_010202 | Lv-Hypp_65 |
| a2 | SPU_010230 | Lv-TnfrsfL1 |
| a2 | SPU_010399 | none |
| a2 | SPU_010444 | none |
| a2 | SPU_010447 | none |
| a2 | SPU_010478 | Lv-Hypp_616 |
| a2 | SPU_010509 | Lv-Prkg1_1 |
| a2 | SPU_010595 | Lv-Cdc6 |
| a2 | SPU_010632 | Lv-Ccdc49 |
| a2 | SPU_010649 | NA |
| a2 | SPU_010675 | none |
| a2 | SPU_010708 | NA |
| a2 | SPU_010784 | Lv-Radr_2 |
| a2 | SPU_010794 | Lv-Oprk1L_1 |

| | | |
|---|---|---|
| a2 | SPU_010812 | Lv-Hypp_627 |
| a2 | SPU_010835 | Lv-Dgcr8L |
| a2 | SPU_010844 | Lv-Nagl |
| a2 | SPU_010917 | none |
| a2 | SPU_010994 | Lv-Srcr98 |
| a2 | SPU_011035 | Lv-Nlr162 |
| a2 | SPU_011042 | Lv-Sarm |
| a2 | SPU_011058 | Lv-Mfsd1_1 |
| a2 | SPU_011059 | Lv-Pla2g4b |
| a2 | SPU_011078 | none |
| a2 | SPU_011095 | Lv-My_hpp13sm |
| a2 | SPU_011110 | Lv-Pp1 |
| a2 | SPU_011118 | NA |
| a2 | SPU_011245 | Lv-Chst11_11 |
| a2 | SPU_011281 | NA |
| a2 | SPU_011335 | Lv-Ptprg/Z1 |
| a2 | SPU_011364 | Lv-LshmanlsnL_2 |
| a2 | SPU_011405 | Lv-Hrh3L_10 |
| a2 | SPU_011449 | Lv-Plcb4_1 |
| a2 | SPU_011531 | Lv-Daxx |
| a2 | SPU_011547 | Lv-Jmjd4l |
| a2 | SPU_011635 | Lv-FoxJ2_1 |
| a2 | SPU_011645 | Lv-Unk_82 |
| a2 | SPU_011673 | Lv-Hypp_1997 |
| a2 | SPU_011691 | Lv-Mrm1 |
| a2 | SPU_011758 | none |
| a2 | SPU_011828 | Lv-Slc16a9_2 |
| a2 | SPU_011895 | none |
| a2 | SPU_011959 | Lv-Pigx |
| a2 | SPU_012004 | Lv-Mrpl |
| a2 | SPU_012122 | Lv-Pknox |
| a2 | SPU_012179 | none |
| a2 | SPU_012318 | Lv-KpLr3 |
| a2 | SPU_012395 | Lv-Toe1 |
| a2 | SPU_012401 | Lv-Hypp_684 |
| a2 | SPU_012505 | Lv-Pald |
| a2 | SPU_012520 | Lv-Dusp4L |
| a2 | SPU_012527 | Lv-Sycp2 |

| | | |
|---|---|---|
| a2 | SPU_012540 | Lv-Gatm |
| a2 | SPU_012553 | none |
| a2 | SPU_012666 | Lv-Lrrc4cL |
| a2 | SPU_012698 | Lv-Inpp5a |
| a2 | SPU_012753 | Lv-Trpa1L_3 |
| a2 | SPU_012813 | none |
| a2 | SPU_012864 | none |
| a2 | SPU_012911 | Lv-Z33 |
| a2 | SPU_012922 | Lv-Ubox5 |
| a2 | SPU_012948 | Lv-Z50 |
| a2 | SPU_012949 | Lv-Z122 |
| a2 | SPU_013179 | Lv-Htf9cL |
| a2 | SPU_013216 | NA |
| a2 | SPU_013300 | Lv-Evi5_1 |
| a2 | SPU_013431 | Lv-Peo1 |
| a2 | SPU_013489 | Lv-C3orf39 |
| a2 | SPU_013588 | Lv-Hypp_2090 |
| a2 | SPU_013590 | Lv-DmpyL |
| a2 | SPU_013603 | Lv-Hypp_93 |
| a2 | SPU_013734 | Lv-Fa58C/Clect/Egf/Cub |
| a2 | SPU_013754 | Lv-Clcn6 |
| a2 | SPU_013817 | Lv-Ttll10L_2 |
| a2 | SPU_013965 | Lv-Mol10l1 |
| a2 | SPU_013974 | Lv-Trmt1h |
| a2 | SPU_014058 | none |
| a2 | SPU_014103 | none |
| a2 | SPU_014104 | none |
| a2 | SPU_014144 | Lv-Ninj1L |
| a2 | SPU_014166 | none |
| a2 | SPU_014256 | Lv-Z411 |
| a2 | SPU_014281 | Lv-Fam48a |
| a2 | SPU_014301 | Lv-Alkbh1 |
| a2 | SPU_014368 | Lv-Klraq1 |
| a2 | SPU_014383 | none |
| a2 | SPU_014404 | Lv-Dapk1 |
| a2 | SPU_014405 | Lv-Pnr |
| a2 | SPU_014448 | Lv-Hypp_2151 |
| a2 | SPU_014497 | Lv-Rab37 |

| | | |
|---|---|---|
| a2 | SPU_014588 | Lv-Mb |
| a2 | SPU_014712 | Lv-Nxf1 |
| a2 | SPU_014769 | Lv-Lrr/Gpcr_3 |
| a2 | SPU_014795 | Lv-Trpc4 |
| a2 | SPU_014820 | Lv-Hypp_2170 |
| a2 | SPU_014920 | Lv-Slc25a44 |
| a2 | SPU_014960 | Lv-Hypp_769 |
| a2 | SPU_015085 | Lv-Fam116a |
| a2 | SPU_015162 | NA |
| a2 | SPU_015263 | Lv-Edc4L |
| a2 | SPU_015285 | Lv-Cycb |
| a2 | SPU_015349 | Lv-Nuak |
| a2 | SPU_015367 | Lv-Hypp_2208 |
| a2 | SPU_015417 | Lv-Hrh1_1 |
| a2 | SPU_015450 | Lv-Cpeb1 |
| a2 | SPU_015527 | NA |
| a2 | SPU_015537 | Lv-Arrdc2L |
| a2 | SPU_015538 | Lv-C1orf85L |
| a2 | SPU_015621 | Lv-Madd_1 |
| a2 | SPU_015627 | Lv-Prmt2 |
| a2 | SPU_015691 | Lv-Impa1 |
| a2 | SPU_015745 | none |
| a2 | SPU_015820 | Lv-Hypp_118 |
| a2 | SPU_015876 | Lv-Git1_1 |
| a2 | SPU_015887 | none |
| a2 | SPU_015895 | Lv-Scnn1aL_2 |
| a2 | SPU_015914 | Lv-Hypp_121 |
| a2 | SPU_015916 | Lv-Calcr |
| a2 | SPU_015949 | none |
| a2 | SPU_015968 | Lv-Hrh2_3 |
| a2 | SPU_016064 | Lv-Gkap1 |
| a2 | SPU_016067 | Lv-Kif24 |
| a2 | SPU_016125 | Lv-Tpst2 |
| a2 | SPU_016284 | none |
| a2 | SPU_016288 | Lv-Cnpy4 |
| a2 | SPU_016293 | Lv-Hypp_820 |
| a2 | SPU_016333 | Lv-Slc46a1 |
| a2 | SPU_016393 | Lv-Saps3_1 |

| | | |
|---|---|---|
| a2 | SPU_016394 | Lv-Saps3 |
| a2 | SPU_016447 | Lv-Exosc10_1 |
| a2 | SPU_016456 | Lv-Gababr2 |
| a2 | SPU_016555 | Lv-Egf/LamGap2 |
| a2 | SPU_016578 | Lv-Jmjd5_1 |
| a2 | SPU_016650 | Lv-Par1 |
| a2 | SPU_016707 | none |
| a2 | SPU_016735 | Lv-PppL_110 |
| a2 | SPU_016762 | Lv-Stom_5 |
| a2 | SPU_016835 | Lv-Megf6_4 |
| a2 | SPU_016842 | Lv-Mpp5-3 |
| a2 | SPU_016843 | Lv-Hypp_134 |
| a2 | SPU_016955 | Lv-Mfsd6l_2 |
| a2 | SPU_016963 | Lv-Elac2-2 |
| a2 | SPU_017020 | none |
| a2 | SPU_017112 | Lv-Rhbdf1 |
| a2 | SPU_017151 | Lv-Hypp_2306 |
| a2 | SPU_017166 | none |
| a2 | SPU_017188 | Lv-BcatL |
| a2 | SPU_017190 | Lv-Rab11fip3L |
| a2 | SPU_017200 | Lv-Sgce |
| a2 | SPU_017206 | Lv-C20orf4 |
| a2 | SPU_017249 | Lv-Arxl |
| a2 | SPU_017369 | Lv-Hypp_1349 |
| a2 | SPU_017401 | none |
| a2 | SPU_017415 | Lv-Dctd_1 |
| a2 | SPU_017419 | none |
| a2 | SPU_017426 | Lv-Mglur8 |
| a2 | SPU_017437 | Lv-Phkg1 |
| a2 | SPU_017478 | Lv-Dgki |
| a2 | SPU_017486 | Lv-Rmi1 |
| a2 | SPU_017492 | Lv-Reverb |
| a2 | SPU_017638 | Lv-ExdL1 |
| a2 | SPU_017746 | Lv-Hypp_893 |
| a2 | SPU_017801 | Lv-C1galt1_2 |
| a2 | SPU_017884 | none |
| a2 | SPU_017926 | Lv-Armc5 |
| a2 | SPU_017974 | Lv-Cttnbp2 |

| | | |
|---|---|---|
| a2 | SPU_018077 | Lv-Kif15L |
| a2 | SPU_018107 | Lv-Plekha7 |
| a2 | SPU_018216 | Lv-Tmem47 |
| a2 | SPU_018249 | Lv-St8sia_2 |
| a2 | SPU_018281 | Lv-Snx19 |
| a2 | SPU_018333 | Lv-Rgs3 |
| a2 | SPU_018355 | Lv-Hypp_142 |
| a2 | SPU_018366 | Lv-Rara |
| a2 | SPU_018417 | Lv-Gramd4 |
| a2 | SPU_018450 | none |
| a2 | SPU_018471 | Lv-Kcnt1 |
| a2 | SPU_018544 | Lv-Mfsd4 |
| a2 | SPU_018549 | Lv-Thada |
| a2 | SPU_018769 | Lv-Cerk |
| a2 | SPU_018777 | NA |
| a2 | SPU_018808 | Lv-Atad4 |
| a2 | SPU_018832 | Lv-Trpa1_5 |
| a2 | SPU_018838 | Lv-Tlr100 |
| a2 | SPU_018915 | Lv-TnfrsfL5 |
| a2 | SPU_018997 | Lv-Rho |
| a2 | SPU_019022 | Lv-Jip3_1 |
| a2 | SPU_019063 | Lv-Nek9-2 |
| a2 | SPU_019080 | NA |
| a2 | SPU_019098 | Lv-Hypp_928 |
| a2 | SPU_019099 | Lv-Znrf3 |
| a2 | SPU_019101 | Lv-Xpapd2Lmb_2 |
| a2 | SPU_019117 | Lv-P90rsk1 |
| a2 | SPU_019119 | Lv-Ap1gbp1L |
| a2 | SPU_019152 | Lv-Ldargampdl |
| a2 | SPU_019226 | Lv-Hypp_151 |
| a2 | SPU_019261 | Lv-Flam2(N-terminus) |
| a2 | SPU_019277 | none |
| a2 | SPU_019283 | none |
| a2 | SPU_019341 | Lv-Kclqt1 |
| a2 | SPU_019365 | Lv-Z23 |
| a2 | SPU_019552 | Lv-PusL1 |
| a2 | SPU_019573 | Lv-Hypp_2450 |
| a2 | SPU_019592 | Lv-Ugt2b33_1 |

| | | |
|---|---|---|
| a2 | SPU_019755 | Lv-Osbpl1a |
| a2 | SPU_019756 | none |
| a2 | SPU_019776 | Lv-Zcchc8 |
| a2 | SPU_019821 | Lv-Orc1 |
| a2 | SPU_019883 | Lv-Cyp1r3 |
| a2 | SPU_019972 | Lv-NmbrL_5 |
| a2 | SPU_020055 | Lv-C14orf159L |
| a2 | SPU_020062 | Lv-Rasgrp3 |
| a2 | SPU_020072 | Lv-Hypp_163 |
| a2 | SPU_020163 | NA |
| a2 | SPU_020179 | Lv-Rpain |
| a2 | SPU_020196 | Lv-Nup35 |
| a2 | SPU_020211 | Lv-Slc7a4_1 |
| a2 | SPU_020213 | Lv-Mglur5 |
| a2 | SPU_020230 | NA |
| a2 | SPU_020332 | Lv-Slc24a3 |
| a2 | SPU_020344 | none |
| a2 | SPU_020434 | Lv-Ankle2 |
| a2 | SPU_020441 | Lv-Uhfr1 |
| a2 | SPU_020459 | Lv-Disp1_2 |
| a2 | SPU_020465 | Lv-Mapkapk2/3 |
| a2 | SPU_020501 | Lv-Plc |
| a2 | SPU_020646 | Lv-RorL |
| a2 | SPU_020702 | Lv-Sash1 |
| a2 | SPU_020740 | Lv-Eda2rL1 |
| a2 | SPU_020774 | Lv-Endrvt40 |
| a2 | SPU_020786 | Lv-Hypp_993 |
| a2 | SPU_020788 | Lv-Pigm |
| a2 | SPU_020897 | Lv-HhatL |
| a2 | SPU_020942 | none |
| a2 | SPU_020985 | Lv-Nxf1_1 |
| a2 | SPU_021160 | Lv-NeurmdnBr_3 |
| a2 | SPU_021191 | Lv-Recql4 |
| a2 | SPU_021198 | Lv-Sirt5_1 |
| a2 | SPU_021214 | NA |
| a2 | SPU_021290 | Lv-Echtcnr |
| a2 | SPU_021317 | Lv-Krp180 |
| a2 | SPU_021324 | none |

| | | |
|---|---|---|
| a2 | SPU_021488 | Lv-Prim2L |
| a2 | SPU_021501 | Lv-Ncapd3-2 |
| a2 | SPU_021517 | Lv-Dclr2 |
| a2 | SPU_021624 | Lv-TpaseL_12 |
| a2 | SPU_021697 | Lv-Dennd4a_2 |
| a2 | SPU_021790 | Lv-122B_1 |
| a2 | SPU_021804 | Lv-Enoyl/CoA/hydrt |
| a2 | SPU_021817 | Lv-Sod2a |
| a2 | SPU_021822 | Lv-Pric285_5 |
| a2 | SPU_021853 | Lv-Tsp1/Ig |
| a2 | SPU_021855 | Lv-Hypp_2579 |
| a2 | SPU_021867 | Lv-Lct |
| a2 | SPU_021941 | Lv-Inpp1_1 |
| a2 | SPU_021950 | none |
| a2 | SPU_021986 | Lv-Angel |
| a2 | SPU_022070 | Lv-Nlr158 |
| a2 | SPU_022141 | Lv-Nufil1L |
| a2 | SPU_022215 | Lv-Ggh |
| a2 | SPU_022231 | none |
| a2 | SPU_022241 | Lv-Pld3 |
| a2 | SPU_022250 | Lv-Lrr/Igr |
| a2 | SPU_022335 | none |
| a2 | SPU_022361 | Lv-Dagla |
| a2 | SPU_022392 | Lv-MaelL |
| a2 | SPU_022517 | Lv-Ankrd16 |
| a2 | SPU_022608 | none |
| a2 | SPU_022625 | Lv-Clspnh |
| a2 | SPU_022678 | Lv-Rora |
| a2 | SPU_022722 | none |
| a2 | SPU_022745 | Lv-Ccdc41 |
| a2 | SPU_022789 | Lv-Lrrcc1_1 |
| a2 | SPU_022827 | Lv-Rint1 |
| a2 | SPU_022834 | Lv-Rhop4 |
| a2 | SPU_022871 | none |
| a2 | SPU_022945 | Lv-Hypp_2640 |
| a2 | SPU_023024 | Lv-Mtf2 |
| a2 | SPU_023124 | Lv-Agpat1_3 |
| a2 | SPU_023155 | none |

| | | |
|---|---|---|
| a2 | SPU_023156 | Lv-C9orf72L |
| a2 | SPU_023164 | none |
| a2 | SPU_023180 | none |
| a2 | SPU_023198 | Lv-Usp36 |
| a2 | SPU_023268 | Lv-Hypp_2662 |
| a2 | SPU_023288 | Lv-Hypp_2210 |
| a2 | SPU_023442 | Lv-3oxoacylcs |
| a2 | SPU_023496 | Lv-Ghdc1_7 |
| a2 | SPU_023500 | Lv-Meid1 |
| a2 | SPU_023504 | Lv-Mib |
| a2 | SPU_023532 | Lv-Nlr6 |
| a2 | SPU_023570 | Lv-Rnf145 |
| a2 | SPU_023593 | Lv-Myhpp11_smph1 |
| a2 | SPU_023596 | Lv-C1orf112L |
| a2 | SPU_023597 | Lv-Slc16a14L_2 |
| a2 | SPU_023627 | none |
| a2 | SPU_023657 | Lv-Hypp_2681 |
| a2 | SPU_023739 | Lv-Pbx_1 |
| a2 | SPU_023817 | NA |
| a2 | SPU_023867 | Lv-Couptf1 |
| a2 | SPU_023995 | NA |
| a2 | SPU_024037 | Lv-C3orf19_1 |
| a2 | SPU_024171 | Lv-Ankfy1-2 |
| a2 | SPU_024172 | Lv-Ankfy1 |
| a2 | SPU_024212 | Lv-Chrna7L |
| a2 | SPU_024463 | Lv-Hypp_2734 |
| a2 | SPU_024485 | Lv-Cdc40 |
| a2 | SPU_024531 | none |
| a2 | SPU_024567 | Lv-ClvhH1 |
| a2 | SPU_024575 | Lv-Crhr1_1 |
| a2 | SPU_024645 | Lv-Gpatch3 |
| a2 | SPU_024671 | Lv-Hypp_2746 |
| a2 | SPU_024780 | Lv-Slbp_1 |
| a2 | SPU_024818 | Lv-Rad54b |
| a2 | SPU_024832 | NA |
| a2 | SPU_024930 | Lv-Msk_1 |
| a2 | SPU_024932 | Lv-Hypp_1131 |
| a2 | SPU_024934 | Lv-Hypp_1132 |

| | | |
|---|---|---|
| a2 | SPU_024957 | Lv-Rndl1 |
| a2 | SPU_024966 | none |
| a2 | SPU_024969 | Lv-Rgs/Px1 |
| a2 | SPU_024970 | Lv-Gpt2 |
| a2 | SPU_025017 | Lv-C1qL_3 |
| a2 | SPU_025023 | Lv-Aadac_2 |
| a2 | SPU_025026 | Lv-Hypp_2764 |
| a2 | SPU_025031 | none |
| a2 | SPU_025067 | none |
| a2 | SPU_025152 | none |
| a2 | SPU_025160 | Lv-Bckdk |
| a2 | SPU_025174 | none |
| a2 | SPU_025183 | Lv-Hypp_2777 |
| a2 | SPU_025191 | Lv-Hypp_1142 |
| a2 | SPU_025354 | Lv-Hypp_228 |
| a2 | SPU_025393 | none |
| a2 | SPU_025451 | Lv-Flcn |
| a2 | SPU_025503 | none |
| a2 | SPU_025611 | Lv-Ate1 |
| a2 | SPU_025626 | Lv-Sgcb |
| a2 | SPU_025659 | Lv-FanccL_1 |
| a2 | SPU_025661 | Lv-Grm3 |
| a2 | SPU_025698 | none |
| a2 | SPU_025725 | Lv-Trpt1 |
| a2 | SPU_025748 | Lv-Fignl1 |
| a2 | SPU_025750 | Lv-Slc16a6L_9 |
| a2 | SPU_025761 | Lv-Parn |
| a2 | SPU_025790 | Lv-Brdg1 |
| a2 | SPU_025833 | Lv-Ttvgcac_1 |
| a2 | SPU_025881 | NA |
| a2 | SPU_026079 | Lv-Yank |
| a2 | SPU_026091 | Lv-Hypp_2816 |
| a2 | SPU_026095 | Lv-Zpd |
| a2 | SPU_026150 | Lv-Hypp_1170 |
| a2 | SPU_026307 | Lv-Fn3/Igf_23 |
| a2 | SPU_026336 | Lv-Dhx34 |
| a2 | SPU_026349 | Lv-Fn3f_14 |
| a2 | SPU_026431 | Lv-Sema/Psi_1 |

| | | |
|---|---|---|
| a2 | SPU_026448 | Lv-Grtp1 |
| a2 | SPU_026455 | Lv-Gpr54L_5 |
| a2 | SPU_026495 | Lv-Traf3 |
| a2 | SPU_026510 | Lv-Z74 |
| a2 | SPU_026553 | Lv-Shoc2L |
| a2 | SPU_026576 | Lv-Cycb_1 |
| a2 | SPU_026730 | Lv-CckbrL_4 |
| a2 | SPU_026760 | Lv-Hypp_2867 |
| a2 | SPU_026784 | NA |
| a2 | SPU_026785 | Lv-Hypp_260 |
| a2 | SPU_026856 | Lv-GrprL |
| a2 | SPU_026857 | Lv-Slco4a1_10 |
| a2 | SPU_026893 | NA |
| a2 | SPU_026905 | Lv-Atf2 |
| a2 | SPU_026945 | Lv-Plcl2 |
| a2 | SPU_027053 | Lv-Ets1-2 |
| a2 | SPU_027062 | Lv-Mett11_1 |
| a2 | SPU_027064 | Lv-Apba1_1 |
| a2 | SPU_027120 | none |
| a2 | SPU_027147 | Lv-Riz |
| a2 | SPU_027218 | Lv-Trx |
| a2 | SPU_027220 | none |
| a2 | SPU_027270 | Lv-Trmt61bL |
| a2 | SPU_027276 | none |
| a2 | SPU_027287 | Lv-Srcr218 |
| a2 | SPU_027303 | Lv-Gatsl2 |
| a2 | SPU_027368 | Lv-Mtmr15L |
| a2 | SPU_027408 | Lv-Exosc10 |
| a2 | SPU_027469 | Lv-Hypp_268 |
| a2 | SPU_027582 | Lv-Siatf |
| a2 | SPU_027638 | Lv-Hypp_1207 |
| a2 | SPU_027845 | Lv-Mpped1 |
| a2 | SPU_027898 | Lv-Hypp_277 |
| a2 | SPU_027953 | Lv-Ppp4c |
| a2 | SPU_027970 | Lv-Gfer |
| a2 | SPU_028047 | Lv-Hypp_3027 |
| a2 | SPU_028092 | none |
| a2 | SPU_028094 | Lv-Rhogef10 |

| | | |
|---|---|---|
| a2 | SPU_028139 | Lv-Bcdo2 |
| a2 | SPU_028149 | Lv-Tkyr |
| a2 | SPU_028162 | NA |
| a2 | SPU_028166 | Lv-Lnp |
| a2 | SPU_028176 | Lv-Slc11a2 |
| a2 | SPU_028218 | Lv-Dicer |
| a2 | SPU_028227 | Lv-Hypp_1223 |
| a2 | SPU_028238 | Lv-Inpp1 |
| a2 | SPU_028313 | Lv-Hypp_1226 |
| a2 | SPU_028460 | Lv-Rufy3 |
| a2 | SPU_028462 | Lv-Flj13072L |
| a2 | SPU_028463 | Lv-Mapkapk5b |
| a2 | SPU_028491 | Lv-Pts |
| a2 | SPU_028494 | Lv-Serrp14 |
| a2 | SPU_028563 | Lv-RabO4b |
| a2 | SPU_028580 | none |
| a2 | SPU_028695 | Lv-Gen1L_2 |
| a2 | SPU_028741 | none |
| a2 | SPU_028776 | Lv-Cenpk |
| a2 | SPU_028777 | Lv-Lemd3 |
| a2 | SPU_028800 | Lv-Bks |
| a2 | SPU_028807 | Lv-Oca2 |
| a2 | SPU_028892 | Lv-IppL |
| a3 | SPU_000096 | Lv-Orbph |
| a3 | SPU_000097 | none |
| a3 | SPU_000217 | Lv-EspL1 |
| a3 | SPU_000228 | Lv-Hcfc2 |
| a3 | SPU_000338 | NA |
| a3 | SPU_000388 | Lv-Hox11/13c |
| a3 | SPU_000569 | NA |
| a3 | SPU_000652 | Lv-Dusp14/18/21 |
| a3 | SPU_000754 | none |
| a3 | SPU_000760 | Lv-Smpd2L |
| a3 | SPU_000767 | Lv-Chrna9-2 |
| a3 | SPU_000790 | Lv-Nars2 |
| a3 | SPU_000794 | Lv-C5orf42L |
| a3 | SPU_000797 | Lv-Fn3_13 |
| a3 | SPU_000898 | Lv-Hypp_316 |

| | | |
|---|---|---|
| a3 | SPU_000933 | none |
| a3 | SPU_001067 | none |
| a3 | SPU_001144 | Lv-Ddo |
| a3 | SPU_001159 | Lv-Fn3/Egff_3 |
| a3 | SPU_001166 | Lv-Gpr112L_8 |
| a3 | SPU_001192 | Lv-Pan3 |
| a3 | SPU_001204 | Lv-Ccdc49 |
| a3 | SPU_001207 | NA |
| a3 | SPU_001253 | NA |
| a3 | SPU_001266 | Lv-Srcr14 |
| a3 | SPU_001289 | Lv-Pddc1 |
| a3 | SPU_001450 | Lv-Rfp150L |
| a3 | SPU_001492 | none |
| a3 | SPU_001514 | Lv-Gyltl1b |
| a3 | SPU_001547 | Lv-Hypp_1335 |
| a3 | SPU_001570 | NA |
| a3 | SPU_001604 | none |
| a3 | SPU_001719 | Lv-Eepd1 |
| a3 | SPU_002041 | Lv-Srcr22 |
| a3 | SPU_002080 | Lv-NotchL_8 |
| a3 | SPU_002123 | Lv-Spop |
| a3 | SPU_002130 | Lv-Rnpc3 |
| a3 | SPU_002444 | Lv-Syt9L |
| a3 | SPU_002480 | Lv-Nupl1 |
| a3 | SPU_002554 | Lv-Z75 |
| a3 | SPU_002687 | Lv-Bat3 |
| a3 | SPU_002842 | Lv-Papd5 |
| a3 | SPU_002878 | Lv-Ofd1L |
| a3 | SPU_002894 | none |
| a3 | SPU_002897 | Lv-Tie1L_3 |
| a3 | SPU_003128 | Lv-PtL |
| a3 | SPU_003153 | Lv-Dirc2 |
| a3 | SPU_003171 | Lv-Ly/Cub/Sr/Zp |
| a3 | SPU_003364 | Lv-Hypp_380 |
| a3 | SPU_003449 | Lv-Hypp_3089 |
| a3 | SPU_003502 | Lv-Hcfc1 |
| a3 | SPU_003518 | Lv-Znhit1 |
| a3 | SPU_003581 | Lv-Ptprmp |

| | | |
|---|---|---|
| a3 | SPU_003687 | Lv-Cyp46L3 |
| a3 | SPU_003690 | Lv-Zfp106L |
| a3 | SPU_003748 | Lv-Mov10l |
| a3 | SPU_003954 | none |
| a3 | SPU_003984 | Lv-Bursicona |
| a3 | SPU_003990 | Lv-Mgat2 |
| a3 | SPU_004203 | Lv-Klhdc8a |
| a3 | SPU_004205 | Lv-Hypp_1506 |
| a3 | SPU_004325 | Lv-Fastkd1 |
| a3 | SPU_004418 | Lv-Lrrc45 |
| a3 | SPU_004436 | Lv-Tmc7 |
| a3 | SPU_004441 | Lv-Tmem185a |
| a3 | SPU_004456 | Lv-Impf |
| a3 | SPU_004563 | NA |
| a3 | SPU_004575 | Lv-Dis3L2 |
| a3 | SPU_004658 | none |
| a3 | SPU_004787 | none |
| a3 | SPU_004881 | Lv-Nhedc2 |
| a3 | SPU_004895 | Lv-Egf/LamGap1 |
| a3 | SPU_004902 | Lv-Ide |
| a3 | SPU_004906 | Lv-Hypp_1552 |
| a3 | SPU_004939 | Lv-Kif25L2 |
| a3 | SPU_005021 | Lv-CycK_1 |
| a3 | SPU_005119 | Lv-Slc23a1 |
| a3 | SPU_005125 | Lv-Hypp_3126 |
| a3 | SPU_005139 | none |
| a3 | SPU_005185 | Lv-Herc4L_2 |
| a3 | SPU_005254 | Lv-Tab1 |
| a3 | SPU_005274 | Lv-Sptlc2 |
| a3 | SPU_005281 | Lv-Kiaa0317L |
| a3 | SPU_005285 | Lv-Gch1 |
| a3 | SPU_005358 | Lv-Creb |
| a3 | SPU_005370 | Lv-Gcn1l1 |
| a3 | SPU_005386 | Lv-Zmat5 |
| a3 | SPU_005394 | none |
| a3 | SPU_005629 | Lv-Pmfbp1L |
| a3 | SPU_005650 | Lv-Galnt10 |
| a3 | SPU_005666 | Lv-AmidL_1 |

| | | |
|---|---|---|
| a3 | SPU_005812 | Lv-Hypp_1605 |
| a3 | SPU_005821 | none |
| a3 | SPU_005834 | Lv-Cts1 |
| a3 | SPU_005848 | Lv-Tmprss6 |
| a3 | SPU_005956 | none |
| a3 | SPU_005958 | Lv-Dnajhb12 |
| a3 | SPU_005986 | Lv-2OGFeII |
| a3 | SPU_006020 | Lv-Adam/TsL9 |
| a3 | SPU_006156 | Lv-Hypp_470 |
| a3 | SPU_006175 | Lv-C10orf88L |
| a3 | SPU_006274 | Lv-Hypp_3078 |
| a3 | SPU_006282 | Lv-Lim2L |
| a3 | SPU_006441 | Lv-Neu1_1 |
| a3 | SPU_006444 | Lv-Cycb3 |
| a3 | SPU_006555 | Lv-C14orf145L |
| a3 | SPU_006571 | none |
| a3 | SPU_006576 | Lv-Spr |
| a3 | SPU_006638 | Lv-Dbr1L |
| a3 | SPU_007091 | Lv-Obfc2a |
| a3 | SPU_007108 | Lv-Z67 |
| a3 | SPU_007141 | Lv-Gpld1 |
| a3 | SPU_007261 | Lv-PtpN13 |
| a3 | SPU_007264 | Lv-Notchh1_6 |
| a3 | SPU_007344 | Lv-Mus81h |
| a3 | SPU_007360 | Lv-Z129 |
| a3 | SPU_007361 | Lv-Z144 |
| a3 | SPU_007365 | Lv-Nbeal1_1 |
| a3 | SPU_007429 | Lv-Tlr037 |
| a3 | SPU_007448 | Lv-Lpp60 |
| a3 | SPU_007460 | none |
| a3 | SPU_007508 | Lv-Qsk |
| a3 | SPU_007512 | Lv-Siae |
| a3 | SPU_007556 | none |
| a3 | SPU_007691 | Lv-Hypp_2988 |
| a3 | SPU_007715 | Lv-Egf/Ig/Fn3 |
| a3 | SPU_007729 | NA |
| a3 | SPU_007744 | NA |
| a3 | SPU_007753 | Lv-Hypp_28 |

| | | |
|---|---|---|
| a3 | SPU_007809 | none |
| a3 | SPU_007956 | Lv-Pus1 |
| a3 | SPU_007973 | Lv-Ck1g |
| a3 | SPU_008008 | Lv-Serrp1 |
| a3 | SPU_008010 | Lv-Dis3l2_1 |
| a3 | SPU_008034 | Lv-Edem3 |
| a3 | SPU_008041 | Lv-Tbc1d9 |
| a3 | SPU_008066 | Lv-Clect/Fn3_9 |
| a3 | SPU_008181 | NA |
| a3 | SPU_008198 | Lv-C5orf37 |
| a3 | SPU_008221 | Lv-Tpx2L1 |
| a3 | SPU_008256 | Lv-Ubr5-2 |
| a3 | SPU_008292 | none |
| a3 | SPU_008315 | Lv-Pgrmc1 |
| a3 | SPU_008326 | NA |
| a3 | SPU_008345 | Lv-Ppfibp1 |
| a3 | SPU_008381 | Lv-Hypp_546 |
| a3 | SPU_008398 | Lv-Epb41l3 |
| a3 | SPU_008528 | Lv-Ets4 |
| a3 | SPU_008530 | Lv-Spire1 |
| a3 | SPU_008665 | none |
| a3 | SPU_008746 | Lv-Cspe6L |
| a3 | SPU_008965 | Lv-Chst14_1 |
| a3 | SPU_008995 | Lv-Hypp_564 |
| a3 | SPU_009263 | Lv-glysynk3b |
| a3 | SPU_009276 | none |
| a3 | SPU_009280 | Lv-Plxnb3L |
| a3 | SPU_009295 | Lv-TgshL |
| a3 | SPU_009363 | Lv-Hypp_1845 |
| a3 | SPU_009442 | Lv-Ctbs_3 |
| a3 | SPU_009463 | Lv-Hypp_46 |
| a3 | SPU_009464 | Lv-Hypp_47 |
| a3 | SPU_009497 | Lv-CspeN2a |
| a3 | SPU_009531 | Lv-Ugt2a2 |
| a3 | SPU_009595 | Lv-Zdhhc16 |
| a3 | SPU_009634 | Lv-Rnase3l_1 |
| a3 | SPU_009741 | none |
| a3 | SPU_009804 | NA |

| | | |
|---|---|---|
| a3 | SPU_009821 | NA |
| a3 | SPU_009938 | Lv-Edil3 |
| a3 | SPU_009955 | Lv-Usp19 |
| a3 | SPU_010113 | Lv-Tatdn2 |
| a3 | SPU_010135 | none |
| a3 | SPU_010146 | none |
| a3 | SPU_010225 | Lv-Hypp_66 |
| a3 | SPU_010287 | Lv-Exo1 |
| a3 | SPU_010292 | Lv-Lonrf3 |
| a3 | SPU_010329 | Lv-Ryk |
| a3 | SPU_010414 | Lv-Hypp_1917 |
| a3 | SPU_010430 | Lv-Unr |
| a3 | SPU_010439 | Lv-Dpy19l1 |
| a3 | SPU_010476 | none |
| a3 | SPU_010552 | none |
| a3 | SPU_010587 | Lv-Fn3/Egf_1 |
| a3 | SPU_010604 | Lv-Pnca |
| a3 | SPU_010643 | none |
| a3 | SPU_010698 | Lv-Ptk7 |
| a3 | SPU_010789 | Lv-Nbeal1 |
| a3 | SPU_010956 | none |
| a3 | SPU_010984 | Lv-Lmf1 |
| a3 | SPU_011020 | Lv-P76 |
| a3 | SPU_011120 | NA |
| a3 | SPU_011143 | Lv-Cecr1_1 |
| a3 | SPU_011200 | Lv-Fbxo28 |
| a3 | SPU_011220 | Lv-Nacha7 |
| a3 | SPU_011222 | Lv-Srcr101 |
| a3 | SPU_011226 | Lv-Z415 |
| a3 | SPU_011378 | Lv-Nudt8 |
| a3 | SPU_011551 | Lv-Bmp1/TldL2 |
| a3 | SPU_011574 | Lv-Edd |
| a3 | SPU_011629 | none |
| a3 | SPU_011723 | Lv-Add1L |
| a3 | SPU_011814 | Lv-Spryd3L |
| a3 | SPU_011839 | Lv-Cpsf3L |
| a3 | SPU_011885 | none |
| a3 | SPU_011921 | Lv-Apc5 |

| | | |
|---|---|---|
| a3 | SPU_012080 | Lv-Cyp4VL4 |
| a3 | SPU_012133 | Lv-Zbed1L |
| a3 | SPU_012171 | Lv-Rfx2 |
| a3 | SPU_012198 | Lv-Z124 |
| a3 | SPU_012206 | Lv-C1d |
| a3 | SPU_012212 | Lv-Fam57a |
| a3 | SPU_012294 | NA |
| a3 | SPU_012466 | Lv-Hypp_77 |
| a3 | SPU_012475 | Lv-Bdnf |
| a3 | SPU_012517 | none |
| a3 | SPU_012528 | Lv-Hypp_3060 |
| a3 | SPU_012558 | none |
| a3 | SPU_012734 | none |
| a3 | SPU_012756 | NA |
| a3 | SPU_012862 | Lv-Apaf1L_1 |
| a3 | SPU_012893 | none |
| a3 | SPU_012933 | none |
| a3 | SPU_012979 | Lv-MachrM3_1 |
| a3 | SPU_013075 | Lv-Ktelc1 |
| a3 | SPU_013145 | none |
| a3 | SPU_013430 | none |
| a3 | SPU_013522 | Lv-Sfk1 |
| a3 | SPU_013611 | Lv-Z192 |
| a3 | SPU_013747 | Lv-C1orf59L |
| a3 | SPU_013812 | NA |
| a3 | SPU_013813 | Lv-Xrn1_1 |
| a3 | SPU_014002 | Lv-Dctd |
| a3 | SPU_014045 | Lv-Z35 |
| a3 | SPU_014132 | Lv-Dpp7 |
| a3 | SPU_014249 | Lv-Trh |
| a3 | SPU_014363 | none |
| a3 | SPU_014373 | Lv-Gpx1 |
| a3 | SPU_014592 | Lv-Pofut1_2 |
| a3 | SPU_014635 | Lv-Hypp_2161 |
| a3 | SPU_014657 | Lv-Dis3 |
| a3 | SPU_014731 | Lv-Unk_35 |
| a3 | SPU_014791 | Lv-Gmps |
| a3 | SPU_015019 | Lv-TectaL |

| | | |
|---|---|---|
| a3 | SPU_015167 | none |
| a3 | SPU_015188 | none |
| a3 | SPU_015301 | Lv-2Pde_1 |
| a3 | SPU_015346 | none |
| a3 | SPU_015424 | Lv-Mdc2 |
| a3 | SPU_015458 | Lv-Slc16a12L_9 |
| a3 | SPU_015461 | none |
| a3 | SPU_015473 | Lv-Phdc |
| a3 | SPU_015765 | Lv-Cdc14A |
| a3 | SPU_015793 | Lv-Rbks |
| a3 | SPU_015853 | Lv-Trpc5 |
| a3 | SPU_016006 | Lv-Vrk |
| a3 | SPU_016085 | Lv-C19orf28L |
| a3 | SPU_016150 | Lv-Aer61gt(kinase) |
| a3 | SPU_016201 | NA |
| a3 | SPU_016234 | none |
| a3 | SPU_016272 | Lv-Alk3-6 |
| a3 | SPU_016291 | Lv-Elach2 |
| a3 | SPU_016292 | none |
| a3 | SPU_016340 | Lv-FancgL |
| a3 | SPU_016557 | Lv-Rfx1L |
| a3 | SPU_016727 | NA |
| a3 | SPU_016775 | Lv-Hypp_842 |
| a3 | SPU_016793 | Lv-Prpf31 |
| a3 | SPU_016807 | Lv-CrumbsL1 |
| a3 | SPU_016871 | Lv-Hypp_2285 |
| a3 | SPU_016914 | Lv-Opo |
| a3 | SPU_016995 | Lv-Esco2L |
| a3 | SPU_017042 | Lv-Gpr133L_4 |
| a3 | SPU_017135 | Lv-Tdrd7L |
| a3 | SPU_017136 | Lv-Rpap2 |
| a3 | SPU_017159 | Lv-Galnact2 |
| a3 | SPU_017251 | Lv-SdL |
| a3 | SPU_017301 | Lv-Lrrc58 |
| a3 | SPU_017309 | none |
| a3 | SPU_017311 | none |
| a3 | SPU_017329 | Lv-Sfe9 |
| a3 | SPU_017404 | Lv-Nr1h6 |

| | | |
|---|---|---|
| a3 | SPU_017500 | Lv-C3orf58 |
| a3 | SPU_017503 | Lv-Psme3 |
| a3 | SPU_017619 | Lv-Snx16 |
| a3 | SPU_017629 | none |
| a3 | SPU_017707 | Lv-Bursiconb |
| a3 | SPU_017749 | Lv-Ppactr2 |
| a3 | SPU_017816 | Lv-Pkd |
| a3 | SPU_017852 | none |
| a3 | SPU_017916 | Lv-Tdrd1 |
| a3 | SPU_017938 | Lv-Fn3_36 |
| a3 | SPU_017963 | Lv-Tnr |
| a3 | SPU_018030 | Lv-Fn3/Igf_15 |
| a3 | SPU_018123 | Lv-Trit1l |
| a3 | SPU_018175 | none |
| a3 | SPU_018309 | Lv-Ncapg |
| a3 | SPU_018342 | Lv-Abcb6 |
| a3 | SPU_018456 | none |
| a3 | SPU_018458 | Lv-Rpusd2 |
| a3 | SPU_018557 | Lv-St6 |
| a3 | SPU_018646 | Lv-Hypp_2399 |
| a3 | SPU_018666 | Lv-Zcchc11 |
| a3 | SPU_018795 | Lv-Sirt3_1 |
| a3 | SPU_018806 | Lv-Magi |
| a3 | SPU_018840 | Lv-Tm2d3 |
| a3 | SPU_018876 | Lv-Sult1c3 |
| a3 | SPU_018880 | none |
| a3 | SPU_019054 | Lv-Ankrd17 |
| a3 | SPU_019092 | none |
| a3 | SPU_019158 | Lv-Ldla/Gpcr_1 |
| a3 | SPU_019171 | Lv-Rictor_1 |
| a3 | SPU_019191 | Lv-Hypp_149 |
| a3 | SPU_019224 | Lv-Frk_1 |
| a3 | SPU_019343 | NA |
| a3 | SPU_019369 | Lv-Rndzvn |
| a3 | SPU_019418 | none |
| a3 | SPU_019472 | Lv-Mul1 |
| a3 | SPU_019473 | none |
| a3 | SPU_019493 | Lv-Hsd17b8 |

| | | |
|---|---|---|
| a3 | SPU_019512 | Lv-Ythdf8 |
| a3 | SPU_019579 | Lv-Imdzlprpn |
| a3 | SPU_019659 | Lv-Fn3/Egff_11 |
| a3 | SPU_019766 | none |
| a3 | SPU_019814 | none |
| a3 | SPU_019818 | Lv-Fn3f_42 |
| a3 | SPU_019819 | Lv-Nthl1 |
| a3 | SPU_019894 | NA |
| a3 | SPU_019899 | Lv-Cyp2L40 |
| a3 | SPU_019964 | Lv-Slc16a12L_15 |
| a3 | SPU_020053 | Lv-Smekh1 |
| a3 | SPU_020056 | none |
| a3 | SPU_020061 | Lv-Drap1 |
| a3 | SPU_020187 | Lv-Hypp_2493 |
| a3 | SPU_020254 | NA |
| a3 | SPU_020266 | Lv-Ppm1d |
| a3 | SPU_020320 | NA |
| a3 | SPU_020342 | Lv-BetaCi_1 |
| a3 | SPU_020366 | Lv-GtpbpL1 |
| a3 | SPU_020381 | Lv-Rxfp2L_5 |
| a3 | SPU_020445 | Lv-Agpat3_2 |
| a3 | SPU_020476 | Lv-Gpr177 |
| a3 | SPU_020493 | NA |
| a3 | SPU_020507 | Lv-Fma38b2 |
| a3 | SPU_020520 | Lv-Slc16a12L_16 |
| a3 | SPU_020568 | NA |
| a3 | SPU_020575 | Lv-Tbc1d30 |
| a3 | SPU_020604 | Lv-Ptprlec3 |
| a3 | SPU_020662 | Lv-AknaL |
| a3 | SPU_020789 | Lv-Zc3h18 |
| a3 | SPU_020852 | Lv-Rab27 |
| a3 | SPU_020873 | Lv-Hypp_3002 |
| a3 | SPU_020889 | Lv-Hypp_2525 |
| a3 | SPU_020910 | Lv-PlxnaL |
| a3 | SPU_021125 | Lv-Hnmt_1 |
| a3 | SPU_021168 | Lv-Utp23h |
| a3 | SPU_021200 | Lv-Whirlin |
| a3 | SPU_021259 | Lv-Dus3l |

| | | |
|---|---|---|
| a3 | SPU_021270 | NA |
| a3 | SPU_021319 | Lv-Sycp1L |
| a3 | SPU_021398 | Lv-Stk |
| a3 | SPU_021446 | Lv-Hypp_1024 |
| a3 | SPU_021505 | Lv-Clect_89 |
| a3 | SPU_021513 | Lv-EgfiiiL_1 |
| a3 | SPU_021519 | Lv-Hypp_2559 |
| a3 | SPU_021566 | Lv-Shrls |
| a3 | SPU_021638 | Lv-BetaDxlL |
| a3 | SPU_021792 | none |
| a3 | SPU_021808 | Lv-Camta1L |
| a3 | SPU_021837 | Lv-Ctdspl2 |
| a3 | SPU_021903 | Lv-Cwf19l2_1 |
| a3 | SPU_021946 | Lv-Mth1 |
| a3 | SPU_021959 | Lv-Grip1 |
| a3 | SPU_022011 | Lv-Pds5b |
| a3 | SPU_022017 | NA |
| a3 | SPU_022020 | Lv-Hypp_2592 |
| a3 | SPU_022048 | Lv-Clint1L |
| a3 | SPU_022190 | none |
| a3 | SPU_022290 | none |
| a3 | SPU_022329 | Lv-Ccdc62_1 |
| a3 | SPU_022367 | none |
| a3 | SPU_022487 | Lv-Z106 |
| a3 | SPU_022516 | Lv-Fbxo18-2 |
| a3 | SPU_022609 | Lv-Txlna |
| a3 | SPU_022662 | Lv-Qsox1 |
| a3 | SPU_022680 | Lv-Lrrkb |
| a3 | SPU_022697 | NA |
| a3 | SPU_022731 | none |
| a3 | SPU_022791 | Lv-Gtsf1 |
| a3 | SPU_022792 | Lv-Ehmt1-2 |
| a3 | SPU_023013 | Lv-Coq3_1 |
| a3 | SPU_023042 | Lv-Lace1 |
| a3 | SPU_023049 | Lv-Fam162a |
| a3 | SPU_023263 | none |
| a3 | SPU_023274 | Lv-Ccdc102a |
| a3 | SPU_023453 | none |

| | | |
|---|---|---|
| a3 | SPU_023590 | Lv-Rej4_1 |
| a3 | SPU_023640 | Lv-Jarid2 |
| a3 | SPU_023659 | Lv-Zc3h15 |
| a3 | SPU_023722 | none |
| a3 | SPU_023761 | none |
| a3 | SPU_023766 | Lv-Cchcr1 |
| a3 | SPU_023779 | Lv-Sec31a_1 |
| a3 | SPU_023790 | Lv-Pip5kl1 |
| a3 | SPU_023805 | none |
| a3 | SPU_023810 | Lv-Slc13a3 |
| a3 | SPU_023854 | Lv-Dlgap5 |
| a3 | SPU_023877 | Lv-Samd9L_9 |
| a3 | SPU_023887 | none |
| a3 | SPU_023907 | Lv-Tmem129 |
| a3 | SPU_023927 | none |
| a3 | SPU_023947 | Lv-Dnajc7 |
| a3 | SPU_023968 | Lv-Setd8 |
| a3 | SPU_023992 | Lv-Rcc1 |
| a3 | SPU_024019 | Lv-NephL |
| a3 | SPU_024022 | NA |
| a3 | SPU_024056 | Lv-Lrrcc1 |
| a3 | SPU_024059 | Lv-Golga3 |
| a3 | SPU_024122 | Lv-Dis3-2 |
| a3 | SPU_024173 | Lv-Scly |
| a3 | SPU_024219 | Lv-Hypp_2717 |
| a3 | SPU_024234 | Lv-Qprt_1 |
| a3 | SPU_024347 | Lv-Dnmt3A |
| a3 | SPU_024367 | Lv-Polyp_6 |
| a3 | SPU_024488 | Lv-Lonp2_1 |
| a3 | SPU_024598 | Lv-Mip1 |
| a3 | SPU_024634 | Lv-Manea |
| a3 | SPU_024647 | NA |
| a3 | SPU_024659 | Lv-Cdc14A_1 |
| a3 | SPU_024690 | none |
| a3 | SPU_024785 | Lv-Abcg2C |
| a3 | SPU_024803 | NA |
| a3 | SPU_024814 | Lv-Hfm1 |
| a3 | SPU_024904 | Lv-Hypp_2757 |

| | | |
|---|---|---|
| a3 | SPU_024933 | Lv-Rrm2_1 |
| a3 | SPU_024987 | NA |
| a3 | SPU_025049 | Lv-Lsm10 |
| a3 | SPU_025093 | none |
| a3 | SPU_025095 | Lv-Hypp_2772 |
| a3 | SPU_025141 | Lv-Mbtps1 |
| a3 | SPU_025201 | Lv-Mib1_2 |
| a3 | SPU_025212 | Lv-Yp302 |
| a3 | SPU_025298 | Lv-Pigs2 |
| a3 | SPU_025301 | Lv-Gluld1_11 |
| a3 | SPU_025321 | NA |
| a3 | SPU_025411 | Lv-Gch1_1 |
| a3 | SPU_025504 | Lv-Mapkap1 |
| a3 | SPU_025721 | Lv-Hypp_2798 |
| a3 | SPU_025726 | Lv-Rtf1 |
| a3 | SPU_025783 | NA |
| a3 | SPU_025848 | NA |
| a3 | SPU_025849 | Lv-Z408_1 |
| a3 | SPU_025908 | Lv-Tmem183a |
| a3 | SPU_025963 | Lv-Pus7 |
| a3 | SPU_025993 | Lv-B3gntl1_1 |
| a3 | SPU_026032 | none |
| a3 | SPU_026063 | Lv-Ccdc76 |
| a3 | SPU_026101 | Lv-Pp4r1_1 |
| a3 | SPU_026156 | Lv-Nadk |
| a3 | SPU_026196 | none |
| a3 | SPU_026272 | Lv-Tk1_1 |
| a3 | SPU_026301 | Lv-Gpbp1L |
| a3 | SPU_026339 | Lv-TfIIh_p44L_1 |
| a3 | SPU_026471 | Lv-Egln1 |
| a3 | SPU_026521 | Lv-Addph |
| a3 | SPU_026592 | Lv-Dnajhc22 |
| a3 | SPU_026597 | Lv-Fam13A1 |
| a3 | SPU_026625 | Lv-Edem1 |
| a3 | SPU_026729 | Lv-Orpk1L_10 |
| a3 | SPU_026737 | Lv-Fhit |
| a3 | SPU_026774 | Lv-Ncapd2 |
| a3 | SPU_026788 | Lv-Dcp2 |

| | | |
|---|---|---|
| a3 | SPU_026850 | Lv-Slc16a6L_11 |
| a3 | SPU_026943 | Lv-Slc2a8 |
| a3 | SPU_027212 | Lv-Npffr1L |
| a3 | SPU_027260 | Lv-PppL_154 |
| a3 | SPU_027310 | Lv-Vps13c_2 |
| a3 | SPU_027311 | Lv-Vgll2L |
| a3 | SPU_027417 | Lv-Hypp_2901 |
| a3 | SPU_027561 | Lv-Isyna1 |
| a3 | SPU_027618 | Lv-Gt |
| a3 | SPU_027657 | Lv-Aip |
| a3 | SPU_027736 | Lv-Snupn |
| a3 | SPU_027792 | Lv-Pex16 |
| a3 | SPU_027872 | Lv-Alg1h |
| a3 | SPU_027975 | Lv-Pms2 |
| a3 | SPU_028217 | Lv-Tmem62 |
| a3 | SPU_028289 | Lv-Slc22a1 |
| a3 | SPU_028326 | Lv-Clect/Ldla_4 |
| a3 | SPU_028329 | none |
| a3 | SPU_028436 | Lv-Smyd3_3 |
| a3 | SPU_028444 | Lv-Sarox_4 |
| a3 | SPU_028521 | Lv-Gfm2 |
| a3 | SPU_028573 | Lv-Slc30a7 |
| a3 | SPU_028600 | Lv-Hypp_288 |
| a3 | SPU_028656 | Lv-EhH4_80 |
| a3 | SPU_028734 | Lv-Hypp_1237 |
| a3 | SPU_028865 | Lv-Mov10 |
| a3 | SPU_028935 | Lv-Slsp_27 |
| a3 | SPU_030090 | Lv-Prtelsn_1 |
| a4 | SPU_000029 | Lv-ErtL |
| a4 | SPU_000127 | Lv-Usp3 |
| a4 | SPU_000145 | Lv-Aimf2l |
| a4 | SPU_000146 | Lv-Kpnb1 |
| a4 | SPU_000227 | Lv-Alg10bh |
| a4 | SPU_000251 | Lv-Tmem41a |
| a4 | SPU_000256 | Lv-Becn1 |
| a4 | SPU_000316 | Lv-Inpp4b |
| a4 | SPU_000408 | NA |
| a4 | SPU_000501 | Lv-Alms1L |

| | | |
|---|---|---|
| a4 | SPU_000631 | Lv-Tex261L |
| a4 | SPU_000655 | none |
| a4 | SPU_001072 | Lv-Axin |
| a4 | SPU_001130 | Lv-Ovca2 |
| a4 | SPU_001360 | Lv-KatB |
| a4 | SPU_001508 | Lv-Atad2 |
| a4 | SPU_001555 | Lv-Inturned |
| a4 | SPU_001642 | Lv-RtL_11 |
| a4 | SPU_001744 | NA |
| a4 | SPU_001952 | Lv-Hypp_340 |
| a4 | SPU_002008 | Lv-Z480 |
| a4 | SPU_002009 | Lv-Z480_1 |
| a4 | SPU_002066 | Lv-Wdr22 |
| a4 | SPU_002143 | Lv-Hypp_1374 |
| a4 | SPU_002226 | Lv-Slc39a3 |
| a4 | SPU_002282 | Lv-Rev3 |
| a4 | SPU_002307 | Lv-Rhogef7 |
| a4 | SPU_002484 | NA |
| a4 | SPU_002847 | none |
| a4 | SPU_002851 | Lv-Usp9Y |
| a4 | SPU_002861 | Lv-Clect/Egf/hyalin |
| a4 | SPU_002941 | Lv-Mtss1L |
| a4 | SPU_003008 | Lv-Tbc1d1 |
| a4 | SPU_003152 | Lv-Fbw7L1 |
| a4 | SPU_003258 | Lv-Macf1 |
| a4 | SPU_003411 | NA |
| a4 | SPU_003558 | none |
| a4 | SPU_003580 | Lv-Siat7e_1 |
| a4 | SPU_003614 | Lv-Ino80d |
| a4 | SPU_003791 | Lv-Wbscr16L |
| a4 | SPU_003864 | Lv-Wdfy3_1 |
| a4 | SPU_003910 | Lv-Polrmt |
| a4 | SPU_003955 | Lv-Tab2/3 |
| a4 | SPU_003971 | Lv-Fam179b |
| a4 | SPU_004037 | Lv-Csk2 |
| a4 | SPU_004147 | Lv-C1orf9L |
| a4 | SPU_004287 | Lv-Smad4 |
| a4 | SPU_004323 | Lv-Usp20L |

| | | |
|---|---|---|
| a4 | SPU_004340 | Lv-Smooth |
| a4 | SPU_004375 | Lv-Abca3L |
| a4 | SPU_004534 | Lv-Fam135a |
| a4 | SPU_004591 | Lv-Sntg2 |
| a4 | SPU_004779 | Lv-Hypp_3084 |
| a4 | SPU_004894 | none |
| a4 | SPU_004932 | none |
| a4 | SPU_005107 | Lv-Ubc9 |
| a4 | SPU_005116 | Lv-Cadm1bL |
| a4 | SPU_005266 | Lv-Rsu1L |
| a4 | SPU_005276 | Lv-Pog |
| a4 | SPU_005470 | Lv-Hypp_448 |
| a4 | SPU_005515 | Lv-Samd9L_5 |
| a4 | SPU_005599 | Lv-Tmem138 |
| a4 | SPU_005604 | Lv-Hypp_457 |
| a4 | SPU_005796 | Lv-PppL_184 |
| a4 | SPU_005804 | Lv-Hypp_1603 |
| a4 | SPU_005847 | Lv-Sorl1L |
| a4 | SPU_005963 | Lv-Hmgcr |
| a4 | SPU_006024 | Lv-Igf_5 |
| a4 | SPU_006026 | Lv-Lmr |
| a4 | SPU_006027 | Lv-Syngr3 |
| a4 | SPU_006073 | Lv-PolppL_54 |
| a4 | SPU_006304 | Lv-Hypp_1654 |
| a4 | SPU_006357 | Lv-Hypp_477 |
| a4 | SPU_006666 | Lv-Ing4 |
| a4 | SPU_006755 | Lv-Cul5 |
| a4 | SPU_006972 | none |
| a4 | SPU_007054 | Lv-Mmaa |
| a4 | SPU_007302 | Lv-CtnaL1 |
| a4 | SPU_007306 | Lv-Cyp2L15 |
| a4 | SPU_007408 | Lv-Hypp_515 |
| a4 | SPU_007481 | Lv-Seh1L |
| a4 | SPU_007551 | Lv-SansL |
| a4 | SPU_007557 | Lv-Svlnph |
| a4 | SPU_007699 | none |
| a4 | SPU_007763 | Lv-Mucpdn |
| a4 | SPU_007940 | Lv-Daglb |

| | | |
|---|---|---|
| a4 | SPU_007964 | Lv-Pias1 |
| a4 | SPU_007965 | Lv-Cep120 |
| a4 | SPU_007978 | Lv-Kiaa0892 |
| a4 | SPU_008062 | none |
| a4 | SPU_008173 | Lv-Suvar_3-9 |
| a4 | SPU_008238 | Lv-Dnajc28 |
| a4 | SPU_008258 | Lv-Ifrd2 |
| a4 | SPU_008280 | Lv-Myt1_1 |
| a4 | SPU_008324 | Lv-Hypp_542 |
| a4 | SPU_008368 | Lv-Sptan1L |
| a4 | SPU_008507 | none |
| a4 | SPU_008562 | NA |
| a4 | SPU_008600 | NA |
| a4 | SPU_008662 | Lv-Fbxo4 |
| a4 | SPU_008710 | Lv-Wdr48 |
| a4 | SPU_008736 | Lv-Usp48b |
| a4 | SPU_008803 | Lv-Slc7a11_8 |
| a4 | SPU_008831 | NA |
| a4 | SPU_008846 | none |
| a4 | SPU_008858 | Lv-Timp1 |
| a4 | SPU_008924 | Lv-Hypp_563 |
| a4 | SPU_009283 | Lv-Topbp1 |
| a4 | SPU_009324 | Lv-Pard3L_2 |
| a4 | SPU_009410 | Lv-Fam179b-2 |
| a4 | SPU_009525 | Lv-Limd1 |
| a4 | SPU_009558 | Lv-Usp44 |
| a4 | SPU_009615 | Lv-Clect_71 |
| a4 | SPU_009851 | Lv-Ogfod1 |
| a4 | SPU_009856 | Lv-Eps8 |
| a4 | SPU_009857 | Lv-Hypp_1876 |
| a4 | SPU_009869 | Lv-Cntrn |
| a4 | SPU_009870 | Lv-Dnajhc9 |
| a4 | SPU_009878 | Lv-Dcamkl2 |
| a4 | SPU_009881 | Lv-Kiaa1632L_1 |
| a4 | SPU_009882 | Lv-Kiaa1632h |
| a4 | SPU_009985 | Lv-CycG |
| a4 | SPU_010199 | Lv-Mect1L |
| a4 | SPU_010277 | Lv-Spca |

| | | |
|---|---|---|
| a4 | SPU_010326 | Lv-Hypp_614 |
| a4 | SPU_010550 | none |
| a4 | SPU_010620 | Lv-Exoc3_1 |
| a4 | SPU_010706 | Lv-Nsmafl |
| a4 | SPU_011072 | Lv-Atm_2 |
| a4 | SPU_011087 | Lv-Hdac4 |
| a4 | SPU_011259 | Lv-Styx |
| a4 | SPU_011298 | Lv-Socs6L |
| a4 | SPU_011380 | Lv-Z368 |
| a4 | SPU_011467 | Lv-Rab40 |
| a4 | SPU_011517 | none |
| a4 | SPU_011581 | Lv-Hypp_72 |
| a4 | SPU_011701 | Lv-Rabgap1 |
| a4 | SPU_011849 | none |
| a4 | SPU_012057 | Lv-Rt33 |
| a4 | SPU_012235 | Lv-Rg9mtd2 |
| a4 | SPU_012299 | Lv-Rhpn1 |
| a4 | SPU_012380 | Lv-Pargl |
| a4 | SPU_012494 | Lv-Lrr/Lnb/7Tm/Gpcr |
| a4 | SPU_012577 | Lv-Tsen34L |
| a4 | SPU_012731 | Lv-Fam126a |
| a4 | SPU_012743 | Lv-Ars_3 |
| a4 | SPU_013280 | Lv-PppL_197 |
| a4 | SPU_013281 | Lv-Eftud1L3 |
| a4 | SPU_013497 | Lv-Fn3/Igf_21 |
| a4 | SPU_013643 | Lv-Hypp_94 |
| a4 | SPU_013715 | Lv-Nup155 |
| a4 | SPU_013767 | Lv-Htr1b_1 |
| a4 | SPU_013835 | Lv-Usp34 |
| a4 | SPU_013850 | NA |
| a4 | SPU_013880 | Lv-Vac8 |
| a4 | SPU_013913 | Lv-Wdfy2 |
| a4 | SPU_014035 | NA |
| a4 | SPU_014223 | none |
| a4 | SPU_014226 | Lv-Lim2 |
| a4 | SPU_014232 | Lv-Cops8 |
| a4 | SPU_014293 | Lv-Pim-2 |
| a4 | SPU_014341 | NA |

| | | |
|---|---|---|
| a4 | SPU_014416 | Lv-Topbp1 |
| a4 | SPU_014417 | NA |
| a4 | SPU_014442 | NA |
| a4 | SPU_014561 | Lv-Hypp_3020 |
| a4 | SPU_014687 | Lv-Z494 |
| a4 | SPU_014803 | Lv-Hypp_104 |
| a4 | SPU_014989 | Lv-Cycl1 |
| a4 | SPU_015008 | Lv-Fn3/Egf/Cub |
| a4 | SPU_015035 | none |
| a4 | SPU_015057 | NA |
| a4 | SPU_015412 | Lv-Whrn |
| a4 | SPU_015419 | Lv-Pask_1 |
| a4 | SPU_015505 | Lv-Rnf40 |
| a4 | SPU_015540 | Lv-Smyd4 |
| a4 | SPU_015545 | Lv-Shark |
| a4 | SPU_015669 | Lv-Hypp_797 |
| a4 | SPU_015683 | Lv-Z326 |
| a4 | SPU_015858 | Lv-Hypp_2235 |
| a4 | SPU_016040 | Lv-Lnx2_1 |
| a4 | SPU_016137 | none |
| a4 | SPU_016633 | Lv-Cog1 |
| a4 | SPU_016683 | Lv-Hypp_839 |
| a4 | SPU_016734 | NA |
| a4 | SPU_016743 | Lv-Atl |
| a4 | SPU_016889 | Lv-Nphs1L_3 |
| a4 | SPU_016967 | Lv-Endrvt95 |
| a4 | SPU_016994 | Lv-Pap2dL |
| a4 | SPU_017036 | NA |
| a4 | SPU_017158 | Lv-Atad1 |
| a4 | SPU_017221 | none |
| a4 | SPU_017363 | Lv-Dna2h |
| a4 | SPU_017390 | Lv-PppL_116 |
| a4 | SPU_017473 | Lv-Ccr6a |
| a4 | SPU_017579 | Lv-GlisC |
| a4 | SPU_017734 | none |
| a4 | SPU_017750 | Lv-Z159 |
| a4 | SPU_017754 | Lv-Hypp_894 |
| a4 | SPU_017841 | Lv-Sfi1L |

| | | |
|---|---|---|
| a4 | SPU_017998 | Lv-Ercc1 |
| a4 | SPU_018026 | Lv-Stx12 |
| a4 | SPU_018065 | Lv-Wdr76 |
| a4 | SPU_018232 | Lv-Tmc7_1 |
| a4 | SPU_018306 | none |
| a4 | SPU_018317 | none |
| a4 | SPU_018326 | Lv-Clip170ph |
| a4 | SPU_018378 | Lv-Krp85 |
| a4 | SPU_018528 | none |
| a4 | SPU_018535 | Lv-Crnkl1 |
| a4 | SPU_018585 | Lv-C9orf125L |
| a4 | SPU_018718 | Lv-Donson |
| a4 | SPU_018723 | Lv-Sirt1 |
| a4 | SPU_018753 | NA |
| a4 | SPU_018772 | Lv-Gon4L |
| a4 | SPU_018787 | Lv-Schip1 |
| a4 | SPU_018807 | Lv-Erc1 |
| a4 | SPU_018809 | Lv-Nwd1_3 |
| a4 | SPU_018931 | none |
| a4 | SPU_019061 | Lv-Polr2dL |
| a4 | SPU_019371 | Lv-Tsc1L |
| a4 | SPU_019537 | Lv-Slsp_10 |
| a4 | SPU_019634 | Lv-Sqle_3 |
| a4 | SPU_019737 | Lv-Rfwd2 |
| a4 | SPU_019744 | NA |
| a4 | SPU_020054 | Lv-Daple |
| a4 | SPU_020060 | Lv-Xdh_1 |
| a4 | SPU_020067 | Lv-Polh-2 |
| a4 | SPU_020085 | none |
| a4 | SPU_020090 | none |
| a4 | SPU_020125 | Lv-Nocturnin |
| a4 | SPU_020239 | Lv-Tmco3 |
| a4 | SPU_020262 | Lv-Rabgef1 |
| a4 | SPU_020282 | Lv-Ift88 |
| a4 | SPU_020769 | none |
| a4 | SPU_020779 | Lv-Hypp_991 |
| a4 | SPU_020816 | Lv-Hbs1l |
| a4 | SPU_020835 | none |

| | | |
|---|---|---|
| a4 | SPU_020918 | Lv-Actinfilin |
| a4 | SPU_020938 | Lv-Insig1_1 |
| a4 | SPU_021066 | Lv-EhH2b_68 |
| a4 | SPU_021148 | Lv-Trpch |
| a4 | SPU_021184 | Lv-Abcg2A |
| a4 | SPU_021430 | Lv-Ncoa |
| a4 | SPU_021440 | Lv-PppL_128 |
| a4 | SPU_021503 | Lv-Kiaa1244L |
| a4 | SPU_021530 | none |
| a4 | SPU_021560 | none |
| a4 | SPU_021609 | Lv-CbpdEL_1 |
| a4 | SPU_021665 | Lv-Hypp_174 |
| a4 | SPU_021717 | Lv-Hypp_3031 |
| a4 | SPU_021910 | Lv-PolypL_954 |
| a4 | SPU_021948 | Lv-Ubr3_2 |
| a4 | SPU_022065 | none |
| a4 | SPU_022076 | Lv-Igfb2 |
| a4 | SPU_022131 | Lv-Hap1 |
| a4 | SPU_022185 | none |
| a4 | SPU_022243 | Lv-Tbc1d5 |
| a4 | SPU_022283 | Lv-Hiat1 |
| a4 | SPU_022315 | Lv-NotchL_92 |
| a4 | SPU_022339 | Lv-Srcr190 |
| a4 | SPU_022363 | Lv-Slc10a2L_4 |
| a4 | SPU_022410 | Lv-Tbcc |
| a4 | SPU_022468 | Lv-KpLr1 |
| a4 | SPU_022536 | none |
| a4 | SPU_022659 | Lv-MplL |
| a4 | SPU_023032 | Lv-Cdc45 |
| a4 | SPU_023044 | Lv-Lrp6 |
| a4 | SPU_023139 | Lv-Fhod3_1 |
| a4 | SPU_023287 | Lv-Garnl1 |
| a4 | SPU_023301 | Lv-Hypp_1080 |
| a4 | SPU_023507 | Lv-Tmem49 |
| a4 | SPU_023785 | Lv-Ccdc44 |
| a4 | SPU_023807 | none |
| a4 | SPU_023928 | Lv-Cca2dB |
| a4 | SPU_024197 | Lv-Endrvt_1 |

| | | |
|---|---|---|
| a4 | SPU_024351 | Lv-Rnf31_1 |
| a4 | SPU_024442 | none |
| a4 | SPU_024475 | Lv-Pon1_1 |
| a4 | SPU_024552 | Lv-Hypp_217 |
| a4 | SPU_024557 | Lv-Trpm3_6 |
| a4 | SPU_024587 | Lv-Tmem98 |
| a4 | SPU_024640 | Lv-Eif4enif1 |
| a4 | SPU_024697 | none |
| a4 | SPU_025169 | Lv-Stk33 |
| a4 | SPU_025242 | none |
| a4 | SPU_025257 | NA |
| a4 | SPU_025511 | Lv-Rit1 |
| a4 | SPU_025524 | Lv-Pi3kclassII |
| a4 | SPU_025573 | Lv-Kiaa0913L |
| a4 | SPU_025639 | Lv-Spg20 |
| a4 | SPU_025672 | Lv-Zfpl1 |
| a4 | SPU_025823 | none |
| a4 | SPU_026105 | none |
| a4 | SPU_026170 | Lv-Stac3 |
| a4 | SPU_026210 | Lv-Klhdc3 |
| a4 | SPU_026224 | Lv-Ube2g1 |
| a4 | SPU_026405 | Lv-Sass6 |
| a4 | SPU_026522 | Lv-Lzts2L |
| a4 | SPU_026610 | none |
| a4 | SPU_026732 | Lv-Hypp_1185 |
| a4 | SPU_026946 | Lv-Rp2 |
| a4 | SPU_026956 | Lv-Tmem127 |
| a4 | SPU_026972 | Lv-Slimb/Btrcp |
| a4 | SPU_027029 | Lv-Tor3aL |
| a4 | SPU_027117 | Lv-Nat13 |
| a4 | SPU_027132 | none |
| a4 | SPU_027332 | Lv-Clect/7Tm |
| a4 | SPU_027391 | Lv-Hypp_1201 |
| a4 | SPU_027413 | Lv-Asf1ha |
| a4 | SPU_027558 | Lv-Ccdc79L |
| a4 | SPU_027804 | none |
| a4 | SPU_027967 | Lv-Unk_33 |
| a4 | SPU_028114 | Lv-Hypp_2932 |

| | | |
|---|---|---|
| a4 | SPU_028214 | Lv-Mtusl |
| a4 | SPU_028353 | Lv-Rtn4ip1_2 |
| a4 | SPU_028409 | Lv-Smyd3_2 |
| a4 | SPU_028591 | Lv-PppL_289 |
| a4 | SPU_028743 | Lv-Hypp_2965 |
| a4 | SPU_028901 | Lv-Papd4 |
| a4 | SPU_030167 | Lv-Mafs |
| a5 | SPU_000009 | Lv-Mfsd1 |
| a5 | SPU_000183 | none |
| a5 | SPU_000722 | Lv-Trp53bp1 |
| a5 | SPU_000733 | Lv-Hypp_1287 |
| a5 | SPU_001030 | Lv-Ttl5 |
| a5 | SPU_001037 | Lv-Setd8L |
| a5 | SPU_001135 | none |
| a5 | SPU_001174 | Lv-Maoa |
| a5 | SPU_001222 | Lv-Fn3_39 |
| a5 | SPU_001320 | Lv-Pacs2 |
| a5 | SPU_001756 | none |
| a5 | SPU_002021 | Lv-Svep1 |
| a5 | SPU_002054 | Lv-Atad5 |
| a5 | SPU_002079 | Lv-Slinky |
| a5 | SPU_002173 | Lv-Znfx1_3 |
| a5 | SPU_002340 | Lv-Herc1L |
| a5 | SPU_002640 | Lv-Evi5 |
| a5 | SPU_002976 | Lv-Hypp_1423 |
| a5 | SPU_003043 | Lv-Cda |
| a5 | SPU_003115 | Lv-Vps54 |
| a5 | SPU_003187 | Lv-TipinL |
| a5 | SPU_003487 | Lv-Anln |
| a5 | SPU_003516 | Lv-Usp31 |
| a5 | SPU_003692 | Lv-Usp34-2 |
| a5 | SPU_003869 | Lv-Pign |
| a5 | SPU_004063 | Lv-Herc2L |
| a5 | SPU_004351 | Lv-Aff3 |
| a5 | SPU_004395 | Lv-Cdc20 |
| a5 | SPU_004516 | Lv-Ppil5 |
| a5 | SPU_004765 | Lv-Ece1L_1 |
| a5 | SPU_004781 | Lv-Polr3c |

| | | |
|---|---|---|
| a5 | SPU_004914 | NA |
| a5 | SPU_004915 | Lv-Hypp_3085 |
| a5 | SPU_004933 | Lv-Hypp_3086 |
| a5 | SPU_005062 | Lv-Trim37 |
| a5 | SPU_005070 | Lv-Asps |
| a5 | SPU_005151 | Lv-Sptan1 |
| a5 | SPU_005211 | Lv-Hypp_3080 |
| a5 | SPU_005225 | Lv-Hypp_4 |
| a5 | SPU_005343 | Lv-Orc6 |
| a5 | SPU_005483 | Lv-Myo-ino2dhl |
| a5 | SPU_005526 | Lv-C14orf135L |
| a5 | SPU_005879 | Lv-Hypp_1612 |
| a5 | SPU_005886 | Lv-Lztr1l |
| a5 | SPU_006144 | Lv-C12orf70 |
| a5 | SPU_006258 | Lv-Ube3a-2 |
| a5 | SPU_006303 | none |
| a5 | SPU_006392 | Lv-Rab33 |
| a5 | SPU_006483 | Lv-Sema/Psi_4 |
| a5 | SPU_006713 | Lv-Pgap1L |
| a5 | SPU_006789 | Lv-Smooth_1 |
| a5 | SPU_007193 | none |
| a5 | SPU_007244 | NA |
| a5 | SPU_007346 | Lv-Hypp_23 |
| a5 | SPU_007414 | Lv-Hypp_24 |
| a5 | SPU_007480 | Lv-CdyL |
| a5 | SPU_008023 | Lv-Hypp_1760 |
| a5 | SPU_008070 | Lv-Inturned_1 |
| a5 | SPU_008106 | Lv-Chrm5L_1 |
| a5 | SPU_008118 | Lv-OptnL |
| a5 | SPU_008122 | Lv-C3orf63L |
| a5 | SPU_008144 | none |
| a5 | SPU_008161 | Lv-Hypp_537 |
| a5 | SPU_008461 | Lv-Tfb2m |
| a5 | SPU_008577 | Lv-PolppL_77 |
| a5 | SPU_008581 | Lv-PolppL_78 |
| a5 | SPU_008626 | none |
| a5 | SPU_008767 | Lv-Jmjd5 |
| a5 | SPU_008904 | Lv-Hypp_1812 |

| | | |
|---|---|---|
| a5 | SPU_009133 | Lv-Galnt1-2 |
| a5 | SPU_009211 | Lv-Hypp_1834 |
| a5 | SPU_009288 | Lv-Pde7aL |
| a5 | SPU_009299 | Lv-Hypp_1841 |
| a5 | SPU_009311 | Lv-Ttrspn_6 |
| a5 | SPU_009520 | Lv-Tcf |
| a5 | SPU_009836 | none |
| a5 | SPU_009918 | Lv-Hypp_1879 |
| a5 | SPU_010010 | Lv-Tpcn2 |
| a5 | SPU_010058 | Lv-Fam62b |
| a5 | SPU_010075 | Lv-Kiaa0195L |
| a5 | SPU_010110 | Lv-Hypp_3064 |
| a5 | SPU_010233 | Lv-Zufsp |
| a5 | SPU_010438 | Lv-Pou6 |
| a5 | SPU_010567 | Lv-Kcnk18 |
| a5 | SPU_010737 | Lv-Usp7 |
| a5 | SPU_010852 | none |
| a5 | SPU_010963 | none |
| a5 | SPU_011082 | NA |
| a5 | SPU_011423 | Lv-Hypp_646 |
| a5 | SPU_011563 | Lv-Ncapd3 |
| a5 | SPU_011565 | Lv-Hypp_651 |
| a5 | SPU_011655 | Lv-Lnb/7Tm/GpcrA |
| a5 | SPU_011735 | Lv-C14orf135L |
| a5 | SPU_011738 | Lv-Hypp_657 |
| a5 | SPU_011782 | Lv-Hypp_74 |
| a5 | SPU_011790 | Lv-Fryh |
| a5 | SPU_011935 | NA |
| a5 | SPU_012027 | Lv-Smarcad1 |
| a5 | SPU_012200 | Lv-Ube20 |
| a5 | SPU_012529 | Lv-Mosa |
| a5 | SPU_012847 | Lv-Hypp_81 |
| a5 | SPU_012962 | Lv-Hypp_2061 |
| a5 | SPU_013245 | Lv-Ppgb_1 |
| a5 | SPU_013360 | none |
| a5 | SPU_013411 | Lv-Rassf9L |
| a5 | SPU_013612 | Lv-Fam46c |
| a5 | SPU_013785 | Lv-Tep1L |

| | | |
|---|---|---|
| a5 | SPU_013842 | Lv-Zir1 |
| a5 | SPU_014182 | Lv-Wdr40a |
| a5 | SPU_014272 | Lv-Sil1 |
| a5 | SPU_014311 | Lv-RigIL7 |
| a5 | SPU_014440 | Lv-Mett10d |
| a5 | SPU_014517 | Lv-Inpp5f |
| a5 | SPU_014629 | NA |
| a5 | SPU_014634 | Lv-Slc16a12_6 |
| a5 | SPU_014663 | Lv-C6orf84L |
| a5 | SPU_014691 | Lv-Pole2 |
| a5 | SPU_014954 | none |
| a5 | SPU_015042 | Lv-Nrbf2L |
| a5 | SPU_015092 | Lv-Baz2b |
| a5 | SPU_015348 | Lv-Slc44a2_2 |
| a5 | SPU_015355 | Lv-Hypp_2206 |
| a5 | SPU_015420 | NA |
| a5 | SPU_015500 | Lv-Harbi1_1 |
| a5 | SPU_015642 | Lv-Fgd4 |
| a5 | SPU_016042 | Lv-Mad2a |
| a5 | SPU_016172 | Lv-Pi4kb |
| a5 | SPU_016266 | Lv-Tmem201 |
| a5 | SPU_016321 | none |
| a5 | SPU_016401 | none |
| a5 | SPU_016426 | none |
| a5 | SPU_016695 | NA |
| a5 | SPU_016830 | Lv-Mcfp |
| a5 | SPU_017154 | Lv-Diva |
| a5 | SPU_017155 | NA |
| a5 | SPU_017219 | Lv-Dhrs7b |
| a5 | SPU_017325 | Lv-Brs3L |
| a5 | SPU_017382 | Lv-Ubr4-3 |
| a5 | SPU_017559 | Lv-Arrdc3_2 |
| a5 | SPU_017829 | none |
| a5 | SPU_017959 | Lv-BcDph |
| a5 | SPU_018016 | Lv-Gpr143 |
| a5 | SPU_018044 | Lv-Ubr4_1 |
| a5 | SPU_018062 | none |
| a5 | SPU_018551 | NA |

| | | |
|---|---|---|
| a5 | SPU_018570 | Lv-Clca_10 |
| a5 | SPU_018736 | Lv-Hypp_3040 |
| a5 | SPU_018821 | NA |
| a5 | SPU_018888 | Lv-Z40 |
| a5 | SPU_018968 | Lv-Fez2 |
| a5 | SPU_019040 | Lv-Naga |
| a5 | SPU_019384 | Lv-Z367 |
| a5 | SPU_019427 | Lv-Trip13 |
| a5 | SPU_019452 | Lv-Hypp_2441 |
| a5 | SPU_019539 | Lv-Tapt1 |
| a5 | SPU_019666 | NA |
| a5 | SPU_019706 | Lv-Pir |
| a5 | SPU_019745 | Lv-Hypp_159 |
| a5 | SPU_019955 | Lv-Reps1L_1 |
| a5 | SPU_020070 | Lv-Sos |
| a5 | SPU_020390 | none |
| a5 | SPU_020478 | NA |
| a5 | SPU_020587 | Lv-Ptger3 |
| a5 | SPU_020664 | Lv-Chd1l |
| a5 | SPU_020864 | Lv-Unk_77 |
| a5 | SPU_020871 | Lv-Lmbr1L |
| a5 | SPU_020939 | none |
| a5 | SPU_020986 | Lv-Ccnr2 |
| a5 | SPU_021048 | Lv-Hypp_2533 |
| a5 | SPU_021704 | Lv-CycJ |
| a5 | SPU_021829 | Lv-3HEXO |
| a5 | SPU_021852 | Lv-Hypp_1031 |
| a5 | SPU_022359 | Lv-C10orf28L |
| a5 | SPU_022422 | none |
| a5 | SPU_022447 | Lv-Slc4a1ap |
| a5 | SPU_022485 | Lv-Mll1L |
| a5 | SPU_022841 | Lv-Z142 |
| a5 | SPU_023034 | Lv-Ufd1l |
| a5 | SPU_023090 | Lv-Pric |
| a5 | SPU_023093 | Lv-Hypp_193 |
| a5 | SPU_023246 | Lv-Trpm_1 |
| a5 | SPU_023253 | Lv-Htr6L_1 |
| a5 | SPU_023408 | Lv-Nek10_1 |

| | | |
|---|---|---|
| a5 | SPU_023539 | Lv-TrimL_2 |
| a5 | SPU_023896 | Lv-C1orf124L |
| a5 | SPU_023982 | Lv-Dbf4 |
| a5 | SPU_024061 | NA |
| a5 | SPU_024177 | Lv-Amn |
| a5 | SPU_024199 | Lv-Sav1l |
| a5 | SPU_024297 | NA |
| a5 | SPU_024528 | Lv-Sr/Fu/Igr_1 |
| a5 | SPU_024660 | Lv-C10orf112 |
| a5 | SPU_024755 | Lv-Maoa_2 |
| a5 | SPU_024783 | Lv-C9orf41 |
| a5 | SPU_024815 | Lv-Tlr133 |
| a5 | SPU_024928 | Lv-Armc9-2 |
| a5 | SPU_024990 | Lv-Dnajb9 |
| a5 | SPU_025047 | Lv-Rif1L |
| a5 | SPU_025131 | Lv-Pole-3 |
| a5 | SPU_025404 | Lv-C3orf38L |
| a5 | SPU_025727 | Lv-Fbxo42 |
| a5 | SPU_025749 | Lv-Pole |
| a5 | SPU_025951 | Lv-Pdp |
| a5 | SPU_025960 | Lv-FryL_2 |
| a5 | SPU_025978 | Lv-Hypp_1163 |
| a5 | SPU_026040 | none |
| a5 | SPU_026216 | Lv-Tnfrsf_cl2 |
| a5 | SPU_026447 | Lv-FragileX |
| a5 | SPU_026627 | Lv-Mglur7 |
| a5 | SPU_026709 | Lv-Srcr214 |
| a5 | SPU_026878 | Lv-Klhl20_2 |
| a5 | SPU_027001 | Lv-Usp4 |
| a5 | SPU_027481 | Lv-Bcar3 |
| a5 | SPU_027525 | Lv-PlexA2_3 |
| a5 | SPU_027685 | Lv-Sh2Brs1 |
| a5 | SPU_028054 | Lv-Ccdc101 |
| a5 | SPU_028109 | Lv-Mre11 |
| a5 | SPU_028266 | Lv-Egf/HyalinL11 |
| a5 | SPU_028378 | Lv-Hypp_2953 |
| a5 | SPU_028451 | Lv-Abcd4 |
| a5 | SPU_028592 | Lv-Ptpn14/21 |

| | | |
|---|---|---|
| a5 | SPU_028724 | Lv-Pik2 |
| a5 | SPU_028926 | Lv-Nf2 |
| b1 | SPU_000043 | Lv-Lrp6l |
| b1 | SPU_000448 | Lv-Hypp_303 |
| b1 | SPU_000751 | Lv-MacpfD1 |
| b1 | SPU_000763 | Lv-Ccdc135-2 |
| b1 | SPU_000841 | Lv-Cryptic |
| b1 | SPU_000968 | Lv-Chrm1 |
| b1 | SPU_001025 | Lv-Hypp_321 |
| b1 | SPU_001376 | Lv-Hypp_3112 |
| b1 | SPU_001778 | Lv-Ttrspn_2 |
| b1 | SPU_001945 | Lv-KazL |
| b1 | SPU_002151 | NA |
| b1 | SPU_002252 | Lv-Rab39 |
| b1 | SPU_002300 | none |
| b1 | SPU_002424 | Lv-Ccdc105 |
| b1 | SPU_002662 | Lv-Bmp5/6/7/8_1 |
| b1 | SPU_002783 | Lv-NotchL_1 |
| b1 | SPU_003042 | NA |
| b1 | SPU_003295 | Lv-Cecr2L |
| b1 | SPU_003637 | Lv-Hypp_391 |
| b1 | SPU_003767 | Lv-Hypp_397 |
| b1 | SPU_003839 | NA |
| b1 | SPU_004288 | Lv-Hypp_1510 |
| b1 | SPU_004438 | none |
| b1 | SPU_004458 | NA |
| b1 | SPU_004486 | none |
| b1 | SPU_004490 | none |
| b1 | SPU_004498 | Lv-DlcL2 |
| b1 | SPU_004558 | none |
| b1 | SPU_004619 | Lv-Pacrg |
| b1 | SPU_004880 | none |
| b1 | SPU_005166 | Lv-Hypp_3079 |
| b1 | SPU_005440 | Lv-Chrna7-3 |
| b1 | SPU_005576 | Lv-Mt6_mmpL |
| b1 | SPU_005924 | Lv-Hypp_12 |
| b1 | SPU_005929 | Lv-Hypp_1618 |
| b1 | SPU_006009 | Lv-Hypp_1624 |

| | | |
|---|---|---|
| b1 | SPU_006284 | Lv-Hypp_15 |
| b1 | SPU_006349 | Lv-Ajpx3 |
| b1 | SPU_006391 | Lv-C20orf85L |
| b1 | SPU_006453 | Lv-Tektin1-4 |
| b1 | SPU_006491 | Lv-Hypp_483 |
| b1 | SPU_006724 | Lv-Lphn3L |
| b1 | SPU_006792 | none |
| b1 | SPU_006796 | Lv-Hypp_1682 |
| b1 | SPU_006844 | NA |
| b1 | SPU_006913 | Lv-Wdr64 |
| b1 | SPU_007051 | none |
| b1 | SPU_007086 | Lv-Bcr_1 |
| b1 | SPU_007215 | NA |
| b1 | SPU_007387 | Lv-Phtf2L |
| b1 | SPU_007419 | none |
| b1 | SPU_007437 | Lv-Cobll |
| b1 | SPU_007643 | Lv-Hypp_1739 |
| b1 | SPU_007758 | none |
| b1 | SPU_007779 | none |
| b1 | SPU_007813 | Lv-Hypp_527 |
| b1 | SPU_008114 | none |
| b1 | SPU_008176 | NA |
| b1 | SPU_008183 | Lv-Cks1B_1 |
| b1 | SPU_008314 | Lv-Rwdd2b |
| b1 | SPU_008348 | Lv-Hypp_1780 |
| b1 | SPU_008500 | none |
| b1 | SPU_008614 | NA |
| b1 | SPU_008777 | Lv-Tektin1-1 |
| b1 | SPU_008842 | Lv-Ust |
| b1 | SPU_009002 | Lv-Rej3 |
| b1 | SPU_009072 | Lv-Efcab8L |
| b1 | SPU_009229 | Lv-Hypp_1835 |
| b1 | SPU_009255 | Lv-Ccdc135 |
| b1 | SPU_009301 | Lv-Ksr2 |
| b1 | SPU_009308 | Lv-Aasdh |
| b1 | SPU_009414 | none |
| b1 | SPU_009619 | Lv-Cry/dash |
| b1 | SPU_009641 | Lv-Hypp_1863 |

| | | |
|---|---|---|
| b1 | SPU_009896 | Lv-Efhc2_1 |
| b1 | SPU_009992 | NA |
| b1 | SPU_010087 | NA |
| b1 | SPU_010098 | Lv-Igfbp7 |
| b1 | SPU_010131 | Lv-Wdr65 |
| b1 | SPU_010239 | none |
| b1 | SPU_010387 | Lv-Chrm3 |
| b1 | SPU_010470 | Lv-Clect/Pan_8 |
| b1 | SPU_010492 | Lv-C14orf45 |
| b1 | SPU_010566 | Lv-Kcnk18L |
| b1 | SPU_010712 | NA |
| b1 | SPU_010725 | none |
| b1 | SPU_010882 | Lv-Htr4L_5 |
| b1 | SPU_010970 | none |
| b1 | SPU_011044 | none |
| b1 | SPU_011116 | NA |
| b1 | SPU_011247 | Lv-Hypp_1969 |
| b1 | SPU_011316 | Lv-Hypp_1972 |
| b1 | SPU_011598 | Lv-Unc44L_74 |
| b1 | SPU_011786 | Lv-ThegL |
| b1 | SPU_011852 | Lv-Hypp_2004 |
| b1 | SPU_011855 | Lv-Nlr130 |
| b1 | SPU_012019 | Lv-Hypp_2012 |
| b1 | SPU_012236 | none |
| b1 | SPU_012442 | NA |
| b1 | SPU_012514 | Lv-Col |
| b1 | SPU_012650 | NA |
| b1 | SPU_012786 | Lv-Bmp5/6/7/8 |
| b1 | SPU_013058 | Lv-Hypp_2066 |
| b1 | SPU_013076 | Lv-Ajpx1 |
| b1 | SPU_013154 | Lv-Hrh3_6 |
| b1 | SPU_013329 | none |
| b1 | SPU_013440 | Lv-Pla2g7 |
| b1 | SPU_013529 | Lv-Slc4a2_1 |
| b1 | SPU_014102 | none |
| b1 | SPU_014155 | Lv-Hypp_2134 |
| b1 | SPU_014186 | none |
| b1 | SPU_014318 | Lv-Tada3L |

| | | |
|---|---|---|
| b1 | SPU_014534 | Lv-Mae |
| b1 | SPU_014550 | Lv-Adra1dL |
| b1 | SPU_014744 | NA |
| b1 | SPU_014776 | Lv-Wdr78 |
| b1 | SPU_015022 | Lv-Pogk_1 |
| b1 | SPU_015109 | Lv-Thap9_1 |
| b1 | SPU_015168 | Lv-PppL_199 |
| b1 | SPU_015272 | Lv-Efcab6 |
| b1 | SPU_015788 | Lv-Hypp_800 |
| b1 | SPU_015854 | none |
| b1 | SPU_016128 | Lv-Delta |
| b1 | SPU_016396 | Lv-Fam60a |
| b1 | SPU_016562 | Lv-Hsp40A |
| b1 | SPU_017278 | Lv-Hypp_2317 |
| b1 | SPU_017444 | Lv-Adprh_2 |
| b1 | SPU_017446 | Lv-Hypp_2324 |
| b1 | SPU_017616 | NA |
| b1 | SPU_017778 | none |
| b1 | SPU_017815 | none |
| b1 | SPU_018152 | none |
| b1 | SPU_018238 | Lv-TlrL_11 |
| b1 | SPU_018250 | Lv-St8sia4_4 |
| b1 | SPU_018290 | Lv-Slc35f1 |
| b1 | SPU_018349 | NA |
| b1 | SPU_018362 | Lv-PppL_120 |
| b1 | SPU_018701 | Lv-Slc4a3 |
| b1 | SPU_019059 | Lv-Mprot1L_1 |
| b1 | SPU_019140 | none |
| b1 | SPU_019148 | Lv-Efcab6L |
| b1 | SPU_019169 | Lv-CttnL |
| b1 | SPU_019217 | Lv-Hypp_2425 |
| b1 | SPU_019491 | none |
| b1 | SPU_019521 | none |
| b1 | SPU_019591 | Lv-Tektin1-2 |
| b1 | SPU_019734 | none |
| b1 | SPU_019897 | NA |
| b1 | SPU_020005 | none |
| b1 | SPU_020193 | Lv-Hypp_2494 |

| | | |
|---|---|---|
| b1 | SPU_020442 | Lv-Cecr2L_2 |
| b1 | SPU_020728 | Lv-Tektin2 |
| b1 | SPU_020729 | NA |
| b1 | SPU_021021 | Lv-Pdgfr/vegfrL_1 |
| b1 | SPU_021034 | Lv-Ccdc37 |
| b1 | SPU_021255 | Lv-Cyp3L5 |
| b1 | SPU_021298 | Lv-Trex2L |
| b1 | SPU_021306 | none |
| b1 | SPU_021388 | Lv-Aqp_2 |
| b1 | SPU_021429 | Lv-Efhc1 |
| b1 | SPU_021664 | Lv-Ccdc81 |
| b1 | SPU_021722 | Lv-Hypp_2573 |
| b1 | SPU_021779 | NA |
| b1 | SPU_021884 | Lv-Hypp_2583 |
| b1 | SPU_021912 | Lv-Slc47a1L |
| b1 | SPU_021929 | Lv-Hypp_2588 |
| b1 | SPU_021964 | NA |
| b1 | SPU_022323 | Lv-Tpase3 |
| b1 | SPU_022360 | Lv-C11orf66L |
| b1 | SPU_022385 | none |
| b1 | SPU_022412 | Lv-Nlr99 |
| b1 | SPU_022444 | Lv-TpaseL_13 |
| b1 | SPU_022514 | Lv-Hypp_186 |
| b1 | SPU_022527 | none |
| b1 | SPU_022852 | Lv-Efcab6L_2 |
| b1 | SPU_023137 | Lv-Slc17a5_10 |
| b1 | SPU_023186 | Lv-Hypp_2654 |
| b1 | SPU_023231 | Lv-Cut8L |
| b1 | SPU_023346 | Lv-E2f7 |
| b1 | SPU_023618 | Lv-Tektin3 |
| b1 | SPU_023743 | Lv-Spata17 |
| b1 | SPU_023909 | Lv-B3galt1_12 |
| b1 | SPU_024029 | NA |
| b1 | SPU_024052 | Lv-Lrr/Gpcr_9 |
| b1 | SPU_024086 | Lv-C6orf97L |
| b1 | SPU_024183 | none |
| b1 | SPU_024352 | NA |
| b1 | SPU_024398 | NA |

| | | |
|---|---|---|
| b1 | SPU_024661 | Lv-Hypp_2745 |
| b1 | SPU_024886 | Lv-ZanL_2 |
| b1 | SPU_024943 | Lv-C10orf112L_2 |
| b1 | SPU_025107 | Lv-Ccdc123 |
| b1 | SPU_025153 | Lv-Fbxl10_1 |
| b1 | SPU_025224 | Lv-Xdh_4 |
| b1 | SPU_025490 | none |
| b1 | SPU_025700 | none |
| b1 | SPU_025936 | Lv-Ccdc151 |
| b1 | SPU_025980 | Lv-Spag8 |
| b1 | SPU_026062 | none |
| b1 | SPU_026143 | none |
| b1 | SPU_026477 | Lv-Cyp2L63 |
| b1 | SPU_026664 | Lv-Pcsk7 |
| b1 | SPU_026963 | Lv-Hypp_2878 |
| b1 | SPU_026986 | none |
| b1 | SPU_027012 | Lv-Wee1 |
| b1 | SPU_027354 | Lv-Hypp_1200 |
| b1 | SPU_027358 | Lv-Hypp_3010 |
| b1 | SPU_027414 | NA |
| b1 | SPU_027681 | Lv-C3orf44L |
| b1 | SPU_027862 | Lv-Mamdc4-15 |
| b1 | SPU_027869 | Lv-Hypp_1217 |
| b1 | SPU_027934 | Lv-Mns1 |
| b1 | SPU_027987 | Lv-Acel_1 |
| b1 | SPU_028063 | NA |
| b1 | SPU_028181 | Lv-CepL |
| b1 | SPU_028183 | Lv-Dnajhc4L |
| b1 | SPU_028240 | none |
| b1 | SPU_028341 | Lv-Hypp_1227 |
| b1 | SPU_028386 | Lv-Ifltd1L |
| b1 | SPU_028648 | Lv-EhH1_41 |
| b2 | SPU_000406 | Lv-Fn3/Igf |
| b2 | SPU_000538 | Lv-Egf/Ccp |
| b2 | SPU_000726 | none |
| b2 | SPU_000752 | Lv-Hypp_313 |
| b2 | SPU_000871 | Lv-Tlr007 |
| b2 | SPU_000989 | Lv-Hypp_1298 |

| | | |
|---|---|---|
| b2 | SPU_001016 | Lv-Nlr113 |
| b2 | SPU_001095 | NA |
| b2 | SPU_001189 | Lv-Hypp_327 |
| b2 | SPU_001588 | Lv-GramarL2 |
| b2 | SPU_001812 | Lv-PolppL-7 |
| b2 | SPU_001987 | none |
| b2 | SPU_002029 | Lv-Unk_14 |
| b2 | SPU_002119 | none |
| b2 | SPU_002689 | Lv-Myt1-2 |
| b2 | SPU_002757 | Lv-Lrr/Igr_6 |
| b2 | SPU_003058 | Lv-PolypL_959 |
| b2 | SPU_003208 | Lv-Wscd2L |
| b2 | SPU_003570 | NA |
| b2 | SPU_005473 | Lv-Sstr5L |
| b2 | SPU_005899 | NA |
| b2 | SPU_005940 | Lv-Hypp_1621 |
| b2 | SPU_006030 | Lv-Megf6-2 |
| b2 | SPU_006423 | none |
| b2 | SPU_007184 | none |
| b2 | SPU_008026 | Lv-Ablim1 |
| b2 | SPU_008137 | Lv-PolppL_65 |
| b2 | SPU_008395 | NA |
| b2 | SPU_008910 | Lv-Glur6_2 |
| b2 | SPU_009079 | Lv-Tk11 |
| b2 | SPU_009465 | Lv-Hey |
| b2 | SPU_009750 | none |
| b2 | SPU_010504 | none |
| b2 | SPU_011472 | NA |
| b2 | SPU_011549 | Lv-Hypp_649 |
| b2 | SPU_012354 | Lv-Igf_3 |
| b2 | SPU_012878 | Lv-Ankib1_1 |
| b2 | SPU_013342 | Lv-Pema/Srcr |
| b2 | SPU_013669 | Lv-Mt1-4/MmpL5 |
| b2 | SPU_013676 | Lv-Tlr072 |
| b2 | SPU_013779 | none |
| b2 | SPU_013803 | none |
| b2 | SPU_013829 | Lv-Sea/Egf/Ccp |
| b2 | SPU_014393 | Lv-PppL_264 |

| | | |
|---|---|---|
| b2 | SPU_014901 | none |
| b2 | SPU_015204 | Lv-Hypp_2197 |
| b2 | SPU_015583 | Lv-Hypp_790 |
| b2 | SPU_015904 | Lv-Hypp_802 |
| b2 | SPU_015984 | Lv-Prox1 |
| b2 | SPU_016089 | Lv-Vwd/Egff_1 |
| b2 | SPU_016232 | none |
| b2 | SPU_016290 | Lv-Hypp_2256 |
| b2 | SPU_016318 | none |
| b2 | SPU_016868 | Lv-Tmprss6_1 |
| b2 | SPU_016997 | Lv-Hypp_2996 |
| b2 | SPU_017075 | NA |
| b2 | SPU_017229 | Lv-Hypp_2312 |
| b2 | SPU_017585 | Lv-Casr |
| b2 | SPU_018703 | none |
| b2 | SPU_020122 | Lv-Hypp_963 |
| b2 | SPU_020176 | Lv-Itpkb |
| b2 | SPU_020430 | Lv-Clca_13 |
| b2 | SPU_020756 | Lv-Cyp2L46 |
| b2 | SPU_021541 | Lv-Hypp_1026 |
| b2 | SPU_021746 | Lv-Hypp_2574 |
| b2 | SPU_021904 | Lv-Igf_22 |
| b2 | SPU_022964 | Lv-Cfdp2L |
| b2 | SPU_023210 | Lv-Hypp_1076 |
| b2 | SPU_023440 | Lv-Abcc |
| b2 | SPU_023633 | none |
| b2 | SPU_023918 | NA |
| b2 | SPU_025074 | Lv-Clect_95 |
| b2 | SPU_025401 | Lv-Sterol_oxd |
| b2 | SPU_025588 | Lv-Endrvt47 |
| b2 | SPU_025738 | Lv-FoxM_1 |
| b2 | SPU_025801 | Lv-Kctd1L |
| b2 | SPU_026006 | Lv-Hypp_237 |
| b2 | SPU_026151 | Lv-Znfx1-15 |
| b2 | SPU_026602 | Lv-Myst3L |
| b2 | SPU_027949 | none |
| b2 | SPU_028397 | Lv-Ddx42 |
| b2 | SPU_028415 | Lv-Unc44_189 |

| | | |
|---|---|---|
| b2 | SPU_028570 | none |
| b2 | SPU_028646 | none |
| b2 | SPU_028700 | none |
| b2 | SPU_028702 | Lv-Hypp_2964 |
| c1 | SPU_000036 | Lv-Acot8 |
| c1 | SPU_000079 | Lv-Hypp_1241 |
| c1 | SPU_000201 | Lv-Hnmtl |
| c1 | SPU_000346 | Lv-Clect/Fn3_17 |
| c1 | SPU_000485 | Lv-Chx10 |
| c1 | SPU_000544 | Lv-Orc3 |
| c1 | SPU_000587 | none |
| c1 | SPU_000626 | Lv-Hypp_1267 |
| c1 | SPU_000661 | Lv-Elovl6 |
| c1 | SPU_001129 | Lv-Lrr/Igr_10 |
| c1 | SPU_001372 | Lv-Hypp_1323 |
| c1 | SPU_001381 | Lv-Slc16a6L_2 |
| c1 | SPU_001482 | none |
| c1 | SPU_001490 | Lv-Hypp_1334 |
| c1 | SPU_001584 | Lv-Airc |
| c1 | SPU_001749 | Lv-TlrL_16 |
| c1 | SPU_001949 | Lv-Galnt1 |
| c1 | SPU_001958 | none |
| c1 | SPU_002016 | Lv-Cubn |
| c1 | SPU_002071 | Lv-C8orf45L |
| c1 | SPU_002263 | Lv-PppL_239 |
| c1 | SPU_002266 | Lv-PolppL_15 |
| c1 | SPU_002449 | Lv-Sec23ip |
| c1 | SPU_002634 | NA |
| c1 | SPU_002876 | Lv-Lrr/Gpcr_14 |
| c1 | SPU_003144 | Lv-Hypp_374 |
| c1 | SPU_003213 | Lv-Erf3 |
| c1 | SPU_003376 | NA |
| c1 | SPU_003422 | none |
| c1 | SPU_003491 | none |
| c1 | SPU_003635 | Lv-Eggbr1L_1 |
| c1 | SPU_003653 | Lv-Gmppa |
| c1 | SPU_004199 | NA |
| c1 | SPU_004240 | Lv-Hypp_407 |

| | | |
|---|---|---|
| c1 | SPU_004343 | Lv-Nlr145 |
| c1 | SPU_004758 | Lv-Fn3f_9 |
| c1 | SPU_004830 | Lv-Tie2L_4 |
| c1 | SPU_004910 | none |
| c1 | SPU_005129 | none |
| c1 | SPU_005237 | Lv-Elovl6_2 |
| c1 | SPU_005390 | Lv-Dcst2-3 |
| c1 | SPU_005538 | Lv-Lrr/Igr_2 |
| c1 | SPU_005556 | Lv-Srcr44 |
| c1 | SPU_005585 | Lv-Aqp |
| c1 | SPU_005594 | Lv-Clect_12 |
| c1 | SPU_005622 | none |
| c1 | SPU_005815 | Lv-Plxna4L_3 |
| c1 | SPU_005858 | Lv-Unc44L_45 |
| c1 | SPU_005863 | Lv-Gpd2 |
| c1 | SPU_005960 | Lv-Fn3_22 |
| c1 | SPU_005996 | none |
| c1 | SPU_006072 | Lv-Dnapp58 |
| c1 | SPU_006151 | none |
| c1 | SPU_006845 | Lv-Slc9a2 |
| c1 | SPU_007070 | Lv-FibpL |
| c1 | SPU_007204 | Lv-Cry1 |
| c1 | SPU_007331 | none |
| c1 | SPU_007428 | Lv-Mfsd6L |
| c1 | SPU_007661 | NA |
| c1 | SPU_007718 | Lv-Srcr57 |
| c1 | SPU_007859 | Lv-Tlr041 |
| c1 | SPU_007865 | Lv-El1L1 |
| c1 | SPU_008189 | Lv-Npl |
| c1 | SPU_008354 | Lv-Kirrel2L_2 |
| c1 | SPU_008379 | none |
| c1 | SPU_008472 | Lv-PlexA4_2 |
| c1 | SPU_008541 | none |
| c1 | SPU_008603 | Lv-Hypp_1799 |
| c1 | SPU_008613 | Lv-Fbn2L_1 |
| c1 | SPU_008815 | Lv-Gcnt2_2 |
| c1 | SPU_008857 | Lv-Timp3b |
| c1 | SPU_009069 | Lv-Mid2L |

| | | |
|---|---|---|
| c1 | SPU_009135 | Lv-Dhyg_2 |
| c1 | SPU_009271 | Lv-Nat10 |
| c1 | SPU_009338 | Lv-Uchl5 |
| c1 | SPU_009527 | Lv-TnfsfL2 |
| c1 | SPU_009702 | Lv-Mdn1_2 |
| c1 | SPU_009897 | Lv-Hypp_592 |
| c1 | SPU_009981 | Lv-Urb2h |
| c1 | SPU_009999 | Lv-Hypp_601 |
| c1 | SPU_010184 | Lv-TekL_1 |
| c1 | SPU_010501 | Lv-Srcr90 |
| c1 | SPU_010638 | NA |
| c1 | SPU_010782 | Lv-Heatr1 |
| c1 | SPU_010826 | Lv-Unc5cL_2 |
| c1 | SPU_010889 | none |
| c1 | SPU_010908 | none |
| c1 | SPU_010991 | Lv-Srcr95 |
| c1 | SPU_011256 | none |
| c1 | SPU_011294 | Lv-Ilkap_1 |
| c1 | SPU_011302 | Lv-Gal3st3L_2 |
| c1 | SPU_011345 | Lv-Kctd12b |
| c1 | SPU_011421 | none |
| c1 | SPU_011837 | Lv-Apobec1 |
| c1 | SPU_011893 | Lv-B3gnt5_1 |
| c1 | SPU_012041 | Lv-Tdrd4 |
| c1 | SPU_012317 | none |
| c1 | SPU_012359 | none |
| c1 | SPU_012404 | Lv-Rbpms2L |
| c1 | SPU_012413 | Lv-A2ml1L |
| c1 | SPU_012485 | none |
| c1 | SPU_012487 | Lv-Ddx56_2 |
| c1 | SPU_013078 | Lv-Hypp_702 |
| c1 | SPU_013100 | Lv-Ars |
| c1 | SPU_013130 | Lv-St8sia2_1 |
| c1 | SPU_013160 | Lv-Mdn1 |
| c1 | SPU_013187 | none |
| c1 | SPU_013261 | Lv-Scye1 |
| c1 | SPU_013472 | Lv-Hypp_718 |
| c1 | SPU_013571 | none |

| | | |
|---|---|---|
| c1 | SPU_013652 | Lv-Hypp_2095 |
| c1 | SPU_014283 | none |
| c1 | SPU_014310 | Lv-RigIL1 |
| c1 | SPU_014392 | NA |
| c1 | SPU_014549 | Lv-Sumf2 |
| c1 | SPU_014665 | Lv-Cbr1_3 |
| c1 | SPU_014798 | Lv-PolypL_8 |
| c1 | SPU_014807 | Lv-Ylpm1 |
| c1 | SPU_014882 | Lv-Eprs |
| c1 | SPU_014943 | NA |
| c1 | SPU_015254 | none |
| c1 | SPU_015295 | Lv-Hypp_779 |
| c1 | SPU_015321 | Lv-Celsr1 |
| c1 | SPU_015630 | Lv-Apextrin_1 |
| c1 | SPU_015675 | Lv-Naca |
| c1 | SPU_015906 | Lv-Hypp_120 |
| c1 | SPU_016144 | Lv-Ptprfn1(novel) |
| c1 | SPU_016449 | Lv-Hnf6 |
| c1 | SPU_016646 | Lv-Prss23_1 |
| c1 | SPU_016702 | Lv-Rdh8_1 |
| c1 | SPU_016836 | Lv-EMI/Egf |
| c1 | SPU_016910 | Lv-Unc44_118 |
| c1 | SPU_016938 | none |
| c1 | SPU_017145 | Lv-Hypp_862 |
| c1 | SPU_017148 | Lv-Transop |
| c1 | SPU_017226 | NA |
| c1 | SPU_017326 | Lv-Trpc5_1 |
| c1 | SPU_017348 | Lv-Atbf1 |
| c1 | SPU_017469 | Lv-Abce1 |
| c1 | SPU_017721 | Lv-Slc39a1 |
| c1 | SPU_017925 | Lv-Nup43_1 |
| c1 | SPU_018029 | Lv-TspN/Tsp1 |
| c1 | SPU_018126 | Lv-Tgif |
| c1 | SPU_018242 | Lv-Cyp46L5 |
| c1 | SPU_018275 | Lv-Notch1-15 |
| c1 | SPU_018671 | Lv-Hypp_143 |
| c1 | SPU_018696 | Lv-Hypp_920 |
| c1 | SPU_018894 | Lv-Gss_2 |

| | | |
|---|---|---|
| c1 | SPU_018947 | Lv-Gpr151L_12 |
| c1 | SPU_018980 | Lv-Arl11 |
| c1 | SPU_019003 | NA |
| c1 | SPU_019739 | Lv-Thsd7b |
| c1 | SPU_019841 | NA |
| c1 | SPU_019911 | Lv-B3galt1_8 |
| c1 | SPU_020024 | Lv-Hypp_2478 |
| c1 | SPU_020066 | Lv-C6orf211 |
| c1 | SPU_020093 | Lv-Hypp_2485 |
| c1 | SPU_020128 | none |
| c1 | SPU_020156 | Lv-Aacs_1 |
| c1 | SPU_020174 | Lv-Osbp2_1 |
| c1 | SPU_020203 | Lv-Egfib_9 |
| c1 | SPU_020603 | Lv-Lrp6L_1 |
| c1 | SPU_020609 | Lv-Fah |
| c1 | SPU_020731 | Lv-Gpr133L_2 |
| c1 | SPU_020840 | Lv-Cyc1_1 |
| c1 | SPU_020868 | NA |
| c1 | SPU_020896 | Lv-Hypp_1003 |
| c1 | SPU_021069 | none |
| c1 | SPU_021360 | Lv-B4galt6_6 |
| c1 | SPU_021628 | Lv-Ucp3 |
| c1 | SPU_021716 | Lv-Gars |
| c1 | SPU_022174 | none |
| c1 | SPU_022480 | Lv-Fam55c_6 |
| c1 | SPU_022562 | Lv-Eif3s9 |
| c1 | SPU_022614 | Lv-MidnL |
| c1 | SPU_022932 | Lv-Hypp_2638 |
| c1 | SPU_023004 | Lv-Unk_39 |
| c1 | SPU_023039 | none |
| c1 | SPU_023216 | Lv-Nacha3_1 |
| c1 | SPU_023677 | Lv-Srcr199 |
| c1 | SPU_023690 | Lv-Hypp_1093 |
| c1 | SPU_024044 | Lv-Tie1/2 |
| c1 | SPU_024210 | Lv-Abcg2 |
| c1 | SPU_024773 | Lv-Dhyd |
| c1 | SPU_024949 | Lv-Siat7e |
| c1 | SPU_025132 | none |

| | | |
|---|---|---|
| c1 | SPU_025161 | Lv-PolypL_123 |
| c1 | SPU_025332 | none |
| c1 | SPU_025437 | none |
| c1 | SPU_025454 | Lv-Thsd7b_2 |
| c1 | SPU_025818 | Lv-Psmd2_1 |
| c1 | SPU_025838 | NA |
| c1 | SPU_025839 | Lv-Hypp_1158 |
| c1 | SPU_026055 | Lv-Fut3L_1 |
| c1 | SPU_026182 | Lv-St8sia9L |
| c1 | SPU_026292 | Lv-Glis1_1 |
| c1 | SPU_026375 | NA |
| c1 | SPU_026476 | none |
| c1 | SPU_026565 | Lv-Hypp_2851 |
| c1 | SPU_026795 | Lv-Dfna5L |
| c1 | SPU_026816 | Lv-Adrb2L_20 |
| c1 | SPU_027188 | Lv-Udp1 |
| c1 | SPU_027682 | Lv-Aacs |
| c1 | SPU_027823 | Lv-St6galnac4_3 |
| c1 | SPU_027972 | Lv-Grwd1 |
| c1 | SPU_028242 | Lv-Ankrd6L |
| c1 | SPU_028252 | Lv-Endrvt120 |
| c1 | SPU_028261 | Lv-Msh6 |
| c1 | SPU_028416 | Lv-Dpgm |
| c1 | SPU_028422 | Lv-Rxr |
| c1 | SPU_028492 | Lv-Gal3st1L_5 |
| c1 | SPU_028535 | Lv-Lrp6L_3 |
| c1 | SPU_028638 | Lv-Paqr6 |
| c1 | SPU_028721 | Lv-Mov34L |
| c1 | SPU_030139 | Lv-Ilgf2 |
| c2 | SPU_000234 | Lv-Ank1L |
| c2 | SPU_000434 | none |
| c2 | SPU_000589 | Lv-Hypp_1264 |
| c2 | SPU_001172 | Lv-Srcr11 |
| c2 | SPU_001667 | Lv-HpseL |
| c2 | SPU_001913 | Lv-Hypp_1353 |
| c2 | SPU_002818 | Lv-Lhh2aj |
| c2 | SPU_003196 | none |
| c2 | SPU_003253 | Lv-Cnot1 |

| | | |
|---|---|---|
| c2 | SPU_003300 | Lv-Edil3L_2 |
| c2 | SPU_003439 | none |
| c2 | SPU_003821 | none |
| c2 | SPU_003973 | Lv-Kcnk5_2 |
| c2 | SPU_004207 | Lv-St6sia_1 |
| c2 | SPU_005118 | none |
| c2 | SPU_005167 | Lv-Col805b_2 |
| c2 | SPU_005178 | Lv-Hypp_439 |
| c2 | SPU_005268 | Lv-Higd1A |
| c2 | SPU_005457 | Lv-Unk_80 |
| c2 | SPU_006858 | Lv-Hypp_19 |
| c2 | SPU_006899 | Lv-Enp |
| c2 | SPU_007083 | Lv-MyIV_1 |
| c2 | SPU_008153 | Lv-Hypp_535 |
| c2 | SPU_008225 | Lv-Hypp_539 |
| c2 | SPU_008726 | Lv-Hypp_1802 |
| c2 | SPU_008772 | Lv-Hypp_559 |
| c2 | SPU_008911 | Lv-Hdpx1 |
| c2 | SPU_009077 | Lv-Echn29 |
| c2 | SPU_009203 | Lv-AnL12 |
| c2 | SPU_010388 | Lv-Hypp_1913 |
| c2 | SPU_010396 | none |
| c2 | SPU_010433 | none |
| c2 | SPU_010549 | Lv-Cyp1r4 |
| c2 | SPU_010841 | NA |
| c2 | SPU_010898 | Lv-RtL_26 |
| c2 | SPU_011021 | Lv-Unc44L_70 |
| c2 | SPU_011500 | Lv-Unc44L_73 |
| c2 | SPU_011726 | Lv-Hypp_656 |
| c2 | SPU_012754 | none |
| c2 | SPU_012936 | Lv-ThrombB1 |
| c2 | SPU_013133 | none |
| c2 | SPU_013750 | Lv-Etfa_1 |
| c2 | SPU_014052 | Lv-Hypp_735 |
| c2 | SPU_014375 | Lv-Clect/Fn3_12 |
| c2 | SPU_014608 | Lv-RtL_37 |
| c2 | SPU_014766 | Lv-Cthpsn6 |
| c2 | SPU_014966 | Lv-Psmc6 |

| | | |
|---|---|---|
| c2 | SPU_015171 | none |
| c2 | SPU_015402 | Lv-RtL_38 |
| c2 | SPU_015584 | Lv-Hypp_791 |
| c2 | SPU_015586 | Lv-Cpox |
| c2 | SPU_015860 | none |
| c2 | SPU_016314 | none |
| c2 | SPU_016596 | Lv-Pdcd11 |
| c2 | SPU_016979 | Lv-Hypp_2296 |
| c2 | SPU_017434 | Lv-Ptpra |
| c2 | SPU_017609 | Lv-Unk_9 |
| c2 | SPU_017618 | none |
| c2 | SPU_018359 | Lv-C10orf4_1 |
| c2 | SPU_018534 | Lv-Tlr099 |
| c2 | SPU_018846 | none |
| c2 | SPU_018953 | Lv-Traf3_1 |
| c2 | SPU_019013 | Lv-Hypp_2416 |
| c2 | SPU_019057 | Lv-Nmral1_1 |
| c2 | SPU_019064 | Lv-Eif4Gg(C-ter) |
| c2 | SPU_019245 | Lv-Hypp_2427 |
| c2 | SPU_019588 | Lv-Hypp_2452 |
| c2 | SPU_019765 | Lv-Hypp_160 |
| c2 | SPU_019767 | Lv-Ank7 |
| c2 | SPU_019826 | Lv-Srcr155 |
| c2 | SPU_020949 | none |
| c2 | SPU_021187 | Lv-Hypp_2541 |
| c2 | SPU_021221 | Lv-Clect_88 |
| c2 | SPU_021738 | Lv-Ftsj3_1 |
| c2 | SPU_022470 | Lv-Clect/Egf_13 |
| c2 | SPU_022742 | none |
| c2 | SPU_022833 | Lv-Tm9sf4_1 |
| c2 | SPU_023392 | Lv-Hypp_1084 |
| c2 | SPU_023428 | none |
| c2 | SPU_023663 | Lv-Lhh2aa |
| c2 | SPU_024125 | Lv-PolypL_121 |
| c2 | SPU_024238 | none |
| c2 | SPU_024339 | Lv-Guanc |
| c2 | SPU_024378 | none |
| c2 | SPU_024740 | Lv-Etfb_1 |

| | | |
|---|---|---|
| c2 | SPU_025057 | Lv-Net |
| c2 | SPU_026065 | Lv-Ebna1bp2_1 |
| c2 | SPU_026501 | Lv-Unk_41 |
| c2 | SPU_026866 | Lv-Kiaa1529L_3 |
| c2 | SPU_027032 | Lv-Pkd2L1 |
| c2 | SPU_027096 | Lv-Sr/Gpcr |
| c2 | SPU_027103 | NA |
| c2 | SPU_027192 | none |
| c2 | SPU_027217 | Lv-Hypp_2895 |
| c2 | SPU_027632 | Lv-SommL_3 |
| c2 | SPU_027989 | Lv-Senp1_1 |
| c2 | SPU_028143 | Lv-Ebr1L_9 |
| c2 | SPU_028399 | Lv-EhH4_79 |
| c2 | SPU_028486 | Lv-Unk_8 |
| c2 | SPU_028611 | Lv-Eif3a_1 |
| c2 | SPU_028716 | Lv-Fn3_37 |
| d1 | SPU_000057 | Lv-Igcam/Igcam/Igcam |
| d1 | SPU_000071 | NA |
| d1 | SPU_000136 | Lv-Mec-2l |
| d1 | SPU_000242 | Lv-Dazapi2 |
| d1 | SPU_000750 | none |
| d1 | SPU_000799 | Lv-Dpyd |
| d1 | SPU_000909 | Lv-Hypp_3119 |
| d1 | SPU_000960 | Lv-Ef1bd |
| d1 | SPU_001088 | Lv-Adfp |
| d1 | SPU_001089 | none |
| d1 | SPU_001181 | Lv-Ggt1 |
| d1 | SPU_001235 | Lv-Vps13a |
| d1 | SPU_001279 | Lv-Slc12a8 |
| d1 | SPU_001296 | Lv-Ser/Thrpp2B/Dirsigc |
| d1 | SPU_001299 | none |
| d1 | SPU_001529 | Lv-Vparpf |
| d1 | SPU_001722 | Lv-C19orf2 |
| d1 | SPU_001752 | Lv-Abcb1A |
| d1 | SPU_001779 | Lv-Amy_9 |
| d1 | SPU_001799 | Lv-Otof |
| d1 | SPU_001864 | Lv-Ggt1_1 |
| d1 | SPU_001934 | Lv-Dhrs4 |

| | | |
|---|---|---|
| d1 | SPU_002106 | none |
| d1 | SPU_002311 | Lv-Opn4L |
| d1 | SPU_002328 | Lv-Nos_1 |
| d1 | SPU_002437 | NA |
| d1 | SPU_002572 | Lv-Pla2g3 |
| d1 | SPU_002900 | Lv-Unc44L_28 |
| d1 | SPU_003009 | Lv-Osgin1 |
| d1 | SPU_003071 | Lv-Ndufb7 |
| d1 | SPU_003079 | Lv-Hypp_1429 |
| d1 | SPU_003094 | none |
| d1 | SPU_003193 | Lv-Kirrel2L_6 |
| d1 | SPU_003394 | NA |
| d1 | SPU_003429 | NA |
| d1 | SPU_003674 | Lv-Hypp_392 |
| d1 | SPU_003713 | Lv-Slc6a13 |
| d1 | SPU_003818 | none |
| d1 | SPU_003840 | Lv-MtnK |
| d1 | SPU_003907 | Lv-Hypp_3098 |
| d1 | SPU_003924 | none |
| d1 | SPU_004052 | Lv-Hypp_1495 |
| d1 | SPU_004143 | NA |
| d1 | SPU_004225 | none |
| d1 | SPU_004390 | Lv-Ltb4dh |
| d1 | SPU_004447 | Lv-Lrp4L_3 |
| d1 | SPU_004594 | none |
| d1 | SPU_004692 | Lv-Abcc10a |
| d1 | SPU_004698 | Lv-Hypp_2981 |
| d1 | SPU_004733 | Lv-Hypp_1540 |
| d1 | SPU_004785 | Lv-Gdi1 |
| d1 | SPU_004816 | Lv-Slc43a3_2 |
| d1 | SPU_004820 | Lv-Sbf1L |
| d1 | SPU_004837 | Lv-Ldla/Lrr/Gpcr_2 |
| d1 | SPU_004916 | Lv-PolppL_40 |
| d1 | SPU_004923 | Lv-Acad11 |
| d1 | SPU_004965 | Lv-Bbox1_2 |
| d1 | SPU_005290 | Lv-Slc7a9 |
| d1 | SPU_005539 | Lv-Atp13a4 |
| d1 | SPU_005547 | NA |

| | | |
|---|---|---|
| d1 | SPU_005554 | Lv-Bhmt_8 |
| d1 | SPU_005565 | NA |
| d1 | SPU_005649 | Lv-Hypp_459 |
| d1 | SPU_005651 | none |
| d1 | SPU_005768 | none |
| d1 | SPU_005854 | Lv-Syt1-1 |
| d1 | SPU_005892 | Lv-Hypp_9 |
| d1 | SPU_005931 | Lv-Cyp4L1 |
| d1 | SPU_005932 | none |
| d1 | SPU_006125 | Lv-Gucy2g |
| d1 | SPU_006127 | Lv-Svep1L_10 |
| d1 | SPU_006221 | Lv-CopineE |
| d1 | SPU_006411 | Lv-Hypp_1662 |
| d1 | SPU_006595 | none |
| d1 | SPU_006759 | Lv-Dhdh |
| d1 | SPU_006770 | none |
| d1 | SPU_007045 | Lv-Gbxf |
| d1 | SPU_007127 | Lv-Abhd3 |
| d1 | SPU_007137 | Lv-Arh/Ldl |
| d1 | SPU_007150 | Lv-Accn1 |
| d1 | SPU_007253 | Lv-Srebp |
| d1 | SPU_007700 | Lv-C6orf103 |
| d1 | SPU_007712 | none |
| d1 | SPU_007770 | Lv-CcLta1.2 |
| d1 | SPU_008095 | Lv-Adar |
| d1 | SPU_008260 | Lv-Hypp_541 |
| d1 | SPU_008629 | Lv-Armc9_1 |
| d1 | SPU_008668 | none |
| d1 | SPU_008739 | none |
| d1 | SPU_009018 | Lv-Lh2/Fn3 |
| d1 | SPU_009124 | Lv-Lisp1 |
| d1 | SPU_009256 | Lv-Hypp_573 |
| d1 | SPU_009444 | Lv-Map1a/1B_Lc3L1 |
| d1 | SPU_009506 | none |
| d1 | SPU_009583 | Lv-Slc16a4L_2 |
| d1 | SPU_009704 | none |
| d1 | SPU_010198 | NA |
| d1 | SPU_010360 | none |

| | | |
|---|---|---|
| d1 | SPU_010440 | Lv-Gpr112L_5 |
| d1 | SPU_010468 | Lv-Apn7 |
| d1 | SPU_010918 | Lv-Sulf1 |
| d1 | SPU_011498 | Lv-Rergl1 |
| d1 | SPU_011832 | Lv-Slc2a1 |
| d1 | SPU_012046 | Lv-Orctl3 |
| d1 | SPU_012089 | none |
| d1 | SPU_012119 | Lv-Dysf_3 |
| d1 | SPU_012169 | Lv-Unk_1 |
| d1 | SPU_012183 | Lv-Cytb561_1 |
| d1 | SPU_012184 | NA |
| d1 | SPU_012203 | Lv-Rel |
| d1 | SPU_012244 | none |
| d1 | SPU_012393 | Lv-Nasp |
| d1 | SPU_012417 | Lv-Unk_103 |
| d1 | SPU_012477 | Lv-Clect/Egf/PanApple |
| d1 | SPU_012568 | Lv-Hypp_2039 |
| d1 | SPU_012573 | Lv-Slc5a8 |
| d1 | SPU_012604 | none |
| d1 | SPU_012670 | Lv-Hypp_690 |
| d1 | SPU_012682 | NA |
| d1 | SPU_012817 | Lv-Pgk1 |
| d1 | SPU_012853 | Lv-Arsb_5 |
| d1 | SPU_013039 | Lv-Cyp2L26 |
| d1 | SPU_013268 | Lv-Llgl1 |
| d1 | SPU_013377 | Lv-SerpL3 |
| d1 | SPU_013451 | none |
| d1 | SPU_013540 | NA |
| d1 | SPU_013912 | Lv-Acs13 |
| d1 | SPU_014092 | Lv-Cyp3L16 |
| d1 | SPU_014111 | none |
| d1 | SPU_014394 | Lv-Hypp_2146 |
| d1 | SPU_014398 | Lv-Cbr1_1 |
| d1 | SPU_014558 | none |
| d1 | SPU_014563 | Lv-Stom_7 |
| d1 | SPU_014572 | Lv-Reeler/Egf/Cub |
| d1 | SPU_014888 | Lv-Pcmt1 |
| d1 | SPU_014912 | Lv-Ache_24 |

| | | |
|---|---|---|
| d1 | SPU_014926 | Lv-Tirc3 |
| d1 | SPU_015677 | none |
| d1 | SPU_015701 | Lv-Hrsp12 |
| d1 | SPU_015867 | Lv-Ef1ba |
| d1 | SPU_015898 | none |
| d1 | SPU_016078 | Lv-Adamts9 |
| d1 | SPU_016099 | Lv-Hypp_812 |
| d1 | SPU_016106 | Lv-Ankrd42 |
| d1 | SPU_016119 | none |
| d1 | SPU_016153 | Lv-Abca5_2 |
| d1 | SPU_016331 | Lv-Atp/Gtpbp1_1 |
| d1 | SPU_016410 | Lv-St8sia6 |
| d1 | SPU_016509 | Lv-Hypp_828 |
| d1 | SPU_016865 | none |
| d1 | SPU_016954 | none |
| d1 | SPU_017109 | Lv-Gal/lec7 |
| d1 | SPU_017317 | Lv-Slc16a6L_4 |
| d1 | SPU_017367 | Lv-Gk5 |
| d1 | SPU_017452 | Lv-NiempC1B |
| d1 | SPU_017640 | Lv-Cpn2L |
| d1 | SPU_017650 | Lv-Hypp_2349 |
| d1 | SPU_017676 | Lv-Tmtc1 |
| d1 | SPU_017768 | Lv-Dhrsx_5 |
| d1 | SPU_018023 | none |
| d1 | SPU_018097 | none |
| d1 | SPU_018237 | Lv-TlrL_9 |
| d1 | SPU_018312 | Lv-Crhr1 |
| d1 | SPU_018365 | Lv-Hypp_2381 |
| d1 | SPU_018491 | NA |
| d1 | SPU_018582 | Lv-Dyn2_lic |
| d1 | SPU_018607 | Lv-Dynlc1-2 |
| d1 | SPU_018647 | Lv-Mvp_1 |
| d1 | SPU_018713 | Lv-Notch2_4 |
| d1 | SPU_019179 | none |
| d1 | SPU_019216 | Lv-Gsto1 |
| d1 | SPU_019333 | Lv-Thnsl2 |
| d1 | SPU_019348 | Lv-Sug |
| d1 | SPU_019356 | Lv-Hpgd |

| | | |
|---|---|---|
| d1 | SPU_019641 | Lv-Agxt |
| d1 | SPU_019697 | Lv-HyalinL_5 |
| d1 | SPU_019763 | Lv-Wdr7-2 |
| d1 | SPU_019812 | Lv-Dnahc2L |
| d1 | SPU_019823 | none |
| d1 | SPU_019837 | Lv-Hunk |
| d1 | SPU_019914 | Lv-Tdo2 |
| d1 | SPU_020002 | Lv-Sgpl1_1 |
| d1 | SPU_020084 | Lv-Rdh12 |
| d1 | SPU_020145 | Lv-Klhl2-2 |
| d1 | SPU_020150 | Lv-Hypp_3024 |
| d1 | SPU_020395 | Lv-Enpp5 |
| d1 | SPU_021043 | Lv-Chrna6L_5 |
| d1 | SPU_021285 | Lv-Plscrl_3 |
| d1 | SPU_021289 | Lv-Ppar2 |
| d1 | SPU_021417 | Lv-B3galt1_18 |
| d1 | SPU_021641 | Lv-Hypp_2570 |
| d1 | SPU_021643 | Lv-Rgs20 |
| d1 | SPU_021653 | Lv-Act_3 |
| d1 | SPU_021751 | Lv-TlrL_4 |
| d1 | SPU_021965 | Lv-Dcdsh_3 |
| d1 | SPU_022370 | Lv-Echs1 |
| d1 | SPU_022501 | Lv-Ptpn13 |
| d1 | SPU_022620 | Lv-Unc44_337 |
| d1 | SPU_022639 | Lv-Alpi |
| d1 | SPU_022861 | Lv-Clect_43 |
| d1 | SPU_022928 | Lv-Rhogap8 |
| d1 | SPU_023132 | Lv-Pcs_1 |
| d1 | SPU_023309 | Lv-PskH1 |
| d1 | SPU_023322 | none |
| d1 | SPU_023331 | none |
| d1 | SPU_023334 | Lv-Pcna |
| d1 | SPU_023508 | Lv-Rabl5 |
| d1 | SPU_023604 | Lv-Adcy3 |
| d1 | SPU_023828 | none |
| d1 | SPU_024231 | Lv-GalLr |
| d1 | SPU_024243 | none |
| d1 | SPU_024307 | Lv-NfIL3 |

| | | |
|---|---|---|
| d1 | SPU_024321 | Lv-GhdcL_3 |
| d1 | SPU_024334 | Lv-Lrrc40 |
| d1 | SPU_024416 | Lv-Neurx |
| d1 | SPU_024967 | Lv-Ndufa10 |
| d1 | SPU_025019 | none |
| d1 | SPU_025179 | Lv-Nlr123 |
| d1 | SPU_025291 | none |
| d1 | SPU_025441 | Lv-Nmur2L_2 |
| d1 | SPU_025532 | Lv-Hypp_1151 |
| d1 | SPU_025620 | Lv-Cntnap2L_2 |
| d1 | SPU_025647 | Lv-Kiaa1720 |
| d1 | SPU_025879 | Lv-Gpr50L_2 |
| d1 | SPU_025929 | Lv-Slc7a14 |
| d1 | SPU_025989 | Lv-Gsr |
| d1 | SPU_026110 | Lv-Z262 |
| d1 | SPU_026129 | NA |
| d1 | SPU_026140 | Lv-Hypp_2821 |
| d1 | SPU_026190 | Lv-Gbas |
| d1 | SPU_026294 | none |
| d1 | SPU_026474 | none |
| d1 | SPU_026650 | Lv-Gsn_1 |
| d1 | SPU_026657 | Lv-Unc44_350 |
| d1 | SPU_026725 | Lv-Fn3/Ig_1 |
| d1 | SPU_026882 | Lv-CtL |
| d1 | SPU_026890 | Lv-C2orf24L |
| d1 | SPU_027344 | Lv-Ephx2_2 |
| d1 | SPU_027560 | Lv-Gnmt_5 |
| d1 | SPU_027652 | Lv-Chmp2b |
| d1 | SPU_027945 | Lv-Acad |
| d1 | SPU_028034 | none |
| d1 | SPU_028216 | Lv-TrimL_1 |
| d1 | SPU_028293 | Lv-Unc44_187 |
| d1 | SPU_028320 | none |
| d1 | SPU_028334 | none |
| d1 | SPU_028443 | Lv-Sarox_3 |
| d1 | SPU_028878 | Lv-Hypp_3025 |
| d1 | SPU_028909 | Lv-Pde3bL |
| d1 | SPU_028938 | Lv-Naalad2L_9 |

| | | |
|---|---|---|
| d1 | SPU_030222 | Lv-Axndhc4h |
| d1 | SPU_030226 | Lv-Axndhc5h |
| d2 | SPU_000063 | Lv-Rdh8 |
| d2 | SPU_000070 | Lv-MucL |
| d2 | SPU_000226 | none |
| d2 | SPU_000469 | Lv-Igcamr |
| d2 | SPU_000646 | Lv-Srcr4 |
| d2 | SPU_000684 | Lv-Hypp_1274 |
| d2 | SPU_000694 | Lv-Ugt2b34 |
| d2 | SPU_000765 | Lv-Chrna9L_2 |
| d2 | SPU_000927 | Lv-Rab22 |
| d2 | SPU_000971 | Lv-Slc6a2_1 |
| d2 | SPU_001090 | none |
| d2 | SPU_001255 | Lv-DmrtA2 |
| d2 | SPU_001273 | Lv-Slc5a12_7 |
| d2 | SPU_001298 | Lv-AcheL_2 |
| d2 | SPU_001449 | Lv-Dmbt1 |
| d2 | SPU_001464 | Lv-Hypp_1330 |
| d2 | SPU_001638 | Lv-ThrombA1 |
| d2 | SPU_001720 | Lv-Hypp_1343 |
| d2 | SPU_002133 | Lv-Mamdc4-5 |
| d2 | SPU_002315 | Lv-Hypp_351 |
| d2 | SPU_002376 | Lv-Bhmt2 |
| d2 | SPU_002384 | none |
| d2 | SPU_002430 | Lv-Hypp_1394 |
| d2 | SPU_002448 | Lv-Sage |
| d2 | SPU_002482 | Lv-Rrm1 |
| d2 | SPU_002515 | Lv-Hypp_1399 |
| d2 | SPU_002632 | Lv-Hox11/13a |
| d2 | SPU_002650 | Lv-Aldh1l1 |
| d2 | SPU_002686 | Lv-Gucy1b2 |
| d2 | SPU_002744 | Lv-St8sia2 |
| d2 | SPU_002755 | Lv-Trim56 |
| d2 | SPU_002782 | none |
| d2 | SPU_002810 | Lv-AmidL |
| d2 | SPU_002965 | Lv-AcheL_3 |
| d2 | SPU_003656 | Lv-Hypp_1467 |
| d2 | SPU_003816 | Lv-Mamdc2-4 |

| | | |
|---|---|---|
| d2 | SPU_003858 | Lv-Hs3st6 |
| d2 | SPU_003894 | Lv-Btub5 |
| d2 | SPU_003914 | Lv-Plscr1 |
| d2 | SPU_004023 | Lv-Fnbp1l |
| d2 | SPU_004032 | Lv-Fam151b |
| d2 | SPU_004180 | Lv-Dur3 |
| d2 | SPU_004181 | Lv-Bche-3 |
| d2 | SPU_004183 | Lv-AchE_3 |
| d2 | SPU_004229 | Lv-Dpp10 |
| d2 | SPU_004246 | Lv-Wdr49L |
| d2 | SPU_004328 | Lv-Ddm36L |
| d2 | SPU_004985 | Lv-Npr1_6 |
| d2 | SPU_005193 | Lv-Tcp2 |
| d2 | SPU_005196 | Lv-Slc22a3_1 |
| d2 | SPU_005239 | Lv-Ldla/Lrr/Gpcr_1 |
| d2 | SPU_005391 | Lv-AapL |
| d2 | SPU_005613 | Lv-Titin |
| d2 | SPU_005767 | Lv-Smpd3 |
| d2 | SPU_005842 | Lv-Glyphp1 |
| d2 | SPU_006107 | Lv-Flj46688l2 |
| d2 | SPU_006241 | Lv-Rhogap6 |
| d2 | SPU_006301 | Lv-Slc18a2 |
| d2 | SPU_006306 | Lv-Gal/lec3 |
| d2 | SPU_006381 | Lv-Slc1a3 |
| d2 | SPU_006382 | Lv-Slc1a1_1 |
| d2 | SPU_006474 | Lv-Hypp_2985 |
| d2 | SPU_006499 | Lv-Sfrp3/4II |
| d2 | SPU_006536 | Lv-Rims2 |
| d2 | SPU_006578 | Lv-C1qL_1 |
| d2 | SPU_006582 | Lv-Hectd2 |
| d2 | SPU_006787 | Lv-Pax6_1 |
| d2 | SPU_006797 | Lv-Msclact |
| d2 | SPU_007014 | Lv-Abcb4a |
| d2 | SPU_007015 | Lv-Abcb1 |
| d2 | SPU_007135 | none |
| d2 | SPU_007162 | Lv-Galr2L_8 |
| d2 | SPU_007225 | Lv-Tin/Ag |
| d2 | SPU_007345 | Lv-Chst11_1 |

| | | |
|---|---|---|
| d2 | SPU_007450 | none |
| d2 | SPU_007451 | Lv-Hypp_1723 |
| d2 | SPU_007507 | Lv-MucA |
| d2 | SPU_007583 | Lv-B3galt1_5 |
| d2 | SPU_007664 | Lv-Esd |
| d2 | SPU_007681 | Lv-Hypp_1744 |
| d2 | SPU_007738 | NA |
| d2 | SPU_007798 | Lv-Asp_9 |
| d2 | SPU_007819 | NA |
| d2 | SPU_008443 | Lv-Abcb4b |
| d2 | SPU_008653 | Lv-Igf_23 |
| d2 | SPU_008701 | Lv-Upb1 |
| d2 | SPU_008762 | Lv-Hypp_36 |
| d2 | SPU_008799 | Lv-Dynlc1-3a |
| d2 | SPU_008860 | Lv-Timp2c |
| d2 | SPU_008865 | Lv-Timp4 |
| d2 | SPU_008988 | Lv-Mglur3_2 |
| d2 | SPU_009139 | Lv-Slc5a8_1 |
| d2 | SPU_009319 | Lv-Tryp_SPc/LDL/Cub |
| d2 | SPU_009495 | Lv-Rgn |
| d2 | SPU_009584 | Lv-Gabbr2L_1 |
| d2 | SPU_009765 | Lv-Accn1L |
| d2 | SPU_009885 | Lv-ManrC1A |
| d2 | SPU_009976 | none |
| d2 | SPU_009990 | Lv-Tk7 |
| d2 | SPU_010052 | Lv-Nlr102_1 |
| d2 | SPU_010054 | Lv-My18A_putative_homolog |
| d2 | SPU_010073 | Lv-Lhfp |
| d2 | SPU_010106 | Lv-Kiaa1529L_1 |
| d2 | SPU_010216 | Lv-Clect/Fn3_19 |
| d2 | SPU_010313 | Lv-Gal/lec5 |
| d2 | SPU_010354 | Lv-Rh50 |
| d2 | SPU_010811 | Lv-Sgarp1 |
| d2 | SPU_010827 | Lv-Unc5ch |
| d2 | SPU_010842 | Lv-Mgl |
| d2 | SPU_010883 | Lv-Htr4L_6 |
| d2 | SPU_010959 | Lv-Slc44a2_4 |
| d2 | SPU_011006 | Lv-Unc44L_69 |

| | | |
|---|---|---|
| d2 | SPU_011061 | Lv-Gcnf1_1 |
| d2 | SPU_011106 | Lv-Anxn |
| d2 | SPU_011152 | Lv-Rps18_1 |
| d2 | SPU_011171 | none |
| d2 | SPU_011202 | Lv-Meis |
| d2 | SPU_011363 | Lv-Neto2L |
| d2 | SPU_011451 | Lv-Glyphp3 |
| d2 | SPU_011529 | Lv-Ttrspn_8 |
| d2 | SPU_011609 | Lv-Chrna7_4 |
| d2 | SPU_011657 | Lv-Mmr2L_1 |
| d2 | SPU_011666 | none |
| d2 | SPU_012014 | Lv-NefhL_1 |
| d2 | SPU_012056 | Lv-Hypp_2990 |
| d2 | SPU_012127 | none |
| d2 | SPU_012186 | Lv-AchE_9 |
| d2 | SPU_012222 | Lv-Aqp4 |
| d2 | SPU_012237 | Lv-Hrh4L_5 |
| d2 | SPU_012260 | Lv-Hypp_2023 |
| d2 | SPU_012479 | Lv-Clect/Tm_12 |
| d2 | SPU_012526 | Lv-Hypp_2038 |
| d2 | SPU_012548 | Lv-Sparc_2 |
| d2 | SPU_012609 | none |
| d2 | SPU_012663 | Lv-Slc22a3_4 |
| d2 | SPU_012664 | none |
| d2 | SPU_012799 | Lv-Orxn/HpcLr1 |
| d2 | SPU_013331 | none |
| d2 | SPU_013615 | Lv-Plscrl_2 |
| d2 | SPU_013917 | Lv-ProjL |
| d2 | SPU_014108 | Lv-Tctex1d2 |
| d2 | SPU_014339 | Lv-Rdh11_2 |
| d2 | SPU_014374 | Lv-Hypp_752 |
| d2 | SPU_014492 | Lv-Msp130r6 |
| d2 | SPU_014739 | Lv-Hypp_763 |
| d2 | SPU_014897 | Lv-Orct_1 |
| d2 | SPU_015069 | Lv-Acsl5_3 |
| d2 | SPU_015111 | Lv-Pnlip_6 |
| d2 | SPU_015290 | Lv-Redt |
| d2 | SPU_015595 | Lv-Calpn |

| | | |
|---|---|---|
| d2 | SPU_015633 | Lv-MucL_5 |
| d2 | SPU_015656 | Lv-Tmem20_8 |
| d2 | SPU_015750 | Lv-Nlgn4y |
| d2 | SPU_015763 | Lv-Msp130r5 |
| d2 | SPU_015826 | Lv-Nagk |
| d2 | SPU_015872 | NA |
| d2 | SPU_016007 | Lv-Btub8 |
| d2 | SPU_016176 | Lv-Gdpd1_1 |
| d2 | SPU_016177 | Lv-MachrM5 |
| d2 | SPU_016282 | Lv-Hypp_2255 |
| d2 | SPU_016300 | Lv-Ccp/Zp_1 |
| d2 | SPU_016492 | NA |
| d2 | SPU_016592 | none |
| d2 | SPU_016736 | Lv-NprA_2 |
| d2 | SPU_016802 | none |
| d2 | SPU_016980 | Lv-Cadherin_1 |
| d2 | SPU_016981 | Lv-Notch1-13 |
| d2 | SPU_017051 | Lv-Slc13a2 |
| d2 | SPU_017143 | Lv-Stard5 |
| d2 | SPU_017252 | Lv-Fmo5_2 |
| d2 | SPU_017488 | Lv-Fn3/Igf_38 |
| d2 | SPU_017568 | Lv-Ankrd50L |
| d2 | SPU_017652 | Lv-Gba_5 |
| d2 | SPU_018190 | Lv-NmbrL_3 |
| d2 | SPU_018442 | Lv-Scnn1bL |
| d2 | SPU_018466 | Lv-AkrL |
| d2 | SPU_018615 | Lv-Slc25a3 |
| d2 | SPU_018814 | Lv-Accn3L |
| d2 | SPU_018910 | none |
| d2 | SPU_019203 | Lv-AchE_6 |
| d2 | SPU_019317 | Lv-Rbp1b |
| d2 | SPU_019408 | Lv-Cklf4 |
| d2 | SPU_019567 | Lv-Clrn1 |
| d2 | SPU_019656 | Lv-Cdxl |
| d2 | SPU_019810 | Lv-Notch_1 |
| d2 | SPU_020076 | Lv-Hypp_164 |
| d2 | SPU_020276 | none |
| d2 | SPU_020744 | Lv-Vwd/Vwc/Ldla |

| | | |
|---|---|---|
| d2 | SPU_020754 | Lv-Cyp2L45 |
| d2 | SPU_020803 | Lv-Trk |
| d2 | SPU_020906 | Lv-QrfprL_5 |
| d2 | SPU_020920 | Lv-Actinin |
| d2 | SPU_021033 | Lv-Ache_25 |
| d2 | SPU_021344 | none |
| d2 | SPU_021465 | Lv-Gucy1b2_1 |
| d2 | SPU_021663 | none |
| d2 | SPU_021775 | Lv-Slc16a6_4 |
| d2 | SPU_022006 | Lv-Nme7 |
| d2 | SPU_022135 | Lv-Sdhc |
| d2 | SPU_022261 | Lv-Hypp_1042 |
| d2 | SPU_022300 | Lv-Stom_9 |
| d2 | SPU_022395 | Lv-Stk2m829 |
| d2 | SPU_022596 | Lv-Fmo5_3 |
| d2 | SPU_022663 | none |
| d2 | SPU_022667 | Lv-ThrombB2 |
| d2 | SPU_022706 | Lv-Vwd/Egf/Fn3_1 |
| d2 | SPU_022712 | Lv-Hypp_2624 |
| d2 | SPU_022896 | Lv-Colf_8 |
| d2 | SPU_022929 | Lv-Lama2f_1 |
| d2 | SPU_023003 | Lv-Nvd |
| d2 | SPU_023030 | Lv-Hypp_2644 |
| d2 | SPU_023068 | Lv-Cyp2L54 |
| d2 | SPU_023096 | Lv-Ptgr1 |
| d2 | SPU_023142 | Lv-Ttn2 |
| d2 | SPU_023330 | Lv-Dpd1L_1 |
| d2 | SPU_023475 | none |
| d2 | SPU_023488 | Lv-Clect/Fn3_7 |
| d2 | SPU_023538 | Lv-Fut4_5 |
| d2 | SPU_023643 | Lv-RasDL |
| d2 | SPU_023660 | Lv-CbpdDL_1 |
| d2 | SPU_023693 | Lv-Anpep_1 |
| d2 | SPU_023777 | none |
| d2 | SPU_023899 | Lv-122Kdsp_1 |
| d2 | SPU_024049 | Lv-Kcnk18 |
| d2 | SPU_024179 | Lv-Mmr2L |
| d2 | SPU_024328 | Lv-Rdh7 |

| | | |
|---|---|---|
| d2 | SPU_024362 | Lv-Ttrspn_18 |
| d2 | SPU_024459 | none |
| d2 | SPU_024498 | Lv-Dynlc2-5b |
| d2 | SPU_024597 | Lv-PcolceL |
| d2 | SPU_025072 | Lv-Hebp2_1 |
| d2 | SPU_025498 | Lv-Hypp_2793 |
| d2 | SPU_025548 | Lv-Gst11 |
| d2 | SPU_025621 | Lv-Cntnap2L_3 |
| d2 | SPU_025791 | Lv-Apcdd1 |
| d2 | SPU_025895 | Lv-Hypp_2807 |
| d2 | SPU_025983 | Lv-Srcr210 |
| d2 | SPU_025992 | Lv-Slc29a4_1 |
| d2 | SPU_025997 | Lv-Scn5a |
| d2 | SPU_025999 | Lv-Mglur3_1 |
| d2 | SPU_026031 | Lv-Thymbh |
| d2 | SPU_026041 | Lv-Thsd4L_2 |
| d2 | SPU_026046 | none |
| d2 | SPU_026135 | Lv-Lph |
| d2 | SPU_026220 | Lv-Slc5a9_9 |
| d2 | SPU_026328 | Lv-Mgst2 |
| d2 | SPU_026371 | Lv-Nckx |
| d2 | SPU_026399 | Lv-Fn3/Ig/Egf |
| d2 | SPU_026470 | Lv-CysKL |
| d2 | SPU_026489 | Lv-Tpcn3 |
| d2 | SPU_026798 | Lv-Adam/TsL3 |
| d2 | SPU_027052 | none |
| d2 | SPU_027083 | Lv-Sap30L |
| d2 | SPU_027119 | Lv-Vwd/Vwcp_1 |
| d2 | SPU_027224 | Lv-Slc5a9_7 |
| d2 | SPU_027378 | Lv-Plscrl |
| d2 | SPU_027390 | Lv-Ewamph |
| d2 | SPU_027392 | Lv-Nipa2_1 |
| d2 | SPU_027400 | Lv-Ph4_1 |
| d2 | SPU_027486 | Lv-Apn5a |
| d2 | SPU_027502 | Lv-Slc22a13_1 |
| d2 | SPU_027507 | none |
| d2 | SPU_027620 | none |
| d2 | SPU_027926 | Lv-EpimL |

| | | |
|---|---|---|
| d2 | SPU_028175 | Lv-Nox5_1 |
| d2 | SPU_028199 | Lv-Amy_8 |
| d2 | SPU_028317 | Lv-Hypp_2950 |
| d2 | SPU_028361 | Lv-Unc44_188 |
| d2 | SPU_028575 | NA |
| d2 | SPU_028585 | Lv-C2orf21L_2 |
| d2 | SPU_028704 | none |
| d2 | SPU_028757 | Lv-Aadac_3 |
| d2 | SPU_028843 | Lv-Ccp/Zp_2 |
| d2 | SPU_028905 | Lv-Pger2ep4 |
| d2 | SPU_030073 | Lv-Nt |
| d2 | SPU_030086 | NA |
| d2 | SPU_030153 | Lv-Dbx |
| d3 | SPU_000105 | Lv-Mapkkk12/13 |
| d3 | SPU_000163 | Lv-Sult4a1 |
| d3 | SPU_000167 | Lv-Chac |
| d3 | SPU_000174 | Lv-Slc30a3_1 |
| d3 | SPU_000362 | Lv-Masp |
| d3 | SPU_000443 | none |
| d3 | SPU_000671 | Lv-Hypp_1272 |
| d3 | SPU_000795 | none |
| d3 | SPU_000839 | Lv-EchL1 |
| d3 | SPU_000941 | none |
| d3 | SPU_001517 | none |
| d3 | SPU_001572 | NA |
| d3 | SPU_001863 | Lv-Srcr20 |
| d3 | SPU_001893 | Lv-Clect/Fn3 |
| d3 | SPU_002010 | Lv-Svep1L_3 |
| d3 | SPU_002061 | Lv-Epdmnrp |
| d3 | SPU_002271 | Lv-Kctd21-2 |
| d3 | SPU_002452 | Lv-Hypp_1397 |
| d3 | SPU_002786 | none |
| d3 | SPU_003056 | NA |
| d3 | SPU_003182 | Lv-Pdcd4 |
| d3 | SPU_003720 | Lv-Hypp_3093 |
| d3 | SPU_003820 | Lv-Tmem195-2 |
| d3 | SPU_004221 | Lv-Hypp_1507 |
| d3 | SPU_004369 | Lv-Hypp_1517 |

| | | |
|---|---|---|
| d3 | SPU_004412 | Lv-Cub/Egf |
| d3 | SPU_004813 | Lv-Slc25a5 |
| d3 | SPU_004859 | Lv-Plekhb2 |
| d3 | SPU_004867 | Lv-Sm30E |
| d3 | SPU_004966 | none |
| d3 | SPU_005014 | Lv-Hypp_1556 |
| d3 | SPU_005153 | none |
| d3 | SPU_005160 | Lv-Cyp2L11 |
| d3 | SPU_005200 | Lv-Hypp_3 |
| d3 | SPU_005256 | Lv-Abat |
| d3 | SPU_005324 | none |
| d3 | SPU_005481 | none |
| d3 | SPU_005782 | Lv-Loc201475 |
| d3 | SPU_006004 | Lv-Tsp1k |
| d3 | SPU_006016 | Lv-Nlr203 |
| d3 | SPU_006041 | none |
| d3 | SPU_006084 | Lv-Ccp/Egf/Tsp1/HyalinL1 |
| d3 | SPU_006732 | Lv-ReelerL_7 |
| d3 | SPU_007004 | Lv-Bmp11 |
| d3 | SPU_007155 | Lv-Gapd |
| d3 | SPU_007174 | Lv-Glxn |
| d3 | SPU_007198 | Lv-Hypp_1704 |
| d3 | SPU_007242 | Lv-Rough |
| d3 | SPU_007251 | Lv-Sqst1L |
| d3 | SPU_007284 | Lv-Aldh2_1 |
| d3 | SPU_007431 | Lv-Acy1l2p |
| d3 | SPU_007677 | Lv-Hypp_27 |
| d3 | SPU_007800 | Lv-Sspo |
| d3 | SPU_007931 | Lv-Hypp_1754 |
| d3 | SPU_007946 | Lv-Pgrp2 |
| d3 | SPU_008050 | none |
| d3 | SPU_008202 | Lv-Hypp_9771 |
| d3 | SPU_008222 | Lv-C10orf112L |
| d3 | SPU_008389 | Lv-Bbox1_3 |
| d3 | SPU_008596 | Lv-Hypp_555 |
| d3 | SPU_008605 | Lv-Gpr126 |
| d3 | SPU_008639 | Lv-Acsf2_1 |
| d3 | SPU_008646 | Lv-Slc5a7_2 |

| | | |
|---|---|---|
| d3 | SPU_008703 | Lv-Xbp1 |
| d3 | SPU_008708 | Lv-Bcnt2L |
| d3 | SPU_008825 | Lv-Hao1_5 |
| d3 | SPU_009020 | Lv-C1qL_2 |
| d3 | SPU_009312 | Lv-Chrna5L |
| d3 | SPU_009378 | Lv-Vtgn1 |
| d3 | SPU_009528 | Lv-TnfsfL1 |
| d3 | SPU_009579 | none |
| d3 | SPU_009798 | none |
| d3 | SPU_009954 | Lv-Tsp1 |
| d3 | SPU_010404 | Lv-Irf |
| d3 | SPU_010579 | Lv-Pkd2 |
| d3 | SPU_010634 | Lv-Z447 |
| d3 | SPU_010967 | Lv-Cub/TolloidL |
| d3 | SPU_011090 | Lv-AadacL_4 |
| d3 | SPU_011210 | Lv-Unc44_293 |
| d3 | SPU_011246 | Lv-IrxB |
| d3 | SPU_011359 | Lv-Unc44L_72 |
| d3 | SPU_011469 | Lv-Ache_20 |
| d3 | SPU_011830 | Lv-Mco1 |
| d3 | SPU_012039 | Lv-Srcr104 |
| d3 | SPU_012126 | Lv-Hypp_2018 |
| d3 | SPU_012181 | NA |
| d3 | SPU_012667 | Lv-Saa-a |
| d3 | SPU_012793 | Lv-Trpc6_1 |
| d3 | SPU_012906 | Lv-Pnlip_2 |
| d3 | SPU_013084 | Lv-Sr/Fa58c/Lnb/7Tm/Gpcr |
| d3 | SPU_013167 | Lv-Vstcnlr |
| d3 | SPU_013243 | none |
| d3 | SPU_013444 | Lv-Arih1L |
| d3 | SPU_013458 | Lv-Cara4LA |
| d3 | SPU_013698 | Lv-Nrl |
| d3 | SPU_013752 | none |
| d3 | SPU_014013 | Lv-Abcg2B |
| d3 | SPU_014095 | Lv-Srcr114 |
| d3 | SPU_014164 | Lv-Barh1 |
| d3 | SPU_014540 | Lv-Trim2-4 |
| d3 | SPU_014840 | Lv-RasD |

| | | |
|---|---|---|
| d3 | SPU_015031 | Lv-Gpr51_4 |
| d3 | SPU_015112 | Lv-Pnlip_7 |
| d3 | SPU_015283 | Lv-Slc27a2_1 |
| d3 | SPU_015323 | Lv-Ckb |
| d3 | SPU_015350 | Lv-Ckm |
| d3 | SPU_015423 | Lv-Selenbp1 |
| d3 | SPU_015429 | Lv-Ef1 |
| d3 | SPU_015471 | Lv-Hypp_2215 |
| d3 | SPU_015716 | Lv-Slc22a3_10 |
| d3 | SPU_015737 | Lv-Bbox1_4 |
| d3 | SPU_015749 | Lv-Unc93ha |
| d3 | SPU_015987 | Lv-Hao1_9 |
| d3 | SPU_016122 | none |
| d3 | SPU_016179 | Lv-Lrp4-2 |
| d3 | SPU_016220 | NA |
| d3 | SPU_016345 | Lv-Pemt_1 |
| d3 | SPU_016407 | none |
| d3 | SPU_016746 | Lv-Atub3 |
| d3 | SPU_016767 | Lv-Aldh5a1_1 |
| d3 | SPU_016816 | Lv-Cyp2L33 |
| d3 | SPU_016853 | Lv-Slc23a2 |
| d3 | SPU_017458 | Lv-Hypp_2326 |
| d3 | SPU_017586 | Lv-AgrnL |
| d3 | SPU_017843 | Lv-Miox |
| d3 | SPU_018000 | none |
| d3 | SPU_018156 | Lv-Mt5/6L1 |
| d3 | SPU_018169 | Lv-Cntnap4L_2 |
| d3 | SPU_018272 | Lv-Txndc16 |
| d3 | SPU_018327 | Lv-Tmem20L |
| d3 | SPU_018433 | Lv-Slc5a9_3 |
| d3 | SPU_018502 | none |
| d3 | SPU_018744 | Lv-GlisB |
| d3 | SPU_019088 | Lv-Clect_112 |
| d3 | SPU_019104 | none |
| d3 | SPU_019176 | none |
| d3 | SPU_019184 | Lv-Rem |
| d3 | SPU_019228 | Lv-MachrM4 |
| d3 | SPU_019318 | Lv-Rbp2 |

| | | |
|---|---|---|
| d3 | SPU_019723 | Lv-Dioxg |
| d3 | SPU_019779 | Lv-Rej5 |
| d3 | SPU_020026 | Lv-Slc5a7_4 |
| d3 | SPU_020229 | Lv-Cyp4VL1 |
| d3 | SPU_020408 | Lv-Rxfp2L_6 |
| d3 | SPU_020413 | Lv-Hunk_1 |
| d3 | SPU_020488 | Lv-R/Ptp/R_2 |
| d3 | SPU_020500 | Lv-Hsp70 |
| d3 | SPU_020597 | Lv-Srcr159 |
| d3 | SPU_020794 | Lv-Lphn1L |
| d3 | SPU_020926 | NA |
| d3 | SPU_021046 | Lv-Arsa_2 |
| d3 | SPU_021060 | Lv-Ccbl2 |
| d3 | SPU_021173 | Lv-Fos |
| d3 | SPU_021281 | none |
| d3 | SPU_021602 | Lv-eGas |
| d3 | SPU_021660 | Lv-Crea_1 |
| d3 | SPU_021688 | Lv-Ypel2 |
| d3 | SPU_021761 | Lv-Accn2L |
| d3 | SPU_021781 | Lv-Slc5a5_1 |
| d3 | SPU_021794 | Lv-Gpr45L |
| d3 | SPU_022053 | Lv-Serrate |
| d3 | SPU_022122 | none |
| d3 | SPU_022148 | Lv-Srcr179 |
| d3 | SPU_022151 | Lv-Srcr182 |
| d3 | SPU_022178 | Lv-185/333E2 |
| d3 | SPU_022476 | Lv-Taldo1 |
| d3 | SPU_022508 | Lv-Sult1c4 |
| d3 | SPU_022552 | Lv-AapL |
| d3 | SPU_022815 | NA |
| d3 | SPU_022825 | Lv-Hypp_191 |
| d3 | SPU_022914 | Lv-Slc17a5_8 |
| d3 | SPU_023145 | Lv-SelpL_1 |
| d3 | SPU_023163 | none |
| d3 | SPU_023233 | Lv-Slsp_16 |
| d3 | SPU_023376 | none |
| d3 | SPU_023388 | Lv-Hypp_2669 |
| d3 | SPU_023409 | Lv-Reelin2 |

| | | |
|---|---|---|
| d3 | SPU_023425 | Lv-Syt4 |
| d3 | SPU_023487 | Lv-Clect/Fn3_10 |
| d3 | SPU_023505 | Lv-Mt5/MmpL1 |
| d3 | SPU_023971 | NA |
| d3 | SPU_023996 | Lv-Unc44_168 |
| d3 | SPU_024010 | Lv-C1qL_9 |
| d3 | SPU_024104 | Lv-Lhfpl3 |
| d3 | SPU_024319 | none |
| d3 | SPU_024381 | none |
| d3 | SPU_024455 | Lv-PlbL_1 |
| d3 | SPU_024458 | Lv-Unc44_344 |
| d3 | SPU_024559 | Lv-LacL_7 |
| d3 | SPU_024578 | Lv-24578 |
| d3 | SPU_024674 | Lv-Rergl5 |
| d3 | SPU_024989 | Lv-Rdh8_9 |
| d3 | SPU_025097 | Lv-Clect_96 |
| d3 | SPU_025128 | Lv-St14L_4 |
| d3 | SPU_025340 | Lv-Sult1c_1 |
| d3 | SPU_025537 | Lv-MethsynthII |
| d3 | SPU_025575 | Lv-Srap |
| d3 | SPU_025701 | Lv-Nbr1 |
| d3 | SPU_025901 | Lv-Hypp_1159 |
| d3 | SPU_025931 | Lv-Slc7a14_1 |
| d3 | SPU_026064 | Lv-GadD45 |
| d3 | SPU_027236 | Lv-Vdac2_1 |
| d3 | SPU_027328 | Lv-Mt5/6L2 |
| d3 | SPU_027436 | none |
| d3 | SPU_027556 | Lv-KlkB |
| d3 | SPU_027635 | Lv-Alpl_2 |
| d3 | SPU_027770 | Lv-Hypp_1213 |
| d3 | SPU_027992 | Lv-Slsp_26 |
| d3 | SPU_028091 | Lv-Echn15 |
| d3 | SPU_028629 | Lv-Rdh8_11 |
| d3 | SPU_030074 | Lv-Ngfffap |
| e01 | SPU_000055 | Lv-Strap |
| e01 | SPU_000116 | Lv-Gucy1a3 |
| e01 | SPU_000168 | Lv-Rangap1 |
| e01 | SPU_000178 | Lv-PP2aB56e |

| | | |
|---|---|---|
| e01 | SPU_000181 | Lv-Rt2L |
| e01 | SPU_000225 | Lv-Mphosph10l |
| e01 | SPU_000344 | none |
| e01 | SPU_000525 | Lv-Cercam |
| e01 | SPU_000536 | Lv-Dom3zL_2 |
| e01 | SPU_000540 | Lv-Z98 |
| e01 | SPU_000551 | Lv-Hypp_306 |
| e01 | SPU_000568 | Lv-Hypp_1261 |
| e01 | SPU_000582 | Lv-Herc4_2 |
| e01 | SPU_000600 | none |
| e01 | SPU_000619 | Lv-ApdpL |
| e01 | SPU_000658 | Lv-Hypp_1270 |
| e01 | SPU_000675 | Lv-NotchL_2 |
| e01 | SPU_000712 | Lv-Hypp_1283 |
| e01 | SPU_000731 | Lv-Cln3 |
| e01 | SPU_000734 | none |
| e01 | SPU_000753 | none |
| e01 | SPU_000814 | Lv-Acvr2_1 |
| e01 | SPU_000820 | Lv-Slcoa4a1 |
| e01 | SPU_001001 | Lv-Hypp_320 |
| e01 | SPU_001068 | Lv-Kctd5 |
| e01 | SPU_001132 | Lv-CoQ10hb/a |
| e01 | SPU_001156 | Lv-Adck5 |
| e01 | SPU_001212 | Lv-Nhedc2-2 |
| e01 | SPU_001237 | Lv-Psgp |
| e01 | SPU_001338 | Lv-PolppL |
| e01 | SPU_001356 | Lv-Slc9a8 |
| e01 | SPU_001425 | Lv-C18orf51L |
| e01 | SPU_001491 | Lv-Fvt1 |
| e01 | SPU_001551 | Lv-ArL6ip1 |
| e01 | SPU_001594 | Lv-Hypp_3115 |
| e01 | SPU_001681 | Lv-Nkapl |
| e01 | SPU_001691 | Lv-Hypp_1342 |
| e01 | SPU_001713 | none |
| e01 | SPU_001724 | none |
| e01 | SPU_001736 | Lv-Snrnp25 |
| e01 | SPU_001768 | Lv-LamB2Lf |
| e01 | SPU_001829 | Lv-Smurf2 |

| | | |
|---|---|---|
| e01 | SPU_001836 | Lv-Xpac |
| e01 | SPU_001872 | Lv-Polr2a-2 |
| e01 | SPU_002070 | Lv-Hagb |
| e01 | SPU_002072 | Lv-Mfsd6 |
| e01 | SPU_002138 | Lv-Hypp_1373 |
| e01 | SPU_002183 | Lv-Adeamc |
| e01 | SPU_002211 | none |
| e01 | SPU_002228 | Lv-Gtf2h1 |
| e01 | SPU_002249 | none |
| e01 | SPU_002374 | Lv-Fermth3 |
| e01 | SPU_002451 | Lv-Hypp_1396 |
| e01 | SPU_002508 | Lv-Trip12 |
| e01 | SPU_002553 | Lv-Rad21 |
| e01 | SPU_002586 | Lv-Z366 |
| e01 | SPU_002695 | none |
| e01 | SPU_002821 | Lv-Serrp6 |
| e01 | SPU_002838 | Lv-Hyal1-2 |
| e01 | SPU_003045 | Lv-Pmyh |
| e01 | SPU_003113 | none |
| e01 | SPU_003159 | NA |
| e01 | SPU_003184 | none |
| e01 | SPU_003189 | none |
| e01 | SPU_003197 | Lv-Snw1 |
| e01 | SPU_003230 | none |
| e01 | SPU_003346 | none |
| e01 | SPU_003500 | Lv-Jarid1d |
| e01 | SPU_003537 | Lv-Usp52_1 |
| e01 | SPU_003633 | Lv-Mcm9 |
| e01 | SPU_003702 | Lv-Sh2d4b |
| e01 | SPU_003742 | Lv-Alox5 |
| e01 | SPU_003758 | Lv-Hypp_3095 |
| e01 | SPU_003772 | Lv-Cenpc1 |
| e01 | SPU_003806 | Lv-Ei2bL |
| e01 | SPU_003857 | Lv-Trip12 |
| e01 | SPU_004002 | Lv-Nat12 |
| e01 | SPU_004046 | none |
| e01 | SPU_004069 | Lv-122E |
| e01 | SPU_004178 | none |

| | | |
|---|---|---|
| e01 | SPU_004189 | Lv-Depdc5-2 |
| e01 | SPU_004226 | Lv-Elovl4_2 |
| e01 | SPU_004336 | NA |
| e01 | SPU_004344 | none |
| e01 | SPU_004397 | Lv-Gpx7 |
| e01 | SPU_004423 | NA |
| e01 | SPU_004489 | Lv-Tbc1d25 |
| e01 | SPU_004518 | Lv-Bub1 |
| e01 | SPU_004541 | none |
| e01 | SPU_004562 | Lv-Slc38a9 |
| e01 | SPU_004588 | Lv-Setd2 |
| e01 | SPU_004595 | Lv-Nek8 |
| e01 | SPU_004645 | Lv-Dgcr14L |
| e01 | SPU_004657 | Lv-Dtlh |
| e01 | SPU_004713 | Lv-Stmn1 |
| e01 | SPU_004717 | Lv-Spry/Socscp |
| e01 | SPU_004722 | Lv-Wdr26 |
| e01 | SPU_004724 | none |
| e01 | SPU_004743 | Lv-Nt5c2L1 |
| e01 | SPU_004794 | none |
| e01 | SPU_004861 | Lv-Srp7 |
| e01 | SPU_004984 | Lv-Sec62_1 |
| e01 | SPU_004988 | Lv-Brwd2 |
| e01 | SPU_005044 | NA |
| e01 | SPU_005063 | Lv-Rio3 |
| e01 | SPU_005180 | Lv-Hmcn1L_2 |
| e01 | SPU_005212 | none |
| e01 | SPU_005215 | Lv-Ripk1-4A |
| e01 | SPU_005222 | NA |
| e01 | SPU_005229 | Lv-Z440 |
| e01 | SPU_005272 | Lv-Rab15 |
| e01 | SPU_005278 | Lv-C15orf58L |
| e01 | SPU_005313 | Lv-Gal3st |
| e01 | SPU_005352 | none |
| e01 | SPU_005364 | Lv-Adf1L |
| e01 | SPU_005378 | Lv-Sirt3 |
| e01 | SPU_005384 | none |
| e01 | SPU_005411 | Lv-Nek8_1 |

| | | |
|---|---|---|
| e01 | SPU_005559 | Lv-Z356 |
| e01 | SPU_005592 | Lv-Enpp3_1 |
| e01 | SPU_005637 | NA |
| e01 | SPU_005661 | NA |
| e01 | SPU_005707 | Lv-Mgat4c |
| e01 | SPU_005714 | Lv-CtdbdL |
| e01 | SPU_005726 | NA |
| e01 | SPU_005747 | Lv-Lrrc56 |
| e01 | SPU_005798 | Lv-TrypsinL_3 |
| e01 | SPU_005888 | Lv-Clect_66 |
| e01 | SPU_005922 | Lv-Hypp_1616 |
| e01 | SPU_005979 | Lv-Gpc1246 |
| e01 | SPU_006063 | Lv-Z69 |
| e01 | SPU_006118 | Lv-Lamg1f |
| e01 | SPU_006186 | Lv-Pms1 |
| e01 | SPU_006234 | none |
| e01 | SPU_006239 | Lv-Pcf11 |
| e01 | SPU_006257 | Lv-Man1a2_2 |
| e01 | SPU_006320 | none |
| e01 | SPU_006366 | none |
| e01 | SPU_006369 | Lv-Fdxacb1L |
| e01 | SPU_006427 | Lv-Z126 |
| e01 | SPU_006428 | Lv-Kiaa1704 |
| e01 | SPU_006472 | none |
| e01 | SPU_006539 | Lv-Srcr52 |
| e01 | SPU_006568 | NA |
| e01 | SPU_006596 | Lv-Swap70 |
| e01 | SPU_006641 | Lv-Trim3L_1 |
| e01 | SPU_006665 | Lv-Manba |
| e01 | SPU_006804 | none |
| e01 | SPU_006894 | Lv-Large |
| e01 | SPU_006950 | Lv-Lphn3 |
| e01 | SPU_007011 | Lv-Z346 |
| e01 | SPU_007025 | Lv-Hypp_1694 |
| e01 | SPU_007154 | Lv-Ube2c |
| e01 | SPU_007157 | none |
| e01 | SPU_007170 | none |
| e01 | SPU_007275 | Lv-CntbL |

| | | |
|---|---|---|
| e01 | SPU_007293 | Lv-Zcchc4 |
| e01 | SPU_007511 | Lv-Aqp_1 |
| e01 | SPU_007513 | Lv-SiaeL |
| e01 | SPU_007522 | Lv-Ube2e |
| e01 | SPU_007524 | Lv-C19orf51 |
| e01 | SPU_007611 | Lv-Rfx3 |
| e01 | SPU_007634 | Lv-St15l_1 |
| e01 | SPU_007667 | Lv-Rbm16/SfrS15L |
| e01 | SPU_007750 | Lv-Pomt1 |
| e01 | SPU_007754 | Lv-N4bp2 |
| e01 | SPU_007773 | Lv-Snf2 |
| e01 | SPU_007831 | none |
| e01 | SPU_007863 | Lv-Cbl |
| e01 | SPU_007914 | Lv-Glt8L |
| e01 | SPU_007972 | NA |
| e01 | SPU_007976 | NA |
| e01 | SPU_007977 | Lv-KatnaL2 |
| e01 | SPU_008002 | Lv-Rpgr |
| e01 | SPU_008072 | Lv-Pcgf3 |
| e01 | SPU_008079 | none |
| e01 | SPU_008205 | Lv-QrfprL |
| e01 | SPU_008216 | Lv-Ube2q1 |
| e01 | SPU_008234 | NA |
| e01 | SPU_008289 | none |
| e01 | SPU_008300 | Lv-NiempC1 |
| e01 | SPU_008317 | Lv-AsxL3 |
| e01 | SPU_008329 | none |
| e01 | SPU_008341 | Lv-Nt5c2L |
| e01 | SPU_008350 | Lv-Hypp_545 |
| e01 | SPU_008410 | Lv-Rab43 |
| e01 | SPU_008422 | NA |
| e01 | SPU_008527 | Lv-Sycp3L |
| e01 | SPU_008559 | Lv-Hypp_3068 |
| e01 | SPU_008563 | Lv-Z42 |
| e01 | SPU_008601 | Lv-Hypp_1798 |
| e01 | SPU_008609 | Lv-Nup214L |
| e01 | SPU_008640 | Lv-Asah1_1 |
| e01 | SPU_008698 | Lv-Shc1 |

| | | |
|---|---|---|
| e01 | SPU_008735 | none |
| e01 | SPU_008794 | Lv-Hypp_1804 |
| e01 | SPU_008852 | Lv-Mrps31L |
| e01 | SPU_008889 | Lv-Emi1 |
| e01 | SPU_008901 | none |
| e01 | SPU_009053 | Lv-Z316 |
| e01 | SPU_009063 | Lv-TpaseL_4 |
| e01 | SPU_009162 | Lv-Rtl_21 |
| e01 | SPU_009257 | Lv-Phldb2 |
| e01 | SPU_009261 | Lv-Hypp_1837 |
| e01 | SPU_009348 | Lv-Lin9 |
| e01 | SPU_009443 | Lv-Osm3 |
| e01 | SPU_009451 | Lv-SpataL |
| e01 | SPU_009532 | Lv-Nup50L |
| e01 | SPU_009556 | Lv-Lphn2L_1 |
| e01 | SPU_009590 | Lv-Rad51 |
| e01 | SPU_009623 | Lv-Mogat2 |
| e01 | SPU_009627 | Lv-Wdr6L |
| e01 | SPU_009650 | Lv-Ston2 |
| e01 | SPU_009654 | Lv-Fn3/Egf_6 |
| e01 | SPU_009672 | Lv-Glt_3 |
| e01 | SPU_009727 | Lv-Abt1 |
| e01 | SPU_009738 | Lv-Ak3l1 |
| e01 | SPU_009744 | Lv-Syt7L |
| e01 | SPU_009822 | Lv-Ehd3 |
| e01 | SPU_009906 | Lv-Xpo4 |
| e01 | SPU_009920 | Lv-Hypp_1880 |
| e01 | SPU_010095 | Lv-C11orf30L |
| e01 | SPU_010101 | Lv-Clect_22 |
| e01 | SPU_010170 | Lv-Hypp_609 |
| e01 | SPU_010180 | Lv-Tnfrsf_cl1 |
| e01 | SPU_010248 | Lv-Map2k7 |
| e01 | SPU_010268 | Lv-Ttll4_2 |
| e01 | SPU_010294 | Lv-Spata5 |
| e01 | SPU_010331 | Lv-Sema |
| e01 | SPU_010338 | Lv-Ddx43 |
| e01 | SPU_010340 | none |
| e01 | SPU_010346 | Lv-Depdc1 |

| | | |
|---|---|---|
| e01 | SPU_010375 | Lv-Os9l |
| e01 | SPU_010453 | none |
| e01 | SPU_010505 | Lv-RasefL |
| e01 | SPU_010515 | NA |
| e01 | SPU_010556 | Lv-RecQ5 |
| e01 | SPU_010576 | Lv-Cyp2L24 |
| e01 | SPU_010627 | Lv-Aof1 |
| e01 | SPU_010652 | NA |
| e01 | SPU_010672 | Lv-Hsp8 |
| e01 | SPU_010684 | Lv-Hypp_68 |
| e01 | SPU_010714 | none |
| e01 | SPU_010922 | Lv-Z458 |
| e01 | SPU_010924 | Lv-Z416 |
| e01 | SPU_010973 | none |
| e01 | SPU_010979 | Lv-Srrm2 |
| e01 | SPU_011124 | Lv-P70rsk |
| e01 | SPU_011175 | Lv-Yaf2l |
| e01 | SPU_011189 | Lv-Z141 |
| e01 | SPU_011196 | Lv-Wars2 |
| e01 | SPU_011234 | Lv-Slc35d1 |
| e01 | SPU_011287 | Lv-Ccdc55 |
| e01 | SPU_011442 | Lv-Ncbp1 |
| e01 | SPU_011499 | Lv-Plcb4_2 |
| e01 | SPU_011524 | Lv-Usp38_1 |
| e01 | SPU_011575 | Lv-Casd1 |
| e01 | SPU_011651 | Lv-Haus4 |
| e01 | SPU_011787 | Lv-Smchd_2 |
| e01 | SPU_011808 | Lv-Unk_58 |
| e01 | SPU_011816 | Lv-Ripk1-4B |
| e01 | SPU_012111 | Lv-Z53 |
| e01 | SPU_012163 | Lv-Ttll7 |
| e01 | SPU_012173 | Lv-Lycat_1 |
| e01 | SPU_012201 | Lv-Mosc2_1 |
| e01 | SPU_012278 | Lv-Btk |
| e01 | SPU_012291 | Lv-Wd/sof1 |
| e01 | SPU_012300 | Lv-Jmjd2c |
| e01 | SPU_012394 | Lv-Mutyh |
| e01 | SPU_012409 | none |

| | | |
|---|---|---|
| e01 | SPU_012470 | Lv-AdarB1 |
| e01 | SPU_012546 | Lv-Ctcf |
| e01 | SPU_012590 | Lv-Pcsk5 |
| e01 | SPU_012621 | Lv-Tatabp |
| e01 | SPU_012622 | Lv-Kiaa0831 |
| e01 | SPU_012633 | none |
| e01 | SPU_012643 | none |
| e01 | SPU_012662 | Lv-C14orf169L |
| e01 | SPU_012685 | Lv-Sppl3 |
| e01 | SPU_012736 | Lv-C21orf66L |
| e01 | SPU_012745 | Lv-St6galnac5_1 |
| e01 | SPU_012779 | Lv-Rnase3l |
| e01 | SPU_012780 | Lv-Slc6a5_5 |
| e01 | SPU_012917 | NA |
| e01 | SPU_012955 | Lv-Z422 |
| e01 | SPU_012963 | none |
| e01 | SPU_012964 | Lv-TtlL2 |
| e01 | SPU_012999 | none |
| e01 | SPU_013070 | Lv-Oma1L |
| e01 | SPU_013091 | Lv-HH3cid |
| e01 | SPU_013139 | Lv-Brd1 |
| e01 | SPU_013184 | Lv-Ttrspn_11 |
| e01 | SPU_013193 | Lv-Hypp_706 |
| e01 | SPU_013251 | none |
| e01 | SPU_013274 | Lv-Hypp_710 |
| e01 | SPU_013282 | Lv-Ikbkap |
| e01 | SPU_013477 | Lv-Imp_2 |
| e01 | SPU_013536 | Lv-Api |
| e01 | SPU_013544 | Lv-Snap29 |
| e01 | SPU_013569 | Lv-Lmo2t |
| e01 | SPU_013692 | NA |
| e01 | SPU_013700 | Lv-Otud7b |
| e01 | SPU_013724 | Lv-Ccdc90a |
| e01 | SPU_013729 | Lv-KifC3L1 |
| e01 | SPU_013753 | NA |
| e01 | SPU_014036 | Lv-Nbc |
| e01 | SPU_014062 | Lv-Dgat1 |
| e01 | SPU_014127 | Lv-DtL_2 |

| | | |
|---|---|---|
| e01 | SPU_014129 | Lv-Fam69a |
| e01 | SPU_014243 | Lv-Stx5 |
| e01 | SPU_014262 | Lv-Endrvt |
| e01 | SPU_014520 | none |
| e01 | SPU_014559 | Lv-Vw3b |
| e01 | SPU_014683 | Lv-Z492 |
| e01 | SPU_014690 | Lv-C19orf47L |
| e01 | SPU_014693 | Lv-Ero1L |
| e01 | SPU_014707 | Lv-Setdb1 |
| e01 | SPU_014710 | Lv-Adssl1 |
| e01 | SPU_014817 | Lv-Z423 |
| e01 | SPU_014907 | Lv-Dcdsh_2 |
| e01 | SPU_014936 | Lv-Nim1 |
| e01 | SPU_014956 | none |
| e01 | SPU_014957 | Lv-Epb4.1L5 |
| e01 | SPU_015009 | Lv-Notchh1_10 |
| e01 | SPU_015032 | Lv-Endrvt89 |
| e01 | SPU_015062 | Lv-Phyhd1_2 |
| e01 | SPU_015088 | Lv-Pdpi(cyclophilin)L6 |
| e01 | SPU_015125 | Lv-Gal3st1_6 |
| e01 | SPU_015530 | Lv-Ctns |
| e01 | SPU_015646 | NA |
| e01 | SPU_015688 | Lv-Z154 |
| e01 | SPU_015700 | Lv-C1qL_8 |
| e01 | SPU_015721 | Lv-Hypp_3056 |
| e01 | SPU_015771 | Lv-Serrp8 |
| e01 | SPU_015885 | Lv-Rnf139 |
| e01 | SPU_016049 | Lv-Bap1 |
| e01 | SPU_016058 | Lv-Rbm15L |
| e01 | SPU_016066 | Lv-Parp6 |
| e01 | SPU_016081 | Lv-Pofut1_1 |
| e01 | SPU_016088 | Lv-Clect/Tm/Duf |
| e01 | SPU_016111 | Lv-Ugt1L_5 |
| e01 | SPU_016127 | Lv-Chrna7_5 |
| e01 | SPU_016214 | Lv-V-crk |
| e01 | SPU_016227 | Lv-Pde4dipl |
| e01 | SPU_016250 | Lv-Z70 |
| e01 | SPU_016326 | Lv-Pro_ser_thr_ppip1_1 |

| | | |
|---|---|---|
| e01 | SPU_016337 | NA |
| e01 | SPU_016350 | Lv-Ptdss2 |
| e01 | SPU_016423 | Lv-Giltl |
| e01 | SPU_016465 | Lv-Hypp_2262 |
| e01 | SPU_016544 | Lv-Hypp_830 |
| e01 | SPU_016667 | Lv-Ccdc130 |
| e01 | SPU_016687 | Lv-Nfyc |
| e01 | SPU_016688 | Lv-Hm13 |
| e01 | SPU_016712 | Lv-Fam76a |
| e01 | SPU_016840 | Lv-Egf/Zp_1 |
| e01 | SPU_016899 | Lv-Rnf32 |
| e01 | SPU_016903 | Lv-Cxxc1 |
| e01 | SPU_016904 | NA |
| e01 | SPU_016946 | none |
| e01 | SPU_016960 | Lv-Hypp_2292 |
| e01 | SPU_016962 | Lv-Hypp_2293 |
| e01 | SPU_016969 | Lv-Hypp_2295 |
| e01 | SPU_017160 | none |
| e01 | SPU_017198 | Lv-Tpp2 |
| e01 | SPU_017259 | Lv-Dnajhc10 |
| e01 | SPU_017280 | Lv-Fuca2 |
| e01 | SPU_017281 | Lv-Fuca2_4 |
| e01 | SPU_017308 | Lv-Ahi1 |
| e01 | SPU_017310 | Lv-Smchd1_3 |
| e01 | SPU_017358 | Lv-Zcchc7L |
| e01 | SPU_017389 | Lv-Chrna6_2 |
| e01 | SPU_017407 | Lv-Clock |
| e01 | SPU_017441 | Lv-Z65 |
| e01 | SPU_017756 | none |
| e01 | SPU_017774 | Lv-Hypp_2354 |
| e01 | SPU_017811 | Lv-Z401 |
| e01 | SPU_017832 | Lv-Arl7 |
| e01 | SPU_017837 | Lv-Nkx3.2 |
| e01 | SPU_017842 | Lv-Clect/VWF_type_C |
| e01 | SPU_017946 | Lv-Vps13bL |
| e01 | SPU_017955 | Lv-Depdc1b |
| e01 | SPU_018025 | Lv-Hypp_905 |
| e01 | SPU_018033 | Lv-Tsen54L_1 |

| | | |
|---|---|---|
| e01 | SPU_018072 | Lv-Ankmy2 |
| e01 | SPU_018106 | none |
| e01 | SPU_018199 | Lv-CycK |
| e01 | SPU_018205 | Lv-Lrwd1 |
| e01 | SPU_018206 | Lv-Hypp_2375 |
| e01 | SPU_018234 | Lv-Nsun2 |
| e01 | SPU_018261 | NA |
| e01 | SPU_018291 | Lv-Fbox/RhoGEF |
| e01 | SPU_018293 | Lv-Rt12 |
| e01 | SPU_018299 | none |
| e01 | SPU_018324 | none |
| e01 | SPU_018375 | Lv-Nek10 |
| e01 | SPU_018524 | Lv-APH |
| e01 | SPU_018529 | none |
| e01 | SPU_018649 | Lv-Plekhf2 |
| e01 | SPU_018673 | Lv-Z17 |
| e01 | SPU_018796 | Lv-Hypp_144 |
| e01 | SPU_018891 | none |
| e01 | SPU_018893 | Lv-Cdo1 |
| e01 | SPU_018960 | Lv-Z317 |
| e01 | SPU_018995 | none |
| e01 | SPU_019165 | Lv-Krp110 |
| e01 | SPU_019239 | Lv-Sr/Lrr/Gpcr |
| e01 | SPU_019282 | Lv-Cep70L |
| e01 | SPU_019287 | Lv-Usp8 |
| e01 | SPU_019306 | Lv-Bin3 |
| e01 | SPU_019315 | Lv-Hypp_935 |
| e01 | SPU_019339 | Lv-Kpna3 |
| e01 | SPU_019385 | Lv-Btg4 |
| e01 | SPU_019391 | Lv-Skp2 |
| e01 | SPU_019406 | Lv-Kiaa1147 |
| e01 | SPU_019515 | Lv-NotchL_16 |
| e01 | SPU_019559 | Lv-Pth2 |
| e01 | SPU_019610 | Lv-Yme1L1 |
| e01 | SPU_019743 | Lv-Speract |
| e01 | SPU_019775 | Lv-Kif12L |
| e01 | SPU_019885 | Lv-Slc12a4_1 |
| e01 | SPU_019893 | NA |

| | | |
|---|---|---|
| e01 | SPU_019900 | Lv-Dyrk1Ai2 |
| e01 | SPU_019960 | Lv-Bud13 |
| e01 | SPU_020017 | NA |
| e01 | SPU_020027 | none |
| e01 | SPU_020089 | Lv-Agpat3_1 |
| e01 | SPU_020147 | Lv-Hypp_2491 |
| e01 | SPU_020167 | NA |
| e01 | SPU_020168 | Lv-Nrbp |
| e01 | SPU_020226 | Lv-Hypp_2497 |
| e01 | SPU_020348 | Lv-Chrm5L_3 |
| e01 | SPU_020364 | Lv-Kiaa0430L |
| e01 | SPU_020409 | Lv-Tmem181 |
| e01 | SPU_020410 | Lv-Zc3h12bL |
| e01 | SPU_020529 | Lv-Rassf8 |
| e01 | SPU_020540 | Lv-Dbxd7 |
| e01 | SPU_020696 | Lv-Tprg1L |
| e01 | SPU_020722 | Lv-Smad1/5/8_1 |
| e01 | SPU_020726 | Lv-Z472 |
| e01 | SPU_020787 | Lv-Hypp_994 |
| e01 | SPU_020793 | Lv-Snx21 |
| e01 | SPU_020900 | Lv-Tmem20_2 |
| e01 | SPU_020909 | Lv-Plxnb1_2 |
| e01 | SPU_020945 | Lv-Foxred2 |
| e01 | SPU_020958 | Lv-Hypp_1006 |
| e01 | SPU_020967 | Lv-Hypp_1007 |
| e01 | SPU_020990 | Lv-Hypp_1008 |
| e01 | SPU_021084 | Lv-Fnip2 |
| e01 | SPU_021085 | Lv-Stoml1 |
| e01 | SPU_021095 | none |
| e01 | SPU_021105 | Lv-Nags |
| e01 | SPU_021128 | none |
| e01 | SPU_021164 | none |
| e01 | SPU_021177 | none |
| e01 | SPU_021333 | NA |
| e01 | SPU_021339 | Lv-Hypp_2546 |
| e01 | SPU_021391 | Lv-ArsaL |
| e01 | SPU_021413 | Lv-Hypp_1023 |
| e01 | SPU_021469 | Lv-Ids_7 |

| | | |
|---|---|---|
| e01 | SPU_021559 | Lv-Man2a2 |
| e01 | SPU_021713 | Lv-Gtpbp1_1 |
| e01 | SPU_021728 | Lv-Hypp_175 |
| e01 | SPU_021778 | Lv-Cir |
| e01 | SPU_021833 | Lv-Zdhhc17_1 |
| e01 | SPU_021851 | Lv-Z434 |
| e01 | SPU_021982 | Lv-Mfsd11_1 |
| e01 | SPU_022015 | NA |
| e01 | SPU_022058 | Lv-MccL |
| e01 | SPU_022069 | Lv-Brpf1 |
| e01 | SPU_022082 | Lv-Hypp_2597 |
| e01 | SPU_022103 | Lv-Glcci1L |
| e01 | SPU_022154 | Lv-Rab22l3 |
| e01 | SPU_022171 | Lv-Itfg2 |
| e01 | SPU_022205 | Lv-Hypp_2604 |
| e01 | SPU_022221 | Lv-Rad51c |
| e01 | SPU_022309 | Lv-Polr2a |
| e01 | SPU_022311 | Lv-C12orf51L_1 |
| e01 | SPU_022713 | Lv-Lpcat3_2 |
| e01 | SPU_022770 | Lv-Slu7 |
| e01 | SPU_022858 | none |
| e01 | SPU_022992 | Lv-Nup98_1 |
| e01 | SPU_023020 | Lv-Tep1L_2 |
| e01 | SPU_023058 | Lv-Golt1B |
| e01 | SPU_023075 | Lv-EhH4_65 |
| e01 | SPU_023100 | none |
| e01 | SPU_023107 | NA |
| e01 | SPU_023123 | Lv-Glul |
| e01 | SPU_023151 | none |
| e01 | SPU_023201 | Lv-Fbxo16 |
| e01 | SPU_023262 | Lv-Arfgap1 |
| e01 | SPU_023370 | Lv-Frmpd4 |
| e01 | SPU_023439 | Lv-Harbi1_2 |
| e01 | SPU_023470 | Lv-C9orf117_1 |
| e01 | SPU_023509 | none |
| e01 | SPU_023511 | Lv-Gusb |
| e01 | SPU_023526 | none |
| e01 | SPU_023530 | Lv-SwiL |

| | | |
|---|---|---|
| e01 | SPU_023720 | Lv-Ncaph |
| e01 | SPU_023904 | none |
| e01 | SPU_023905 | Lv-Fringe |
| e01 | SPU_023991 | Lv-Srcr201 |
| e01 | SPU_024028 | Lv-Asl_1 |
| e01 | SPU_024109 | Lv-CerklL |
| e01 | SPU_024114 | Lv-Mad2l1bp |
| e01 | SPU_024126 | Lv-PolypL_122 |
| e01 | SPU_024165 | none |
| e01 | SPU_024235 | Lv-Brap |
| e01 | SPU_024291 | Lv-Hypp_3030 |
| e01 | SPU_024397 | Lv-Sf4 |
| e01 | SPU_024407 | Lv-Ankle1 |
| e01 | SPU_024511 | Lv-Hypp_2738 |
| e01 | SPU_024581 | NA |
| e01 | SPU_024613 | Lv-LsampL_2 |
| e01 | SPU_024623 | Lv-Dhx38 |
| e01 | SPU_024635 | Lv-Sac3 |
| e01 | SPU_024722 | Lv-Tada2b |
| e01 | SPU_024776 | Lv-Hypp_1126 |
| e01 | SPU_024927 | Lv-Med17 |
| e01 | SPU_025030 | Lv-Grid2ipL |
| e01 | SPU_025032 | Lv-C1orf55 |
| e01 | SPU_025062 | Lv-Muc4dL3 |
| e01 | SPU_025084 | Lv-Hypp_2769 |
| e01 | SPU_025137 | Lv-Kiaa1310L |
| e01 | SPU_025157 | Lv-Egfib_2 |
| e01 | SPU_025182 | Lv-Ppp2ca |
| e01 | SPU_025202 | none |
| e01 | SPU_025241 | Lv-Ck1a2 |
| e01 | SPU_025286 | Lv-Zc3h7b_2 |
| e01 | SPU_025306 | NA |
| e01 | SPU_025356 | Lv-Stim1 |
| e01 | SPU_025421 | NA |
| e01 | SPU_025475 | Lv-Akirin2 |
| e01 | SPU_025478 | Lv-March3 |
| e01 | SPU_025483 | Lv-Stard3 |
| e01 | SPU_025487 | Lv-Bin1 |

| | | |
|---|---|---|
| e01 | SPU_025502 | Lv-Neog/DccL |
| e01 | SPU_025536 | Lv-GagpolL_91 |
| e01 | SPU_025589 | Lv-Nfya_1 |
| e01 | SPU_025629 | Lv-Twister |
| e01 | SPU_025644 | Lv-Gltpd1 |
| e01 | SPU_025645 | none |
| e01 | SPU_025679 | Lv-Neh1 |
| e01 | SPU_025691 | Lv-Zfyve28 |
| e01 | SPU_025713 | Lv-DnmbpL |
| e01 | SPU_025819 | Lv-U1snrnpbp_1 |
| e01 | SPU_025856 | Lv-Smurf |
| e01 | SPU_025927 | none |
| e01 | SPU_025949 | Lv-Pigo |
| e01 | SPU_026011 | Lv-Wdr49 |
| e01 | SPU_026034 | Lv-Pex14 |
| e01 | SPU_026260 | Lv-Gal3st1_12 |
| e01 | SPU_026324 | Lv-Hypp_2835 |
| e01 | SPU_026332 | Lv-Prpf38b |
| e01 | SPU_026404 | Lv-Sec31a |
| e01 | SPU_026427 | Lv-Top1 |
| e01 | SPU_026496 | Lv-Socs4/5 |
| e01 | SPU_026535 | Lv-Ube4B |
| e01 | SPU_026611 | Lv-Thap4 |
| e01 | SPU_026620 | Lv-Sip1 |
| e01 | SPU_026666 | none |
| e01 | SPU_026716 | Lv-Lonp2 |
| e01 | SPU_026758 | Lv-Hif2a |
| e01 | SPU_026835 | Lv-Fuca2_2 |
| e01 | SPU_026839 | Lv-Gys1 |
| e01 | SPU_026865 | Lv-Hypp_1189 |
| e01 | SPU_027124 | Lv-Unc44_352 |
| e01 | SPU_027189 | Lv-Hypp_2893 |
| e01 | SPU_027234 | Lv-Bindin_1 |
| e01 | SPU_027245 | Lv-Ulk1 |
| e01 | SPU_027388 | Lv-Nkcc1 |
| e01 | SPU_027389 | Lv-Lama5Lf |
| e01 | SPU_027420 | Lv-Dph5 |
| e01 | SPU_027426 | Lv-Tbl2 |

| | | |
|---|---|---|
| e01 | SPU_027432 | Lv-Wdr68 |
| e01 | SPU_027446 | Lv-Enz2 |
| e01 | SPU_027520 | Lv-Ihpk2_1 |
| e01 | SPU_027529 | Lv-Hypp_3122 |
| e01 | SPU_027625 | Lv-LrpL_8 |
| e01 | SPU_027669 | Lv-P4ha1_1 |
| e01 | SPU_027670 | none |
| e01 | SPU_027726 | Lv-Pex3_1 |
| e01 | SPU_027811 | none |
| e01 | SPU_027834 | none |
| e01 | SPU_027844 | Lv-Madl1 |
| e01 | SPU_027935 | Lv-Bmal |
| e01 | SPU_027998 | Lv-Rfxdc2L |
| e01 | SPU_028029 | Lv-Hypp_279 |
| e01 | SPU_028066 | Lv-Alk4/5/7 |
| e01 | SPU_028137 | Lv-F8a1L |
| e01 | SPU_028249 | Lv-Akap13 |
| e01 | SPU_028368 | Lv-Lamb1L_2 |
| e01 | SPU_028375 | Lv-Golph3 |
| e01 | SPU_028429 | none |
| e01 | SPU_028439 | Lv-Hypp_284 |
| e01 | SPU_028476 | Lv-Dhx8 |
| e01 | SPU_028503 | Lv-PnplL_1 |
| e01 | SPU_028526 | Lv-Lypla3 |
| e01 | SPU_028531 | Lv-Ppp2r3c |
| e01 | SPU_028622 | NA |
| e01 | SPU_028680 | Lv-Srcr227 |
| e01 | SPU_028715 | Lv-Kif23 |
| e01 | SPU_028769 | none |
| e01 | SPU_028852 | none |
| e01 | SPU_028854 | none |
| e01 | SPU_028855 | Lv-Mphosph10 |
| e01 | SPU_028867 | Lv-C12orf48L |
| e01 | SPU_028885 | Lv-Stx16_1 |
| e01 | SPU_028891 | none |
| e01 | SPU_028915 | Lv-Hypp_2974 |
| e01 | SPU_028916 | Lv-Hypp_1991 |
| e01 | SPU_028932 | Lv-EhH4_82 |

| | | |
|---|---|---|
| e01 | SPU_030082 | Lv-Gg2 |
| e02 | SPU_000059 | Lv-Ddx55L |
| e02 | SPU_000189 | Lv-C3orf31L |
| e02 | SPU_000191 | Lv-Cmpk |
| e02 | SPU_000193 | Lv-Stip |
| e02 | SPU_000261 | Lv-Hypp_1247 |
| e02 | SPU_000283 | Lv-Tho1 |
| e02 | SPU_000305 | Lv-Ankzf1 |
| e02 | SPU_000318 | Lv-Prhoxnb |
| e02 | SPU_000342 | Lv-Utp20h |
| e02 | SPU_000356 | Lv-Slc16a6L |
| e02 | SPU_000357 | Lv-Gnmt |
| e02 | SPU_000450 | Lv-Kctd1 |
| e02 | SPU_000654 | Lv-Srcr5 |
| e02 | SPU_000776 | Lv-Ptpn9 |
| e02 | SPU_000832 | Lv-MyV |
| e02 | SPU_000848 | none |
| e02 | SPU_000925 | Lv-Fn3_14 |
| e02 | SPU_001012 | Lv-Trabd |
| e02 | SPU_001061 | Lv-Utp18 |
| e02 | SPU_001164 | Lv-Bxdc5 |
| e02 | SPU_001291 | Lv-Glmn |
| e02 | SPU_001304 | Lv-Xrcc1L |
| e02 | SPU_001341 | Lv-Wdr45L |
| e02 | SPU_001346 | Lv-Hypp_329 |
| e02 | SPU_001479 | Lv-Tusc4 |
| e02 | SPU_001559 | Lv-Cutc |
| e02 | SPU_001648 | Lv-Z355 |
| e02 | SPU_001686 | Lv-Aars_1 |
| e02 | SPU_001811 | NA |
| e02 | SPU_001814 | NA |
| e02 | SPU_001907 | Lv-Fn3/Egff_7 |
| e02 | SPU_002162 | none |
| e02 | SPU_002225 | Lv-Fem1hc |
| e02 | SPU_002352 | none |
| e02 | SPU_002387 | Lv-Gbf1L |
| e02 | SPU_002465 | Lv-Atp13a3_2 |
| e02 | SPU_002466 | none |

| | | |
|---|---|---|
| e02 | SPU_002483 | Lv-Fam114a2 |
| e02 | SPU_002522 | Lv-B3gnt6 |
| e02 | SPU_002546 | Lv-Coq2 |
| e02 | SPU_002559 | Lv-Phf3 |
| e02 | SPU_002623 | Lv-Hypp_3104 |
| e02 | SPU_002643 | Lv-Renbp |
| e02 | SPU_002774 | Lv-Mina |
| e02 | SPU_003049 | Lv-Hcfc2_1 |
| e02 | SPU_003085 | Lv-Smp3 |
| e02 | SPU_003091 | Lv-PolypL_933 |
| e02 | SPU_003093 | Lv-Muc4 |
| e02 | SPU_003255 | Lv-Galnac4s-6stL_3 |
| e02 | SPU_003292 | Lv-Dap3 |
| e02 | SPU_003307 | none |
| e02 | SPU_003365 | Lv-Hrh2L |
| e02 | SPU_003540 | Lv-Prom1 |
| e02 | SPU_003632 | Lv-Kynu |
| e02 | SPU_003751 | NA |
| e02 | SPU_003948 | Lv-Hypp_1488 |
| e02 | SPU_004011 | Lv-Srcr33 |
| e02 | SPU_004025 | Lv-Slc5a7_1 |
| e02 | SPU_004048 | Lv-Vps36 |
| e02 | SPU_004192 | NA |
| e02 | SPU_004216 | Lv-Aga |
| e02 | SPU_004236 | NA |
| e02 | SPU_004286 | Lv-Lman1 |
| e02 | SPU_004298 | Lv-elof1h |
| e02 | SPU_004434 | Lv-Card11 |
| e02 | SPU_004443 | Lv-Adck2 |
| e02 | SPU_004520 | Lv-C20orf72L |
| e02 | SPU_004648 | Lv-Lpll |
| e02 | SPU_004669 | Lv-Gosr2 |
| e02 | SPU_004862 | Lv-Ccdc9 |
| e02 | SPU_004893 | Lv-Aars |
| e02 | SPU_004912 | Lv-Ddx47 |
| e02 | SPU_004944 | Lv-Mvk |
| e02 | SPU_005066 | Lv-Unc44L_40 |
| e02 | SPU_005123 | Lv-Scfd1 |

| | | |
|---|---|---|
| e02 | SPU_005179 | Lv-Hypp_1569 |
| e02 | SPU_005206 | Lv-Notch2L |
| e02 | SPU_005218 | Lv-Oxa1L |
| e02 | SPU_005428 | Lv-Kptn |
| e02 | SPU_005505 | Lv-Gpr52L_2 |
| e02 | SPU_005696 | Lv-PppL_183 |
| e02 | SPU_005790 | Lv-Tmem104 |
| e02 | SPU_005808 | NA |
| e02 | SPU_006044 | Lv-Surf4 |
| e02 | SPU_006176 | NA |
| e02 | SPU_006212 | Lv-Ste24pL_1 |
| e02 | SPU_006276 | Lv-Gpcr126a1 |
| e02 | SPU_006325 | Lv-Dem1 |
| e02 | SPU_006482 | Lv-Tpase6 |
| e02 | SPU_006580 | Lv-Tial1 |
| e02 | SPU_006634 | none |
| e02 | SPU_006687 | Lv-Hypp_1674 |
| e02 | SPU_006766 | none |
| e02 | SPU_006767 | Lv-Sar1 |
| e02 | SPU_006798 | Lv-Hypp_1683 |
| e02 | SPU_006853 | Lv-Cars |
| e02 | SPU_007130 | Lv-Hypp_506 |
| e02 | SPU_007318 | none |
| e02 | SPU_007349 | Lv-Srcr39 |
| e02 | SPU_007386 | Lv-Ddx54 |
| e02 | SPU_007442 | Lv-Hypp_1721 |
| e02 | SPU_007458 | Lv-Hypp_516 |
| e02 | SPU_007733 | Lv-ScyL1 |
| e02 | SPU_007745 | Lv-Cahb |
| e02 | SPU_007795 | Lv-Surreal/Gpcr84 |
| e02 | SPU_007828 | Lv-Fam160b1L |
| e02 | SPU_007840 | Lv-Srcr45 |
| e02 | SPU_008054 | Lv-Sec63h |
| e02 | SPU_008058 | Lv-VarsiA |
| e02 | SPU_008087 | Lv-Hypp_1764 |
| e02 | SPU_008088 | Lv-Hypp_1765 |
| e02 | SPU_008178 | none |
| e02 | SPU_008214 | Lv-Smc5 |

| | | |
|---|---|---|
| e02 | SPU_008219 | Lv-Pomgnt1 |
| e02 | SPU_008400 | NA |
| e02 | SPU_008465 | Lv-Dohh |
| e02 | SPU_008670 | Lv-Ptprr |
| e02 | SPU_008680 | none |
| e02 | SPU_008786 | Lv-Dhx29 |
| e02 | SPU_008797 | Lv-Hypp_1805 |
| e02 | SPU_008886 | Lv-Ubxd2 |
| e02 | SPU_008956 | NA |
| e02 | SPU_009000 | none |
| e02 | SPU_009021 | NA |
| e02 | SPU_009036 | Lv-Lrrka |
| e02 | SPU_009067 | Lv-Slc25a27_1 |
| e02 | SPU_009109 | Lv-Znrd1 |
| e02 | SPU_009437 | Lv-Ctps |
| e02 | SPU_009477 | Lv-Hsp701C |
| e02 | SPU_009571 | Lv-Fn3/Igf_35 |
| e02 | SPU_009678 | Lv-Btbd8 |
| e02 | SPU_009711 | Lv-Ncdn |
| e02 | SPU_009726 | Lv-Abhd13 |
| e02 | SPU_009742 | Lv-Rbm21L |
| e02 | SPU_009747 | none |
| e02 | SPU_009778 | Lv-Tbl3_1 |
| e02 | SPU_009983 | Lv-Hypp_3067 |
| e02 | SPU_010013 | Lv-Wdr74 |
| e02 | SPU_010061 | Lv-Hypp_1896 |
| e02 | SPU_010088 | Lv-Hypp_1897 |
| e02 | SPU_010119 | Lv-Nol9 |
| e02 | SPU_010226 | Lv-Srcr84 |
| e02 | SPU_010281 | Lv-Armc7 |
| e02 | SPU_010304 | Lv-Psme4 |
| e02 | SPU_010323 | Lv-Mars |
| e02 | SPU_010580 | Lv-Ube1 |
| e02 | SPU_010743 | Lv-Eif2Bd |
| e02 | SPU_010773 | Lv-Dhx57L |
| e02 | SPU_010881 | none |
| e02 | SPU_011199 | Lv-Pola2 |
| e02 | SPU_011253 | Lv-Snx30L |

| | | |
|---|---|---|
| e02 | SPU_011395 | Lv-Hsp701G |
| e02 | SPU_011416 | Lv-Ugt8_1 |
| e02 | SPU_011577 | Lv-Ap2b1_1 |
| e02 | SPU_011615 | Lv-Samd9lL |
| e02 | SPU_011643 | Lv-PppL_259 |
| e02 | SPU_011774 | none |
| e02 | SPU_011954 | Lv-Rb |
| e02 | SPU_012087 | Lv-Cebpz |
| e02 | SPU_012090 | none |
| e02 | SPU_012096 | Lv-Ecsit |
| e02 | SPU_012152 | Lv-C19orf61 |
| e02 | SPU_012239 | Lv-Cern |
| e02 | SPU_012324 | NA |
| e02 | SPU_012858 | Lv-Myto/Isoleu/Synth |
| e02 | SPU_012938 | Lv-Hypp_697 |
| e02 | SPU_012941 | NA |
| e02 | SPU_013036 | NA |
| e02 | SPU_013042 | Lv-Cog5 |
| e02 | SPU_013062 | Lv-Hypp_701 |
| e02 | SPU_013090 | Lv-Ttrap |
| e02 | SPU_013140 | Lv-Ron |
| e02 | SPU_013347 | Lv-Hypp_3059 |
| e02 | SPU_013376 | Lv-Txndc |
| e02 | SPU_013406 | Lv-Z118 |
| e02 | SPU_013435 | NA |
| e02 | SPU_013486 | none |
| e02 | SPU_013667 | Lv-Fn3f_40 |
| e02 | SPU_013735 | Lv-Slc3a1 |
| e02 | SPU_013749 | none |
| e02 | SPU_013810 | Lv-Ptpn6/11 |
| e02 | SPU_013815 | Lv-Abcf3 |
| e02 | SPU_013859 | Lv-GtpbpL2(nucleolar) |
| e02 | SPU_013963 | Lv-C2orf44L |
| e02 | SPU_013966 | Lv-Cacna2d2 |
| e02 | SPU_014063 | Lv-Prmt7 |
| e02 | SPU_014197 | Lv-Z55 |
| e02 | SPU_014250 | Lv-Mat1 |
| e02 | SPU_014258 | Lv-Sned1_3 |

| | | |
|---|---|---|
| e02 | SPU_014304 | Lv-Chst3L_4 |
| e02 | SPU_014361 | Lv-Prpf4 |
| e02 | SPU_014424 | Lv-Utp20_4 |
| e02 | SPU_014426 | Lv-Hypp_101 |
| e02 | SPU_014504 | Lv-Gluld1_6 |
| e02 | SPU_014589 | Lv-Plxnb1L |
| e02 | SPU_014625 | Lv-Ppm1gL |
| e02 | SPU_014650 | Lv-Gabra1 |
| e02 | SPU_014714 | NA |
| e02 | SPU_014752 | Lv-Nck |
| e02 | SPU_014806 | Lv-Yif1B |
| e02 | SPU_014899 | none |
| e02 | SPU_014950 | none |
| e02 | SPU_015221 | Lv-Pdcl3 |
| e02 | SPU_015247 | Lv-MmsL19 |
| e02 | SPU_015304 | Lv-Tsnl |
| e02 | SPU_015319 | none |
| e02 | SPU_015472 | Lv-Alg3 |
| e02 | SPU_015562 | Lv-PolypL_949 |
| e02 | SPU_015649 | Lv-Hypp_2223 |
| e02 | SPU_015827 | none |
| e02 | SPU_015850 | Lv-Cugbp1 |
| e02 | SPU_015901 | none |
| e02 | SPU_015923 | Lv-Ptpase_1 |
| e02 | SPU_015985 | Lv-Fam82a |
| e02 | SPU_016002 | Lv-Abcc9 |
| e02 | SPU_016041 | Lv-EgfiL_94 |
| e02 | SPU_016090 | Lv-Slc35e3 |
| e02 | SPU_016327 | Lv-DppdL_2 |
| e02 | SPU_016343 | Lv-E12 |
| e02 | SPU_016404 | Lv-Metaxin1 |
| e02 | SPU_016446 | Lv-C20orf59L |
| e02 | SPU_016484 | Lv-Hypp_825 |
| e02 | SPU_016575 | none |
| e02 | SPU_016616 | Lv-Unc44L_115 |
| e02 | SPU_016634 | Lv-Dap3_1 |
| e02 | SPU_016645 | Lv-Heatr6-3 |
| e02 | SPU_016663 | none |

| | | |
|---|---|---|
| e02 | SPU_016678 | Lv-Egfi_4 |
| e02 | SPU_016810 | Lv-Nlr138_1 |
| e02 | SPU_017162 | none |
| e02 | SPU_017202 | Lv-Ig/Egf/Vwc |
| e02 | SPU_017255 | NA |
| e02 | SPU_017438 | Lv-Srrd |
| e02 | SPU_017578 | Lv-Ghdc |
| e02 | SPU_017617 | none |
| e02 | SPU_017633 | Lv-Nudt18 |
| e02 | SPU_017899 | Lv-Hs3st |
| e02 | SPU_018153 | Lv-Fn3/Egf_3 |
| e02 | SPU_018210 | Lv-RcL1 |
| e02 | SPU_018335 | Lv-Iswi |
| e02 | SPU_018394 | Lv-Osgepl1L |
| e02 | SPU_018481 | Lv-Exosc7 |
| e02 | SPU_018505 | Lv-GbA3 |
| e02 | SPU_018556 | Lv-Pex1 |
| e02 | SPU_018685 | none |
| e02 | SPU_018748 | Lv-Flad1 |
| e02 | SPU_018758 | Lv-Pmpca |
| e02 | SPU_018802 | Lv-Dlp |
| e02 | SPU_018922 | Lv-Srm |
| e02 | SPU_018996 | Lv-Nom1_1 |
| e02 | SPU_019096 | Lv-AdprhL2 |
| e02 | SPU_019123 | Lv-Ddx52_1 |
| e02 | SPU_019162 | Lv-Rrp8L |
| e02 | SPU_019164 | NA |
| e02 | SPU_019168 | Lv-Tgfbrap1_1 |
| e02 | SPU_019197 | Lv-Chaf1bL |
| e02 | SPU_019280 | none |
| e02 | SPU_019311 | Lv-Svep1L_6 |
| e02 | SPU_019396 | Lv-Cophs4 |
| e02 | SPU_019505 | Lv-Adpgk_1 |
| e02 | SPU_019560 | Lv-Hypp_2449 |
| e02 | SPU_019563 | Lv-Slc16a12_2 |
| e02 | SPU_019726 | none |
| e02 | SPU_019736 | Lv-Mrps2l_1 |
| e02 | SPU_019781 | Lv-B3galt1_11 |

| | | |
|---|---|---|
| e02 | SPU_019858 | Lv-Patl1 |
| e02 | SPU_019929 | Lv-Pdia6 |
| e02 | SPU_019996 | Lv-AAT-amino_acid_transporter |
| e02 | SPU_020011 | Lv-Pwp1 |
| e02 | SPU_020040 | Lv-N_wasp_bp |
| e02 | SPU_020136 | Lv-Tbrg4 |
| e02 | SPU_020241 | Lv-Hypp_2498 |
| e02 | SPU_020248 | Lv-C14orf21L |
| e02 | SPU_020269 | Lv-Appbp2_1 |
| e02 | SPU_020302 | Lv-Lias |
| e02 | SPU_020303 | Lv-Tmem144 |
| e02 | SPU_020310 | Lv-Psmc3ip |
| e02 | SPU_020402 | Lv-Prodh |
| e02 | SPU_020456 | Lv-Unc44_244 |
| e02 | SPU_020463 | Lv-Trnt1 |
| e02 | SPU_020466 | Lv-Hypp_974 |
| e02 | SPU_020484 | Lv-Hypp_167 |
| e02 | SPU_020750 | none |
| e02 | SPU_020796 | Lv-Sdccag3L |
| e02 | SPU_020886 | Lv-Gulo |
| e02 | SPU_020944 | Lv-SelpL |
| e02 | SPU_021167 | Lv-Pak1ip1 |
| e02 | SPU_021359 | Lv-B4galt5_2 |
| e02 | SPU_021412 | Lv-Bag1 |
| e02 | SPU_021525 | Lv-Nbn |
| e02 | SPU_021645 | Lv-NatL_1 |
| e02 | SPU_021656 | Lv-Khc |
| e02 | SPU_021721 | Lv-AT_DegT |
| e02 | SPU_021795 | Lv-Timm22 |
| e02 | SPU_021796 | NA |
| e02 | SPU_022010 | none |
| e02 | SPU_022040 | Lv-My7A |
| e02 | SPU_022088 | Lv-Prei4L |
| e02 | SPU_022116 | Lv-Colf_6 |
| e02 | SPU_022239 | Lv-Uck2 |
| e02 | SPU_022324 | Lv-Hypp_1043 |
| e02 | SPU_022478 | Lv-Slc25a26_3 |
| e02 | SPU_022479 | Lv-Pdpi(cyclophilin)L9 |

| | | |
|---|---|---|
| e02 | SPU_022502 | Lv-Nfu1 |
| e02 | SPU_022532 | none |
| e02 | SPU_022575 | Lv-Sh3bp2 |
| e02 | SPU_022650 | none |
| e02 | SPU_022675 | Lv-Gpatch4 |
| e02 | SPU_022980 | Lv-Ids_6 |
| e02 | SPU_023060 | Lv-Mamdc4_9 |
| e02 | SPU_023168 | Lv-Rxfp1L_5 |
| e02 | SPU_023190 | Lv-C10orf137L_2 |
| e02 | SPU_023194 | Lv-Endrvt45 |
| e02 | SPU_023217 | Lv-Ran |
| e02 | SPU_023272 | Lv-Slc16a6L_7 |
| e02 | SPU_023273 | Lv-Slc16a12L_17 |
| e02 | SPU_023285 | none |
| e02 | SPU_023302 | Lv-PppL_136 |
| e02 | SPU_023313 | Lv-Pi4ka |
| e02 | SPU_023412 | Lv-Ranbp2L_4 |
| e02 | SPU_023556 | Lv-Hypp_2676 |
| e02 | SPU_023572 | Lv-Hdac6 |
| e02 | SPU_023601 | Lv-Mpsk1 |
| e02 | SPU_023603 | NA |
| e02 | SPU_023796 | Lv-Mamdc4-11 |
| e02 | SPU_023812 | NA |
| e02 | SPU_023815 | Lv-Slsp_18 |
| e02 | SPU_023830 | none |
| e02 | SPU_023861 | Lv-Wdr77 |
| e02 | SPU_023997 | NA |
| e02 | SPU_024033 | Lv-C1qL_6 |
| e02 | SPU_024203 | none |
| e02 | SPU_024464 | NA |
| e02 | SPU_024805 | NA |
| e02 | SPU_024865 | Lv-Tpase9 |
| e02 | SPU_025155 | none |
| e02 | SPU_025185 | Lv-Hus1 |
| e02 | SPU_025349 | NA |
| e02 | SPU_025373 | Lv-Tie1_8 |
| e02 | SPU_025459 | Lv-ApdL1 |
| e02 | SPU_025466 | NA |

| | | |
|---|---|---|
| e02 | SPU_025637 | Lv-Alg11 |
| e02 | SPU_025646 | Lv-Dsh |
| e02 | SPU_025702 | Lv-Guk1 |
| e02 | SPU_025770 | Lv-Pah_1 |
| e02 | SPU_025795 | Lv-Hypp_2801 |
| e02 | SPU_025806 | Lv-Tram1 |
| e02 | SPU_025884 | Lv-Ubfd1 |
| e02 | SPU_025903 | Lv-Abcc9a |
| e02 | SPU_025956 | Lv-Cyp2L60 |
| e02 | SPU_026118 | Lv-Ssu72 |
| e02 | SPU_026237 | Lv-Kif16L |
| e02 | SPU_026238 | Lv-Pigw |
| e02 | SPU_026271 | Lv-B3galt6_1 |
| e02 | SPU_026338 | Lv-Perlc |
| e02 | SPU_026389 | Lv-Metap1 |
| e02 | SPU_026502 | none |
| e02 | SPU_026518 | none |
| e02 | SPU_026759 | Lv-Tdrd9 |
| e02 | SPU_026783 | Lv-Atr |
| e02 | SPU_026864 | Lv-Noc4l |
| e02 | SPU_026892 | Lv-Tubgcp3 |
| e02 | SPU_026955 | none |
| e02 | SPU_026969 | Lv-Ank1L_1 |
| e02 | SPU_027185 | none |
| e02 | SPU_027239 | Lv-Fit2L |
| e02 | SPU_027384 | Lv-Bbs2_2 |
| e02 | SPU_027415 | Lv-RoboL_2 |
| e02 | SPU_027524 | Lv-Raf |
| e02 | SPU_027601 | Lv-PppL_160 |
| e02 | SPU_027606 | none |
| e02 | SPU_027663 | none |
| e02 | SPU_027691 | Lv-PppL_162 |
| e02 | SPU_027756 | Lv-Abcc5f |
| e02 | SPU_027779 | Lv-Polr3f |
| e02 | SPU_027901 | Lv-Mthfr |
| e02 | SPU_027921 | Lv-Dmc1 |
| e02 | SPU_027954 | Lv-Polr1e |
| e02 | SPU_028019 | Lv-Atxn10 |

| | | |
|---|---|---|
| e02 | SPU_028106 | Lv-Mfsd5 |
| e02 | SPU_028235 | Lv-Tmem214_1 |
| e02 | SPU_028493 | Lv-B3galt5_6 |
| e02 | SPU_028567 | Lv-Fbn2 |
| e02 | SPU_028620 | Lv-Perlc_1 |
| e02 | SPU_028621 | Lv-Hypp_1235 |
| e02 | SPU_028637 | none |
| e02 | SPU_028706 | Lv-Eif2a_3 |
| e02 | SPU_028710 | Lv-Diablo |
| e02 | SPU_028744 | Lv-Wdr21 |
| e02 | SPU_028824 | NA |
| e02 | SPU_028837 | Lv-Mrpl21 |
| e03 | SPU_000006 | Lv-RigIL4 |
| e03 | SPU_000017 | Lv-Z307 |
| e03 | SPU_000138 | Lv-Maeal |
| e03 | SPU_000184 | NA |
| e03 | SPU_000188 | none |
| e03 | SPU_000205 | Lv-Mtg1 |
| e03 | SPU_000244 | Lv-Prp4 |
| e03 | SPU_000249 | Lv-Mtbpl |
| e03 | SPU_000290 | Lv-Clect_63 |
| e03 | SPU_000297 | Lv-Ints5 |
| e03 | SPU_000313 | Lv-Vps37ha |
| e03 | SPU_000322 | Lv-Ubn2 |
| e03 | SPU_000324 | Lv-Asnsd1 |
| e03 | SPU_000403 | none |
| e03 | SPU_000405 | Lv-Rrp1L |
| e03 | SPU_000432 | Lv-Actl16b |
| e03 | SPU_000490 | none |
| e03 | SPU_000539 | none |
| e03 | SPU_000572 | Lv-Creld1 |
| e03 | SPU_000578 | Lv-Z460 |
| e03 | SPU_000602 | Lv-Narfl |
| e03 | SPU_000748 | NA |
| e03 | SPU_000940 | Lv-Ccdc93 |
| e03 | SPU_000982 | Lv-Ccdc5L |
| e03 | SPU_001101 | Lv-Rbm19 |
| e03 | SPU_001106 | Lv-TlrL_3 |

| | | |
|---|---|---|
| e03 | SPU_001165 | Lv-Arrdc3 |
| e03 | SPU_001239 | none |
| e03 | SPU_001262 | Lv-Birc6 |
| e03 | SPU_001280 | Lv-Ptpl |
| e03 | SPU_001377 | Lv-Atg3L |
| e03 | SPU_001412 | Lv-Vta1 |
| e03 | SPU_001489 | Lv-CalcneurL_2 |
| e03 | SPU_001505 | Lv-Gtpbp4L |
| e03 | SPU_001542 | Lv-C9orf78l |
| e03 | SPU_001545 | Lv-Spcat3l |
| e03 | SPU_001583 | Lv-Sugt1L |
| e03 | SPU_001606 | Lv-Hypp_337 |
| e03 | SPU_001641 | Lv-Lrrc57L |
| e03 | SPU_001646 | Lv-Heatr6-2 |
| e03 | SPU_001656 | none |
| e03 | SPU_001660 | NA |
| e03 | SPU_001690 | Lv-Cdk8 |
| e03 | SPU_001714 | NA |
| e03 | SPU_001723 | none |
| e03 | SPU_001789 | none |
| e03 | SPU_001798 | none |
| e03 | SPU_001830 | Lv-Trm11 |
| e03 | SPU_001931 | Lv-PrpK |
| e03 | SPU_001978 | Lv-Ttll11 |
| e03 | SPU_001979 | Lv-Stx6 |
| e03 | SPU_001994 | Lv-Fn3/Egf/Igf_1 |
| e03 | SPU_002044 | none |
| e03 | SPU_002058 | NA |
| e03 | SPU_002059 | none |
| e03 | SPU_002115 | Lv-Tud/Kh |
| e03 | SPU_002118 | Lv-RalBP1 |
| e03 | SPU_002127 | NA |
| e03 | SPU_002147 | Lv-Dslp |
| e03 | SPU_002191 | Lv-Slc35c |
| e03 | SPU_002254 | NA |
| e03 | SPU_002257 | Lv-Txnl4A |
| e03 | SPU_002294 | Lv-Nampt |
| e03 | SPU_002388 | Lv-Pik3ap1L |

| | | |
|---|---|---|
| e03 | SPU_002395 | Lv-Hypp_1392 |
| e03 | SPU_002474 | Lv-Shc1_1 |
| e03 | SPU_002511 | Lv-Taf6ll |
| e03 | SPU_002530 | Lv-Telo2 |
| e03 | SPU_002536 | Lv-Frmd4a |
| e03 | SPU_002561 | Lv-Arsj |
| e03 | SPU_002617 | Lv-Cg31140pa |
| e03 | SPU_002664 | Lv-Spag1 |
| e03 | SPU_002692 | none |
| e03 | SPU_002707 | Lv-Z388 |
| e03 | SPU_002741 | NA |
| e03 | SPU_002839 | Lv-Akap1 |
| e03 | SPU_002935 | Lv-Shoc2 |
| e03 | SPU_003026 | Lv-PolppL_19 |
| e03 | SPU_003029 | NA |
| e03 | SPU_003050 | Lv-Rad52 |
| e03 | SPU_003070 | Lv-Ap3d1 |
| e03 | SPU_003131 | none |
| e03 | SPU_003190 | Lv-Slc35b3 |
| e03 | SPU_003195 | none |
| e03 | SPU_003237 | Lv-Ankzf1-2 |
| e03 | SPU_003282 | Lv-Nup62 |
| e03 | SPU_003373 | NA |
| e03 | SPU_003589 | Lv-Mical3 |
| e03 | SPU_003596 | NA |
| e03 | SPU_003662 | Lv-Hypp_1468 |
| e03 | SPU_003685 | Lv-Z450 |
| e03 | SPU_003783 | none |
| e03 | SPU_003871 | Lv-Hypp_1485 |
| e03 | SPU_003942 | NA |
| e03 | SPU_003945 | NA |
| e03 | SPU_003970 | Lv-Tsnax |
| e03 | SPU_003979 | Lv-Fbxl7 |
| e03 | SPU_004042 | Lv-Sra1L |
| e03 | SPU_004071 | none |
| e03 | SPU_004081 | NA |
| e03 | SPU_004095 | Lv-Ankrd40 |
| e03 | SPU_004166 | Lv-PppL_174 |

| | | |
|---|---|---|
| e03 | SPU_004202 | Lv-C9orf64L |
| e03 | SPU_004206 | Lv-Prcp |
| e03 | SPU_004267 | none |
| e03 | SPU_004300 | Lv-Ppm1gl_1 |
| e03 | SPU_004318 | Lv-Prss16L |
| e03 | SPU_004330 | Lv-Rg9mtd1 |
| e03 | SPU_004354 | Lv-Nudt6 |
| e03 | SPU_004365 | Lv-Nif3l1 |
| e03 | SPU_004433 | Lv-Smpd4 |
| e03 | SPU_004454 | Lv-Thumpd3 |
| e03 | SPU_004470 | none |
| e03 | SPU_004487 | Lv-Ube2m |
| e03 | SPU_004493 | Lv-Gcnt2 |
| e03 | SPU_004513 | Lv-Ufsp2 |
| e03 | SPU_004519 | Lv-Strn |
| e03 | SPU_004687 | Lv-Tbpaf1 |
| e03 | SPU_004705 | none |
| e03 | SPU_004771 | Lv-Uxs1 |
| e03 | SPU_004796 | Lv-Prosch |
| e03 | SPU_004818 | Lv-Clect_65 |
| e03 | SPU_004868 | Lv-Sesn2 |
| e03 | SPU_004921 | Lv-IunhL_2 |
| e03 | SPU_004943 | Lv-Dscr3 |
| e03 | SPU_004994 | Lv-EgfiaL |
| e03 | SPU_005018 | Lv-Tie1L_4 |
| e03 | SPU_005059 | NA |
| e03 | SPU_005071 | Lv-Z87 |
| e03 | SPU_005103 | none |
| e03 | SPU_005137 | none |
| e03 | SPU_005174 | Lv-Dnajc3 |
| e03 | SPU_005190 | Lv-Ints10 |
| e03 | SPU_005221 | none |
| e03 | SPU_005264 | Lv-Pdella |
| e03 | SPU_005332 | none |
| e03 | SPU_005340 | none |
| e03 | SPU_005436 | Lv-Heatr5 |
| e03 | SPU_005464 | Lv-Srcr43 |
| e03 | SPU_005652 | Lv-Atm_3 |

| | | |
|---|---|---|
| e03 | SPU_005690 | Lv-Alkbh7 |
| e03 | SPU_005693 | Lv-Wdr41 |
| e03 | SPU_005727 | Lv-Hypp_1617 |
| e03 | SPU_005786 | NA |
| e03 | SPU_005874 | Lv-C12orf56L |
| e03 | SPU_005878 | Lv-CamkK |
| e03 | SPU_005881 | Lv-Sphk1 |
| e03 | SPU_005887 | Lv-Tmem177 |
| e03 | SPU_005930 | none |
| e03 | SPU_005978 | Lv-Mpi |
| e03 | SPU_006092 | Lv-Asph |
| e03 | SPU_006230 | Lv-Tim1 |
| e03 | SPU_006260 | none |
| e03 | SPU_006270 | none |
| e03 | SPU_006377 | none |
| e03 | SPU_006440 | Lv-Enox1 |
| e03 | SPU_006451 | Lv-Rmnd1h |
| e03 | SPU_006505 | Lv-Hypp_484 |
| e03 | SPU_006599 | Lv-Atpaf2 |
| e03 | SPU_006626 | Lv-Zfyve19 |
| e03 | SPU_006635 | none |
| e03 | SPU_006681 | Lv-Hypp_1673 |
| e03 | SPU_006795 | Lv-Neura |
| e03 | SPU_006811 | Lv-Mylph |
| e03 | SPU_006882 | Lv-Grx1L |
| e03 | SPU_006905 | Lv-Golga1L |
| e03 | SPU_006949 | Lv-Kiaa0528L |
| e03 | SPU_007007 | Lv-Mcm8 |
| e03 | SPU_007055 | Lv-Smu1L |
| e03 | SPU_007065 | Lv-TlrL_7 |
| e03 | SPU_007074 | Lv-PolppL_57 |
| e03 | SPU_007112 | none |
| e03 | SPU_007151 | Lv-Ctsb |
| e03 | SPU_007163 | Lv-Shc1_2 |
| e03 | SPU_007171 | Lv-Hypp_20 |
| e03 | SPU_007179 | Lv-Itpk1 |
| e03 | SPU_007196 | Lv-ZwilchL |
| e03 | SPU_007222 | Lv-Sgk |

| | | |
|---|---|---|
| e03 | SPU_007363 | Lv-Ppm1h/J/M |
| e03 | SPU_007369 | Lv-Efr3a |
| e03 | SPU_007379 | Lv-Ddb1_1 |
| e03 | SPU_007395 | Lv-Lrch3 |
| e03 | SPU_007457 | none |
| e03 | SPU_007505 | Lv-Kif6 |
| e03 | SPU_007519 | NA |
| e03 | SPU_007543 | Lv-Hypp_3071 |
| e03 | SPU_007579 | Lv-Fgfr1op2L |
| e03 | SPU_007613 | Lv-Cnot3 |
| e03 | SPU_007623 | none |
| e03 | SPU_007725 | none |
| e03 | SPU_007732 | none |
| e03 | SPU_007772 | Lv-Hdac11 |
| e03 | SPU_007864 | none |
| e03 | SPU_007868 | none |
| e03 | SPU_007869 | Lv-Eif2b2 |
| e03 | SPU_007876 | Lv-Slc35 |
| e03 | SPU_007887 | Lv-Form1ph |
| e03 | SPU_007904 | Lv-Hypp_1753 |
| e03 | SPU_007966 | Lv-PrccL |
| e03 | SPU_007970 | Lv-Ints3_1 |
| e03 | SPU_008103 | Lv-Scyl2_1 |
| e03 | SPU_008110 | Lv-Kif4_1 |
| e03 | SPU_008142 | Lv-Hsd17b12b |
| e03 | SPU_008163 | Lv-Pofut2 |
| e03 | SPU_008218 | Lv-Mark3 |
| e03 | SPU_008250 | Lv-Elovl2 |
| e03 | SPU_008253 | Lv-R/Ptp/A |
| e03 | SPU_008271 | Lv-Mpdu1 |
| e03 | SPU_008311 | Lv-Pex13 |
| e03 | SPU_008385 | Lv-Dph2h |
| e03 | SPU_008556 | Lv-Znf622 |
| e03 | SPU_008561 | Lv-Exosc2 |
| e03 | SPU_008575 | Lv-Pdia4 |
| e03 | SPU_008651 | Lv-Eif2Bg |
| e03 | SPU_008669 | Lv-TfIID11 |
| e03 | SPU_008692 | Lv-Slc35a5 |

| | | |
|---|---|---|
| e03 | SPU_008700 | Lv-Pp1L |
| e03 | SPU_008721 | Lv-Kiaa1109L_1 |
| e03 | SPU_008728 | Lv-Ggt1_8 |
| e03 | SPU_008760 | Lv-Hypp_558 |
| e03 | SPU_008840 | Lv-GtfIIA1 |
| e03 | SPU_008893 | Lv-Kiaa0319L |
| e03 | SPU_008916 | Lv-Phgdhl1 |
| e03 | SPU_008958 | Lv-Hypp_3005 |
| e03 | SPU_008959 | Lv-Cpa3L |
| e03 | SPU_008986 | Lv-Gpr107 |
| e03 | SPU_009003 | Lv-Mkrn1 |
| e03 | SPU_009016 | Lv-Sdf4 |
| e03 | SPU_009055 | NA |
| e03 | SPU_009084 | Lv-Mrpl13 |
| e03 | SPU_009108 | Lv-Hsd17b7 |
| e03 | SPU_009115 | Lv-Nup107 |
| e03 | SPU_009189 | Lv-Asb7 |
| e03 | SPU_009207 | Lv-Hypp_1833 |
| e03 | SPU_009240 | Lv-Apc7 |
| e03 | SPU_009270 | Lv-Hreif2ak |
| e03 | SPU_009318 | Lv-SnsL |
| e03 | SPU_009328 | none |
| e03 | SPU_009345 | Lv-Tfiihp44 |
| e03 | SPU_009377 | none |
| e03 | SPU_009399 | Lv-Map2k5 |
| e03 | SPU_009407 | Lv-Pdxk |
| e03 | SPU_009426 | Lv-C10orf4 |
| e03 | SPU_009429 | Lv-DhpmdL5 |
| e03 | SPU_009545 | none |
| e03 | SPU_009638 | Lv-Prss7_1 |
| e03 | SPU_009660 | NA |
| e03 | SPU_009683 | Lv-Wdr47 |
| e03 | SPU_009685 | Lv-Z227 |
| e03 | SPU_009700 | none |
| e03 | SPU_009716 | Lv-Frmd3 |
| e03 | SPU_009743 | Lv-Msto1 |
| e03 | SPU_009761 | Lv-Sae1 |
| e03 | SPU_009941 | Lv-Mios |

| | | |
|---|---|---|
| e03 | SPU_009984 | none |
| e03 | SPU_010039 | Lv-Nlr199 |
| e03 | SPU_010102 | Lv-Lrr/Kr/Fa58c |
| e03 | SPU_010117 | Lv-Setmar |
| e03 | SPU_010150 | Lv-C12orf4 |
| e03 | SPU_010201 | Lv-Aggf1 |
| e03 | SPU_010221 | Lv-Kras |
| e03 | SPU_010307 | Lv-Srcab |
| e03 | SPU_010383 | Lv-Z137 |
| e03 | SPU_010386 | Lv-Fbxo22 |
| e03 | SPU_010427 | Lv-Coq6 |
| e03 | SPU_010428 | none |
| e03 | SPU_010451 | NA |
| e03 | SPU_010510 | Lv-Rapgef2_1 |
| e03 | SPU_010522 | Lv-Rsrc2 |
| e03 | SPU_010543 | Lv-Ehmt1 |
| e03 | SPU_010658 | none |
| e03 | SPU_010825 | NA |
| e03 | SPU_010872 | Lv-Mapkkk5 |
| e03 | SPU_010879 | Lv-RtL_25 |
| e03 | SPU_010909 | Lv-Srcr93 |
| e03 | SPU_010938 | Lv-Hypp_633 |
| e03 | SPU_010939 | Lv-Slc25a38 |
| e03 | SPU_011001 | Lv-Z257 |
| e03 | SPU_011022 | Lv-Dlg |
| e03 | SPU_011129 | NA |
| e03 | SPU_011242 | Lv-Pex5 |
| e03 | SPU_011317 | Lv-Wdr32 |
| e03 | SPU_011399 | Lv-Bag2 |
| e03 | SPU_011495 | Lv-N6amt1 |
| e03 | SPU_011496 | none |
| e03 | SPU_011527 | Lv-Tiam2 |
| e03 | SPU_011550 | Lv-Med14 |
| e03 | SPU_011580 | Lv-Mgat4b |
| e03 | SPU_011603 | Lv-Usp25 |
| e03 | SPU_011695 | Lv-Mcm3apL |
| e03 | SPU_011789 | Lv-Nup54 |
| e03 | SPU_011807 | Lv-Htr1fL_2 |

| | | |
|---|---|---|
| e03 | SPU_011856 | Lv-Wdhd1 |
| e03 | SPU_011879 | Lv-Z412 |
| e03 | SPU_011908 | Lv-Hypp_75 |
| e03 | SPU_011968 | Lv-Tmem135 |
| e03 | SPU_011983 | Lv-Ppp6c_1 |
| e03 | SPU_012026 | none |
| e03 | SPU_012082 | Lv-Vash2 |
| e03 | SPU_012094 | Lv-Hypp_2016 |
| e03 | SPU_012150 | Lv-Ncapg |
| e03 | SPU_012303 | Lv-Shank1L |
| e03 | SPU_012335 | none |
| e03 | SPU_012367 | none |
| e03 | SPU_012451 | NA |
| e03 | SPU_012489 | Lv-Omi/HtrA2 |
| e03 | SPU_012530 | Lv-Fn3/Igf_8 |
| e03 | SPU_012543 | Lv-Fbxl8 |
| e03 | SPU_012545 | Lv-Dhrs1 |
| e03 | SPU_012547 | NA |
| e03 | SPU_012600 | Lv-Lss_1 |
| e03 | SPU_012605 | Lv-Ck2a |
| e03 | SPU_012675 | Lv-Slc31a1_1 |
| e03 | SPU_012846 | Lv-Ppgb |
| e03 | SPU_012870 | Lv-Gripap1 |
| e03 | SPU_012898 | Lv-Cnnm2 |
| e03 | SPU_012909 | Lv-Eif6 |
| e03 | SPU_012921 | Lv-Sgpp2 |
| e03 | SPU_012959 | Lv-D123l_1 |
| e03 | SPU_012990 | NA |
| e03 | SPU_013093 | Lv-Nmd3 |
| e03 | SPU_013132 | Lv-Mgc10433L |
| e03 | SPU_013197 | none |
| e03 | SPU_013226 | Lv-ToporsL_2 |
| e03 | SPU_013399 | Lv-Churc1 |
| e03 | SPU_013457 | none |
| e03 | SPU_013523 | Lv-RtL_34 |
| e03 | SPU_013581 | Lv-C9orf114 |
| e03 | SPU_013602 | Lv-Ccdc22 |
| e03 | SPU_013722 | none |

| | | |
|---|---|---|
| e03 | SPU_013786 | Lv-Ddx10 |
| e03 | SPU_013796 | Lv-Tmem107 |
| e03 | SPU_013922 | none |
| e03 | SPU_013931 | Lv-Scpep1 |
| e03 | SPU_013953 | Lv-Ascc2 |
| e03 | SPU_013970 | Lv-Hypp_2117 |
| e03 | SPU_014011 | Lv-P4ha1 |
| e03 | SPU_014020 | Lv-Cgi09l |
| e03 | SPU_014061 | Lv-C8orf38 |
| e03 | SPU_014076 | Lv-Sidt1 |
| e03 | SPU_014114 | Lv-Srbeta |
| e03 | SPU_014167 | Lv-CaskL |
| e03 | SPU_014237 | Lv-C12orf10L |
| e03 | SPU_014282 | Lv-Alg5 |
| e03 | SPU_014369 | Lv-Cabin1L |
| e03 | SPU_014378 | NA |
| e03 | SPU_014412 | Lv-Notchh2_3 |
| e03 | SPU_014415 | Lv-Hypp_100 |
| e03 | SPU_014462 | none |
| e03 | SPU_014474 | NA |
| e03 | SPU_014553 | Lv-Ppan |
| e03 | SPU_014575 | Lv-Dgkb |
| e03 | SPU_014627 | none |
| e03 | SPU_014628 | Lv-Erf |
| e03 | SPU_014655 | Lv-Exoc1 |
| e03 | SPU_014695 | Lv-Wdr85 |
| e03 | SPU_014727 | Lv-Trdmt1 |
| e03 | SPU_014732 | Lv-TfIID13 |
| e03 | SPU_014733 | none |
| e03 | SPU_014757 | Lv-Ranbp2_2 |
| e03 | SPU_014852 | Lv-Ubr4_2 |
| e03 | SPU_015014 | Lv-Mcm3apL_2 |
| e03 | SPU_015205 | Lv-Nlr165 |
| e03 | SPU_015213 | Lv-Mlh1 |
| e03 | SPU_015215 | Lv-3Bp1 |
| e03 | SPU_015226 | Lv-Vps13hb_3 |
| e03 | SPU_015322 | Lv-Gtmbp |
| e03 | SPU_015327 | none |

| | | |
|---|---|---|
| e03 | SPU_015329 | none |
| e03 | SPU_015369 | none |
| e03 | SPU_015380 | Lv-Fuk |
| e03 | SPU_015396 | Lv-Mboat7 |
| e03 | SPU_015443 | Lv-Eif2Be |
| e03 | SPU_015452 | Lv-Methiapd2L_1 |
| e03 | SPU_015582 | none |
| e03 | SPU_015615 | Lv-Gal3st1_7 |
| e03 | SPU_015622 | Lv-C12orf11L |
| e03 | SPU_015626 | Lv-Adrb2L_9 |
| e03 | SPU_015682 | Lv-Z417_1 |
| e03 | SPU_015697 | Lv-Rnf144a |
| e03 | SPU_015738 | Lv-Bbox1_5 |
| e03 | SPU_015873 | Lv-Rsad1 |
| e03 | SPU_016025 | Lv-CrtapL |
| e03 | SPU_016032 | Lv-Serrp5 |
| e03 | SPU_016190 | Lv-Dtnb |
| e03 | SPU_016208 | Lv-Eif1 |
| e03 | SPU_016228 | none |
| e03 | SPU_016271 | Lv-Hsd11b1 |
| e03 | SPU_016296 | none |
| e03 | SPU_016322 | Lv-Hypp_127 |
| e03 | SPU_016342 | none |
| e03 | SPU_016385 | Lv-Pctaire |
| e03 | SPU_016392 | NA |
| e03 | SPU_016495 | none |
| e03 | SPU_016503 | Lv-Hypp_3052 |
| e03 | SPU_016508 | Lv-Rbbp5 |
| e03 | SPU_016530 | Lv-Fn3/Egf_9 |
| e03 | SPU_016560 | Lv-TpaseL_8 |
| e03 | SPU_016714 | Lv-Hypp_840 |
| e03 | SPU_016799 | Lv-Arsb |
| e03 | SPU_016848 | Lv-Eapp |
| e03 | SPU_017029 | Lv-Dhx35 |
| e03 | SPU_017070 | Lv-AnL8_1 |
| e03 | SPU_017088 | Lv-Ccdc111 |
| e03 | SPU_017137 | Lv-Alg12h |
| e03 | SPU_017285 | Lv-Lias_1 |

| | | |
|---|---|---|
| e03 | SPU_017424 | Lv-Ttc36L |
| e03 | SPU_017433 | Lv-Tmem53 |
| e03 | SPU_017450 | Lv-Mare |
| e03 | SPU_017509 | Lv-Igbp1l |
| e03 | SPU_017528 | Lv-Tmem180 |
| e03 | SPU_017535 | Lv-Hypp_3042 |
| e03 | SPU_017546 | Lv-Gpkow |
| e03 | SPU_017597 | none |
| e03 | SPU_017621 | Lv-Slc6a9 |
| e03 | SPU_017642 | Lv-Smad2/3 |
| e03 | SPU_017711 | Lv-Hypp_138 |
| e03 | SPU_017920 | none |
| e03 | SPU_017937 | Lv-Fn3_35 |
| e03 | SPU_017980 | Lv-Ulk3 |
| e03 | SPU_018037 | Lv-Mios_1 |
| e03 | SPU_018145 | Lv-Hypp_2371 |
| e03 | SPU_018162 | Lv-Med30 |
| e03 | SPU_018191 | Lv-Mapkkk9_1 |
| e03 | SPU_018321 | Lv-Tor1A |
| e03 | SPU_018354 | Lv-TfIIh_p44L |
| e03 | SPU_018372 | Lv-Cyp2L35 |
| e03 | SPU_018383 | Lv-Gr |
| e03 | SPU_018423 | Lv-Ash2 |
| e03 | SPU_018439 | Lv-Commd5 |
| e03 | SPU_018498 | Lv-Als2_1 |
| e03 | SPU_018616 | Lv-Txndc11 |
| e03 | SPU_018692 | Lv-Groucho |
| e03 | SPU_018694 | Lv-C1orf43L |
| e03 | SPU_018697 | Lv-Bbs4 |
| e03 | SPU_018702 | Lv-Txndc4 |
| e03 | SPU_018710 | none |
| e03 | SPU_018750 | Lv-Slc46a1L_6 |
| e03 | SPU_018764 | Lv-Klp6L2 |
| e03 | SPU_018771 | Lv-Polr3gL |
| e03 | SPU_018774 | none |
| e03 | SPU_018861 | Lv-Thrb |
| e03 | SPU_018899 | Lv-Men1 |
| e03 | SPU_018901 | Lv-Oxch_1 |

| | | |
|---|---|---|
| e03 | SPU_018902 | Lv-M6pr |
| e03 | SPU_018978 | Lv-C1qbp |
| e03 | SPU_018987 | Lv-Tpmtl_1 |
| e03 | SPU_019004 | Lv-Lhpp |
| e03 | SPU_019017 | Lv-Cenpp |
| e03 | SPU_019111 | Lv-Xpr1_1 |
| e03 | SPU_019177 | Lv-Tprt |
| e03 | SPU_019253 | Lv-Itsn2 |
| e03 | SPU_019267 | Lv-Kiaa0543L |
| e03 | SPU_019305 | Lv-PppL_122 |
| e03 | SPU_019316 | Lv-Haus3 |
| e03 | SPU_019397 | Lv-Rwdd4a |
| e03 | SPU_019436 | Lv-Vcpip1 |
| e03 | SPU_019508 | Lv-Z371 |
| e03 | SPU_019530 | none |
| e03 | SPU_019587 | Lv-Vav |
| e03 | SPU_019627 | Lv-HsK |
| e03 | SPU_019637 | Lv-Slc22a15 |
| e03 | SPU_019646 | none |
| e03 | SPU_019748 | Lv-Tbc1d20 |
| e03 | SPU_019791 | Lv-RtL_48 |
| e03 | SPU_019824 | Lv-Z425 |
| e03 | SPU_019840 | Lv-Hypp_162 |
| e03 | SPU_019920 | Lv-PtpN12/18/22 |
| e03 | SPU_019971 | Lv-Hypp_2476 |
| e03 | SPU_020098 | Lv-Hypp_962 |
| e03 | SPU_020120 | none |
| e03 | SPU_020172 | none |
| e03 | SPU_020180 | Lv-Tbc1d10b |
| e03 | SPU_020242 | Lv-Sepsecs |
| e03 | SPU_020275 | none |
| e03 | SPU_020407 | Lv-Parp16 |
| e03 | SPU_020450 | none |
| e03 | SPU_020480 | Lv-Z77 |
| e03 | SPU_020494 | Lv-NrdlsnL |
| e03 | SPU_020528 | Lv-Tcea1 |
| e03 | SPU_020560 | Lv-C5orf21 |
| e03 | SPU_020569 | Lv-Nlr171 |

| | | |
|---|---|---|
| e03 | SPU_020570 | NA |
| e03 | SPU_020600 | Lv-Gpm6b |
| e03 | SPU_020639 | Lv-Slc39a7 |
| e03 | SPU_020640 | Lv-Ubr4_6 |
| e03 | SPU_020699 | Lv-Slc20a2 |
| e03 | SPU_020713 | Lv-Hypp_2520 |
| e03 | SPU_020808 | Lv-Man2c1 |
| e03 | SPU_020825 | NA |
| e03 | SPU_020963 | Lv-WrnL_1 |
| e03 | SPU_020974 | none |
| e03 | SPU_020978 | Lv-LrpL_3 |
| e03 | SPU_020999 | none |
| e03 | SPU_021014 | Lv-Ank3_1 |
| e03 | SPU_021135 | Lv-Pdzd8 |
| e03 | SPU_021149 | none |
| e03 | SPU_021182 | Lv-Xrcc2 |
| e03 | SPU_021195 | Lv-Apc4 |
| e03 | SPU_021350 | Lv-Trpm3_1 |
| e03 | SPU_021444 | Lv-Pelp1 |
| e03 | SPU_021473 | Lv-Ids_4 |
| e03 | SPU_021498 | Lv-Cabin1 |
| e03 | SPU_021531 | none |
| e03 | SPU_021538 | none |
| e03 | SPU_021585 | Lv-Snx29 |
| e03 | SPU_021593 | Lv-Atm_1 |
| e03 | SPU_021618 | Lv-Cep164L |
| e03 | SPU_021652 | Lv-Sat2_5 |
| e03 | SPU_021666 | Lv-Dok2 |
| e03 | SPU_021725 | Lv-Fn3/Igf_16 |
| e03 | SPU_021937 | Lv-Kiaa1524L |
| e03 | SPU_021988 | Lv-Srcr173 |
| e03 | SPU_022112 | Lv-Csk |
| e03 | SPU_022157 | Lv-Rab24 |
| e03 | SPU_022213 | Lv-Pon1 |
| e03 | SPU_022232 | Lv-Ikbkap_1 |
| e03 | SPU_022253 | Lv-Hgsnat |
| e03 | SPU_022297 | Lv-PlxnaL_1 |
| e03 | SPU_022327 | none |

| | | |
|---|---|---|
| e03 | SPU_022334 | none |
| e03 | SPU_022344 | Lv-PppL_274 |
| e03 | SPU_022394 | Lv-Nlr60 |
| e03 | SPU_022439 | none |
| e03 | SPU_022440 | Lv-Wdr55_1 |
| e03 | SPU_022504 | Lv-RvtL/End/Exn/Pp |
| e03 | SPU_022518 | Lv-Fn3f_18 |
| e03 | SPU_022593 | Lv-Cyp2L52 |
| e03 | SPU_022646 | Lv-Ogt |
| e03 | SPU_022672 | Lv-GnsL |
| e03 | SPU_022687 | Lv-Dncic2L |
| e03 | SPU_022711 | Lv-RtL_58 |
| e03 | SPU_022762 | Lv-Hrb2 |
| e03 | SPU_022765 | Lv-Fmo2_3 |
| e03 | SPU_022800 | Lv-Z236 |
| e03 | SPU_022813 | Lv-Mettl2b |
| e03 | SPU_022839 | Lv-Ptprlec2(novel) |
| e03 | SPU_022973 | Lv-Ccdc52L |
| e03 | SPU_022976 | Lv-Phka2 |
| e03 | SPU_023022 | NA |
| e03 | SPU_023074 | Lv-C8orf41L |
| e03 | SPU_023102 | Lv-Hypp_194 |
| e03 | SPU_023204 | Lv-Gpaa1 |
| e03 | SPU_023293 | Lv-Cmah |
| e03 | SPU_023326 | Lv-Naf1L |
| e03 | SPU_023402 | Lv-Tsc2_1 |
| e03 | SPU_023414 | Lv-Fggy |
| e03 | SPU_023492 | none |
| e03 | SPU_023547 | Lv-C1galt1_3 |
| e03 | SPU_023631 | Lv-Ppapdc1 |
| e03 | SPU_023632 | Lv-Mfsd10 |
| e03 | SPU_023728 | Lv-Hypp_1095 |
| e03 | SPU_023740 | Lv-Thap9 |
| e03 | SPU_024040 | none |
| e03 | SPU_024079 | Lv-Adora2b_1 |
| e03 | SPU_024135 | Lv-Gpr155 |
| e03 | SPU_024136 | Lv-Narg2L |
| e03 | SPU_024182 | none |

| | | |
|---|---|---|
| e03 | SPU_024312 | Lv-Gns |
| e03 | SPU_024380 | Lv-Hypp_2726 |
| e03 | SPU_024392 | Lv-Sfrs8_1 |
| e03 | SPU_024401 | Lv-NfkbizL |
| e03 | SPU_024428 | Lv-C7orf42 |
| e03 | SPU_024441 | Lv-Snapc4L |
| e03 | SPU_024532 | Lv-Neu1 |
| e03 | SPU_024644 | Lv-Tada2l_1 |
| e03 | SPU_024748 | Lv-Z276 |
| e03 | SPU_024750 | Lv-Rfc2 |
| e03 | SPU_024801 | Lv-Brrn1 |
| e03 | SPU_024852 | Lv-Hypp_1128 |
| e03 | SPU_024866 | Lv-Slc16a12_5 |
| e03 | SPU_024997 | Lv-Prrc1 |
| e03 | SPU_025045 | Lv-Slc39a1_1 |
| e03 | SPU_025051 | NA |
| e03 | SPU_025059 | Lv-Fbln2L |
| e03 | SPU_025082 | Lv-RagC |
| e03 | SPU_025117 | Lv-Hypp_225 |
| e03 | SPU_025138 | Lv-Nlr68 |
| e03 | SPU_025176 | Lv-Atm |
| e03 | SPU_025186 | none |
| e03 | SPU_025226 | Lv-Gpr54_2 |
| e03 | SPU_025247 | Lv-Ccdc149 |
| e03 | SPU_025260 | none |
| e03 | SPU_025275 | NA |
| e03 | SPU_025303 | Lv-Adcy |
| e03 | SPU_025428 | Lv-Ilk |
| e03 | SPU_025470 | Lv-Slc25a19 |
| e03 | SPU_025489 | Lv-Smug1 |
| e03 | SPU_025518 | Lv-Tim2 |
| e03 | SPU_025625 | Lv-Slc10a7 |
| e03 | SPU_025654 | Lv-Armc1 |
| e03 | SPU_025678 | none |
| e03 | SPU_025739 | Lv-RtL_62 |
| e03 | SPU_025744 | Lv-Apc10 |
| e03 | SPU_025788 | Lv-Fn3/Egf/Lnb/Tm7 |
| e03 | SPU_025810 | none |

| | | |
|---|---|---|
| e03 | SPU_025824 | NA |
| e03 | SPU_025825 | NA |
| e03 | SPU_025868 | Lv-C20orf11L |
| e03 | SPU_025940 | Lv-Inoc1 |
| e03 | SPU_025988 | Lv-Ngly1L |
| e03 | SPU_026005 | Lv-Cop9scs5_1 |
| e03 | SPU_026027 | Lv-Btbd10 |
| e03 | SPU_026124 | Lv-Ppp1r16b |
| e03 | SPU_026169 | Lv-Kiaa0766 |
| e03 | SPU_026226 | Lv-Fh1 |
| e03 | SPU_026382 | Lv-Aup1l |
| e03 | SPU_026401 | Lv-Polr3a_1 |
| e03 | SPU_026410 | Lv-Hypp_2839 |
| e03 | SPU_026445 | Lv-Agfg1L |
| e03 | SPU_026446 | none |
| e03 | SPU_026684 | none |
| e03 | SPU_026766 | Lv-Sfk5 |
| e03 | SPU_026777 | Lv-Mt |
| e03 | SPU_026810 | none |
| e03 | SPU_026887 | Lv-Hypp_1190 |
| e03 | SPU_026888 | Lv-Aamp |
| e03 | SPU_026924 | Lv-Chst11_2 |
| e03 | SPU_026961 | Lv-Rhogap19 |
| e03 | SPU_026962 | Lv-Ldb2 |
| e03 | SPU_027075 | none |
| e03 | SPU_027144 | Lv-Pi3kp110 |
| e03 | SPU_027174 | Lv-Ccdc51 |
| e03 | SPU_027209 | Lv-Mgll |
| e03 | SPU_027237 | Lv-Napg |
| e03 | SPU_027242 | Lv-Kiaa1609l |
| e03 | SPU_027286 | Lv-GagpolL_15 |
| e03 | SPU_027320 | Lv-IlvbL_1 |
| e03 | SPU_027443 | Lv-PlexA4_1 |
| e03 | SPU_027489 | NA |
| e03 | SPU_027493 | none |
| e03 | SPU_027683 | Lv-Pgbd4_1 |
| e03 | SPU_027687 | Lv-Tie2L_1 |
| e03 | SPU_027697 | NA |

| | | |
|---|---|---|
| e03 | SPU_027720 | Lv-Hypp_1210 |
| e03 | SPU_027766 | Lv-Mettl9 |
| e03 | SPU_027768 | Lv-Prmt5 |
| e03 | SPU_027826 | Lv-GbB |
| e03 | SPU_027890 | Lv-Gfm1 |
| e03 | SPU_028059 | none |
| e03 | SPU_028079 | none |
| e03 | SPU_028164 | Lv-Endogl1_1 |
| e03 | SPU_028193 | NA |
| e03 | SPU_028213 | Lv-Rc74 |
| e03 | SPU_028273 | Lv-My10f_3 |
| e03 | SPU_028300 | Lv-B7L4 |
| e03 | SPU_028305 | Lv-Fech |
| e03 | SPU_028311 | none |
| e03 | SPU_028332 | Lv-Smarca3 |
| e03 | SPU_028338 | Lv-Abca1L |
| e03 | SPU_028360 | Lv-Rabggtb |
| e03 | SPU_028396 | Lv-Tex10L |
| e03 | SPU_028425 | Lv-Blzf1 |
| e03 | SPU_028496 | Lv-Thnsl1_1 |
| e03 | SPU_028548 | Lv-Elovl6_4 |
| e03 | SPU_028711 | Lv-RdbpL |
| e03 | SPU_028726 | Lv-PlexA4 |
| e03 | SPU_028848 | Lv-Ing1 |
| e04 | SPU_000011 | none |
| e04 | SPU_000125 | none |
| e04 | SPU_000137 | Lv-destabilaseL |
| e04 | SPU_000222 | Lv-Pgrp3 |
| e04 | SPU_000223 | Lv-Slc16a12_10 |
| e04 | SPU_000240 | Lv-Ube2zl |
| e04 | SPU_000367 | NA |
| e04 | SPU_000392 | none |
| e04 | SPU_000410 | Lv-Egfib_3 |
| e04 | SPU_000484 | Lv-Cytb561 |
| e04 | SPU_000548 | Lv-Vps41h |
| e04 | SPU_000560 | Lv-March8l |
| e04 | SPU_000606 | none |
| e04 | SPU_000784 | Lv-Twnfln |

| | | |
|---|---|---|
| e04 | SPU_000938 | none |
| e04 | SPU_000939 | Lv-Hpcl |
| e04 | SPU_000948 | Lv-Lass6 |
| e04 | SPU_001009 | Lv-Xpf_1 |
| e04 | SPU_001046 | Lv-Reps1L |
| e04 | SPU_001077 | Lv-Chst11-5 |
| e04 | SPU_001082 | Lv-Hypp_1302 |
| e04 | SPU_001150 | NA |
| e04 | SPU_001240 | Lv-Tep1 |
| e04 | SPU_001297 | Lv-Hypp_1320 |
| e04 | SPU_001333 | Lv-C14orf124 |
| e04 | SPU_001535 | Lv-Ankdd1a |
| e04 | SPU_001558 | Lv-Poll |
| e04 | SPU_001621 | Lv-Gprk1 |
| e04 | SPU_001695 | Lv-Eea1 |
| e04 | SPU_001698 | Lv-Gat_1 |
| e04 | SPU_001710 | none |
| e04 | SPU_001726 | Lv-Amt3 |
| e04 | SPU_001810 | Lv-PolppL-5 |
| e04 | SPU_001959 | none |
| e04 | SPU_001990 | none |
| e04 | SPU_002002 | Lv-Tim21L |
| e04 | SPU_002057 | Lv-Eap1L |
| e04 | SPU_002167 | Lv-Drd2L_12 |
| e04 | SPU_002174 | Lv-CpsiL |
| e04 | SPU_002189 | Lv-Adap1 |
| e04 | SPU_002347 | none |
| e04 | SPU_002425 | Lv-Lrsam1 |
| e04 | SPU_002534 | Lv-Tubgcp6L |
| e04 | SPU_002672 | Lv-Mtif3L |
| e04 | SPU_002696 | Lv-Tak1 |
| e04 | SPU_002727 | Lv-Wwox_2 |
| e04 | SPU_002781 | Lv-Spg21 |
| e04 | SPU_002835 | NA |
| e04 | SPU_002865 | Lv-Tep1L_4 |
| e04 | SPU_002882 | Lv-Usp46L |
| e04 | SPU_003072 | Lv-Elovl7L |
| e04 | SPU_003158 | Lv-Ssh |

| | | |
|---|---|---|
| e04 | SPU_003241 | Lv-Abcb7 |
| e04 | SPU_003284 | Lv-Kringle/Clect/Egf |
| e04 | SPU_003294 | Lv-Samd9L_3 |
| e04 | SPU_003296 | Lv-Snf7/cytda |
| e04 | SPU_003357 | Lv-Carkl |
| e04 | SPU_003381 | NA |
| e04 | SPU_003450 | Lv-Sorbs1 |
| e04 | SPU_003523 | Lv-Hypp_1460 |
| e04 | SPU_003554 | Lv-Hypp_3090 |
| e04 | SPU_003590 | Lv-Kiaa1841L |
| e04 | SPU_003827 | Lv-Mat2b |
| e04 | SPU_003915 | Lv-Insr |
| e04 | SPU_003938 | none |
| e04 | SPU_004107 | Lv-Sarmr12 |
| e04 | SPU_004146 | none |
| e04 | SPU_004156 | NA |
| e04 | SPU_004256 | Lv-Gnb1L |
| e04 | SPU_004269 | Lv-Mycbp2 |
| e04 | SPU_004279 | Lv-Ankrd10 |
| e04 | SPU_004320 | Lv-Itfg1l |
| e04 | SPU_004398 | Lv-Tm2d1 |
| e04 | SPU_004442 | Lv-Rab9 |
| e04 | SPU_004482 | Lv-Ugt2b37L |
| e04 | SPU_004500 | Lv-TrypsinL_1 |
| e04 | SPU_004548 | Lv-Adipor2 |
| e04 | SPU_004583 | Lv-Hypp_1535 |
| e04 | SPU_004640 | none |
| e04 | SPU_004683 | none |
| e04 | SPU_004702 | Lv-Ebf3 |
| e04 | SPU_004718 | Lv-Mtif2L2 |
| e04 | SPU_004782 | none |
| e04 | SPU_005102 | Lv-Nhlrc2 |
| e04 | SPU_005186 | NA |
| e04 | SPU_005197 | Lv-Mcm3apL_3 |
| e04 | SPU_005241 | Lv-Gne |
| e04 | SPU_005338 | Lv-Ears2 |
| e04 | SPU_005401 | none |
| e04 | SPU_005454 | Lv-Endrvt14 |

| | | |
|---|---|---|
| e04 | SPU_005489 | Lv-Fhod3 |
| e04 | SPU_005546 | Lv-Hypp_1588 |
| e04 | SPU_005606 | Lv-Pus7hL |
| e04 | SPU_005686 | Lv-Pcbp3 |
| e04 | SPU_005711 | Lv-Sars2-2 |
| e04 | SPU_005751 | Lv-Ptchd3 |
| e04 | SPU_005792 | Lv-Hypp_88 |
| e04 | SPU_005817 | Lv-Cc2d2a |
| e04 | SPU_005852 | Lv-Impdh2 |
| e04 | SPU_005901 | none |
| e04 | SPU_005997 | Lv-Unc44L_49 |
| e04 | SPU_006106 | none |
| e04 | SPU_006134 | NA |
| e04 | SPU_006174 | Lv-Tm2d2_1 |
| e04 | SPU_006317 | Lv-Ddrgk1_1 |
| e04 | SPU_006332 | Lv-Fbxl12 |
| e04 | SPU_006335 | Lv-Hypp_476 |
| e04 | SPU_006388 | Lv-Rab30l |
| e04 | SPU_006398 | Lv-Notch2-10 |
| e04 | SPU_006528 | Lv-Rptp_mu_1 |
| e04 | SPU_006683 | Lv-Pax1-9 |
| e04 | SPU_006778 | Lv-Serca |
| e04 | SPU_006802 | none |
| e04 | SPU_006812 | Lv-Kirrel2L |
| e04 | SPU_006881 | Lv-Fbn2L |
| e04 | SPU_006886 | Lv-Bet1 |
| e04 | SPU_007023 | Lv-My1a/My1b/My1c |
| e04 | SPU_007027 | none |
| e04 | SPU_007052 | Lv-St14L |
| e04 | SPU_007069 | Lv-Slit1bL |
| e04 | SPU_007081 | Lv-Serrp15 |
| e04 | SPU_007097 | Lv-Rassf5 |
| e04 | SPU_007277 | Lv-UredL |
| e04 | SPU_007501 | Lv-Hypp_1725 |
| e04 | SPU_007514 | none |
| e04 | SPU_007567 | Lv-Gpr161 |
| e04 | SPU_007635 | Lv-Hypp_1738 |
| e04 | SPU_007847 | none |

| | | |
|---|---|---|
| e04 | SPU_007935 | Lv-Gde1 |
| e04 | SPU_007952 | Lv-Tirc2_1 |
| e04 | SPU_007954 | Lv-Bivm |
| e04 | SPU_007959 | Lv-Hypp_1756 |
| e04 | SPU_007961 | none |
| e04 | SPU_008139 | Lv-PolppL_67 |
| e04 | SPU_008174 | none |
| e04 | SPU_008180 | Lv-Pkci |
| e04 | SPU_008339 | Lv-Slc6a5_1 |
| e04 | SPU_008364 | Lv-Chst1 |
| e04 | SPU_008406 | Lv-Acads_1 |
| e04 | SPU_008608 | Lv-C9orf126 |
| e04 | SPU_008616 | Lv-Gmpr |
| e04 | SPU_008652 | Lv-Psat1_1 |
| e04 | SPU_008677 | Lv-Msr |
| e04 | SPU_008758 | Lv-RtL_20 |
| e04 | SPU_008888 | Lv-Hypp_38 |
| e04 | SPU_008930 | Lv-Isa |
| e04 | SPU_008951 | NA |
| e04 | SPU_008977 | Lv-Fbxo6 |
| e04 | SPU_008999 | Lv-Hypp_1819 |
| e04 | SPU_009170 | none |
| e04 | SPU_009190 | Lv-Ensa |
| e04 | SPU_009289 | Lv-4Cl_1 |
| e04 | SPU_009303 | none |
| e04 | SPU_009317 | none |
| e04 | SPU_009342 | NA |
| e04 | SPU_009404 | Lv-Trpc4L |
| e04 | SPU_009547 | Lv-Mtx2 |
| e04 | SPU_009554 | Lv-Herc4-4 |
| e04 | SPU_009691 | Lv-Plceta |
| e04 | SPU_009706 | NA |
| e04 | SPU_009707 | Lv-Cars2 |
| e04 | SPU_009841 | Lv-Pprppsl |
| e04 | SPU_010033 | Lv-Rhogef4 |
| e04 | SPU_010056 | Lv-Khdrbh2 |
| e04 | SPU_010185 | none |
| e04 | SPU_010452 | Lv-Alkbh8 |

| | | |
|---|---|---|
| e04 | SPU_010517 | Lv-Unk_73 |
| e04 | SPU_010569 | Lv-Gpr105L |
| e04 | SPU_010761 | Lv-Slc35a1_1 |
| e04 | SPU_010763 | Lv-Eph_1 |
| e04 | SPU_010777 | Lv-Fadd |
| e04 | SPU_010781 | Lv-Alg2 |
| e04 | SPU_010808 | Lv-LshmanlsnL_1 |
| e04 | SPU_010851 | Lv-CdoL |
| e04 | SPU_010903 | Lv-Hypp_1945 |
| e04 | SPU_010954 | Lv-Kap |
| e04 | SPU_011017 | Lv-Atr_1 |
| e04 | SPU_011083 | none |
| e04 | SPU_011093 | Lv-Mccc1 |
| e04 | SPU_011186 | Lv-Sdr16c5 |
| e04 | SPU_011261 | none |
| e04 | SPU_011300 | none |
| e04 | SPU_011351 | Lv-Ppm1g |
| e04 | SPU_011443 | none |
| e04 | SPU_011473 | Lv-Fn3 |
| e04 | SPU_011508 | Lv-Hypp_69 |
| e04 | SPU_011523 | Lv-Ttrl |
| e04 | SPU_011689 | Lv-Ttc15 |
| e04 | SPU_011705 | Lv-Ppm1k |
| e04 | SPU_011728 | Lv-Hypp_1999 |
| e04 | SPU_011941 | NA |
| e04 | SPU_011963 | Lv-Tsg101 |
| e04 | SPU_011976 | Lv-Notchl3 |
| e04 | SPU_012044 | Lv-Parp4l |
| e04 | SPU_012075 | Lv-Apeh-2 |
| e04 | SPU_012078 | Lv-Ip3r |
| e04 | SPU_012199 | Lv-Ugt1L_3 |
| e04 | SPU_012229 | Lv-Apg7l |
| e04 | SPU_012270 | none |
| e04 | SPU_012275 | Lv-Rbck1 |
| e04 | SPU_012329 | Lv-Stx1 |
| e04 | SPU_012357 | Lv-Zdhhc24_1 |
| e04 | SPU_012428 | none |
| e04 | SPU_012560 | Lv-Bbs8 |

| | | |
|---|---|---|
| e04 | SPU_012644 | Lv-Acox1_1 |
| e04 | SPU_012690 | Lv-Slc15a2 |
| e04 | SPU_012792 | Lv-Trpc3 |
| e04 | SPU_012828 | none |
| e04 | SPU_012915 | none |
| e04 | SPU_012934 | Lv-Hypp_2057 |
| e04 | SPU_012985 | NA |
| e04 | SPU_013073 | Lv-PolypL_5 |
| e04 | SPU_013116 | none |
| e04 | SPU_013138 | none |
| e04 | SPU_013219 | NA |
| e04 | SPU_013229 | Lv-C10orf118L |
| e04 | SPU_013364 | Lv-Asci1cs3 |
| e04 | SPU_013525 | Lv-Lrrc16a_1 |
| e04 | SPU_013634 | Lv-Osbpl8 |
| e04 | SPU_013675 | Lv-Ap2b1 |
| e04 | SPU_013716 | Lv-Sorc |
| e04 | SPU_013918 | Lv-Drd1L_4 |
| e04 | SPU_013919 | Lv-Fank1 |
| e04 | SPU_013950 | Lv-Il1r/Rs1 |
| e04 | SPU_013991 | Lv-Tmem30a |
| e04 | SPU_014027 | Lv-Ociad1 |
| e04 | SPU_014037 | Lv-Par3ph1 |
| e04 | SPU_014060 | Lv-Mrpl3l |
| e04 | SPU_014319 | Lv-Pxmp4 |
| e04 | SPU_014493 | Lv-Hypp_2154 |
| e04 | SPU_014610 | Lv-Ipo4 |
| e04 | SPU_014659 | Lv-Nek11_1 |
| e04 | SPU_014713 | none |
| e04 | SPU_014835 | none |
| e04 | SPU_014951 | Lv-SfxL |
| e04 | SPU_014997 | Lv-PolypL_9 |
| e04 | SPU_015037 | Lv-SacSl_6 |
| e04 | SPU_015091 | none |
| e04 | SPU_015103 | Lv-CesL |
| e04 | SPU_015120 | Lv-Cyp20Lp |
| e04 | SPU_015241 | Lv-Gnpat |
| e04 | SPU_015436 | Lv-Unc44L_109 |

| | | |
|---|---|---|
| e04 | SPU_015495 | Lv-Z224 |
| e04 | SPU_015496 | Lv-Hypp_2217 |
| e04 | SPU_015514 | Lv-Rdh14 |
| e04 | SPU_015698 | none |
| e04 | SPU_015776 | Lv-Kri1 |
| e04 | SPU_015794 | Lv-Hypp_801 |
| e04 | SPU_015856 | Lv-Lamp1 |
| e04 | SPU_015894 | Lv-Scnn1g |
| e04 | SPU_015919 | Lv-Fn3_32 |
| e04 | SPU_016013 | Lv-Dgkq |
| e04 | SPU_016027 | none |
| e04 | SPU_016045 | Lv-Pka |
| e04 | SPU_016065 | none |
| e04 | SPU_016115 | Lv-Dnajhc13 |
| e04 | SPU_016216 | Lv-Kiaa1841 |
| e04 | SPU_016287 | Lv-Hagh |
| e04 | SPU_016305 | none |
| e04 | SPU_016409 | Lv-Dsp3_1 |
| e04 | SPU_016435 | Lv-Kntc1L |
| e04 | SPU_016597 | Lv-Galr2L_22 |
| e04 | SPU_016605 | Lv-Rab14 |
| e04 | SPU_016722 | Lv-Hypp_2272 |
| e04 | SPU_016808 | none |
| e04 | SPU_016811 | Lv-Endrvt94 |
| e04 | SPU_016934 | Lv-Cask |
| e04 | SPU_017006 | Lv-Clect/Tm_5 |
| e04 | SPU_017077 | Lv-Ccndbp1 |
| e04 | SPU_017103 | Lv-Frrs1 |
| e04 | SPU_017212 | Lv-Slc45a2_1 |
| e04 | SPU_017260 | Lv-Hypp_868 |
| e04 | SPU_017393 | Lv-PppL_209 |
| e04 | SPU_017491 | Lv-Nr1m1 |
| e04 | SPU_017627 | Lv-GliA |
| e04 | SPU_017646 | none |
| e04 | SPU_017680 | Lv-Dhps_3 |
| e04 | SPU_017705 | Lv-Cdk5 |
| e04 | SPU_017745 | Lv-Bola1 |
| e04 | SPU_017755 | Lv-C3orf1 |

| | | |
|---|---|---|
| e04 | SPU_017921 | none |
| e04 | SPU_018060 | Lv-Fut7 |
| e04 | SPU_018168 | Lv-Sarmr9 |
| e04 | SPU_018171 | Lv-Adam17L1 |
| e04 | SPU_018271 | Lv-Ccs |
| e04 | SPU_018330 | Lv-Commd8 |
| e04 | SPU_018357 | Lv-Gpr21L_12 |
| e04 | SPU_018377 | Lv-Reep5 |
| e04 | SPU_018477 | Lv-Egfib_8 |
| e04 | SPU_018490 | Lv-Guf1 |
| e04 | SPU_018513 | Lv-PappA1L |
| e04 | SPU_018545 | Lv-Acox1_2 |
| e04 | SPU_018598 | Lv-Ubc13 |
| e04 | SPU_018620 | Lv-Gpr51_1 |
| e04 | SPU_018653 | Lv-Elovl6_7 |
| e04 | SPU_018791 | Lv-Slc5a12_2 |
| e04 | SPU_018872 | Lv-Kcnk5_1 |
| e04 | SPU_018948 | NA |
| e04 | SPU_019027 | Lv-Aadat_5 |
| e04 | SPU_019030 | Lv-Mbd5 |
| e04 | SPU_019166 | Lv-Gnpda1 |
| e04 | SPU_019185 | Lv-Abcb1_2 |
| e04 | SPU_019225 | Lv-Spata20 |
| e04 | SPU_019259 | Lv-ActinL7 |
| e04 | SPU_019300 | Lv-Napa |
| e04 | SPU_019373 | Lv-Surf5 |
| e04 | SPU_019405 | Lv-Pnkp_1 |
| e04 | SPU_019414 | none |
| e04 | SPU_019470 | Lv-Ugt1L_6 |
| e04 | SPU_019484 | Lv-Hspbap1 |
| e04 | SPU_019574 | Lv-Hypp_2451 |
| e04 | SPU_019686 | Lv-Fak |
| e04 | SPU_019738 | NA |
| e04 | SPU_019768 | none |
| e04 | SPU_019794 | Lv-Cas |
| e04 | SPU_019813 | Lv-Ghdc1L |
| e04 | SPU_020000 | none |
| e04 | SPU_020001 | Lv-Sgpl1 |

| | | |
|---|---|---|
| e04 | SPU_020014 | Lv-Ogdh_2 |
| e04 | SPU_020131 | Lv-Tirc7 |
| e04 | SPU_020137 | Lv-Hypp_2490 |
| e04 | SPU_020138 | Lv-Psat1 |
| e04 | SPU_020201 | Lv-Dennd2a |
| e04 | SPU_020231 | Lv-Fbxl15 |
| e04 | SPU_020243 | Lv-Sh3d19L_1 |
| e04 | SPU_020309 | Lv-Pip4k2b |
| e04 | SPU_020397 | Lv-Elovl4 |
| e04 | SPU_020426 | Lv-Abca5_1 |
| e04 | SPU_020578 | Lv-Gdpd5 |
| e04 | SPU_020615 | Lv-Cqrrf1 |
| e04 | SPU_020623 | Lv-Cspe8L4 |
| e04 | SPU_020629 | none |
| e04 | SPU_020667 | Lv-Htr4L_8 |
| e04 | SPU_020829 | Lv-Hypp_998 |
| e04 | SPU_020946 | Lv-Cbxh4 |
| e04 | SPU_021008 | Lv-Aki |
| e04 | SPU_021032 | NA |
| e04 | SPU_021212 | Lv-Hypp_1016 |
| e04 | SPU_021296 | Lv-PppL_127 |
| e04 | SPU_021335 | Lv-SacSl_7 |
| e04 | SPU_021378 | none |
| e04 | SPU_021409 | none |
| e04 | SPU_021416 | Lv-Boc |
| e04 | SPU_021451 | Lv-Hypp_2555 |
| e04 | SPU_021504 | Lv-Mek |
| e04 | SPU_021581 | Lv-Dyspph2 |
| e04 | SPU_021693 | Lv-Mcee |
| e04 | SPU_021727 | Lv-Ipo9 |
| e04 | SPU_021971 | Lv-Golga1 |
| e04 | SPU_021979 | Lv-Mthfd1 |
| e04 | SPU_021991 | Lv-Cpa2L_4 |
| e04 | SPU_022004 | Lv-Rhogef9 |
| e04 | SPU_022107 | Lv-AmotL |
| e04 | SPU_022127 | NA |
| e04 | SPU_022164 | Lv-Mast2 |
| e04 | SPU_022563 | Lv-Endrvt8 |

| | | |
|---|---|---|
| e04 | SPU_022588 | Lv-Dnase1_2 |
| e04 | SPU_022692 | none |
| e04 | SPU_022756 | Lv-Dctn6 |
| e04 | SPU_022788 | none |
| e04 | SPU_022823 | Lv-Hypp_1058 |
| e04 | SPU_022873 | Lv-Egf/Zp_2 |
| e04 | SPU_022946 | Lv-Fam8a1 |
| e04 | SPU_022996 | Lv-Hypp_2642 |
| e04 | SPU_023073 | Lv-Unc44_338 |
| e04 | SPU_023258 | Lv-PolypL_119 |
| e04 | SPU_023341 | none |
| e04 | SPU_023423 | Lv-Wasf2 |
| e04 | SPU_023497 | NA |
| e04 | SPU_023542 | Lv-RtL_4 |
| e04 | SPU_023599 | Lv-Dnamth |
| e04 | SPU_023609 | Lv-Pla2g6 |
| e04 | SPU_023613 | Lv-Ubc |
| e04 | SPU_023645 | none |
| e04 | SPU_023676 | Lv-Mkp5 |
| e04 | SPU_023764 | Lv-Atp6v0a1_2 |
| e04 | SPU_023809 | none |
| e04 | SPU_023903 | Lv-PapL1 |
| e04 | SPU_023981 | Lv-Hypp_2702 |
| e04 | SPU_024025 | NA |
| e04 | SPU_024038 | none |
| e04 | SPU_024077 | Lv-TtlL |
| e04 | SPU_024327 | Lv-PppL_221 |
| e04 | SPU_024481 | none |
| e04 | SPU_024509 | none |
| e04 | SPU_024523 | Lv-Amds_4 |
| e04 | SPU_024772 | Lv-Cox11l |
| e04 | SPU_024822 | Lv-Hypp_2754 |
| e04 | SPU_024827 | Lv-NehL |
| e04 | SPU_024893 | Lv-Ap2a2 |
| e04 | SPU_024912 | Lv-Czph |
| e04 | SPU_025012 | Lv-Hypp_223 |
| e04 | SPU_025058 | Lv-Odin/CajL |
| e04 | SPU_025081 | Lv-Tmcc |

| | | |
|---|---|---|
| e04 | SPU_025159 | Lv-Mapk15 |
| e04 | SPU_025223 | Lv-Fh1/Fh2cph |
| e04 | SPU_025276 | Lv-Mtmr12L |
| e04 | SPU_025339 | Lv-Rej8L |
| e04 | SPU_025465 | Lv-Rt |
| e04 | SPU_025554 | Lv-Dnmbp |
| e04 | SPU_025618 | Lv-Hypp_232 |
| e04 | SPU_025798 | Lv-Smile_1 |
| e04 | SPU_025867 | Lv-Slc16a12L_18 |
| e04 | SPU_025928 | Lv-Lrrc47 |
| e04 | SPU_026033 | Lv-Hypp_1166 |
| e04 | SPU_026125 | none |
| e04 | SPU_026154 | Lv-Crea |
| e04 | SPU_026171 | Lv-Hypp_2824 |
| e04 | SPU_026202 | Lv-Mmab |
| e04 | SPU_026244 | Lv-Wdr69 |
| e04 | SPU_026278 | Lv-Thnsl1 |
| e04 | SPU_026284 | NA |
| e04 | SPU_026308 | none |
| e04 | SPU_026309 | Lv-Aifm3 |
| e04 | SPU_026467 | Lv-Wdr17 |
| e04 | SPU_026505 | Lv-Fut8-4 |
| e04 | SPU_026548 | Lv-Ppfia1 |
| e04 | SPU_026558 | Lv-Dhrs6 |
| e04 | SPU_026649 | Lv-Notch1b |
| e04 | SPU_026773 | Lv-Grm6L |
| e04 | SPU_027013 | Lv-Unc13dL |
| e04 | SPU_027042 | Lv-C3orf33L |
| e04 | SPU_027225 | none |
| e04 | SPU_027265 | Lv-PppL_157 |
| e04 | SPU_027271 | Lv-Zfyve21L |
| e04 | SPU_027279 | Lv-Cc2d1a |
| e04 | SPU_027346 | Lv-Hypp_2899 |
| e04 | SPU_027359 | Lv-Fbxo8 |
| e04 | SPU_027526 | Lv-PlexA2 |
| e04 | SPU_027528 | Lv-Pdzrn4 |
| e04 | SPU_027599 | Lv-Decr1_1 |
| e04 | SPU_027608 | none |

| | | |
|---|---|---|
| e04 | SPU_027674 | Lv-Ip3r_5 |
| e04 | SPU_027727 | none |
| e04 | SPU_027782 | Lv-R3hdm1 |
| e04 | SPU_027809 | Lv-Hypp_2920 |
| e04 | SPU_027889 | Lv-Bgal |
| e04 | SPU_027996 | Lv-Hypp_2925 |
| e04 | SPU_028018 | NA |
| e04 | SPU_028068 | Lv-Kcnma1 |
| e04 | SPU_028129 | Lv-Aoah |
| e04 | SPU_028147 | Lv-Hyd |
| e04 | SPU_028178 | Lv-Gaa_1 |
| e04 | SPU_028263 | Lv-Exoc2 |
| e04 | SPU_028291 | Lv-Unc44_185 |
| e04 | SPU_028302 | NA |
| e04 | SPU_028388 | Lv-PolypL_129 |
| e04 | SPU_028412 | Lv-Ack1_1 |
| e04 | SPU_028413 | NA |
| e04 | SPU_028457 | Lv-Itsn2_1 |
| e04 | SPU_028782 | NA |
| e04 | SPU_028925 | none |
| e04 | SPU_028939 | Lv-Naalad2L_10 |
| e04 | SPU_030066 | Lv-Jak_2 |
| e04 | SPU_030085 | Lv-Gg3 |
| e04 | SPU_030204 | Lv-Il17-24 |
| e05 | SPU_000066 | Lv-Nsun7l |
| e05 | SPU_000092 | Lv-Kcnh1 |
| e05 | SPU_000341 | Lv-Hypp_3124 |
| e05 | SPU_000429 | none |
| e05 | SPU_000667 | Lv-Cret |
| e05 | SPU_000889 | none |
| e05 | SPU_001002 | none |
| e05 | SPU_001079 | Lv-Mocos |
| e05 | SPU_001202 | none |
| e05 | SPU_001330 | Lv-Epb4.1l1 |
| e05 | SPU_001427 | Lv-Syne1L |
| e05 | SPU_001785 | Lv-Eggbr1L |
| e05 | SPU_001906 | Lv-Slc16a12L_2 |
| e05 | SPU_001916 | Lv-Mil2 |

| | | |
|---|---|---|
| e05 | SPU_001942 | Lv-Vwd/Vwc |
| e05 | SPU_002060 | Lv-Tnrc9 |
| e05 | SPU_002073 | Lv-Ryr3L_2 |
| e05 | SPU_002148 | Lv-Dnajb1 |
| e05 | SPU_002177 | none |
| e05 | SPU_002289 | Lv-Nubp1-2 |
| e05 | SPU_002410 | Lv-EchnL_3 |
| e05 | SPU_002533 | Lv-Melkl |
| e05 | SPU_002589 | Lv-AchE_2 |
| e05 | SPU_002590 | Lv-Cyp46L1 |
| e05 | SPU_002657 | Lv-Fn3f_8 |
| e05 | SPU_002802 | Lv-Fn3_17 |
| e05 | SPU_002840 | Lv-Ilgfr |
| e05 | SPU_002853 | none |
| e05 | SPU_002917 | NA |
| e05 | SPU_003078 | Lv-Idua |
| e05 | SPU_003177 | none |
| e05 | SPU_003238 | Lv-Hypp_1437 |
| e05 | SPU_003257 | Lv-Dystoh3 |
| e05 | SPU_003344 | Lv-Wdr59 |
| e05 | SPU_003361 | Lv-Hypp_1447 |
| e05 | SPU_003371 | Lv-PolppL_31 |
| e05 | SPU_003412 | Lv-C7orf58L |
| e05 | SPU_003417 | Lv-Trim2 |
| e05 | SPU_003613 | Lv-Kcnk10 |
| e05 | SPU_003669 | Lv-Plcb4 |
| e05 | SPU_003766 | none |
| e05 | SPU_003786 | NA |
| e05 | SPU_003844 | Lv-Dcamkl |
| e05 | SPU_003862 | none |
| e05 | SPU_003921 | NA |
| e05 | SPU_003986 | Lv-Pard6b |
| e05 | SPU_004022 | Lv-Gabbr1_1 |
| e05 | SPU_004144 | Lv-C2orf21L |
| e05 | SPU_004194 | none |
| e05 | SPU_004272 | Lv-Eif2ak |
| e05 | SPU_004273 | Lv-Lct-9 |
| e05 | SPU_004590 | Lv-Glysh |

| | | |
|---|---|---|
| e05 | SPU_004671 | Lv-Tbk1 |
| e05 | SPU_004740 | Lv-Hypp_3125 |
| e05 | SPU_004798 | Lv-Hao1_3 |
| e05 | SPU_004892 | Lv-Oplah |
| e05 | SPU_005041 | Lv-Hypp_437 |
| e05 | SPU_005056 | Lv-Slc24a5 |
| e05 | SPU_005127 | Lv-EchL4 |
| e05 | SPU_005195 | Lv-Slc22a3 |
| e05 | SPU_005204 | none |
| e05 | SPU_005226 | Lv-Galk2_1 |
| e05 | SPU_005295 | Lv-Raph1L |
| e05 | SPU_005348 | Lv-Tex11L |
| e05 | SPU_005402 | Lv-C14orf18L |
| e05 | SPU_005407 | Lv-Adcy9 |
| e05 | SPU_005471 | Lv-Fam45a |
| e05 | SPU_005541 | Lv-Slc7a4_3 |
| e05 | SPU_005648 | Lv-Bhmt-9 |
| e05 | SPU_005787 | Lv-Mlx |
| e05 | SPU_005845 | Lv-AK |
| e05 | SPU_005885 | Lv-Ptpn3/4 |
| e05 | SPU_005935 | Lv-Hypp_2983 |
| e05 | SPU_005938 | Lv-AdamtsL |
| e05 | SPU_005957 | NA |
| e05 | SPU_005959 | Lv-Lrpap1L |
| e05 | SPU_005988 | Lv-Frs3L |
| e05 | SPU_006035 | Lv-Megf6-3 |
| e05 | SPU_006055 | Lv-Srcr47 |
| e05 | SPU_006058 | Lv-Z464 |
| e05 | SPU_006259 | Lv-Hypp_472 |
| e05 | SPU_006318 | Lv-Wipi2 |
| e05 | SPU_006686 | Lv-C6orf70L |
| e05 | SPU_006988 | Lv-Syk/Zap70 |
| e05 | SPU_007129 | Lv-Adi1 |
| e05 | SPU_007210 | Lv-Derp13 |
| e05 | SPU_007409 | Lv-Cyp17L1 |
| e05 | SPU_007697 | none |
| e05 | SPU_007703 | Lv-Hypp_524 |
| e05 | SPU_007830 | none |

| | | |
|---|---|---|
| e05 | SPU_007905 | Lv-Slc7a11_3 |
| e05 | SPU_008273 | Lv-Hypp_1775 |
| e05 | SPU_008353 | Lv-Sim |
| e05 | SPU_008361 | NA |
| e05 | SPU_008414 | Lv-Unc93a_1 |
| e05 | SPU_008503 | Lv-Crocc |
| e05 | SPU_008555 | Lv-Fam134cL |
| e05 | SPU_008853 | Lv-Mib2_2 |
| e05 | SPU_008944 | Lv-Tbc1d19 |
| e05 | SPU_009005 | NA |
| e05 | SPU_009118 | Lv-Cyp2L17 |
| e05 | SPU_009220 | Lv-Srcr184 |
| e05 | SPU_009267 | NA |
| e05 | SPU_009268 | Lv-Nae1 |
| e05 | SPU_009279 | NA |
| e05 | SPU_009599 | Lv-Trpv6L |
| e05 | SPU_009692 | Lv-Cyp2L19 |
| e05 | SPU_009710 | Lv-Hypp_1867 |
| e05 | SPU_009752 | NA |
| e05 | SPU_009818 | Lv-DscamL_1 |
| e05 | SPU_009943 | NA |
| e05 | SPU_009991 | none |
| e05 | SPU_010085 | Lv-Tmem184b |
| e05 | SPU_010089 | none |
| e05 | SPU_010151 | Lv-Doc2Lb |
| e05 | SPU_010374 | Lv-Fn3rch |
| e05 | SPU_010475 | Lv-Lrrc42 |
| e05 | SPU_010555 | Lv-Trpc5_4 |
| e05 | SPU_010618 | Lv-Slco4a1_1 |
| e05 | SPU_010709 | Lv-Hypp_1935 |
| e05 | SPU_010770 | Lv-KifC3L2 |
| e05 | SPU_010775 | Lv-Hao1_6 |
| e05 | SPU_010787 | none |
| e05 | SPU_010876 | Lv-Pkc1 |
| e05 | SPU_010892 | Lv-Akrf_1 |
| e05 | SPU_010958 | Lv-Srd5a1 |
| e05 | SPU_011257 | Lv-A2m |
| e05 | SPU_011274 | Lv-Unc44_295 |

| | | |
|---|---|---|
| e05 | SPU_011278 | Lv-Hypp_1971 |
| e05 | SPU_011348 | Lv-Fxr |
| e05 | SPU_011583 | Lv-Z421 |
| e05 | SPU_011588 | Lv-Spc39L |
| e05 | SPU_011717 | Lv-Hypp_655 |
| e05 | SPU_012003 | Lv-Abcc9e |
| e05 | SPU_012071 | Lv-MifL1 |
| e05 | SPU_012072 | Lv-Napepld |
| e05 | SPU_012076 | Lv-Cca2dA |
| e05 | SPU_012097 | Lv-Sgsh_1 |
| e05 | SPU_012120 | Lv-Adk_2 |
| e05 | SPU_012192 | NA |
| e05 | SPU_012211 | Lv-Tnfrsf1a |
| e05 | SPU_012293 | none |
| e05 | SPU_012702 | Lv-Srr_2 |
| e05 | SPU_012704 | Lv-Slc22a18 |
| e05 | SPU_012712 | Lv-Sco1 |
| e05 | SPU_012798 | none |
| e05 | SPU_012856 | Lv-Syt12 |
| e05 | SPU_012871 | Lv-Kcnd2 |
| e05 | SPU_013013 | none |
| e05 | SPU_013089 | none |
| e05 | SPU_013095 | Lv-Nacha7_1 |
| e05 | SPU_013152 | Lv-Hypp_705 |
| e05 | SPU_013156 | Lv-HdigdcL |
| e05 | SPU_013237 | Lv-Syne1 |
| e05 | SPU_013302 | NA |
| e05 | SPU_013323 | Lv-CadherinL4 |
| e05 | SPU_013358 | Lv-Serrp9 |
| e05 | SPU_013380 | Lv-Srd5a2l |
| e05 | SPU_013563 | none |
| e05 | SPU_013660 | Lv-Npepps |
| e05 | SPU_013784 | Lv-Hypp_728 |
| e05 | SPU_013791 | Lv-Hypp_2109 |
| e05 | SPU_013793 | none |
| e05 | SPU_013869 | Lv-Eya |
| e05 | SPU_013936 | Lv-Paox |
| e05 | SPU_013988 | Lv-Nudt22 |

| | | |
|---|---|---|
| e05 | SPU_014146 | Lv-Hypp_2130 |
| e05 | SPU_014316 | Lv-Syp |
| e05 | SPU_014320 | none |
| e05 | SPU_014381 | Lv-Deaf1 |
| e05 | SPU_014431 | none |
| e05 | SPU_014574 | Lv-Dgkb_1 |
| e05 | SPU_014787 | none |
| e05 | SPU_015308 | none |
| e05 | SPU_015341 | Lv-Apn1 |
| e05 | SPU_015365 | none |
| e05 | SPU_015465 | Lv-Hypp_113 |
| e05 | SPU_015479 | none |
| e05 | SPU_015504 | none |
| e05 | SPU_015551 | Lv-Garnl3 |
| e05 | SPU_015770 | Lv-Prrdh_1 |
| e05 | SPU_015797 | Lv-Hypp_115 |
| e05 | SPU_015842 | Lv-Gpam |
| e05 | SPU_015961 | Lv-Clect/Pan_4 |
| e05 | SPU_016171 | Lv-Sea/Ldla/Cub |
| e05 | SPU_016241 | Lv-Gyg1 |
| e05 | SPU_016359 | Lv-Crocc2 |
| e05 | SPU_016378 | Lv-Tm6sf1 |
| e05 | SPU_016559 | none |
| e05 | SPU_016692 | Lv-Hypp_2268 |
| e05 | SPU_016708 | none |
| e05 | SPU_016717 | Lv-Dhx58 |
| e05 | SPU_016723 | Lv-Ccp/Lnb/7Tm/Gpcr |
| e05 | SPU_016854 | none |
| e05 | SPU_016922 | none |
| e05 | SPU_016948 | Lv-Glyctk1 |
| e05 | SPU_017043 | Lv-Tbc1d2 |
| e05 | SPU_017128 | Lv-Necap1 |
| e05 | SPU_017256 | Lv-Mylip |
| e05 | SPU_017346 | Lv-Ta/Pp2c |
| e05 | SPU_017360 | Lv-Hypp_3050 |
| e05 | SPU_017368 | Lv-Herc4-5 |
| e05 | SPU_017470 | Lv-Klf3/8/12 |
| e05 | SPU_017561 | Lv-Acbd6 |

| | | |
|---|---|---|
| e05 | SPU_017573 | Lv-Cdipt |
| e05 | SPU_017601 | Lv-Cpvl |
| e05 | SPU_017612 | Lv-Pde6d |
| e05 | SPU_017671 | Lv-Gckr |
| e05 | SPU_017752 | Lv-Slc2a13 |
| e05 | SPU_017792 | Lv-Ap4b1 |
| e05 | SPU_018155 | Lv-Vwd/Egff_2 |
| e05 | SPU_018176 | Lv-Lekr1L |
| e05 | SPU_018254 | Lv-LancL |
| e05 | SPU_018563 | Lv-Slc16a14_5 |
| e05 | SPU_018676 | none |
| e05 | SPU_018706 | Lv-Hypp_2400 |
| e05 | SPU_018762 | Lv-Sv2c |
| e05 | SPU_018820 | Lv-Slfnl1 |
| e05 | SPU_018975 | Lv-St6galnac5_2 |
| e05 | SPU_019035 | Lv-Ccdc60L |
| e05 | SPU_019085 | Lv-Rapsn |
| e05 | SPU_019097 | Lv-Hdpx2 |
| e05 | SPU_019121 | Lv-Egfiii_1 |
| e05 | SPU_019125 | Lv-Ttc30b |
| e05 | SPU_019183 | Lv-Mib1L_5 |
| e05 | SPU_019238 | Lv-Sr/Lrr/Gpcr_1 |
| e05 | SPU_019270 | Lv-Trpc5L |
| e05 | SPU_019291 | Lv-Srcr151 |
| e05 | SPU_019332 | Lv-Ppar1 |
| e05 | SPU_019377 | Lv-Hypp_2436 |
| e05 | SPU_019388 | Lv-Slco4a1_12 |
| e05 | SPU_019502 | none |
| e05 | SPU_019503 | Lv-Pla2_1 |
| e05 | SPU_019535 | Lv-Ankrd55 |
| e05 | SPU_019620 | Lv-Qik |
| e05 | SPU_019694 | Lv-Lnb7tmr |
| e05 | SPU_019873 | none |
| e05 | SPU_019982 | Lv-D2hgdh |
| e05 | SPU_020063 | none |
| e05 | SPU_020065 | none |
| e05 | SPU_020100 | Lv-Oach_1 |
| e05 | SPU_020220 | NA |

| | | |
|---|---|---|
| e05 | SPU_020307 | Lv-Unc44_144 |
| e05 | SPU_020317 | Lv-Mfsd7 |
| e05 | SPU_020388 | Lv-PolypL_951 |
| e05 | SPU_020470 | Lv-MylkL |
| e05 | SPU_020564 | Lv-Dehal1 |
| e05 | SPU_020602 | Lv-Raph1L_2 |
| e05 | SPU_020709 | Lv-Mat2a |
| e05 | SPU_020859 | none |
| e05 | SPU_020955 | Lv-TnfrsfL2 |
| e05 | SPU_021073 | Lv-C19orf50L |
| e05 | SPU_021086 | Lv-Cadherin_5 |
| e05 | SPU_021092 | Lv-Chrm3L |
| e05 | SPU_021120 | Lv-Ccdc87 |
| e05 | SPU_021186 | Lv-Mdh |
| e05 | SPU_021209 | Lv-Parvb |
| e05 | SPU_021351 | Lv-Acel_4 |
| e05 | SPU_021462 | Lv-Ece1L |
| e05 | SPU_021470 | Lv-Ids_3 |
| e05 | SPU_021580 | Lv-Dysth1 |
| e05 | SPU_021588 | Lv-Adra1a |
| e05 | SPU_021596 | Lv-Hypp_2566 |
| e05 | SPU_021598 | Lv-Slc16a14_6 |
| e05 | SPU_022002 | Lv-Uros |
| e05 | SPU_022005 | Lv-Rbx2 |
| e05 | SPU_022326 | Lv-Hdhd2 |
| e05 | SPU_022340 | none |
| e05 | SPU_022456 | Lv-My15ph |
| e05 | SPU_022509 | Lv-PolypL_117 |
| e05 | SPU_022576 | none |
| e05 | SPU_023085 | Lv-Chdh |
| e05 | SPU_023121 | Lv-Rgn_2 |
| e05 | SPU_023153 | Lv-Cradd2 |
| e05 | SPU_023247 | Lv-Pgrp1 |
| e05 | SPU_023270 | Lv-Etnk1 |
| e05 | SPU_023393 | Lv-Frag1 |
| e05 | SPU_023460 | Lv-Acsm3 |
| e05 | SPU_023498 | none |
| e05 | SPU_023575 | Lv-Pde5a |

| | | |
|---|---|---|
| e05 | SPU_023638 | Lv-Abca3_1 |
| e05 | SPU_023641 | Lv-Srcr198 |
| e05 | SPU_023662 | Lv-Dhdh_2 |
| e05 | SPU_023792 | Lv-Iunh |
| e05 | SPU_023866 | none |
| e05 | SPU_023972 | Lv-Gpr51_2 |
| e05 | SPU_024137 | Lv-Mppe1 |
| e05 | SPU_024335 | Lv-Egfi_7_1 |
| e05 | SPU_024437 | Lv-Hypp_1115 |
| e05 | SPU_024503 | Lv-Hypp_2737 |
| e05 | SPU_024624 | Lv-Hypp_1118 |
| e05 | SPU_024629 | Lv-Slc5a9_8 |
| e05 | SPU_024642 | none |
| e05 | SPU_024693 | Lv-Nudt19 |
| e05 | SPU_024826 | Lv-Slc16a6L_8 |
| e05 | SPU_025129 | Lv-Rhogef3 |
| e05 | SPU_025245 | none |
| e05 | SPU_025328 | none |
| e05 | SPU_025715 | NA |
| e05 | SPU_025724 | Lv-Fyve1 |
| e05 | SPU_025745 | Lv-Fbxl20L |
| e05 | SPU_025763 | Lv-Muts4 |
| e05 | SPU_025782 | Lv-Acdh |
| e05 | SPU_025882 | Lv-Dapk1_1 |
| e05 | SPU_025922 | NA |
| e05 | SPU_025950 | Lv-Ankar_1 |
| e05 | SPU_026052 | none |
| e05 | SPU_026073 | Lv-Plekhg5 |
| e05 | SPU_026092 | Lv-Egf/Ig/Lnb/7Tm/Gpcr |
| e05 | SPU_026180 | Lv-Tctn2 |
| e05 | SPU_026225 | Lv-Cyp3L4 |
| e05 | SPU_026358 | Lv-Ano10 |
| e05 | SPU_026376 | Lv-CtrhLr |
| e05 | SPU_026377 | Lv-Crhr2L_1 |
| e05 | SPU_026437 | Lv-Ugt2b33_4 |
| e05 | SPU_026555 | Lv-122B |
| e05 | SPU_026567 | NA |
| e05 | SPU_026577 | Lv-Adsda |

| | | |
|---|---|---|
| e05 | SPU_026631 | Lv-C7orf23L |
| e05 | SPU_026689 | Lv-PstkL |
| e05 | SPU_026776 | Lv-Sntb1 |
| e05 | SPU_026807 | Lv-H6pd |
| e05 | SPU_026860 | Lv-Lis1 |
| e05 | SPU_027019 | none |
| e05 | SPU_027304 | Lv-Prodh2 |
| e05 | SPU_027338 | Lv-Hypp_266 |
| e05 | SPU_027604 | Lv-Mmp18/19L7 |
| e05 | SPU_027616 | Lv-Kis |
| e05 | SPU_027690 | Lv-PppL_234 |
| e05 | SPU_027891 | Lv-Sat2_2 |
| e05 | SPU_028082 | Lv-Crocc_4 |
| e05 | SPU_028138 | Lv-Hypp_2936 |
| e05 | SPU_028152 | Lv-Cyp4VL6 |
| e05 | SPU_028211 | Lv-Gabbr2L |
| e05 | SPU_028345 | none |
| e05 | SPU_028351 | Lv-Drd1_3 |
| e05 | SPU_028369 | Lv-Slc16a14L_7 |
| e05 | SPU_028376 | none |
| e05 | SPU_028495 | Lv-Hypp_1231 |
| e05 | SPU_028501 | Lv-Vwc |
| e05 | SPU_028539 | Lv-Clect/kring |
| e05 | SPU_028579 | Lv-Rfc1 |
| e05 | SPU_028617 | Lv-RhoL |
| e05 | SPU_028649 | Lv-Drak |
| e05 | SPU_028730 | NA |
| e05 | SPU_028732 | Lv-C1qL_5 |
| e05 | SPU_028833 | Lv-Cep110l |
| e05 | SPU_028913 | Lv-Myo1b |
| e05 | SPU_028914 | Lv-Megf11 |
| e05 | SPU_030043 | Lv-Naalad2L_18 |
| e05 | SPU_030141 | Lv-IL17rL |
| e06 | SPU_000117 | Lv-Gucy1b3 |
| e06 | SPU_000241 | Lv-Tal1ph |
| e06 | SPU_000247 | Lv-Tppp2/3 |
| e06 | SPU_000471 | Lv-Rdh10_1 |
| e06 | SPU_000498 | Lv-Gpr161L_2 |

| | | |
|---|---|---|
| e06 | SPU_000523 | Lv-Nlr148 |
| e06 | SPU_000564 | Lv-Mib1L_2 |
| e06 | SPU_000588 | none |
| e06 | SPU_000611 | Lv-Plce |
| e06 | SPU_000649 | Lv-Tnfaip3 |
| e06 | SPU_000719 | Lv-Oprk1L |
| e06 | SPU_000730 | Lv-Gpr64L_2 |
| e06 | SPU_000792 | Lv-Rxfp1L |
| e06 | SPU_000856 | Lv-Kcns3 |
| e06 | SPU_000872 | Lv-Galc |
| e06 | SPU_000959 | NA |
| e06 | SPU_000963 | none |
| e06 | SPU_001060 | Lv-Asah1 |
| e06 | SPU_001246 | Lv-Hypp_1318 |
| e06 | SPU_001328 | Lv-ReelerL |
| e06 | SPU_001495 | none |
| e06 | SPU_001561 | Lv-Sqrdl |
| e06 | SPU_001614 | Lv-Fbxl7-2 |
| e06 | SPU_001624 | Lv-Ccb2A |
| e06 | SPU_001846 | Lv-MtoL1_1 |
| e06 | SPU_001876 | Lv-Hypp_1351 |
| e06 | SPU_001935 | Lv-Vcl |
| e06 | SPU_001975 | Lv-Lhx3-4 |
| e06 | SPU_002297 | Lv-Slc45a2_2 |
| e06 | SPU_002344 | Lv-Hypp_1388 |
| e06 | SPU_002393 | Lv-Hypp_1390 |
| e06 | SPU_002401 | none |
| e06 | SPU_002497 | Lv-Hypp_357 |
| e06 | SPU_002499 | Lv-Gucy2gL |
| e06 | SPU_002537 | none |
| e06 | SPU_002585 | Lv-Ephx1 |
| e06 | SPU_002604 | none |
| e06 | SPU_002626 | none |
| e06 | SPU_002642 | Lv-Pm20d1 |
| e06 | SPU_002737 | Lv-Nacha2 |
| e06 | SPU_002926 | Lv-Mb_1 |
| e06 | SPU_003347 | NA |
| e06 | SPU_003499 | Lv-Hypp_386 |

| | | |
|---|---|---|
| e06 | SPU_003577 | Lv-EchnL_5 |
| e06 | SPU_003712 | Lv-Lyplal1 |
| e06 | SPU_003730 | Lv-Protocad15L |
| e06 | SPU_003875 | Lv-Htr1aL_4 |
| e06 | SPU_003917 | none |
| e06 | SPU_004274 | Lv-Lct-10 |
| e06 | SPU_004322 | Lv-Vapa |
| e06 | SPU_004324 | Lv-Ankrd29 |
| e06 | SPU_004406 | Lv-Neuglbn |
| e06 | SPU_004699 | none |
| e06 | SPU_004710 | Lv-Adam/Ts2/3L2 |
| e06 | SPU_004850 | Lv-Clca_1 |
| e06 | SPU_004920 | none |
| e06 | SPU_004949 | Lv-Reck |
| e06 | SPU_005097 | Lv-5ht7 |
| e06 | SPU_005181 | Lv-Endrvt61 |
| e06 | SPU_005265 | Lv-Hat1 |
| e06 | SPU_005406 | Lv-Muc4dL1 |
| e06 | SPU_005513 | none |
| e06 | SPU_005668 | Lv-Cyp2L13 |
| e06 | SPU_005737 | Lv-Vegf2 |
| e06 | SPU_006068 | Lv-Sr/Fu/Igr |
| e06 | SPU_006522 | NA |
| e06 | SPU_006526 | none |
| e06 | SPU_006530 | Lv-Gnbp1/2/3B_1 |
| e06 | SPU_006592 | Lv-Hypp_487 |
| e06 | SPU_006850 | Lv-Myh9/Myh10/Myh11 |
| e06 | SPU_006902 | Lv-Mib2 |
| e06 | SPU_006924 | NA |
| e06 | SPU_007206 | none |
| e06 | SPU_007218 | Lv-Mmr2l |
| e06 | SPU_007280 | Lv-Plscrl_1 |
| e06 | SPU_007412 | Lv-Unc13aL |
| e06 | SPU_007413 | NA |
| e06 | SPU_007537 | Lv-Na/Cae |
| e06 | SPU_007614 | Lv-Actg1 |
| e06 | SPU_007624 | Lv-Ros_1 |
| e06 | SPU_007625 | Lv-Nmur2 |

| | | |
|---|---|---|
| e06 | SPU_007673 | Lv-Rapgef4_1 |
| e06 | SPU_007696 | Lv-Ada |
| e06 | SPU_008462 | Lv-Hypp_548 |
| e06 | SPU_008484 | Lv-SetL |
| e06 | SPU_008595 | Lv-Egfr |
| e06 | SPU_008789 | Lv-Adsr |
| e06 | SPU_008820 | Lv-Hypp_37 |
| e06 | SPU_009097 | Lv-Xkr6_1 |
| e06 | SPU_009122 | Lv-Matn2L |
| e06 | SPU_009217 | Lv-Slc23a2_1 |
| e06 | SPU_009336 | Lv-Slc16a4_2 |
| e06 | SPU_009445 | Lv-SaroxL |
| e06 | SPU_009866 | Lv-Pctp |
| e06 | SPU_009921 | Lv-Hypp_59 |
| e06 | SPU_010067 | Lv-Kcnh8 |
| e06 | SPU_010142 | Lv-Slc16a9L_1 |
| e06 | SPU_010172 | Lv-FAD-dep_OxRdtase |
| e06 | SPU_010275 | Lv-Plcg_2 |
| e06 | SPU_010347 | Lv-Ctr1pk |
| e06 | SPU_010496 | Lv-Leng9L |
| e06 | SPU_010592 | Lv-LacL_6 |
| e06 | SPU_010648 | Lv-RabO1 |
| e06 | SPU_010682 | Lv-Npr1L_1 |
| e06 | SPU_010755 | Lv-PtprN2p |
| e06 | SPU_010948 | Lv-AnL5 |
| e06 | SPU_011009 | Lv-Cyp3L3 |
| e06 | SPU_011056 | Lv-Hypp_1956 |
| e06 | SPU_011197 | Lv-IkB |
| e06 | SPU_011241 | none |
| e06 | SPU_011299 | Lv-Mif1 |
| e06 | SPU_011305 | Lv-Apoa1bp |
| e06 | SPU_011315 | Lv-Ato |
| e06 | SPU_011430 | Lv-Chrm3L_8 |
| e06 | SPU_011464 | none |
| e06 | SPU_011554 | none |
| e06 | SPU_011784 | Lv-Slco2a1 |
| e06 | SPU_011937 | Lv-Ataxia1 |
| e06 | SPU_011980 | Lv-Dmgdh |

| | | |
|---|---|---|
| e06 | SPU_011985 | Lv-Far1 |
| e06 | SPU_012224 | Lv-Unc44_227 |
| e06 | SPU_012550 | Lv-FibB |
| e06 | SPU_012725 | Lv-Mpst |
| e06 | SPU_012727 | Lv-Tst_1 |
| e06 | SPU_013024 | Lv-Surreal/Gpcr5 |
| e06 | SPU_013105 | Lv-Abcg4 |
| e06 | SPU_013114 | Lv-Hypp_2069 |
| e06 | SPU_013178 | Lv-Nr1m3 |
| e06 | SPU_013231 | Lv-Slc7a11_5 |
| e06 | SPU_013254 | Lv-Taui1ph1 |
| e06 | SPU_013374 | Lv-Hypp_2080 |
| e06 | SPU_013404 | none |
| e06 | SPU_013463 | NA |
| e06 | SPU_013485 | Lv-Kchn2L |
| e06 | SPU_013574 | none |
| e06 | SPU_013583 | Lv-Stxbp5 |
| e06 | SPU_013723 | Lv-Neto1 |
| e06 | SPU_013800 | Lv-Fn3_29 |
| e06 | SPU_013990 | Lv-Pink1L |
| e06 | SPU_014038 | none |
| e06 | SPU_014046 | Lv-Sord_1 |
| e06 | SPU_014260 | none |
| e06 | SPU_014263 | Lv-Dish2(Dab2) |
| e06 | SPU_014300 | Lv-Hypp_748 |
| e06 | SPU_014377 | Lv-Hypp_753 |
| e06 | SPU_014573 | NA |
| e06 | SPU_014641 | NA |
| e06 | SPU_014781 | Lv-NotchL_12 |
| e06 | SPU_014876 | Lv-Taut |
| e06 | SPU_015059 | Lv-Slc7a4_4 |
| e06 | SPU_015081 | NA |
| e06 | SPU_015403 | Lv-Anktm1 |
| e06 | SPU_015449 | Lv-Gtlf3b |
| e06 | SPU_015483 | Lv-Glbn |
| e06 | SPU_015544 | Lv-Llgl2 |
| e06 | SPU_015600 | none |
| e06 | SPU_015603 | Lv-ReelerL_2 |

| | | |
|---|---|---|
| e06 | SPU_015636 | Lv-Bai3L |
| e06 | SPU_015678 | Lv-Slc13a2_1 |
| e06 | SPU_015727 | Lv-Mamdc4 |
| e06 | SPU_015836 | Lv-Slc27a6 |
| e06 | SPU_015864 | Lv-Lphn3L_2 |
| e06 | SPU_015989 | Lv-Srcr130 |
| e06 | SPU_016249 | Lv-SommL_1 |
| e06 | SPU_016585 | Lv-Kcnh7 |
| e06 | SPU_016651 | Lv-Dao1 |
| e06 | SPU_016709 | Lv-SacSl_9 |
| e06 | SPU_016752 | none |
| e06 | SPU_016898 | Lv-Neurmdnr |
| e06 | SPU_016983 | Lv-Kif1a |
| e06 | SPU_017000 | none |
| e06 | SPU_017107 | Lv-Slc9a7 |
| e06 | SPU_017139 | Lv-Bicc1 |
| e06 | SPU_017211 | Lv-Glra1_1 |
| e06 | SPU_017303 | Lv-Macf1L |
| e06 | SPU_017321 | Lv-Vwa_1 |
| e06 | SPU_017445 | Lv-ImcvL |
| e06 | SPU_017467 | Lv-Sat2_4 |
| e06 | SPU_017485 | none |
| e06 | SPU_017582 | Lv-Cyp1r2 |
| e06 | SPU_017634 | Lv-Fmn/Abp |
| e06 | SPU_017988 | Lv-Adcy5 |
| e06 | SPU_018078 | Lv-Psmg2 |
| e06 | SPU_018098 | Lv-Unk_26 |
| e06 | SPU_018101 | Lv-Rhop1 |
| e06 | SPU_018125 | Lv-Fbxo32 |
| e06 | SPU_018165 | Lv-Hypp_2373 |
| e06 | SPU_018221 | Lv-Bicc1 |
| e06 | SPU_018340 | Lv-Mfsd2 |
| e06 | SPU_018684 | Lv-BphL |
| e06 | SPU_018853 | Lv-Vmd2l |
| e06 | SPU_019288 | Lv-Treh |
| e06 | SPU_019295 | Lv-Angptl1 |
| e06 | SPU_019541 | none |
| e06 | SPU_019615 | none |

| | | |
|---|---|---|
| e06 | SPU_019708 | Lv-SfrL |
| e06 | SPU_019742 | Lv-Capn9 |
| e06 | SPU_019799 | Lv-Tk6 |
| e06 | SPU_019884 | NA |
| e06 | SPU_019949 | Lv-Accn2_3 |
| e06 | SPU_020218 | none |
| e06 | SPU_020249 | Lv-Brsk |
| e06 | SPU_020281 | Lv-Ptpn1/2 |
| e06 | SPU_020363 | Lv-Enase |
| e06 | SPU_020398 | Lv-Pkx1 |
| e06 | SPU_020411 | none |
| e06 | SPU_020579 | Lv-Slc16a3L_2 |
| e06 | SPU_020673 | Lv-Slc10a2_10 |
| e06 | SPU_020732 | Lv-Hypp_988 |
| e06 | SPU_020780 | none |
| e06 | SPU_020960 | none |
| e06 | SPU_021079 | Lv-Unc44_151 |
| e06 | SPU_021199 | none |
| e06 | SPU_021262 | Lv-Fn3_7 |
| e06 | SPU_021621 | Lv-MyX |
| e06 | SPU_021987 | Lv-Srcr172 |
| e06 | SPU_022101 | NA |
| e06 | SPU_022108 | Lv-Slc39a12_2 |
| e06 | SPU_022175 | Lv-Dab2ip |
| e06 | SPU_022223 | Lv-Clect_59 |
| e06 | SPU_022303 | Lv-Sult1d1 |
| e06 | SPU_022358 | Lv-Hk1_2 |
| e06 | SPU_022674 | NA |
| e06 | SPU_022683 | Lv-Drd1_2 |
| e06 | SPU_022787 | Lv-Nwd1_4 |
| e06 | SPU_022799 | Lv-RabO3a |
| e06 | SPU_022887 | none |
| e06 | SPU_022901 | Lv-Abhd7_2 |
| e06 | SPU_022997 | Lv-Ch25h |
| e06 | SPU_023094 | none |
| e06 | SPU_023113 | none |
| e06 | SPU_023329 | Lv-Dpd1L |
| e06 | SPU_023339 | Lv-Nfia |

| | | |
|---|---|---|
| e06 | SPU_023377 | Lv-Plxna4L_2 |
| e06 | SPU_023535 | Lv-Hypp_2675 |
| e06 | SPU_023875 | NA |
| e06 | SPU_023980 | Lv-C3orf32 |
| e06 | SPU_024015 | Lv-Hypp_2704 |
| e06 | SPU_024024 | NA |
| e06 | SPU_024051 | Lv-Hypp_2705 |
| e06 | SPU_024076 | Lv-PkdccL |
| e06 | SPU_024080 | Lv-Pam_1 |
| e06 | SPU_024186 | Lv-Adrb2L_19 |
| e06 | SPU_024215 | none |
| e06 | SPU_024251 | Lv-Lrr/Igr_18 |
| e06 | SPU_024325 | Lv-Rab7l1 |
| e06 | SPU_024516 | Lv-Impact |
| e06 | SPU_024765 | Lv-Rdh8L_1 |
| e06 | SPU_024845 | Lv-Mett11L_2 |
| e06 | SPU_024996 | Lv-Hypp_1133 |
| e06 | SPU_025046 | Lv-Tmem26L |
| e06 | SPU_025123 | Lv-Alpl_4 |
| e06 | SPU_025315 | Lv-Mglur3 |
| e06 | SPU_025361 | NA |
| e06 | SPU_025378 | Lv-Rhop2 |
| e06 | SPU_025501 | none |
| e06 | SPU_025508 | none |
| e06 | SPU_025623 | Lv-Hypp_3127 |
| e06 | SPU_025829 | Lv-Cyp46L6 |
| e06 | SPU_026077 | Lv-My7 |
| e06 | SPU_026458 | Lv-CcK/GLr |
| e06 | SPU_026589 | Lv-Hypp_2857 |
| e06 | SPU_026805 | none |
| e06 | SPU_026853 | Lv-Hrh3_9 |
| e06 | SPU_026900 | Lv-Hs3st5L_2 |
| e06 | SPU_027005 | Lv-Hypp_1195 |
| e06 | SPU_027078 | Lv-Csad |
| e06 | SPU_027108 | Lv-PolypL_89 |
| e06 | SPU_027281 | Lv-Tpi1 |
| e06 | SPU_027288 | Lv-Srcr219 |
| e06 | SPU_027289 | Lv-Efha1 |

| | | |
|---|---|---|
| e06 | SPU_027316 | none |
| e06 | SPU_027510 | Lv-Npr1_3 |
| e06 | SPU_027631 | NA |
| e06 | SPU_027703 | Lv-Dnase2_2 |
| e06 | SPU_027852 | Lv-Mttp |
| e06 | SPU_027995 | Lv-Slc6a5_4 |
| e06 | SPU_028141 | Lv-Acaa1b_1 |
| e06 | SPU_028455 | Lv-Glur6 |
| e06 | SPU_028475 | none |
| e06 | SPU_028586 | Lv-Fmo5_5 |
| e06 | SPU_028685 | Lv-Vwdp |
| e06 | SPU_028745 | Lv-Unc44_192 |
| e06 | SPU_028920 | Lv-D130067i03riklp |
| e07 | SPU_000054 | Lv-Dus2l |
| e07 | SPU_000245 | Lv-Hypp_297 |
| e07 | SPU_000288 | NA |
| e07 | SPU_000304 | Lv-Vps4h |
| e07 | SPU_000371 | Lv-P300/Cbpaf |
| e07 | SPU_000423 | Lv-Hypp_301 |
| e07 | SPU_000452 | NA |
| e07 | SPU_000511 | none |
| e07 | SPU_000547 | Lv-Aci_2 |
| e07 | SPU_000561 | Lv-Fut11 |
| e07 | SPU_000668 | Lv-Unvn |
| e07 | SPU_000838 | Lv-Tulp3L |
| e07 | SPU_000861 | Lv-Myb |
| e07 | SPU_000875 | Lv-Kif9L2 |
| e07 | SPU_000906 | Lv-EchL2 |
| e07 | SPU_000950 | Lv-Api5 |
| e07 | SPU_001006 | Lv-Zranb2 |
| e07 | SPU_001148 | Lv-Clptm1 |
| e07 | SPU_001310 | Lv-Syf2h |
| e07 | SPU_001367 | Lv-Pwwp2bL |
| e07 | SPU_001436 | Lv-TlrL_10 |
| e07 | SPU_001498 | Lv-Mllt |
| e07 | SPU_001571 | none |
| e07 | SPU_001665 | Lv-Dusp11 |
| e07 | SPU_001668 | Lv-As3mt |

| | | |
|---|---|---|
| e07 | SPU_001751 | Lv-Ctsl |
| e07 | SPU_001762 | Lv-Hypp_338 |
| e07 | SPU_001797 | Lv-MacpfC.2 |
| e07 | SPU_001804 | Lv-Krfp1L |
| e07 | SPU_001963 | Lv-Hypp_1360 |
| e07 | SPU_001971 | Lv-Tlr012 |
| e07 | SPU_002023 | none |
| e07 | SPU_002178 | Lv-Astacin5 |
| e07 | SPU_002219 | Lv-Nefl |
| e07 | SPU_002319 | none |
| e07 | SPU_002647 | NA |
| e07 | SPU_002719 | none |
| e07 | SPU_002761 | Lv-Hypp_1412 |
| e07 | SPU_002830 | Lv-Unc119hb |
| e07 | SPU_002849 | Lv-Gpiasp |
| e07 | SPU_002859 | none |
| e07 | SPU_003046 | NA |
| e07 | SPU_003116 | none |
| e07 | SPU_003164 | none |
| e07 | SPU_003198 | none |
| e07 | SPU_003209 | Lv-Clptm1_1 |
| e07 | SPU_003269 | Lv-Polq |
| e07 | SPU_003285 | Lv-Sea/Egf_2 |
| e07 | SPU_003480 | Lv-Fbox11 |
| e07 | SPU_003517 | none |
| e07 | SPU_003629 | NA |
| e07 | SPU_003709 | Lv-RcE1 |
| e07 | SPU_003744 | Lv-Unk_4 |
| e07 | SPU_003803 | Lv-Ppat |
| e07 | SPU_003831 | Lv-Cramp1L |
| e07 | SPU_003926 | Lv-Lama1 |
| e07 | SPU_003953 | Lv-Leo1L |
| e07 | SPU_003996 | NA |
| e07 | SPU_004008 | Lv-Fha |
| e07 | SPU_004145 | none |
| e07 | SPU_004171 | none |
| e07 | SPU_004217 | Lv-SoxD1 |
| e07 | SPU_004276 | none |

| | | |
|---|---|---|
| e07 | SPU_004509 | Lv-Lmn |
| e07 | SPU_004611 | Lv-Ly/Mam/Zp |
| e07 | SPU_004701 | Lv-Hypp_3083 |
| e07 | SPU_004704 | Lv-Sfxn2 |
| e07 | SPU_004806 | Lv-Z197 |
| e07 | SPU_004807 | Lv-Z32 |
| e07 | SPU_004858 | Lv-Sap18 |
| e07 | SPU_005083 | Lv-Ndst |
| e07 | SPU_005228 | Lv-CalsynteninL1 |
| e07 | SPU_005259 | none |
| e07 | SPU_005267 | none |
| e07 | SPU_005350 | none |
| e07 | SPU_005362 | Lv-Tmem106L |
| e07 | SPU_005544 | Lv-EraL1L |
| e07 | SPU_005654 | Lv-PromL1b |
| e07 | SPU_005664 | Lv-Hypp_1594 |
| e07 | SPU_005869 | Lv-Hypp_465 |
| e07 | SPU_005914 | Lv-Sepx1 |
| e07 | SPU_006103 | none |
| e07 | SPU_006264 | Lv-Srcr49 |
| e07 | SPU_006307 | Lv-Arsi |
| e07 | SPU_006373 | Lv-Hypp_478 |
| e07 | SPU_006378 | Lv-Ppm1e/f |
| e07 | SPU_006534 | Lv-Ebr1_5 |
| e07 | SPU_006564 | none |
| e07 | SPU_006772 | Lv-Clcn7 |
| e07 | SPU_006871 | Lv-Z45 |
| e07 | SPU_007039 | Lv-Echn24 |
| e07 | SPU_007078 | Lv-Topk |
| e07 | SPU_007136 | Lv-Dmbt1L_1 |
| e07 | SPU_007574 | Lv-Hypp_1731 |
| e07 | SPU_007576 | Lv-Clect_17 |
| e07 | SPU_007598 | none |
| e07 | SPU_007607 | NA |
| e07 | SPU_007768 | Lv-Papk |
| e07 | SPU_007833 | Lv-Fads2 |
| e07 | SPU_007872 | Lv-Chst5 |
| e07 | SPU_008293 | NA |

| | | |
|---|---|---|
| e07 | SPU_008501 | Lv-Fbox31 |
| e07 | SPU_008537 | Lv-Dpp9 |
| e07 | SPU_008592 | Lv-Hypp_553 |
| e07 | SPU_008617 | Lv-Glyt_5 |
| e07 | SPU_008654 | Lv-Sstr4L_2 |
| e07 | SPU_008766 | Lv-Kif14Lp |
| e07 | SPU_008878 | NA |
| e07 | SPU_008992 | none |
| e07 | SPU_009082 | Lv-Ugt8 |
| e07 | SPU_009147 | Lv-Siatf_2 |
| e07 | SPU_009155 | Lv-Ctnnb |
| e07 | SPU_009174 | Lv-Clect/Egf/hyalin_2 |
| e07 | SPU_009185 | none |
| e07 | SPU_009196 | Lv-Cnot6L |
| e07 | SPU_009218 | Lv-Ags |
| e07 | SPU_009254 | none |
| e07 | SPU_009259 | Lv-Hypp_43 |
| e07 | SPU_009499 | Lv-Rtel1 |
| e07 | SPU_009646 | Lv-Z223 |
| e07 | SPU_009665 | none |
| e07 | SPU_009675 | Lv-Hypp_1865 |
| e07 | SPU_009756 | Lv-Tsg |
| e07 | SPU_009817 | Lv-Wap/Wap_1 |
| e07 | SPU_009831 | Lv-Sp2 |
| e07 | SPU_009832 | Lv-Sp2_1 |
| e07 | SPU_009838 | Lv-Cbp20 |
| e07 | SPU_009995 | none |
| e07 | SPU_010116 | Lv-DagL |
| e07 | SPU_010126 | Lv-TpaseL_5 |
| e07 | SPU_010246 | Lv-Cyp2L22 |
| e07 | SPU_010343 | Lv-Ddx5_17L |
| e07 | SPU_010426 | Lv-Z128 |
| e07 | SPU_010516 | Lv-Hypp_620 |
| e07 | SPU_010710 | Lv-Mogat1_1 |
| e07 | SPU_010985 | NA |
| e07 | SPU_011178 | Lv-Stt3b_1 |
| e07 | SPU_011194 | Lv-Wmp1 |
| e07 | SPU_011434 | none |

| | | |
|---|---|---|
| e07 | SPU_011585 | Lv-Hypp_652 |
| e07 | SPU_011595 | Lv-Pigg |
| e07 | SPU_011619 | Lv-Hypp_1992 |
| e07 | SPU_011720 | Lv-Z90 |
| e07 | SPU_011815 | Lv-Aif_1 |
| e07 | SPU_011850 | Lv-Hypp_2003 |
| e07 | SPU_011871 | Lv-Chrna1L_1 |
| e07 | SPU_012114 | Lv-C3orf48L |
| e07 | SPU_012136 | Lv-Nedd8 |
| e07 | SPU_012469 | Lv-Elk |
| e07 | SPU_012612 | Lv-Hypp_78 |
| e07 | SPU_012744 | none |
| e07 | SPU_012788 | Lv-Qrich2L |
| e07 | SPU_012834 | Lv-Kcnj4 |
| e07 | SPU_012920 | Lv-Reep2 |
| e07 | SPU_013057 | Lv-TpaseL_7 |
| e07 | SPU_013223 | NA |
| e07 | SPU_013244 | Lv-Top1_1 |
| e07 | SPU_013286 | Lv-Hypp_88 |
| e07 | SPU_013382 | Lv-Hypp_714 |
| e07 | SPU_013439 | Lv-Hypp_91 |
| e07 | SPU_013494 | Lv-Mekk4b |
| e07 | SPU_013543 | Lv-Hypp_2088 |
| e07 | SPU_013642 | Lv-Hcn2 |
| e07 | SPU_013689 | Lv-L3mbt_1 |
| e07 | SPU_013765 | Lv-Glur2 |
| e07 | SPU_013836 | NA |
| e07 | SPU_013891 | none |
| e07 | SPU_014230 | NA |
| e07 | SPU_014290 | Lv-Larp6 |
| e07 | SPU_014329 | Lv-Hypp_2141 |
| e07 | SPU_014332 | Lv-Usf |
| e07 | SPU_014422 | Lv-Egfi_8 |
| e07 | SPU_014489 | Lv-Iws1 |
| e07 | SPU_014772 | NA |
| e07 | SPU_014788 | NA |
| e07 | SPU_014843 | Lv-Cyp1r1 |
| e07 | SPU_015067 | Lv-Slc25a16 |

| | | |
|---|---|---|
| e07 | SPU_015071 | Lv-Z398 |
| e07 | SPU_015114 | NA |
| e07 | SPU_015224 | none |
| e07 | SPU_015271 | Lv-Sclt1 |
| e07 | SPU_015281 | Lv-Mekk4 |
| e07 | SPU_015351 | Lv-Sfxn1 |
| e07 | SPU_015375 | Lv-Bloc1S1 |
| e07 | SPU_015442 | Lv-Cyp2L30 |
| e07 | SPU_015532 | NA |
| e07 | SPU_015648 | none |
| e07 | SPU_015825 | Lv-Mgmt |
| e07 | SPU_015832 | Lv-Sts |
| e07 | SPU_015934 | Lv-Hypp_803 |
| e07 | SPU_015939 | Lv-Ppgb_3 |
| e07 | SPU_016169 | Lv-Pgs1 |
| e07 | SPU_016262 | NA |
| e07 | SPU_016320 | Lv-RtL_41 |
| e07 | SPU_016353 | Lv-Hypp_2258 |
| e07 | SPU_016368 | Lv-Nol8L_1 |
| e07 | SPU_016491 | NA |
| e07 | SPU_016534 | Lv-Hypp_829 |
| e07 | SPU_016574 | Lv-Dhrsx_2 |
| e07 | SPU_016579 | none |
| e07 | SPU_016643 | none |
| e07 | SPU_016713 | Lv-Por |
| e07 | SPU_016730 | Lv-PppL_109 |
| e07 | SPU_016949 | none |
| e07 | SPU_016999 | none |
| e07 | SPU_017039 | Lv-Cadherin_2 |
| e07 | SPU_017046 | none |
| e07 | SPU_017099 | Lv-Crot_1 |
| e07 | SPU_017110 | none |
| e07 | SPU_017172 | Lv-Myst2_1 |
| e07 | SPU_017177 | Lv-C2orf65 |
| e07 | SPU_017264 | none |
| e07 | SPU_017300 | Lv-Kiaa1529L_2 |
| e07 | SPU_017338 | Lv-Ihpk2 |
| e07 | SPU_017679 | NA |

| | | |
|---|---|---|
| e07 | SPU_017690 | Lv-Snx25 |
| e07 | SPU_017736 | Lv-Slc22a13 |
| e07 | SPU_017819 | Lv-C14orf50 |
| e07 | SPU_017840 | Lv-Fam73b_1 |
| e07 | SPU_017957 | Lv-Clca1_5 |
| e07 | SPU_017961 | Lv-Ghmpk |
| e07 | SPU_017990 | Lv-Hexb |
| e07 | SPU_018022 | Lv-Sf1 |
| e07 | SPU_018043 | NA |
| e07 | SPU_018256 | none |
| e07 | SPU_018264 | NA |
| e07 | SPU_018282 | NA |
| e07 | SPU_018329 | Lv-Atox1 |
| e07 | SPU_018397 | Lv-Lkap |
| e07 | SPU_018473 | NA |
| e07 | SPU_018652 | Lv-Hypp_3039 |
| e07 | SPU_018707 | Lv-Hypp_2401 |
| e07 | SPU_018887 | Lv-Adam/TsL4 |
| e07 | SPU_018889 | none |
| e07 | SPU_019107 | none |
| e07 | SPU_019223 | none |
| e07 | SPU_019276 | Lv-Xrn2_1 |
| e07 | SPU_019289 | Lv-Hypp_2430 |
| e07 | SPU_019380 | Lv-Ccdc33 |
| e07 | SPU_019392 | Lv-Cdh1 |
| e07 | SPU_019404 | none |
| e07 | SPU_019407 | Lv-Z101 |
| e07 | SPU_019440 | Lv-Del |
| e07 | SPU_019443 | NA |
| e07 | SPU_019606 | none |
| e07 | SPU_019720 | Lv-Unc44_324 |
| e07 | SPU_019913 | Lv-Hypp_951 |
| e07 | SPU_019987 | NA |
| e07 | SPU_020094 | Lv-Chst5L |
| e07 | SPU_020096 | Lv-Chst4 |
| e07 | SPU_020097 | Lv-C1orf173L |
| e07 | SPU_020215 | Lv-PppL_213 |
| e07 | SPU_020222 | Lv-Ptar1 |

| | | |
|---|---|---|
| e07 | SPU_020267 | NA |
| e07 | SPU_020333 | NA |
| e07 | SPU_020588 | none |
| e07 | SPU_020645 | Lv-Endrvt1271 |
| e07 | SPU_020784 | none |
| e07 | SPU_020830 | Lv-Ss18L |
| e07 | SPU_020904 | Lv-PromL1a |
| e07 | SPU_020907 | Lv-Rassf2 |
| e07 | SPU_021207 | Lv-Pde8A |
| e07 | SPU_021383 | Lv-Z56 |
| e07 | SPU_021397 | Lv-EraL1 |
| e07 | SPU_021491 | Lv-Arxl2 |
| e07 | SPU_021547 | Lv-Sfmbt2 |
| e07 | SPU_021604 | NA |
| e07 | SPU_021627 | none |
| e07 | SPU_021670 | Lv-Atub11 |
| e07 | SPU_021703 | Lv-Polr2b |
| e07 | SPU_021887 | none |
| e07 | SPU_022027 | Lv-Hypp_2594 |
| e07 | SPU_022045 | Lv-Slc36a1 |
| e07 | SPU_022137 | Lv-CubnL_4 |
| e07 | SPU_022142 | Lv-Hypp_2599 |
| e07 | SPU_022284 | Lv-Lkb1/Xeek1 |
| e07 | SPU_022475 | Lv-Prelid2 |
| e07 | SPU_022629 | Lv-Lyk5 |
| e07 | SPU_022670 | Lv-CdkL5-1 |
| e07 | SPU_022701 | Lv-Znf/Cchct/Rnad_dnap/I/Cat |
| e07 | SPU_022737 | Lv-Hmbox1 |
| e07 | SPU_022922 | Lv-Rbm22 |
| e07 | SPU_022968 | Lv-Dcdsh_4 |
| e07 | SPU_022988 | Lv-Tcp1 |
| e07 | SPU_023057 | Lv-Chst9_3 |
| e07 | SPU_023063 | Lv-Hypp_1068 |
| e07 | SPU_023122 | none |
| e07 | SPU_023214 | NA |
| e07 | SPU_023218 | Lv-Hypp_2658 |
| e07 | SPU_023260 | none |
| e07 | SPU_023283 | Lv-Colf_10 |

| | | |
|---|---|---|
| e07 | SPU_023289 | Lv-Hypp_3000 |
| e07 | SPU_023298 | NA |
| e07 | SPU_023338 | NA |
| e07 | SPU_023355 | none |
| e07 | SPU_023371 | Lv-Rgs22L_1 |
| e07 | SPU_023417 | Lv-EhH1_29 |
| e07 | SPU_023587 | NA |
| e07 | SPU_023607 | Lv-Clect/Egf_15 |
| e07 | SPU_023710 | Lv-RtL_5 |
| e07 | SPU_023848 | Lv-Hypp_2690 |
| e07 | SPU_023950 | Lv-Gtf2ird2 |
| e07 | SPU_023975 | Lv-Gulp1 |
| e07 | SPU_024217 | Lv-Ccp/Clect/Zp |
| e07 | SPU_024466 | Lv-Hypp_215 |
| e07 | SPU_024806 | Lv-Chst11_13 |
| e07 | SPU_024834 | Lv-Mamdc4-12 |
| e07 | SPU_025008 | Lv-Hypp_2763 |
| e07 | SPU_025115 | Lv-Wdr45 |
| e07 | SPU_025187 | Lv-GagpL_5 |
| e07 | SPU_025192 | Lv-CholdhL |
| e07 | SPU_025207 | NA |
| e07 | SPU_025208 | Lv-Odf2 |
| e07 | SPU_025309 | Lv-Supt5h_1 |
| e07 | SPU_025405 | Lv-Ubqln1 |
| e07 | SPU_025453 | Lv-Gpr27L |
| e07 | SPU_025499 | Lv-Hypp_1150 |
| e07 | SPU_025541 | Lv-Smc6-2 |
| e07 | SPU_025634 | Lv-Eif4e3 |
| e07 | SPU_025845 | Lv-Ubxn11 |
| e07 | SPU_026039 | Lv-Lama5L |
| e07 | SPU_026048 | none |
| e07 | SPU_026223 | Lv-ReelerL_6 |
| e07 | SPU_026255 | Lv-Uqub |
| e07 | SPU_026362 | Lv-Sh2d5 |
| e07 | SPU_026379 | NA |
| e07 | SPU_026434 | NA |
| e07 | SPU_026663 | Lv-Tep1L_6 |
| e07 | SPU_026806 | Lv-Sf3a1_1 |

| | | |
|---|---|---|
| e07 | SPU_026813 | NA |
| e07 | SPU_026879 | Lv-C20orf86L |
| e07 | SPU_026898 | Lv-Glutamyl/tRNAsynth |
| e07 | SPU_026915 | Lv-Hmcn1L_3 |
| e07 | SPU_026994 | NA |
| e07 | SPU_027007 | none |
| e07 | SPU_027038 | Lv-Catsperg |
| e07 | SPU_027307 | Lv-Hypp_1199 |
| e07 | SPU_027353 | none |
| e07 | SPU_027547 | Lv-Tbc1d16 |
| e07 | SPU_027592 | Lv-Trngltam |
| e07 | SPU_027613 | Lv-Cyp17L2 |
| e07 | SPU_027630 | none |
| e07 | SPU_027778 | Lv-Tropmph |
| e07 | SPU_027801 | Lv-EhH4_78 |
| e07 | SPU_027895 | Lv-Sf3a1_2 |
| e07 | SPU_027939 | Lv-Atg4cL |
| e07 | SPU_028136 | Lv-Hypp_2935 |
| e07 | SPU_028146 | Lv-Slc15a4_2 |
| e07 | SPU_028184 | Lv-SmocL |
| e07 | SPU_028275 | Lv-Ost |
| e07 | SPU_028323 | NA |
| e07 | SPU_028359 | Lv-Nsda |
| e07 | SPU_028419 | Lv-Hs3st5L_3 |
| e07 | SPU_028479 | Lv-Tel |
| e07 | SPU_028520 | NA |
| e07 | SPU_028929 | Lv-EhH2a_69 |
| e07 | SPU_030132 | Lv-TlrP76 |
| e08 | SPU_000252 | Lv-Med1 |
| e08 | SPU_000260 | Lv-PbL1 |
| e08 | SPU_000282 | Lv-Cry1/2L |
| e08 | SPU_000334 | Lv-Elovl4_1 |
| e08 | SPU_000345 | NA |
| e08 | SPU_000383 | Lv-Man1b1 |
| e08 | SPU_000664 | none |
| e08 | SPU_000687 | Lv-Ambra1 |
| e08 | SPU_000793 | Lv-Hmg2l1 |
| e08 | SPU_000818 | Lv-GakL_1 |

| | | |
|---|---|---|
| e08 | SPU_000847 | Lv-Spnsh1 |
| e08 | SPU_000878 | Lv-Hypp_1289 |
| e08 | SPU_000902 | Lv-Caprin1L |
| e08 | SPU_000964 | Lv-Bbs2 |
| e08 | SPU_000988 | Lv-Pigc |
| e08 | SPU_001168 | Lv-Alox12b-2 |
| e08 | SPU_001178 | Lv-Sp2mpL |
| e08 | SPU_001186 | Lv-Wdr42c |
| e08 | SPU_001217 | Lv-Btbd5_1 |
| e08 | SPU_001244 | Lv-M_transf |
| e08 | SPU_001252 | Lv-Kiaa1109L |
| e08 | SPU_001283 | none |
| e08 | SPU_001307 | Lv-Cdc5L |
| e08 | SPU_001325 | none |
| e08 | SPU_001541 | Lv-Bud31_1 |
| e08 | SPU_001582 | none |
| e08 | SPU_001596 | Lv-Atp11 |
| e08 | SPU_001643 | Lv-C20orf152L_1 |
| e08 | SPU_001680 | Lv-Hypp_3116 |
| e08 | SPU_001709 | NA |
| e08 | SPU_001718 | Lv-Cfdp1 |
| e08 | SPU_001742 | Lv-Zmynd8 |
| e08 | SPU_001890 | Lv-Kif14L2 |
| e08 | SPU_001900 | NA |
| e08 | SPU_001909 | Lv-Lancl3 |
| e08 | SPU_001937 | Lv-Z19 |
| e08 | SPU_001954 | Lv-Tor1 |
| e08 | SPU_002063 | NA |
| e08 | SPU_002065 | Lv-C10orf97L_1 |
| e08 | SPU_002114 | Lv-Notch1 |
| e08 | SPU_002197 | none |
| e08 | SPU_002286 | Lv-Uqcc |
| e08 | SPU_002290 | none |
| e08 | SPU_002320 | Lv-FoxN1/4 |
| e08 | SPU_002326 | Lv-Phc2L_1 |
| e08 | SPU_002443 | Lv-Eif3s4 |
| e08 | SPU_002445 | none |
| e08 | SPU_002525 | Lv-Spryd3 |

| | | |
|---|---|---|
| e08 | SPU_002532 | NA |
| e08 | SPU_002545 | NA |
| e08 | SPU_002638 | Lv-Rap2 |
| e08 | SPU_002681 | Lv-Sfrs6 |
| e08 | SPU_002684 | Lv-Sv2b |
| e08 | SPU_002879 | Lv-Endrvt_3 |
| e08 | SPU_002946 | Lv-Hypp_371 |
| e08 | SPU_003083 | Lv-Bcs1L |
| e08 | SPU_003249 | Lv-Kiaa1076l |
| e08 | SPU_003262 | Lv-Faim |
| e08 | SPU_003288 | Lv-Hypp_1440 |
| e08 | SPU_003358 | Lv-Smg6h |
| e08 | SPU_003399 | Lv-Hypp_3088 |
| e08 | SPU_003440 | Lv-Hypp_382 |
| e08 | SPU_003478 | Lv-Sf3B2 |
| e08 | SPU_003481 | none |
| e08 | SPU_003485 | Lv-MPRL11 |
| e08 | SPU_003519 | NA |
| e08 | SPU_003530 | none |
| e08 | SPU_003682 | Lv-Hypp_1472 |
| e08 | SPU_003743 | Lv-Prp19 |
| e08 | SPU_004197 | Lv-PppL_176 |
| e08 | SPU_004224 | Lv-Kif22 |
| e08 | SPU_004381 | Lv-Bnip1 |
| e08 | SPU_004392 | Lv-RtL_14 |
| e08 | SPU_004407 | Lv-Slco5a1 |
| e08 | SPU_004445 | Lv-Frg1_1 |
| e08 | SPU_004462 | none |
| e08 | SPU_004488 | Lv-Hypp_1529 |
| e08 | SPU_004561 | Lv-Pgm1 |
| e08 | SPU_004566 | none |
| e08 | SPU_004621 | Lv-Lfng |
| e08 | SPU_004672 | Lv-Hypp_423 |
| e08 | SPU_004773 | none |
| e08 | SPU_004827 | Lv-Sh3md2Lp |
| e08 | SPU_004844 | Lv-Lztf1_1 |
| e08 | SPU_004860 | Lv-Cwc_Cwf_15 |
| e08 | SPU_005000 | Lv-Srcr40 |

| | | |
|---|---|---|
| e08 | SPU_005054 | Lv-Sgsh |
| e08 | SPU_005135 | Lv-Sea_1 |
| e08 | SPU_005219 | Lv-TpaseL_34 |
| e08 | SPU_005236 | none |
| e08 | SPU_005277 | Lv-Nek2 |
| e08 | SPU_005306 | none |
| e08 | SPU_005416 | none |
| e08 | SPU_005443 | Lv-Tatdn3 |
| e08 | SPU_005490 | Lv-Hypp_1584 |
| e08 | SPU_005509 | Lv-Kif3AL |
| e08 | SPU_005537 | Lv-Plod2 |
| e08 | SPU_005567 | Lv-Amdhd2 |
| e08 | SPU_005630 | Lv-RtL_66 |
| e08 | SPU_005687 | Lv-Pcid2 |
| e08 | SPU_005689 | none |
| e08 | SPU_005710 | Lv-Caprin1 |
| e08 | SPU_005721 | Lv-Echn23 |
| e08 | SPU_005840 | Lv-Tep1L_5 |
| e08 | SPU_005868 | Lv-Hp_Loc200403L |
| e08 | SPU_005911 | Lv-Hypp_467 |
| e08 | SPU_005916 | NA |
| e08 | SPU_006039 | Lv-Erbb2ip |
| e08 | SPU_006043 | none |
| e08 | SPU_006057 | Lv-Zc3h12d |
| e08 | SPU_006064 | Lv-Z79 |
| e08 | SPU_006093 | Lv-Ubtf |
| e08 | SPU_006119 | Lv-Pigk |
| e08 | SPU_006280 | Lv-Hypp_473 |
| e08 | SPU_006294 | Lv-Polg |
| e08 | SPU_006308 | Lv-Atp6v1c1 |
| e08 | SPU_006322 | none |
| e08 | SPU_006548 | none |
| e08 | SPU_006680 | Lv-Sel1L |
| e08 | SPU_006684 | none |
| e08 | SPU_006685 | Lv-Mipol1L |
| e08 | SPU_006720 | Lv-Brd |
| e08 | SPU_006726 | Lv-Hypp_492 |
| e08 | SPU_006749 | Lv-Mett11L |

| | | |
|---|---|---|
| e08 | SPU_006768 | Lv-Polb |
| e08 | SPU_006826 | Lv-Arl3 |
| e08 | SPU_006847 | Lv-Pum |
| e08 | SPU_006873 | Lv-Fam81a |
| e08 | SPU_006979 | Lv-PppL_188 |
| e08 | SPU_007033 | Lv-Akrf |
| e08 | SPU_007060 | none |
| e08 | SPU_007145 | Lv-Hypp_507 |
| e08 | SPU_007213 | NA |
| e08 | SPU_007268 | none |
| e08 | SPU_007341 | Lv-Caspr/C-term |
| e08 | SPU_007410 | Lv-Sybl1 |
| e08 | SPU_007482 | Lv-Hypp_25 |
| e08 | SPU_007589 | Lv-C14orf104L |
| e08 | SPU_007618 | Lv-Srcr55 |
| e08 | SPU_007722 | Lv-Cyfip2 |
| e08 | SPU_007726 | Lv-Clca3_1 |
| e08 | SPU_007823 | none |
| e08 | SPU_007836 | NA |
| e08 | SPU_007837 | none |
| e08 | SPU_007848 | Lv-PppL_250 |
| e08 | SPU_007911 | Lv-Adam15L1 |
| e08 | SPU_007933 | Lv-CycF_1 |
| e08 | SPU_007979 | Lv-Gata2a |
| e08 | SPU_008001 | Lv-Clect_69 |
| e08 | SPU_008042 | Lv-Higd2A |
| e08 | SPU_008101 | Lv-Fam78a |
| e08 | SPU_008134 | none |
| e08 | SPU_008158 | Lv-Pbf6L |
| e08 | SPU_008165 | Lv-RtL_17 |
| e08 | SPU_008206 | Lv-Tarbp2L |
| e08 | SPU_008252 | Lv-Cox4L |
| e08 | SPU_008272 | none |
| e08 | SPU_008312 | Lv-Erh |
| e08 | SPU_008319 | none |
| e08 | SPU_008334 | Lv-Hypp_543 |
| e08 | SPU_008441 | Lv-Hps3L |
| e08 | SPU_008448 | Lv-Mgc16943L |

| | | |
|---|---|---|
| e08 | SPU_008480 | Lv-Pgbd4L_2 |
| e08 | SPU_008517 | none |
| e08 | SPU_008573 | Lv-Hypp_1793 |
| e08 | SPU_008648 | Lv-Dmrt2 |
| e08 | SPU_008682 | Lv-C1orf77L |
| e08 | SPU_008734 | NA |
| e08 | SPU_008780 | Lv-Ttc14L |
| e08 | SPU_008835 | Lv-Hypp_562 |
| e08 | SPU_008866 | Lv-Timp4b |
| e08 | SPU_008875 | Lv-Gcnt2_3 |
| e08 | SPU_008892 | Lv-Rngtt |
| e08 | SPU_008895 | Lv-Ptp4a |
| e08 | SPU_008906 | Lv-Rad54 |
| e08 | SPU_008914 | Lv-Ctr9 |
| e08 | SPU_008919 | Lv-Sfrs3 |
| e08 | SPU_008979 | Lv-P53L |
| e08 | SPU_009233 | Lv-Pcif1 |
| e08 | SPU_009241 | Lv-Cuta |
| e08 | SPU_009245 | none |
| e08 | SPU_009305 | Lv-Pmvk |
| e08 | SPU_009389 | Lv-Cg15216pa |
| e08 | SPU_009394 | Lv-Ints7 |
| e08 | SPU_009413 | Lv-Nxf |
| e08 | SPU_009431 | Lv-Hypp_45 |
| e08 | SPU_009455 | Lv-C1orf102L |
| e08 | SPU_009542 | Lv-RtL_22 |
| e08 | SPU_009551 | Lv-Hypp_1855 |
| e08 | SPU_009567 | Lv-Hypp_1856 |
| e08 | SPU_009568 | Lv-Hypp_1857 |
| e08 | SPU_009628 | Lv-Slc5a7_5 |
| e08 | SPU_009667 | Lv-Hypp_53 |
| e08 | SPU_009802 | Lv-Znf862 |
| e08 | SPU_009837 | none |
| e08 | SPU_009840 | NA |
| e08 | SPU_009846 | Lv-Lamb2f |
| e08 | SPU_009873 | Lv-Hypp_591 |
| e08 | SPU_009908 | Lv-Rag1l5 |
| e08 | SPU_009986 | Lv-Hypp_600 |

| | | |
|---|---|---|
| e08 | SPU_009996 | Lv-Kiaa1161L |
| e08 | SPU_010022 | Lv-Lrp1b |
| e08 | SPU_010059 | Lv-Mdm1L |
| e08 | SPU_010109 | Lv-Zdhhc2 |
| e08 | SPU_010157 | Lv-Tm9sf2_1 |
| e08 | SPU_010175 | none |
| e08 | SPU_010190 | none |
| e08 | SPU_010197 | Lv-ZfrL |
| e08 | SPU_010240 | Lv-Srcr87 |
| e08 | SPU_010256 | Lv-Endrvt75 |
| e08 | SPU_010295 | Lv-Yyi |
| e08 | SPU_010311 | Lv-Nifun |
| e08 | SPU_010393 | Lv-Hypp_67 |
| e08 | SPU_010415 | none |
| e08 | SPU_010420 | Lv-Nusap1L |
| e08 | SPU_010424 | Lv-Otx |
| e08 | SPU_010472 | Lv-Ccp/Sea |
| e08 | SPU_010521 | Lv-Polq_1 |
| e08 | SPU_010529 | Lv-Ttf1 |
| e08 | SPU_010596 | Lv-Cwc22 |
| e08 | SPU_010856 | Lv-Bub3h |
| e08 | SPU_010894 | Lv-Alox5_5 |
| e08 | SPU_010937 | Lv-C17orf101 |
| e08 | SPU_010981 | Lv-Alox-4 |
| e08 | SPU_010986 | Lv-Klhl7L |
| e08 | SPU_011018 | Lv-Polh |
| e08 | SPU_011168 | Lv-Unk_74 |
| e08 | SPU_011183 | Lv-Hypp_1967 |
| e08 | SPU_011266 | Lv-Ythdc1 |
| e08 | SPU_011354 | none |
| e08 | SPU_011387 | Lv-Hypp_1978 |
| e08 | SPU_011426 | Lv-ExonepL |
| e08 | SPU_011427 | none |
| e08 | SPU_011458 | none |
| e08 | SPU_011483 | Lv-Sf3b2L |
| e08 | SPU_011505 | Lv-Pgbd4 |
| e08 | SPU_011542 | Lv-Glipr1l1 |
| e08 | SPU_011611 | Lv-Z134 |

| | | |
|---|---|---|
| e08 | SPU_011628 | Lv-Hypp_1993 |
| e08 | SPU_011646 | Lv-TpaseL_6 |
| e08 | SPU_011659 | none |
| e08 | SPU_011683 | Lv-Dynlc2-4c |
| e08 | SPU_011687 | Lv-Z258 |
| e08 | SPU_011711 | Lv-Bmpr2_1 |
| e08 | SPU_011901 | Lv-Znf474 |
| e08 | SPU_011905 | Lv-Hypp_664 |
| e08 | SPU_011956 | Lv-Slc26a11 |
| e08 | SPU_011984 | Lv-Trove2 |
| e08 | SPU_011991 | NA |
| e08 | SPU_011993 | Lv-Sf3a2 |
| e08 | SPU_012131 | none |
| e08 | SPU_012194 | Lv-Naalad2L_4 |
| e08 | SPU_012202 | Lv-Xpo1h |
| e08 | SPU_012208 | none |
| e08 | SPU_012311 | Lv-Hypp_2026 |
| e08 | SPU_012471 | Lv-Dhfr |
| e08 | SPU_012492 | Lv-PolypL_4 |
| e08 | SPU_012493 | none |
| e08 | SPU_012576 | none |
| e08 | SPU_012579 | Lv-Adpgk |
| e08 | SPU_012691 | Lv-Cep290_1 |
| e08 | SPU_012728 | Lv-Wdr16-2 |
| e08 | SPU_012816 | Lv-Supt5h |
| e08 | SPU_012851 | Lv-Exos6 |
| e08 | SPU_012890 | Lv-Hypp_2054 |
| e08 | SPU_012891 | none |
| e08 | SPU_012903 | NA |
| e08 | SPU_012930 | Lv-Prpf8_1 |
| e08 | SPU_012969 | none |
| e08 | SPU_013008 | Lv-Hypp_2064 |
| e08 | SPU_013043 | Lv-Ctr1aL |
| e08 | SPU_013048 | none |
| e08 | SPU_013134 | Lv-Nr2C |
| e08 | SPU_013166 | Lv-Slc27a2 |
| e08 | SPU_013169 | Lv-Tecp5 |
| e08 | SPU_013170 | Lv-Tecp4 |

| | | |
|---|---|---|
| e08 | SPU_013172 | Lv-Fn3/Egf_2 |
| e08 | SPU_013209 | Lv-Otcba |
| e08 | SPU_013238 | NA |
| e08 | SPU_013298 | Lv-Sds |
| e08 | SPU_013341 | Lv-Hypp_711 |
| e08 | SPU_013391 | Lv-Endrvt85 |
| e08 | SPU_013416 | Lv-Ccdc47 |
| e08 | SPU_013493 | none |
| e08 | SPU_013565 | Lv-Gcs1 |
| e08 | SPU_013576 | Lv-Z96 |
| e08 | SPU_013601 | Lv-Gm2a |
| e08 | SPU_013638 | Lv-RbmX2L |
| e08 | SPU_013656 | Lv-Cachd1 |
| e08 | SPU_013713 | Lv-Hypp_2099 |
| e08 | SPU_013794 | Lv-Rab32l |
| e08 | SPU_013795 | Lv-Harbi1 |
| e08 | SPU_013845 | Lv-PppL_263 |
| e08 | SPU_013851 | Lv-Nudt13 |
| e08 | SPU_013855 | none |
| e08 | SPU_013942 | Lv-Slc7a7 |
| e08 | SPU_014019 | Lv-Ddhd1 |
| e08 | SPU_014072 | Lv-Hypp_97 |
| e08 | SPU_014145 | none |
| e08 | SPU_014403 | Lv-Amt1 |
| e08 | SPU_014482 | none |
| e08 | SPU_014509 | Lv-CopineL_1 |
| e08 | SPU_014557 | none |
| e08 | SPU_014580 | Lv-GrpE |
| e08 | SPU_014590 | Lv-Liph |
| e08 | SPU_014597 | none |
| e08 | SPU_014643 | Lv-Z372 |
| e08 | SPU_014653 | Lv-Blmh |
| e08 | SPU_014718 | Lv-Slc30a1 |
| e08 | SPU_014728 | Lv-Ctnna2L |
| e08 | SPU_014793 | Lv-Spalt |
| e08 | SPU_014812 | NA |
| e08 | SPU_014821 | Lv-Adck1 |
| e08 | SPU_014832 | Lv-Cpsf5 |

| | | |
|---|---|---|
| e08 | SPU_014925 | Lv-Klhl12L |
| e08 | SPU_014962 | Lv-Hecw2 |
| e08 | SPU_014995 | Lv-Tnpo1 |
| e08 | SPU_015041 | Lv-Rmnd5ha |
| e08 | SPU_015055 | none |
| e08 | SPU_015094 | none |
| e08 | SPU_015123 | Lv-Srcr123 |
| e08 | SPU_015177 | Lv-Prr11 |
| e08 | SPU_015180 | Lv-Sbnh1 |
| e08 | SPU_015195 | Lv-Gcc2 |
| e08 | SPU_015230 | Lv-Mfap1 |
| e08 | SPU_015249 | Lv-Ubtd2 |
| e08 | SPU_015358 | Lv-Egr |
| e08 | SPU_015377 | Lv-Aci_3 |
| e08 | SPU_015388 | Lv-Wdr33-2 |
| e08 | SPU_015414 | Lv-Cenpi |
| e08 | SPU_015430 | NA |
| e08 | SPU_015445 | Lv-Ugdh |
| e08 | SPU_015489 | Lv-Nit1 |
| e08 | SPU_015574 | Lv-Hypp_7899 |
| e08 | SPU_015575 | Lv-Gucy1 |
| e08 | SPU_015614 | NA |
| e08 | SPU_015647 | none |
| e08 | SPU_015652 | Lv-Sara/Ny/SAR41 |
| e08 | SPU_015676 | Lv-Hnrpa |
| e08 | SPU_015736 | Lv-Cpsf6L |
| e08 | SPU_015812 | Lv-HyalinL |
| e08 | SPU_015913 | none |
| e08 | SPU_015932 | Lv-QtrtD1 |
| e08 | SPU_015953 | Lv-Pnlip |
| e08 | SPU_015971 | Lv-Hypp_806 |
| e08 | SPU_015972 | NA |
| e08 | SPU_016022 | Lv-Hypp_2243 |
| e08 | SPU_016069 | Lv-Fbxl13 |
| e08 | SPU_016087 | Lv-LhH1a |
| e08 | SPU_016336 | Lv-Hypp_2257 |
| e08 | SPU_016521 | none |
| e08 | SPU_016588 | Lv-Phf8 |

| | | |
|---|---|---|
| e08 | SPU_016600 | NA |
| e08 | SPU_016691 | none |
| e08 | SPU_016696 | Lv-RtL_43 |
| e08 | SPU_016748 | Lv-Cdc2l1 |
| e08 | SPU_016772 | Lv-Clect_110 |
| e08 | SPU_016831 | Lv-Shmt2 |
| e08 | SPU_016859 | Lv-Hypp_846 |
| e08 | SPU_016921 | Lv-Nlr49 |
| e08 | SPU_017047 | none |
| e08 | SPU_017079 | Lv-Hypp_2302 |
| e08 | SPU_017100 | Lv-Ttc19 |
| e08 | SPU_017113 | Lv-Unc44_122 |
| e08 | SPU_017118 | Lv-Endrvt33 |
| e08 | SPU_017165 | Lv-CenpnL |
| e08 | SPU_017286 | none |
| e08 | SPU_017405 | Lv-Fibcd1_1 |
| e08 | SPU_017508 | Lv-Snx4 |
| e08 | SPU_017532 | Lv-Hypp_135 |
| e08 | SPU_017684 | Lv-Acmsd |
| e08 | SPU_017725 | Lv-P3a2 |
| e08 | SPU_017786 | Lv-L3mbtl3 |
| e08 | SPU_017874 | Lv-Cript |
| e08 | SPU_017942 | Lv-Wbp11 |
| e08 | SPU_018053 | Lv-Hypp_3043 |
| e08 | SPU_018068 | Lv-Wdr67 |
| e08 | SPU_018108 | Lv-TwistnbL |
| e08 | SPU_018157 | Lv-Zbed1-4 |
| e08 | SPU_018218 | Lv-Myef2 |
| e08 | SPU_018220 | Lv-MsrL |
| e08 | SPU_018348 | Lv-CasprL |
| e08 | SPU_018351 | Lv-PaxB |
| e08 | SPU_018363 | Lv-Mdm1L_1 |
| e08 | SPU_018367 | Lv-Bxdc2 |
| e08 | SPU_018459 | Lv-Hypp_2387 |
| e08 | SPU_018460 | Lv-Endrvt98 |
| e08 | SPU_018497 | none |
| e08 | SPU_018670 | Lv-EhH4_54 |
| e08 | SPU_018799 | Lv-Adrb1L_6 |

| | | |
|---|---|---|
| e08 | SPU_018864 | Lv-Zbed1_1 |
| e08 | SPU_019044 | Lv-Hypp_2417 |
| e08 | SPU_019095 | Lv-Slc25a28 |
| e08 | SPU_019205 | Lv-XpapdL |
| e08 | SPU_019231 | Lv-Cub/Egf/Lnb/7Tm/Gpcr_2 |
| e08 | SPU_019257 | Lv-Slc25a14 |
| e08 | SPU_019314 | NA |
| e08 | SPU_019389 | Lv-Ago |
| e08 | SPU_019395 | Lv-Hypp_2437 |
| e08 | SPU_019429 | none |
| e08 | SPU_019459 | Lv-Chrom1 |
| e08 | SPU_019555 | Lv-Srrm1 |
| e08 | SPU_019600 | NA |
| e08 | SPU_019625 | none |
| e08 | SPU_019640 | Lv-Glce |
| e08 | SPU_019674 | Lv-Hypp_2459 |
| e08 | SPU_019682 | Lv-Hypp_2460 |
| e08 | SPU_019692 | Lv-Ppt2 |
| e08 | SPU_019785 | Lv-Cr1L |
| e08 | SPU_019951 | Lv-Fbw7 |
| e08 | SPU_019953 | Lv-Cep290 |
| e08 | SPU_019976 | Lv-Naalad2L_5 |
| e08 | SPU_020019 | Lv-Ahsah1 |
| e08 | SPU_020020 | Lv-RigIL3 |
| e08 | SPU_020117 | Lv-Higd1A_1 |
| e08 | SPU_020118 | Lv-Slc5a12_5 |
| e08 | SPU_020186 | NA |
| e08 | SPU_020191 | Lv-Bud31 |
| e08 | SPU_020427 | Lv-Tm9sf2_2 |
| e08 | SPU_020495 | Lv-Hypp_2509 |
| e08 | SPU_020509 | none |
| e08 | SPU_020628 | Lv-Eif2c2 |
| e08 | SPU_020659 | Lv-Sap130L |
| e08 | SPU_020761 | Lv-Golga4 |
| e08 | SPU_020766 | Lv-SnrpC |
| e08 | SPU_020767 | NA |
| e08 | SPU_020992 | none |
| e08 | SPU_021050 | Lv-Tpase2 |

| | | |
|---|---|---|
| e08 | SPU_021107 | Lv-Hypp_1010 |
| e08 | SPU_021118 | Lv-Rab6ip1l |
| e08 | SPU_021123 | Lv-L3mbt |
| e08 | SPU_021210 | Lv-Tead4 |
| e08 | SPU_021282 | none |
| e08 | SPU_021316 | Lv-Ppp1r10 |
| e08 | SPU_021328 | Lv-Contactin2 |
| e08 | SPU_021352 | Lv-Acel_8 |
| e08 | SPU_021366 | Lv-Snrp70 |
| e08 | SPU_021382 | Lv-Z202 |
| e08 | SPU_021487 | Lv-Oraov1 |
| e08 | SPU_021496 | Lv-Polyq |
| e08 | SPU_021506 | none |
| e08 | SPU_021543 | Lv-Gtpbp1 |
| e08 | SPU_021616 | Lv-Psf2 |
| e08 | SPU_021648 | none |
| e08 | SPU_021687 | Lv-Recq |
| e08 | SPU_021695 | Lv-Apc11 |
| e08 | SPU_021700 | Lv-Apla_1 |
| e08 | SPU_021757 | Lv-Z347 |
| e08 | SPU_021807 | Lv-Ccdc53 |
| e08 | SPU_021835 | none |
| e08 | SPU_022030 | Lv-RtL_55 |
| e08 | SPU_022071 | Lv-C3orf21 |
| e08 | SPU_022078 | Lv-LamG/Egff4 |
| e08 | SPU_022081 | none |
| e08 | SPU_022084 | none |
| e08 | SPU_022143 | Lv-Hypp_1039 |
| e08 | SPU_022163 | Lv-Max |
| e08 | SPU_022322 | Lv-Apc3 |
| e08 | SPU_022331 | Lv-Kiaa0517L_1 |
| e08 | SPU_022372 | Lv-Jmjd1b |
| e08 | SPU_022379 | Lv-EchnL_8 |
| e08 | SPU_022482 | Lv-Loxhd1 |
| e08 | SPU_022494 | Lv-Nphs1L |
| e08 | SPU_022515 | Lv-Dcun1d3 |
| e08 | SPU_022714 | Lv-Ldla/Egf/Ig/Gpcr |
| e08 | SPU_022766 | Lv-Trrap_2 |

| | | |
|---|---|---|
| e08 | SPU_022784 | Lv-Lactb |
| e08 | SPU_022808 | Lv-Hypp_189 |
| e08 | SPU_022819 | Lv-Hypp_190 |
| e08 | SPU_022866 | Lv-Endrvt105 |
| e08 | SPU_022907 | Lv-Kdelr2 |
| e08 | SPU_023035 | Lv-Tlr119 |
| e08 | SPU_023037 | Lv-Bbx |
| e08 | SPU_023148 | Lv-Chordc1 |
| e08 | SPU_023160 | Lv-Heatr4 |
| e08 | SPU_023296 | Lv-Chd3 |
| e08 | SPU_023333 | Lv-Sec13h |
| e08 | SPU_023381 | none |
| e08 | SPU_023506 | Lv-Ino80b |
| e08 | SPU_023527 | Lv-Wdr37 |
| e08 | SPU_023537 | none |
| e08 | SPU_023559 | none |
| e08 | SPU_023695 | Lv-Nab1L |
| e08 | SPU_023704 | Lv-PppL_220 |
| e08 | SPU_023718 | NA |
| e08 | SPU_023774 | Lv-Smg6L |
| e08 | SPU_023863 | NA |
| e08 | SPU_023906 | Lv-Rnf38_1 |
| e08 | SPU_024002 | Lv-Seawi_1 |
| e08 | SPU_024014 | none |
| e08 | SPU_024039 | Lv-Lix1L |
| e08 | SPU_024060 | Lv-Sfrs1L |
| e08 | SPU_024065 | none |
| e08 | SPU_024074 | Lv-PtprW2 |
| e08 | SPU_024087 | Lv-Reeler/Ring |
| e08 | SPU_024097 | Lv-Ssbp3L |
| e08 | SPU_024130 | Lv-Tmem165 |
| e08 | SPU_024145 | Lv-Pde4dipL_1 |
| e08 | SPU_024160 | Lv-Hypp_209 |
| e08 | SPU_024175 | Lv-Hypp_210 |
| e08 | SPU_024180 | Lv-Tfip11 |
| e08 | SPU_024213 | Lv-Hypp_2716 |
| e08 | SPU_024223 | Lv-Hypp_212 |
| e08 | SPU_024256 | Lv-Hprp8bp |

| | | |
|---|---|---|
| e08 | SPU_024281 | Lv-PolqL |
| e08 | SPU_024309 | Lv-Fastkd1L |
| e08 | SPU_024332 | Lv-Ack2 |
| e08 | SPU_024344 | Lv-Ftsj2 |
| e08 | SPU_024383 | Lv-Atp7a |
| e08 | SPU_024394 | Lv-Camk4 |
| e08 | SPU_024399 | Lv-Zbed1 |
| e08 | SPU_024400 | Lv-Zbed1_5 |
| e08 | SPU_024421 | none |
| e08 | SPU_024450 | Lv-Slc26a11_1 |
| e08 | SPU_024508 | Lv-LacL |
| e08 | SPU_024565 | Lv-Cd109L |
| e08 | SPU_024570 | none |
| e08 | SPU_024591 | Lv-Hypp_1117 |
| e08 | SPU_024689 | Lv-Hypp_1123 |
| e08 | SPU_024743 | Lv-Hypp_2751 |
| e08 | SPU_024798 | Lv-U2af1 |
| e08 | SPU_024830 | none |
| e08 | SPU_024867 | none |
| e08 | SPU_024950 | Lv-Hypp_221 |
| e08 | SPU_024999 | NA |
| e08 | SPU_025000 | Lv-Jtbl |
| e08 | SPU_025022 | Lv-Lrrc63 |
| e08 | SPU_025094 | Lv-Hypp_2771 |
| e08 | SPU_025101 | none |
| e08 | SPU_025165 | none |
| e08 | SPU_025172 | NA |
| e08 | SPU_025184 | Lv-Clect_48 |
| e08 | SPU_025218 | none |
| e08 | SPU_025255 | Lv-Hypp_227 |
| e08 | SPU_025363 | Lv-Ttrspn_20 |
| e08 | SPU_025365 | none |
| e08 | SPU_025380 | Lv-Galnt7_3 |
| e08 | SPU_025460 | Lv-Rnf11 |
| e08 | SPU_025556 | Lv-Hypp_2795 |
| e08 | SPU_025657 | Lv-Brg1 |
| e08 | SPU_025686 | Lv-Hccs |
| e08 | SPU_025705 | Lv-Sec14L2_2 |

| | | |
|---|---|---|
| e08 | SPU_025716 | Lv-Helz-3 |
| e08 | SPU_025846 | Lv-PppL_145 |
| e08 | SPU_025857 | Lv-Bat4L |
| e08 | SPU_025883 | Lv-C16orf73L |
| e08 | SPU_025906 | Lv-Rxfp2_6 |
| e08 | SPU_025909 | Lv-Sfrs7 |
| e08 | SPU_025959 | Lv-Z314 |
| e08 | SPU_026004 | Lv-Hypp_2810 |
| e08 | SPU_026007 | none |
| e08 | SPU_026109 | NA |
| e08 | SPU_026116 | Lv-Wdr33 |
| e08 | SPU_026155 | NA |
| e08 | SPU_026159 | Lv-Arid |
| e08 | SPU_026163 | Lv-Hypp_2823 |
| e08 | SPU_026194 | Lv-Hypp_1171 |
| e08 | SPU_026205 | Lv-Mnt |
| e08 | SPU_026262 | Lv-Zmym4 |
| e08 | SPU_026287 | none |
| e08 | SPU_026465 | Lv-Mll3 |
| e08 | SPU_026549 | Lv-NeurlsnL_1 |
| e08 | SPU_026559 | NA |
| e08 | SPU_026594 | Lv-Cul2 |
| e08 | SPU_026616 | Lv-Atg16L1 |
| e08 | SPU_026644 | Lv-Gcf1 |
| e08 | SPU_026661 | none |
| e08 | SPU_026679 | Lv-Fbxl6L |
| e08 | SPU_026698 | Lv-Rag1l6 |
| e08 | SPU_026714 | Lv-Adpgk_2 |
| e08 | SPU_026796 | NA |
| e08 | SPU_026897 | Lv-Lsm12L |
| e08 | SPU_026950 | Lv-Klhl24 |
| e08 | SPU_027079 | Lv-Clect_54 |
| e08 | SPU_027133 | Lv-Cadherin_8 |
| e08 | SPU_027156 | Lv-Wrch |
| e08 | SPU_027181 | none |
| e08 | SPU_027195 | none |
| e08 | SPU_027196 | Lv-Trmt12 |
| e08 | SPU_027250 | Lv-A2(IV)L |

| | | |
|---|---|---|
| e08 | SPU_027258 | Lv-GagpolL_11 |
| e08 | SPU_027277 | none |
| e08 | SPU_027324 | Lv-Endrvt116 |
| e08 | SPU_027365 | Lv-Endrvt117 |
| e08 | SPU_027403 | Lv-Hypp_267 |
| e08 | SPU_027451 | Lv-CbpdDL |
| e08 | SPU_027471 | Lv-Hypp_2906 |
| e08 | SPU_027658 | none |
| e08 | SPU_027673 | Lv-Gldc |
| e08 | SPU_027675 | Lv-Hsat |
| e08 | SPU_027719 | none |
| e08 | SPU_027874 | none |
| e08 | SPU_027876 | Lv-Ppgl/Mut |
| e08 | SPU_027937 | Lv-Dynlc2-6a |
| e08 | SPU_027963 | Lv-Ubr7 |
| e08 | SPU_027981 | none |
| e08 | SPU_028009 | Lv-Phldb1L |
| e08 | SPU_028045 | Lv-Z110 |
| e08 | SPU_028061 | Lv-Dac |
| e08 | SPU_028150 | Lv-Kndc1 |
| e08 | SPU_028283 | Lv-Hypp_2947 |
| e08 | SPU_028398 | Lv-Gpatch1 |
| e08 | SPU_028434 | Lv-Unk_98 |
| e08 | SPU_028508 | NA |
| e08 | SPU_028614 | Lv-Tex9 |
| e08 | SPU_028653 | none |
| e08 | SPU_028671 | Lv-Crmp1 |
| e08 | SPU_028719 | Lv-Clca4L_2 |
| e08 | SPU_028722 | Lv-Msl1L |
| e08 | SPU_028739 | Lv-Syyn1 |
| e08 | SPU_028753 | Lv-Z359 |
| e08 | SPU_028835 | Lv-Tfip11 |
| e08 | SPU_028906 | none |
| e08 | SPU_028919 | Lv-Hsp40E |
| e08 | SPU_030075 | NA |
| e08 | SPU_030143 | Lv-Cd59/Sca2L2 |
| e09 | SPU_000101 | Lv-Sf3B3 |
| e09 | SPU_000118 | Lv-Aasdhppt |

| | | |
|---|---|---|
| e09 | SPU_000142 | Lv-Col18a1Lf |
| e09 | SPU_000144 | Lv-Nop5 |
| e09 | SPU_000216 | Lv-Egf/HyalinL14 |
| e09 | SPU_000236 | Lv-Idh1 |
| e09 | SPU_000264 | Lv-Tf2L |
| e09 | SPU_000319 | Lv-Nudcd2l |
| e09 | SPU_000333 | Lv-Zbed1-2 |
| e09 | SPU_000440 | Lv-Z115 |
| e09 | SPU_000456 | Lv-Ppcs |
| e09 | SPU_000565 | Lv-Hypp_307 |
| e09 | SPU_000573 | Lv-Hypp_309 |
| e09 | SPU_000637 | Lv-Pdpi(cyclophilin)L5 |
| e09 | SPU_000788 | none |
| e09 | SPU_000830 | Lv-Gss |
| e09 | SPU_000835 | Lv-Pah |
| e09 | SPU_000843 | none |
| e09 | SPU_000860 | Lv-Slc16a12L_4 |
| e09 | SPU_001185 | Lv-Tmem147 |
| e09 | SPU_001265 | Lv-Dpf2 |
| e09 | SPU_001355 | none |
| e09 | SPU_001387 | Lv-Ugt2b1 |
| e09 | SPU_001401 | Lv-Trrap |
| e09 | SPU_001428 | none |
| e09 | SPU_001446 | Lv-Hectd1 |
| e09 | SPU_001462 | Lv-Mylip-2 |
| e09 | SPU_001595 | Lv-Slc26a5 |
| e09 | SPU_001603 | Lv-Reptin52 |
| e09 | SPU_001704 | NA |
| e09 | SPU_001795 | none |
| e09 | SPU_001826 | Lv-Cpsf1 |
| e09 | SPU_001915 | none |
| e09 | SPU_002090 | Lv-Z342 |
| e09 | SPU_002159 | Lv-Znf862L |
| e09 | SPU_002176 | none |
| e09 | SPU_002193 | Lv-Apc6 |
| e09 | SPU_002284 | Lv-Npac |
| e09 | SPU_002563 | none |
| e09 | SPU_002578 | Lv-Acin1 |

| | | |
|---|---|---|
| e09 | SPU_002603 | Lv-SoxC |
| e09 | SPU_002680 | Lv-Abcf2 |
| e09 | SPU_002716 | Lv-Hypp_1408 |
| e09 | SPU_002725 | Lv-Hgd |
| e09 | SPU_002749 | NA |
| e09 | SPU_002939 | Lv-Acin1l |
| e09 | SPU_003087 | Lv-Cpsf3 |
| e09 | SPU_003124 | none |
| e09 | SPU_003133 | Lv-Sfrs7L |
| e09 | SPU_003154 | NA |
| e09 | SPU_003166 | Lv-Myc |
| e09 | SPU_003348 | Lv-Dusp16 |
| e09 | SPU_003370 | Lv-Unc44L_30 |
| e09 | SPU_003479 | Lv-Cdkn3 |
| e09 | SPU_003697 | Lv-Hypp_395 |
| e09 | SPU_003705 | Lv-Mta1_1 |
| e09 | SPU_003739 | Lv-Hypp_1479 |
| e09 | SPU_003746 | Lv-Unk_108 |
| e09 | SPU_003750 | Lv-Ncapg2 |
| e09 | SPU_003796 | none |
| e09 | SPU_003923 | Lv-Scrnph2 |
| e09 | SPU_003943 | NA |
| e09 | SPU_003951 | none |
| e09 | SPU_003997 | Lv-Adra1L_2 |
| e09 | SPU_004373 | none |
| e09 | SPU_004476 | Lv-HyalinL_1 |
| e09 | SPU_004514 | Lv-Hypp_1532 |
| e09 | SPU_004524 | Lv-Nup210 |
| e09 | SPU_004596 | Lv-U2A |
| e09 | SPU_004635 | Lv-Crop |
| e09 | SPU_004663 | Lv-Kiaa1731L |
| e09 | SPU_004664 | none |
| e09 | SPU_004706 | Lv-Hypp_1537 |
| e09 | SPU_004715 | Lv-Bcas2 |
| e09 | SPU_004836 | Lv-Ewsr1l_1 |
| e09 | SPU_004879 | Lv-Fam55cL_2 |
| e09 | SPU_004942 | Lv-Samd1L |
| e09 | SPU_004983 | Lv-Chordin |

| | | |
|---|---|---|
| e09 | SPU_004998 | Lv-Ttc17 |
| e09 | SPU_005026 | Lv-Nlr124 |
| e09 | SPU_005082 | none |
| e09 | SPU_005336 | Lv-PolppL_44 |
| e09 | SPU_005339 | Lv-Tlr026 |
| e09 | SPU_005419 | Lv-Sfk3 |
| e09 | SPU_005528 | none |
| e09 | SPU_005572 | Lv-Hmg2 |
| e09 | SPU_005617 | Lv-Slc16a1L |
| e09 | SPU_005766 | NA |
| e09 | SPU_005856 | Lv-Ckip1 |
| e09 | SPU_006145 | Lv-Kiaa1324L |
| e09 | SPU_006160 | none |
| e09 | SPU_006430 | none |
| e09 | SPU_006432 | NA |
| e09 | SPU_006433 | Lv-Kiaa0907L |
| e09 | SPU_006446 | Lv-Rbm28 |
| e09 | SPU_006493 | none |
| e09 | SPU_006623 | Lv-Satb1 |
| e09 | SPU_006716 | Lv-Hypp_491 |
| e09 | SPU_006739 | none |
| e09 | SPU_006743 | none |
| e09 | SPU_006775 | Lv-Tnc |
| e09 | SPU_006825 | NA |
| e09 | SPU_006868 | Lv-Endrvt63 |
| e09 | SPU_006872 | Lv-Kiaa0907L_1 |
| e09 | SPU_006876 | Lv-Abcc5B |
| e09 | SPU_007031 | none |
| e09 | SPU_007096 | Lv-Hmgb3 |
| e09 | SPU_007142 | Lv-Mfn |
| e09 | SPU_007336 | none |
| e09 | SPU_007389 | Lv-Mta1 |
| e09 | SPU_007483 | Lv-Cnot10 |
| e09 | SPU_007528 | Lv-PolppL_60 |
| e09 | SPU_007580 | none |
| e09 | SPU_007592 | Lv-Dnajhc17 |
| e09 | SPU_007622 | none |
| e09 | SPU_007668 | Lv-Hypp_26 |

| | | |
|---|---|---|
| e09 | SPU_007723 | Lv-Tcfap4L |
| e09 | SPU_007850 | Lv-Tlr040 |
| e09 | SPU_007891 | none |
| e09 | SPU_007941 | Lv-Lsm/Unk_1 |
| e09 | SPU_008011 | Lv-Hnrnpk |
| e09 | SPU_008240 | none |
| e09 | SPU_008304 | Lv-Myst2 |
| e09 | SPU_008374 | Lv-Farp |
| e09 | SPU_008532 | Lv-Dhx37 |
| e09 | SPU_008533 | Lv-Ints2 |
| e09 | SPU_008689 | Lv-Wash1_1 |
| e09 | SPU_008693 | none |
| e09 | SPU_008704 | Lv-Prkrip |
| e09 | SPU_008723 | Lv-Gopc |
| e09 | SPU_008753 | Lv-Mib1_1 |
| e09 | SPU_008768 | Lv-Hdact |
| e09 | SPU_008781 | Lv-Hypp_560 |
| e09 | SPU_009014 | NA |
| e09 | SPU_009033 | Lv-Echn28 |
| e09 | SPU_009042 | Lv-Cts2 |
| e09 | SPU_009061 | Lv-Avl19L |
| e09 | SPU_009087 | Lv-Gal3st1_4 |
| e09 | SPU_009107 | Lv-2310067g05rikh |
| e09 | SPU_009150 | Lv-Eif2beta |
| e09 | SPU_009156 | Lv-Hypp_1827 |
| e09 | SPU_009193 | none |
| e09 | SPU_009226 | Lv-Nt5c3 |
| e09 | SPU_009242 | Lv-Cub/Fa58C/Ldla/Gpcr |
| e09 | SPU_009282 | Lv-Smarce1 |
| e09 | SPU_009372 | Lv-B3galt1_7 |
| e09 | SPU_009373 | Lv-Hypp_1846 |
| e09 | SPU_009383 | none |
| e09 | SPU_009729 | none |
| e09 | SPU_009739 | Lv-Rnf38 |
| e09 | SPU_009759 | Lv-Sf3B5 |
| e09 | SPU_009809 | Lv-Hypp_1874 |
| e09 | SPU_009813 | Lv-Hypp_589 |
| e09 | SPU_009861 | Lv-Gagpol4 |

| | | |
|---|---|---|
| e09 | SPU_009891 | Lv-Nop56 |
| e09 | SPU_009894 | none |
| e09 | SPU_009919 | Lv-Fkbp4L |
| e09 | SPU_010118 | Lv-P38mapk_1 |
| e09 | SPU_010163 | Lv-Phf12 |
| e09 | SPU_010254 | Lv-Yeats4 |
| e09 | SPU_010285 | Lv-Tprjmjc |
| e09 | SPU_010300 | Lv-Hypp_1908 |
| e09 | SPU_010312 | Lv-Utp11L |
| e09 | SPU_010328 | Lv-Ebna1bp2 |
| e09 | SPU_010520 | Lv-Aof2 |
| e09 | SPU_010659 | Lv-Ski |
| e09 | SPU_010704 | Lv-Ugt1L_2 |
| e09 | SPU_010766 | Lv-Rnf8 |
| e09 | SPU_010862 | Lv-Enpp7_1 |
| e09 | SPU_010902 | Lv-Strongylocentrotus_purpuratus-specific_proteinl2b |
| e09 | SPU_010957 | Lv-Rskl |
| e09 | SPU_011033 | none |
| e09 | SPU_011043 | Lv-Endrvt23 |
| e09 | SPU_011094 | Lv-Noc3l |
| e09 | SPU_011117 | none |
| e09 | SPU_011146 | Lv-Srcr100 |
| e09 | SPU_011169 | Lv-Hypp_1965 |
| e09 | SPU_011229 | NA |
| e09 | SPU_011271 | Lv-Sfrp1/5 |
| e09 | SPU_011296 | Lv-Lsm/Unk |
| e09 | SPU_011334 | Lv-Pdxdc1 |
| e09 | SPU_011404 | Lv-Rergl2 |
| e09 | SPU_011606 | Lv-Chrna6L |
| e09 | SPU_011617 | Lv-Abcc4d |
| e09 | SPU_011676 | Lv-Dmap1_1 |
| e09 | SPU_011725 | Lv-Tm9sf3 |
| e09 | SPU_011727 | Lv-Entpd5 |
| e09 | SPU_011988 | Lv-Endrvt25 |
| e09 | SPU_012134 | Lv-Zbed1-3 |
| e09 | SPU_012182 | Lv-Chst12L |
| e09 | SPU_012225 | Lv-Hypp_677 |

| | | |
|---|---|---|
| e09 | SPU_012257 | Lv-Tlr068 |
| e09 | SPU_012337 | Lv-Thrap4L |
| e09 | SPU_012376 | Lv-Sld5 |
| e09 | SPU_012447 | Lv-Hypp_2032 |
| e09 | SPU_012490 | Lv-Hmx |
| e09 | SPU_012599 | Lv-Imp |
| e09 | SPU_012637 | Lv-Caskin1 |
| e09 | SPU_012671 | Lv-Tirc4 |
| e09 | SPU_012711 | none |
| e09 | SPU_012825 | NA |
| e09 | SPU_012838 | Lv-Creb3l1 |
| e09 | SPU_012927 | Lv-Hypp_2056 |
| e09 | SPU_013059 | none |
| e09 | SPU_013115 | Lv-Hypp_703 |
| e09 | SPU_013196 | Lv-Hypp_2075 |
| e09 | SPU_013267 | Lv-Hypp_2077 |
| e09 | SPU_013303 | Lv-Nrarp |
| e09 | SPU_013400 | Lv-Hypp_2081 |
| e09 | SPU_013478 | Lv-Nwd1_2 |
| e09 | SPU_013507 | Lv-Pigf |
| e09 | SPU_013509 | Lv-Odadc1L |
| e09 | SPU_013582 | none |
| e09 | SPU_013680 | Lv-Endrvt29 |
| e09 | SPU_013686 | Lv-Hmmr |
| e09 | SPU_013861 | Lv-Z190 |
| e09 | SPU_013915 | Lv-Slc35f4 |
| e09 | SPU_013971 | NA |
| e09 | SPU_013984 | Lv-Hypp_97 |
| e09 | SPU_013992 | Lv-Catsper4L |
| e09 | SPU_014248 | Lv-Akap6L |
| e09 | SPU_014279 | Lv-Nav3l |
| e09 | SPU_014428 | Lv-Yars2 |
| e09 | SPU_014430 | Lv-U5116kD |
| e09 | SPU_014439 | Lv-Kiaa0664L |
| e09 | SPU_014498 | Lv-Gg1 |
| e09 | SPU_014513 | NA |
| e09 | SPU_014536 | NA |
| e09 | SPU_014542 | Lv-Samd8 |

| | | |
|---|---|---|
| e09 | SPU_014543 | Lv-Hypp_103 |
| e09 | SPU_014579 | Lv-Eif3a |
| e09 | SPU_014611 | Lv-Rbm35A |
| e09 | SPU_014631 | Lv-Rt6 |
| e09 | SPU_014645 | Lv-Nomo |
| e09 | SPU_014808 | Lv-Ddx48 |
| e09 | SPU_014889 | none |
| e09 | SPU_014959 | Lv-Ape |
| e09 | SPU_014969 | none |
| e09 | SPU_015145 | Lv-SfpqL |
| e09 | SPU_015151 | Lv-Hypp_2193 |
| e09 | SPU_015174 | Lv-Rt36 |
| e09 | SPU_015214 | none |
| e09 | SPU_015331 | Lv-Utp6L |
| e09 | SPU_015386 | Lv-Supt6h |
| e09 | SPU_015400 | Lv-Unc44_308 |
| e09 | SPU_015404 | Lv-LamG/Egff3 |
| e09 | SPU_015407 | none |
| e09 | SPU_015468 | NA |
| e09 | SPU_015554 | Lv-TyrL_1 |
| e09 | SPU_015594 | Lv-Gopc_1 |
| e09 | SPU_015643 | Lv-Pot1h |
| e09 | SPU_015708 | Lv-Col4a2 |
| e09 | SPU_015729 | none |
| e09 | SPU_015799 | none |
| e09 | SPU_015800 | Lv-Z246 |
| e09 | SPU_015877 | NA |
| e09 | SPU_015970 | NA |
| e09 | SPU_016000 | Lv-GagpL_2 |
| e09 | SPU_016096 | Lv-Cycs_2 |
| e09 | SPU_016139 | NA |
| e09 | SPU_016157 | Lv-Papss |
| e09 | SPU_016352 | Lv-Sak |
| e09 | SPU_016383 | Lv-Slco4a1_3 |
| e09 | SPU_016395 | Lv-Ckap2L |
| e09 | SPU_016552 | Lv-Ddx42_1 |
| e09 | SPU_016599 | NA |
| e09 | SPU_016676 | Lv-Hypp_838 |

| | | |
|---|---|---|
| e09 | SPU_016699 | Lv-Usp39 |
| e09 | SPU_016733 | Lv-Med4 |
| e09 | SPU_016777 | Lv-Endrvt32 |
| e09 | SPU_016876 | none |
| e09 | SPU_016928 | Lv-Unc44_120 |
| e09 | SPU_017087 | Lv-Ttc19L |
| e09 | SPU_017199 | Lv-Hypp_2310 |
| e09 | SPU_017220 | Lv-Cmas |
| e09 | SPU_017237 | NA |
| e09 | SPU_017266 | Lv-Tpase5 |
| e09 | SPU_017319 | Lv-Cenpo |
| e09 | SPU_017342 | Lv-Slc7a9_4 |
| e09 | SPU_017377 | NA |
| e09 | SPU_017430 | Lv-Hypp_879 |
| e09 | SPU_017596 | none |
| e09 | SPU_017603 | Lv-Slsp_4 |
| e09 | SPU_017777 | Lv-Preb |
| e09 | SPU_017802 | Lv-SnrpB_1 |
| e09 | SPU_017806 | Lv-Fam44a |
| e09 | SPU_017836 | Lv-Acp2_1 |
| e09 | SPU_017861 | Lv-Spo11 |
| e09 | SPU_017911 | none |
| e09 | SPU_017935 | Lv-Unk_102 |
| e09 | SPU_017943 | none |
| e09 | SPU_018004 | Lv-Mcrs1 |
| e09 | SPU_018027 | Lv-Hgd_1 |
| e09 | SPU_018131 | none |
| e09 | SPU_018265 | Lv-Hypp_141 |
| e09 | SPU_018278 | Lv-Endrvt36 |
| e09 | SPU_018304 | Lv-Larp7 |
| e09 | SPU_018576 | Lv-Gatad1 |
| e09 | SPU_018602 | Lv-PppL_121 |
| e09 | SPU_018605 | none |
| e09 | SPU_018626 | Lv-Nphp4 |
| e09 | SPU_018662 | none |
| e09 | SPU_018792 | Lv-Pb1_1 |
| e09 | SPU_018805 | Lv-Fbl14 |
| e09 | SPU_018841 | Lv-Ddx49 |

| | | |
|---|---|---|
| e09 | SPU_018897 | Lv-If5 |
| e09 | SPU_018924 | Lv-Cybb |
| e09 | SPU_018943 | Lv-Hirha |
| e09 | SPU_019113 | Lv-RapL |
| e09 | SPU_019278 | Lv-Slc9a10 |
| e09 | SPU_019357 | Lv-Agpat5 |
| e09 | SPU_019361 | Lv-Endrvt100 |
| e09 | SPU_019425 | Lv-Brd7 |
| e09 | SPU_019534 | NA |
| e09 | SPU_019557 | Lv-Hnrpr |
| e09 | SPU_019577 | none |
| e09 | SPU_019657 | Lv-Sr140 |
| e09 | SPU_019701 | Lv-AcpL2 |
| e09 | SPU_019718 | Lv-Pof1 |
| e09 | SPU_019724 | Lv-Hypp_157 |
| e09 | SPU_019851 | Lv-Hypp_950 |
| e09 | SPU_019866 | none |
| e09 | SPU_019980 | Lv-Parp2 |
| e09 | SPU_020010 | Lv-Top3A |
| e09 | SPU_020058 | Lv-Vti1bh |
| e09 | SPU_020105 | none |
| e09 | SPU_020169 | NA |
| e09 | SPU_020173 | Lv-Hypp_3034 |
| e09 | SPU_020184 | Lv-RtL_49 |
| e09 | SPU_020233 | Lv-Cyp2L41 |
| e09 | SPU_020251 | Lv-Fk506bp4 |
| e09 | SPU_020368 | Lv-Atp13a1 |
| e09 | SPU_020468 | Lv-PppL_216 |
| e09 | SPU_020472 | Lv-Hypp_975 |
| e09 | SPU_020491 | Lv-Thredh |
| e09 | SPU_020582 | Lv-Hypp_2513 |
| e09 | SPU_020598 | Lv-Ltmd1 |
| e09 | SPU_020677 | Lv-Fgfr |
| e09 | SPU_020679 | Lv-Stard7 |
| e09 | SPU_020693 | Lv-Supt6h |
| e09 | SPU_020872 | Lv-G6pc_1 |
| e09 | SPU_020931 | Lv-Tm9sf2 |
| e09 | SPU_020948 | Lv-Rt15 |

| | | |
|---|---|---|
| e09 | SPU_020980 | Lv-Ankib1 |
| e09 | SPU_021019 | Lv-Fn3f_29 |
| e09 | SPU_021044 | Lv-Notchl6 |
| e09 | SPU_021076 | Lv-Unc44_150 |
| e09 | SPU_021081 | Lv-Endrvt41 |
| e09 | SPU_021115 | Lv-PppL_217 |
| e09 | SPU_021286 | NA |
| e09 | SPU_021294 | Lv-Endrvt43 |
| e09 | SPU_021301 | none |
| e09 | SPU_021308 | Lv-Sra |
| e09 | SPU_021340 | Lv-Lrr/Fn3f |
| e09 | SPU_021341 | none |
| e09 | SPU_021479 | none |
| e09 | SPU_021510 | Lv-Hectd1_1 |
| e09 | SPU_021512 | Lv-Papol |
| e09 | SPU_021556 | Lv-Hypp_3033 |
| e09 | SPU_021608 | Lv-HesC |
| e09 | SPU_021622 | Lv-Unc44_155 |
| e09 | SPU_021629 | Lv-Smc1L1 |
| e09 | SPU_021631 | Lv-Trspap1 |
| e09 | SPU_021766 | NA |
| e09 | SPU_021777 | Lv-C17orf53L |
| e09 | SPU_021791 | Lv-Adra1aL |
| e09 | SPU_021864 | Lv-Ndnl2 |
| e09 | SPU_021889 | Lv-Notch2L_1 |
| e09 | SPU_021966 | Lv-Hsd17b2_1 |
| e09 | SPU_021989 | Lv-AcotL |
| e09 | SPU_022052 | Lv-Odz3 |
| e09 | SPU_022068 | none |
| e09 | SPU_022134 | Lv-Gal3st3L_4 |
| e09 | SPU_022354 | Lv-Ldlrap1 |
| e09 | SPU_022393 | Lv-Hypp_182 |
| e09 | SPU_022398 | Lv-Ipo11 |
| e09 | SPU_022477 | Lv-Slc25a26_2 |
| e09 | SPU_022496 | Lv-RtL_57 |
| e09 | SPU_022583 | none |
| e09 | SPU_022626 | Lv-Lin52 |
| e09 | SPU_022734 | Lv-U2af2 |

| | | |
|---|---|---|
| e09 | SPU_022746 | Lv-Tmem115 |
| e09 | SPU_022869 | Lv-Wdr25 |
| e09 | SPU_022876 | none |
| e09 | SPU_022878 | Lv-Rbm14 |
| e09 | SPU_022937 | Lv-Hypp_2639 |
| e09 | SPU_022940 | Lv-TardbpL2 |
| e09 | SPU_022970 | Lv-Kiaa1429L |
| e09 | SPU_022974 | Lv-Polyd |
| e09 | SPU_023014 | Lv-Frizz9/10 |
| e09 | SPU_023177 | Lv-Hbn |
| e09 | SPU_023196 | none |
| e09 | SPU_023203 | Lv-Zranb1 |
| e09 | SPU_023221 | none |
| e09 | SPU_023276 | Lv-Sec14L_1 |
| e09 | SPU_023344 | Lv-Rbm17L |
| e09 | SPU_023399 | Lv-Hyr_1 |
| e09 | SPU_023424 | Lv-Bmi1 |
| e09 | SPU_023466 | Lv-Calu |
| e09 | SPU_023571 | Lv-Glex3 |
| e09 | SPU_023674 | Lv-EndrvtB |
| e09 | SPU_023778 | Lv-Glt28d2 |
| e09 | SPU_023789 | Lv-Sfrs11 |
| e09 | SPU_023881 | Lv-Nolc1 |
| e09 | SPU_023888 | Lv-Cstf3_1 |
| e09 | SPU_023889 | Lv-Arrb1 |
| e09 | SPU_023941 | Lv-Myt1 |
| e09 | SPU_023951 | NA |
| e09 | SPU_024128 | Lv-Agapd |
| e09 | SPU_024341 | Lv-Ankrd54 |
| e09 | SPU_024491 | Lv-Hypp_216 |
| e09 | SPU_024675 | Lv-Yeats2 |
| e09 | SPU_024692 | Lv-Hypp_2747 |
| e09 | SPU_024703 | none |
| e09 | SPU_024733 | Lv-Tlr132 |
| e09 | SPU_024854 | Lv-RtLp |
| e09 | SPU_024890 | Lv-HH2az |
| e09 | SPU_025068 | Lv-Ttrspn_19 |
| e09 | SPU_025089 | Lv-Krcp1 |

| | | |
|---|---|---|
| e09 | SPU_025124 | Lv-Hypp_1138 |
| e09 | SPU_025139 | Lv-Tdg |
| e09 | SPU_025144 | Lv-Cstf3 |
| e09 | SPU_025205 | Lv-TpaseL_17 |
| e09 | SPU_025281 | none |
| e09 | SPU_025325 | Lv-DhpmdL5_1 |
| e09 | SPU_025381 | NA |
| e09 | SPU_025385 | Lv-PppL_142 |
| e09 | SPU_025424 | Lv-Hsp100A |
| e09 | SPU_025444 | none |
| e09 | SPU_025467 | none |
| e09 | SPU_025515 | Lv-Slc4a10 |
| e09 | SPU_025572 | Lv-Dis |
| e09 | SPU_025595 | Lv-Cyp51 |
| e09 | SPU_025653 | none |
| e09 | SPU_025688 | none |
| e09 | SPU_025693 | Lv-Golga7 |
| e09 | SPU_025733 | Lv-Par3L |
| e09 | SPU_025784 | none |
| e09 | SPU_025815 | Lv-Atp1a3 |
| e09 | SPU_025821 | Lv-Hrh3L_19 |
| e09 | SPU_025864 | Lv-Ncaph1 |
| e09 | SPU_025979 | Lv-Yl1 |
| e09 | SPU_026001 | none |
| e09 | SPU_026050 | Lv-Ndufv1 |
| e09 | SPU_026096 | Lv-Calnexin |
| e09 | SPU_026147 | Lv-Dmap1 |
| e09 | SPU_026160 | Lv-Lipf_1 |
| e09 | SPU_026204 | Lv-Hypp_2827 |
| e09 | SPU_026207 | Lv-Pigl |
| e09 | SPU_026355 | none |
| e09 | SPU_026763 | Lv-Scml1 |
| e09 | SPU_026782 | Lv-Nudt2 |
| e09 | SPU_026801 | Lv-Amy_6 |
| e09 | SPU_026804 | Lv-PppL_150 |
| e09 | SPU_026875 | Lv-Trim2L_2 |
| e09 | SPU_026881 | Lv-Ndufv2 |
| e09 | SPU_026896 | Lv-Hypp_262 |

| | | |
|---|---|---|
| e09 | SPU_026906 | Lv-Hnrnpa1L |
| e09 | SPU_026933 | Lv-Pus3 |
| e09 | SPU_026935 | Lv-Dnajc11 |
| e09 | SPU_027065 | none |
| e09 | SPU_027191 | none |
| e09 | SPU_027235 | Lv-Blimp1 |
| e09 | SPU_027255 | none |
| e09 | SPU_027257 | Lv-PppL_152 |
| e09 | SPU_027438 | none |
| e09 | SPU_027460 | Lv-Spata22L |
| e09 | SPU_027464 | Lv-Hypp_1203 |
| e09 | SPU_027517 | Lv-Tet3L |
| e09 | SPU_027521 | Lv-Hypp_2908 |
| e09 | SPU_027562 | Lv-Msp/Orc |
| e09 | SPU_027704 | Lv-Adra1aL_11 |
| e09 | SPU_027708 | Lv-Z491 |
| e09 | SPU_027790 | Lv-Hypp_275 |
| e09 | SPU_027982 | Lv-Exosc9 |
| e09 | SPU_027993 | none |
| e09 | SPU_028010 | Lv-Erh_1 |
| e09 | SPU_028165 | Lv-Hypp_2939 |
| e09 | SPU_028192 | Lv-PppL_288 |
| e09 | SPU_028219 | Lv-C1orf107 |
| e09 | SPU_028258 | Lv-Zc3m11a |
| e09 | SPU_028259 | Lv-Hypp_1224 |
| e09 | SPU_028306 | Lv-FechL |
| e09 | SPU_028310 | Lv-Slc29a1 |
| e09 | SPU_028340 | Lv-Hypp_283 |
| e09 | SPU_028482 | NA |
| e09 | SPU_028517 | Lv-Drd4L |
| e09 | SPU_028632 | none |
| e09 | SPU_028737 | Lv-Tmem145L |
| e09 | SPU_028738 | Lv-Hypp_1238 |
| e09 | SPU_028761 | Lv-Aadac_6 |
| e09 | SPU_028774 | Lv-Xpc |
| e09 | SPU_028783 | Lv-Glur3_1 |
| e09 | SPU_028784 | Lv-Glur2_2 |
| e09 | SPU_028827 | Lv-E2f4 |

| | | |
|---|---|---|
| e09 | SPU_030106 | NA |
| e10 | SPU_000031 | Lv-Clect/Pan/Kringle |
| e10 | SPU_000339 | Lv-Hypp_1250 |
| e10 | SPU_000512 | Lv-Uco2 |
| e10 | SPU_000514 | Lv-Ksr1 |
| e10 | SPU_000630 | Lv-Fam20c |
| e10 | SPU_000903 | Lv-Hypp_1291 |
| e10 | SPU_000996 | Lv-Hypp_3007 |
| e10 | SPU_001035 | none |
| e10 | SPU_001452 | Lv-Dachs |
| e10 | SPU_001454 | Lv-Fat |
| e10 | SPU_001550 | Lv-Ulk3_1 |
| e10 | SPU_001597 | Lv-Cry |
| e10 | SPU_001598 | NA |
| e10 | SPU_001727 | NA |
| e10 | SPU_001853 | Lv-Hypp_3108 |
| e10 | SPU_001898 | Lv-Clect/Apple_1 |
| e10 | SPU_001998 | Lv-Smad6/7 |
| e10 | SPU_002091 | NA |
| e10 | SPU_002099 | Lv-Hypp_1371 |
| e10 | SPU_002161 | Lv-Sf3B4 |
| e10 | SPU_002246 | Lv-Mekk4a |
| e10 | SPU_002262 | NA |
| e10 | SPU_002461 | Lv-Aplysia/Adp/Ribcyc |
| e10 | SPU_002614 | Lv-Erk3/4 |
| e10 | SPU_002646 | none |
| e10 | SPU_002728 | Lv-Hypp_368 |
| e10 | SPU_002983 | Lv-Nkd2 |
| e10 | SPU_003066 | Lv-Hypp_1428 |
| e10 | SPU_003234 | Lv-Zc3h10 |
| e10 | SPU_003243 | Lv-Hypp_377 |
| e10 | SPU_003483 | none |
| e10 | SPU_003501 | Lv-Plek |
| e10 | SPU_003609 | Lv-Pnkp |
| e10 | SPU_003710 | Lv-Clik1 |
| e10 | SPU_003856 | Lv-Rsf1 |
| e10 | SPU_003889 | Lv-Hs3st2 |
| e10 | SPU_004049 | Lv-Cdc73 |

| | | |
|---|---|---|
| e10 | SPU_004073 | Lv-Hypp_404 |
| e10 | SPU_004080 | NA |
| e10 | SPU_004301 | none |
| e10 | SPU_004307 | Lv-Hypp_1512 |
| e10 | SPU_004310 | Lv-Nt5c2 |
| e10 | SPU_004399 | none |
| e10 | SPU_004538 | Lv-Galnt6 |
| e10 | SPU_005145 | none |
| e10 | SPU_005168 | Lv-Ache-13 |
| e10 | SPU_005224 | NA |
| e10 | SPU_005234 | Lv-Astacin7 |
| e10 | SPU_005400 | none |
| e10 | SPU_005405 | Lv-Hypp_446 |
| e10 | SPU_005524 | none |
| e10 | SPU_005818 | Lv-Ucm |
| e10 | SPU_006117 | none |
| e10 | SPU_006139 | Lv-Hypp_469 |
| e10 | SPU_006183 | Lv-Hypp_2984 |
| e10 | SPU_006215 | Lv-Rbbp6L |
| e10 | SPU_006300 | Lv-Hypp_474 |
| e10 | SPU_006312 | Lv-Dp1 |
| e10 | SPU_006333 | Lv-Sil |
| e10 | SPU_006336 | Lv-Moap1L |
| e10 | SPU_006656 | Lv-Tmtc4 |
| e10 | SPU_006707 | none |
| e10 | SPU_006781 | NA |
| e10 | SPU_006801 | Lv-Rhogap21 |
| e10 | SPU_007019 | Lv-B4galt6_8 |
| e10 | SPU_007233 | Lv-Nkain1 |
| e10 | SPU_007255 | Lv-Lrp1bL |
| e10 | SPU_007325 | Lv-PolppL_58 |
| e10 | SPU_007418 | Lv-Tlr036 |
| e10 | SPU_007646 | Lv-Galr2L_9 |
| e10 | SPU_007686 | Lv-Ptchd2 |
| e10 | SPU_007749 | Lv-Atf6b |
| e10 | SPU_007777 | none |
| e10 | SPU_007785 | Lv-Pxmp3 |
| e10 | SPU_007841 | Lv-Fbxl4 |

| | | |
|---|---|---|
| e10 | SPU_007842 | Lv-Ankrd28 |
| e10 | SPU_007958 | Lv-Fam55cL |
| e10 | SPU_008043 | Lv-Wdr54 |
| e10 | SPU_008107 | Lv-Hypp_31 |
| e10 | SPU_008117 | Lv-Shr2 |
| e10 | SPU_008299 | Lv-DachL |
| e10 | SPU_008499 | Lv-Serrp13 |
| e10 | SPU_008745 | Lv-Gpr133_1 |
| e10 | SPU_008832 | Lv-Rt43 |
| e10 | SPU_008934 | Lv-Slc4a2 |
| e10 | SPU_008941 | Lv-Adrb1L_1 |
| e10 | SPU_008954 | Lv-Map1a/1B_Lc3L2 |
| e10 | SPU_009073 | Lv-FatL2 |
| e10 | SPU_009093 | NA |
| e10 | SPU_009151 | Lv-Oatflp |
| e10 | SPU_009250 | Lv-Fjx1 |
| e10 | SPU_009427 | Lv-Z225 |
| e10 | SPU_009469 | none |
| e10 | SPU_009523 | Lv-Klhl2_1 |
| e10 | SPU_009553 | Lv-Z263 |
| e10 | SPU_009559 | Lv-Plk1_1 |
| e10 | SPU_009563 | Lv-Srcr12_1 |
| e10 | SPU_009605 | none |
| e10 | SPU_009630 | NA |
| e10 | SPU_009636 | Lv-Clect/Egf_9 |
| e10 | SPU_009909 | Lv-Rag1l3 |
| e10 | SPU_010065 | Lv-Ph4_2 |
| e10 | SPU_010188 | Lv-Hsp100C |
| e10 | SPU_010241 | Lv-Srcr88 |
| e10 | SPU_010623 | Lv-Sfrs2 |
| e10 | SPU_010644 | Lv-B3galt1_9 |
| e10 | SPU_010656 | Lv-Ncag1 |
| e10 | SPU_010685 | Lv-Thoc4 |
| e10 | SPU_010687 | Lv-Dhcr7 |
| e10 | SPU_010803 | none |
| e10 | SPU_011027 | Lv-Brca1L |
| e10 | SPU_011068 | NA |
| e10 | SPU_011074 | Lv-Fam55c |

| | | |
|---|---|---|
| e10 | SPU_011123 | Lv-Adrb1L_3 |
| e10 | SPU_011130 | Lv-Wnt16 |
| e10 | SPU_011195 | Lv-Hypp_2207 |
| e10 | SPU_011489 | NA |
| e10 | SPU_011512 | none |
| e10 | SPU_011599 | Lv-Unc44L_75 |
| e10 | SPU_011605 | Lv-Glo1 |
| e10 | SPU_011994 | Lv-Hypp_2010 |
| e10 | SPU_011998 | none |
| e10 | SPU_012175 | NA |
| e10 | SPU_012430 | Lv-Sat2_3 |
| e10 | SPU_012506 | Lv-Dkk3 |
| e10 | SPU_012515 | none |
| e10 | SPU_012584 | Lv-TlrP09 |
| e10 | SPU_012755 | Lv-PolppL_96 |
| e10 | SPU_012913 | Lv-Z92 |
| e10 | SPU_012943 | none |
| e10 | SPU_012980 | Lv-Parp1_1 |
| e10 | SPU_013005 | Lv-Th1L |
| e10 | SPU_013063 | Lv-Hypp_86 |
| e10 | SPU_013305 | Lv-Grf |
| e10 | SPU_013320 | Lv-Dnl3 |
| e10 | SPU_013508 | Lv-Mitosin |
| e10 | SPU_013520 | Lv-FarpL_1 |
| e10 | SPU_013672 | Lv-UacaL |
| e10 | SPU_013695 | Lv-Hypp_2098 |
| e10 | SPU_013757 | Lv-Slco4a1_2 |
| e10 | SPU_013951 | none |
| e10 | SPU_014069 | Lv-Rag1l2 |
| e10 | SPU_014125 | Lv-Endrvt87 |
| e10 | SPU_014185 | Lv-Wee1_1 |
| e10 | SPU_014308 | Lv-Hypp_112 |
| e10 | SPU_014384 | Lv-Mtnr1bL |
| e10 | SPU_014494 | Lv-Pex10 |
| e10 | SPU_014548 | Lv-Tlr189 |
| e10 | SPU_014638 | Lv-Hypp_2162 |
| e10 | SPU_014914 | Lv-Cts10 |
| e10 | SPU_014981 | Lv-Hypp_105 |

| | | |
|---|---|---|
| e10 | SPU_015002 | Lv-Zmym3 |
| e10 | SPU_015021 | none |
| e10 | SPU_015058 | Lv-Unc44L_106 |
| e10 | SPU_015306 | Lv-Nipa2 |
| e10 | SPU_015339 | NA |
| e10 | SPU_015466 | Lv-Numb |
| e10 | SPU_015580 | none |
| e10 | SPU_015696 | Lv-Hypp_3055 |
| e10 | SPU_015720 | Lv-Samhd1 |
| e10 | SPU_016016 | Lv-Notchl5_1 |
| e10 | SPU_016470 | Lv-PppL_204 |
| e10 | SPU_016517 | none |
| e10 | SPU_016594 | NA |
| e10 | SPU_016864 | Lv-Unk_61 |
| e10 | SPU_016872 | Lv-PppL_205 |
| e10 | SPU_016958 | Lv-Dnajhc30 |
| e10 | SPU_016989 | none |
| e10 | SPU_017246 | Lv-Hypp_3049 |
| e10 | SPU_017307 | Lv-NkrfL |
| e10 | SPU_017314 | none |
| e10 | SPU_017396 | none |
| e10 | SPU_017427 | Lv-Z155 |
| e10 | SPU_017590 | Lv-Hypp_2346 |
| e10 | SPU_017630 | Lv-Hypp_885 |
| e10 | SPU_017641 | Lv-Hypp_887 |
| e10 | SPU_017657 | NA |
| e10 | SPU_017724 | none |
| e10 | SPU_017930 | none |
| e10 | SPU_017932 | Lv-PppL_211 |
| e10 | SPU_017947 | Lv-Z116 |
| e10 | SPU_017949 | Lv-Plk1 |
| e10 | SPU_018142 | Lv-Trpm3L |
| e10 | SPU_018315 | Lv-CspeN1a |
| e10 | SPU_018331 | Lv-Transaa_1 |
| e10 | SPU_018446 | Lv-Scnn1g_1 |
| e10 | SPU_018566 | Lv-Hypp_917 |
| e10 | SPU_018612 | NA |
| e10 | SPU_018740 | Lv-Hypp_2404 |

| | | |
|---|---|---|
| e10 | SPU_018761 | NA |
| e10 | SPU_018871 | Lv-Hypp_2409 |
| e10 | SPU_018946 | Lv-TpaseL_11 |
| e10 | SPU_019055 | Lv-Hypp_926 |
| e10 | SPU_019187 | Lv-Rxfp2_2 |
| e10 | SPU_019299 | Lv-C1orf101L |
| e10 | SPU_019550 | Lv-Scrnph1 |
| e10 | SPU_019611 | Lv-Jarid1aL |
| e10 | SPU_019797 | Lv-Hypp_2466 |
| e10 | SPU_019815 | none |
| e10 | SPU_019846 | Lv-Sepn1 |
| e10 | SPU_020077 | Lv-Hypp_2483 |
| e10 | SPU_020154 | none |
| e10 | SPU_020181 | Lv-Vwdf_1 |
| e10 | SPU_020596 | Lv-Actb_3 |
| e10 | SPU_020727 | NA |
| e10 | SPU_020746 | Lv-Nol10 |
| e10 | SPU_020798 | none |
| e10 | SPU_020812 | Lv-Tuba1c |
| e10 | SPU_020894 | Lv-Hypp_1002 |
| e10 | SPU_020998 | NA |
| e10 | SPU_021129 | Lv-Hypp_1012 |
| e10 | SPU_021154 | Lv-Hsp40F |
| e10 | SPU_021194 | Lv-TlrP23 |
| e10 | SPU_021225 | Lv-Tlr109 |
| e10 | SPU_021253 | Lv-Fn3/Egf/Ccp/HyalinL |
| e10 | SPU_021266 | Lv-PppL_126 |
| e10 | SPU_021338 | Lv-Hypp_1022 |
| e10 | SPU_021387 | Lv-Aqp1 |
| e10 | SPU_021445 | none |
| e10 | SPU_021557 | Lv-Gabp |
| e10 | SPU_021575 | Lv-Hypp_2562 |
| e10 | SPU_021577 | none |
| e10 | SPU_021607 | Lv-HnmtL_2 |
| e10 | SPU_021699 | NA |
| e10 | SPU_021838 | Lv-Non/MmyIIhcph |
| e10 | SPU_021969 | Lv-DomiL |
| e10 | SPU_022033 | none |

| | | |
|---|---|---|
| e10 | SPU_022037 | Lv-Hypp_2595 |
| e10 | SPU_022056 | Lv-Ciz1 |
| e10 | SPU_022169 | Lv-Fam55c_5 |
| e10 | SPU_022235 | Lv-Samhd1_1 |
| e10 | SPU_022274 | Lv-Bcl9 |
| e10 | SPU_022302 | Lv-Aji |
| e10 | SPU_022528 | Lv-Srcr193 |
| e10 | SPU_022836 | Lv-Fam86 |
| e10 | SPU_023011 | Lv-Slc39a8 |
| e10 | SPU_023047 | Lv-Adrb3L_1 |
| e10 | SPU_023197 | Lv-NudE |
| e10 | SPU_023259 | none |
| e10 | SPU_023549 | NA |
| e10 | SPU_023673 | Lv-EndrvtA |
| e10 | SPU_023788 | Lv-Tcp11 |
| e10 | SPU_024011 | Lv-Chst11L_3 |
| e10 | SPU_024091 | none |
| e10 | SPU_024127 | Lv-Clect_45 |
| e10 | SPU_024129 | Lv-Phf17 |
| e10 | SPU_024150 | Lv-Fbxo9 |
| e10 | SPU_024317 | none |
| e10 | SPU_024462 | Lv-RtL_6 |
| e10 | SPU_024569 | none |
| e10 | SPU_024880 | NA |
| e10 | SPU_024896 | Lv-Ercc6b |
| e10 | SPU_024903 | Lv-Ese |
| e10 | SPU_024983 | Lv-Maf1 |
| e10 | SPU_025175 | Lv-Tuba3 |
| e10 | SPU_025193 | none |
| e10 | SPU_025302 | Lv-Alx1 |
| e10 | SPU_025496 | Lv-Hypp_231 |
| e10 | SPU_025590 | Lv-FoxM |
| e10 | SPU_025615 | Lv-Rbl1_1 |
| e10 | SPU_025720 | Lv-Larp4L |
| e10 | SPU_025767 | Lv-Kins8l |
| e10 | SPU_025858 | Lv-Slc35a1 |
| e10 | SPU_025869 | Lv-Hypp_234 |
| e10 | SPU_025998 | none |

| | | |
|---|---|---|
| e10 | SPU_026017 | none |
| e10 | SPU_026022 | Lv-Hypp_2813 |
| e10 | SPU_026117 | Lv-Lrp1L_3 |
| e10 | SPU_026145 | Lv-Endrvt_2 |
| e10 | SPU_026250 | Lv-Zfc3h1 |
| e10 | SPU_026251 | none |
| e10 | SPU_026261 | Lv-Amy_5 |
| e10 | SPU_026277 | Lv-Wnt5 |
| e10 | SPU_026356 | Lv-Ankrd12 |
| e10 | SPU_026573 | Lv-Klkb1L |
| e10 | SPU_026603 | Lv-Hypp_2858 |
| e10 | SPU_026727 | Lv-Fam55d_1 |
| e10 | SPU_026886 | Lv-Rbm12L |
| e10 | SPU_026958 | none |
| e10 | SPU_027105 | Lv-PppL_283 |
| e10 | SPU_027383 | none |
| e10 | SPU_027595 | Lv-Agl_1 |
| e10 | SPU_027774 | Lv-Srf |
| e10 | SPU_027816 | Lv-Tdo2_1 |
| e10 | SPU_027851 | none |
| e10 | SPU_028303 | Lv-HelbL |
| e10 | SPU_028435 | Lv-Hypp_2955 |
| e10 | SPU_028481 | Lv-Hypp_285 |
| e10 | SPU_028577 | Lv-Hypp_1234 |
| e10 | SPU_028676 | Lv-Klhl2 |
| e10 | SPU_028831 | none |
| e10 | SPU_030133 | Lv-TlrP77 |
| e11 | SPU_000049 | Lv-TrypL |
| e11 | SPU_000082 | Lv-Pif1h |
| e11 | SPU_000233 | Lv-Reo_6L |
| e11 | SPU_000243 | Lv-Wdr60 |
| e11 | SPU_000298 | none |
| e11 | SPU_000445 | Lv-EhH2a |
| e11 | SPU_000545 | Lv-Gtf3c1L |
| e11 | SPU_000590 | Lv-Adhfe1 |
| e11 | SPU_000591 | Lv-Adhfe1_1 |
| e11 | SPU_000640 | Lv-Hypp_1268 |
| e11 | SPU_000645 | Lv-Cyp27L_1 |

| | | |
|---|---|---|
| e11 | SPU_000669 | Lv-Bmp2/4 |
| e11 | SPU_000870 | none |
| e11 | SPU_001232 | Lv-Nin |
| e11 | SPU_001249 | Lv-Unc44L_10 |
| e11 | SPU_001336 | Lv-Mamdc2 |
| e11 | SPU_001403 | none |
| e11 | SPU_001411 | Lv-Hypp_1325 |
| e11 | SPU_001488 | Lv-C20orf54L |
| e11 | SPU_001499 | Lv-Chst2L_2 |
| e11 | SPU_001552 | Lv-Kiaa1409L |
| e11 | SPU_001685 | Lv-Unc44L_17 |
| e11 | SPU_002101 | Lv-Kiaa1529L |
| e11 | SPU_002268 | Lv-Unc44_2651 |
| e11 | SPU_002378 | none |
| e11 | SPU_002386 | none |
| e11 | SPU_002468 | Lv-Unc44L_25 |
| e11 | SPU_002535 | Lv-Lrguk |
| e11 | SPU_002552 | Lv-Stxbp1 |
| e11 | SPU_002562 | Lv-Kiaa1751L |
| e11 | SPU_002602 | NA |
| e11 | SPU_002738 | Lv-Ccdc92 |
| e11 | SPU_002780 | Lv-Ndufs4 |
| e11 | SPU_002792 | Lv-Socs2/3 |
| e11 | SPU_002809 | Lv-Lhh2bj |
| e11 | SPU_002966 | Lv-Slc25a42L |
| e11 | SPU_003015 | Lv-Chrm3L_3 |
| e11 | SPU_003063 | Lv-Zc3p26_1 |
| e11 | SPU_003123 | Lv-Wdr16 |
| e11 | SPU_003156 | none |
| e11 | SPU_003157 | NA |
| e11 | SPU_003188 | NA |
| e11 | SPU_003218 | Lv-PolypL_34 |
| e11 | SPU_003244 | none |
| e11 | SPU_003461 | Lv-Unk |
| e11 | SPU_003564 | NA |
| e11 | SPU_003654 | Lv-Pftaire-1 |
| e11 | SPU_003865 | Lv-Pf2L1 |
| e11 | SPU_003888 | Lv-Ccdc89 |

| | | |
|---|---|---|
| e11 | SPU_003929 | Lv-Gal/Lect/Lnb/7Tm/Gpcr_1 |
| e11 | SPU_004012 | Lv-Hypp_7993 |
| e11 | SPU_004121 | NA |
| e11 | SPU_004248 | Lv-Cluap1 |
| e11 | SPU_004466 | Lv-Shroom4L |
| e11 | SPU_004674 | Lv-Hypp_424 |
| e11 | SPU_004685 | Lv-PolppL_38 |
| e11 | SPU_004691 | Lv-Hypp_3082 |
| e11 | SPU_004819 | Lv-Hypp_1545 |
| e11 | SPU_004934 | Lv-Dnah6 |
| e11 | SPU_005132 | Lv-Cub/Clect/Ldla/Gpcr |
| e11 | SPU_005171 | Lv-Hox6 |
| e11 | SPU_005191 | Lv-Rtdr1L |
| e11 | SPU_005300 | Lv-Abcc5D |
| e11 | SPU_005303 | Lv-Fnbp4L |
| e11 | SPU_005354 | none |
| e11 | SPU_005467 | NA |
| e11 | SPU_005482 | Lv-Asah2 |
| e11 | SPU_005501 | Lv-Crispld2L |
| e11 | SPU_005507 | Lv-Ipla2(gamma) |
| e11 | SPU_005511 | Lv-Baxi1 |
| e11 | SPU_005586 | none |
| e11 | SPU_005739 | Lv-Commd6 |
| e11 | SPU_005769 | Lv-Hypp_463 |
| e11 | SPU_005806 | Lv-Slco3a1 |
| e11 | SPU_005975 | Lv-Unc44L_48 |
| e11 | SPU_006078 | NA |
| e11 | SPU_006110 | Lv-Rundc1 |
| e11 | SPU_006278 | NA |
| e11 | SPU_006344 | Lv-Hypp_3013 |
| e11 | SPU_006438 | Lv-Abcc5h |
| e11 | SPU_006503 | none |
| e11 | SPU_006519 | Lv-Lrrc50-2 |
| e11 | SPU_006556 | Lv-Maoa_5 |
| e11 | SPU_006699 | Lv-Ia1/Ic140-1 |
| e11 | SPU_006747 | Lv-Fam154b |
| e11 | SPU_006756 | Lv-Atub9 |
| e11 | SPU_006862 | Lv-Hypp_499 |

| | | |
|---|---|---|
| e11 | SPU_006922 | Lv-TnL2 |
| e11 | SPU_006926 | NA |
| e11 | SPU_007167 | Lv-PppL_189 |
| e11 | SPU_007205 | Lv-Glyt_2 |
| e11 | SPU_007252 | Lv-Ttc29 |
| e11 | SPU_007376 | Lv-Pdk2 |
| e11 | SPU_007743 | none |
| e11 | SPU_007801 | NA |
| e11 | SPU_007843 | Lv-PolppL_63 |
| e11 | SPU_007908 | none |
| e11 | SPU_007924 | Lv-Sult1c |
| e11 | SPU_007926 | NA |
| e11 | SPU_007984 | Lv-Atub8 |
| e11 | SPU_007999 | Lv-Hypp_1759 |
| e11 | SPU_008131 | Lv-PppL_190 |
| e11 | SPU_008203 | Lv-Hypp_1772 |
| e11 | SPU_008246 | none |
| e11 | SPU_008367 | Lv-Hypp_1782 |
| e11 | SPU_008547 | Lv-Nlr50 |
| e11 | SPU_008679 | Lv-Zmym1-2 |
| e11 | SPU_008727 | Lv-Tgm1 |
| e11 | SPU_008792 | NA |
| e11 | SPU_008890 | Lv-Psmf1 |
| e11 | SPU_008989 | none |
| e11 | SPU_008990 | Lv-Gdap2 |
| e11 | SPU_009059 | Lv-Nuf2 |
| e11 | SPU_009131 | Lv-Pnpo |
| e11 | SPU_009260 | Lv-Ift57_1 |
| e11 | SPU_009337 | Lv-Hypp_576 |
| e11 | SPU_009425 | none |
| e11 | SPU_009611 | none |
| e11 | SPU_009671 | Lv-Hypp_1864 |
| e11 | SPU_009695 | none |
| e11 | SPU_009746 | Lv-Endrvt70 |
| e11 | SPU_010002 | Lv-NfascL |
| e11 | SPU_010079 | Lv-Hypp_605 |
| e11 | SPU_010140 | Lv-Arl11l1 |
| e11 | SPU_010169 | Lv-Hypp_1901 |

| | | |
|---|---|---|
| e11 | SPU_010280 | Lv-Ampk |
| e11 | SPU_010316 | none |
| e11 | SPU_010349 | none |
| e11 | SPU_010410 | Lv-Endrvt20 |
| e11 | SPU_010445 | Lv-Ccp/Egf/Tsp1/HyalinL2 |
| e11 | SPU_010469 | none |
| e11 | SPU_010578 | NA |
| e11 | SPU_010686 | Lv-B4galt6_2 |
| e11 | SPU_010728 | Lv-Ccdc11 |
| e11 | SPU_010767 | none |
| e11 | SPU_010993 | Lv-Srcr97 |
| e11 | SPU_011067 | Lv-Hypp_1957 |
| e11 | SPU_011092 | Lv-Akap14 |
| e11 | SPU_011158 | none |
| e11 | SPU_011239 | Lv-Ift80L1 |
| e11 | SPU_011446 | Lv-Bbs1 |
| e11 | SPU_011468 | Lv-Kiaa0141L_1 |
| e11 | SPU_011669 | none |
| e11 | SPU_011699 | Lv-Spcaph |
| e11 | SPU_011700 | Lv-Nelf |
| e11 | SPU_011739 | none |
| e11 | SPU_011882 | none |
| e11 | SPU_011931 | Lv-Ankrd13 |
| e11 | SPU_011952 | Lv-Unk_75 |
| e11 | SPU_011962 | Lv-Ldha |
| e11 | SPU_012032 | Lv-Unk_54 |
| e11 | SPU_012045 | Lv-rsp3h |
| e11 | SPU_012330 | Lv-Iqcd |
| e11 | SPU_012336 | NA |
| e11 | SPU_012356 | Lv-C1orf58 |
| e11 | SPU_012382 | Lv-Clect/Ldla/Lrr/Gpcr |
| e11 | SPU_012467 | Lv-C21orf2L |
| e11 | SPU_012521 | Lv-Nme5 |
| e11 | SPU_012606 | Lv-Ccdc113 |
| e11 | SPU_012648 | Lv-Zhangfei |
| e11 | SPU_012984 | Lv-Endrvt27 |
| e11 | SPU_013001 | Lv-Rt34 |
| e11 | SPU_013085 | Lv-Rt5 |

| | | |
|---|---|---|
| e11 | SPU_013266 | NA |
| e11 | SPU_013742 | Lv-Acad10 |
| e11 | SPU_013764 | none |
| e11 | SPU_013798 | Lv-Adora2a |
| e11 | SPU_013805 | Lv-Armc9 |
| e11 | SPU_013828 | Lv-Fchsd2 |
| e11 | SPU_013998 | Lv-Polyp_1 |
| e11 | SPU_013999 | Lv-Ank1_2 |
| e11 | SPU_014003 | Lv-Pogk |
| e11 | SPU_014067 | Lv-Got2 |
| e11 | SPU_014083 | Lv-Ccdc19 |
| e11 | SPU_014107 | Lv-Spef2_1 |
| e11 | SPU_014269 | Lv-Htr1a |
| e11 | SPU_014285 | Lv-Tubb2a |
| e11 | SPU_014449 | Lv-Aadac_1 |
| e11 | SPU_014531 | none |
| e11 | SPU_014564 | Lv-Stom_1 |
| e11 | SPU_014568 | Lv-Stom_4 |
| e11 | SPU_014609 | none |
| e11 | SPU_014801 | Lv-RshL3 |
| e11 | SPU_014804 | Lv-Man2b2 |
| e11 | SPU_014909 | Lv-Rogdih |
| e11 | SPU_015216 | Lv-Hypp_110 |
| e11 | SPU_015275 | Lv-Hypp_2204 |
| e11 | SPU_015320 | Lv-Ia/Ic28 |
| e11 | SPU_015384 | NA |
| e11 | SPU_015608 | Lv-Tuba1b_1 |
| e11 | SPU_015625 | Lv-Oad_bp58 |
| e11 | SPU_015657 | NA |
| e11 | SPU_015668 | Lv-Cts11 |
| e11 | SPU_015874 | Lv-Polyp_2 |
| e11 | SPU_015928 | Lv-Mdh1b |
| e11 | SPU_016014 | Lv-Tirc1 |
| e11 | SPU_016086 | Lv-Hypp_2246 |
| e11 | SPU_016102 | NA |
| e11 | SPU_016166 | Lv-Wdr13 |
| e11 | SPU_016429 | Lv-Dlec1 |
| e11 | SPU_016483 | Lv-Endrvt30 |

| | | |
|---|---|---|
| e11 | SPU_016490 | Lv-Z145 |
| e11 | SPU_016602 | Lv-PolypL_13 |
| e11 | SPU_016614 | none |
| e11 | SPU_016655 | Lv-Kif19 |
| e11 | SPU_016766 | Lv-Endrvt31 |
| e11 | SPU_016944 | none |
| e11 | SPU_016992 | Lv-C19orf22L |
| e11 | SPU_017044 | Lv-Hypp_858 |
| e11 | SPU_017383 | Lv-PolypL_78 |
| e11 | SPU_017550 | Lv-Hypp_882 |
| e11 | SPU_017584 | Lv-C12orf63L |
| e11 | SPU_017600 | Lv-Rbm10 |
| e11 | SPU_017604 | Lv-Retsat |
| e11 | SPU_017631 | Lv-Hypp_2348 |
| e11 | SPU_017823 | Lv-Unc44_129 |
| e11 | SPU_017892 | Lv-Slc25a36 |
| e11 | SPU_017898 | Lv-Hypp_2360 |
| e11 | SPU_017953 | Lv-Hypp_139 |
| e11 | SPU_017989 | Lv-Mt1 |
| e11 | SPU_017996 | NA |
| e11 | SPU_018100 | Lv-Tlr091 |
| e11 | SPU_018136 | Lv-Notch1-14 |
| e11 | SPU_018419 | Lv-Prkar2a |
| e11 | SPU_018565 | none |
| e11 | SPU_018715 | none |
| e11 | SPU_018783 | none |
| e11 | SPU_018784 | Lv-Unk_106 |
| e11 | SPU_018788 | Lv-C2orf56 |
| e11 | SPU_018822 | NA |
| e11 | SPU_018854 | Lv-Dlc2 |
| e11 | SPU_018906 | none |
| e11 | SPU_018908 | Lv-Six3 |
| e11 | SPU_019032 | Lv-Vps13c |
| e11 | SPU_019248 | Lv-Hypp_2428 |
| e11 | SPU_019258 | Lv-Slc25a14_1 |
| e11 | SPU_019271 | Lv-Hypp_3000 |
| e11 | SPU_019304 | Lv-Ccdc146 |
| e11 | SPU_019456 | Lv-Hypp_2442 |

| | | |
|---|---|---|
| e11 | SPU_019546 | NA |
| e11 | SPU_019553 | Lv-Adk |
| e11 | SPU_019556 | Lv-C6orf199L |
| e11 | SPU_019571 | none |
| e11 | SPU_019598 | Lv-Endrvt101 |
| e11 | SPU_019749 | Lv-Klhl20L |
| e11 | SPU_020048 | NA |
| e11 | SPU_020073 | Lv-Ccdc104 |
| e11 | SPU_020086 | Lv-Hypp_961 |
| e11 | SPU_020157 | Lv-Lrmp |
| e11 | SPU_020420 | Lv-Cubn_1 |
| e11 | SPU_020533 | Lv-Inpp5e |
| e11 | SPU_020605 | Lv-Ndufv1L |
| e11 | SPU_020681 | Lv-Fgfrl1 |
| e11 | SPU_020783 | Lv-Hypp_992 |
| e11 | SPU_020795 | none |
| e11 | SPU_020813 | none |
| e11 | SPU_020925 | Lv-Daam1 |
| e11 | SPU_020956 | Lv-Dmbt1L_8 |
| e11 | SPU_021026 | Lv-Hypp_1009 |
| e11 | SPU_021172 | Lv-Fra2 |
| e11 | SPU_021222 | Lv-Hypp_171 |
| e11 | SPU_021236 | Lv-Rabl4 |
| e11 | SPU_021331 | Lv-Trex2 |
| e11 | SPU_021354 | Lv-Acel_3 |
| e11 | SPU_021369 | Lv-C15orf33L |
| e11 | SPU_021489 | Lv-Hypp_2557 |
| e11 | SPU_021574 | Lv-Hypp_2561 |
| e11 | SPU_021592 | Lv-Cpc1 |
| e11 | SPU_021667 | none |
| e11 | SPU_021709 | Lv-Clect/Pan_12 |
| e11 | SPU_021760 | Lv-Ift172 |
| e11 | SPU_022196 | Lv-Hypp_2602 |
| e11 | SPU_022298 | Lv-Stom_8 |
| e11 | SPU_022389 | none |
| e11 | SPU_022466 | Lv-Hypp_2609 |
| e11 | SPU_022560 | Lv-AnubL1 |
| e11 | SPU_022561 | Lv-Mipphp1 |

| | | |
|---|---|---|
| e11 | SPU_022749 | Lv-C9orf128 |
| e11 | SPU_022860 | Lv-Ank1_3 |
| e11 | SPU_022898 | Lv-Abhd7 |
| e11 | SPU_022925 | Lv-TpaseL_15 |
| e11 | SPU_023027 | Lv-Rad9_1 |
| e11 | SPU_023205 | Lv-GhdcL_2 |
| e11 | SPU_023486 | Lv-FibcystL_1 |
| e11 | SPU_023489 | Lv-Hypp_2673 |
| e11 | SPU_023672 | Lv-EndrvtC |
| e11 | SPU_023775 | Lv-Hypp_206 |
| e11 | SPU_023957 | Lv-Mok1 |
| e11 | SPU_024093 | Lv-Phb2 |
| e11 | SPU_024368 | Lv-PolypL_955 |
| e11 | SPU_024384 | Lv-Aqp3 |
| e11 | SPU_024435 | Lv-Hypp_214 |
| e11 | SPU_024449 | Lv-Oma |
| e11 | SPU_024465 | Lv-Hypp_2735 |
| e11 | SPU_024548 | NA |
| e11 | SPU_024595 | NA |
| e11 | SPU_024605 | Lv-Plakg/GcL |
| e11 | SPU_024616 | NA |
| e11 | SPU_024618 | NA |
| e11 | SPU_024650 | none |
| e11 | SPU_024688 | Lv-Enpp3 |
| e11 | SPU_024768 | none |
| e11 | SPU_024895 | Lv-Aldh7a1 |
| e11 | SPU_024944 | none |
| e11 | SPU_025027 | Lv-Hypp_224 |
| e11 | SPU_025118 | Lv-Nos2L |
| e11 | SPU_025292 | Lv-Cic |
| e11 | SPU_025319 | Lv-PppL_226 |
| e11 | SPU_025370 | NA |
| e11 | SPU_025413 | Lv-C/Mycbp |
| e11 | SPU_025617 | none |
| e11 | SPU_025641 | Lv-Adra1dL_2 |
| e11 | SPU_025808 | none |
| e11 | SPU_025942 | none |
| e11 | SPU_025994 | Lv-C15orf33 |

| | | |
|---|---|---|
| e11 | SPU_026380 | none |
| e11 | SPU_026429 | Lv-Hypp_1178 |
| e11 | SPU_026433 | Lv-C10orf123L |
| e11 | SPU_026444 | none |
| e11 | SPU_026547 | Lv-AnL10b |
| e11 | SPU_026632 | Lv-Iqca1L |
| e11 | SPU_026953 | Lv-C16orf70L |
| e11 | SPU_027016 | Lv-Rt19 |
| e11 | SPU_027020 | Lv-Prpf39 |
| e11 | SPU_027106 | Lv-Hypp_2887 |
| e11 | SPU_027423 | Lv-Mekk1_1 |
| e11 | SPU_027455 | Lv-Hypp_2904 |
| e11 | SPU_027483 | Lv-Hbld2 |
| e11 | SPU_027692 | Lv-Tmem62_1 |
| e11 | SPU_027784 | Lv-Klp6L1 |
| e11 | SPU_027786 | Lv-Unk_85 |
| e11 | SPU_027850 | Lv-Glud1 |
| e11 | SPU_027931 | Lv-Hypp_2922 |
| e11 | SPU_028096 | Lv-St6galnac4_2 |
| e11 | SPU_028169 | Lv-RasO |
| e11 | SPU_028354 | Lv-Pdhb |
| e11 | SPU_028362 | NA |
| e11 | SPU_028373 | Lv-Il16L |
| e11 | SPU_028819 | NA |
| e11 | SPU_028851 | Lv-Tmem68 |
| e11 | SPU_028896 | Lv-Pdpi(cyclophilin)L8 |
| e11 | SPU_030223 | Lv-Axndhc1h |
| e11 | SPU_030224 | Lv-Axndhc2h |
| e11 | SPU_030227 | Lv-Axndhc6h |
| e11 | SPU_030230 | Lv-Axndhc9h |
| e11 | SPU_030231 | Lv-Axndhc10h |
| e11 | SPU_030232 | Lv-Axndhc12h |
| e12 | SPU_000003 | Lv-Hypp_1239 |
| e12 | SPU_000042 | Lv-Krfp1 |
| e12 | SPU_000044 | Lv-Lrps5l |
| e12 | SPU_000135 | Lv-Zf(mynd)_13L |
| e12 | SPU_000253 | Lv-RbjL1 |
| e12 | SPU_000488 | Lv-C11orf65L |

| | | |
|---|---|---|
| e12 | SPU_000628 | Lv-Ankrd53 |
| e12 | SPU_000632 | Lv-C10orf134L |
| e12 | SPU_000739 | Lv-Smad5 |
| e12 | SPU_001075 | none |
| e12 | SPU_001257 | Lv-Hypp_328 |
| e12 | SPU_001293 | Lv-Cpeb2 |
| e12 | SPU_001395 | Lv-Hypp_3113 |
| e12 | SPU_001448 | Lv-Hypp_1329 |
| e12 | SPU_001717 | Lv-Kiaa0895L |
| e12 | SPU_001824 | Lv-Hypp_3107 |
| e12 | SPU_001985 | NA |
| e12 | SPU_002011 | Lv-Dcdc2 |
| e12 | SPU_002097 | Lv-Hypp_1369 |
| e12 | SPU_002121 | Lv-Hypp_343 |
| e12 | SPU_002175 | none |
| e12 | SPU_002180 | Lv-Hypp_346 |
| e12 | SPU_002372 | Lv-Nlr55 |
| e12 | SPU_002494 | Lv-Hypp_356 |
| e12 | SPU_002517 | none |
| e12 | SPU_002568 | NA |
| e12 | SPU_002651 | Lv-Hypp_361 |
| e12 | SPU_002678 | NA |
| e12 | SPU_002775 | Lv-Ccdc83 |
| e12 | SPU_002820 | Lv-Fn3_18 |
| e12 | SPU_002877 | Lv-Unken3-5dL |
| e12 | SPU_002944 | Lv-CalcneurL |
| e12 | SPU_003044 | Lv-Iqsec1 |
| e12 | SPU_003068 | Lv-Wdr88 |
| e12 | SPU_003074 | Lv-Lrrc69 |
| e12 | SPU_003329 | Lv-Slc44a2 |
| e12 | SPU_003409 | Lv-C19orf45L |
| e12 | SPU_003608 | Lv-Tirc5 |
| e12 | SPU_003909 | Lv-Rt23 |
| e12 | SPU_003918 | Lv-Lrp8 |
| e12 | SPU_004261 | Lv-Ncf2l |
| e12 | SPU_004299 | NA |
| e12 | SPU_004360 | Lv-Tlr022 |
| e12 | SPU_004437 | Lv-Tmc5 |

| | | |
|---|---|---|
| e12 | SPU_004530 | Lv-FbsL_4 |
| e12 | SPU_004585 | none |
| e12 | SPU_004762 | Lv-Odadc2L1 |
| e12 | SPU_004882 | Lv-C2orf77L |
| e12 | SPU_004945 | Lv-Sea/Hyr/Clect |
| e12 | SPU_004989 | Lv-Brwd2_2 |
| e12 | SPU_005169 | Lv-Hox5 |
| e12 | SPU_005287 | Lv-ArsbL |
| e12 | SPU_005312 | Lv-Bbs7 |
| e12 | SPU_005408 | Lv-Armc2 |
| e12 | SPU_005442 | none |
| e12 | SPU_005757 | Lv-Slc16a14 |
| e12 | SPU_006050 | Lv-TpaseL_27 |
| e12 | SPU_006285 | Lv-Hypp_16 |
| e12 | SPU_006343 | none |
| e12 | SPU_006353 | Lv-Pde4Bi1 |
| e12 | SPU_006464 | Lv-Dmbt1L_3 |
| e12 | SPU_006598 | Lv-Lrrc48 |
| e12 | SPU_006629 | NA |
| e12 | SPU_006884 | Lv-Sea/Egf_3 |
| e12 | SPU_007128 | Lv-Hypp_1698 |
| e12 | SPU_007140 | Lv-Wd66 |
| e12 | SPU_007211 | NA |
| e12 | SPU_007351 | Lv-Rt27 |
| e12 | SPU_007463 | Lv-Lrr3 |
| e12 | SPU_007504 | Lv-Xcat2_1 |
| e12 | SPU_007632 | Lv-Rt28 |
| e12 | SPU_007671 | none |
| e12 | SPU_007771 | Lv-Flj46688l1_1 |
| e12 | SPU_007786 | Lv-Hypp_1746 |
| e12 | SPU_007821 | none |
| e12 | SPU_008067 | none |
| e12 | SPU_008188 | Lv-Unk_99 |
| e12 | SPU_008243 | Lv-C10orf79 |
| e12 | SPU_008285 | Lv-Slc6a7 |
| e12 | SPU_008322 | none |
| e12 | SPU_008426 | Lv-Rvt/Exn/End/Pp |
| e12 | SPU_008545 | Lv-PolppL_75 |

| | | |
|---|---|---|
| e12 | SPU_008645 | Lv-Pafah1L |
| e12 | SPU_008731 | none |
| e12 | SPU_008830 | Lv-Otud3_1 |
| e12 | SPU_009119 | Lv-PppL_252 |
| e12 | SPU_009397 | Lv-PolppL_81 |
| e12 | SPU_009400 | Lv-KinsC |
| e12 | SPU_009484 | none |
| e12 | SPU_009543 | Lv-Hypp_580 |
| e12 | SPU_009561 | Lv-Wdr78_1 |
| e12 | SPU_009640 | Lv-Tsnaxip1 |
| e12 | SPU_009686 | none |
| e12 | SPU_009853 | NA |
| e12 | SPU_009930 | Lv-LrpL_6 |
| e12 | SPU_009932 | Lv-Heldr5L |
| e12 | SPU_010008 | Lv-Hypp_1890 |
| e12 | SPU_010134 | Lv-Hypp_62 |
| e12 | SPU_010165 | NA |
| e12 | SPU_010341 | none |
| e12 | SPU_010493 | Lv-Endrvt76 |
| e12 | SPU_010637 | Lv-Wdr35 |
| e12 | SPU_010718 | Lv-EgfiL_3 |
| e12 | SPU_010731 | Lv-Ube2q1L |
| e12 | SPU_010744 | none |
| e12 | SPU_010891 | none |
| e12 | SPU_010905 | Lv-Hypp_632 |
| e12 | SPU_010928 | Lv-Endrvt22 |
| e12 | SPU_010992 | Lv-Srcr96 |
| e12 | SPU_011153 | Lv-Hypp_641 |
| e12 | SPU_011179 | Lv-Wdr52-2 |
| e12 | SPU_011765 | none |
| e12 | SPU_011769 | Lv-Hypp_659 |
| e12 | SPU_011781 | Lv-Fic3 |
| e12 | SPU_011805 | Lv-C20orf54_1 |
| e12 | SPU_012023 | Lv-Hypp_2013 |
| e12 | SPU_012115 | Lv-Efhb |
| e12 | SPU_012139 | Lv-Unk_100 |
| e12 | SPU_012151 | none |
| e12 | SPU_012178 | none |

| | | |
|---|---|---|
| e12 | SPU_012193 | Lv-Krt78L_2 |
| e12 | SPU_012223 | Lv-Fam167a |
| e12 | SPU_012290 | Lv-Dlec1L |
| e12 | SPU_012328 | Lv-Tpcn1 |
| e12 | SPU_012446 | Lv-Gpr50L_1 |
| e12 | SPU_012450 | none |
| e12 | SPU_012575 | Lv-Lhh2ae |
| e12 | SPU_012628 | Lv-Atp6v1d |
| e12 | SPU_012697 | NA |
| e12 | SPU_012726 | Lv-Tst |
| e12 | SPU_012730 | Lv-Hypp_691 |
| e12 | SPU_012809 | Lv-Ia1/Ic140-2 |
| e12 | SPU_012823 | Lv-Zf(cchc)_26 |
| e12 | SPU_012879 | Lv-Hypp_83 |
| e12 | SPU_012923 | Lv-C10orf93 |
| e12 | SPU_013103 | Lv-Spag17 |
| e12 | SPU_013119 | Lv-Sage_1 |
| e12 | SPU_013177 | Lv-Kiaa0895L_1 |
| e12 | SPU_013199 | Lv-Cyp2L27 |
| e12 | SPU_013295 | Lv-Iqca1 |
| e12 | SPU_013315 | Lv-Caps2 |
| e12 | SPU_013321 | Lv-Lrrc50 |
| e12 | SPU_013348 | none |
| e12 | SPU_013349 | Lv-Shpk |
| e12 | SPU_013385 | none |
| e12 | SPU_013487 | Lv-Kirrel2L_3 |
| e12 | SPU_013538 | NA |
| e12 | SPU_013539 | Lv-Slc39a13 |
| e12 | SPU_013562 | none |
| e12 | SPU_013875 | Lv-C6orf165L |
| e12 | SPU_013890 | Lv-Rpgrip1L |
| e12 | SPU_014008 | Lv-C16orf48L |
| e12 | SPU_014014 | Lv-Xpf |
| e12 | SPU_014194 | Lv-Hypp_741 |
| e12 | SPU_014245 | Lv-Unk_76 |
| e12 | SPU_014414 | Lv-Hypp_99 |
| e12 | SPU_014529 | Lv-Klhl2_2 |
| e12 | SPU_014743 | Lv-Adrb2L_5 |

| | | |
|---|---|---|
| e12 | SPU_014750 | none |
| e12 | SPU_014805 | none |
| e12 | SPU_015276 | Lv-Ubq/RL40fp |
| e12 | SPU_015280 | Lv-Hypp_778 |
| e12 | SPU_015332 | NA |
| e12 | SPU_015356 | Lv-Btbd16 |
| e12 | SPU_015389 | none |
| e12 | SPU_015515 | Lv-Zmym1L_2 |
| e12 | SPU_015549 | Lv-Fbxo15 |
| e12 | SPU_015628 | Lv-Hypp_793 |
| e12 | SPU_015692 | none |
| e12 | SPU_015767 | Lv-Z57 |
| e12 | SPU_015915 | NA |
| e12 | SPU_015976 | Lv-Fbxw7L |
| e12 | SPU_016103 | Lv-Clect_108 |
| e12 | SPU_016156 | none |
| e12 | SPU_016246 | Lv-Hypp_2254 |
| e12 | SPU_016258 | Lv-Ccdc40-2 |
| e12 | SPU_016411 | Lv-Ptprorph1 |
| e12 | SPU_016480 | Lv-Unc44_312 |
| e12 | SPU_016601 | Lv-PolypL_12 |
| e12 | SPU_016737 | Lv-Hypp_2273 |
| e12 | SPU_016738 | none |
| e12 | SPU_016930 | Lv-Zswim2 |
| e12 | SPU_016931 | NA |
| e12 | SPU_017012 | Lv-Hypp_2298 |
| e12 | SPU_017013 | Lv-Wdr52 |
| e12 | SPU_017572 | Lv-Xpg_1 |
| e12 | SPU_017648 | Lv-Hypp_137 |
| e12 | SPU_017658 | Lv-Hypp_2997 |
| e12 | SPU_017780 | Lv-Lrrc67 |
| e12 | SPU_017895 | none |
| e12 | SPU_017987 | Lv-Arl6 |
| e12 | SPU_018018 | Lv-C4orf47 |
| e12 | SPU_018028 | Lv-Bbs5 |
| e12 | SPU_018124 | Lv-Hypp_2370 |
| e12 | SPU_018135 | Lv-Hypp_908 |
| e12 | SPU_018137 | NA |

| | | |
|---|---|---|
| e12 | SPU_018233 | Lv-Ttc26 |
| e12 | SPU_018243 | Lv-Dnl4 |
| e12 | SPU_018280 | Lv-Rt11 |
| e12 | SPU_018337 | Lv-Ift74 |
| e12 | SPU_018447 | Lv-Ttl13 |
| e12 | SPU_018584 | Lv-Hypp_2392 |
| e12 | SPU_018587 | Lv-Dcdc5 |
| e12 | SPU_018608 | Lv-Lrr/Igr_11 |
| e12 | SPU_018618 | NA |
| e12 | SPU_018714 | Lv-Hypp_2402 |
| e12 | SPU_018789 | Lv-Hypp_921 |
| e12 | SPU_018962 | Lv-Pmm1 |
| e12 | SPU_018965 | none |
| e12 | SPU_019106 | Lv-Slsp_7 |
| e12 | SPU_019120 | Lv-Opn5L |
| e12 | SPU_019139 | Lv-Anxa7 |
| e12 | SPU_019155 | none |
| e12 | SPU_019198 | none |
| e12 | SPU_019301 | Lv-Pap |
| e12 | SPU_019310 | Lv-Hypp_152 |
| e12 | SPU_019531 | Lv-Ccdc63_1 |
| e12 | SPU_019561 | Lv-Rtkn |
| e12 | SPU_019654 | Lv-Hypp_2457 |
| e12 | SPU_019722 | Lv-Cry2 |
| e12 | SPU_019729 | Lv-Hypp_158 |
| e12 | SPU_019778 | Lv-Cdipt_1 |
| e12 | SPU_019958 | none |
| e12 | SPU_019997 | Lv-Mgrn1 |
| e12 | SPU_020198 | Lv-Rgs22L |
| e12 | SPU_020526 | Lv-Ccdc39 |
| e12 | SPU_020622 | none |
| e12 | SPU_020641 | NA |
| e12 | SPU_020719 | Lv-C1orf201 |
| e12 | SPU_020799 | Lv-ScavrB2L1 |
| e12 | SPU_020850 | Lv-B4galt6_5 |
| e12 | SPU_020863 | none |
| e12 | SPU_020991 | none |
| e12 | SPU_021089 | Lv-Hypp_169 |

| | | |
|---|---|---|
| e12 | SPU_021320 | Lv-Rt40 |
| e12 | SPU_021384 | none |
| e12 | SPU_021548 | Lv-Bctg1 |
| e12 | SPU_021668 | Lv-Atub10 |
| e12 | SPU_021843 | Lv-Z64 |
| e12 | SPU_021847 | Lv-Hrppk |
| e12 | SPU_021913 | NA |
| e12 | SPU_021944 | Lv-Kiaa1095L |
| e12 | SPU_022138 | Lv-CubnL_5 |
| e12 | SPU_022155 | none |
| e12 | SPU_022183 | Lv-Shprh |
| e12 | SPU_022198 | Lv-Hypp_2603 |
| e12 | SPU_022224 | none |
| e12 | SPU_022249 | Lv-Klhl9 |
| e12 | SPU_022256 | Lv-Morn1 |
| e12 | SPU_022266 | none |
| e12 | SPU_022268 | Lv-Ankrd5 |
| e12 | SPU_022343 | none |
| e12 | SPU_022418 | Lv-Hypp_2608 |
| e12 | SPU_022700 | Lv-PppL_277 |
| e12 | SPU_022830 | Lv-Hypp_2634 |
| e12 | SPU_022939 | none |
| e12 | SPU_023192 | none |
| e12 | SPU_023213 | Lv-Hypp_2656 |
| e12 | SPU_023354 | none |
| e12 | SPU_023410 | Lv-Hypp_1085 |
| e12 | SPU_023438 | none |
| e12 | SPU_023624 | Lv-Fbxw10L |
| e12 | SPU_023714 | Lv-Clect_113 |
| e12 | SPU_023959 | NA |
| e12 | SPU_023974 | Lv-Granln |
| e12 | SPU_024050 | Lv-Lrriq4 |
| e12 | SPU_024240 | NA |
| e12 | SPU_024270 | Lv-PppL_137 |
| e12 | SPU_024451 | Lv-Hypp_2733 |
| e12 | SPU_024582 | Lv-Tmem26_5 |
| e12 | SPU_024766 | Lv-Unk_32 |
| e12 | SPU_024767 | NA |

| | | |
|---|---|---|
| e12 | SPU_024771 | Lv-Hypp_220 |
| e12 | SPU_024797 | Lv-Trappc4 |
| e12 | SPU_024940 | none |
| e12 | SPU_024980 | none |
| e12 | SPU_025111 | Lv-Hypp_1137 |
| e12 | SPU_025368 | Lv-Redt_Aldo/Keto |
| e12 | SPU_025457 | Lv-TpaseL_18 |
| e12 | SPU_025528 | Lv-Zmynd12 |
| e12 | SPU_025593 | none |
| e12 | SPU_025787 | Lv-Spag6 |
| e12 | SPU_025876 | Lv-Hypp_235 |
| e12 | SPU_025920 | Lv-Hypp_3120 |
| e12 | SPU_025939 | Lv-C7orf67L |
| e12 | SPU_025995 | none |
| e12 | SPU_026067 | Lv-Ctl1 |
| e12 | SPU_026134 | Lv-C6orf60 |
| e12 | SPU_026146 | Lv-C1orf177L |
| e12 | SPU_026192 | Lv-aminotransferase |
| e12 | SPU_026212 | Lv-Aldh3a2 |
| e12 | SPU_026218 | Lv-Hypp_2828 |
| e12 | SPU_026381 | Lv-Pkdh1l1 |
| e12 | SPU_026388 | none |
| e12 | SPU_026523 | none |
| e12 | SPU_026545 | none |
| e12 | SPU_026569 | Lv-Notch1-21 |
| e12 | SPU_026629 | Lv-Fibulin |
| e12 | SPU_026665 | NA |
| e12 | SPU_026691 | Lv-C14orf148L |
| e12 | SPU_026735 | Lv-Slmoh2 |
| e12 | SPU_026757 | none |
| e12 | SPU_026768 | none |
| e12 | SPU_026775 | Lv-Thap9L |
| e12 | SPU_026840 | Lv-Smpd3L |
| e12 | SPU_026842 | none |
| e12 | SPU_026895 | none |
| e12 | SPU_026977 | none |
| e12 | SPU_027198 | Lv-Hypp_2894 |
| e12 | SPU_027227 | Lv-Ift20 |

| | | |
|---|---|---|
| e12 | SPU_027317 | NA |
| e12 | SPU_027371 | Lv-Vlgr1 |
| e12 | SPU_027448 | Lv-Ppox |
| e12 | SPU_027579 | Lv-Rib43A |
| e12 | SPU_027594 | NA |
| e12 | SPU_027711 | NA |
| e12 | SPU_027818 | Lv-Ctso |
| e12 | SPU_027893 | Lv-Iqcg |
| e12 | SPU_028185 | Lv-Pollp |
| e12 | SPU_028270 | none |
| e12 | SPU_028605 | none |
| e12 | SPU_028607 | Lv-Bpr |
| e12 | SPU_030060 | Lv-Pu1 |
| e12 | SPU_030233 | Lv-Axndhch |
| e13 | SPU_000026 | Lv-Adsl |
| e13 | SPU_000068 | NA |
| e13 | SPU_000075 | Lv-Methiapd2L |
| e13 | SPU_000084 | Lv-Tbcd |
| e13 | SPU_000104 | Lv-Trim33l |
| e13 | SPU_000121 | NA |
| e13 | SPU_000229 | Lv-CopineC |
| e13 | SPU_000232 | none |
| e13 | SPU_000287 | Lv-Vparpf_3 |
| e13 | SPU_000311 | Lv-Gemin5 |
| e13 | SPU_000327 | Lv-Unc44L_2 |
| e13 | SPU_000329 | NA |
| e13 | SPU_000382 | Lv-Man1b1L |
| e13 | SPU_000477 | Lv-Hypp_304 |
| e13 | SPU_000502 | Lv-Upf3 |
| e13 | SPU_000510 | Lv-Gltscr2L |
| e13 | SPU_000586 | Lv-Hypp_1263 |
| e13 | SPU_000627 | Lv-SnrpG |
| e13 | SPU_000648 | Lv-Urb1h |
| e13 | SPU_000771 | Lv-B3galt1 |
| e13 | SPU_000801 | Lv-Smarcc2 |
| e13 | SPU_000821 | Lv-Pycr1L |
| e13 | SPU_000842 | Lv-Kpnb3 |
| e13 | SPU_000865 | Lv-Trrap_1 |

| | | |
|---|---|---|
| e13 | SPU_000884 | Lv-Ttrspn |
| e13 | SPU_000904 | Lv-Ter1 |
| e13 | SPU_000966 | Lv-Timm17B |
| e13 | SPU_000973 | NA |
| e13 | SPU_000995 | Lv-Ghr |
| e13 | SPU_001041 | Lv-B3galt1_16 |
| e13 | SPU_001062 | Lv-ElgC |
| e13 | SPU_001224 | Lv-Hypp_1316 |
| e13 | SPU_001261 | Lv-Mia3 |
| e13 | SPU_001335 | Lv-AtripL |
| e13 | SPU_001366 | Lv-Arp1/Centph |
| e13 | SPU_001390 | Lv-Hdac2 |
| e13 | SPU_001435 | none |
| e13 | SPU_001457 | Lv-Med2 |
| e13 | SPU_001478 | NA |
| e13 | SPU_001586 | Lv-Hsp902A |
| e13 | SPU_001632 | none |
| e13 | SPU_001733 | Lv-Ubap2L |
| e13 | SPU_001786 | Lv-Nup133 |
| e13 | SPU_001801 | Lv-Fut6-2 |
| e13 | SPU_001813 | Lv-PolppL-8 |
| e13 | SPU_001924 | Lv-Cant1 |
| e13 | SPU_002194 | Lv-Dkc1 |
| e13 | SPU_002301 | Lv-Sec61a2 |
| e13 | SPU_002330 | NA |
| e13 | SPU_002338 | Lv-Ext2 |
| e13 | SPU_002355 | Lv-Sea/Egf/Hyalinl |
| e13 | SPU_002366 | Lv-Ssh_1 |
| e13 | SPU_002478 | Lv-Ddx51 |
| e13 | SPU_002518 | none |
| e13 | SPU_002588 | Lv-Cenpj |
| e13 | SPU_002637 | Lv-Dnad_rnapIIppE |
| e13 | SPU_002715 | Lv-Tmem161bh |
| e13 | SPU_002730 | NA |
| e13 | SPU_002789 | Lv-Ppie_1 |
| e13 | SPU_002790 | Lv-Ppie_2 |
| e13 | SPU_002799 | Lv-Hypp_1413 |
| e13 | SPU_002920 | Lv-Smc6-3 |

| | | |
|---|---|---|
| e13 | SPU_002942 | Lv-Rwdd1 |
| e13 | SPU_002949 | Lv-Clca1-4 |
| e13 | SPU_003028 | Lv-PolppL_21 |
| e13 | SPU_003039 | NA |
| e13 | SPU_003073 | Lv-Urod |
| e13 | SPU_003107 | Lv-Msy |
| e13 | SPU_003225 | none |
| e13 | SPU_003233 | Lv-Smarcd1L |
| e13 | SPU_003279 | Lv-Mterfd1 |
| e13 | SPU_003290 | Lv-Rkhd |
| e13 | SPU_003323 | none |
| e13 | SPU_003383 | Lv-Gal3st1L_3 |
| e13 | SPU_003387 | Lv-Ercc8 |
| e13 | SPU_003415 | Lv-Ampd2 |
| e13 | SPU_003488 | Lv-Wdhd1_2 |
| e13 | SPU_003541 | Lv-Th0c3 |
| e13 | SPU_003582 | Lv-Herc1 |
| e13 | SPU_003591 | Lv-Nan2 |
| e13 | SPU_003627 | NA |
| e13 | SPU_003683 | Lv-Nup85 |
| e13 | SPU_003719 | Lv-Recql |
| e13 | SPU_003724 | Lv-Cnot2 |
| e13 | SPU_003754 | none |
| e13 | SPU_003879 | Lv-Hdac3 |
| e13 | SPU_003899 | Lv-Hypp_2208 |
| e13 | SPU_003949 | Lv-Hypp_402 |
| e13 | SPU_004062 | Lv-Sfrp3/4l |
| e13 | SPU_004077 | none |
| e13 | SPU_004078 | Lv-Hpse |
| e13 | SPU_004118 | Lv-Rgp1h |
| e13 | SPU_004130 | Lv-Tmf1 |
| e13 | SPU_004154 | Lv-Nle1 |
| e13 | SPU_004195 | Lv-Sec22_2 |
| e13 | SPU_004219 | Lv-Hypp_406 |
| e13 | SPU_004247 | Lv-Sin |
| e13 | SPU_004262 | none |
| e13 | SPU_004277 | Lv-Polr1b |
| e13 | SPU_004386 | Lv-Hypp_416 |

| | | |
|---|---|---|
| e13 | SPU_004403 | Lv-Hnrpl |
| e13 | SPU_004540 | Lv-Hsp70bp |
| e13 | SPU_004545 | Lv-Fubp1 |
| e13 | SPU_004653 | none |
| e13 | SPU_004808 | Lv-Hypp_430 |
| e13 | SPU_004814 | Lv-Rio |
| e13 | SPU_004843 | Lv-HlfL |
| e13 | SPU_004975 | Lv-Chk2 |
| e13 | SPU_005163 | Lv-Cacna1h |
| e13 | SPU_005434 | Lv-HnrpcL_1 |
| e13 | SPU_005519 | Lv-Arl13bl |
| e13 | SPU_005520 | none |
| e13 | SPU_005550 | Lv-Ttc27 |
| e13 | SPU_005615 | Lv-Pofut1-3 |
| e13 | SPU_005628 | Lv-TardbpL1 |
| e13 | SPU_005659 | Lv-Eaf2L |
| e13 | SPU_005667 | Lv-Mcm10 |
| e13 | SPU_005679 | none |
| e13 | SPU_005681 | Lv-Cherp_1 |
| e13 | SPU_005694 | NA |
| e13 | SPU_005712 | Lv-Ftsjd1 |
| e13 | SPU_005788 | Lv-Mrpl45 |
| e13 | SPU_005795 | Lv-Qtrt1 |
| e13 | SPU_005861 | Lv-Zmym1L_4 |
| e13 | SPU_005987 | Lv-Nuc205 |
| e13 | SPU_006031 | Lv-Ernst1 |
| e13 | SPU_006113 | Lv-Safb2L |
| e13 | SPU_006180 | Lv-Nsun5 |
| e13 | SPU_006181 | Lv-Cstf1 |
| e13 | SPU_006210 | Lv-Impad1 |
| e13 | SPU_006275 | none |
| e13 | SPU_006414 | Lv-Pc |
| e13 | SPU_006457 | Lv-Abcc10 |
| e13 | SPU_006478 | Lv-Ergic3 |
| e13 | SPU_006584 | Lv-Hypp_1667 |
| e13 | SPU_006636 | Lv-Zmynd8_2 |
| e13 | SPU_006692 | none |
| e13 | SPU_006790 | Lv-Pgm3 |

| | | |
|---|---|---|
| e13 | SPU_006840 | Lv-Baz1a |
| e13 | SPU_006897 | Lv-Brd8 |
| e13 | SPU_006987 | NA |
| e13 | SPU_006990 | Lv-Rbm19_1 |
| e13 | SPU_007111 | Lv-Banf1 |
| e13 | SPU_007144 | Lv-Hypp_1701 |
| e13 | SPU_007287 | none |
| e13 | SPU_007315 | Lv-Nomo_1 |
| e13 | SPU_007374 | Lv-Psmc3 |
| e13 | SPU_007382 | NA |
| e13 | SPU_007385 | Lv-Hypp_512 |
| e13 | SPU_007479 | Lv-Arp6 |
| e13 | SPU_007536 | Lv-Atpbd4 |
| e13 | SPU_007540 | none |
| e13 | SPU_007675 | Lv-Sf3a3 |
| e13 | SPU_007694 | Lv-Hypp_1745 |
| e13 | SPU_007728 | Lv-Endrvt15 |
| e13 | SPU_007736 | Lv-Ilf2 |
| e13 | SPU_007902 | NA |
| e13 | SPU_008013 | Lv-Nsun4 |
| e13 | SPU_008017 | Lv-Rrn3 |
| e13 | SPU_008080 | Lv-B4galt6 |
| e13 | SPU_008113 | none |
| e13 | SPU_008195 | Lv-Hpd_1 |
| e13 | SPU_008274 | none |
| e13 | SPU_008298 | Lv-Gemin8 |
| e13 | SPU_008365 | Lv-Ogg1 |
| e13 | SPU_008388 | Lv-Ints1 |
| e13 | SPU_008505 | Lv-Cys_rich_fgfr |
| e13 | SPU_008638 | Lv-Narg1 |
| e13 | SPU_008881 | Lv-C10orf76L |
| e13 | SPU_008899 | Lv-Bard1L |
| e13 | SPU_008902 | Lv-Hypp_1811 |
| e13 | SPU_008928 | Lv-Mettl1 |
| e13 | SPU_008938 | Lv-Usp37L |
| e13 | SPU_009023 | Lv-Cnot8 |
| e13 | SPU_009086 | Lv-Qki |
| e13 | SPU_009205 | Lv-Ints7_1 |

| | | |
|---|---|---|
| e13 | SPU_009216 | none |
| e13 | SPU_009274 | Lv-C5orf22L |
| e13 | SPU_009277 | Lv-Med_rnapIIh |
| e13 | SPU_009358 | Lv-Dtymk_1 |
| e13 | SPU_009428 | Lv-Tmem45a |
| e13 | SPU_009450 | Lv-Tlr141 |
| e13 | SPU_009513 | Lv-Sec24c |
| e13 | SPU_009572 | none |
| e13 | SPU_009653 | Lv-CspeN3c |
| e13 | SPU_009735 | Lv-Farsa |
| e13 | SPU_009797 | Lv-Ku70l |
| e13 | SPU_009922 | Lv-Cys_rich_fgfr_1 |
| e13 | SPU_009980 | Lv-TdTl/Pol/Mu |
| e13 | SPU_010009 | Lv-Zmat2 |
| e13 | SPU_010162 | Lv-Trim56L |
| e13 | SPU_010171 | Lv-Hypp_610 |
| e13 | SPU_010278 | none |
| e13 | SPU_010392 | Lv-Fut8_2 |
| e13 | SPU_010533 | Lv-Orc2 |
| e13 | SPU_010572 | NA |
| e13 | SPU_010573 | none |
| e13 | SPU_010628 | Lv-Gorasp2l |
| e13 | SPU_010674 | Lv-Tmem43 |
| e13 | SPU_010690 | Lv-Gal3st1 |
| e13 | SPU_010691 | Lv-Ddx3Y |
| e13 | SPU_010788 | Lv-Fn3_27 |
| e13 | SPU_010824 | Lv-Pwp2-2 |
| e13 | SPU_010926 | Lv-Polr1a |
| e13 | SPU_010975 | Lv-Rrbp1L |
| e13 | SPU_011216 | none |
| e13 | SPU_011396 | Lv-Znf294 |
| e13 | SPU_011429 | Lv-Znf593 |
| e13 | SPU_011567 | Lv-Ssr3 |
| e13 | SPU_011569 | none |
| e13 | SPU_011665 | Lv-Mrto |
| e13 | SPU_011712 | Lv-Prmt3 |
| e13 | SPU_011777 | Lv-Hypp_660 |
| e13 | SPU_011824 | Lv-PolypL_70 |

| | | |
|---|---|---|
| e13 | SPU_011904 | Lv-Nudt15 |
| e13 | SPU_011916 | Lv-Ncoa5L |
| e13 | SPU_011934 | Lv-Qprt |
| e13 | SPU_012051 | Lv-Cpsf_100kDa |
| e13 | SPU_012099 | Lv-Ict1L |
| e13 | SPU_012124 | Lv-Nubp1 |
| e13 | SPU_012164 | Lv-Hypp_671 |
| e13 | SPU_012276 | Lv-Fanca |
| e13 | SPU_012563 | Lv-Rhogap11A |
| e13 | SPU_012607 | Lv-SnrpD1 |
| e13 | SPU_012696 | Lv-Cdc23 |
| e13 | SPU_012778 | Lv-Tatdn2L |
| e13 | SPU_012804 | Lv-Uckl1 |
| e13 | SPU_012820 | Lv-Slc25a24 |
| e13 | SPU_012961 | Lv-Phlp |
| e13 | SPU_012998 | Lv-Fn3_1 |
| e13 | SPU_013007 | Lv-SnrpB |
| e13 | SPU_013088 | NA |
| e13 | SPU_013113 | none |
| e13 | SPU_013144 | Lv-Mprot1L_2 |
| e13 | SPU_013375 | Lv-Ublcp1_1 |
| e13 | SPU_013379 | Lv-Kiaa1456L |
| e13 | SPU_013398 | Lv-Eif3j |
| e13 | SPU_013468 | none |
| e13 | SPU_013557 | Lv-Fcolf |
| e13 | SPU_013566 | Lv-Mecr |
| e13 | SPU_013681 | Lv-Kiaa0090L |
| e13 | SPU_013725 | Lv-Rnmt7 |
| e13 | SPU_013804 | Lv-Rere |
| e13 | SPU_013889 | Lv-Pxdn |
| e13 | SPU_013972 | Lv-Tmf1L |
| e13 | SPU_014105 | Lv-Exosc1 |
| e13 | SPU_014147 | Lv-Hypp_2131 |
| e13 | SPU_014154 | Lv-NicalinL |
| e13 | SPU_014323 | Lv-Pigv |
| e13 | SPU_014525 | Lv-Par6ph |
| e13 | SPU_014612 | Lv-Hypp_759 |
| e13 | SPU_014644 | Lv-Hypp_2163 |

| | | |
|---|---|---|
| e13 | SPU_014716 | Lv-PolypL_6 |
| e13 | SPU_014735 | none |
| e13 | SPU_014762 | Lv-Exosc3 |
| e13 | SPU_014775 | Lv-Dimt1 |
| e13 | SPU_014794 | Lv-Nob1 |
| e13 | SPU_015064 | Lv-Ddx23 |
| e13 | SPU_015198 | Lv-Lnb/7Tm/GpcrB |
| e13 | SPU_015300 | Lv-Gpr115_5 |
| e13 | SPU_015305 | Lv-Vparpf_2 |
| e13 | SPU_015318 | Lv-Ttc35 |
| e13 | SPU_015529 | Lv-Dna_pkcs |
| e13 | SPU_015577 | none |
| e13 | SPU_015607 | Lv-Ddx18 |
| e13 | SPU_015634 | Lv-Hypp_794 |
| e13 | SPU_015653 | Lv-Pcnt |
| e13 | SPU_015706 | Lv-Nadrin |
| e13 | SPU_015754 | none |
| e13 | SPU_015851 | Lv-Cpt1a |
| e13 | SPU_015969 | Lv-Exoc4 |
| e13 | SPU_016010 | Lv-Slc6a5-2 |
| e13 | SPU_016024 | Lv-Fdps_1 |
| e13 | SPU_016082 | Lv-Slc7a5 |
| e13 | SPU_016215 | none |
| e13 | SPU_016341 | Lv-Ube2j1 |
| e13 | SPU_016381 | Lv-C5orf44 |
| e13 | SPU_016467 | Lv-Ddx24_1 |
| e13 | SPU_016546 | Lv-MacpfB3 |
| e13 | SPU_016553 | Lv-Utp14A |
| e13 | SPU_016626 | Lv-Sec24c_1 |
| e13 | SPU_016632 | none |
| e13 | SPU_016671 | Lv-Ddx31 |
| e13 | SPU_016844 | NA |
| e13 | SPU_016888 | Lv-Hypp_849 |
| e13 | SPU_016892 | none |
| e13 | SPU_016936 | Lv-Fn3_3 |
| e13 | SPU_016937 | Lv-RptphL1 |
| e13 | SPU_017217 | none |
| e13 | SPU_017222 | none |

| | | |
|---|---|---|
| e13 | SPU_017265 | Lv-Unc44_124 |
| e13 | SPU_017413 | Lv-Larp1 |
| e13 | SPU_017547 | Lv-Tm9sf4 |
| e13 | SPU_017583 | Lv-Ddx59_1 |
| e13 | SPU_017602 | Lv-Prpf40B |
| e13 | SPU_017831 | Lv-Ints1_2 |
| e13 | SPU_018009 | Lv-AnpepL |
| e13 | SPU_018039 | Lv-MettL5_2 |
| e13 | SPU_018067 | NA |
| e13 | SPU_018172 | Lv-Utp14A_1 |
| e13 | SPU_018203 | Lv-Ddx55 |
| e13 | SPU_018350 | Lv-Sec24c_2 |
| e13 | SPU_018356 | Lv-Cub/HyalinL3 |
| e13 | SPU_018387 | Lv-Endrvt37 |
| e13 | SPU_018464 | Lv-Lrpprc-2 |
| e13 | SPU_018546 | Lv-Aspscr1 |
| e13 | SPU_018589 | none |
| e13 | SPU_018592 | none |
| e13 | SPU_018599 | Lv-Hypp_2394 |
| e13 | SPU_018634 | none |
| e13 | SPU_018645 | Lv-Rbm13 |
| e13 | SPU_018688 | Lv-Farsb |
| e13 | SPU_018765 | Lv-Spr_1 |
| e13 | SPU_019010 | Lv-Hypp_3010 |
| e13 | SPU_019147 | Lv-SnrpE |
| e13 | SPU_019210 | Lv-Ngdn |
| e13 | SPU_019326 | none |
| e13 | SPU_019340 | Lv-Xpo6_1 |
| e13 | SPU_019431 | Lv-Dcps |
| e13 | SPU_019435 | Lv-Z114 |
| e13 | SPU_019475 | Lv-Rrp15h |
| e13 | SPU_019569 | Lv-Brd8_1 |
| e13 | SPU_019580 | Lv-SnrpF |
| e13 | SPU_019601 | Lv-Wap/Ty/Egf |
| e13 | SPU_019607 | none |
| e13 | SPU_019664 | Lv-Tsr1_1 |
| e13 | SPU_019673 | Lv-Rbm8A |
| e13 | SPU_019695 | Lv-Igf2r |

| | | |
|---|---|---|
| e13 | SPU_019888 | Lv-Mad2b |
| e13 | SPU_019933 | none |
| e13 | SPU_019992 | Lv-Unc44_326 |
| e13 | SPU_020108 | none |
| e13 | SPU_020373 | none |
| e13 | SPU_020422 | Lv-Neurl4-2 |
| e13 | SPU_020440 | Lv-Ankrd13c |
| e13 | SPU_020483 | Lv-Hypp_976 |
| e13 | SPU_020490 | none |
| e13 | SPU_020631 | Lv-C20orf24L |
| e13 | SPU_020674 | Lv-Tlcd1 |
| e13 | SPU_020720 | Lv-Gale |
| e13 | SPU_020739 | Lv-Gluld1_10 |
| e13 | SPU_020755 | none |
| e13 | SPU_020771 | NA |
| e13 | SPU_020815 | Lv-Ncoa5L_1 |
| e13 | SPU_020820 | Lv-Unc44_148 |
| e13 | SPU_020851 | Lv-Nudt3 |
| e13 | SPU_020922 | none |
| e13 | SPU_020924 | Lv-Gtf2h4 |
| e13 | SPU_020994 | Lv-Eid3 |
| e13 | SPU_021000 | Lv-Rdh14_1 |
| e13 | SPU_021010 | none |
| e13 | SPU_021022 | Lv-Osgepl1L_1 |
| e13 | SPU_021144 | Lv-Tmed4 |
| e13 | SPU_021175 | Lv-HspC152l |
| e13 | SPU_021277 | Lv-Sca2L |
| e13 | SPU_021302 | Lv-Iars |
| e13 | SPU_021312 | none |
| e13 | SPU_021323 | Lv-Hypp_2545 |
| e13 | SPU_021481 | Lv-Bbs10L |
| e13 | SPU_021615 | none |
| e13 | SPU_021684 | Lv-Tdrd3L_1 |
| e13 | SPU_021715 | Lv-Endrvt103 |
| e13 | SPU_021759 | Lv-Tat |
| e13 | SPU_021902 | NA |
| e13 | SPU_022080 | Lv-Bzw1 |
| e13 | SPU_022156 | Lv-Supv3L1 |

| | | |
|---|---|---|
| e13 | SPU_022206 | Lv-Rpusd4 |
| e13 | SPU_022238 | NA |
| e13 | SPU_022260 | Lv-Prps1 |
| e13 | SPU_022264 | Lv-Hnrpul1 |
| e13 | SPU_022265 | Lv-Hnrpul1_1 |
| e13 | SPU_022281 | none |
| e13 | SPU_022429 | Lv-Cdk5rap3 |
| e13 | SPU_022430 | Lv-Hypp_183 |
| e13 | SPU_022606 | Lv-Hypp_1052 |
| e13 | SPU_022728 | Lv-TpaseL_14 |
| e13 | SPU_022735 | Lv-Ppie |
| e13 | SPU_022779 | Lv-Pdcd6 |
| e13 | SPU_022850 | Lv-Mettl14 |
| e13 | SPU_022908 | Lv-Gnptgl |
| e13 | SPU_022923 | Lv-Hnrph2L |
| e13 | SPU_022926 | Lv-Abca1_2 |
| e13 | SPU_022993 | Lv-Hypp_1064 |
| e13 | SPU_023019 | Lv-Atad3 |
| e13 | SPU_023112 | NA |
| e13 | SPU_023200 | Lv-Staufen1 |
| e13 | SPU_023222 | none |
| e13 | SPU_023235 | Lv-Hypp_1079 |
| e13 | SPU_023282 | Lv-Hypp_201 |
| e13 | SPU_023318 | Lv-Imp4 |
| e13 | SPU_023401 | Lv-Pdk1 |
| e13 | SPU_023403 | Lv-St6galnac4_1 |
| e13 | SPU_023427 | none |
| e13 | SPU_023457 | Lv-Abcc9l |
| e13 | SPU_023653 | Lv-Lactb2 |
| e13 | SPU_023712 | none |
| e13 | SPU_023727 | Lv-Klf13 |
| e13 | SPU_023855 | Lv-Glt8d1 |
| e13 | SPU_023882 | Lv-Ptenh |
| e13 | SPU_023895 | Lv-Srp14 |
| e13 | SPU_023948 | Lv-Erf1 |
| e13 | SPU_023962 | NA |
| e13 | SPU_023985 | Lv-Naalad2L_14 |
| e13 | SPU_024030 | Lv-SnrpD3 |

| | | |
|---|---|---|
| e13 | SPU_024063 | Lv-Tmed3 |
| e13 | SPU_024155 | Lv-Hypp_2712 |
| e13 | SPU_024178 | none |
| e13 | SPU_024255 | none |
| e13 | SPU_024284 | Lv-PppL_139 |
| e13 | SPU_024288 | none |
| e13 | SPU_024306 | Lv-Dazap_1 |
| e13 | SPU_024324 | none |
| e13 | SPU_024348 | Lv-Srp68 |
| e13 | SPU_024349 | Lv-Pp1r8 |
| e13 | SPU_024412 | none |
| e13 | SPU_024549 | none |
| e13 | SPU_024568 | none |
| e13 | SPU_024658 | none |
| e13 | SPU_024704 | Lv-Bat1 |
| e13 | SPU_024820 | Lv-Ptprlec6(novel) |
| e13 | SPU_024829 | none |
| e13 | SPU_024948 | Lv-Elmo |
| e13 | SPU_024953 | Lv-Z34 |
| e13 | SPU_024979 | Lv-Endrvt112 |
| e13 | SPU_025105 | NA |
| e13 | SPU_025163 | none |
| e13 | SPU_025318 | Lv-Mrpl19 |
| e13 | SPU_025320 | Lv-Utp20-2 |
| e13 | SPU_025324 | none |
| e13 | SPU_025367 | Lv-Cpt1a_1 |
| e13 | SPU_025376 | none |
| e13 | SPU_025423 | Lv-Kiaa0196 |
| e13 | SPU_025456 | Lv-Hypp_230 |
| e13 | SPU_025469 | Lv-Slc25a19_1 |
| e13 | SPU_025527 | Lv-Slc33a1 |
| e13 | SPU_025633 | Lv-Dnaja3 |
| e13 | SPU_025714 | none |
| e13 | SPU_025785 | Lv-Nom1 |
| e13 | SPU_025889 | Lv-Hypp_233 |
| e13 | SPU_026082 | Lv-Eed |
| e13 | SPU_026157 | Lv-Ahctf1 |
| e13 | SPU_026240 | Lv-Nagpa |

| | | |
|---|---|---|
| e13 | SPU_026247 | Lv-Lrpprc |
| e13 | SPU_026499 | Lv-Ndufaf1 |
| e13 | SPU_026937 | none |
| e13 | SPU_026954 | Lv-Lyrm4 |
| e13 | SPU_026980 | Lv-Kbtbd8 |
| e13 | SPU_027004 | Lv-Sfrs2_1 |
| e13 | SPU_027022 | Lv-Yipf4l |
| e13 | SPU_027050 | none |
| e13 | SPU_027067 | Lv-Pollp_1 |
| e13 | SPU_027089 | Lv-Lmf2 |
| e13 | SPU_027158 | Lv-CapsL |
| e13 | SPU_027183 | none |
| e13 | SPU_027216 | Lv-Pycs |
| e13 | SPU_027335 | Lv-Gage2 |
| e13 | SPU_027548 | Lv-Yipf6 |
| e13 | SPU_027712 | Lv-Pcbp1 |
| e13 | SPU_027740 | none |
| e13 | SPU_027802 | Lv-Apoe |
| e13 | SPU_027810 | Lv-Tmem18 |
| e13 | SPU_027833 | Lv-Aur |
| e13 | SPU_027843 | Lv-Hel308 |
| e13 | SPU_027976 | Lv-Ecd |
| e13 | SPU_028020 | Lv-Nef3 |
| e13 | SPU_028036 | Lv-Rasgrf1 |
| e13 | SPU_028130 | Lv-Spsc3h |
| e13 | SPU_028390 | Lv-Tmprss2L_2 |
| e13 | SPU_028524 | Lv-Unk_11 |
| e13 | SPU_028571 | Lv-Hypp_287 |
| e13 | SPU_028604 | Lv-Polr3b |
| e13 | SPU_028636 | none |
| e13 | SPU_028664 | Lv-Bat5 |
| e13 | SPU_028672 | Lv-Tysnd1_1 |
| e13 | SPU_028674 | Lv-Hypp_2963 |
| e13 | SPU_028679 | Lv-Z481 |
| e13 | SPU_028771 | Lv-Prpf18 |
| e13 | SPU_028772 | Lv-Rad18 |
| e13 | SPU_028779 | Lv-Gcn1l1_2 |
| e14 | SPU_000166 | Lv-Slc35f3L |

| | | |
|---|---|---|
| e14 | SPU_000317 | Lv-Chst9 |
| e14 | SPU_000446 | Lv-Adrm1L |
| e14 | SPU_000486 | Lv-Tydp1 |
| e14 | SPU_000526 | Lv-Ebr1 |
| e14 | SPU_000556 | none |
| e14 | SPU_000595 | Lv-Ef1a |
| e14 | SPU_000724 | none |
| e14 | SPU_000759 | Lv-Psmd13 |
| e14 | SPU_000879 | none |
| e14 | SPU_001010 | none |
| e14 | SPU_001038 | Lv-C2gnt3 |
| e14 | SPU_001078 | Lv-Znf830L |
| e14 | SPU_001119 | Lv-Ssrp1 |
| e14 | SPU_001124 | none |
| e14 | SPU_001125 | Lv-Rbms3 |
| e14 | SPU_001203 | none |
| e14 | SPU_001206 | Lv-Unc44_196 |
| e14 | SPU_001216 | none |
| e14 | SPU_001476 | Lv-Hypp_1332 |
| e14 | SPU_001599 | none |
| e14 | SPU_001652 | Lv-PolppL-4 |
| e14 | SPU_001776 | NA |
| e14 | SPU_001823 | none |
| e14 | SPU_001879 | Lv-Abcc5C |
| e14 | SPU_001933 | Lv-Dad1 |
| e14 | SPU_002050 | Lv-Ef1a(Cter) |
| e14 | SPU_002124 | Lv-Suclg2 |
| e14 | SPU_002153 | Lv-Gpsn2 |
| e14 | SPU_002234 | Lv-Psmc |
| e14 | SPU_002260 | Lv-Nckap1 |
| e14 | SPU_002278 | Lv-Prg4_2 |
| e14 | SPU_002299 | Lv-Psma5 |
| e14 | SPU_002348 | NA |
| e14 | SPU_002432 | Lv-Unk_30 |
| e14 | SPU_002521 | Lv-Rgs7 |
| e14 | SPU_002616 | Lv-Clta |
| e14 | SPU_002674 | Lv-TfIIIa |
| e14 | SPU_002713 | Lv-Fut10 |

| | | |
|---|---|---|
| e14 | SPU_002724 | Lv-Mrps9 |
| e14 | SPU_002747 | Lv-Unk_94 |
| e14 | SPU_002760 | Lv-Kiaa1033L |
| e14 | SPU_002788 | Lv-Btub1 |
| e14 | SPU_002813 | none |
| e14 | SPU_002815 | Lv-Dlx |
| e14 | SPU_002908 | Lv-VarsiB |
| e14 | SPU_002993 | Lv-Ddx1 |
| e14 | SPU_003062 | Lv-Zc3p26_7 |
| e14 | SPU_003114 | Lv-Nova1 |
| e14 | SPU_003120 | Lv-Ppp2r2aL |
| e14 | SPU_003206 | Lv-Lrr/Gpcr_1 |
| e14 | SPU_003240 | Lv-Slc34a2 |
| e14 | SPU_003311 | Lv-Osbpl6 |
| e14 | SPU_003313 | Lv-NpC2L |
| e14 | SPU_003318 | Lv-PolppL_26 |
| e14 | SPU_003324 | Lv-Psd4 |
| e14 | SPU_003372 | Lv-PolppL_32 |
| e14 | SPU_003402 | Lv-Sdc |
| e14 | SPU_003514 | Lv-Rhol3 |
| e14 | SPU_003625 | Lv-Osbpl6_1 |
| e14 | SPU_003825 | Lv-14-3-3e |
| e14 | SPU_003835 | Lv-Tgfb2 |
| e14 | SPU_003867 | Lv-Gcsh |
| e14 | SPU_003913 | Lv-Oat |
| e14 | SPU_004007 | Lv-Galnt7 |
| e14 | SPU_004021 | Lv-Limc1 |
| e14 | SPU_004057 | Lv-Elovl6_1 |
| e14 | SPU_004065 | none |
| e14 | SPU_004198 | NA |
| e14 | SPU_004223 | Lv-Hypp_1508 |
| e14 | SPU_004245 | Lv-Netrin1 |
| e14 | SPU_004417 | Lv-Prpf6 |
| e14 | SPU_004526 | Lv-Nr1h6c |
| e14 | SPU_004582 | none |
| e14 | SPU_004587 | Lv-Top2b |
| e14 | SPU_004628 | Lv-Map1aph |
| e14 | SPU_004630 | Lv-Uba6 |

| | | |
|---|---|---|
| e14 | SPU_004654 | Lv-Acsf3 |
| e14 | SPU_004749 | Lv-Wdtc1-2 |
| e14 | SPU_004750 | Lv-Acsbg1 |
| e14 | SPU_004767 | Lv-Otop2L |
| e14 | SPU_004866 | Lv-Rin2L |
| e14 | SPU_005051 | Lv-PolppL_43 |
| e14 | SPU_005100 | Lv-Rnf103 |
| e14 | SPU_005115 | Lv-Clect/Fn3_5 |
| e14 | SPU_005271 | Lv-Arl2 |
| e14 | SPU_005304 | Lv-C5orf33 |
| e14 | SPU_005355 | Lv-Grf1 |
| e14 | SPU_005373 | Lv-Hs3st3b1 |
| e14 | SPU_005397 | Lv-Rttn |
| e14 | SPU_005447 | Lv-Rap1gap |
| e14 | SPU_005508 | Lv-Fn3/Ig_2 |
| e14 | SPU_005534 | Lv-Slc17a5_2 |
| e14 | SPU_005602 | Lv-Tensin |
| e14 | SPU_005680 | Lv-Hypp_460 |
| e14 | SPU_005703 | none |
| e14 | SPU_005715 | none |
| e14 | SPU_005776 | Lv-Ift122h |
| e14 | SPU_005835 | none |
| e14 | SPU_005851 | Lv-Haao |
| e14 | SPU_005880 | Lv-TnL1 |
| e14 | SPU_006015 | Lv-Prim1 |
| e14 | SPU_006104 | Lv-Hypp_1630 |
| e14 | SPU_006130 | Lv-Rt26 |
| e14 | SPU_006143 | Lv-Slc25a20 |
| e14 | SPU_006152 | none |
| e14 | SPU_006372 | Lv-Polr3h |
| e14 | SPU_006590 | Lv-Hypp_486 |
| e14 | SPU_006752 | Lv-Deadc1 |
| e14 | SPU_006848 | Lv-Mcm3 |
| e14 | SPU_006857 | none |
| e14 | SPU_006967 | Lv-Tmem164 |
| e14 | SPU_007013 | Lv-CycD |
| e14 | SPU_007077 | NA |
| e14 | SPU_007257 | none |

| | | |
|---|---|---|
| e14 | SPU_007406 | Lv-Cyp1L1 |
| e14 | SPU_007447 | Lv-Hypp_1722 |
| e14 | SPU_007464 | Lv-Tomm70a |
| e14 | SPU_007563 | none |
| e14 | SPU_007577 | Lv-Hypp_1732 |
| e14 | SPU_007674 | none |
| e14 | SPU_007767 | Lv-Gabrb3L |
| e14 | SPU_007803 | Lv-Hs3st5L |
| e14 | SPU_007825 | Lv-Ndufs2 |
| e14 | SPU_007829 | Lv-Endrvt64 |
| e14 | SPU_007851 | Lv-Ccdc21L |
| e14 | SPU_007909 | Lv-Ndufs2_1 |
| e14 | SPU_007945 | Lv-Hypp_1755 |
| e14 | SPU_007947 | Lv-Kirrel2L_1 |
| e14 | SPU_008045 | none |
| e14 | SPU_008046 | Lv-Arl9/10 |
| e14 | SPU_008063 | Lv-Sbno1L |
| e14 | SPU_008073 | Lv-Uqcrb |
| e14 | SPU_008083 | Lv-Dpagt1 |
| e14 | SPU_008111 | Lv-Polr1d |
| e14 | SPU_008175 | Lv-Mitf |
| e14 | SPU_008193 | Lv-RtL_18 |
| e14 | SPU_008282 | Lv-Ibtk |
| e14 | SPU_008294 | Lv-Taf5 |
| e14 | SPU_008318 | Lv-Sel1 |
| e14 | SPU_008488 | Lv-Rt30 |
| e14 | SPU_008549 | none |
| e14 | SPU_008553 | Lv-Hadhb |
| e14 | SPU_008719 | Lv-Polk |
| e14 | SPU_008870 | none |
| e14 | SPU_008905 | Lv-Rad23 |
| e14 | SPU_008942 | NA |
| e14 | SPU_009044 | Lv-Pel |
| e14 | SPU_009103 | Lv-Kiaa0564L |
| e14 | SPU_009164 | Lv-Hsp701E |
| e14 | SPU_009165 | Lv-Hsp701A |
| e14 | SPU_009272 | Lv-Chst11L_1 |
| e14 | SPU_009285 | none |

| | | |
|---|---|---|
| e14 | SPU_009330 | Lv-PppL_253 |
| e14 | SPU_009351 | none |
| e14 | SPU_009417 | Lv-Ugt2b33 |
| e14 | SPU_009430 | Lv-Slc22a3_8 |
| e14 | SPU_009461 | Lv-Rpn2 |
| e14 | SPU_009713 | Lv-Stoml2_1 |
| e14 | SPU_009714 | Lv-Fastkd5 |
| e14 | SPU_009734 | Lv-Yipf1l |
| e14 | SPU_009800 | none |
| e14 | SPU_009923 | none |
| e14 | SPU_009946 | Lv-Mrps14 |
| e14 | SPU_010024 | Lv-AspLm |
| e14 | SPU_010074 | Lv-Hypp_61 |
| e14 | SPU_010076 | NA |
| e14 | SPU_010214 | NA |
| e14 | SPU_010283 | Lv-Ser_asprp |
| e14 | SPU_010442 | Lv-Hypp_615 |
| e14 | SPU_010541 | none |
| e14 | SPU_010545 | Lv-Racl1 |
| e14 | SPU_010722 | Lv-Furin1 |
| e14 | SPU_010747 | Lv-Copa |
| e14 | SPU_010800 | Lv-Actr10h |
| e14 | SPU_010830 | none |
| e14 | SPU_010932 | Lv-Hypp_1946 |
| e14 | SPU_010944 | Lv-Ntan1 |
| e14 | SPU_010976 | Lv-Pxmp2 |
| e14 | SPU_011105 | none |
| e14 | SPU_011108 | Lv-Hypp_1959 |
| e14 | SPU_011136 | Lv-Asb8 |
| e14 | SPU_011243 | none |
| e14 | SPU_011307 | Lv-Abcd3 |
| e14 | SPU_011367 | Lv-Uba/Ubx |
| e14 | SPU_011511 | Lv-Exoc3L |
| e14 | SPU_011535 | Lv-Dopey1l |
| e14 | SPU_011546 | Lv-Lsm5 |
| e14 | SPU_011560 | Lv-Pla2g1b_6 |
| e14 | SPU_011612 | Lv-PolppL_92 |
| e14 | SPU_011662 | Lv-Hypp_1996 |

| | | |
|---|---|---|
| e14 | SPU_011688 | Lv-Fram1l |
| e14 | SPU_011779 | Lv-Zrsr2 |
| e14 | SPU_011862 | NA |
| e14 | SPU_011874 | Lv-C3orf58L |
| e14 | SPU_011878 | none |
| e14 | SPU_011884 | Lv-Idh3b |
| e14 | SPU_011932 | Lv-Slc16a14L_3 |
| e14 | SPU_012049 | Lv-Fdx1 |
| e14 | SPU_012102 | Lv-PolypL_72 |
| e14 | SPU_012204 | none |
| e14 | SPU_012250 | Lv-Asna1 |
| e14 | SPU_012263 | Lv-Hypp_2024 |
| e14 | SPU_012297 | Lv-Hypp_2991 |
| e14 | SPU_012304 | Lv-C1orf84L |
| e14 | SPU_012373 | Lv-Pdhx |
| e14 | SPU_012396 | Lv-Btf3l4 |
| e14 | SPU_012422 | Lv-PolypL_92 |
| e14 | SPU_012504 | Lv-Asphm2 |
| e14 | SPU_012519 | Lv-PolppL_94 |
| e14 | SPU_012601 | Lv-Hypp_2040 |
| e14 | SPU_012618 | Lv-Emaps |
| e14 | SPU_012678 | Lv-Atub14 |
| e14 | SPU_012733 | none |
| e14 | SPU_012739 | none |
| e14 | SPU_012770 | Lv-Chl1l |
| e14 | SPU_012784 | Lv-Atp6v1f |
| e14 | SPU_012822 | NA |
| e14 | SPU_012840 | Lv-Traf1_1 |
| e14 | SPU_012874 | Lv-Abcg9 |
| e14 | SPU_012939 | none |
| e14 | SPU_012957 | Lv-Hypp_2060 |
| e14 | SPU_013010 | Lv-Glud1_1 |
| e14 | SPU_013097 | Lv-Psma6 |
| e14 | SPU_013194 | Lv-Siah |
| e14 | SPU_013414 | Lv-Gi |
| e14 | SPU_013511 | Lv-Fam63b |
| e14 | SPU_013515 | Lv-Hypp_2086 |
| e14 | SPU_013617 | Lv-Smc4 |

| | | |
|---|---|---|
| e14 | SPU_013699 | Lv-Lap3L |
| e14 | SPU_013711 | none |
| e14 | SPU_013727 | Lv-Lsm14A |
| e14 | SPU_013728 | NA |
| e14 | SPU_013808 | Lv-PolppL_100 |
| e14 | SPU_013857 | Lv-Vps28 |
| e14 | SPU_013935 | Lv-Dynact1ph |
| e14 | SPU_014138 | Lv-Slc10a2L_1 |
| e14 | SPU_014140 | Lv-Ttrspn_12 |
| e14 | SPU_014156 | NA |
| e14 | SPU_014233 | Lv-Mpnd |
| e14 | SPU_014343 | Lv-Aco1 |
| e14 | SPU_014432 | Lv-HmH2a.1 |
| e14 | SPU_014460 | Lv-Acp2 |
| e14 | SPU_014594 | Lv-14-3-3_1 |
| e14 | SPU_014630 | none |
| e14 | SPU_014660 | Lv-Slc25a4 |
| e14 | SPU_014706 | none |
| e14 | SPU_014742 | Lv-Lrrfip2 |
| e14 | SPU_014784 | Lv-Fgd6 |
| e14 | SPU_014831 | Lv-Gtpbp5 |
| e14 | SPU_014885 | Lv-Rbm25 |
| e14 | SPU_014942 | NA |
| e14 | SPU_014999 | Lv-Gagpol7 |
| e14 | SPU_015012 | Lv-Nsfl1C |
| e14 | SPU_015013 | Lv-Psmd2 |
| e14 | SPU_015016 | none |
| e14 | SPU_015039 | none |
| e14 | SPU_015044 | Lv-Ssr4 |
| e14 | SPU_015148 | Lv-Hypp_2192 |
| e14 | SPU_015184 | Lv-Kiaa1797L |
| e14 | SPU_015237 | none |
| e14 | SPU_015360 | Lv-Ado |
| e14 | SPU_015406 | Lv-Kiaa0467L |
| e14 | SPU_015418 | Lv-Copb1 |
| e14 | SPU_015441 | Lv-Ranbp3 |
| e14 | SPU_015491 | none |
| e14 | SPU_015506 | Lv-Gosr1 |

| | | |
|---|---|---|
| e14 | SPU_015783 | Lv-Paip1L |
| e14 | SPU_015843 | Lv-Ube4A |
| e14 | SPU_015902 | Lv-Hypp_2238 |
| e14 | SPU_015950 | Lv-Hypp_122 |
| e14 | SPU_016030 | Lv-RtL_2 |
| e14 | SPU_016205 | Lv-Ndufs7 |
| e14 | SPU_016226 | Lv-Mif6 |
| e14 | SPU_016245 | Lv-Mipep_1 |
| e14 | SPU_016294 | Lv-Tmem189 |
| e14 | SPU_016403 | Lv-DmxL2 |
| e14 | SPU_016422 | Lv-Hypp_2261 |
| e14 | SPU_016474 | Lv-L1td1L_1 |
| e14 | SPU_016540 | none |
| e14 | SPU_016642 | Lv-Hypp_132 |
| e14 | SPU_016644 | Lv-Rhogef11l |
| e14 | SPU_016731 | Lv-Gdpd5-2 |
| e14 | SPU_016776 | NA |
| e14 | SPU_016873 | Lv-Zf(cchc)26_2 |
| e14 | SPU_016976 | Lv-Rhogef2 |
| e14 | SPU_017009 | Lv-Slc46a1L_4 |
| e14 | SPU_017153 | none |
| e14 | SPU_017293 | Lv-Pric285 |
| e14 | SPU_017302 | Lv-Rbm10_1 |
| e14 | SPU_017417 | Lv-Gcat_1 |
| e14 | SPU_017453 | Lv-Srcr139 |
| e14 | SPU_017695 | Lv-Ganab |
| e14 | SPU_017727 | Lv-IrbitL |
| e14 | SPU_017798 | Lv-Sdha_1 |
| e14 | SPU_017845 | Lv-PppL_117 |
| e14 | SPU_017944 | Lv-Vps53 |
| e14 | SPU_017993 | Lv-Nlr18 |
| e14 | SPU_018019 | Lv-Ighmbp2 |
| e14 | SPU_018059 | Lv-PlexB1_1 |
| e14 | SPU_018197 | Lv-Muc/hyalinL |
| e14 | SPU_018244 | Lv-Clta_1 |
| e14 | SPU_018246 | Lv-Smad6/7_1 |
| e14 | SPU_018322 | Lv-Agpat1 |
| e14 | SPU_018386 | NA |

| | | |
|---|---|---|
| e14 | SPU_018428 | Lv-Zzz3L |
| e14 | SPU_018430 | Lv-Srcr143 |
| e14 | SPU_018500 | Lv-Dynamnp50ph |
| e14 | SPU_018674 | Lv-Mrpl20l |
| e14 | SPU_018677 | Lv-Oach |
| e14 | SPU_018730 | Lv-Top2a |
| e14 | SPU_018879 | Lv-C12orf66 |
| e14 | SPU_018904 | Lv-ScavrB1L |
| e14 | SPU_018907 | Lv-Rpe |
| e14 | SPU_018932 | Lv-AhnakL |
| e14 | SPU_018944 | Lv-Med31 |
| e14 | SPU_018981 | Lv-Arl11l2 |
| e14 | SPU_019012 | Lv-Cope |
| e14 | SPU_019056 | Lv-Hypp_927 |
| e14 | SPU_019127 | Lv-Syt6-2 |
| e14 | SPU_019138 | Lv-Lhh2be |
| e14 | SPU_019409 | Lv-Abca2 |
| e14 | SPU_019430 | none |
| e14 | SPU_019494 | Lv-Cdc42 |
| e14 | SPU_019543 | Lv-Ube2w |
| e14 | SPU_019644 | Lv-Sgk493 |
| e14 | SPU_019679 | Lv-RtL_47 |
| e14 | SPU_019691 | Lv-Flot1 |
| e14 | SPU_019809 | Lv-Mtrr |
| e14 | SPU_020116 | NA |
| e14 | SPU_020199 | Lv-Syt7 |
| e14 | SPU_020202 | Lv-Hypp_2495 |
| e14 | SPU_020274 | Lv-Msi1 |
| e14 | SPU_020321 | Lv-Kctd1_2 |
| e14 | SPU_020539 | Lv-Hypp_977 |
| e14 | SPU_020544 | NA |
| e14 | SPU_020571 | NA |
| e14 | SPU_020658 | none |
| e14 | SPU_020687 | none |
| e14 | SPU_020730 | none |
| e14 | SPU_020817 | Lv-Hypp_997 |
| e14 | SPU_020827 | NA |
| e14 | SPU_020844 | Lv-Fcho2 |

| | | |
|---|---|---|
| e14 | SPU_020853 | Lv-Ubxd1 |
| e14 | SPU_020861 | Lv-Chst3L_5 |
| e14 | SPU_020874 | Lv-Rt4 |
| e14 | SPU_020943 | Lv-Gtub2f |
| e14 | SPU_021078 | Lv-Pmpcb |
| e14 | SPU_021088 | Lv-Ddx46 |
| e14 | SPU_021100 | Lv-Unc44_334 |
| e14 | SPU_021193 | Lv-Adam/Ts16/18L |
| e14 | SPU_021226 | Lv-Cybb_1 |
| e14 | SPU_021404 | Lv-Mfcad |
| e14 | SPU_021426 | Lv-Fam175b |
| e14 | SPU_021511 | Lv-Calm2 |
| e14 | SPU_021527 | Lv-Mppe1L |
| e14 | SPU_021536 | none |
| e14 | SPU_021655 | Lv-C21orf57 |
| e14 | SPU_021679 | Lv-Kiaa1055L |
| e14 | SPU_021800 | Lv-Clc2 |
| e14 | SPU_021810 | Lv-Actr5h |
| e14 | SPU_021816 | Lv-Nsf |
| e14 | SPU_021821 | Lv-PppL_131 |
| e14 | SPU_021882 | none |
| e14 | SPU_021933 | Lv-Pfas |
| e14 | SPU_021962 | Lv-Gabpb |
| e14 | SPU_022023 | Lv-Jak_1 |
| e14 | SPU_022059 | none |
| e14 | SPU_022132 | Lv-LrpL_5 |
| e14 | SPU_022172 | Lv-Cndp2 |
| e14 | SPU_022267 | Lv-Slc35b2 |
| e14 | SPU_022549 | Lv-Ivd_2 |
| e14 | SPU_022757 | none |
| e14 | SPU_022798 | Lv-Hunkc |
| e14 | SPU_022902 | Lv-Ippk |
| e14 | SPU_022912 | Lv-Flj37953l |
| e14 | SPU_023012 | Lv-Gtf2E1 |
| e14 | SPU_023025 | Lv-Amacr |
| e14 | SPU_023092 | Lv-Nudt12-2 |
| e14 | SPU_023134 | Lv-Ada_1 |
| e14 | SPU_023208 | Lv-ChbpL_2 |

| | | |
|---|---|---|
| e14 | SPU_023215 | none |
| e14 | SPU_023226 | Lv-Laptm4a |
| e14 | SPU_023237 | Lv-Fut4_3 |
| e14 | SPU_023286 | Lv-Psma1 |
| e14 | SPU_023307 | none |
| e14 | SPU_023327 | Lv-Trap5 |
| e14 | SPU_023332 | Lv-Hal |
| e14 | SPU_023397 | Lv-Fn3/Egff_4 |
| e14 | SPU_023433 | none |
| e14 | SPU_023434 | Lv-Uox |
| e14 | SPU_023461 | Lv-Calpain |
| e14 | SPU_023468 | Lv-PolypL_82 |
| e14 | SPU_023482 | Lv-Hypp_1088 |
| e14 | SPU_023523 | none |
| e14 | SPU_023531 | Lv-Gart_1 |
| e14 | SPU_023573 | Lv-Mtnr1b_2 |
| e14 | SPU_023622 | none |
| e14 | SPU_023626 | Lv-Prim2 |
| e14 | SPU_023647 | Lv-Ap3s1 |
| e14 | SPU_023725 | Lv-Samd9L_8 |
| e14 | SPU_023754 | Lv-Adora2bL_6 |
| e14 | SPU_023762 | Lv-Idi1 |
| e14 | SPU_023820 | Lv-Dym |
| e14 | SPU_023834 | Lv-Gfpt1 |
| e14 | SPU_023990 | Lv-Sart3 |
| e14 | SPU_024047 | Lv-Bcnt_1 |
| e14 | SPU_024140 | Lv-Psmd7 |
| e14 | SPU_024214 | Lv-Mrpl12-like |
| e14 | SPU_024239 | Lv-Hypp_2718 |
| e14 | SPU_024311 | Lv-Depdc6 |
| e14 | SPU_024389 | Lv-Hypp_2728 |
| e14 | SPU_024402 | Lv-Hyou1 |
| e14 | SPU_024518 | Lv-Hypp_2740 |
| e14 | SPU_024544 | Lv-Hypp_1116 |
| e14 | SPU_024579 | none |
| e14 | SPU_024646 | none |
| e14 | SPU_024669 | Lv-Wnt4_1 |
| e14 | SPU_024702 | Lv-Ptges2 |

| | | |
|---|---|---|
| e14 | SPU_024726 | none |
| e14 | SPU_024760 | Lv-PolypL_956 |
| e14 | SPU_024841 | none |
| e14 | SPU_024849 | Lv-Rft1 |
| e14 | SPU_024985 | Lv-Pfdn4 |
| e14 | SPU_024995 | Lv-KirrelL |
| e14 | SPU_025243 | Lv-Hypp_2783 |
| e14 | SPU_025256 | none |
| e14 | SPU_025287 | Lv-Hypp_2786 |
| e14 | SPU_025317 | none |
| e14 | SPU_025366 | Lv-XyltL |
| e14 | SPU_025399 | Lv-Hypp_229 |
| e14 | SPU_025410 | none |
| e14 | SPU_025482 | Lv-Ash1 |
| e14 | SPU_025491 | Lv-Hypp_2792 |
| e14 | SPU_025492 | Lv-Gbx |
| e14 | SPU_025557 | Lv-Pde1C |
| e14 | SPU_025559 | Lv-Col15/18r |
| e14 | SPU_025591 | none |
| e14 | SPU_025609 | Lv-PppL_143 |
| e14 | SPU_025622 | Lv-Cadherin_6 |
| e14 | SPU_025640 | Lv-Klhl5 |
| e14 | SPU_025676 | Lv-Spred |
| e14 | SPU_025685 | Lv-C2orf29 |
| e14 | SPU_025689 | Lv-Siahbp1 |
| e14 | SPU_025703 | NA |
| e14 | SPU_025704 | Lv-Sec14l2 |
| e14 | SPU_025706 | Lv-Fn3krp_1 |
| e14 | SPU_025888 | Lv-Maf |
| e14 | SPU_025924 | none |
| e14 | SPU_025948 | Lv-Nlr97 |
| e14 | SPU_026020 | Lv-Nlr59 |
| e14 | SPU_026068 | none |
| e14 | SPU_026071 | Lv-Nlr57 |
| e14 | SPU_026215 | Lv-Wwc1 |
| e14 | SPU_026295 | Lv-Sdhb |
| e14 | SPU_026341 | Lv-Hypp_1173 |
| e14 | SPU_026395 | Lv-Abcc1b |

| | | |
|---|---|---|
| e14 | SPU_026396 | Lv-Rab3iL1 |
| e14 | SPU_026407 | Lv-Hypp_2838 |
| e14 | SPU_026426 | Lv-Hypp_1177 |
| e14 | SPU_026475 | Lv-Ganab_1 |
| e14 | SPU_026544 | Lv-Ftcd_2 |
| e14 | SPU_026550 | Lv-Reeler/Egf/Cub_1 |
| e14 | SPU_026618 | Lv-Ndufa5 |
| e14 | SPU_026637 | none |
| e14 | SPU_026750 | Lv-Hypp_2866 |
| e14 | SPU_026815 | Lv-Gal3st1_13 |
| e14 | SPU_026960 | Lv-Ttc1 |
| e14 | SPU_026971 | none |
| e14 | SPU_026984 | Lv-QpctlL |
| e14 | SPU_027169 | Lv-Hmcn1L_4 |
| e14 | SPU_027199 | none |
| e14 | SPU_027226 | Lv-Mto1 |
| e14 | SPU_027240 | Lv-Cyb5 |
| e14 | SPU_027314 | Lv-Lpin2 |
| e14 | SPU_027334 | Lv-PaxA |
| e14 | SPU_027340 | Lv-Parp3 |
| e14 | SPU_027369 | none |
| e14 | SPU_027370 | Lv-Cdk7 |
| e14 | SPU_027419 | Lv-Slc30a7_1 |
| e14 | SPU_027430 | Lv-Alg14h |
| e14 | SPU_027494 | none |
| e14 | SPU_027523 | Lv-Mcl1 |
| e14 | SPU_027710 | Lv-Idh3a |
| e14 | SPU_027730 | Lv-Ddost |
| e14 | SPU_027734 | Lv-Cbfb |
| e14 | SPU_027780 | none |
| e14 | SPU_027783 | NA |
| e14 | SPU_027964 | Lv-Setd3 |
| e14 | SPU_028003 | Lv-Miro1 |
| e14 | SPU_028049 | Lv-Cub/Clect/Ldla/Lrr/Gpcr |
| e14 | SPU_028105 | Lv-Gbe1 |
| e14 | SPU_028135 | Lv-H1-0 |
| e14 | SPU_028140 | none |
| e14 | SPU_028156 | Lv-Hypp_2938 |

| | | |
|---|---|---|
| e14 | SPU_028209 | Lv-Rab6 |
| e14 | SPU_028221 | Lv-Atub4 |
| e14 | SPU_028251 | Lv-Mctp1L |
| e14 | SPU_028401 | Lv-Fn3/Ig |
| e14 | SPU_028405 | Lv-Tuba1c_2 |
| e14 | SPU_028441 | Lv-Pipox_4 |
| e14 | SPU_028477 | Lv-Eif4e1 |
| e14 | SPU_028504 | Lv-Atp7a_1 |
| e14 | SPU_028593 | Lv-Ptss1L |
| e14 | SPU_028659 | Lv-Psmd11 |
| e14 | SPU_028673 | none |
| e14 | SPU_028689 | Lv-Lrig1 |
| e14 | SPU_028770 | NA |
| e14 | SPU_028813 | NA |
| e14 | SPU_028829 | Lv-Trappc2L |
| e14 | SPU_028876 | Lv-Pgm2l1 |
| e14 | SPU_030062 | Lv-AnL7 |
| e14 | SPU_030142 | Lv-Cd59/Sca2L1 |
| e15 | SPU_000078 | Lv-Gpcr88 |
| e15 | SPU_000187 | Lv-Gmppb |
| e15 | SPU_000195 | NA |
| e15 | SPU_000396 | Lv-Hypp_300 |
| e15 | SPU_000397 | Lv-C4orf23L |
| e15 | SPU_000468 | Lv-Plk2 |
| e15 | SPU_000476 | Lv-Unc44L_3 |
| e15 | SPU_000607 | Lv-Acaca |
| e15 | SPU_000914 | Lv-Npfa2 |
| e15 | SPU_001020 | Lv-Surreal/Gpcr109 |
| e15 | SPU_001083 | Lv-Exoc4_2 |
| e15 | SPU_001091 | Lv-Ier3ip1 |
| e15 | SPU_001306 | none |
| e15 | SPU_001511 | Lv-Anp32bL |
| e15 | SPU_001772 | Lv-Rpl24al |
| e15 | SPU_001780 | Lv-Set |
| e15 | SPU_001796 | Lv-Frem2 |
| e15 | SPU_002052 | Lv-Hypp_3109 |
| e15 | SPU_002235 | Lv-Rfx4L |
| e15 | SPU_002341 | Lv-Fam120c |

| | | |
|---|---|---|
| e15 | SPU_002360 | Lv-Isot |
| e15 | SPU_002510 | Lv-C20orf3 |
| e15 | SPU_002714 | Lv-PicalmL |
| e15 | SPU_002779 | Lv-Hsp703B |
| e15 | SPU_002817 | Lv-Tmem111 |
| e15 | SPU_002819 | Lv-Wif |
| e15 | SPU_002869 | NA |
| e15 | SPU_002951 | none |
| e15 | SPU_003005 | NA |
| e15 | SPU_003054 | NA |
| e15 | SPU_003135 | Lv-Umps |
| e15 | SPU_003173 | Lv-Cog2-2 |
| e15 | SPU_003434 | Lv-Hypp_381 |
| e15 | SPU_003496 | Lv-Arf4/5 |
| e15 | SPU_003521 | Lv-Pes1 |
| e15 | SPU_003548 | Lv-E78a |
| e15 | SPU_003574 | none |
| e15 | SPU_003646 | Lv-Eif2a |
| e15 | SPU_003994 | Lv-Clect/Fn3_3 |
| e15 | SPU_004005 | Lv-Adap1L |
| e15 | SPU_004093 | Lv-Rsl1d1 |
| e15 | SPU_004126 | Lv-NclL_2 |
| e15 | SPU_004140 | Lv-Nox4 |
| e15 | SPU_004187 | Lv-Dip2b |
| e15 | SPU_004218 | none |
| e15 | SPU_004254 | Lv-PolypL_961 |
| e15 | SPU_004319 | Lv-CtnnbL1 |
| e15 | SPU_004337 | Lv-Nop58 |
| e15 | SPU_004348 | Lv-Hypp_413 |
| e15 | SPU_004415 | Lv-Rrp12 |
| e15 | SPU_004446 | none |
| e15 | SPU_004492 | Lv-Irsp53/58 |
| e15 | SPU_004673 | Lv-Osbp |
| e15 | SPU_004747 | Lv-Tk8 |
| e15 | SPU_004757 | Lv-Sars |
| e15 | SPU_004870 | Lv-Gcnt1 |
| e15 | SPU_004911 | Lv-Nadsyn1_2 |
| e15 | SPU_004918 | Lv-Snapc1 |

| | | |
|---|---|---|
| e15 | SPU_004919 | none |
| e15 | SPU_005096 | Lv-Dhx9 |
| e15 | SPU_005111 | Lv-Clect_11 |
| e15 | SPU_005141 | Lv-RtL_15 |
| e15 | SPU_005238 | Lv-Plxdc2 |
| e15 | SPU_005307 | Lv-Zinc318L |
| e15 | SPU_005433 | Lv-HnrpcL |
| e15 | SPU_005456 | none |
| e15 | SPU_005560 | none |
| e15 | SPU_005564 | Lv-Hypp_453 |
| e15 | SPU_005625 | Lv-Hdlbp_1 |
| e15 | SPU_005862 | Lv-Sec61b_1 |
| e15 | SPU_005873 | none |
| e15 | SPU_005991 | Lv-Clect_14 |
| e15 | SPU_006001 | Lv-Notchh1_5 |
| e15 | SPU_006048 | Lv-Me1 |
| e15 | SPU_006367 | none |
| e15 | SPU_006401 | Lv-Ugcgl1 |
| e15 | SPU_006484 | Lv-Z204 |
| e15 | SPU_006509 | Lv-Acly |
| e15 | SPU_006577 | none |
| e15 | SPU_006672 | Lv-Canx |
| e15 | SPU_006718 | Lv-Tmtc2 |
| e15 | SPU_006773 | Lv-Eif3d |
| e15 | SPU_006992 | Lv-Fut7_1 |
| e15 | SPU_006994 | Lv-C12orf30 |
| e15 | SPU_006997 | none |
| e15 | SPU_007247 | Lv-Oaz1 |
| e15 | SPU_007595 | Lv-Yars |
| e15 | SPU_007680 | Lv-Hypp_1743 |
| e15 | SPU_007687 | Lv-Ugt2b31 |
| e15 | SPU_008049 | NA |
| e15 | SPU_008059 | Lv-Kctd21_1 |
| e15 | SPU_008115 | Lv-Pontin52 |
| e15 | SPU_008413 | Lv-Mybbp1 |
| e15 | SPU_008576 | NA |
| e15 | SPU_008709 | none |
| e15 | SPU_008717 | Lv-Rio2 |

| | | |
|---|---|---|
| e15 | SPU_008720 | Lv-Hypp_557 |
| e15 | SPU_008908 | Lv-Vasa |
| e15 | SPU_008981 | Lv-Lyss |
| e15 | SPU_008993 | Lv-EhH3_21 |
| e15 | SPU_009075 | Lv-Timm10 |
| e15 | SPU_009089 | none |
| e15 | SPU_009090 | none |
| e15 | SPU_009214 | Lv-Slc36a1_1 |
| e15 | SPU_009248 | Lv-Eif3e_1 |
| e15 | SPU_009310 | none |
| e15 | SPU_009335 | Lv-Abcf1 |
| e15 | SPU_009548 | none |
| e15 | SPU_009682 | Lv-Nup188 |
| e15 | SPU_009737 | Lv-Kiaa0020 |
| e15 | SPU_009848 | Lv-Vparpf_1 |
| e15 | SPU_009862 | none |
| e15 | SPU_009872 | Lv-Slc25a3_2 |
| e15 | SPU_009951 | none |
| e15 | SPU_010030 | Lv-Cog6 |
| e15 | SPU_010114 | Lv-GtpbpL3(nucleolar) |
| e15 | SPU_010227 | Lv-Srcr85 |
| e15 | SPU_010282 | Lv-Nol1 |
| e15 | SPU_010299 | none |
| e15 | SPU_010303 | Lv-Eif3k |
| e15 | SPU_010362 | none |
| e15 | SPU_010379 | Lv-Sec23ip |
| e15 | SPU_010480 | none |
| e15 | SPU_010585 | Lv-Wdr43 |
| e15 | SPU_010636 | Lv-Mftc |
| e15 | SPU_010699 | Lv-Polr1C |
| e15 | SPU_010758 | Lv-Sdad1 |
| e15 | SPU_010759 | Lv-Slc16a12L_1 |
| e15 | SPU_010823 | Lv-Rptp_mu |
| e15 | SPU_010849 | none |
| e15 | SPU_010853 | Lv-Dhx15 |
| e15 | SPU_010941 | Lv-Rrs1 |
| e15 | SPU_011013 | Lv-Hypp_635 |
| e15 | SPU_011016 | Lv-ColmL |

| | | |
|---|---|---|
| e15 | SPU_011744 | Lv-Ltv1 |
| e15 | SPU_012000 | Lv-Pwp2h |
| e15 | SPU_012013 | Lv-Hypp_2011 |
| e15 | SPU_012031 | Lv-Frem2_1 |
| e15 | SPU_012091 | Lv-Nfkbil1L |
| e15 | SPU_012161 | Lv-Gpr52L_5 |
| e15 | SPU_012189 | Lv-Lpcat3_1 |
| e15 | SPU_012216 | Lv-Eif3i |
| e15 | SPU_012350 | Lv-Igf_14 |
| e15 | SPU_012366 | Lv-Pnlip_4 |
| e15 | SPU_012455 | none |
| e15 | SPU_012533 | none |
| e15 | SPU_012614 | NA |
| e15 | SPU_012747 | none |
| e15 | SPU_012807 | Lv-NclL |
| e15 | SPU_012988 | Lv-Tiprl |
| e15 | SPU_013061 | none |
| e15 | SPU_013077 | Lv-Hypp_87 |
| e15 | SPU_013171 | Lv-Hp_Loc81556L |
| e15 | SPU_013174 | Lv-Ssr1 |
| e15 | SPU_013346 | Lv-Chml |
| e15 | SPU_013384 | Lv-MyXII_1 |
| e15 | SPU_013434 | none |
| e15 | SPU_013533 | Lv-B3gnt5_4 |
| e15 | SPU_013567 | none |
| e15 | SPU_013577 | Lv-Smarcb1 |
| e15 | SPU_013584 | none |
| e15 | SPU_013659 | Lv-C7orf30L |
| e15 | SPU_013719 | Lv-Cog2 |
| e15 | SPU_013755 | Lv-Nip7 |
| e15 | SPU_013856 | Lv-Usp14 |
| e15 | SPU_013905 | NA |
| e15 | SPU_014066 | Lv-Bop1 |
| e15 | SPU_014113 | none |
| e15 | SPU_014261 | Lv-Wdr75 |
| e15 | SPU_014267 | Lv-Lcmt2_1 |
| e15 | SPU_014312 | Lv-Pmm2 |
| e15 | SPU_014348 | Lv-Dio1L |

| | | |
|---|---|---|
| e15 | SPU_014505 | Lv-Ddx56_1 |
| e15 | SPU_014823 | Lv-Hypp_764 |
| e15 | SPU_014998 | Lv-Adrb2L_8 |
| e15 | SPU_015001 | Lv-Nphs1L_1 |
| e15 | SPU_015048 | Lv-Acss2 |
| e15 | SPU_015068 | NA |
| e15 | SPU_015142 | Lv-Atp13a3_1 |
| e15 | SPU_015336 | none |
| e15 | SPU_015342 | Lv-Aqrh |
| e15 | SPU_015352 | Lv-Dcst2 |
| e15 | SPU_015353 | Lv-Dcst2_1 |
| e15 | SPU_015397 | none |
| e15 | SPU_015448 | Lv-Stk11ip |
| e15 | SPU_015598 | Lv-Chst14 |
| e15 | SPU_015693 | Lv-Hypp_2225 |
| e15 | SPU_015705 | Lv-Fn3/Igf_13 |
| e15 | SPU_015743 | Lv-Rnf121 |
| e15 | SPU_015804 | Lv-Mphosph6 |
| e15 | SPU_015811 | Lv-Fkbp39L |
| e15 | SPU_015822 | Lv-Chst14_2 |
| e15 | SPU_015920 | Lv-Afi |
| e15 | SPU_015927 | Lv-Mki67ipL |
| e15 | SPU_016029 | none |
| e15 | SPU_016101 | Lv-Chaf1a |
| e15 | SPU_016170 | Lv-Hypp_816 |
| e15 | SPU_016267 | Lv-Slc45 |
| e15 | SPU_016408 | Lv-Slc16a6_3 |
| e15 | SPU_016547 | Lv-Sfrs7l |
| e15 | SPU_016652 | Lv-Ddo_1 |
| e15 | SPU_016653 | Lv-Xpmc2hL |
| e15 | SPU_016665 | Lv-Lgtnl |
| e15 | SPU_016726 | Lv-Lars |
| e15 | SPU_016729 | Lv-4Ehp |
| e15 | SPU_016739 | Lv-Hdlbp |
| e15 | SPU_016774 | Lv-Hypp_2279 |
| e15 | SPU_016822 | Lv-Eif3c_1 |
| e15 | SPU_016959 | Lv-Hcrtr2L_1 |
| e15 | SPU_017028 | Lv-Ddx27 |

| | | |
|---|---|---|
| e15 | SPU_017030 | none |
| e15 | SPU_017052 | Lv-Stag3 |
| e15 | SPU_017059 | Lv-Top3B |
| e15 | SPU_017076 | Lv-Heatr3 |
| e15 | SPU_017082 | Lv-Nhp2l1 |
| e15 | SPU_017209 | none |
| e15 | SPU_017215 | Lv-Scy1L1bp1 |
| e15 | SPU_017333 | Lv-Bxdc1 |
| e15 | SPU_017347 | Lv-Gmds |
| e15 | SPU_017374 | Lv-Fmo3 |
| e15 | SPU_017412 | NA |
| e15 | SPU_017644 | NA |
| e15 | SPU_017781 | Lv-Asps_1 |
| e15 | SPU_017854 | none |
| e15 | SPU_017872 | NA |
| e15 | SPU_017896 | Lv-St8sia3 |
| e15 | SPU_018054 | Lv-Frem2_2 |
| e15 | SPU_018070 | Lv-Fn3_5 |
| e15 | SPU_018081 | Lv-Rpb7 |
| e15 | SPU_018185 | Lv-Tmem87a |
| e15 | SPU_018189 | Lv-Hypp_90 |
| e15 | SPU_018219 | Lv-Hnrph1 |
| e15 | SPU_018338 | none |
| e15 | SPU_018440 | Lv-Snd1 |
| e15 | SPU_018468 | Lv-Cyp2L36 |
| e15 | SPU_018469 | Lv-Adrb2L_11 |
| e15 | SPU_018596 | Lv-Vprbp |
| e15 | SPU_018797 | Lv-Evi1 |
| e15 | SPU_018800 | Lv-Nup37 |
| e15 | SPU_018801 | Lv-Mrpl23 |
| e15 | SPU_018810 | Lv-Sm32 |
| e15 | SPU_018982 | Lv-My10f_1 |
| e15 | SPU_019006 | none |
| e15 | SPU_019011 | Lv-Gpn1 |
| e15 | SPU_019091 | NA |
| e15 | SPU_019323 | Lv-MifL2 |
| e15 | SPU_019415 | Lv-Nvll_1 |
| e15 | SPU_019462 | Lv-Polr2h |

| | | |
|---|---|---|
| e15 | SPU_019584 | none |
| e15 | SPU_019647 | Lv-C20orf3_1 |
| e15 | SPU_019802 | none |
| e15 | SPU_019803 | Lv-Bamacan |
| e15 | SPU_019881 | NA |
| e15 | SPU_019904 | none |
| e15 | SPU_019931 | Lv-Ptcd3 |
| e15 | SPU_019947 | Lv-Hypp_2474 |
| e15 | SPU_020162 | Lv-Slc16a2_1 |
| e15 | SPU_020322 | Lv-Hsp902a1 |
| e15 | SPU_020412 | Lv-Eif2g |
| e15 | SPU_020492 | Lv-Atpaf1 |
| e15 | SPU_020584 | Lv-Acac |
| e15 | SPU_020778 | Lv-Emg1h |
| e15 | SPU_020801 | Lv-Tars |
| e15 | SPU_020989 | Lv-Unc44_248 |
| e15 | SPU_021137 | Lv-Hypp_2539 |
| e15 | SPU_021305 | none |
| e15 | SPU_021311 | Lv-Copb2 |
| e15 | SPU_021419 | Lv-Phf10 |
| e15 | SPU_021449 | Lv-Eif3eip |
| e15 | SPU_021472 | none |
| e15 | SPU_021718 | none |
| e15 | SPU_021747 | Lv-Igf_20 |
| e15 | SPU_021801 | none |
| e15 | SPU_021802 | Lv-Mvd |
| e15 | SPU_021883 | Lv-SnrpE_1 |
| e15 | SPU_021934 | NA |
| e15 | SPU_022003 | NA |
| e15 | SPU_022158 | Lv-Hsp701H |
| e15 | SPU_022286 | Lv-Srcr186 |
| e15 | SPU_022537 | Lv-14-3-3_2 |
| e15 | SPU_022539 | Lv-Noc2L |
| e15 | SPU_022557 | Lv-FibropellinL2 |
| e15 | SPU_022603 | Lv-Uprt |
| e15 | SPU_022635 | none |
| e15 | SPU_022676 | none |
| e15 | SPU_022783 | Lv-Rfk |

| | | |
|---|---|---|
| e15 | SPU_022803 | Lv-Hypp_2632 |
| e15 | SPU_023016 | Lv-Frem1L |
| e15 | SPU_023021 | none |
| e15 | SPU_023101 | Lv-Slsp_15 |
| e15 | SPU_023143 | Lv-Cirh1a |
| e15 | SPU_023207 | none |
| e15 | SPU_023209 | Lv-Hypp_2655 |
| e15 | SPU_023290 | NA |
| e15 | SPU_023295 | Lv-Ddx6 |
| e15 | SPU_023420 | Lv-Slc35a3 |
| e15 | SPU_023819 | Lv-Mcm6 |
| e15 | SPU_023852 | Lv-Mgst3_1 |
| e15 | SPU_023965 | Lv-Eif3s3L |
| e15 | SPU_024067 | Lv-Thoc5 |
| e15 | SPU_024082 | Lv-Hsp90L |
| e15 | SPU_024244 | none |
| e15 | SPU_024260 | Lv-Bysl |
| e15 | SPU_024279 | none |
| e15 | SPU_024436 | Lv-Unc44_253 |
| e15 | SPU_024514 | Lv-Nudcd1 |
| e15 | SPU_024553 | Lv-Tinp1 |
| e15 | SPU_024680 | Lv-C20orf54L_2 |
| e15 | SPU_024696 | Lv-Tmem186 |
| e15 | SPU_024859 | Lv-Eif4Gg |
| e15 | SPU_024911 | Lv-Psmd1_1 |
| e15 | SPU_024926 | none |
| e15 | SPU_025016 | Lv-Arcn1 |
| e15 | SPU_025036 | Lv-B3galt1_14 |
| e15 | SPU_025140 | Lv-Pb1 |
| e15 | SPU_025213 | none |
| e15 | SPU_025234 | Lv-Rbm26 |
| e15 | SPU_025251 | Lv-Fbl |
| e15 | SPU_025416 | Lv-Ddx50 |
| e15 | SPU_025570 | Lv-Wdr12 |
| e15 | SPU_025699 | Lv-P75ntr |
| e15 | SPU_025766 | Lv-Rptp_mu_2 |
| e15 | SPU_025777 | Lv-Tomm40 |
| e15 | SPU_025797 | Lv-Gtpbp4L_1 |

| | | |
|---|---|---|
| e15 | SPU_025814 | Lv-Ranbp1 |
| e15 | SPU_025913 | Lv-PppL_146 |
| e15 | SPU_025967 | Lv-Gpr52 |
| e15 | SPU_026178 | Lv-Abcb9C |
| e15 | SPU_026263 | Lv-Rbbp4 |
| e15 | SPU_026340 | Lv-Atic |
| e15 | SPU_026454 | Lv-Mamdc4-13 |
| e15 | SPU_026485 | none |
| e15 | SPU_026552 | Lv-Slc17a5_3 |
| e15 | SPU_026628 | Lv-Smc1L1_1 |
| e15 | SPU_026693 | Lv-Tmem48 |
| e15 | SPU_026703 | Lv-Hypp_2862 |
| e15 | SPU_026791 | none |
| e15 | SPU_026862 | Lv-Ftsj3 |
| e15 | SPU_026863 | Lv-Eif3c |
| e15 | SPU_026932 | none |
| e15 | SPU_026967 | NA |
| e15 | SPU_027047 | Lv-Nol6 |
| e15 | SPU_027107 | Lv-Txndc14 |
| e15 | SPU_027149 | Lv-Mrpl4 |
| e15 | SPU_027229 | Lv-Ugt2b1_4 |
| e15 | SPU_027331 | Lv-Tk5 |
| e15 | SPU_027472 | Lv-Ia1 |
| e15 | SPU_027518 | none |
| e15 | SPU_027551 | none |
| e15 | SPU_027637 | Lv-Xpot1 |
| e15 | SPU_027646 | NA |
| e15 | SPU_027684 | none |
| e15 | SPU_027943 | Lv-Hypp_2923 |
| e15 | SPU_027984 | none |
| e15 | SPU_027994 | none |
| e15 | SPU_028005 | none |
| e15 | SPU_028123 | none |
| e15 | SPU_028148 | Lv-Ac/Sc |
| e15 | SPU_028265 | Lv-Usp10 |
| e15 | SPU_028312 | Lv-Pp5c |
| e15 | SPU_028447 | none |
| e15 | SPU_028467 | Lv-Pdcd10 |

| | | |
|---|---|---|
| e15 | SPU_028677 | Lv-Tfb1m |
| e15 | SPU_028803 | Lv-GhdcL_5 |
| e15 | SPU_028828 | Lv-Pabpn1 |
| e15 | SPU_028860 | none |
| e15 | SPU_028897 | Lv-Apn6 |
| e15 | SPU_030138 | NA |
| e16 | SPU_000091 | Lv-Fimbph |
| e16 | SPU_000129 | Lv-Arnt |
| e16 | SPU_000177 | Lv-Pcsk9l |
| e16 | SPU_000185 | Lv-Hypp_1245 |
| e16 | SPU_000265 | none |
| e16 | SPU_000296 | Lv-Gelslnph |
| e16 | SPU_000302 | none |
| e16 | SPU_000479 | Lv-Ccdc63 |
| e16 | SPU_000552 | Lv-Arf1 |
| e16 | SPU_000620 | Lv-Lrp12L_2 |
| e16 | SPU_000723 | Lv-76p |
| e16 | SPU_000729 | Lv-PlexA4_3 |
| e16 | SPU_000934 | Lv-Hypp_318 |
| e16 | SPU_000961 | Lv-Ptb |
| e16 | SPU_001094 | Lv-Rab18 |
| e16 | SPU_001123 | Lv-Xdh |
| e16 | SPU_001154 | Lv-Clip170ph3 |
| e16 | SPU_001191 | Lv-Gluld1 |
| e16 | SPU_001269 | Lv-Cpox_1 |
| e16 | SPU_001332 | Lv-Ap1g1 |
| e16 | SPU_001344 | Lv-Pr65a |
| e16 | SPU_001392 | Lv-Eif5B |
| e16 | SPU_001512 | Lv-Gtf2E2 |
| e16 | SPU_001513 | Lv-TxnL1 |
| e16 | SPU_001579 | Lv-Vdp |
| e16 | SPU_001640 | Lv-Syt14 |
| e16 | SPU_001822 | Lv-Rpa1 |
| e16 | SPU_001927 | Lv-Eps15l1l |
| e16 | SPU_002003 | Lv-Hnrpa2B1L |
| e16 | SPU_002093 | NA |
| e16 | SPU_002213 | none |
| e16 | SPU_002302 | Lv-Trmu |

| | | |
|---|---|---|
| e16 | SPU_002389 | Lv-Cox18h |
| e16 | SPU_002414 | Lv-Gemin4L |
| e16 | SPU_002417 | NA |
| e16 | SPU_002475 | Lv-AccsL |
| e16 | SPU_002488 | Lv-Galnac4s-6stL_2 |
| e16 | SPU_002520 | Lv-Ugp2 |
| e16 | SPU_002608 | Lv-Vc2_1 |
| e16 | SPU_002649 | Lv-Pfkfb1 |
| e16 | SPU_002927 | Lv-Slc25a15 |
| e16 | SPU_003022 | Lv-Ugt1L_10 |
| e16 | SPU_003040 | Lv-Hypp_373 |
| e16 | SPU_003130 | Lv-Spg11 |
| e16 | SPU_003140 | Lv-Bet1_1 |
| e16 | SPU_003194 | Lv-Sim_wsk_ald_syn_ph |
| e16 | SPU_003321 | Lv-PolppL_29 |
| e16 | SPU_003447 | none |
| e16 | SPU_003552 | Lv-Zzef1 |
| e16 | SPU_003556 | Lv-Clic6 |
| e16 | SPU_003585 | Lv-Drk |
| e16 | SPU_003586 | NA |
| e16 | SPU_003617 | none |
| e16 | SPU_003811 | NA |
| e16 | SPU_003822 | Lv-Lamb2 |
| e16 | SPU_003848 | Lv-Z298 |
| e16 | SPU_003877 | Lv-Acsf2-2 |
| e16 | SPU_003906 | Lv-Yipf5 |
| e16 | SPU_003992 | Lv-Clect/Fn3_1 |
| e16 | SPU_004064 | Lv-Ttc37 |
| e16 | SPU_004165 | Lv-Nlr84 |
| e16 | SPU_004266 | Lv-Deltat |
| e16 | SPU_004268 | Lv-Akap10 |
| e16 | SPU_004297 | Lv-Ap1m1 |
| e16 | SPU_004367 | Lv-Med23 |
| e16 | SPU_004378 | Lv-Hypp_414 |
| e16 | SPU_004388 | Lv-Cog8_1 |
| e16 | SPU_004468 | Lv-Rfc3 |
| e16 | SPU_004522 | Lv-Tbce |
| e16 | SPU_004592 | Lv-Slc16a4 |

| | | |
|---|---|---|
| e16 | SPU_004604 | Lv-Gtpbp |
| e16 | SPU_004662 | Lv-Commd2 |
| e16 | SPU_004697 | none |
| e16 | SPU_004760 | Lv-Zer1 |
| e16 | SPU_004821 | Lv-Cnot1_1 |
| e16 | SPU_004841 | Lv-Crhbp |
| e16 | SPU_004854 | Lv-Sord |
| e16 | SPU_004857 | Lv-Paaf1 |
| e16 | SPU_004873 | Lv-Spg11L_2 |
| e16 | SPU_004936 | NA |
| e16 | SPU_004958 | none |
| e16 | SPU_005217 | NA |
| e16 | SPU_005227 | Lv-Mrps30 |
| e16 | SPU_005387 | none |
| e16 | SPU_005393 | Lv-Gtpbp6 |
| e16 | SPU_005553 | none |
| e16 | SPU_005672 | Lv-St3gal3 |
| e16 | SPU_005677 | none |
| e16 | SPU_005832 | Lv-Tlr028 |
| e16 | SPU_005836 | Lv-Lipk |
| e16 | SPU_005839 | Lv-Aco2 |
| e16 | SPU_005883 | Lv-Pkdcc |
| e16 | SPU_005919 | Lv-Hypp_3075 |
| e16 | SPU_005955 | Lv-Notchl1 |
| e16 | SPU_006000 | Lv-Notchh1_4 |
| e16 | SPU_006047 | Lv-Cct8 |
| e16 | SPU_006153 | Lv-Galk1 |
| e16 | SPU_006165 | Lv-LrpL_1 |
| e16 | SPU_006261 | Lv-Aprt |
| e16 | SPU_006296 | Lv-06296 |
| e16 | SPU_006348 | Lv-Def8 |
| e16 | SPU_006412 | Lv-Cophs3 |
| e16 | SPU_006466 | Lv-Fanci |
| e16 | SPU_006473 | none |
| e16 | SPU_006558 | Lv-Lamg1 |
| e16 | SPU_006682 | Lv-Slc25a21 |
| e16 | SPU_006930 | Lv-Egfia_1 |
| e16 | SPU_007057 | Lv-Mrpl22 |

| | | |
|---|---|---|
| e16 | SPU_007067 | none |
| e16 | SPU_007209 | Lv-Heatr7a |
| e16 | SPU_007254 | Lv-Ccdc40 |
| e16 | SPU_007258 | none |
| e16 | SPU_007327 | Lv-Mrpl2l |
| e16 | SPU_007343 | Lv-Myd88 |
| e16 | SPU_007362 | Lv-C1orf27L |
| e16 | SPU_007377 | Lv-Polr3d |
| e16 | SPU_007411 | Lv-C6orf57L |
| e16 | SPU_007427 | Lv-Mett5d1 |
| e16 | SPU_007472 | Lv-PolypL_937 |
| e16 | SPU_007502 | Lv-Glb1 |
| e16 | SPU_007515 | none |
| e16 | SPU_007539 | none |
| e16 | SPU_007586 | Lv-Acbd3L |
| e16 | SPU_007610 | none |
| e16 | SPU_007655 | Lv-Cdk2 |
| e16 | SPU_007670 | Lv-Psmc5 |
| e16 | SPU_007845 | NA |
| e16 | SPU_007937 | Lv-Ldla/Gpcr_2 |
| e16 | SPU_007962 | Lv-Nedd4 |
| e16 | SPU_008009 | Lv-Aldh4a1 |
| e16 | SPU_008069 | Lv-Aco2_1 |
| e16 | SPU_008128 | Lv-Ttc33 |
| e16 | SPU_008133 | Lv-Porcupine |
| e16 | SPU_008152 | Lv-Cyp2L16 |
| e16 | SPU_008264 | Lv-Kiaa0564pL |
| e16 | SPU_008506 | none |
| e16 | SPU_008678 | Lv-Pkcd |
| e16 | SPU_008779 | Lv-Mapko1 |
| e16 | SPU_008834 | none |
| e16 | SPU_008844 | Lv-Ppef2 |
| e16 | SPU_008877 | Lv-Alkbh5 |
| e16 | SPU_008880 | Lv-Clec16a |
| e16 | SPU_008970 | Lv-Bckdhb |
| e16 | SPU_009015 | Lv-ProteoliaisinL |
| e16 | SPU_009160 | Lv-Nlr125 |
| e16 | SPU_009227 | Lv-Fam119a |

| | | |
|---|---|---|
| e16 | SPU_009246 | Lv-Cobra1 |
| e16 | SPU_009304 | Lv-DolkL |
| e16 | SPU_009360 | Lv-Acadvl |
| e16 | SPU_009387 | Lv-Scfd2 |
| e16 | SPU_009446 | Lv-Asns_4 |
| e16 | SPU_009449 | Lv-Sarox |
| e16 | SPU_009452 | none |
| e16 | SPU_009454 | Lv-Mgat5 |
| e16 | SPU_009530 | Lv-B3galt1L_2 |
| e16 | SPU_009592 | Lv-Tpt1l |
| e16 | SPU_009728 | none |
| e16 | SPU_009775 | Lv-Sult1a4 |
| e16 | SPU_009795 | Lv-Kiaa0556_1 |
| e16 | SPU_009858 | none |
| e16 | SPU_009912 | NA |
| e16 | SPU_010005 | NA |
| e16 | SPU_010014 | Lv-Ubr2 |
| e16 | SPU_010023 | NA |
| e16 | SPU_010035 | Lv-Cox4nb |
| e16 | SPU_010337 | none |
| e16 | SPU_010342 | none |
| e16 | SPU_010418 | Lv-Commd1 |
| e16 | SPU_010423 | Lv-Ehbp1L |
| e16 | SPU_010446 | none |
| e16 | SPU_010563 | Lv-Cyp2L23 |
| e16 | SPU_010670 | Lv-Sbdsh |
| e16 | SPU_010671 | Lv-Tmpitl |
| e16 | SPU_010679 | Lv-Hypp_1934 |
| e16 | SPU_010688 | none |
| e16 | SPU_010733 | Lv-Hspc105 |
| e16 | SPU_010805 | Lv-Akt |
| e16 | SPU_010866 | Lv-Mmd2l |
| e16 | SPU_010875 | none |
| e16 | SPU_010914 | Lv-Z25_1 |
| e16 | SPU_010952 | Lv-Mrpl16 |
| e16 | SPU_010966 | Lv-Afg3L |
| e16 | SPU_010988 | Lv-Hrh4L_4 |
| e16 | SPU_011025 | Lv-Hypp_637 |

| | | |
|---|---|---|
| e16 | SPU_011063 | Lv-Irs1 |
| e16 | SPU_011213 | Lv-Srr_3 |
| e16 | SPU_011279 | Lv-Plekhg4 |
| e16 | SPU_011303 | Lv-Dlg2 |
| e16 | SPU_011308 | Lv-Egfi3 |
| e16 | SPU_011309 | Lv-Svep1L_2 |
| e16 | SPU_011365 | Lv-Fibp |
| e16 | SPU_011564 | Lv-Hypp_70 |
| e16 | SPU_011594 | none |
| e16 | SPU_011618 | Lv-Hypp_8991 |
| e16 | SPU_011652 | Lv-FancbL |
| e16 | SPU_011694 | Lv-Fnbp1l_1 |
| e16 | SPU_011801 | Lv-PolypL_941 |
| e16 | SPU_011829 | Lv-Clect/Pan_11 |
| e16 | SPU_011861 | Lv-Arfip1 |
| e16 | SPU_011964 | Lv-Pqlc1 |
| e16 | SPU_012009 | Lv-Dock180 |
| e16 | SPU_012047 | Lv-Jagnh1L |
| e16 | SPU_012088 | none |
| e16 | SPU_012116 | Lv-Tsfm |
| e16 | SPU_012125 | none |
| e16 | SPU_012215 | Lv-Clect/Fn3_6 |
| e16 | SPU_012301 | Lv-Dhps_1 |
| e16 | SPU_012320 | Lv-Ccm2 |
| e16 | SPU_012333 | Lv-Hypp_2027 |
| e16 | SPU_012349 | Lv-Psmd5 |
| e16 | SPU_012654 | Lv-Egfib |
| e16 | SPU_012714 | Lv-Hypp_2046 |
| e16 | SPU_012814 | Lv-Slc25a11 |
| e16 | SPU_012887 | Lv-Msh2 |
| e16 | SPU_013045 | Lv-Exoc6 |
| e16 | SPU_013050 | Lv-Unc44L_86 |
| e16 | SPU_013161 | Lv-My15 |
| e16 | SPU_013242 | Lv-Lypla1 |
| e16 | SPU_013257 | Lv-Elach1 |
| e16 | SPU_013265 | none |
| e16 | SPU_013326 | Lv-Ak2_1 |
| e16 | SPU_013338 | Lv-Gluld1_14 |

| | | |
|---|---|---|
| e16 | SPU_013527 | Lv-Ints8 |
| e16 | SPU_013564 | none |
| e16 | SPU_013661 | none |
| e16 | SPU_013740 | Lv-Hypp_2100 |
| e16 | SPU_013756 | Lv-Ppib |
| e16 | SPU_013797 | none |
| e16 | SPU_013811 | Lv-Lpcat3 |
| e16 | SPU_013849 | Lv-PolppL_104 |
| e16 | SPU_013887 | none |
| e16 | SPU_014059 | Lv-Ids_2 |
| e16 | SPU_014190 | none |
| e16 | SPU_014198 | Lv-Hypp_2136 |
| e16 | SPU_014207 | NA |
| e16 | SPU_014255 | NA |
| e16 | SPU_014259 | Lv-Tfg |
| e16 | SPU_014313 | Lv-Tmed2 |
| e16 | SPU_014365 | Lv-Sec22_1 |
| e16 | SPU_014515 | Lv-Gi[cterm] |
| e16 | SPU_014582 | none |
| e16 | SPU_014778 | NA |
| e16 | SPU_014797 | Lv-PolypL_7 |
| e16 | SPU_014833 | none |
| e16 | SPU_014968 | NA |
| e16 | SPU_015004 | Lv-Dscc1h |
| e16 | SPU_015152 | Lv-Dtwd1 |
| e16 | SPU_015158 | Lv-PolypL_948 |
| e16 | SPU_015187 | Lv-Eif3L |
| e16 | SPU_015316 | NA |
| e16 | SPU_015317 | Lv-Pabpc1 |
| e16 | SPU_015363 | Lv-TincL |
| e16 | SPU_015370 | Lv-C1orf2L |
| e16 | SPU_015399 | Lv-Mkln1 |
| e16 | SPU_015482 | Lv-Lamb1L |
| e16 | SPU_015522 | NA |
| e16 | SPU_015644 | Lv-Snx8 |
| e16 | SPU_015687 | Lv-Fars2 |
| e16 | SPU_015759 | Lv-Hypp_799 |
| e16 | SPU_015762 | Lv-RtL_40 |

| | | |
|---|---|---|
| e16 | SPU_015936 | Lv-Wdr4 |
| e16 | SPU_015962 | Lv-Clect/Pan_5 |
| e16 | SPU_015981 | Lv-Dlst |
| e16 | SPU_016056 | Lv-Cyp3L8 |
| e16 | SPU_016142 | Lv-Glt_1 |
| e16 | SPU_016193 | none |
| e16 | SPU_016198 | Lv-Ptss4L |
| e16 | SPU_016361 | Lv-GBF1 |
| e16 | SPU_016382 | Lv-Ard1 |
| e16 | SPU_016415 | none |
| e16 | SPU_016439 | Lv-Eef1e1 |
| e16 | SPU_016606 | Lv-Ocrl1 |
| e16 | SPU_016609 | Lv-Bcdo |
| e16 | SPU_016617 | Lv-Tulp4L |
| e16 | SPU_016639 | Lv-Tmem38b |
| e16 | SPU_016761 | Lv-Pin1 |
| e16 | SPU_016886 | Lv-Drg2 |
| e16 | SPU_016978 | Lv-RtL_45 |
| e16 | SPU_017040 | Lv-Fbxw5 |
| e16 | SPU_017058 | Lv-Cbs |
| e16 | SPU_017277 | Lv-Hypp_2316 |
| e16 | SPU_017350 | Lv-Pdia5 |
| e16 | SPU_017357 | Lv-Hypp_877 |
| e16 | SPU_017414 | Lv-RarS |
| e16 | SPU_017484 | Lv-Sashp |
| e16 | SPU_017594 | Lv-Dis3L |
| e16 | SPU_017606 | Lv-Ddx59 |
| e16 | SPU_017651 | Lv-Hypp_2350 |
| e16 | SPU_017653 | Lv-Hypp_888 |
| e16 | SPU_017726 | Lv-Lktnh |
| e16 | SPU_017891 | Lv-Hypp_2359 |
| e16 | SPU_017897 | Lv-Gtf2E2_1 |
| e16 | SPU_017945 | Lv-Aig1 |
| e16 | SPU_017954 | Lv-Prkcshl |
| e16 | SPU_017960 | none |
| e16 | SPU_017968 | Lv-Slc46a1L_5 |
| e16 | SPU_018102 | Lv-Hs6st2 |
| e16 | SPU_018146 | Lv-Rpp38 |

| | | |
|---|---|---|
| e16 | SPU_018320 | Lv-Rt13 |
| e16 | SPU_018429 | Lv-Srcr142 |
| e16 | SPU_018438 | Lv-Coro7 |
| e16 | SPU_018454 | Lv-Klhl12L_2 |
| e16 | SPU_018510 | none |
| e16 | SPU_018532 | Lv-EgfibL |
| e16 | SPU_018595 | NA |
| e16 | SPU_018656 | Lv-Magt1 |
| e16 | SPU_018780 | Lv-Mapkbp1L_1 |
| e16 | SPU_018823 | Lv-Orc5 |
| e16 | SPU_018955 | Lv-Cnot1_3 |
| e16 | SPU_018992 | Lv-Hemk1 |
| e16 | SPU_019286 | Lv-Copg2 |
| e16 | SPU_019378 | Lv-Hk1_1 |
| e16 | SPU_019393 | Lv-Rexo2 |
| e16 | SPU_019423 | Lv-Lsm6 |
| e16 | SPU_019511 | Lv-Myof |
| e16 | SPU_019613 | Lv-Notch1_16 |
| e16 | SPU_019653 | Lv-Z309 |
| e16 | SPU_019731 | Lv-Ivnslabp |
| e16 | SPU_019855 | Lv-PppL_123 |
| e16 | SPU_019864 | Lv-Pfas_2 |
| e16 | SPU_019957 | Lv-Etfb |
| e16 | SPU_020018 | none |
| e16 | SPU_020083 | Lv-Ranbp2L_3 |
| e16 | SPU_020130 | Lv-Nup43 |
| e16 | SPU_020149 | Lv-Rpn1 |
| e16 | SPU_020204 | Lv-Endrvt39 |
| e16 | SPU_020264 | Lv-Rfc5 |
| e16 | SPU_020291 | Lv-Igf_4 |
| e16 | SPU_020315 | Lv-Tmprss2L |
| e16 | SPU_020327 | Lv-Hypp_970 |
| e16 | SPU_020335 | Lv-Fn3f_11 |
| e16 | SPU_020601 | none |
| e16 | SPU_020616 | Lv-Lrrc68 |
| e16 | SPU_020624 | Lv-Trappc3 |
| e16 | SPU_020663 | NA |
| e16 | SPU_020802 | Lv-Snx27 |

| | | |
|---|---|---|
| e16 | SPU_020968 | Lv-MucD |
| e16 | SPU_021003 | NA |
| e16 | SPU_021071 | Lv-PMF1 |
| e16 | SPU_021138 | Lv-Hypp_1013 |
| e16 | SPU_021139 | Lv-Hypp_1014 |
| e16 | SPU_021171 | Lv-Bpnt1 |
| e16 | SPU_021203 | Lv-AT |
| e16 | SPU_021303 | Lv-Pmca |
| e16 | SPU_021394 | Lv-Ehd1 |
| e16 | SPU_021422 | none |
| e16 | SPU_021552 | Lv-Srpk2 |
| e16 | SPU_021558 | Lv-Caf1 |
| e16 | SPU_021634 | none |
| e16 | SPU_021736 | Lv-Nxn |
| e16 | SPU_021768 | Lv-Hypp_2577 |
| e16 | SPU_021806 | Lv-Mett11d1 |
| e16 | SPU_021872 | Lv-Hypp_2581 |
| e16 | SPU_021942 | Lv-Grb14 |
| e16 | SPU_021968 | none |
| e16 | SPU_022140 | Lv-Mtr_1 |
| e16 | SPU_022293 | Lv-Unc44_252 |
| e16 | SPU_022305 | Lv-Thoc2 |
| e16 | SPU_022316 | NA |
| e16 | SPU_022351 | Lv-Mrps35 |
| e16 | SPU_022628 | Lv-Fads2_1 |
| e16 | SPU_022636 | Lv-Dock3 |
| e16 | SPU_022696 | none |
| e16 | SPU_022877 | Lv-Degs2 |
| e16 | SPU_022924 | Lv-Grhpr |
| e16 | SPU_022930 | Lv-Coro7_1 |
| e16 | SPU_023029 | none |
| e16 | SPU_023083 | Lv-Eif4A |
| e16 | SPU_023149 | Lv-Ercc2_1 |
| e16 | SPU_023174 | none |
| e16 | SPU_023471 | Lv-Trip4 |
| e16 | SPU_023519 | Lv-Cd2bp2 |
| e16 | SPU_023670 | Lv-Ciapin1 |
| e16 | SPU_023769 | Lv-Cbwd1 |

| | | |
|---|---|---|
| e16 | SPU_023829 | Lv-Alg6 |
| e16 | SPU_023952 | Lv-Abl |
| e16 | SPU_023978 | none |
| e16 | SPU_024008 | Lv-PhaxL |
| e16 | SPU_024103 | Lv-HspD1 |
| e16 | SPU_024110 | Lv-Act_4 |
| e16 | SPU_024275 | NA |
| e16 | SPU_024295 | NA |
| e16 | SPU_024310 | Lv-Myeov2L |
| e16 | SPU_024388 | Lv-Hypp_2727 |
| e16 | SPU_024445 | Lv-Arih1 |
| e16 | SPU_024510 | Lv-Ccdc6 |
| e16 | SPU_024529 | Lv-Dnah5 |
| e16 | SPU_024536 | none |
| e16 | SPU_024560 | Lv-Hdhd1a |
| e16 | SPU_024620 | Lv-Cttn |
| e16 | SPU_024666 | Lv-Abcb8 |
| e16 | SPU_024719 | Lv-Glrx5 |
| e16 | SPU_024777 | Lv-Fn3_10 |
| e16 | SPU_024810 | Lv-Setd6 |
| e16 | SPU_024931 | none |
| e16 | SPU_025086 | Lv-Fer |
| e16 | SPU_025090 | Lv-Exosc5 |
| e16 | SPU_025188 | NA |
| e16 | SPU_025296 | none |
| e16 | SPU_025313 | Lv-Acadsb_1 |
| e16 | SPU_025335 | none |
| e16 | SPU_025342 | Lv-AtfL |
| e16 | SPU_025374 | Lv-Sema/Fn3 |
| e16 | SPU_025435 | Lv-Psmd1 |
| e16 | SPU_025450 | Lv-Ssr3_1 |
| e16 | SPU_025555 | Lv-Arf6 |
| e16 | SPU_025569 | Lv-Mrpl17 |
| e16 | SPU_025635 | Lv-Wasl |
| e16 | SPU_025670 | none |
| e16 | SPU_025674 | Lv-Gcn2 |
| e16 | SPU_025677 | Lv-NphsL_5 |
| e16 | SPU_025812 | Lv-Ndusf8 |

| | | |
|---|---|---|
| e16 | SPU_025831 | Lv-Prps1_1 |
| e16 | SPU_025896 | Lv-Dedd |
| e16 | SPU_025952 | Lv-Slc7a11_6 |
| e16 | SPU_025975 | Lv-Dcc/NeogL |
| e16 | SPU_026074 | NA |
| e16 | SPU_026201 | Lv-Fam109a |
| e16 | SPU_026219 | Lv-Centd |
| e16 | SPU_026302 | Lv-Lamb2_1 |
| e16 | SPU_026390 | Lv-Z426 |
| e16 | SPU_026394 | none |
| e16 | SPU_026487 | none |
| e16 | SPU_026497 | Lv-Sympk |
| e16 | SPU_026568 | Lv-Hypp_2853 |
| e16 | SPU_026583 | Lv-Pycr1 |
| e16 | SPU_026612 | Lv-Tert |
| e16 | SPU_026643 | Lv-EmapL2 |
| e16 | SPU_026706 | Lv-Psmg1 |
| e16 | SPU_026740 | Lv-Rqcd1 |
| e16 | SPU_026990 | Lv-Rpa2 |
| e16 | SPU_027008 | Lv-Fkbp15 |
| e16 | SPU_027172 | none |
| e16 | SPU_027329 | Lv-TstL |
| e16 | SPU_027355 | Lv-Rnf14 |
| e16 | SPU_027591 | Lv-Aass_1 |
| e16 | SPU_027693 | Lv-Trappc6B |
| e16 | SPU_027725 | none |
| e16 | SPU_027777 | Lv-Stt3hb |
| e16 | SPU_027827 | Lv-Ssr2 |
| e16 | SPU_027832 | Lv-CholdhL_1 |
| e16 | SPU_027841 | Lv-Ccdc25 |
| e16 | SPU_027900 | NA |
| e16 | SPU_027962 | Lv-Fut8_3 |
| e16 | SPU_027965 | Lv-Rdh13 |
| e16 | SPU_028095 | Lv-C7orf25 |
| e16 | SPU_028191 | none |
| e16 | SPU_028286 | none |
| e16 | SPU_028294 | Lv-Nlr42 |
| e16 | SPU_028346 | none |

| | | |
|---|---|---|
| e16 | SPU_028394 | Lv-Gstcd |
| e16 | SPU_028430 | Lv-Gtf2h3 |
| e16 | SPU_028464 | Lv-C4orf41_2 |
| e16 | SPU_028555 | Lv-Coq7 |
| e16 | SPU_028556 | Lv-Hypp_1233 |
| e16 | SPU_028560 | Lv-GabrA_1 |
| e16 | SPU_028562 | Lv-Pisd |
| e16 | SPU_028587 | Lv-Psph |
| e16 | SPU_028588 | NA |
| e16 | SPU_028613 | Lv-Fcolf_2 |
| e16 | SPU_028652 | none |
| e16 | SPU_028850 | NA |
| e16 | SPU_028904 | Lv-Serpa |
| e16 | SPU_028912 | Lv-Psmd3 |
| e17 | SPU_000008 | Lv-Plcxd3L |
| e17 | SPU_000022 | Lv-Atp5f1 |
| e17 | SPU_000040 | Lv-C7orf60L |
| e17 | SPU_000061 | Lv-Btub2 |
| e17 | SPU_000062 | Lv-Btub3 |
| e17 | SPU_000202 | Lv-Chn2 |
| e17 | SPU_000255 | Lv-Nurr1 |
| e17 | SPU_000281 | Lv-Cat |
| e17 | SPU_000361 | Lv-Tcf19L |
| e17 | SPU_000455 | Lv-Hibch |
| e17 | SPU_000509 | none |
| e17 | SPU_000603 | Lv-Z135 |
| e17 | SPU_000660 | Lv-Faah2 |
| e17 | SPU_000735 | NA |
| e17 | SPU_000796 | NA |
| e17 | SPU_000886 | Lv-Mtch2l |
| e17 | SPU_000919 | Lv-Hypp_317 |
| e17 | SPU_000937 | Lv-Perlc_3 |
| e17 | SPU_000974 | Lv-Crn1Aph |
| e17 | SPU_000997 | Lv-C3-2 |
| e17 | SPU_001000 | Lv-KatsA |
| e17 | SPU_001131 | none |
| e17 | SPU_001141 | NA |
| e17 | SPU_001183 | Lv-Atp5j |

| | | |
|---|---|---|
| e17 | SPU_001413 | Lv-Hypp_1326 |
| e17 | SPU_001441 | Lv-Phol |
| e17 | SPU_001515 | Lv-Chmp5 |
| e17 | SPU_001581 | Lv-Hypp_1337 |
| e17 | SPU_001608 | Lv-Nlr181 |
| e17 | SPU_001817 | Lv-Pkm2 |
| e17 | SPU_001886 | Lv-Wsb1 |
| e17 | SPU_001925 | Lv-Ndufa6 |
| e17 | SPU_001986 | Lv-Maoa_1 |
| e17 | SPU_001989 | Lv-Hypp_1362 |
| e17 | SPU_002030 | Lv-Slc25a10 |
| e17 | SPU_002074 | Lv-Enosf1L |
| e17 | SPU_002110 | Lv-Dnahc8 |
| e17 | SPU_002128 | Lv-Ndufa12 |
| e17 | SPU_002184 | none |
| e17 | SPU_002188 | Lv-Hypp_1377 |
| e17 | SPU_002250 | NA |
| e17 | SPU_002293 | Lv-Acadl |
| e17 | SPU_002336 | Lv-Hypp_353 |
| e17 | SPU_002354 | Lv-EhH2b_12 |
| e17 | SPU_002356 | Lv-Atp6v1g3 |
| e17 | SPU_002373 | Lv-Kiaa1219L |
| e17 | SPU_002385 | Lv-Ugcg |
| e17 | SPU_002671 | Lv-Hypp_364 |
| e17 | SPU_002673 | Lv-Graf |
| e17 | SPU_002758 | Lv-Ndbcp |
| e17 | SPU_002807 | Lv-Idh3g |
| e17 | SPU_002909 | Lv-Gclm |
| e17 | SPU_002937 | Lv-Psma7 |
| e17 | SPU_003047 | none |
| e17 | SPU_003057 | Lv-Zc3p26_3 |
| e17 | SPU_003102 | Lv-Jun |
| e17 | SPU_003104 | Lv-Atp6ap2 |
| e17 | SPU_003112 | Lv-Unc44_269 |
| e17 | SPU_003260 | Lv-Abp620h |
| e17 | SPU_003310 | Lv-Mpp6 |
| e17 | SPU_003408 | Lv-Ndufa13_1 |
| e17 | SPU_003464 | Lv-Uroc1 |

| | | |
|---|---|---|
| e17 | SPU_003495 | Lv-Sarmr10 |
| e17 | SPU_003752 | Lv-Lrp4L_2 |
| e17 | SPU_003755 | Lv-Hypp_394 |
| e17 | SPU_004214 | Lv-Xyli |
| e17 | SPU_004303 | Lv-Hypp_410 |
| e17 | SPU_004356 | Lv-Ankrd50 |
| e17 | SPU_004358 | Lv-Immt |
| e17 | SPU_004472 | Lv-Hypp_1527 |
| e17 | SPU_004485 | Lv-Prep |
| e17 | SPU_004506 | Lv-Slc7a11_2 |
| e17 | SPU_004515 | Lv-Gal3st1-3 |
| e17 | SPU_004532 | Lv-Fat1L |
| e17 | SPU_004533 | Lv-Dusp23(partial) |
| e17 | SPU_004612 | Lv-C22orf28 |
| e17 | SPU_004695 | Lv-MgL_2 |
| e17 | SPU_004732 | Lv-Rgma |
| e17 | SPU_004774 | Lv-Hypp_1543 |
| e17 | SPU_004797 | Lv-Pank2 |
| e17 | SPU_004823 | Lv-Ndufb8 |
| e17 | SPU_004855 | Lv-Acat |
| e17 | SPU_004872 | Lv-Nlr19 |
| e17 | SPU_004901 | Lv-Lsm7 |
| e17 | SPU_005050 | Lv-PolppL_42 |
| e17 | SPU_005216 | Lv-Ttc5 |
| e17 | SPU_005269 | Lv-Slc25a12 |
| e17 | SPU_005289 | NA |
| e17 | SPU_005308 | Lv-C2orf63L |
| e17 | SPU_005458 | Lv-Bpi |
| e17 | SPU_005527 | Lv-Dhrs7 |
| e17 | SPU_005644 | Lv-Hypp_3073 |
| e17 | SPU_005742 | none |
| e17 | SPU_005993 | Lv-Nlr92 |
| e17 | SPU_006172 | Lv-Cfln |
| e17 | SPU_006211 | Lv-Pxr4 |
| e17 | SPU_006271 | Lv-Gabarap |
| e17 | SPU_006395 | none |
| e17 | SPU_006434 | none |
| e17 | SPU_006443 | Lv-Acp1 |

| | | |
|---|---|---|
| e17 | SPU_006513 | Lv-Dusp10 |
| e17 | SPU_006662 | Lv-Hypp_18 |
| e17 | SPU_006693 | NA |
| e17 | SPU_006954 | none |
| e17 | SPU_006986 | Lv-PolppL_56 |
| e17 | SPU_006993 | none |
| e17 | SPU_007044 | Lv-FMO6L |
| e17 | SPU_007085 | Lv-Mvp |
| e17 | SPU_007092 | Lv-Dic1 |
| e17 | SPU_007107 | none |
| e17 | SPU_007114 | Lv-Hmgcs1 |
| e17 | SPU_007161 | Lv-Hsd17b4 |
| e17 | SPU_007231 | Lv-Ttrspn_3 |
| e17 | SPU_007282 | Lv-PtpN23 |
| e17 | SPU_007283 | Lv-Aldh2 |
| e17 | SPU_007286 | Lv-Hypp_511 |
| e17 | SPU_007405 | Lv-Tom1l |
| e17 | SPU_007407 | Lv-Gcnt2_1 |
| e17 | SPU_007422 | Lv-Gpr54_1 |
| e17 | SPU_007435 | Lv-Atub1 |
| e17 | SPU_007436 | none |
| e17 | SPU_007562 | Lv-Hypp_521 |
| e17 | SPU_007676 | Lv-Oxidrd |
| e17 | SPU_007683 | Lv-Acsl5 |
| e17 | SPU_007853 | NA |
| e17 | SPU_007969 | Lv-Spag1L |
| e17 | SPU_008140 | NA |
| e17 | SPU_008199 | none |
| e17 | SPU_008671 | Lv-Hypp_1801 |
| e17 | SPU_008818 | Lv-DisLB2 |
| e17 | SPU_008828 | none |
| e17 | SPU_008833 | Lv-Nlr112 |
| e17 | SPU_008971 | Lv-Gria1 |
| e17 | SPU_008994 | Lv-Fap133L |
| e17 | SPU_008996 | Lv-Ttrspn_5 |
| e17 | SPU_009006 | Lv-Hypp_3069 |
| e17 | SPU_009080 | Lv-Hypp_568 |
| e17 | SPU_009146 | none |

| | | |
|---|---|---|
| e17 | SPU_009243 | Lv-Lmlm |
| e17 | SPU_009380 | Lv-Endrvt69 |
| e17 | SPU_009488 | Lv-Nlr22 |
| e17 | SPU_009518 | Lv-Gst |
| e17 | SPU_009549 | none |
| e17 | SPU_009588 | Lv-Vha26 |
| e17 | SPU_009643 | Lv-Pdcd5 |
| e17 | SPU_009803 | Lv-Hypp_588 |
| e17 | SPU_009871 | Lv-Slc10a2_6 |
| e17 | SPU_009901 | Lv-NeurmdnBLr |
| e17 | SPU_010000 | Lv-PolppL_86 |
| e17 | SPU_010237 | Lv-Endrvt74 |
| e17 | SPU_010238 | Lv-Tppp3 |
| e17 | SPU_010247 | Lv-Acadm |
| e17 | SPU_010373 | Lv-Crk1L |
| e17 | SPU_010397 | Lv-Hypp_1914 |
| e17 | SPU_010437 | none |
| e17 | SPU_010551 | none |
| e17 | SPU_010631 | Lv-Arp3ph |
| e17 | SPU_010677 | Lv-Arsb_3 |
| e17 | SPU_010762 | Lv-Rab1 |
| e17 | SPU_010765 | none |
| e17 | SPU_010778 | Lv-Phyh |
| e17 | SPU_010785 | Lv-Slc24a2_1 |
| e17 | SPU_010797 | Lv-Rej7 |
| e17 | SPU_010974 | none |
| e17 | SPU_011047 | Lv-Hypp_638 |
| e17 | SPU_011126 | Lv-Nxn_1 |
| e17 | SPU_011138 | Lv-Hypp_1961 |
| e17 | SPU_011174 | Lv-Nfe2 |
| e17 | SPU_011203 | Lv-PolypL_940 |
| e17 | SPU_011332 | Lv-Clca1_3 |
| e17 | SPU_011372 | Lv-Aox |
| e17 | SPU_011374 | Lv-Ndufs3_1 |
| e17 | SPU_011494 | none |
| e17 | SPU_011507 | Lv-Hypp_1984 |
| e17 | SPU_011566 | Lv-Slc5a12 |
| e17 | SPU_011724 | Lv-Tacc |

| | | |
|---|---|---|
| e17 | SPU_011911 | none |
| e17 | SPU_012155 | Lv-Unc44_302 |
| e17 | SPU_012228 | Lv-Arp2/3cs4 |
| e17 | SPU_012262 | Lv-Ttc18 |
| e17 | SPU_012361 | Lv-Kctd3_1 |
| e17 | SPU_012368 | Lv-Ndufb9 |
| e17 | SPU_012386 | Lv-Pacsin2 |
| e17 | SPU_012498 | Lv-Slc39a12 |
| e17 | SPU_012542 | none |
| e17 | SPU_012630 | Lv-Lbr |
| e17 | SPU_012679 | Lv-Atub13 |
| e17 | SPU_012687 | Lv-Gal3st3L_1 |
| e17 | SPU_012732 | Lv-Ndufb3 |
| e17 | SPU_012761 | none |
| e17 | SPU_012762 | Lv-Hypp_7048 |
| e17 | SPU_012983 | Lv-Mcm3_1 |
| e17 | SPU_013006 | Lv-Hydin |
| e17 | SPU_013037 | Lv-Ndufa8 |
| e17 | SPU_013079 | none |
| e17 | SPU_013082 | none |
| e17 | SPU_013087 | Lv-Sucla2 |
| e17 | SPU_013092 | Lv-TspO |
| e17 | SPU_013094 | Lv-Wipf1L |
| e17 | SPU_013143 | Lv-Hypp_2072 |
| e17 | SPU_013225 | Lv-Uqcrh |
| e17 | SPU_013372 | NA |
| e17 | SPU_013383 | none |
| e17 | SPU_013608 | Lv-Dj1p |
| e17 | SPU_013644 | Lv-Ndufs1_1 |
| e17 | SPU_013738 | none |
| e17 | SPU_013769 | Lv-Arsa |
| e17 | SPU_013948 | none |
| e17 | SPU_014171 | Lv-Hypp_740 |
| e17 | SPU_014295 | Lv-Patched |
| e17 | SPU_014302 | Lv-Chst3L_3 |
| e17 | SPU_014336 | Lv-Stambp |
| e17 | SPU_014456 | Lv-Dgcr6L_2 |
| e17 | SPU_014479 | none |

| | | |
|---|---|---|
| e17 | SPU_014681 | Lv-Hypp_761 |
| e17 | SPU_014773 | none |
| e17 | SPU_014862 | none |
| e17 | SPU_014886 | Lv-Ypel5 |
| e17 | SPU_014973 | Lv-Endrvt88 |
| e17 | SPU_014975 | Lv-Xdh_3 |
| e17 | SPU_014978 | Lv-Vegf |
| e17 | SPU_015149 | none |
| e17 | SPU_015209 | Lv-Hypp_109 |
| e17 | SPU_015238 | Lv-Cox15 |
| e17 | SPU_015310 | Lv-Accn1_1 |
| e17 | SPU_015361 | Lv-Unk_86 |
| e17 | SPU_015368 | Lv-Slsp |
| e17 | SPU_015408 | Lv-Hypp_3054 |
| e17 | SPU_015619 | Lv-Unc44L_111 |
| e17 | SPU_015959 | Lv-Ndufs3 |
| e17 | SPU_015977 | Lv-Cnnm2_1 |
| e17 | SPU_016061 | NA |
| e17 | SPU_016070 | Lv-Btbd2 |
| e17 | SPU_016105 | none |
| e17 | SPU_016186 | Lv-MaoaL |
| e17 | SPU_016217 | none |
| e17 | SPU_016390 | Lv-Shfm1 |
| e17 | SPU_016523 | Lv-Rar |
| e17 | SPU_016680 | Lv-C16orf62L |
| e17 | SPU_016706 | Lv-Prkab2 |
| e17 | SPU_016784 | none |
| e17 | SPU_016806 | none |
| e17 | SPU_016817 | none |
| e17 | SPU_016832 | Lv-Fut6_1 |
| e17 | SPU_016849 | Lv-Snx6 |
| e17 | SPU_016882 | Lv-Atp6v1a2 |
| e17 | SPU_016993 | Lv-Atp6v0b |
| e17 | SPU_017108 | NA |
| e17 | SPU_017119 | none |
| e17 | SPU_017138 | Lv-ABCC9g |
| e17 | SPU_017157 | Lv-Ghitm |
| e17 | SPU_017409 | Lv-Slc16a12L_14 |

| | | |
|---|---|---|
| e17 | SPU_017553 | Lv-Abca1_1 |
| e17 | SPU_017669 | Lv-Them2 |
| e17 | SPU_017714 | Lv-CTI |
| e17 | SPU_017870 | Lv-Kiaa0100L |
| e17 | SPU_017881 | none |
| e17 | SPU_017941 | Lv-Hypp_2362 |
| e17 | SPU_018061 | none |
| e17 | SPU_018079 | Lv-Drps27a |
| e17 | SPU_018267 | none |
| e17 | SPU_018479 | Lv-Cdc42l |
| e17 | SPU_018624 | Lv-Mdh2 |
| e17 | SPU_018717 | Lv-Arp2 |
| e17 | SPU_018719 | NA |
| e17 | SPU_018794 | none |
| e17 | SPU_018860 | Lv-Wdr7 |
| e17 | SPU_018892 | Lv-Ndufa11 |
| e17 | SPU_019068 | Lv-Hhat-2 |
| e17 | SPU_019081 | Lv-Cct6A |
| e17 | SPU_019087 | Lv-Rab7 |
| e17 | SPU_019118 | Lv-Cox6a1 |
| e17 | SPU_019130 | Lv-Hypp_147 |
| e17 | SPU_019206 | Lv-Ndufs6 |
| e17 | SPU_019214 | Lv-Tmem66 |
| e17 | SPU_019275 | NA |
| e17 | SPU_019330 | Lv-Allc |
| e17 | SPU_019424 | Lv-Nedd8_2 |
| e17 | SPU_019466 | NA |
| e17 | SPU_019645 | Lv-Bckdha |
| e17 | SPU_019715 | Lv-Erk1/2 |
| e17 | SPU_019806 | NA |
| e17 | SPU_019831 | Lv-Hypp_949 |
| e17 | SPU_019939 | none |
| e17 | SPU_019952 | none |
| e17 | SPU_019989 | none |
| e17 | SPU_020006 | Lv-Gk |
| e17 | SPU_020123 | Lv-ElfB |
| e17 | SPU_020124 | Lv-ElfA |
| e17 | SPU_020140 | Lv-Pdcd6ip |

| | | |
|---|---|---|
| e17 | SPU_020197 | Lv-Profilin |
| e17 | SPU_020328 | Lv-Hypp_971 |
| e17 | SPU_020389 | Lv-Dnah17 |
| e17 | SPU_020625 | Lv-C6orf224L |
| e17 | SPU_020642 | Lv-Mut |
| e17 | SPU_020716 | Lv-Hypp_986 |
| e17 | SPU_020749 | Lv-Aldh16aL |
| e17 | SPU_020765 | Lv-Ighmbp2-2 |
| e17 | SPU_020856 | Lv-Tbc1d24 |
| e17 | SPU_020860 | Lv-Slco4a1_8 |
| e17 | SPU_020888 | Lv-Ankrd27L |
| e17 | SPU_020905 | Lv-Hypp_1004 |
| e17 | SPU_021007 | Lv-CapZB |
| e17 | SPU_021103 | Lv-Abcc3a |
| e17 | SPU_021229 | Lv-Cf5/8L2 |
| e17 | SPU_021307 | Lv-Prkag2 |
| e17 | SPU_021332 | Lv-Trex2L_1 |
| e17 | SPU_021343 | none |
| e17 | SPU_021518 | Lv-C5orf41L |
| e17 | SPU_021625 | Lv-Clns1a |
| e17 | SPU_021677 | none |
| e17 | SPU_021701 | Lv-Uba1 |
| e17 | SPU_021825 | none |
| e17 | SPU_021926 | Lv-Wdr23_1 |
| e17 | SPU_021927 | Lv-Runx1t1 |
| e17 | SPU_021973 | Lv-Hypp_178 |
| e17 | SPU_021995 | none |
| e17 | SPU_022031 | Lv-Pnkd |
| e17 | SPU_022187 | Lv-Atub7 |
| e17 | SPU_022355 | Lv-Hypp_9809 |
| e17 | SPU_022397 | Lv-Crym |
| e17 | SPU_022529 | Lv-Tpx |
| e17 | SPU_022591 | Lv-Hypp_2618 |
| e17 | SPU_022761 | Lv-Ndufa9 |
| e17 | SPU_022797 | none |
| e17 | SPU_022807 | Lv-Pdha1 |
| e17 | SPU_022837 | Lv-Rgs9 |
| e17 | SPU_022857 | Lv-Hrh1 |

| | | |
|---|---|---|
| e17 | SPU_022865 | none |
| e17 | SPU_022870 | Lv-Selb_1 |
| e17 | SPU_022944 | NA |
| e17 | SPU_023048 | Lv-Supt16hL_1 |
| e17 | SPU_023103 | Lv-Hypp_1071 |
| e17 | SPU_023109 | none |
| e17 | SPU_023158 | Lv-Tp53i3 |
| e17 | SPU_023159 | Lv-Ttc18_1 |
| e17 | SPU_023265 | Lv-Gcdh |
| e17 | SPU_023284 | Lv-Hypp_2663 |
| e17 | SPU_023328 | Lv-Tmem143 |
| e17 | SPU_023367 | none |
| e17 | SPU_023369 | Lv-Nek4 |
| e17 | SPU_023610 | Lv-Baiap3 |
| e17 | SPU_023733 | Lv-Scaper |
| e17 | SPU_023776 | Lv-Apaf |
| e17 | SPU_023782 | Lv-Sept6 |
| e17 | SPU_023958 | Lv-Tmem63b |
| e17 | SPU_024020 | Lv-Fbgp |
| e17 | SPU_024084 | Lv-Srcr202 |
| e17 | SPU_024115 | Lv-Ndufa13 |
| e17 | SPU_024168 | Lv-Psmb7 |
| e17 | SPU_024202 | Lv-Hypp2715 |
| e17 | SPU_024221 | Lv-Snx2 |
| e17 | SPU_024261 | Lv-Fmo5_4 |
| e17 | SPU_024267 | Lv-Maob |
| e17 | SPU_024278 | Lv-Ust_3 |
| e17 | SPU_024294 | NA |
| e17 | SPU_024366 | Lv-FerHc |
| e17 | SPU_024373 | NA |
| e17 | SPU_024420 | none |
| e17 | SPU_024515 | Lv-Mcm |
| e17 | SPU_024526 | Lv-Rhoapk1 |
| e17 | SPU_024723 | Lv-Hypp_1124 |
| e17 | SPU_024752 | Lv-Mtmr3/4 |
| e17 | SPU_024800 | none |
| e17 | SPU_024873 | Lv-Gls |
| e17 | SPU_025005 | Lv-Ddx1_1 |

| | | |
|---|---|---|
| e17 | SPU_025025 | none |
| e17 | SPU_025100 | Lv-Cth |
| e17 | SPU_025110 | Lv-Xab2 |
| e17 | SPU_025307 | Lv-Adk_1 |
| e17 | SPU_025427 | Lv-Pcm1 |
| e17 | SPU_025507 | Lv-Udx2 |
| e17 | SPU_025614 | Lv-Fum1 |
| e17 | SPU_025732 | Lv-Rad50 |
| e17 | SPU_025771 | Lv-Rhogdi |
| e17 | SPU_025863 | Lv-Cyp2L59 |
| e17 | SPU_025900 | Lv-Hypp_236 |
| e17 | SPU_025921 | Lv-Mboat2 |
| e17 | SPU_025961 | Lv-Uchl |
| e17 | SPU_026000 | Lv-Hypp_1164 |
| e17 | SPU_026111 | Lv-Hypp_240 |
| e17 | SPU_026131 | Lv-Ugt2b17L |
| e17 | SPU_026173 | Lv-Hypp_3009 |
| e17 | SPU_026206 | Lv-Ldhd |
| e17 | SPU_026217 | Lv-PolydL |
| e17 | SPU_026291 | Lv-Gclc |
| e17 | SPU_026359 | none |
| e17 | SPU_026386 | Lv-Adh3 |
| e17 | SPU_026418 | Lv-Klf11 |
| e17 | SPU_026490 | Lv-Sepw1 |
| e17 | SPU_026491 | none |
| e17 | SPU_026580 | Lv-Atp16 |
| e17 | SPU_026624 | Lv-Scu |
| e17 | SPU_026827 | none |
| e17 | SPU_026949 | Lv-Tf |
| e17 | SPU_026982 | Lv-Serinc1 |
| e17 | SPU_027069 | Lv-Qpc |
| e17 | SPU_027165 | NA |
| e17 | SPU_027343 | Lv-Itih_6 |
| e17 | SPU_027428 | none |
| e17 | SPU_027627 | Lv-Txndc5 |
| e17 | SPU_027655 | Lv-Tufm |
| e17 | SPU_027660 | none |
| e17 | SPU_027849 | NA |

| | | |
|---|---|---|
| e17 | SPU_027974 | Lv-Arpc1a |
| e17 | SPU_028100 | Lv-Fn3/Igf_25 |
| e17 | SPU_028173 | Lv-Adam10 |
| e17 | SPU_028319 | none |
| e17 | SPU_028357 | Lv-Wdr1 |
| e17 | SPU_028445 | Lv-Polsrs |
| e17 | SPU_028483 | Lv-Nlr80 |
| e17 | SPU_028554 | Lv-Pcyt2_1 |
| e17 | SPU_028728 | Lv-Dld |
| e17 | SPU_028733 | Lv-C20orf103L |
| e17 | SPU_028781 | Lv-Vinph |
| e17 | SPU_028814 | none |
| e17 | SPU_028874 | Lv-Pdpi(cyclophilin)L3 |
| e17 | SPU_028880 | none |
| e17 | SPU_028917 | Lv-Gpd1 |
| e17 | SPU_030072 | Lv-TnfsfL4 |
| e17 | SPU_030234 | Lv-Axndhc15h |
| e18 | SPU_000268 | Lv-Mon2l |
| e18 | SPU_000278 | Lv-Tubgcp2 |
| e18 | SPU_000437 | Lv-Z211 |
| e18 | SPU_000503 | Lv-Rab11 |
| e18 | SPU_000555 | Lv-C20orf7L |
| e18 | SPU_000597 | Lv-Hsdl |
| e18 | SPU_000676 | Lv-Lipf_2 |
| e18 | SPU_000779 | Lv-Gba2L |
| e18 | SPU_000805 | NA |
| e18 | SPU_000812 | Lv-Cct2 |
| e18 | SPU_001161 | Lv-Ctnnd2 |
| e18 | SPU_001188 | Lv-Gluld1_12 |
| e18 | SPU_001195 | Lv-Ttrspn_1 |
| e18 | SPU_001211 | Lv-Slc16a9 |
| e18 | SPU_001225 | Lv-Ggdef |
| e18 | SPU_001258 | Lv-Clpp |
| e18 | SPU_001264 | Lv-Pyg |
| e18 | SPU_001324 | Lv-Samd9L_2 |
| e18 | SPU_001404 | Lv-Nmt1 |
| e18 | SPU_001480 | none |
| e18 | SPU_001564 | Lv-Slc6a5 |

| | | |
|---|---|---|
| e18 | SPU_001634 | Lv-MyIV |
| e18 | SPU_001774 | Lv-Nacha2_1 |
| e18 | SPU_002013 | Lv-Fre1L |
| e18 | SPU_002015 | Lv-Dennd3 |
| e18 | SPU_002264 | Lv-PolppL_13 |
| e18 | SPU_002295 | Lv-Sephs2 |
| e18 | SPU_002556 | none |
| e18 | SPU_002560 | Lv-Pla2L |
| e18 | SPU_002654 | Lv-Hypp_362 |
| e18 | SPU_002660 | Lv-Cyp46L2 |
| e18 | SPU_002717 | NA |
| e18 | SPU_002735 | Lv-Hypp_3106 |
| e18 | SPU_002764 | none |
| e18 | SPU_002812 | Lv-StablnL |
| e18 | SPU_002887 | Lv-B4galt5 |
| e18 | SPU_002998 | Lv-Hprt1 |
| e18 | SPU_003067 | Lv-Fen1 |
| e18 | SPU_003110 | Lv-Eif5A |
| e18 | SPU_003274 | none |
| e18 | SPU_003281 | NA |
| e18 | SPU_003332 | Lv-Acy1l2L_3 |
| e18 | SPU_003418 | Lv-Fn3/Egff_15 |
| e18 | SPU_003522 | Lv-Hypp_387 |
| e18 | SPU_003536 | none |
| e18 | SPU_003559 | Lv-Acot9 |
| e18 | SPU_003689 | none |
| e18 | SPU_003765 | Lv-Ogfod2 |
| e18 | SPU_003769 | Lv-Tmprss6L |
| e18 | SPU_003832 | Lv-Glyt |
| e18 | SPU_003887 | Lv-Galnt13 |
| e18 | SPU_003922 | Lv-Soat1 |
| e18 | SPU_004372 | Lv-Svep1L_5 |
| e18 | SPU_004379 | Lv-Flj33220 |
| e18 | SPU_004480 | Lv-Ugtla9 |
| e18 | SPU_004512 | Lv-Unc44L_36 |
| e18 | SPU_004643 | Lv-Dlg2_1 |
| e18 | SPU_004682 | Lv-Ap3b2-2 |
| e18 | SPU_004694 | Lv-Adcy2 |

| | | |
|---|---|---|
| e18 | SPU_004700 | Lv-Cct7 |
| e18 | SPU_004964 | Lv-Pkg2 |
| e18 | SPU_004969 | Lv-Tnrc4 |
| e18 | SPU_004974 | Lv-Chchd2_1 |
| e18 | SPU_005068 | none |
| e18 | SPU_005112 | NA |
| e18 | SPU_005208 | NA |
| e18 | SPU_005251 | Lv-C10orf33L |
| e18 | SPU_005282 | Lv-C14orf166b |
| e18 | SPU_005337 | Lv-Rdh11 |
| e18 | SPU_005409 | NA |
| e18 | SPU_005480 | Lv-Camk2a_1 |
| e18 | SPU_005493 | Lv-Mtvr2l |
| e18 | SPU_005660 | Lv-Hypp_1593 |
| e18 | SPU_005942 | Lv-Taut_2 |
| e18 | SPU_005977 | Lv-Phca |
| e18 | SPU_006017 | Lv-Atp6v1b2 |
| e18 | SPU_006022 | Lv-Igf_15 |
| e18 | SPU_006054 | Lv-Frap_1 |
| e18 | SPU_006168 | Lv-Gal3st1_1 |
| e18 | SPU_006205 | Lv-EhH4_16 |
| e18 | SPU_006225 | Lv-Notchh2 |
| e18 | SPU_006231 | NA |
| e18 | SPU_006283 | Lv-Hypp_14 |
| e18 | SPU_006407 | Lv-Hypp_1659 |
| e18 | SPU_006664 | Lv-Ugt1L |
| e18 | SPU_006742 | Lv-Tectonic |
| e18 | SPU_006794 | Lv-Xpapd |
| e18 | SPU_006803 | Lv-Creb3l3 |
| e18 | SPU_006909 | Lv-Gcsa |
| e18 | SPU_006931 | Lv-Vps18h |
| e18 | SPU_006944 | Lv-Ndufb5 |
| e18 | SPU_007050 | Lv-Atg2hb |
| e18 | SPU_007201 | Lv-Unc44_282 |
| e18 | SPU_007202 | none |
| e18 | SPU_007266 | Lv-Hypp_1709 |
| e18 | SPU_007397 | Lv-Mem_trans |
| e18 | SPU_007456 | Lv-Nek9 |

| | | |
|---|---|---|
| e18 | SPU_007462 | Lv-Hypp_3070 |
| e18 | SPU_007546 | Lv-Hypp_1727 |
| e18 | SPU_008033 | Lv-Chst9_1 |
| e18 | SPU_008075 | Lv-Fdft1 |
| e18 | SPU_008302 | Lv-Sarmr13 |
| e18 | SPU_008320 | NA |
| e18 | SPU_008622 | Lv-Igf_2 |
| e18 | SPU_008687 | Lv-Hypp_35 |
| e18 | SPU_008917 | Lv-Pex11 |
| e18 | SPU_008918 | Lv-Megf10L |
| e18 | SPU_008929 | Lv-Hypp_1814 |
| e18 | SPU_008953 | Lv-Talpid3 |
| e18 | SPU_009340 | Lv-MyIIIph |
| e18 | SPU_009456 | Lv-Htatsf1L_1 |
| e18 | SPU_009494 | Lv-Slc16a6_2 |
| e18 | SPU_009534 | Lv-Txnrd3 |
| e18 | SPU_009705 | Lv-Hypp_586 |
| e18 | SPU_009708 | Lv-Trim2 |
| e18 | SPU_009961 | Lv-Oxct1_1 |
| e18 | SPU_010193 | none |
| e18 | SPU_010208 | Lv-Agxt2l1 |
| e18 | SPU_010382 | Lv-Ascc1 |
| e18 | SPU_010441 | Lv-Csmd2L |
| e18 | SPU_010608 | none |
| e18 | SPU_010630 | Lv-Pa2g4L |
| e18 | SPU_010664 | Lv-Zizimin1 |
| e18 | SPU_010776 | Lv-Unc5 |
| e18 | SPU_010791 | Lv-Ces2 |
| e18 | SPU_010913 | none |
| e18 | SPU_011029 | Lv-RhoA |
| e18 | SPU_011031 | NA |
| e18 | SPU_011091 | Lv-Gtf2F2 |
| e18 | SPU_011173 | Lv-Fyco1 |
| e18 | SPU_011283 | Lv-Pitpna_1 |
| e18 | SPU_011313 | Lv-Nudfc2 |
| e18 | SPU_011444 | Lv-Lrba |
| e18 | SPU_011479 | Lv-Pde9a |
| e18 | SPU_011509 | Lv-Tk12 |

| | | |
|---|---|---|
| e18 | SPU_011722 | Lv-RhoBTB |
| e18 | SPU_011729 | Lv-Nphp1 |
| e18 | SPU_011863 | Lv-RtL_30 |
| e18 | SPU_012024 | Lv-PitpnaL |
| e18 | SPU_012066 | none |
| e18 | SPU_012147 | Lv-Hypp_670 |
| e18 | SPU_012372 | Lv-Lrp4 |
| e18 | SPU_012431 | Lv-Mcm5 |
| e18 | SPU_012611 | Lv-NeurlsnL |
| e18 | SPU_012695 | Lv-Atp6ap1 |
| e18 | SPU_012765 | Lv-Folr1 |
| e18 | SPU_012819 | Lv-Lrr/Igr_4 |
| e18 | SPU_012875 | Lv-Acsl1 |
| e18 | SPU_012971 | Lv-Hypp_2062 |
| e18 | SPU_013074 | Lv-PolypL_945 |
| e18 | SPU_013200 | Lv-Dynlc1 |
| e18 | SPU_013293 | Lv-Svep1L_11 |
| e18 | SPU_013332 | none |
| e18 | SPU_013333 | Lv-Hypp_90 |
| e18 | SPU_013427 | NA |
| e18 | SPU_013450 | Lv-Tpd52L2 |
| e18 | SPU_013454 | Lv-Psap |
| e18 | SPU_013516 | Lv-Hypp_2087 |
| e18 | SPU_013620 | Lv-Sd |
| e18 | SPU_013697 | none |
| e18 | SPU_013710 | Lv-Hypp_95 |
| e18 | SPU_013866 | Lv-Rxfp1L_4 |
| e18 | SPU_013895 | none |
| e18 | SPU_014189 | Lv-Acot2L |
| e18 | SPU_014372 | Lv-Ogdh |
| e18 | SPU_014397 | Lv-My1d/My1e |
| e18 | SPU_014400 | none |
| e18 | SPU_014490 | none |
| e18 | SPU_014746 | Lv-Akrf_3 |
| e18 | SPU_014789 | Lv-Tsta3 |
| e18 | SPU_014826 | Lv-Surreal/Gpcr111 |
| e18 | SPU_014875 | Lv-Wbscr27 |
| e18 | SPU_014884 | Lv-Plekhg1 |

| | | |
|---|---|---|
| e18 | SPU_014948 | NA |
| e18 | SPU_014986 | Lv-Cbr1_4 |
| e18 | SPU_015108 | Lv-Stat |
| e18 | SPU_015159 | Lv-Hypp_2194 |
| e18 | SPU_015194 | Lv-Trpc3_1 |
| e18 | SPU_015359 | Lv-FibcystL |
| e18 | SPU_015415 | Lv-Rcbtb1 |
| e18 | SPU_015456 | Lv-Nr1h6b |
| e18 | SPU_015587 | Lv-Notum |
| e18 | SPU_015674 | Lv-NprA_1 |
| e18 | SPU_015760 | none |
| e18 | SPU_015769 | Lv-Hypp_2229 |
| e18 | SPU_015815 | NA |
| e18 | SPU_015903 | Lv-LsampL_1 |
| e18 | SPU_015973 | Lv-Ank4 |
| e18 | SPU_016062 | Lv-TbcaL1 |
| e18 | SPU_016077 | Lv-Nisch |
| e18 | SPU_016130 | Lv-Nt5dc3 |
| e18 | SPU_016197 | Lv-Slc16a3L_1 |
| e18 | SPU_016273 | Lv-Sapd |
| e18 | SPU_016334 | Lv-AdcyL |
| e18 | SPU_016514 | Lv-Sept3 |
| e18 | SPU_016631 | Lv-Nap1l1 |
| e18 | SPU_016791 | Lv-Dnl1 |
| e18 | SPU_016798 | none |
| e18 | SPU_016827 | none |
| e18 | SPU_016847 | Lv-Memo1 |
| e18 | SPU_016862 | Lv-Clect/Ftp |
| e18 | SPU_017032 | Lv-Ppa1 |
| e18 | SPU_017068 | NA |
| e18 | SPU_017073 | none |
| e18 | SPU_017233 | Lv-Hypp_3048 |
| e18 | SPU_017244 | Lv-Ufc1 |
| e18 | SPU_017370 | Lv-ThromE |
| e18 | SPU_017533 | Lv-Pde5_1 |
| e18 | SPU_017593 | Lv-Birc1 |
| e18 | SPU_017628 | Lv-Nudt5 |
| e18 | SPU_017877 | Lv-Gephph |

| | | |
|---|---|---|
| e18 | SPU_017912 | Lv-Achra9L |
| e18 | SPU_018011 | none |
| e18 | SPU_018063 | Lv-C20orf194 |
| e18 | SPU_018376 | none |
| e18 | SPU_018492 | Lv-YjeFk |
| e18 | SPU_018705 | Lv-Moesin |
| e18 | SPU_018786 | Lv-Pin4 |
| e18 | SPU_018898 | Lv-Klc |
| e18 | SPU_018905 | Lv-Alkbh8L |
| e18 | SPU_019009 | Lv-Dmd |
| e18 | SPU_019078 | Lv-Rpa3 |
| e18 | SPU_019082 | Lv-Cyp2L38 |
| e18 | SPU_019153 | none |
| e18 | SPU_019157 | Lv-Chmp6 |
| e18 | SPU_019347 | Lv-Hypp_154 |
| e18 | SPU_019372 | none |
| e18 | SPU_019394 | Lv-Cadherin_3 |
| e18 | SPU_019401 | Lv-Crim1 |
| e18 | SPU_019428 | Lv-Dhrsx_4 |
| e18 | SPU_019454 | NA |
| e18 | SPU_019458 | Lv-Rab2 |
| e18 | SPU_019548 | none |
| e18 | SPU_019586 | Lv-Lmo4 |
| e18 | SPU_019696 | Lv-Nlr95 |
| e18 | SPU_019703 | Lv-Bmi1L |
| e18 | SPU_019773 | none |
| e18 | SPU_019853 | Lv-NotchL_13 |
| e18 | SPU_019999 | Lv-Phscd1 |
| e18 | SPU_020151 | none |
| e18 | SPU_020225 | none |
| e18 | SPU_020268 | Lv-Cnp |
| e18 | SPU_020294 | Lv-Hsp20.2 |
| e18 | SPU_020548 | Lv-Hypp_2511 |
| e18 | SPU_020591 | Lv-Hypp_2514 |
| e18 | SPU_020762 | Lv-Slc |
| e18 | SPU_020875 | Lv-Dysf_1 |
| e18 | SPU_020911 | Lv-Galr2L_26 |
| e18 | SPU_021013 | none |

| | | |
|---|---|---|
| e18 | SPU_021217 | Lv-Dysf2 |
| e18 | SPU_021232 | Lv-Hypp_1018 |
| e18 | SPU_021572 | none |
| e18 | SPU_021605 | none |
| e18 | SPU_021658 | Lv-Bhmt_5 |
| e18 | SPU_021680 | NA |
| e18 | SPU_021773 | Lv-Asns_1 |
| e18 | SPU_021854 | Lv-Mocs2 |
| e18 | SPU_021931 | Lv-Aadc |
| e18 | SPU_021970 | Lv-Slc16a12_8 |
| e18 | SPU_022201 | Lv-Crbn |
| e18 | SPU_022207 | Lv-Limk |
| e18 | SPU_022332 | Lv-Tpmtl |
| e18 | SPU_022333 | none |
| e18 | SPU_022368 | Lv-Hypp_181 |
| e18 | SPU_022378 | Lv-Ehhadh |
| e18 | SPU_022399 | Lv-Cbr1_5 |
| e18 | SPU_022413 | Lv-Phlpb_2 |
| e18 | SPU_022431 | Lv-Rhogap24 |
| e18 | SPU_022707 | Lv-Myd88L2 |
| e18 | SPU_022987 | Lv-Slc35e4 |
| e18 | SPU_022991 | Lv-Rab21l |
| e18 | SPU_023104 | NA |
| e18 | SPU_023171 | Lv-Hypp_2653 |
| e18 | SPU_023232 | Lv-Rdh8_2 |
| e18 | SPU_023310 | none |
| e18 | SPU_023336 | Lv-Dlc1 |
| e18 | SPU_023380 | Lv-Sec14L2_4 |
| e18 | SPU_023514 | none |
| e18 | SPU_023568 | none |
| e18 | SPU_023669 | Lv-Galns |
| e18 | SPU_023808 | Lv-Cyp3L7 |
| e18 | SPU_023831 | none |
| e18 | SPU_023839 | Lv-DspchdLA |
| e18 | SPU_023886 | Lv-Sh3glb1 |
| e18 | SPU_024066 | NA |
| e18 | SPU_024112 | Lv-Xdh_2 |
| e18 | SPU_024201 | none |

| | | |
|---|---|---|
| e18 | SPU_024241 | Lv-Cdc37 |
| e18 | SPU_024259 | none |
| e18 | SPU_024473 | Lv-Ampd |
| e18 | SPU_024525 | Lv-Sfk7 |
| e18 | SPU_024540 | Lv-Pigt |
| e18 | SPU_024599 | Lv-Flnc |
| e18 | SPU_024604 | Lv-Rdh11_5 |
| e18 | SPU_024607 | Lv-Vps11 |
| e18 | SPU_024626 | Lv-Heatr7a_2 |
| e18 | SPU_024668 | NA |
| e18 | SPU_024709 | Lv-Nlr173 |
| e18 | SPU_024731 | Lv-Tlr131 |
| e18 | SPU_024741 | Lv-Sele |
| e18 | SPU_024775 | Lv-Nsdhl |
| e18 | SPU_024821 | none |
| e18 | SPU_024984 | Lv-Ndrg1L |
| e18 | SPU_025077 | Lv-Nlr46 |
| e18 | SPU_025196 | Lv-Acnn1 |
| e18 | SPU_025198 | Lv-Mycbpap |
| e18 | SPU_025279 | NA |
| e18 | SPU_025484 | Lv-Cct5 |
| e18 | SPU_025486 | Lv-Mox |
| e18 | SPU_025513 | Lv-Duoxa1 |
| e18 | SPU_025520 | Lv-ApdL1_1 |
| e18 | SPU_025577 | Lv-Secisbp2l_1 |
| e18 | SPU_025817 | Lv-Rfxc1l |
| e18 | SPU_025953 | Lv-Pme1 |
| e18 | SPU_026222 | Lv-ReelerL_5 |
| e18 | SPU_026273 | Lv-Slc24a6 |
| e18 | SPU_026344 | Lv-Hypp_1174 |
| e18 | SPU_026460 | none |
| e18 | SPU_026554 | Lv-Hypp_1181 |
| e18 | SPU_026797 | Lv-Vwde |
| e18 | SPU_026833 | Lv-Gaa_2 |
| e18 | SPU_026928 | Lv-Dci |
| e18 | SPU_026944 | Lv-Arf1l1 |
| e18 | SPU_026968 | Lv-Kiaa1598L |
| e18 | SPU_026978 | NA |

| | | |
|---|---|---|
| e18 | SPU_027115 | NA |
| e18 | SPU_027186 | Lv-Endrvt52 |
| e18 | SPU_027318 | none |
| e18 | SPU_027341 | Lv-Itih_4 |
| e18 | SPU_027362 | Lv-Top1L |
| e18 | SPU_027406 | Lv-Oprs1_1 |
| e18 | SPU_027650 | Lv-Gaa |
| e18 | SPU_027796 | NA |
| e18 | SPU_027932 | Lv-Psma2 |
| e18 | SPU_027990 | Lv-Herc4_6 |
| e18 | SPU_028001 | Lv-Mccc2 |
| e18 | SPU_028015 | Lv-Hypp_2927 |
| e18 | SPU_028069 | Lv-Sam8 |
| e18 | SPU_028414 | Lv-Hypp_2954 |
| e18 | SPU_028630 | Lv-Nlr107 |
| e18 | SPU_028881 | NA |
| e18 | SPU_028886 | NA |
| e19 | SPU_000012 | Lv-Agmat |
| e19 | SPU_000047 | Lv-Egfl |
| e19 | SPU_000077 | Lv-Atp5c1 |
| e19 | SPU_000094 | Lv-Hypp_293 |
| e19 | SPU_000186 | Lv-Lrr/Igr_3 |
| e19 | SPU_000275 | Lv-Hao1 |
| e19 | SPU_000299 | Lv-RasaL2 |
| e19 | SPU_000364 | Lv-Atp6v1h |
| e19 | SPU_000435 | Lv-MlckL |
| e19 | SPU_000458 | Lv-Siat7b |
| e19 | SPU_000680 | Lv-Adam/Ts6/10L1 |
| e19 | SPU_000692 | Lv-Ugt2b17 |
| e19 | SPU_000721 | none |
| e19 | SPU_000913 | Lv-Htr4L |
| e19 | SPU_001054 | Lv-Nlr48 |
| e19 | SPU_001138 | Lv-Cara14LA |
| e19 | SPU_001613 | Lv-MyIIelc |
| e19 | SPU_001758 | Lv-Paqr8 |
| e19 | SPU_001807 | none |
| e19 | SPU_001885 | Lv-Myl9 |
| e19 | SPU_002012 | Lv-Atpif1L |

| | | |
|---|---|---|
| e19 | SPU_002185 | none |
| e19 | SPU_002380 | Lv-Cyp4VL2 |
| e19 | SPU_002438 | Lv-Rps21 |
| e19 | SPU_002458 | Lv-Rpl11 |
| e19 | SPU_002531 | Lv-Fn3f_20 |
| e19 | SPU_002635 | Lv-Hypp_360 |
| e19 | SPU_002693 | none |
| e19 | SPU_002855 | none |
| e19 | SPU_002886 | none |
| e19 | SPU_002932 | Lv-Ckmt2 |
| e19 | SPU_003134 | Lv-C2gnt3_1 |
| e19 | SPU_003200 | Lv-Nlr9 |
| e19 | SPU_003232 | Lv-Cyp17r2 |
| e19 | SPU_003696 | Lv-Sept4 |
| e19 | SPU_003718 | Lv-Rps16 |
| e19 | SPU_003849 | Lv-Rpl32 |
| e19 | SPU_004000 | none |
| e19 | SPU_004017 | Lv-Wap/Ig/Ku/C345c |
| e19 | SPU_004105 | none |
| e19 | SPU_004294 | Lv-Pde2aL |
| e19 | SPU_004607 | Lv-Agmat_1 |
| e19 | SPU_004625 | Lv-Bhmt_1 |
| e19 | SPU_004703 | Lv-Acnat2L |
| e19 | SPU_004924 | Lv-Wnt10 |
| e19 | SPU_005032 | Lv-Calm |
| e19 | SPU_005033 | Lv-Calm1 |
| e19 | SPU_005194 | none |
| e19 | SPU_005201 | Lv-Idh2 |
| e19 | SPU_005252 | Lv-Rpl26 |
| e19 | SPU_005296 | Lv-Atp5b |
| e19 | SPU_005325 | Lv-Gnmt_2 |
| e19 | SPU_005418 | Lv-Gdap1L |
| e19 | SPU_005581 | NA |
| e19 | SPU_005691 | Lv-Pdia3 |
| e19 | SPU_005709 | none |
| e19 | SPU_005785 | Lv-AcheL_4 |
| e19 | SPU_005844 | Lv-Hypp_3074 |
| e19 | SPU_005849 | Lv-Gpr112L |

| | | |
|---|---|---|
| e19 | SPU_005904 | Lv-Sr/So/Gpcr |
| e19 | SPU_006214 | Lv-Sod |
| e19 | SPU_006354 | Lv-Dynlt2 |
| e19 | SPU_006374 | Lv-Atp11b |
| e19 | SPU_006383 | Lv-Scp2 |
| e19 | SPU_006437 | NA |
| e19 | SPU_006460 | Lv-Vps52L |
| e19 | SPU_006569 | Lv-Rpl10 |
| e19 | SPU_006583 | Lv-Mad |
| e19 | SPU_006637 | Lv-Z193 |
| e19 | SPU_006701 | none |
| e19 | SPU_006721 | Lv-Lrr1_1 |
| e19 | SPU_006816 | none |
| e19 | SPU_006852 | Lv-Tyms |
| e19 | SPU_006859 | Lv-SNAP25 |
| e19 | SPU_006974 | none |
| e19 | SPU_007003 | Lv-Hypp_504 |
| e19 | SPU_007006 | Lv-Pde2a |
| e19 | SPU_007098 | none |
| e19 | SPU_007175 | Lv-Hypp_21 |
| e19 | SPU_007335 | Lv-Cyp4L2 |
| e19 | SPU_007359 | Lv-Surreal/Gpcr102 |
| e19 | SPU_007424 | Lv-Btub9 |
| e19 | SPU_007601 | Lv-Slco4a1_7 |
| e19 | SPU_007881 | Lv-Tmed6 |
| e19 | SPU_007960 | Lv-Atp5o |
| e19 | SPU_008000 | Lv-Odadc3L2 |
| e19 | SPU_008305 | Lv-Pdpi(cyclophilin)L7 |
| e19 | SPU_008437 | Lv-Unc44L_60 |
| e19 | SPU_008464 | Lv-Acaa2 |
| e19 | SPU_008543 | Lv-AkrL_3 |
| e19 | SPU_008606 | Lv-Xpnpep1_1 |
| e19 | SPU_008778 | Lv-Rpl3 |
| e19 | SPU_009300 | Lv-Cct3 |
| e19 | SPU_009302 | Lv-Chst11_9 |
| e19 | SPU_009415 | Lv-Iqgap1 |
| e19 | SPU_009423 | Lv-Cap1 |
| e19 | SPU_009433 | Lv-CplcaIIIb |

| | | |
|---|---|---|
| e19 | SPU_009460 | Lv-Hypp_1851 |
| e19 | SPU_009481 | Lv-CskaI |
| e19 | SPU_009483 | Lv-actin IIb |
| e19 | SPU_009644 | Lv-Np |
| e19 | SPU_009698 | Lv-Abtb1 |
| e19 | SPU_009880 | Lv-Hypp_1877 |
| e19 | SPU_009959 | Lv-Hypp_596 |
| e19 | SPU_010031 | Lv-Nalcn |
| e19 | SPU_010148 | NA |
| e19 | SPU_010156 | Lv-Mark4 |
| e19 | SPU_010273 | Lv-Slc16a14L_1 |
| e19 | SPU_010274 | none |
| e19 | SPU_010528 | Lv-Rpl35 |
| e19 | SPU_010727 | NA |
| e19 | SPU_010750 | Lv-Protease |
| e19 | SPU_010756 | Lv-Chst11_10 |
| e19 | SPU_010822 | NA |
| e19 | SPU_010829 | Lv-Ef2 |
| e19 | SPU_010887 | Lv-Hypp_631 |
| e19 | SPU_010936 | Lv-Rps2 |
| e19 | SPU_010943 | Lv-Fascin1 |
| e19 | SPU_011187 | Lv-Smpd1 |
| e19 | SPU_011237 | Lv-Bche-4 |
| e19 | SPU_011333 | Lv-Cyfip2_2 |
| e19 | SPU_011649 | Lv-Peci |
| e19 | SPU_011693 | Lv-Hemic1 |
| e19 | SPU_011713 | Lv-Sardh |
| e19 | SPU_011823 | Lv-Tlr066 |
| e19 | SPU_011917 | Lv-Tagln2_1 |
| e19 | SPU_012282 | Lv-Actph6 |
| e19 | SPU_012383 | Lv-Kr/Fa58C/Clect/Egf/Gpcr |
| e19 | SPU_012414 | Lv-Mcf2l |
| e19 | SPU_012486 | Lv-Hypp_2035 |
| e19 | SPU_012561 | Lv-OstaL |
| e19 | SPU_012610 | Lv-Lrr/Gpcr_4 |
| e19 | SPU_012636 | Lv-Dbi |
| e19 | SPU_012677 | Lv-Mdh1 |
| e19 | SPU_012839 | Lv-Gpi_1 |

| | | |
|---|---|---|
| e19 | SPU_012956 | none |
| e19 | SPU_013017 | none |
| e19 | SPU_013041 | Lv-Rdh11_1 |
| e19 | SPU_013052 | Lv-Atp5g1 |
| e19 | SPU_013064 | none |
| e19 | SPU_013106 | Lv-ThrombD |
| e19 | SPU_013125 | Lv-Slc5a12_1 |
| e19 | SPU_013135 | Lv-Atp5l |
| e19 | SPU_013183 | Lv-Trop1 |
| e19 | SPU_013235 | Lv-Hypp_709 |
| e19 | SPU_013273 | Lv-Btub4 |
| e19 | SPU_013322 | none |
| e19 | SPU_013353 | Lv-Atp5h |
| e19 | SPU_013386 | NA |
| e19 | SPU_013428 | Lv-Dcxr |
| e19 | SPU_013432 | Lv-Rpl5 |
| e19 | SPU_013452 | Lv-Rpsa |
| e19 | SPU_013599 | Lv-Fam124a |
| e19 | SPU_013772 | Lv-Vhaac39 |
| e19 | SPU_013819 | Lv-Eno1 |
| e19 | SPU_013896 | NA |
| e19 | SPU_013909 | none |
| e19 | SPU_013959 | Lv-B3galt1L_3 |
| e19 | SPU_014128 | Lv-Nlr11 |
| e19 | SPU_014174 | none |
| e19 | SPU_014276 | none |
| e19 | SPU_014342 | Lv-Suox |
| e19 | SPU_014421 | none |
| e19 | SPU_014476 | Lv-Slc6a6 |
| e19 | SPU_014478 | Lv-Cox4i1 |
| e19 | SPU_014501 | Lv-Slc43a3L |
| e19 | SPU_014584 | Lv-Phgdh |
| e19 | SPU_014596 | Lv-Trpc7 |
| e19 | SPU_014637 | Lv-Tagln2 |
| e19 | SPU_014800 | Lv-Rps10 |
| e19 | SPU_014841 | Lv-Radil |
| e19 | SPU_014869 | Lv-Prxdxn |
| e19 | SPU_015150 | Lv-Hypp_772 |

| | | |
|---|---|---|
| e19 | SPU_015274 | Lv-Got2_1 |
| e19 | SPU_015335 | Lv-Vamp1 |
| e19 | SPU_015757 | Lv-QrfprL_2 |
| e19 | SPU_015768 | Lv-Nlr179 |
| e19 | SPU_015772 | Lv-Klf15 |
| e19 | SPU_015997 | Lv-Ndufs5 |
| e19 | SPU_016098 | Lv-Hypp_811 |
| e19 | SPU_016141 | none |
| e19 | SPU_016204 | NA |
| e19 | SPU_016312 | none |
| e19 | SPU_016362 | Lv-Acsf2 |
| e19 | SPU_016363 | Lv-Fn3/Egf/Lnb/Tm7_1 |
| e19 | SPU_016414 | Lv-Vha55 |
| e19 | SPU_016441 | Lv-Tkt |
| e19 | SPU_016442 | Lv-Cyp2L32 |
| e19 | SPU_016443 | Lv-Brn1-2-4 |
| e19 | SPU_016532 | NA |
| e19 | SPU_016611 | Lv-Dnajhc5b |
| e19 | SPU_016640 | Lv-Fut8L_2 |
| e19 | SPU_016657 | Lv-Cnbs |
| e19 | SPU_016705 | Lv-Prdx5 |
| e19 | SPU_016824 | Lv-Fut3 |
| e19 | SPU_016829 | Lv-C20orf152L |
| e19 | SPU_017124 | none |
| e19 | SPU_017132 | Lv-Chrna9 |
| e19 | SPU_017242 | none |
| e19 | SPU_017318 | Lv-Fads |
| e19 | SPU_017527 | Lv-Ppglcp |
| e19 | SPU_017605 | Lv-Atp5a1 |
| e19 | SPU_017664 | none |
| e19 | SPU_017719 | Lv-Pde5_2 |
| e19 | SPU_017751 | Lv-Ebp |
| e19 | SPU_018074 | none |
| e19 | SPU_018194 | Lv-Pus10 |
| e19 | SPU_018195 | Lv-Arfl5 |
| e19 | SPU_018224 | Lv-Sarox_1 |
| e19 | SPU_018287 | Lv-Peps |
| e19 | SPU_018435 | none |

| | | |
|---|---|---|
| e19 | SPU_018486 | Lv-Hs3st1L |
| e19 | SPU_018562 | Lv-Hypp_2391 |
| e19 | SPU_018567 | Lv-Dynlc1-1 |
| e19 | SPU_018848 | Lv-Fjx1l |
| e19 | SPU_018934 | Lv-Nvll |
| e19 | SPU_018970 | Lv-Hypp_2414 |
| e19 | SPU_019338 | Lv-Alx |
| e19 | SPU_019514 | Lv-Scdr9 |
| e19 | SPU_019628 | Lv-Dom3zL_4 |
| e19 | SPU_019685 | Lv-Cox5b |
| e19 | SPU_019898 | Lv-Cyp2L39 |
| e19 | SPU_020292 | Lv-Glrx |
| e19 | SPU_020323 | Lv-Slc16a6L_6 |
| e19 | SPU_020487 | none |
| e19 | SPU_020505 | Lv-Cyhr1 |
| e19 | SPU_020607 | Lv-Gsn |
| e19 | SPU_020627 | Lv-Rpl19 |
| e19 | SPU_020971 | Lv-Hyalin15 |
| e19 | SPU_021016 | NA |
| e19 | SPU_021017 | Lv-Msr1L |
| e19 | SPU_021030 | none |
| e19 | SPU_021037 | Lv-AspatL |
| e19 | SPU_021087 | Lv-Cyp2L47 |
| e19 | SPU_021110 | Lv-Slc5a9_5 |
| e19 | SPU_021176 | Lv-Adrb2L_13 |
| e19 | SPU_021271 | Lv-Ush1gL |
| e19 | SPU_021288 | none |
| e19 | SPU_021322 | Lv-Rps7 |
| e19 | SPU_021377 | Lv-G6pdx |
| e19 | SPU_021550 | none |
| e19 | SPU_021551 | Lv-Hypp_2560 |
| e19 | SPU_021673 | Lv-Tollip |
| e19 | SPU_021682 | Lv-Rdh1_1 |
| e19 | SPU_021719 | Lv-Ugt2b7_1 |
| e19 | SPU_021724 | NA |
| e19 | SPU_021726 | Lv-Admp |
| e19 | SPU_021743 | Lv-Gal3st3L_3 |
| e19 | SPU_021857 | Lv-Clca1-2 |

| | | |
|---|---|---|
| e19 | SPU_021871 | Lv-Rdh1_2 |
| e19 | SPU_021924 | Lv-Rpl34 |
| e19 | SPU_022038 | Lv-Capk |
| e19 | SPU_022186 | Lv-Nt5e_1 |
| e19 | SPU_022258 | none |
| e19 | SPU_022270 | Lv-Tcp1_1 |
| e19 | SPU_022325 | Lv-Lphn3-2 |
| e19 | SPU_022682 | Lv-Rps11 |
| e19 | SPU_022684 | Lv-Drd1_1 |
| e19 | SPU_022750 | Lv-Rlp2 |
| e19 | SPU_022957 | Lv-Rps5 |
| e19 | SPU_023066 | Lv-C11orf54 |
| e19 | SPU_023135 | Lv-Hypp_1074 |
| e19 | SPU_023347 | Lv-RasGEF |
| e19 | SPU_023361 | Lv-Slco4a1_13 |
| e19 | SPU_023362 | Lv-Slco4a1_9 |
| e19 | SPU_023368 | Lv-Asb5 |
| e19 | SPU_023651 | Lv-Trpc_1 |
| e19 | SPU_023667 | none |
| e19 | SPU_023835 | none |
| e19 | SPU_023917 | Lv-Slc46a1L_7 |
| e19 | SPU_024102 | Lv-Cs |
| e19 | SPU_024138 | Lv-Abhd4 |
| e19 | SPU_024149 | Lv-Muc5bL |
| e19 | SPU_024379 | Lv-Pgd |
| e19 | SPU_024408 | Lv-Srcr204 |
| e19 | SPU_024439 | Lv-Sirpb/GL1 |
| e19 | SPU_024446 | Lv-Cox5a |
| e19 | SPU_024497 | Lv-Dynlc2-5a |
| e19 | SPU_024541 | Lv-Rrm1_1 |
| e19 | SPU_024657 | Lv-Sod2 |
| e19 | SPU_024877 | Lv-Osr |
| e19 | SPU_024884 | Lv-Kcnj12_1 |
| e19 | SPU_024909 | none |
| e19 | SPU_024910 | Lv-Slsp_23 |
| e19 | SPU_024976 | Lv-Slc5a9L |
| e19 | SPU_024998 | Lv-C9orf56L |
| e19 | SPU_025222 | none |

| | | |
|---|---|---|
| e19 | SPU_025249 | Lv-Krn2 |
| e19 | SPU_025331 | Lv-EchnL |
| e19 | SPU_025364 | Lv-Hypp_1145 |
| e19 | SPU_025397 | Lv-Suclg1 |
| e19 | SPU_025414 | Lv-Gal/lec15 |
| e19 | SPU_025436 | Lv-5ht2 |
| e19 | SPU_025449 | none |
| e19 | SPU_025574 | Lv-Rps14 |
| e19 | SPU_025630 | Lv-Adcy10 |
| e19 | SPU_025681 | none |
| e19 | SPU_025886 | NA |
| e19 | SPU_025954 | Lv-Aif |
| e19 | SPU_026026 | none |
| e19 | SPU_026072 | Lv-EceL |
| e19 | SPU_026078 | Lv-Act_5 |
| e19 | SPU_026090 | Lv-Gal3st1_11 |
| e19 | SPU_026094 | Lv-CathepsinL |
| e19 | SPU_026148 | Lv-Pnlip_12 |
| e19 | SPU_026172 | Lv-Hypp_2825 |
| e19 | SPU_026183 | Lv-Slc8a3 |
| e19 | SPU_026199 | Lv-Cltc |
| e19 | SPU_026229 | none |
| e19 | SPU_026436 | Lv-Ugt1L_8 |
| e19 | SPU_026438 | Lv-Abch1 |
| e19 | SPU_026493 | Lv-Aldh6a1 |
| e19 | SPU_026541 | Lv-Rps23 |
| e19 | SPU_026640 | Lv-Rpl7_1 |
| e19 | SPU_027219 | Lv-Mmp21L |
| e19 | SPU_027283 | Lv-Oxch_3 |
| e19 | SPU_027462 | Lv-Plcg |
| e19 | SPU_027555 | none |
| e19 | SPU_027576 | none |
| e19 | SPU_027748 | Lv-Vgcnl1 |
| e19 | SPU_027812 | none |
| e19 | SPU_027885 | Lv-Cp |
| e19 | SPU_028022 | Lv-HnmtL_3 |
| e19 | SPU_028060 | Lv-Nlr122 |
| e19 | SPU_028104 | Lv-Rpl15 |

| | | |
|---|---|---|
| e19 | SPU_028168 | Lv-Slc25a35 |
| e19 | SPU_028179 | Lv-Hypp_3011 |
| e19 | SPU_028226 | Lv-Rpl23 |
| e19 | SPU_028442 | Lv-Sarox_2 |
| e19 | SPU_028628 | Lv-Rdh8_6 |
| e19 | SPU_028633 | Lv-Gsto1_1 |
| e19 | SPU_028754 | Lv-Ggct |
| e19 | SPU_028760 | Lv-Aadac_5 |
| e19 | SPU_028790 | Lv-Slc16a3L |
| e19 | SPU_028866 | Lv-Rps4_1 |
| e19 | SPU_028873 | Lv-Atp5d |
| e19 | SPU_028940 | NA |
| e20 | SPU_000014 | Lv-Ube2a |
| e20 | SPU_000073 | Lv-Pik1 |
| e20 | SPU_000074 | Lv-Tsga14L |
| e20 | SPU_000128 | Lv-Tropmy1 |
| e20 | SPU_000300 | none |
| e20 | SPU_000402 | Lv-Gpr158_1 |
| e20 | SPU_000430 | Lv-Rnasek |
| e20 | SPU_000491 | Lv-Ppap2cL |
| e20 | SPU_000493 | Lv-Bbs9 |
| e20 | SPU_000613 | none |
| e20 | SPU_000670 | NA |
| e20 | SPU_000742 | Lv-Ube2v1/2 |
| e20 | SPU_000931 | Lv-St8sia |
| e20 | SPU_001242 | Lv-Hypp_1317 |
| e20 | SPU_001391 | Lv-Trim3L_2 |
| e20 | SPU_001497 | none |
| e20 | SPU_001540 | Lv-Kiaa0415L |
| e20 | SPU_001591 | Lv-Hypp_336 |
| e20 | SPU_001730 | none |
| e20 | SPU_001905 | Lv-Myd88L1 |
| e20 | SPU_001955 | NA |
| e20 | SPU_002105 | Lv-Camk1 |
| e20 | SPU_002236 | Lv-Hypp_3101 |
| e20 | SPU_002238 | Lv-Abcb9B |
| e20 | SPU_002247 | Lv-Unc44L_23 |
| e20 | SPU_002306 | Lv-Egf/Ccp/Zp |

| | | |
|---|---|---|
| e20 | SPU_002361 | NA |
| e20 | SPU_002371 | Lv-Cyp2L4 |
| e20 | SPU_002420 | Lv-Clect/Ly |
| e20 | SPU_002620 | Lv-Ttc21b-2 |
| e20 | SPU_002777 | Lv-Ugt2b34_1 |
| e20 | SPU_002850 | Lv-C10orf79L |
| e20 | SPU_002901 | Lv-Aldh9a1 |
| e20 | SPU_002945 | Lv-CalcneurL_1 |
| e20 | SPU_002992 | Lv-TyrL |
| e20 | SPU_003048 | Lv-Wdr19 |
| e20 | SPU_003065 | Lv-ArL2bp |
| e20 | SPU_003119 | Lv-Pp2asB55 |
| e20 | SPU_003146 | Lv-Hypp_375 |
| e20 | SPU_003174 | Lv-Tasp1 |
| e20 | SPU_003239 | Lv-Z373 |
| e20 | SPU_003261 | Lv-Macf1-2 |
| e20 | SPU_003263 | Lv-Spag16 |
| e20 | SPU_003404 | Lv-Dnahc9 |
| e20 | SPU_003425 | Lv-Chmp1a |
| e20 | SPU_003458 | none |
| e20 | SPU_003467 | Lv-Chst1_3 |
| e20 | SPU_003565 | none |
| e20 | SPU_003634 | Lv-Herc4-3 |
| e20 | SPU_003661 | Lv-Chrna7-2 |
| e20 | SPU_003732 | Lv-Hypp_3094 |
| e20 | SPU_003749 | Lv-Aspapd |
| e20 | SPU_003817 | Lv-Hypp_1484 |
| e20 | SPU_003830 | none |
| e20 | SPU_003919 | Lv-Efcab5 |
| e20 | SPU_003927 | none |
| e20 | SPU_004172 | Lv-SelcysL |
| e20 | SPU_004238 | Lv-Rgs12 |
| e20 | SPU_004242 | Lv-Ass_1 |
| e20 | SPU_004302 | Lv-Hypp_1511 |
| e20 | SPU_004329 | Lv-Hypp_412 |
| e20 | SPU_004349 | none |
| e20 | SPU_004459 | none |
| e20 | SPU_004521 | Lv-Hypp_419 |

| | | |
|---|---|---|
| e20 | SPU_004620 | Lv-Endrvt13 |
| e20 | SPU_004652 | none |
| e20 | SPU_004712 | Lv-Hypp_1539 |
| e20 | SPU_004751 | Lv-Atp8b2 |
| e20 | SPU_004835 | Lv-PppL_179 |
| e20 | SPU_004863 | Lv-Ckap1 |
| e20 | SPU_004897 | NA |
| e20 | SPU_004977 | Lv-Hypp_1 |
| e20 | SPU_005005 | Lv-Uhrf1bp1L |
| e20 | SPU_005017 | Lv-Plscr1_1 |
| e20 | SPU_005335 | Lv-Ddx58 |
| e20 | SPU_005468 | none |
| e20 | SPU_005485 | none |
| e20 | SPU_005607 | none |
| e20 | SPU_005608 | Lv-Hypp_1589 |
| e20 | SPU_005671 | Lv-Hypp_1595 |
| e20 | SPU_005697 | none |
| e20 | SPU_005719 | Lv-C1orf125 |
| e20 | SPU_005730 | Lv-Cops2 |
| e20 | SPU_005743 | Lv-Amd1 |
| e20 | SPU_005756 | Lv-C7orf63 |
| e20 | SPU_005791 | Lv-Iqck |
| e20 | SPU_005937 | Lv-Hypp_1619 |
| e20 | SPU_005973 | Lv-Oda/Ic1L |
| e20 | SPU_006042 | Lv-Chchd2 |
| e20 | SPU_006202 | none |
| e20 | SPU_006244 | none |
| e20 | SPU_006254 | Lv-Srcr48 |
| e20 | SPU_006266 | Lv-Hypp_1648 |
| e20 | SPU_006269 | Lv-Lphn3L_1 |
| e20 | SPU_006302 | Lv-Lrrc6 |
| e20 | SPU_006305 | Lv-Zfyve26 |
| e20 | SPU_006419 | Lv-Ccdc132 |
| e20 | SPU_006771 | Lv-Hypp_495 |
| e20 | SPU_006782 | Lv-C17orf28L |
| e20 | SPU_007158 | Lv-Hypp_508 |
| e20 | SPU_007200 | none |
| e20 | SPU_007208 | Lv-PpL |

| | | |
|---|---|---|
| e20 | SPU_007223 | Lv-Hephl1 |
| e20 | SPU_007260 | Lv-Hadhsc |
| e20 | SPU_007288 | Lv-Hypp_1711 |
| e20 | SPU_007334 | Lv-Hs2stL |
| e20 | SPU_007503 | NA |
| e20 | SPU_007516 | Lv-Mtpn |
| e20 | SPU_007665 | Lv-Pruneh |
| e20 | SPU_007788 | Lv-C6org186L |
| e20 | SPU_007827 | Lv-Slc5_1 |
| e20 | SPU_007927 | Lv-Sat2_1 |
| e20 | SPU_007997 | Lv-Dnhd1L |
| e20 | SPU_008085 | Lv-Mlck |
| e20 | SPU_008170 | none |
| e20 | SPU_008192 | Lv-Slc23a1_2 |
| e20 | SPU_008266 | none |
| e20 | SPU_008267 | Lv-Tlr044 |
| e20 | SPU_008424 | Lv-PolypL_938 |
| e20 | SPU_008434 | Lv-5/Lo |
| e20 | SPU_008574 | Lv-Hypp_1794 |
| e20 | SPU_008599 | Lv-Vbp1 |
| e20 | SPU_008657 | Lv-Kiaa1875L |
| e20 | SPU_008673 | Lv-Sema6 |
| e20 | SPU_008771 | NA |
| e20 | SPU_008774 | none |
| e20 | SPU_008819 | none |
| e20 | SPU_008871 | Lv-Tha1 |
| e20 | SPU_008872 | Lv-Ccdc96 |
| e20 | SPU_008883 | Lv-PppL_251 |
| e20 | SPU_008976 | Lv-Clect_70 |
| e20 | SPU_009094 | Lv-Hypp_1823 |
| e20 | SPU_009099 | Lv-Odc1_1 |
| e20 | SPU_009110 | NA |
| e20 | SPU_009136 | Lv-XyltL_1 |
| e20 | SPU_009161 | Lv-Nlr125_1 |
| e20 | SPU_009182 | Lv-Sdhd |
| e20 | SPU_009264 | Lv-Endrvt18 |
| e20 | SPU_009485 | Lv-Hypp_48 |
| e20 | SPU_009591 | Lv-Hypp_51 |

| | | |
|---|---|---|
| e20 | SPU_009883 | Lv-Bnip2 |
| e20 | SPU_009884 | NA |
| e20 | SPU_009886 | Lv-Clect_72 |
| e20 | SPU_009949 | Lv-Ef1a_1 |
| e20 | SPU_009950 | none |
| e20 | SPU_010123 | Lv-Igf_10 |
| e20 | SPU_010182 | Lv-RtL_23 |
| e20 | SPU_010244 | none |
| e20 | SPU_010377 | Lv-Ipgm |
| e20 | SPU_010384 | Lv-Paqr3 |
| e20 | SPU_010570 | Lv-Kif20A |
| e20 | SPU_010629 | Lv-Mekk2/3 |
| e20 | SPU_010707 | Lv-PolppL_89 |
| e20 | SPU_010786 | Lv-Mcl |
| e20 | SPU_011125 | NA |
| e20 | SPU_011336 | Lv-Glt_3 |
| e20 | SPU_011369 | Lv-AP |
| e20 | SPU_011375 | NA |
| e20 | SPU_011510 | Lv-Hypp_1985 |
| e20 | SPU_011538 | Lv-Rhogap20 |
| e20 | SPU_011562 | Lv-Pla2 |
| e20 | SPU_011571 | Lv-Hypp_71 |
| e20 | SPU_011648 | Lv-Fn3f_39 |
| e20 | SPU_011767 | Lv-Hypp_658 |
| e20 | SPU_011909 | none |
| e20 | SPU_012002 | Lv-Ttc21b |
| e20 | SPU_012055 | none |
| e20 | SPU_012061 | Lv-Unc44_226 |
| e20 | SPU_012067 | NA |
| e20 | SPU_012140 | none |
| e20 | SPU_012254 | Lv-Bzrap1L |
| e20 | SPU_012488 | Lv-Qdpr |
| e20 | SPU_012501 | Lv-Tmem131 |
| e20 | SPU_012512 | none |
| e20 | SPU_012703 | none |
| e20 | SPU_012715 | Lv-Stx17 |
| e20 | SPU_012777 | Lv-C3orf37L |
| e20 | SPU_012877 | Lv-Costal2 |

| | | |
|---|---|---|
| e20 | SPU_012881 | none |
| e20 | SPU_012895 | Lv-Ccdc108 |
| e20 | SPU_012896 | none |
| e20 | SPU_012918 | Lv-Hes1 |
| e20 | SPU_012925 | Lv-Endrvt82 |
| e20 | SPU_013009 | Lv-Klhl29-2 |
| e20 | SPU_013083 | Lv-Hypp_3063 |
| e20 | SPU_013117 | none |
| e20 | SPU_013287 | none |
| e20 | SPU_013343 | Lv-PolypL_946 |
| e20 | SPU_013387 | Lv-C6orf103L |
| e20 | SPU_013395 | Lv-Spg11_1 |
| e20 | SPU_013408 | none |
| e20 | SPU_013425 | Lv-Endrvt28 |
| e20 | SPU_013640 | Lv-Hypp_726 |
| e20 | SPU_013645 | Lv-Hypp_2794 |
| e20 | SPU_013730 | none |
| e20 | SPU_013761 | Lv-Hypp_2104 |
| e20 | SPU_013846 | NA |
| e20 | SPU_013954 | Lv-Fn3f_4 |
| e20 | SPU_014012 | Lv-C2orf81L |
| e20 | SPU_014228 | Lv-Bsdc1 |
| e20 | SPU_014252 | none |
| e20 | SPU_014453 | Lv-Slc6a3 |
| e20 | SPU_014463 | Lv-Iqch |
| e20 | SPU_014658 | Lv-Pae |
| e20 | SPU_014673 | none |
| e20 | SPU_014729 | Lv-C6orf118L |
| e20 | SPU_014810 | Lv-Lrrc23 |
| e20 | SPU_014830 | Lv-AnL11_1 |
| e20 | SPU_014922 | none |
| e20 | SPU_014967 | Lv-Tp53bp2 |
| e20 | SPU_015079 | Lv-Clect_33 |
| e20 | SPU_015084 | Lv-Acbd5 |
| e20 | SPU_015090 | Lv-Aacy1 |
| e20 | SPU_015124 | none |
| e20 | SPU_015437 | Lv-Kif2L |
| e20 | SPU_015478 | Lv-Tsnaxip1_1 |

| | | |
|---|---|---|
| e20 | SPU_015581 | Lv-Igf_11 |
| e20 | SPU_015605 | Lv-Rab23 |
| e20 | SPU_015654 | Lv-TnfsfL3 |
| e20 | SPU_015685 | Lv-Rnf123 |
| e20 | SPU_015714 | Lv-Hypp_114 |
| e20 | SPU_015722 | Lv-C3orf23L |
| e20 | SPU_015816 | Lv-Rt37 |
| e20 | SPU_015918 | Lv-Fn3_31 |
| e20 | SPU_015951 | Lv-Hypp_123 |
| e20 | SPU_016051 | Lv-Arg_1 |
| e20 | SPU_016108 | Lv-Invs |
| e20 | SPU_016118 | NA |
| e20 | SPU_016121 | Lv-Polyp |
| e20 | SPU_016140 | Lv-Usp6nL |
| e20 | SPU_016158 | Lv-PolppL_105 |
| e20 | SPU_016224 | none |
| e20 | SPU_016311 | none |
| e20 | SPU_016344 | Lv-Pemt |
| e20 | SPU_016478 | Lv-Kiaa0329 |
| e20 | SPU_016649 | none |
| e20 | SPU_016769 | none |
| e20 | SPU_016825 | Lv-Pecr |
| e20 | SPU_016841 | NA |
| e20 | SPU_016895 | none |
| e20 | SPU_017083 | Lv-Plscr3 |
| e20 | SPU_017114 | Lv-Unc44_123 |
| e20 | SPU_017173 | Lv-Dnah9L |
| e20 | SPU_017262 | Lv-Hypp_869 |
| e20 | SPU_017339 | Lv-Hypp_875 |
| e20 | SPU_017656 | Lv-Z10 |
| e20 | SPU_017667 | none |
| e20 | SPU_017692 | NA |
| e20 | SPU_017775 | NA |
| e20 | SPU_017869 | Lv-Endrvt97 |
| e20 | SPU_017978 | Lv-Rgs12-2 |
| e20 | SPU_018012 | Lv-Dtd1h |
| e20 | SPU_018038 | Lv-Dirc2_1 |
| e20 | SPU_018150 | Lv-Hypp_909 |

| | | |
|---|---|---|
| e20 | SPU_018163 | Lv-Ap1s2 |
| e20 | SPU_018186 | Lv-Mosc2_2 |
| e20 | SPU_018193 | Lv-Sipp1 |
| e20 | SPU_018370 | none |
| e20 | SPU_018381 | Lv-Hunt(C_terminal) |
| e20 | SPU_018434 | Lv-Vps25 |
| e20 | SPU_018536 | Lv-Bahd1 |
| e20 | SPU_018537 | Lv-C20orf26L |
| e20 | SPU_018575 | none |
| e20 | SPU_018642 | Lv-Hypp_2398 |
| e20 | SPU_018732 | Lv-Sh3bgrL3 |
| e20 | SPU_018751 | Lv-Rdh10 |
| e20 | SPU_018837 | Lv-Unc44_137 |
| e20 | SPU_018923 | none |
| e20 | SPU_018950 | Lv-Unc44_138 |
| e20 | SPU_018956 | Lv-Endrvt38 |
| e20 | SPU_018958 | Lv-SmtnL |
| e20 | SPU_019141 | Lv-Atp8a1_1 |
| e20 | SPU_019506 | Lv-Dyi2 |
| e20 | SPU_019510 | Lv-Rdh8_5 |
| e20 | SPU_019558 | Lv-Arl2l1 |
| e20 | SPU_019570 | Lv-Ttll3 |
| e20 | SPU_019590 | Lv-Ift52_1 |
| e20 | SPU_019693 | Lv-Ssna1 |
| e20 | SPU_019707 | Lv-Hypp_942 |
| e20 | SPU_019713 | NA |
| e20 | SPU_019730 | none |
| e20 | SPU_019741 | Lv-EndrvtL |
| e20 | SPU_019854 | Lv-Notch1-17 |
| e20 | SPU_019984 | Lv-Ptpn9_1 |
| e20 | SPU_020028 | Lv-Hrh2L_8 |
| e20 | SPU_020044 | Lv-Unc44_142 |
| e20 | SPU_020232 | none |
| e20 | SPU_020252 | Lv-PppL_214 |
| e20 | SPU_020475 | none |
| e20 | SPU_020581 | Lv-DpdL |
| e20 | SPU_020617 | Lv-Tymp |
| e20 | SPU_020634 | Lv-Klp6L3 |

| | | |
|---|---|---|
| e20 | SPU_020757 | Lv-Dlat |
| e20 | SPU_020804 | Lv-Commd10 |
| e20 | SPU_020881 | Lv-Echdc1 |
| e20 | SPU_021180 | Lv-Endrvt42 |
| e20 | SPU_021196 | Lv-Hypp_170 |
| e20 | SPU_021206 | NA |
| e20 | SPU_021227 | Lv-Dnah8 |
| e20 | SPU_021238 | Lv-Big3 |
| e20 | SPU_021407 | Lv-Hbp1 |
| e20 | SPU_021411 | Lv-Ndc80L |
| e20 | SPU_021428 | Lv-L1 |
| e20 | SPU_021463 | Lv-Wdr19_2 |
| e20 | SPU_021573 | Lv-Ct |
| e20 | SPU_021623 | Lv-Slc17a5_7 |
| e20 | SPU_021651 | Lv-Ift172_2 |
| e20 | SPU_021683 | Lv-Surreal/Gpcr91 |
| e20 | SPU_021813 | Lv-Agps |
| e20 | SPU_021918 | Lv-Ift140_1 |
| e20 | SPU_022092 | none |
| e20 | SPU_022204 | Lv-PppL_132 |
| e20 | SPU_022225 | Lv-Pex12 |
| e20 | SPU_022445 | Lv-Chmp1b |
| e20 | SPU_022484 | none |
| e20 | SPU_022493 | Lv-Pcs |
| e20 | SPU_022587 | Lv-Nphp3 |
| e20 | SPU_022604 | Lv-Syap1 |
| e20 | SPU_022664 | none |
| e20 | SPU_022704 | Lv-Cnt3 |
| e20 | SPU_022727 | NA |
| e20 | SPU_022778 | Lv-Chmp4b |
| e20 | SPU_022842 | Lv-Dbt |
| e20 | SPU_023015 | none |
| e20 | SPU_023054 | Lv-Hypp_2647 |
| e20 | SPU_023245 | none |
| e20 | SPU_023315 | Lv-RtnpLp |
| e20 | SPU_023483 | Lv-PolypL_120 |
| e20 | SPU_023583 | Lv-Nek4b |
| e20 | SPU_023617 | Lv-Abca3a |

| | | |
|---|---|---|
| e20 | SPU_023683 | Lv-Hypp_2682 |
| e20 | SPU_023688 | Lv-Rej8A |
| e20 | SPU_024069 | Lv-Gramd1b |
| e20 | SPU_024071 | Lv-Pcca |
| e20 | SPU_024073 | Lv-Ly/Egf/Sea |
| e20 | SPU_024085 | Lv-Kcnk18L_2 |
| e20 | SPU_024117 | Lv-Hypp_2710 |
| e20 | SPU_024356 | NA |
| e20 | SPU_024431 | Lv-Cryl1_1 |
| e20 | SPU_024490 | Lv-Mamdc4_2 |
| e20 | SPU_024507 | Lv-Skp1 |
| e20 | SPU_024513 | Lv-RagA |
| e20 | SPU_024521 | Lv-Ttc3L |
| e20 | SPU_024535 | Lv-Agl |
| e20 | SPU_024554 | Lv-Abhd8 |
| e20 | SPU_024610 | Lv-MuskL |
| e20 | SPU_024617 | Lv-Tuba1c_1 |
| e20 | SPU_024681 | none |
| e20 | SPU_024739 | Lv-AnL9 |
| e20 | SPU_024764 | Lv-R/Ptp/B |
| e20 | SPU_024793 | Lv-Gol/Dup |
| e20 | SPU_025305 | Lv-Kiaa0562L_1 |
| e20 | SPU_025398 | none |
| e20 | SPU_025420 | Lv-C10orf92L |
| e20 | SPU_025432 | Lv-Igf_9 |
| e20 | SPU_025567 | Lv-Agxt2 |
| e20 | SPU_025585 | Lv-Cand1_1 |
| e20 | SPU_025816 | Lv-Tbcel_1 |
| e20 | SPU_025915 | Lv-Kiaa0174L |
| e20 | SPU_026293 | Lv-Unc44_179 |
| e20 | SPU_026331 | Lv-C1orf125L |
| e20 | SPU_026342 | Lv-Dnhd1L_2 |
| e20 | SPU_026365 | Lv-Sri |
| e20 | SPU_026492 | NA |
| e20 | SPU_026504 | Lv-Hypp_2848 |
| e20 | SPU_026529 | none |
| e20 | SPU_026634 | Lv-Chmp2a |
| e20 | SPU_026814 | Lv-Slc35f5 |

| | | |
|---|---|---|
| e20 | SPU_027153 | Lv-Cyp2L64 |
| e20 | SPU_027167 | Lv-Kiaa1012L |
| e20 | SPU_027313 | Lv-Hypp_265 |
| e20 | SPU_027326 | Lv-Dnah9 |
| e20 | SPU_027559 | Lv-Yap1 |
| e20 | SPU_027629 | none |
| e20 | SPU_027717 | none |
| e20 | SPU_027751 | Lv-Mfs |
| e20 | SPU_027799 | Lv-Slc7a11_12 |
| e20 | SPU_027820 | Lv-PppL_163 |
| e20 | SPU_027840 | Lv-Fkbp12 |
| e20 | SPU_027988 | Lv-Endrvt119 |
| e20 | SPU_028073 | Lv-Ipp2 |
| e20 | SPU_028102 | none |
| e20 | SPU_028172 | Lv-Hypp_2940 |
| e20 | SPU_028205 | Lv-Dynact_p25ph |
| e20 | SPU_028280 | none |
| e20 | SPU_028336 | Lv-Lppr |
| e20 | SPU_028478 | Lv-Jnk |
| e20 | SPU_028532 | Lv-Gpx1_1 |
| e20 | SPU_028599 | NA |
| e20 | SPU_028623 | Lv-Unc44_354 |
| e20 | SPU_028661 | none |
| e20 | SPU_028665 | Lv-Fn3f_47 |
| e20 | SPU_028682 | NA |
| e20 | SPU_028775 | Lv-Dag1 |
| e20 | SPU_028812 | Lv-PolypL_130 |
| e20 | SPU_028887 | none |
| e20 | SPU_030014 | NA |
| e20 | SPU_030225 | Lv-Axndhc3h |
| e20 | SPU_030228 | Lv-Axndhc7h |
| e20 | SPU_030235 | Lv-Cytd2hc |
| f1 | SPU_000021 | Lv-HscbL |
| f1 | SPU_000046 | Lv-Cdc7 |
| f1 | SPU_000067 | Lv-Crj1L |
| f1 | SPU_000069 | Lv-Glyt1L |
| f1 | SPU_000122 | Lv-Nmcp1l |
| f1 | SPU_000170 | Lv-Ars2 |

| | | |
|---|---|---|
| f1 | SPU_000171 | Lv-Ars2_1 |
| f1 | SPU_000212 | Lv-Gnptabl |
| f1 | SPU_000213 | Lv-Tmem69 |
| f1 | SPU_000214 | Lv-C20orf31 |
| f1 | SPU_000320 | Lv-Hypp_1248 |
| f1 | SPU_000326 | Lv-Elp2h |
| f1 | SPU_000352 | none |
| f1 | SPU_000480 | Lv-Zfp509L |
| f1 | SPU_000554 | Lv-Fn3/Igf_19 |
| f1 | SPU_000570 | Lv-Hypp_308 |
| f1 | SPU_000604 | none |
| f1 | SPU_000614 | Lv-SacSl_3 |
| f1 | SPU_000622 | Lv-Sap130 |
| f1 | SPU_000650 | Lv-Snip1 |
| f1 | SPU_000688 | NA |
| f1 | SPU_000689 | Lv-Pop1 |
| f1 | SPU_000707 | Lv-Srbd1 |
| f1 | SPU_000714 | Lv-PppL_292 |
| f1 | SPU_000755 | Lv-B3gat3 |
| f1 | SPU_000761 | Lv-B3gnt5 |
| f1 | SPU_000770 | none |
| f1 | SPU_000786 | Lv-Ddx20 |
| f1 | SPU_000806 | Lv-Tk9 |
| f1 | SPU_000850 | Lv-Tbccd1 |
| f1 | SPU_000887 | NA |
| f1 | SPU_001031 | Lv-Zmiz1L |
| f1 | SPU_001044 | none |
| f1 | SPU_001085 | Lv-Rpia |
| f1 | SPU_001120 | none |
| f1 | SPU_001121 | NA |
| f1 | SPU_001122 | NA |
| f1 | SPU_001311 | Lv-Qrsl1 |
| f1 | SPU_001371 | none |
| f1 | SPU_001409 | Lv-Dmbt1L_2 |
| f1 | SPU_001421 | none |
| f1 | SPU_001439 | Lv-Polr2c |
| f1 | SPU_001477 | none |
| f1 | SPU_001605 | Lv-Sfrs10 |

| | | |
|---|---|---|
| f1 | SPU_001669 | Lv-Slc4a11 |
| f1 | SPU_001734 | none |
| f1 | SPU_001777 | Lv-Sirt6_1 |
| f1 | SPU_001815 | Lv-PolppL_10 |
| f1 | SPU_001816 | Lv-Phf14 |
| f1 | SPU_001819 | Lv-Faastkd2L |
| f1 | SPU_001889 | Lv-Srp72 |
| f1 | SPU_001897 | none |
| f1 | SPU_001976 | Lv-Surf6L |
| f1 | SPU_002117 | Lv-LshmanlsnL |
| f1 | SPU_002132 | Lv-Nol8 |
| f1 | SPU_002150 | Lv-Prp38a |
| f1 | SPU_002210 | Lv-Cdk1 |
| f1 | SPU_002214 | Lv-Lon |
| f1 | SPU_002217 | Lv-Hypp_350 |
| f1 | SPU_002220 | Lv-Smn1 |
| f1 | SPU_002242 | Lv-Hypp_1380 |
| f1 | SPU_002285 | Lv-Tceb3 |
| f1 | SPU_002332 | Lv-C9orf97L |
| f1 | SPU_002353 | none |
| f1 | SPU_002358 | Lv-Hypp_1389 |
| f1 | SPU_002467 | Lv-Agrin |
| f1 | SPU_002628 | none |
| f1 | SPU_002629 | Lv-Hypp_1405 |
| f1 | SPU_002644 | Lv-Apex2 |
| f1 | SPU_002709 | Lv-Cept1 |
| f1 | SPU_002808 | Lv-Fut |
| f1 | SPU_002891 | none |
| f1 | SPU_002914 | none |
| f1 | SPU_002918 | Lv-Hypp_1420 |
| f1 | SPU_002925 | Lv-Lrr |
| f1 | SPU_002936 | Lv-Tmed5 |
| f1 | SPU_002938 | Lv-Acp5a |
| f1 | SPU_002953 | none |
| f1 | SPU_003025 | Lv-PolppL_18 |
| f1 | SPU_003030 | Lv-PolppL_23 |
| f1 | SPU_003122 | none |
| f1 | SPU_003256 | NA |

| | | |
|---|---|---|
| f1 | SPU_003278 | Lv-Hypp_379 |
| f1 | SPU_003390 | none |
| f1 | SPU_003403 | Lv-Mst |
| f1 | SPU_003436 | none |
| f1 | SPU_003468 | Lv-Endrvt12 |
| f1 | SPU_003470 | Lv-Spz12 |
| f1 | SPU_003545 | Lv-Mysm1 |
| f1 | SPU_003555 | none |
| f1 | SPU_003561 | Lv-Ptdss1 |
| f1 | SPU_003573 | none |
| f1 | SPU_003593 | Lv-Fbxl10 |
| f1 | SPU_003630 | Lv-Wrnip1L |
| f1 | SPU_003644 | Lv-Tpr |
| f1 | SPU_003655 | Lv-Frizz1/2/7 |
| f1 | SPU_003672 | Lv-Trappc9 |
| f1 | SPU_003807 | Lv-Hypp_398 |
| f1 | SPU_003861 | none |
| f1 | SPU_003897 | Lv-Prc1L |
| f1 | SPU_003989 | Lv-Hypp_403 |
| f1 | SPU_004001 | Lv-Polr2i |
| f1 | SPU_004004 | Lv-AsrgL1 |
| f1 | SPU_004055 | none |
| f1 | SPU_004075 | Lv-Abcc5i |
| f1 | SPU_004213 | none |
| f1 | SPU_004249 | Lv-Stx8 |
| f1 | SPU_004296 | none |
| f1 | SPU_004304 | Lv-Hpd |
| f1 | SPU_004326 | Lv-Tpk1 |
| f1 | SPU_004353 | Lv-Ndfip1l |
| f1 | SPU_004383 | Lv-Entpd4 |
| f1 | SPU_004494 | Lv-Msh5 |
| f1 | SPU_004535 | Lv-PppL_177 |
| f1 | SPU_004555 | none |
| f1 | SPU_004802 | Lv-Z232 |
| f1 | SPU_004840 | Lv-Etif4BL |
| f1 | SPU_004853 | Lv-Hypp_432 |
| f1 | SPU_004875 | Lv-Apip |
| f1 | SPU_004940 | Lv-Ecm/VWC/Vwd |

| | | |
|---|---|---|
| f1 | SPU_004960 | none |
| f1 | SPU_004970 | NA |
| f1 | SPU_005035 | Lv-Z339 |
| f1 | SPU_005037 | NA |
| f1 | SPU_005205 | Lv-Psrl |
| f1 | SPU_005275 | Lv-Mjd |
| f1 | SPU_005314 | Lv-Zcsl2 |
| f1 | SPU_005333 | none |
| f1 | SPU_005369 | none |
| f1 | SPU_005415 | Lv-Pde10a |
| f1 | SPU_005445 | Lv-Fem1bL |
| f1 | SPU_005525 | Lv-Paf1-2 |
| f1 | SPU_005548 | Lv-ScribL |
| f1 | SPU_005610 | Lv-Taok2i1 |
| f1 | SPU_005612 | Lv-Hypp_458 |
| f1 | SPU_005655 | Lv-Cyp2L12 |
| f1 | SPU_005729 | Lv-Cks1B |
| f1 | SPU_005778 | Lv-Qars |
| f1 | SPU_005805 | none |
| f1 | SPU_005962 | Lv-Mbd4 |
| f1 | SPU_005998 | Lv-Ddx52 |
| f1 | SPU_006040 | Lv-Pla2p |
| f1 | SPU_006162 | Lv-Rsafd1 |
| f1 | SPU_006288 | Lv-Hypp_1651 |
| f1 | SPU_006365 | Lv-Elmod3 |
| f1 | SPU_006370 | Lv-Ncam |
| f1 | SPU_006375 | Lv-Atg4hb |
| f1 | SPU_006393 | Lv-Usp38 |
| f1 | SPU_006447 | Lv-TpaseL_1 |
| f1 | SPU_006543 | Lv-PppL_186 |
| f1 | SPU_006566 | Lv-PppL_187 |
| f1 | SPU_006651 | none |
| f1 | SPU_006668 | Lv-Rrp9 |
| f1 | SPU_006746 | NA |
| f1 | SPU_006788 | Lv-Ttl1 |
| f1 | SPU_006813 | Lv-Hairy |
| f1 | SPU_006846 | none |
| f1 | SPU_006861 | Lv-Ino80e |

| | | |
|---|---|---|
| f1 | SPU_006888 | Lv-Slc30a6 |
| f1 | SPU_006900 | Lv-Fusip1 |
| f1 | SPU_006907 | Lv-Gtf3c4L |
| f1 | SPU_006937 | Lv-Piga |
| f1 | SPU_006938 | Lv-Ube1dc1 |
| f1 | SPU_006939 | Lv-Tlr034 |
| f1 | SPU_006965 | none |
| f1 | SPU_006966 | Lv-Hypp_1690 |
| f1 | SPU_006977 | Lv-Hypp_3072 |
| f1 | SPU_007017 | none |
| f1 | SPU_007018 | Lv-Hypp_505 |
| f1 | SPU_007041 | Lv-B3galt1_4 |
| f1 | SPU_007089 | Lv-Hypp_1696 |
| f1 | SPU_007190 | Lv-Z185 |
| f1 | SPU_007192 | none |
| f1 | SPU_007219 | Lv-Rnf146l |
| f1 | SPU_007232 | Lv-Ica1 |
| f1 | SPU_007267 | Lv-Pla2g1b |
| f1 | SPU_007273 | Lv-PPcdc |
| f1 | SPU_007309 | Lv-Lypd6b |
| f1 | SPU_007392 | Lv-Pigb |
| f1 | SPU_007403 | Lv-Slc5a6 |
| f1 | SPU_007417 | Lv-Dhx16_1 |
| f1 | SPU_007533 | Lv-Lhh2af |
| f1 | SPU_007588 | Lv-Ttyh2 |
| f1 | SPU_007604 | Lv-Slco_1 |
| f1 | SPU_007656 | none |
| f1 | SPU_007657 | Lv-Large_2 |
| f1 | SPU_007672 | Lv-Css2 |
| f1 | SPU_007684 | Lv-Sec62h |
| f1 | SPU_007808 | NA |
| f1 | SPU_007849 | none |
| f1 | SPU_007974 | Lv-Aggf1L |
| f1 | SPU_007995 | Lv-Ttrspn_4 |
| f1 | SPU_008012 | NA |
| f1 | SPU_008040 | Lv-Aatf |
| f1 | SPU_008071 | Lv-Ccdc94 |
| f1 | SPU_008120 | Lv-Slmap |

| | | |
|---|---|---|
| f1 | SPU_008187 | Lv-Prdm7 |
| f1 | SPU_008215 | Lv-Upf1 |
| f1 | SPU_008241 | Lv-Trip11L |
| f1 | SPU_008242 | NA |
| f1 | SPU_008287 | Lv-Mtmr1/2 |
| f1 | SPU_008340 | Lv-Hypp_544 |
| f1 | SPU_008412 | Lv-Gpn3 |
| f1 | SPU_008508 | Lv-Cox4L_1 |
| f1 | SPU_008519 | Lv-Hypp_1788 |
| f1 | SPU_008520 | none |
| f1 | SPU_008521 | none |
| f1 | SPU_008526 | Lv-GbA2 |
| f1 | SPU_008565 | Lv-Slc16a4L_1 |
| f1 | SPU_008625 | Lv-TbL1xr1 |
| f1 | SPU_008656 | Lv-Hsp40D |
| f1 | SPU_008702 | Lv-Kiaa2013L |
| f1 | SPU_008730 | NA |
| f1 | SPU_008793 | Lv-Bind1 |
| f1 | SPU_008839 | none |
| f1 | SPU_008856 | Lv-Fbxo7 |
| f1 | SPU_008945 | Lv-Notchh1_7 |
| f1 | SPU_009012 | Lv-Chchd4 |
| f1 | SPU_009028 | Lv-Tchp |
| f1 | SPU_009029 | Lv-Abhd11 |
| f1 | SPU_009068 | NA |
| f1 | SPU_009095 | Lv-Bms1l1 |
| f1 | SPU_009140 | Lv-Pygopos |
| f1 | SPU_009184 | none |
| f1 | SPU_009213 | Lv-Nup98 |
| f1 | SPU_009225 | Lv-Unk_49 |
| f1 | SPU_009262 | Lv-Oct1-2 |
| f1 | SPU_009266 | Lv-Cpsf1_1 |
| f1 | SPU_009326 | none |
| f1 | SPU_009555 | Lv-Drg1 |
| f1 | SPU_009632 | Lv-Samd9L_6 |
| f1 | SPU_009642 | Lv-Rreb1 |
| f1 | SPU_009645 | Lv-Clptm1_2 |
| f1 | SPU_009657 | none |

| | | |
|---|---|---|
| f1 | SPU_009794 | Lv-Hypp_55 |
| f1 | SPU_009810 | Lv-Ppm1a/b |
| f1 | SPU_009843 | none |
| f1 | SPU_009847 | Lv-Z156 |
| f1 | SPU_009889 | Lv-Vps13hb2 |
| f1 | SPU_009903 | Lv-Pdss2_1 |
| f1 | SPU_009948 | NA |
| f1 | SPU_010103 | Lv-Clca2_1 |
| f1 | SPU_010158 | NA |
| f1 | SPU_010174 | Lv-Tk2 |
| f1 | SPU_010272 | Lv-Oxch |
| f1 | SPU_010339 | Lv-Mrgbp |
| f1 | SPU_010422 | Lv-Ppp4r4 |
| f1 | SPU_010495 | none |
| f1 | SPU_010497 | Lv-Drd1L_1 |
| f1 | SPU_010565 | Lv-Slit1(C-terminus) |
| f1 | SPU_010726 | NA |
| f1 | SPU_010735 | none |
| f1 | SPU_010736 | Lv-Ccdc45 |
| f1 | SPU_010783 | Lv-Wdr70 |
| f1 | SPU_010846 | Lv-Nlk |
| f1 | SPU_010847 | Lv-Asl |
| f1 | SPU_010874 | Lv-Lcmt1 |
| f1 | SPU_010946 | Lv-G12 |
| f1 | SPU_011065 | Lv-BetaGi |
| f1 | SPU_011080 | Lv-SoxH |
| f1 | SPU_011159 | Lv-Foxred1 |
| f1 | SPU_011161 | NA |
| f1 | SPU_011214 | Lv-Mnd1 |
| f1 | SPU_011321 | NA |
| f1 | SPU_011323 | Lv-Hypp_3066 |
| f1 | SPU_011357 | Lv-Cds2 |
| f1 | SPU_011578 | Lv-Hypp_1990 |
| f1 | SPU_011640 | Lv-Arid5b |
| f1 | SPU_011658 | Lv-CycH |
| f1 | SPU_011661 | NA |
| f1 | SPU_011715 | none |
| f1 | SPU_011734 | Lv-Paf1 |

| | | |
|---|---|---|
| f1 | SPU_011799 | Lv-PolypL_2 |
| f1 | SPU_011848 | Lv-Dnajc2 |
| f1 | SPU_011886 | Lv-Lsm16 |
| f1 | SPU_011902 | Lv-Ythdc2_1 |
| f1 | SPU_011903 | Lv-Cep120-2 |
| f1 | SPU_011906 | none |
| f1 | SPU_011939 | Lv-C7orf20 |
| f1 | SPU_011940 | Lv-Usp22 |
| f1 | SPU_011970 | Lv-Rabl3 |
| f1 | SPU_011974 | Lv-Hypp_2009 |
| f1 | SPU_012018 | Lv-Ccdc97 |
| f1 | SPU_012070 | Lv-Cpt2 |
| f1 | SPU_012084 | Lv-Sac |
| f1 | SPU_012109 | Lv-Ttrspn_9 |
| f1 | SPU_012167 | Lv-Hypp_672 |
| f1 | SPU_012221 | Lv-UdpN_act_a_Dgalspp |
| f1 | SPU_012240 | Lv-Znf259l |
| f1 | SPU_012283 | Lv-Sgsm1 |
| f1 | SPU_012288 | Lv-Sdr_3 |
| f1 | SPU_012315 | Lv-Z215 |
| f1 | SPU_012321 | NA |
| f1 | SPU_012360 | NA |
| f1 | SPU_012457 | Lv-Taf3 |
| f1 | SPU_012507 | Lv-Lyz |
| f1 | SPU_012536 | Lv-Endrvt80 |
| f1 | SPU_012544 | none |
| f1 | SPU_012549 | Lv-Mt1-4/MmpL4 |
| f1 | SPU_012565 | none |
| f1 | SPU_012583 | Lv-Z383 |
| f1 | SPU_012639 | Lv-Exoc7 |
| f1 | SPU_012674 | Lv-Cdk9 |
| f1 | SPU_012724 | Lv-Z218 |
| f1 | SPU_012775 | none |
| f1 | SPU_012787 | none |
| f1 | SPU_012848 | Lv-NprA |
| f1 | SPU_012902 | Lv-Hypp_2055 |
| f1 | SPU_012912 | Lv-Z150 |
| f1 | SPU_012914 | Lv-Z62 |

| | | |
|---|---|---|
| f1 | SPU_012965 | Lv-Senp6L |
| f1 | SPU_013163 | Lv-Suz12 |
| f1 | SPU_013195 | Lv-Balz1b |
| f1 | SPU_013277 | none |
| f1 | SPU_013310 | Lv-Pdpr_1 |
| f1 | SPU_013344 | Lv-PolypL_947 |
| f1 | SPU_013396 | Lv-Timm44 |
| f1 | SPU_013397 | none |
| f1 | SPU_013415 | Lv-Z328 |
| f1 | SPU_013442 | Lv-Hypp_92 |
| f1 | SPU_013446 | Lv-Hypp_2082 |
| f1 | SPU_013460 | Lv-Slc19a2 |
| f1 | SPU_013482 | none |
| f1 | SPU_013491 | Lv-Notchh1_9 |
| f1 | SPU_013541 | Lv-Hypp_720 |
| f1 | SPU_013551 | Lv-Alad_1 |
| f1 | SPU_013575 | Lv-Dnajc8l |
| f1 | SPU_013598 | NA |
| f1 | SPU_013668 | Lv-Clect/Fn3_11 |
| f1 | SPU_013706 | Lv-Mrps27 |
| f1 | SPU_013707 | none |
| f1 | SPU_013807 | Lv-Rer1h |
| f1 | SPU_013853 | Lv-Ggps1 |
| f1 | SPU_013863 | Lv-Rab20 |
| f1 | SPU_013870 | Lv-Hypp_2112 |
| f1 | SPU_013871 | Lv-Thap2 |
| f1 | SPU_013893 | Lv-Cts5 |
| f1 | SPU_013902 | Lv-Z386 |
| f1 | SPU_013945 | Lv-Chrna9_1 |
| f1 | SPU_014028 | Lv-Bak |
| f1 | SPU_014070 | Lv-Hypp_2125 |
| f1 | SPU_014130 | NA |
| f1 | SPU_014131 | Lv-Notch |
| f1 | SPU_014168 | Lv-AT_V |
| f1 | SPU_014205 | Lv-Taar7aL |
| f1 | SPU_014225 | NA |
| f1 | SPU_014288 | Lv-Rpp14 |
| f1 | SPU_014330 | Lv-Mak10 |

| | | |
|---|---|---|
| f1 | SPU_014360 | Lv-Zgpat |
| f1 | SPU_014362 | Lv-Zdhhc12 |
| f1 | SPU_014370 | Lv-Tysnd1 |
| f1 | SPU_014410 | Lv-Hypp_2993 |
| f1 | SPU_014435 | none |
| f1 | SPU_014452 | Lv-Hypp_2152 |
| f1 | SPU_014459 | Lv-Rcc2 |
| f1 | SPU_014477 | Lv-Cnp_1 |
| f1 | SPU_014483 | none |
| f1 | SPU_014621 | Lv-Pdcd2 |
| f1 | SPU_014836 | Lv-Cp2 |
| f1 | SPU_014923 | Lv-Z28 |
| f1 | SPU_014924 | Lv-C2orf37 |
| f1 | SPU_014974 | Lv-Btaf1 |
| f1 | SPU_015017 | Lv-Btbd11 |
| f1 | SPU_015182 | NA |
| f1 | SPU_015222 | none |
| f1 | SPU_015223 | Lv-Vps13hb |
| f1 | SPU_015260 | Lv-Mrps23 |
| f1 | SPU_015373 | Lv-Pex3 |
| f1 | SPU_015411 | Lv-Skiv2l |
| f1 | SPU_015416 | Lv-IncenpL |
| f1 | SPU_015751 | Lv-Z220 |
| f1 | SPU_015782 | none |
| f1 | SPU_015814 | Lv-Fasn_1 |
| f1 | SPU_015869 | Lv-Oaf |
| f1 | SPU_015889 | Lv-Rev1 |
| f1 | SPU_015935 | Lv-SnrpA/U2Bl |
| f1 | SPU_015965 | Lv-Iqwd1 |
| f1 | SPU_016008 | Lv-Alk2 |
| f1 | SPU_016017 | Lv-Med13 |
| f1 | SPU_016018 | Lv-B3gtl |
| f1 | SPU_016031 | none |
| f1 | SPU_016047 | Lv-EhH3_33 |
| f1 | SPU_016126 | Lv-Hypp_814 |
| f1 | SPU_016151 | none |
| f1 | SPU_016251 | Lv-Cyp2L31 |
| f1 | SPU_016265 | Lv-PppL_268 |

| | | |
|---|---|---|
| f1 | SPU_016289 | Lv-Dom3hz |
| f1 | SPU_016310 | none |
| f1 | SPU_016367 | NA |
| f1 | SPU_016372 | Lv-Med13l |
| f1 | SPU_016485 | Lv-Hypp_131 |
| f1 | SPU_016527 | Lv-Tsp1/LamGap |
| f1 | SPU_016538 | Lv-Slc39a9 |
| f1 | SPU_016608 | Lv-Hypp_833 |
| f1 | SPU_016654 | NA |
| f1 | SPU_016689 | Lv-Seawi |
| f1 | SPU_016757 | Lv-Klhl5-2 |
| f1 | SPU_016765 | Lv-Hypp_2277 |
| f1 | SPU_016805 | Lv-Hypp_133 |
| f1 | SPU_016850 | Lv-Abcb10A |
| f1 | SPU_016869 | Lv-Kctd1_1 |
| f1 | SPU_016957 | Lv-Arsa_1 |
| f1 | SPU_017018 | Lv-Hypp_857 |
| f1 | SPU_017048 | Lv-Lmbrd2 |
| f1 | SPU_017055 | none |
| f1 | SPU_017117 | Lv-Ddx41 |
| f1 | SPU_017121 | Lv-Solh |
| f1 | SPU_017142 | Lv-Mrpl15 |
| f1 | SPU_017163 | Lv-Zdhhc7 |
| f1 | SPU_017175 | Lv-Slc10a2_9 |
| f1 | SPU_017216 | Lv-Wdr46 |
| f1 | SPU_017289 | Lv-KifC1 |
| f1 | SPU_017313 | Lv-Tmem5 |
| f1 | SPU_017385 | Lv-Cdc2l1_1 |
| f1 | SPU_017455 | Lv-Upk_1 |
| f1 | SPU_017515 | Lv-Upf2 |
| f1 | SPU_017558 | Lv-Mettl3 |
| f1 | SPU_017702 | none |
| f1 | SPU_017761 | Lv-XerdL |
| f1 | SPU_017813 | Lv-Hypp_2357 |
| f1 | SPU_017827 | Lv-G3bp2L |
| f1 | SPU_017830 | Lv-Cstf2 |
| f1 | SPU_017999 | Lv-Hypp_2365 |
| f1 | SPU_018024 | Lv-Tsen2 |

| | | |
|---|---|---|
| f1 | SPU_018036 | Lv-Endrvt35 |
| f1 | SPU_018082 | Lv-PrepL |
| f1 | SPU_018109 | NA |
| f1 | SPU_018117 | Lv-Dnajc16 |
| f1 | SPU_018119 | Lv-Tbpl1 |
| f1 | SPU_018149 | Lv-Rpusd1 |
| f1 | SPU_018174 | Lv-Cep57 |
| f1 | SPU_018187 | Lv-Mosc2 |
| f1 | SPU_018255 | Lv-Hypp_912 |
| f1 | SPU_018260 | Lv-Hypp_2376 |
| f1 | SPU_018390 | Lv-C16orf7L |
| f1 | SPU_018414 | Lv-Hypp_914 |
| f1 | SPU_018530 | Lv-Ctbp |
| f1 | SPU_018581 | Lv-Dach1 |
| f1 | SPU_018586 | NA |
| f1 | SPU_018590 | Lv-Tpr1_1 |
| f1 | SPU_018790 | Lv-Esf1 |
| f1 | SPU_018844 | none |
| f1 | SPU_018921 | none |
| f1 | SPU_019019 | none |
| f1 | SPU_019033 | Lv-Nstrn |
| f1 | SPU_019062 | Lv-Polrmt_1 |
| f1 | SPU_019072 | NA |
| f1 | SPU_019128 | Lv-Cspg4L |
| f1 | SPU_019156 | Lv-Hypp_148 |
| f1 | SPU_019254 | Lv-Pdcd11_1 |
| f1 | SPU_019383 | Lv-C12orf51L |
| f1 | SPU_019387 | Lv-Rab32 |
| f1 | SPU_019439 | Lv-Tmem57-2 |
| f1 | SPU_019461 | NA |
| f1 | SPU_019599 | Lv-Adh_Z |
| f1 | SPU_019651 | Lv-Z391 |
| f1 | SPU_019725 | Lv-Hypp_3037 |
| f1 | SPU_019769 | Lv-Prpf3_1 |
| f1 | SPU_019772 | none |
| f1 | SPU_019777 | Lv-CbpdDL_3 |
| f1 | SPU_019816 | Lv-Pih1d1 |
| f1 | SPU_019915 | Lv-Apa |

| | | |
|---|---|---|
| f1 | SPU_019921 | Lv-Rad26 |
| f1 | SPU_019959 | Lv-Z14 |
| f1 | SPU_020171 | Lv-Tll1_2 |
| f1 | SPU_020192 | Lv-LamaLf |
| f1 | SPU_020217 | Lv-Slc16a7L_1 |
| f1 | SPU_020224 | NA |
| f1 | SPU_020250 | Lv-Lipt1 |
| f1 | SPU_020272 | Lv-Hypp_969 |
| f1 | SPU_020288 | Lv-Tnpo1_2 |
| f1 | SPU_020329 | Lv-Srp54 |
| f1 | SPU_020414 | Lv-Krp170 |
| f1 | SPU_020530 | Lv-Sms |
| f1 | SPU_020562 | NA |
| f1 | SPU_020563 | Lv-Z395 |
| f1 | SPU_020773 | Lv-B3galt5_4 |
| f1 | SPU_020869 | Lv-Poldip3 |
| f1 | SPU_020880 | Lv-Tdh |
| f1 | SPU_020951 | none |
| f1 | SPU_020987 | Lv-Adss2 |
| f1 | SPU_021028 | Lv-Fbxo30 |
| f1 | SPU_021165 | Lv-LamL |
| f1 | SPU_021166 | none |
| f1 | SPU_021231 | NA |
| f1 | SPU_021237 | none |
| f1 | SPU_021461 | Lv-Polyp_5 |
| f1 | SPU_021524 | Lv-Paox_1 |
| f1 | SPU_021533 | NA |
| f1 | SPU_021537 | NA |
| f1 | SPU_021564 | Lv-Comt |
| f1 | SPU_021603 | Lv-Hypp_172 |
| f1 | SPU_021723 | Lv-Mphosph8 |
| f1 | SPU_021769 | Lv-Rt41 |
| f1 | SPU_021859 | Lv-Gscrta1s |
| f1 | SPU_021880 | Lv-Hypp_2582 |
| f1 | SPU_022029 | none |
| f1 | SPU_022194 | none |
| f1 | SPU_022236 | Lv-Hsp40B |
| f1 | SPU_022257 | Lv-Tbc1d22b |

| | | |
|---|---|---|
| f1 | SPU_022306 | none |
| f1 | SPU_022352 | none |
| f1 | SPU_022417 | Lv-Hypp_1045 |
| f1 | SPU_022446 | Lv-Sdccag1 |
| f1 | SPU_022467 | Lv-C2orf69L |
| f1 | SPU_022469 | Lv-Rnf126 |
| f1 | SPU_022544 | Lv-Cd163L |
| f1 | SPU_022556 | none |
| f1 | SPU_022623 | Lv-Pla2g1b_9 |
| f1 | SPU_022777 | none |
| f1 | SPU_022809 | Lv-Tlk |
| f1 | SPU_022812 | Lv-Mettl2a |
| f1 | SPU_022835 | Lv-Znf598 |
| f1 | SPU_022843 | NA |
| f1 | SPU_022847 | Lv-Med12L_2 |
| f1 | SPU_022856 | Lv-Rcor3 |
| f1 | SPU_022892 | Lv-Z61 |
| f1 | SPU_022894 | Lv-Hypp_2636 |
| f1 | SPU_023007 | Lv-Hypp_1065 |
| f1 | SPU_023243 | Lv-Nphs1L_4 |
| f1 | SPU_023312 | none |
| f1 | SPU_023314 | none |
| f1 | SPU_023316 | Lv-Ist1l |
| f1 | SPU_023366 | Lv-Enz1 |
| f1 | SPU_023452 | Lv-Dcun1d4 |
| f1 | SPU_023495 | none |
| f1 | SPU_023510 | Lv-Z444 |
| f1 | SPU_023600 | Lv-Hypp_1091 |
| f1 | SPU_023616 | Lv-Dnttip2 |
| f1 | SPU_023700 | Lv-Polr2j |
| f1 | SPU_023708 | Lv-EllL |
| f1 | SPU_023752 | Lv-Ftsf1l |
| f1 | SPU_023916 | none |
| f1 | SPU_023929 | Lv-Hypp_207 |
| f1 | SPU_023932 | Lv-Dap5_1 |
| f1 | SPU_023956 | Lv-Hypp_208 |
| f1 | SPU_023960 | Lv-Hypp_2699 |
| f1 | SPU_024072 | Lv-Pars2 |

| | | |
|---|---|---|
| f1 | SPU_024094 | Lv-Orc4 |
| f1 | SPU_024095 | Lv-SnRNP-100kDa_1 |
| f1 | SPU_024143 | NA |
| f1 | SPU_024158 | Lv-Cog3 |
| f1 | SPU_024364 | none |
| f1 | SPU_024495 | Lv-Khk |
| f1 | SPU_024517 | Lv-Xrcc4 |
| f1 | SPU_024551 | none |
| f1 | SPU_024555 | Lv-Fkbp8 |
| f1 | SPU_024633 | Lv-Med12L_3 |
| f1 | SPU_024691 | Lv-Dot1LL |
| f1 | SPU_024735 | none |
| f1 | SPU_024745 | Lv-Z52 |
| f1 | SPU_024802 | Lv-Mblk1 |
| f1 | SPU_024840 | Lv-Mob1ph |
| f1 | SPU_024897 | Lv-CoilL |
| f1 | SPU_024900 | Lv-Drd2 |
| f1 | SPU_024929 | Lv-Mocs3 |
| f1 | SPU_024941 | Lv-PppL_222 |
| f1 | SPU_024964 | none |
| f1 | SPU_025001 | none |
| f1 | SPU_025028 | Lv-Rrp7a |
| f1 | SPU_025029 | Lv-C7orf26L_1 |
| f1 | SPU_025040 | Lv-Htr1fL_1 |
| f1 | SPU_025063 | Lv-Prrdh_3 |
| f1 | SPU_025127 | none |
| f1 | SPU_025154 | Lv-PgbtpaseL |
| f1 | SPU_025206 | none |
| f1 | SPU_025246 | Lv-Lman2 |
| f1 | SPU_025418 | Lv-Golga2 |
| f1 | SPU_025580 | Lv-Med8 |
| f1 | SPU_025581 | Lv-Carm1 |
| f1 | SPU_025675 | none |
| f1 | SPU_025740 | Lv-RtL_7 |
| f1 | SPU_025743 | NA |
| f1 | SPU_025832 | Lv-Bptf |
| f1 | SPU_025834 | Lv-Aimp2 |
| f1 | SPU_025850 | Lv-Z408 |

| | | |
|---|---|---|
| f1 | SPU_025872 | Lv-Scyl2 |
| f1 | SPU_025923 | Lv-Alg13 |
| f1 | SPU_025981 | Lv-Ccdc112 |
| f1 | SPU_026053 | Lv-Hypp_239 |
| f1 | SPU_026115 | Lv-Hypp_2820 |
| f1 | SPU_026239 | none |
| f1 | SPU_026383 | Lv-Sdccag1-2 |
| f1 | SPU_026411 | Lv-Ppwd1 |
| f1 | SPU_026412 | Lv-Hypp_244 |
| f1 | SPU_026419 | none |
| f1 | SPU_026500 | none |
| f1 | SPU_026517 | Lv-Hypp_245 |
| f1 | SPU_026712 | Lv-Masa |
| f1 | SPU_026723 | Lv-Serrp1a |
| f1 | SPU_026733 | Lv-Hypp_259 |
| f1 | SPU_026778 | Lv-Plsc |
| f1 | SPU_026781 | Lv-Cldn12 |
| f1 | SPU_026824 | none |
| f1 | SPU_026836 | Lv-Hypp_261 |
| f1 | SPU_026845 | none |
| f1 | SPU_026867 | Lv-Vps13d_1 |
| f1 | SPU_026925 | Lv-UbiadL |
| f1 | SPU_026936 | Lv-Wdr8 |
| f1 | SPU_026959 | Lv-Pi4k2b |
| f1 | SPU_026970 | Lv-Nop14 |
| f1 | SPU_026983 | none |
| f1 | SPU_026993 | Lv-HpcaL1 |
| f1 | SPU_026999 | Lv-Lyrm1 |
| f1 | SPU_027051 | Lv-Hypp_2883 |
| f1 | SPU_027057 | Lv-Rpap1 |
| f1 | SPU_027066 | Lv-Sub1l |
| f1 | SPU_027070 | Lv-Plrg1 |
| f1 | SPU_027082 | NA |
| f1 | SPU_027109 | Lv-PolypL_90 |
| f1 | SPU_027111 | Lv-Swift |
| f1 | SPU_027176 | Lv-Fut4_8 |
| f1 | SPU_027233 | none |
| f1 | SPU_027325 | Lv-RtL_64 |

| | | |
|---|---|---|
| f1 | SPU_027356 | Lv-Cadherin_9 |
| f1 | SPU_027363 | Lv-St6L |
| f1 | SPU_027376 | NA |
| f1 | SPU_027380 | Lv-Tgfbr3 |
| f1 | SPU_027394 | Lv-Wrb |
| f1 | SPU_027461 | Lv-Supt4h1 |
| f1 | SPU_027478 | Lv-PppL_285 |
| f1 | SPU_027516 | Lv-Tex264L |
| f1 | SPU_027597 | none |
| f1 | SPU_027611 | Lv-Polg2 |
| f1 | SPU_027641 | Lv-Stxbp4 |
| f1 | SPU_027651 | Lv-Clpx |
| f1 | SPU_027656 | Lv-Nup88 |
| f1 | SPU_027666 | Lv-Ark2 |
| f1 | SPU_027672 | NA |
| f1 | SPU_027678 | NA |
| f1 | SPU_027679 | Lv-PolypL_128 |
| f1 | SPU_027716 | Lv-PppL_287 |
| f1 | SPU_027752 | NA |
| f1 | SPU_027830 | Lv-C12orf26 |
| f1 | SPU_027848 | NA |
| f1 | SPU_027918 | Lv-Z288 |
| f1 | SPU_027966 | Lv-Sf3b14b |
| f1 | SPU_027978 | none |
| f1 | SPU_027979 | none |
| f1 | SPU_027983 | none |
| f1 | SPU_028033 | Lv-Sdr_1 |
| f1 | SPU_028110 | Lv-TerfaL |
| f1 | SPU_028333 | Lv-HltfL |
| f1 | SPU_028385 | none |
| f1 | SPU_028411 | Lv-TimL |
| f1 | SPU_028449 | none |
| f1 | SPU_028545 | Lv-Zdhhc6 |
| f1 | SPU_028559 | Lv-Med16 |
| f1 | SPU_028561 | none |
| f1 | SPU_028606 | Lv-Ubl7 |
| f1 | SPU_028666 | Lv-C9orf28 |
| f1 | SPU_028759 | Lv-Aadac_4 |

| | | |
|---|---|---|
| f1 | SPU_028809 | Lv-Taf6 |
| f1 | SPU_028838 | Lv-Hypp_2972 |
| f1 | SPU_028861 | Lv-Dap5 |
| f2 | SPU_000019 | Lv-CspL_1 |
| f2 | SPU_000051 | Lv-Srcr |
| f2 | SPU_000124 | Lv-Hypp_294 |
| f2 | SPU_000176 | Lv-Reeler/Egf/Foln |
| f2 | SPU_000194 | Lv-TransposaseI |
| f2 | SPU_000210 | Lv-Pola |
| f2 | SPU_000246 | Lv-Cep72 |
| f2 | SPU_000267 | none |
| f2 | SPU_000303 | Lv-Srp19 |
| f2 | SPU_000312 | Lv-Fbxl21 |
| f2 | SPU_000417 | Lv-Slc27a1 |
| f2 | SPU_000478 | Lv-Glod4L |
| f2 | SPU_000508 | Lv-GbA |
| f2 | SPU_000639 | Lv-Mcart1 |
| f2 | SPU_000673 | Lv-Fn3/Egf |
| f2 | SPU_000862 | NA |
| f2 | SPU_000912 | Lv-Pfdn1 |
| f2 | SPU_000936 | none |
| f2 | SPU_001146 | Lv-Med6 |
| f2 | SPU_001184 | none |
| f2 | SPU_001198 | Lv-Seta |
| f2 | SPU_001208 | Lv-Herc2 |
| f2 | SPU_001256 | none |
| f2 | SPU_001277 | Lv-Fancd2 |
| f2 | SPU_001340 | Lv-Unc44_197 |
| f2 | SPU_001352 | Lv-Ggnbp2 |
| f2 | SPU_001368 | none |
| f2 | SPU_001442 | Lv-C22orf9L |
| f2 | SPU_001533 | Lv-FibA_1 |
| f2 | SPU_001567 | Lv-Oprk1L_3 |
| f2 | SPU_001568 | none |
| f2 | SPU_001569 | Lv-Fk506bp12 |
| f2 | SPU_001644 | Lv-Hypp_1340 |
| f2 | SPU_001645 | Lv-Dixdc1L |
| f2 | SPU_001679 | none |

| | | |
|---|---|---|
| f2 | SPU_001788 | Lv-Ppp1cb |
| f2 | SPU_001962 | Lv-Rnf207 |
| f2 | SPU_002024 | NA |
| f2 | SPU_002083 | Lv-C7orf27L |
| f2 | SPU_002112 | Lv-Fn3/Egf_11 |
| f2 | SPU_002125 | Lv-Hypp_344 |
| f2 | SPU_002181 | Lv-Setd8l-3 |
| f2 | SPU_002241 | Lv-Megf8 |
| f2 | SPU_002267 | NA |
| f2 | SPU_002296 | Lv-Hypp_1387 |
| f2 | SPU_002329 | Lv-Ercc6a |
| f2 | SPU_002335 | Lv-Tm9sf1 |
| f2 | SPU_002363 | Lv-Whirl |
| f2 | SPU_002398 | Lv-Gpi_pigU |
| f2 | SPU_002428 | Lv-Mb_2 |
| f2 | SPU_002477 | none |
| f2 | SPU_002498 | Lv-Ttc28 |
| f2 | SPU_002507 | Lv-Rbj |
| f2 | SPU_002613 | Lv-Vps45A |
| f2 | SPU_002648 | Lv-Appl1 |
| f2 | SPU_002723 | Lv-My6 |
| f2 | SPU_002825 | Lv-Tbc1d23 |
| f2 | SPU_002826 | Lv-Hypp_1414 |
| f2 | SPU_002885 | Lv-Cnih |
| f2 | SPU_002889 | Lv-Mdm1 |
| f2 | SPU_002899 | Lv-Cyp2Lf |
| f2 | SPU_002933 | Lv-Cul1 |
| f2 | SPU_002962 | Lv-Nlr66 |
| f2 | SPU_002984 | Lv-Wdr27 |
| f2 | SPU_003027 | Lv-PolppL_20 |
| f2 | SPU_003033 | Lv-Slc25A17 |
| f2 | SPU_003059 | Lv-Zc3p26_4 |
| f2 | SPU_003060 | Lv-Zc3p26_5 |
| f2 | SPU_003109 | Lv-Ppih |
| f2 | SPU_003132 | Lv-Rap1gds1 |
| f2 | SPU_003142 | Lv-Pten_1 |
| f2 | SPU_003181 | Lv-Plekha1 |
| f2 | SPU_003185 | Lv-Hypp_376 |

| | | |
|---|---|---|
| f2 | SPU_003319 | Lv-PolppL_27 |
| f2 | SPU_003322 | Lv-PolppL_30 |
| f2 | SPU_003379 | Lv-Hypp_1448 |
| f2 | SPU_003380 | Lv-Hsd3b |
| f2 | SPU_003410 | Lv-Kiaa0226L |
| f2 | SPU_003446 | none |
| f2 | SPU_003457 | Lv-Mpp5 |
| f2 | SPU_003497 | Lv-Zmynd10 |
| f2 | SPU_003509 | Lv-Ncor2 |
| f2 | SPU_003529 | Lv-PolppL_34 |
| f2 | SPU_003584 | none |
| f2 | SPU_003595 | Lv-Cutl |
| f2 | SPU_003651 | Lv-Pold |
| f2 | SPU_003652 | Lv-Fam49b |
| f2 | SPU_003717 | Lv-Kif6L |
| f2 | SPU_003753 | Lv-Hypp_1480 |
| f2 | SPU_003761 | Lv-Hypp_3096 |
| f2 | SPU_003805 | Lv-Hebp2 |
| f2 | SPU_003834 | Lv-Unc44L_33 |
| f2 | SPU_003845 | Lv-Rabep1 |
| f2 | SPU_003969 | none |
| f2 | SPU_004010 | none |
| f2 | SPU_004079 | Lv-Unc45h |
| f2 | SPU_004097 | Lv-Rapgef2 |
| f2 | SPU_004129 | Lv-Hypp_1501 |
| f2 | SPU_004132 | Lv-Kiaa0556 |
| f2 | SPU_004188 | Lv-Disph1 |
| f2 | SPU_004193 | NA |
| f2 | SPU_004255 | Lv-Zw10 |
| f2 | SPU_004284 | none |
| f2 | SPU_004306 | Lv-Hypp_411 |
| f2 | SPU_004424 | Lv-Fam91a1 |
| f2 | SPU_004444 | Lv-Mals |
| f2 | SPU_004525 | Lv-Mbip |
| f2 | SPU_004554 | NA |
| f2 | SPU_004639 | Lv-Clk2 |
| f2 | SPU_004719 | Lv-Ttc7b |
| f2 | SPU_004776 | Lv-Hmg20a |

| | | |
|---|---|---|
| f2 | SPU_004789 | Lv-Ccrk |
| f2 | SPU_004809 | Lv-Pi4k |
| f2 | SPU_004811 | Lv-PolppL_39 |
| f2 | SPU_004812 | none |
| f2 | SPU_004839 | NA |
| f2 | SPU_004889 | Lv-C19orf62L |
| f2 | SPU_004900 | Lv-Hypp_1550 |
| f2 | SPU_005009 | Lv-Fbxw7 |
| f2 | SPU_005029 | Lv-Fam21a |
| f2 | SPU_005109 | Lv-Hypp_1566 |
| f2 | SPU_005134 | Lv-Prom1L |
| f2 | SPU_005149 | Lv-Gprk3 |
| f2 | SPU_005273 | none |
| f2 | SPU_005309 | Lv-Lrrc9 |
| f2 | SPU_005328 | Lv-Hypp_443 |
| f2 | SPU_005361 | Lv-Lcmt2 |
| f2 | SPU_005588 | NA |
| f2 | SPU_005597 | Lv-Stub1_1 |
| f2 | SPU_005598 | Lv-Nfrkbl |
| f2 | SPU_005618 | NA |
| f2 | SPU_005708 | Lv-Ectdrn |
| f2 | SPU_005740 | Lv-Sorl1 |
| f2 | SPU_005793 | Lv-Hypp_1602 |
| f2 | SPU_005964 | Lv-Rnf10 |
| f2 | SPU_006028 | none |
| f2 | SPU_006069 | Lv-Cdk10 |
| f2 | SPU_006082 | Lv-Bbs2_1 |
| f2 | SPU_006169 | Lv-Brms1L |
| f2 | SPU_006238 | NA |
| f2 | SPU_006396 | Lv-Bmp5-8 |
| f2 | SPU_006405 | Lv-Ube3c |
| f2 | SPU_006560 | none |
| f2 | SPU_006585 | Lv-Man2b1 |
| f2 | SPU_006670 | Lv-Lin37l |
| f2 | SPU_006748 | Lv-Lrrc28 |
| f2 | SPU_006851 | Lv-SaaL1 |
| f2 | SPU_006883 | none |
| f2 | SPU_006890 | Lv-Galm |

| | | |
|---|---|---|
| f2 | SPU_006901 | Lv-Tkl1a |
| f2 | SPU_006995 | none |
| f2 | SPU_007058 | Lv-Wdr61 |
| f2 | SPU_007120 | Lv-Ubap1L |
| f2 | SPU_007169 | Lv-Z219 |
| f2 | SPU_007173 | Lv-C17orf71 |
| f2 | SPU_007180 | Lv-Pnn |
| f2 | SPU_007185 | Lv-Cdk5rap1 |
| f2 | SPU_007220 | Lv-Hypp_1707 |
| f2 | SPU_007248 | Lv-My10_putative_homolog |
| f2 | SPU_007357 | Lv-Abcb10B |
| f2 | SPU_007368 | none |
| f2 | SPU_007485 | Lv-Gs |
| f2 | SPU_007550 | Lv-Pef1_1 |
| f2 | SPU_007564 | Lv-Elmod2 |
| f2 | SPU_007568 | Lv-Spry1 |
| f2 | SPU_007658 | Lv-Hormad1 |
| f2 | SPU_007659 | Lv-Kctd1L_1 |
| f2 | SPU_007663 | Lv-Hypp_1742 |
| f2 | SPU_007862 | Lv-Cblr1 |
| f2 | SPU_007890 | Lv-Hypp_530 |
| f2 | SPU_007903 | NA |
| f2 | SPU_007944 | Lv-Ppil4 |
| f2 | SPU_007957 | Lv-Z370 |
| f2 | SPU_007981 | Lv-Lmpt |
| f2 | SPU_008018 | Lv-Apc1_1 |
| f2 | SPU_008031 | Lv-Hypp_1762 |
| f2 | SPU_008065 | Lv-Clect/Egf_10 |
| f2 | SPU_008172 | Lv-Dazap |
| f2 | SPU_008286 | Lv-Mgst2 |
| f2 | SPU_008290 | Lv-Tud |
| f2 | SPU_008328 | Lv-PolppL_71 |
| f2 | SPU_008331 | Lv-Paxillin |
| f2 | SPU_008360 | NA |
| f2 | SPU_008407 | Lv-Denr |
| f2 | SPU_008427 | none |
| f2 | SPU_008454 | Lv-Ggcx |
| f2 | SPU_008458 | Lv-Manba |

| | | |
|---|---|---|
| f2 | SPU_008466 | Lv-Ptprorph2 |
| f2 | SPU_008538 | Lv-Lhfpl2 |
| f2 | SPU_008586 | NA |
| f2 | SPU_008611 | NA |
| f2 | SPU_008621 | Lv-Ulk4 |
| f2 | SPU_008694 | Lv-Pcyt1b |
| f2 | SPU_008697 | none |
| f2 | SPU_008811 | Lv-Hypp_561 |
| f2 | SPU_008827 | Lv-Galnt7_1 |
| f2 | SPU_008845 | Lv-Mlx/Ip |
| f2 | SPU_008851 | none |
| f2 | SPU_008896 | none |
| f2 | SPU_008897 | Lv-Ddef1 |
| f2 | SPU_008900 | Lv-Hypp_1810 |
| f2 | SPU_008909 | none |
| f2 | SPU_008960 | Lv-Ttl |
| f2 | SPU_009013 | Lv-Otud5 |
| f2 | SPU_009027 | Lv-Rab35 |
| f2 | SPU_009039 | Lv-Fn3/Igf_20 |
| f2 | SPU_009125 | Lv-Tmub2 |
| f2 | SPU_009130 | Lv-Endogl1 |
| f2 | SPU_009134 | Lv-Mras |
| f2 | SPU_009163 | none |
| f2 | SPU_009168 | Lv-Socax |
| f2 | SPU_009228 | Lv-Nphp3_1 |
| f2 | SPU_009249 | Lv-Ppt1 |
| f2 | SPU_009251 | none |
| f2 | SPU_009252 | Lv-Lec7a6_9 |
| f2 | SPU_009322 | none |
| f2 | SPU_009357 | Lv-tauD |
| f2 | SPU_009376 | none |
| f2 | SPU_009418 | none |
| f2 | SPU_009470 | Lv-Lats |
| f2 | SPU_009517 | Lv-CopineB |
| f2 | SPU_009521 | none |
| f2 | SPU_009524 | Lv-PolppL_82 |
| f2 | SPU_009722 | Lv-Banp |
| f2 | SPU_009736 | Lv-Zir2 |

| | | |
|---|---|---|
| f2 | SPU_009772 | Lv-Sars2 |
| f2 | SPU_009835 | none |
| f2 | SPU_009839 | Lv-Hypp_1875 |
| f2 | SPU_009892 | Lv-Rabepkl |
| f2 | SPU_009902 | Lv-Hypp_58 |
| f2 | SPU_009962 | none |
| f2 | SPU_010016 | Lv-Unc44_220 |
| f2 | SPU_010025 | Lv-Endrvt71 |
| f2 | SPU_010125 | Lv-Fn3/Igf_2 |
| f2 | SPU_010196 | Lv-Gpr64L_5 |
| f2 | SPU_010209 | Lv-Mthfd2 |
| f2 | SPU_010219 | Lv-Slc46a3L |
| f2 | SPU_010302 | Lv-Acyp1 |
| f2 | SPU_010315 | Lv-Rnf160 |
| f2 | SPU_010336 | Lv-Myo9B |
| f2 | SPU_010363 | Lv-Ptcd1L |
| f2 | SPU_010506 | none |
| f2 | SPU_010530 | Lv-Ttk |
| f2 | SPU_010553 | none |
| f2 | SPU_010641 | Lv-Bax |
| f2 | SPU_010840 | Lv-G-cadL1 |
| f2 | SPU_010906 | Lv-Chtf18 |
| f2 | SPU_011060 | Lv-Tpase4 |
| f2 | SPU_011135 | Lv-PolppL_91 |
| f2 | SPU_011140 | none |
| f2 | SPU_011154 | Lv-PnpLa7 |
| f2 | SPU_011190 | Lv-Stam |
| f2 | SPU_011218 | none |
| f2 | SPU_011267 | Lv-AtminL |
| f2 | SPU_011295 | Lv-Cycl |
| f2 | SPU_011301 | Lv-Rnpc2 |
| f2 | SPU_011306 | Lv-Keap1 |
| f2 | SPU_011311 | none |
| f2 | SPU_011368 | none |
| f2 | SPU_011460 | none |
| f2 | SPU_011559 | Lv-Hypp_650 |
| f2 | SPU_011576 | Lv-Nr1m2 |
| f2 | SPU_011610 | Lv-Hypp_653 |

| | | |
|---|---|---|
| f2 | SPU_011641 | Lv-Ef2k |
| f2 | SPU_011707 | none |
| f2 | SPU_011753 | Lv-Uba3 |
| f2 | SPU_011822 | Lv-Cophs7b |
| f2 | SPU_011847 | Lv-C3orf60L |
| f2 | SPU_011854 | NA |
| f2 | SPU_011877 | Lv-Ahi |
| f2 | SPU_011891 | NA |
| f2 | SPU_011896 | Lv-Kin |
| f2 | SPU_011919 | none |
| f2 | SPU_011986 | Lv-Phf21A |
| f2 | SPU_011997 | Lv-Wdr79 |
| f2 | SPU_012081 | Lv-Cyp4L3 |
| f2 | SPU_012085 | Lv-Rabl3_1 |
| f2 | SPU_012098 | Lv-Map2k4 |
| f2 | SPU_012233 | Lv-Egfib_6 |
| f2 | SPU_012370 | Lv-Hypp_2029 |
| f2 | SPU_012385 | Lv-Ttrspn_10 |
| f2 | SPU_012441 | Lv-Pdrg1 |
| f2 | SPU_012482 | Lv-Taf1l_1 |
| f2 | SPU_012581 | Lv-SetxL |
| f2 | SPU_012632 | Lv-Z417 |
| f2 | SPU_012642 | Lv-Aldh3b1 |
| f2 | SPU_012729 | Lv-Hypp_2047 |
| f2 | SPU_012751 | none |
| f2 | SPU_012808 | Lv-Dhrs11L |
| f2 | SPU_012827 | none |
| f2 | SPU_012866 | Lv-Wdr24 |
| f2 | SPU_012919 | none |
| f2 | SPU_012946 | Lv-Thumpd1 |
| f2 | SPU_012977 | Lv-PlexB1 |
| f2 | SPU_013098 | NA |
| f2 | SPU_013101 | Lv-Tmem168 |
| f2 | SPU_013104 | Lv-Glt |
| f2 | SPU_013128 | Lv-Fbxw9 |
| f2 | SPU_013201 | Lv-Dhx36_1 |
| f2 | SPU_013304 | Lv-Unc44_229 |
| f2 | SPU_013337 | Lv-Rab34 |

| | | |
|---|---|---|
| f2 | SPU_013370 | Lv-Hypp_713 |
| f2 | SPU_013413 | Lv-Pcbd2 |
| f2 | SPU_013441 | Lv-Pafah2_1 |
| f2 | SPU_013488 | Lv-Myo18a_1 |
| f2 | SPU_013503 | Lv-Htr7L_1 |
| f2 | SPU_013517 | Lv-Rgs9bp |
| f2 | SPU_013610 | Lv-C7orf26L |
| f2 | SPU_013619 | Lv-Nlr5 |
| f2 | SPU_013655 | Lv-Gtf3c3 |
| f2 | SPU_013702 | Lv-PolppL_99 |
| f2 | SPU_013818 | Lv-Kiaa0406L |
| f2 | SPU_013831 | NA |
| f2 | SPU_013833 | Lv-Plekhj1 |
| f2 | SPU_013928 | NA |
| f2 | SPU_014071 | Lv-Chn2l |
| f2 | SPU_014251 | Lv-Mtr |
| f2 | SPU_014299 | NA |
| f2 | SPU_014305 | Lv-Kiaa0082 |
| f2 | SPU_014324 | Lv-Nupb2 |
| f2 | SPU_014386 | Lv-Hypp_2145 |
| f2 | SPU_014444 | Lv-pABCt_ |
| f2 | SPU_014507 | Lv-Itpa |
| f2 | SPU_014576 | Lv-Pea |
| f2 | SPU_014591 | none |
| f2 | SPU_014670 | Lv-Synj1 |
| f2 | SPU_014672 | Lv-Clect/Cub |
| f2 | SPU_014684 | Lv-Z94 |
| f2 | SPU_014751 | none |
| f2 | SPU_014759 | Lv-Fn3/Igf_37 |
| f2 | SPU_014763 | Lv-Sara |
| f2 | SPU_014774 | Lv-Mier3 |
| f2 | SPU_014790 | Lv-Ppp1r7 |
| f2 | SPU_014811 | Lv-Copb2L |
| f2 | SPU_014837 | Lv-Hypp_766 |
| f2 | SPU_014850 | Lv-Fn3f_54 |
| f2 | SPU_014896 | Lv-RabO5 |
| f2 | SPU_014902 | Lv-Rnf157 |
| f2 | SPU_014918 | none |

| | | |
|---|---|---|
| f2 | SPU_014938 | Lv-Gba_1 |
| f2 | SPU_015018 | Lv-Hypp_106 |
| f2 | SPU_015135 | Lv-Pex19L |
| f2 | SPU_015217 | Lv-Hypp-2198 |
| f2 | SPU_015311 | Lv-Nktr |
| f2 | SPU_015328 | none |
| f2 | SPU_015376 | none |
| f2 | SPU_015392 | Lv-Hypp_782 |
| f2 | SPU_015394 | none |
| f2 | SPU_015428 | none |
| f2 | SPU_015454 | none |
| f2 | SPU_015494 | Lv-Ccny |
| f2 | SPU_015507 | Lv-Txnrd2 |
| f2 | SPU_015662 | none |
| f2 | SPU_015671 | Lv-Dip2hc |
| f2 | SPU_015724 | Lv-Vac14 |
| f2 | SPU_015855 | Lv-Fut3L |
| f2 | SPU_015954 | Lv-Hypp_804 |
| f2 | SPU_015955 | none |
| f2 | SPU_015966 | Lv-Slc28a3_1 |
| f2 | SPU_016009 | Lv-BetaLi |
| f2 | SPU_016012 | Lv-Cytb |
| f2 | SPU_016028 | Lv-Mil1 |
| f2 | SPU_016035 | Lv-Smg7 |
| f2 | SPU_016053 | Lv-Ptprlec4 |
| f2 | SPU_016136 | Lv-March5 |
| f2 | SPU_016219 | Lv-C10orf137L |
| f2 | SPU_016286 | Lv-Nudc3 |
| f2 | SPU_016339 | Lv-Slc17 |
| f2 | SPU_016416 | Lv-Hypp_2260 |
| f2 | SPU_016466 | Lv-Hypp_824 |
| f2 | SPU_016471 | Lv-PolppL_108 |
| f2 | SPU_016473 | Lv-Hypp_130 |
| f2 | SPU_016580 | Lv-PolypL_10 |
| f2 | SPU_016635 | none |
| f2 | SPU_016763 | Lv-CkIIbi1 |
| f2 | SPU_016770 | none |
| f2 | SPU_016780 | Lv-Endrvt93 |

| | | |
|---|---|---|
| f2 | SPU_016828 | Lv-Tigd4 |
| f2 | SPU_016857 | Lv-Rt38 |
| f2 | SPU_016877 | Lv-Prpf40a |
| f2 | SPU_016893 | Lv-C10orf42 |
| f2 | SPU_016909 | NA |
| f2 | SPU_016953 | Lv-FancaL |
| f2 | SPU_016956 | Lv-Vps37hb |
| f2 | SPU_016982 | Lv-PppL_113 |
| f2 | SPU_017081 | none |
| f2 | SPU_017105 | Lv-Slc34a2_1 |
| f2 | SPU_017204 | Lv-Hypp_3046 |
| f2 | SPU_017272 | Lv-Hypp_870 |
| f2 | SPU_017320 | Lv-Ccp |
| f2 | SPU_017428 | none |
| f2 | SPU_017436 | Lv-Mks1L |
| f2 | SPU_017439 | Lv-Rad51L3 |
| f2 | SPU_017451 | Lv-Ptcd1 |
| f2 | SPU_017507 | Lv-Fam40a |
| f2 | SPU_017531 | Lv-PolypL_114 |
| f2 | SPU_017539 | Lv-Hypp_881 |
| f2 | SPU_017541 | Lv-DeadbpuknI |
| f2 | SPU_017542 | Lv-Setd4 |
| f2 | SPU_017569 | Lv-Fcmd |
| f2 | SPU_017649 | Lv-Hgf |
| f2 | SPU_017715 | Lv-Tmem33 |
| f2 | SPU_017716 | Lv-Hypp_892 |
| f2 | SPU_017728 | Lv-Wdr90-3 |
| f2 | SPU_017846 | Lv-Z313 |
| f2 | SPU_017886 | Lv-Vps15 |
| f2 | SPU_017904 | Lv-Nf1 |
| f2 | SPU_017910 | Lv-Ciaoh1 |
| f2 | SPU_017923 | Lv-C10orf58L |
| f2 | SPU_017948 | Lv-Lars2 |
| f2 | SPU_017972 | Lv-RalA |
| f2 | SPU_017973 | none |
| f2 | SPU_018020 | Lv-C6orf70 |
| f2 | SPU_018021 | Lv-Park2 |
| f2 | SPU_018085 | Lv-Pck2 |

| | | |
|---|---|---|
| f2 | SPU_018132 | Lv-Fdxr |
| f2 | SPU_018209 | Lv-Hmha1 |
| f2 | SPU_018239 | none |
| f2 | SPU_018284 | Lv-Hypp_2379 |
| f2 | SPU_018300 | Lv-Cep135 |
| f2 | SPU_018344 | none |
| f2 | SPU_018395 | NA |
| f2 | SPU_018413 | Lv-Lrr_2 |
| f2 | SPU_018418 | Lv-Pxk |
| f2 | SPU_018488 | Lv-Kiaa0564pL_1 |
| f2 | SPU_018515 | NA |
| f2 | SPU_018577 | none |
| f2 | SPU_018643 | Lv-L2hgdh |
| f2 | SPU_018678 | none |
| f2 | SPU_018933 | Lv-Sipa1l1l |
| f2 | SPU_018971 | Lv-Lrrc34 |
| f2 | SPU_018991 | Lv-Cyld |
| f2 | SPU_019090 | Lv-Taz_1 |
| f2 | SPU_019142 | none |
| f2 | SPU_019207 | none |
| f2 | SPU_019256 | Lv-Phip_1 |
| f2 | SPU_019274 | none |
| f2 | SPU_019399 | none |
| f2 | SPU_019513 | NA |
| f2 | SPU_019526 | NA |
| f2 | SPU_019532 | Lv-Kiaa1530L |
| f2 | SPU_019583 | NA |
| f2 | SPU_019604 | none |
| f2 | SPU_019608 | Lv-Hypp_2455 |
| f2 | SPU_019621 | Lv-Unc44_241 |
| f2 | SPU_019626 | none |
| f2 | SPU_019633 | none |
| f2 | SPU_019650 | Lv-Pigq |
| f2 | SPU_019662 | Lv-Hypp_2458 |
| f2 | SPU_019750 | Lv-Hypp_2463 |
| f2 | SPU_019771 | Lv-Dut |
| f2 | SPU_019782 | Lv-Cys_rich_fgfr_3 |
| f2 | SPU_019844 | Lv-Mudeng |

| | | |
|---|---|---|
| f2 | SPU_019856 | Lv-Snx26 |
| f2 | SPU_019910 | Lv-Tpase7 |
| f2 | SPU_019923 | Lv-C3orf20L |
| f2 | SPU_019963 | Lv-Sh3pxd2a |
| f2 | SPU_019966 | Lv-Slc16a14L_6 |
| f2 | SPU_020050 | Lv-PtprD/F/S |
| f2 | SPU_020095 | none |
| f2 | SPU_020111 | Lv-Atp9b-2 |
| f2 | SPU_020293 | none |
| f2 | SPU_020306 | Lv-Cnot4L |
| f2 | SPU_020347 | none |
| f2 | SPU_020365 | Lv-CrumbsL2 |
| f2 | SPU_020367 | Lv-C6orf134 |
| f2 | SPU_020429 | Lv-Rab28 |
| f2 | SPU_020435 | Lv-Hk1 |
| f2 | SPU_020443 | Lv-Cecr1L |
| f2 | SPU_020541 | Lv-Creg1 |
| f2 | SPU_020636 | Lv-CopineD |
| f2 | SPU_020675 | none |
| f2 | SPU_020698 | Lv-Taf5L |
| f2 | SPU_020768 | Lv-Hypp_990 |
| f2 | SPU_020847 | Lv-Slsp_13 |
| f2 | SPU_021025 | Lv-Endrvt6 |
| f2 | SPU_021106 | Lv-PolypL_116 |
| f2 | SPU_021143 | Lv-Dusp1 |
| f2 | SPU_021233 | Lv-GbL |
| f2 | SPU_021235 | Lv-ColL |
| f2 | SPU_021274 | Lv-PkdL |
| f2 | SPU_021280 | Lv-Uba2 |
| f2 | SPU_021300 | Lv-Hypp_2544 |
| f2 | SPU_021327 | Lv-Cntn5L |
| f2 | SPU_021342 | none |
| f2 | SPU_021367 | none |
| f2 | SPU_021386 | NA |
| f2 | SPU_021448 | Lv-Vps13d |
| f2 | SPU_021532 | Lv-Tmem128 |
| f2 | SPU_021619 | Lv-PppL_129 |
| f2 | SPU_021689 | Lv-Eif2Ba |

| | | |
|---|---|---|
| f2 | SPU_021737 | Lv-Hvcn1 |
| f2 | SPU_021874 | Lv-Dennd1a |
| f2 | SPU_021879 | Lv-Nucb2 |
| f2 | SPU_021928 | Lv-Taf9 |
| f2 | SPU_022153 | Lv-Ck1a1 |
| f2 | SPU_022160 | Lv-Kif18 |
| f2 | SPU_022195 | Lv-Smicl |
| f2 | SPU_022245 | none |
| f2 | SPU_022259 | Lv-E2 |
| f2 | SPU_022262 | Lv-Kiaa0391L |
| f2 | SPU_022295 | Lv-Rb1cc |
| f2 | SPU_022388 | Lv-Unc44_160 |
| f2 | SPU_022414 | Lv-Mocs1_2 |
| f2 | SPU_022426 | none |
| f2 | SPU_022435 | Lv-Tsc22d2 |
| f2 | SPU_022513 | Lv-Dhx40_1 |
| f2 | SPU_022531 | NA |
| f2 | SPU_022534 | Lv-Deth1 |
| f2 | SPU_022550 | Lv-Hypp_2617 |
| f2 | SPU_022627 | Lv-Slc38a3 |
| f2 | SPU_022632 | Lv-Srgap3 |
| f2 | SPU_022633 | Lv-Agrin_1 |
| f2 | SPU_022647 | Lv-Igf_7 |
| f2 | SPU_022652 | Lv-Agpat1_2 |
| f2 | SPU_022658 | Lv-Chst12_1 |
| f2 | SPU_022679 | Lv-Ttc23 |
| f2 | SPU_022702 | Lv-PppL_219 |
| f2 | SPU_022740 | Lv-Hypp_2629 |
| f2 | SPU_022771 | Lv-Lpp |
| f2 | SPU_022794 | Lv-Slain2 |
| f2 | SPU_022845 | Lv-Hyi |
| f2 | SPU_023010 | Lv-Pola1 |
| f2 | SPU_023091 | Lv-Pric2 |
| f2 | SPU_023126 | Lv-Cenpe |
| f2 | SPU_023144 | Lv-Abca1_3 |
| f2 | SPU_023195 | none |
| f2 | SPU_023224 | Lv-Nme6 |
| f2 | SPU_023240 | Lv-Endrvt46 |

| | | |
|---|---|---|
| f2 | SPU_023324 | Lv-NotchL_94 |
| f2 | SPU_023343 | Lv-Ccnr1 |
| f2 | SPU_023378 | Lv-RtL_59 |
| f2 | SPU_023422 | Lv-Mobkl3 |
| f2 | SPU_023443 | Lv-Unk_44 |
| f2 | SPU_023464 | Lv-RtL_60 |
| f2 | SPU_023586 | Lv-Unc44_952 |
| f2 | SPU_023605 | Lv-Ift122Ap |
| f2 | SPU_023701 | Lv-Gpr89 |
| f2 | SPU_023729 | Lv-Smc6 |
| f2 | SPU_023768 | Lv-Dtnbp1 |
| f2 | SPU_023795 | Lv-Serac1 |
| f2 | SPU_023922 | none |
| f2 | SPU_023977 | Lv-Hypp_2701 |
| f2 | SPU_024005 | Lv-Ccdc57_1 |
| f2 | SPU_024034 | Lv-Cbara1 |
| f2 | SPU_024124 | Lv-Hypp_1107 |
| f2 | SPU_024142 | Lv-Ankrd27 |
| f2 | SPU_024323 | Lv-Map2Lph |
| f2 | SPU_024345 | Lv-Rptor |
| f2 | SPU_024409 | none |
| f2 | SPU_024418 | Lv-Fam120bL |
| f2 | SPU_024419 | Lv-Bbs6 |
| f2 | SPU_024468 | none |
| f2 | SPU_024506 | none |
| f2 | SPU_024524 | NA |
| f2 | SPU_024546 | Lv-Smc2 |
| f2 | SPU_024574 | Lv-Lrrc29 |
| f2 | SPU_024580 | none |
| f2 | SPU_024627 | Lv-Leng1L |
| f2 | SPU_024667 | Lv-Dnajc25 |
| f2 | SPU_024677 | none |
| f2 | SPU_024684 | Lv-Hypp_1122 |
| f2 | SPU_024718 | Lv-Ccdc61 |
| f2 | SPU_024795 | Lv-Hypp_2753 |
| f2 | SPU_024846 | Lv-St8sia2_4 |
| f2 | SPU_024919 | none |
| f2 | SPU_024945 | Lv-PppL_140 |

| | | |
|---|---|---|
| f2 | SPU_025010 | Lv-FoxK |
| f2 | SPU_025015 | Lv-Hmbs |
| f2 | SPU_025108 | Lv-Hypp_2773 |
| f2 | SPU_025112 | none |
| f2 | SPU_025126 | Lv-Large_1 |
| f2 | SPU_025265 | Lv-Polyp_8 |
| f2 | SPU_025310 | Lv-Echn12 |
| f2 | SPU_025415 | Lv-Hypp_1146 |
| f2 | SPU_025488 | Lv-Iqcb1 |
| f2 | SPU_025506 | none |
| f2 | SPU_025587 | Lv-Hypp_1155 |
| f2 | SPU_025599 | none |
| f2 | SPU_025697 | Lv-Dera |
| f2 | SPU_025717 | none |
| f2 | SPU_025754 | Lv-Aaas |
| f2 | SPU_025758 | Lv-Coasy |
| f2 | SPU_025779 | Lv-Slc28a2 |
| f2 | SPU_025840 | Lv-Snx14 |
| f2 | SPU_026018 | Lv-Hypp_238 |
| f2 | SPU_026024 | Lv-Hypp_1165 |
| f2 | SPU_026030 | Lv-Megf8-2 |
| f2 | SPU_026097 | Lv-C2cd3_2 |
| f2 | SPU_026102 | Lv-Unc44_347 |
| f2 | SPU_026107 | none |
| f2 | SPU_026133 | Lv-Hypp_1169 |
| f2 | SPU_026297 | Lv-Vps39 |
| f2 | SPU_026303 | Lv-Lamb1_3 |
| f2 | SPU_026337 | none |
| f2 | SPU_026415 | Lv-Tbrg1L |
| f2 | SPU_026456 | Lv-Smg5 |
| f2 | SPU_026468 | Lv-Hypp_2845 |
| f2 | SPU_026478 | Lv-Hypp_2846 |
| f2 | SPU_026488 | Lv-HesBL |
| f2 | SPU_026556 | Lv-Atp9b |
| f2 | SPU_026557 | Lv-Hypp_1182 |
| f2 | SPU_026562 | Lv-Inf2L |
| f2 | SPU_026575 | Lv-Hypp_2854 |
| f2 | SPU_026633 | Lv-Dmtf |

| | | |
|---|---|---|
| f2 | SPU_026715 | NA |
| f2 | SPU_026726 | Lv-Scnm1L |
| f2 | SPU_026731 | Lv-Ddr1 |
| f2 | SPU_026772 | Lv-Pib5pa |
| f2 | SPU_026780 | Lv-NgrnL |
| f2 | SPU_026787 | none |
| f2 | SPU_026826 | Lv-Cep63 |
| f2 | SPU_026907 | Lv-Tjp1 |
| f2 | SPU_026952 | Lv-Kiaa1468L |
| f2 | SPU_026957 | Lv-Rnaseh2a |
| f2 | SPU_026975 | none |
| f2 | SPU_026987 | Lv-Ppp2AdirsB56 |
| f2 | SPU_027006 | NA |
| f2 | SPU_027068 | Lv-Dgcr6ll_1 |
| f2 | SPU_027101 | Lv-Ptp69Dl |
| f2 | SPU_027110 | none |
| f2 | SPU_027146 | Lv-Hypp_3006 |
| f2 | SPU_027238 | Lv-Acot8_1 |
| f2 | SPU_027266 | Lv-PppL_158 |
| f2 | SPU_027305 | NA |
| f2 | SPU_027382 | Lv-Alox5_2 |
| f2 | SPU_027387 | Lv-Camsap1L |
| f2 | SPU_027401 | NA |
| f2 | SPU_027404 | Lv-Brf1 |
| f2 | SPU_027418 | Lv-Pomgnt1L |
| f2 | SPU_027425 | none |
| f2 | SPU_027427 | Lv-Hypp_2902 |
| f2 | SPU_027435 | Lv-Gpr175 |
| f2 | SPU_027437 | Lv-B3gnt2L |
| f2 | SPU_027485 | Lv-PppL_286 |
| f2 | SPU_027519 | none |
| f2 | SPU_027602 | Lv-PppL_161 |
| f2 | SPU_027603 | Lv-Gmfb |
| f2 | SPU_027654 | Lv-Fuzzy2 |
| f2 | SPU_027665 | Lv-PolypL_126 |
| f2 | SPU_027788 | Lv-Hypp_2919 |
| f2 | SPU_027875 | Lv-Brp16 |
| f2 | SPU_027919 | Lv-Z344 |

| | | |
|---|---|---|
| f2 | SPU_027936 | Lv-Lyrm5 |
| f2 | SPU_028030 | Lv-Furin_1 |
| f2 | SPU_028039 | NA |
| f2 | SPU_028124 | Lv-Dusp6 |
| f2 | SPU_028144 | Lv-Grina |
| f2 | SPU_028174 | Lv-Unk_18 |
| f2 | SPU_028228 | none |
| f2 | SPU_028234 | Lv-Tatdn1 |
| f2 | SPU_028248 | Lv-Hypp_2943 |
| f2 | SPU_028262 | Lv-Thred_1 |
| f2 | SPU_028284 | Lv-Ddx19a |
| f2 | SPU_028348 | Lv-Nek7 |
| f2 | SPU_028383 | Lv-PppL_166 |
| f2 | SPU_028437 | none |
| f2 | SPU_028458 | Lv-Mgea5 |
| f2 | SPU_028529 | none |
| f2 | SPU_028543 | none |
| f2 | SPU_028546 | Lv-Hypp_1232 |
| f2 | SPU_028601 | Lv-Indol |
| f2 | SPU_028609 | Lv-Ccdc99 |
| f2 | SPU_028647 | none |
| f2 | SPU_028663 | Lv-Hypp_2962 |
| f2 | SPU_028713 | Lv-Hypp_291 |
| f2 | SPU_028826 | Lv-Kif1aL |
| f2 | SPU_028832 | Lv-TpaseL_23 |
| f2 | SPU_028888 | Lv-Brwd1 |
| f2 | SPU_030229 | Lv-Axndhc8h |
| f3 | SPU_000034 | Lv-Fhad1L |
| f3 | SPU_000045 | Lv-Fic1 |
| f3 | SPU_000048 | Lv-Slc31a1 |
| f3 | SPU_000172 | Lv-Naprt1L |
| f3 | SPU_000262 | Lv-Smcr7L |
| f3 | SPU_000277 | Lv-Ttll9 |
| f3 | SPU_000295 | Lv-Aht1L |
| f3 | SPU_000301 | Lv-Rdh |
| f3 | SPU_000328 | Lv-Cycl2 |
| f3 | SPU_000363 | none |
| f3 | SPU_000563 | Lv-Cdkl2 |

| | | |
|---|---|---|
| f3 | SPU_000577 | Lv-Igvf |
| f3 | SPU_000764 | Lv-Sarmr4 |
| f3 | SPU_000802 | Lv-Hypp_314 |
| f3 | SPU_000858 | none |
| f3 | SPU_000859 | Lv-Slc16a12_9 |
| f3 | SPU_000890 | Lv-Osbpl9 |
| f3 | SPU_000908 | Lv-Ank2h |
| f3 | SPU_000957 | Lv-Ecm/Tspn/Vwc/Vwd |
| f3 | SPU_001092 | Lv-Fn3/Igf_43 |
| f3 | SPU_001127 | none |
| f3 | SPU_001136 | Lv-Pgbd4L |
| f3 | SPU_001214 | Lv-Crat |
| f3 | SPU_001288 | Lv-Rab31 |
| f3 | SPU_001329 | Lv-RtL_10 |
| f3 | SPU_001402 | none |
| f3 | SPU_001538 | Lv-Fn3f |
| f3 | SPU_001562 | Lv-Hypp_335 |
| f3 | SPU_001631 | Lv-C5orf36L |
| f3 | SPU_001672 | Lv-Xrra1L |
| f3 | SPU_001716 | none |
| f3 | SPU_001831 | Lv-Pfkm |
| f3 | SPU_001983 | Lv-NmbrL |
| f3 | SPU_002004 | Lv-Hdpx3 |
| f3 | SPU_002146 | Lv-Hypp_1375 |
| f3 | SPU_002155 | NA |
| f3 | SPU_002165 | Lv-122C |
| f3 | SPU_002256 | Lv-Mlf2 |
| f3 | SPU_002292 | Lv-Syt1-2 |
| f3 | SPU_002431 | Lv-Abcb1b |
| f3 | SPU_002435 | Lv-Sdk2 |
| f3 | SPU_002440 | Lv-Hypp_1395 |
| f3 | SPU_002659 | Lv-CspeN4 |
| f3 | SPU_002694 | none |
| f3 | SPU_002700 | Lv-Hypp_366 |
| f3 | SPU_002811 | none |
| f3 | SPU_002857 | Lv-Gad |
| f3 | SPU_002961 | NA |
| f3 | SPU_003077 | none |

| | | |
|---|---|---|
| f3 | SPU_003086 | Lv-Fn3_19 |
| f3 | SPU_003139 | Lv-C22orf25L |
| f3 | SPU_003210 | Lv-Lycat |
| f3 | SPU_003221 | none |
| f3 | SPU_003235 | Lv-Dcakd |
| f3 | SPU_003493 | Lv-Hypp_1457 |
| f3 | SPU_003691 | Lv-Gtpbp2 |
| f3 | SPU_003762 | Lv-Nlr117 |
| f3 | SPU_003850 | Lv-Ggt1_6 |
| f3 | SPU_003890 | Lv-Hypp_400 |
| f3 | SPU_003959 | none |
| f3 | SPU_004027 | Lv-NotchL_9 |
| f3 | SPU_004220 | Lv-Kctd10 |
| f3 | SPU_004334 | Lv-Endrvt59 |
| f3 | SPU_004455 | Lv-Ankrd26L |
| f3 | SPU_004467 | Lv-Zadh1 |
| f3 | SPU_004471 | Lv-C3orf15 |
| f3 | SPU_004478 | none |
| f3 | SPU_004641 | Lv-Zbtb8os |
| f3 | SPU_004874 | Lv-Hypp_433 |
| f3 | SPU_004976 | Lv-PolppL_41 |
| f3 | SPU_004982 | Lv-C20orf54L_3 |
| f3 | SPU_004993 | Lv-C17orf66L |
| f3 | SPU_005008 | Lv-Xpapd2Lmb_3 |
| f3 | SPU_005085 | NA |
| f3 | SPU_005164 | none |
| f3 | SPU_005231 | NA |
| f3 | SPU_005240 | NA |
| f3 | SPU_005297 | Lv-Hypp_1577 |
| f3 | SPU_005298 | Lv-Racgap1 |
| f3 | SPU_005321 | Lv-Supt3h |
| f3 | SPU_005360 | Lv-Pcdp1L |
| f3 | SPU_005403 | NA |
| f3 | SPU_005439 | Lv-Cyp46L4 |
| f3 | SPU_005448 | Lv-Mgst5 |
| f3 | SPU_005469 | Lv-Nans |
| f3 | SPU_005543 | Lv-Ccdc13 |
| f3 | SPU_005552 | none |

| | | |
|---|---|---|
| f3 | SPU_005558 | none |
| f3 | SPU_005764 | Lv-Rnf41L |
| f3 | SPU_005779 | Lv-Rap1 |
| f3 | SPU_006019 | Lv-Nlr183 |
| f3 | SPU_006074 | Lv-RtL_68 |
| f3 | SPU_006086 | none |
| f3 | SPU_006246 | Lv-Hypp_1641 |
| f3 | SPU_006400 | Lv-Lzic |
| f3 | SPU_006410 | Lv-Pan2 |
| f3 | SPU_006529 | Lv-Gnbp1/2/3B |
| f3 | SPU_006544 | Lv-Xylb_1 |
| f3 | SPU_006650 | Lv-Gba |
| f3 | SPU_006710 | none |
| f3 | SPU_006723 | NA |
| f3 | SPU_006793 | none |
| f3 | SPU_006810 | Lv-Slc8a1 |
| f3 | SPU_006836 | Lv-PolppL_55 |
| f3 | SPU_006947 | Lv-Mnk |
| f3 | SPU_007029 | none |
| f3 | SPU_007068 | Lv-Dhyg_1 |
| f3 | SPU_007071 | Lv-Ccdc65 |
| f3 | SPU_007116 | NA |
| f3 | SPU_007312 | Lv-CycI |
| f3 | SPU_007321 | NA |
| f3 | SPU_007340 | Lv-Wdyhv1 |
| f3 | SPU_007354 | Lv-Tsen54L |
| f3 | SPU_007461 | Lv-Hypp_517 |
| f3 | SPU_007493 | Lv-App |
| f3 | SPU_007495 | Lv-Cd36antigenL |
| f3 | SPU_007689 | Lv-Rt29 |
| f3 | SPU_007720 | Lv-Bstrnph |
| f3 | SPU_007802 | Lv-PolppL_61 |
| f3 | SPU_007878 | Lv-Adprh_1 |
| f3 | SPU_008039 | Lv-Endrvt17 |
| f3 | SPU_008121 | Lv-Slob |
| f3 | SPU_008162 | Lv-Hypp_1769 |
| f3 | SPU_008184 | none |
| f3 | SPU_008231 | none |

| | | |
|---|---|---|
| f3 | SPU_008239 | Lv-Hypp_1773 |
| f3 | SPU_008683 | Lv-Upp1 |
| f3 | SPU_008747 | Lv-Fused |
| f3 | SPU_008804 | Lv-Slc7a11_9 |
| f3 | SPU_009031 | Lv-Acss3 |
| f3 | SPU_009123 | Lv-Nebnph |
| f3 | SPU_009138 | Lv-Hypp_571 |
| f3 | SPU_009230 | Lv-Mprot1L |
| f3 | SPU_009278 | none |
| f3 | SPU_009364 | Lv-Lactb3 |
| f3 | SPU_009370 | Lv-C3orf15_1 |
| f3 | SPU_009411 | NA |
| f3 | SPU_009467 | Lv-Hypp_1852 |
| f3 | SPU_009516 | Lv-CopineA |
| f3 | SPU_009620 | NA |
| f3 | SPU_009674 | Lv-Mfnl2 |
| f3 | SPU_009926 | NA |
| f3 | SPU_009964 | Lv-CnBsi1 |
| f3 | SPU_010186 | Lv-Ttc16 |
| f3 | SPU_010359 | Lv-Hypp_1911 |
| f3 | SPU_010412 | none |
| f3 | SPU_010421 | none |
| f3 | SPU_010443 | Lv-Ankar |
| f3 | SPU_010799 | Lv-Cdkl1 |
| f3 | SPU_010877 | Lv-Iqce |
| f3 | SPU_010947 | NA |
| f3 | SPU_010977 | none |
| f3 | SPU_011002 | Lv-Cebpg |
| f3 | SPU_011034 | Lv-Tropmy2 |
| f3 | SPU_011176 | Lv-Clect/Scop/7Tm |
| f3 | SPU_011249 | Lv-Mtnr1aL_1 |
| f3 | SPU_011407 | Lv-Tbc1d15 |
| f3 | SPU_011480 | Lv-Hypp_1980 |
| f3 | SPU_011596 | Lv-Bdh |
| f3 | SPU_011800 | Lv-PolypL_3 |
| f3 | SPU_012259 | Lv-Fbxw10 |
| f3 | SPU_012319 | Lv-Notchh1_8 |
| f3 | SPU_012481 | Lv-Hypp_2033 |

| | | |
|---|---|---|
| f3 | SPU_012589 | Lv-Slc10a2_3 |
| f3 | SPU_012620 | none |
| f3 | SPU_012659 | Lv-Cryz_2 |
| f3 | SPU_012768 | Lv-Lucf_3 |
| f3 | SPU_012785 | Lv-Pp1A |
| f3 | SPU_012833 | Lv-Hypp_695 |
| f3 | SPU_012850 | Lv-Spata4 |
| f3 | SPU_012897 | none |
| f3 | SPU_012992 | Lv-Dscam/DccL |
| f3 | SPU_013016 | none |
| f3 | SPU_013301 | Lv-Myp |
| f3 | SPU_013352 | Lv-Tirc6(partial) |
| f3 | SPU_013362 | none |
| f3 | SPU_013455 | Lv-Mib2_6 |
| f3 | SPU_013471 | Lv-Mib2_1 |
| f3 | SPU_013843 | Lv-FtzF |
| f3 | SPU_013874 | Lv-HnmtL_1 |
| f3 | SPU_014101 | Lv-TbceL |
| f3 | SPU_014135 | Lv-Slc10a2_7 |
| f3 | SPU_014165 | Lv-EnRT |
| f3 | SPU_014344 | Lv-Arg |
| f3 | SPU_014364 | Lv-Rsb66L |
| f3 | SPU_014468 | Lv-LrpL |
| f3 | SPU_014524 | Lv-Hypp_2156 |
| f3 | SPU_014618 | Lv-Col805b_1 |
| f3 | SPU_014747 | Lv-Surreal/Gpcr117 |
| f3 | SPU_014796 | Lv-Lrriq1 |
| f3 | SPU_014883 | Lv-Unc44L_103 |
| f3 | SPU_014895 | Lv-Wdr38 |
| f3 | SPU_014996 | none |
| f3 | SPU_015121 | none |
| f3 | SPU_015385 | Lv-Lcn7 |
| f3 | SPU_015508 | Lv-Vwa |
| f3 | SPU_015523 | Lv-Hypp_2219 |
| f3 | SPU_015576 | Lv-Decr1 |
| f3 | SPU_015746 | Lv-Fn3_2 |
| f3 | SPU_015789 | Lv-TlrP16 |
| f3 | SPU_015937 | Lv-Srcr128 |

| | | |
|---|---|---|
| f3 | SPU_015975 | Lv-Lca5 |
| f3 | SPU_015996 | Lv-Capn5 |
| f3 | SPU_016095 | Lv-Galnt11 |
| f3 | SPU_016168 | Lv-Mef2 |
| f3 | SPU_016212 | none |
| f3 | SPU_016223 | Lv-Hypp_125 |
| f3 | SPU_016264 | none |
| f3 | SPU_016475 | Lv-RtL_69 |
| f3 | SPU_016504 | none |
| f3 | SPU_016563 | Lv-Cxorf22L |
| f3 | SPU_016668 | Lv-Hypp_837 |
| f3 | SPU_016801 | Lv-Hypp_844 |
| f3 | SPU_016998 | Lv-Hypp_853 |
| f3 | SPU_017062 | none |
| f3 | SPU_017133 | Lv-Pef1 |
| f3 | SPU_017283 | Lv-Ttc6 |
| f3 | SPU_017327 | Lv-Stbd1 |
| f3 | SPU_017334 | none |
| f3 | SPU_017654 | Lv-Hypp_889 |
| f3 | SPU_017733 | Lv-UshL_1 |
| f3 | SPU_017743 | Lv-PogkL |
| f3 | SPU_017964 | Lv-Slc16a14_3 |
| f3 | SPU_017975 | Lv-C3orf25L |
| f3 | SPU_018133 | Lv-Evc2L |
| f3 | SPU_018222 | Lv-Slc16a7L |
| f3 | SPU_018285 | none |
| f3 | SPU_018508 | NA |
| f3 | SPU_018522 | Lv-Slc16a14_4 |
| f3 | SPU_018623 | Lv-Dusp24 |
| f3 | SPU_018739 | Lv-Mapk15_2 |
| f3 | SPU_018825 | Lv-Prardh |
| f3 | SPU_018850 | Lv-Z63 |
| f3 | SPU_018882 | Lv-Zadh2 |
| f3 | SPU_018916 | none |
| f3 | SPU_018986 | Lv-Ccdc42 |
| f3 | SPU_019202 | Lv-C1orf57L |
| f3 | SPU_019354 | NA |
| f3 | SPU_019416 | Lv-GagpolL_16 |

| | | |
|---|---|---|
| f3 | SPU_019528 | NA |
| f3 | SPU_019684 | Lv-Hypp_2461 |
| f3 | SPU_019759 | none |
| f3 | SPU_019793 | Lv-Xcat2 |
| f3 | SPU_019807 | Lv-C18orf34L |
| f3 | SPU_019817 | Lv-LRP2BP |
| f3 | SPU_019842 | none |
| f3 | SPU_019896 | Lv-Arsb_1 |
| f3 | SPU_019917 | Lv-PlexA2_2 |
| f3 | SPU_019922 | Lv-Hypp_2472 |
| f3 | SPU_019926 | Lv-Slsp_11 |
| f3 | SPU_019927 | Lv-Traf3ip1 |
| f3 | SPU_020012 | Lv-UshL |
| f3 | SPU_020059 | none |
| f3 | SPU_020238 | Lv-Ankrd26L_2 |
| f3 | SPU_020263 | Lv-PppL_215 |
| f3 | SPU_020496 | Lv-PolypL_952 |
| f3 | SPU_020547 | Lv-Adam15L3 |
| f3 | SPU_020550 | Lv-Ctbs_1 |
| f3 | SPU_020608 | Lv-Hypp_981 |
| f3 | SPU_020678 | Lv-Nan1 |
| f3 | SPU_020735 | Lv-Unk_64 |
| f3 | SPU_020898 | Lv-Xrra1L_1 |
| f3 | SPU_020969 | Lv-DscamL_2 |
| f3 | SPU_021142 | Lv-Evc2 |
| f3 | SPU_021234 | Lv-Tex2L |
| f3 | SPU_021402 | Lv-Cub/Egf/Lnb/7Tm/Gpcr_1 |
| f3 | SPU_021582 | none |
| f3 | SPU_021692 | Lv-Srcr168 |
| f3 | SPU_021809 | NA |
| f3 | SPU_021856 | Lv-Serinc3 |
| f3 | SPU_021870 | none |
| f3 | SPU_021967 | Lv-Gucy1b2_3 |
| f3 | SPU_022034 | NA |
| f3 | SPU_022035 | Lv-Dyrk4_1 |
| f3 | SPU_022129 | Lv-Rdh1 |
| f3 | SPU_022282 | NA |
| f3 | SPU_022366 | Lv-CubnL_6 |

| | | |
|---|---|---|
| f3 | SPU_022371 | Lv-Rej6 |
| f3 | SPU_022441 | Lv-Nlr163 |
| f3 | SPU_022605 | Lv-Hecah |
| f3 | SPU_022681 | none |
| f3 | SPU_022725 | none |
| f3 | SPU_022759 | Lv-SacSl_10 |
| f3 | SPU_022776 | NA |
| f3 | SPU_022948 | Lv-Ttc39c |
| f3 | SPU_022972 | none |
| f3 | SPU_022986 | Lv-Zadh1_1 |
| f3 | SPU_023086 | Lv-Herc4_1 |
| f3 | SPU_023211 | none |
| f3 | SPU_023416 | none |
| f3 | SPU_023463 | Lv-Wnt4 |
| f3 | SPU_023516 | Lv-Dnm1 |
| f3 | SPU_023646 | Lv-Wbp2nL |
| f3 | SPU_023876 | Lv-Mlcka |
| f3 | SPU_023880 | Lv-Hypp_2694 |
| f3 | SPU_023925 | Lv-Srxn1 |
| f3 | SPU_024016 | Lv-Alas2 |
| f3 | SPU_024111 | Lv-Frmd8 |
| f3 | SPU_024181 | Lv-Nell1L |
| f3 | SPU_024198 | Lv-Hypp_2714 |
| f3 | SPU_024337 | Lv-Tranglutamnt1_1 |
| f3 | SPU_024537 | NA |
| f3 | SPU_024545 | none |
| f3 | SPU_024564 | Lv-Cd109L_2 |
| f3 | SPU_024606 | Lv-Flj20487l |
| f3 | SPU_024641 | none |
| f3 | SPU_024652 | Lv-Fbxl20 |
| f3 | SPU_024874 | Lv-C2orf39L |
| f3 | SPU_024876 | NA |
| f3 | SPU_024958 | Lv-Hypp_222 |
| f3 | SPU_025113 | Lv-Soxb2 |
| f3 | SPU_025156 | Lv-Egfi_6 |
| f3 | SPU_025162 | Lv-Diversin |
| f3 | SPU_025426 | Lv-Hypp_1147 |
| f3 | SPU_025683 | none |

| | | |
|---|---|---|
| f3 | SPU_025741 | Lv-RtL_8 |
| f3 | SPU_025747 | Lv-Kcnk18L_3 |
| f3 | SPU_025822 | Lv-Hypp_2803 |
| f3 | SPU_025893 | Lv-Mak |
| f3 | SPU_025943 | Lv-Tmem145_1 |
| f3 | SPU_026126 | Lv-Hypp_241 |
| f3 | SPU_026184 | Lv-Pgap1 |
| f3 | SPU_026280 | Lv-Krp95 |
| f3 | SPU_026314 | Lv-Ap2s1_1 |
| f3 | SPU_026378 | none |
| f3 | SPU_026498 | Lv-Pkce |
| f3 | SPU_026531 | Lv-ENPP51 |
| f3 | SPU_026570 | Lv-Slsp_24 |
| f3 | SPU_026673 | Lv-Nox5 |
| f3 | SPU_026704 | Lv-Cxorf30L_1 |
| f3 | SPU_026724 | Lv-Eif2ak3perk |
| f3 | SPU_026825 | Lv-Abcb9A |
| f3 | SPU_026885 | Lv-Fam92a |
| f3 | SPU_026899 | Lv-Dyrk2 |
| f3 | SPU_027261 | NA |
| f3 | SPU_027296 | Lv-Chdh_1 |
| f3 | SPU_027352 | Lv-Ccdc147 |
| f3 | SPU_027375 | Lv-Tsh_r |
| f3 | SPU_027515 | Lv-Hypp_1205 |
| f3 | SPU_027535 | Lv-Ly/Zp |
| f3 | SPU_027821 | Lv-PppL_164 |
| f3 | SPU_027969 | Lv-FoxJ1 |
| f3 | SPU_028064 | Lv-Hypp_280 |
| f3 | SPU_028108 | Lv-SacSl_5 |
| f3 | SPU_028301 | Lv-Z163 |
| f3 | SPU_028693 | none |
| f3 | SPU_028708 | Lv-Dhyg_4 |
| f3 | SPU_028736 | Lv-NaaladlL |
| f3 | SPU_028817 | Lv-SybL1 |
| f3 | SPU_028899 | none |
| f3 | SPU_030165 | Lv-Nek1c |
| f4 | SPU_000114 | Lv-Fut6 |
| f4 | SPU_000141 | Lv-Clect/Egf/lamininl/fibronectin3/Tm |

| | | |
|---|---|---|
| f4 | SPU_000286 | Lv-Commd4 |
| f4 | SPU_000291 | Lv-Tusc2L |
| f4 | SPU_000354 | Lv-Tmem14C |
| f4 | SPU_000522 | Lv-Aldh8a1 |
| f4 | SPU_000641 | Lv-Echdc3 |
| f4 | SPU_000758 | Lv-Slc16 |
| f4 | SPU_000868 | Lv-Hypp_315 |
| f4 | SPU_000920 | Lv-Ambra1L |
| f4 | SPU_000943 | Lv-Rvt/Exn/End/Pp_1 |
| f4 | SPU_000952 | Lv-Cables1 |
| f4 | SPU_000956 | Lv-Hypp_1297 |
| f4 | SPU_000987 | none |
| f4 | SPU_001021 | Lv-EhH3_4 |
| f4 | SPU_001034 | Lv-Endrvt55 |
| f4 | SPU_001081 | Lv-Heatr2 |
| f4 | SPU_001102 | Lv-Hypp_1305 |
| f4 | SPU_001263 | Lv-Thada |
| f4 | SPU_001281 | Lv-Zmym1 |
| f4 | SPU_001308 | Lv-Taf12 |
| f4 | SPU_001347 | Lv-Hypp_330 |
| f4 | SPU_001365 | Lv-Su(fu) |
| f4 | SPU_001393 | Lv-Eif5B_1 |
| f4 | SPU_001420 | Lv-Spag7L |
| f4 | SPU_001429 | Lv-Endrvt56 |
| f4 | SPU_001493 | Lv-Tmem67_1 |
| f4 | SPU_001539 | none |
| f4 | SPU_001651 | Lv-PolppL-3 |
| f4 | SPU_001721 | Lv-Lin41 |
| f4 | SPU_001747 | Lv-Hypp_1344 |
| f4 | SPU_001787 | none |
| f4 | SPU_001832 | Lv-Sf3B14L |
| f4 | SPU_001883 | Lv-Fut4L |
| f4 | SPU_001888 | Lv-Ccp_1 |
| f4 | SPU_001980 | Lv-Rc3h2 |
| f4 | SPU_002018 | none |
| f4 | SPU_002103 | Lv-Taf |
| f4 | SPU_002281 | Lv-Rev3_1 |
| f4 | SPU_002364 | none |

| | | |
|---|---|---|
| f4 | SPU_002397 | Lv-Hypp_355 |
| f4 | SPU_002457 | none |
| f4 | SPU_002490 | Lv-Pgls |
| f4 | SPU_002551 | Lv-Egfiii |
| f4 | SPU_002593 | none |
| f4 | SPU_002645 | Lv-Srpk1L |
| f4 | SPU_002655 | Lv-Spf30 |
| f4 | SPU_002769 | Lv-Scap |
| f4 | SPU_002785 | Lv-Dda1 |
| f4 | SPU_002904 | Lv-C7orf43L |
| f4 | SPU_002930 | Lv-Pres |
| f4 | SPU_002968 | Lv-CalmL4 |
| f4 | SPU_003061 | Lv-Zc3p26_6 |
| f4 | SPU_003075 | Lv-Nup93 |
| f4 | SPU_003095 | Lv-PolppL_24 |
| f4 | SPU_003137 | Lv-Rbph |
| f4 | SPU_003149 | Lv-Dusp12 |
| f4 | SPU_003151 | Lv-EnRTL3 |
| f4 | SPU_003162 | Lv-Arf1l2 |
| f4 | SPU_003223 | Lv-Ift81h |
| f4 | SPU_003275 | Lv-Hypp_1439 |
| f4 | SPU_003317 | Lv-PolppL_25 |
| f4 | SPU_003340 | Lv-Endrvt57 |
| f4 | SPU_003350 | Lv-Lrp12L |
| f4 | SPU_003397 | Lv-Chka |
| f4 | SPU_003406 | Lv-Arfgef1 |
| f4 | SPU_003438 | Lv-Pak1 |
| f4 | SPU_003472 | Lv-Brip1 |
| f4 | SPU_003475 | Lv-Mrpl14L |
| f4 | SPU_003494 | Lv-Hypp_385 |
| f4 | SPU_003503 | Lv-Hypp_1459 |
| f4 | SPU_003642 | Lv-Agpat3 |
| f4 | SPU_003645 | Lv-Rae1 |
| f4 | SPU_003647 | Lv-Hypp_1466 |
| f4 | SPU_003673 | Lv-Gpr126L |
| f4 | SPU_003795 | none |
| f4 | SPU_003829 | Lv-Txndc9 |
| f4 | SPU_003847 | Lv-Psmc3_1 |

| | | |
|---|---|---|
| f4 | SPU_003892 | Lv-Otubl |
| f4 | SPU_003903 | Lv-Hypp_2209 |
| f4 | SPU_003936 | none |
| f4 | SPU_003937 | Lv-Kiaa1822 |
| f4 | SPU_003962 | none |
| f4 | SPU_003999 | none |
| f4 | SPU_004003 | Lv-Galnt7-2 |
| f4 | SPU_004061 | Lv-Nido/Egf/Zp |
| f4 | SPU_004168 | Lv-Hypp_1502 |
| f4 | SPU_004278 | Lv-PolppL_36 |
| f4 | SPU_004291 | NA |
| f4 | SPU_004314 | Lv-Setd8l-4 |
| f4 | SPU_004327 | Lv-Sec61g |
| f4 | SPU_004335 | Lv-Susd2-4 |
| f4 | SPU_004385 | Lv-Hypp_415 |
| f4 | SPU_004408 | Lv-Pick1 |
| f4 | SPU_004414 | Lv-Hlf |
| f4 | SPU_004463 | Lv-Ei24 |
| f4 | SPU_004465 | Lv-Hypp_1525 |
| f4 | SPU_004474 | Lv-Nubpl |
| f4 | SPU_004517 | Lv-Pellino |
| f4 | SPU_004584 | Lv-Sdf2 |
| f4 | SPU_004689 | Lv-Asteh1 |
| f4 | SPU_004723 | Lv-Err |
| f4 | SPU_004735 | Lv-Chd1 |
| f4 | SPU_004736 | none |
| f4 | SPU_004769 | Lv-Ass |
| f4 | SPU_004786 | Lv-Madp1 |
| f4 | SPU_004826 | none |
| f4 | SPU_004842 | none |
| f4 | SPU_004865 | Lv-Wdr20a |
| f4 | SPU_004878 | Lv-Hypp_1549 |
| f4 | SPU_004908 | none |
| f4 | SPU_005001 | none |
| f4 | SPU_005048 | Lv-Myo19L |
| f4 | SPU_005061 | Lv-Flot2h |
| f4 | SPU_005110 | Lv-Tomm22L_2 |
| f4 | SPU_005114 | Lv-TpaseL_33 |

| | | |
|---|---|---|
| f4 | SPU_005133 | Lv-PppL_180 |
| f4 | SPU_005136 | Lv-Msh5_1 |
| f4 | SPU_005192 | Lv-Pip5k1a |
| f4 | SPU_005232 | none |
| f4 | SPU_005258 | Lv-Sgsm3 |
| f4 | SPU_005292 | none |
| f4 | SPU_005372 | Lv-Hypp_445 |
| f4 | SPU_005438 | none |
| f4 | SPU_005496 | Lv-Hypp_1586 |
| f4 | SPU_005582 | Lv-PolppL_51 |
| f4 | SPU_005584 | Lv-Gtf3c5 |
| f4 | SPU_005621 | none |
| f4 | SPU_005640 | Lv-Sdccag10L |
| f4 | SPU_005641 | Lv-Hectd3 |
| f4 | SPU_005675 | Lv-Nudt9 |
| f4 | SPU_005722 | Lv-Atg5 |
| f4 | SPU_005875 | Lv-CspeN5 |
| f4 | SPU_005876 | Lv-Med15 |
| f4 | SPU_005941 | Lv-Csmd3L |
| f4 | SPU_006079 | Lv-Chst11_8 |
| f4 | SPU_006080 | Lv-Hypp_1628 |
| f4 | SPU_006163 | Lv-Dctn4 |
| f4 | SPU_006189 | Lv-Mis12l |
| f4 | SPU_006194 | Lv-L1td1L |
| f4 | SPU_006243 | Lv-Hypp_471 |
| f4 | SPU_006310 | Lv-Clect/Egf_8 |
| f4 | SPU_006358 | none |
| f4 | SPU_006404 | none |
| f4 | SPU_006471 | Lv-Int4 |
| f4 | SPU_006475 | Lv-Trb |
| f4 | SPU_006606 | none |
| f4 | SPU_006809 | Lv-Fbp2 |
| f4 | SPU_006864 | Lv-BolaL3 |
| f4 | SPU_006911 | Lv-77kDamap |
| f4 | SPU_006923 | Lv-PolypL_936 |
| f4 | SPU_006936 | Lv-Thoc6h |
| f4 | SPU_006953 | none |
| f4 | SPU_006957 | Lv-Hypp_1689 |

| | | |
|---|---|---|
| f4 | SPU_007036 | Lv-Vps35h |
| f4 | SPU_007072 | Lv-LrpL_4 |
| f4 | SPU_007119 | none |
| f4 | SPU_007197 | NA |
| f4 | SPU_007207 | Lv-Hypp_1705 |
| f4 | SPU_007214 | Lv-Nmt2 |
| f4 | SPU_007243 | Lv-Commd7 |
| f4 | SPU_007317 | Lv-Bmp1/TldL1 |
| f4 | SPU_007486 | Lv-NeurL2 |
| f4 | SPU_007523 | Lv-FlvcrL |
| f4 | SPU_007531 | Lv-Hypp_519 |
| f4 | SPU_007569 | Lv-Mars2 |
| f4 | SPU_007724 | NA |
| f4 | SPU_007727 | none |
| f4 | SPU_007760 | Lv-Naglu |
| f4 | SPU_007818 | Lv-PolppL_62 |
| f4 | SPU_007875 | Lv-Ccdc150 |
| f4 | SPU_007942 | Lv-Epn2 |
| f4 | SPU_007971 | Lv-Mrps16l |
| f4 | SPU_007998 | Lv-Hypp_1758 |
| f4 | SPU_008007 | Lv-PolppL_64 |
| f4 | SPU_008051 | none |
| f4 | SPU_008077 | Lv-Parg |
| f4 | SPU_008157 | Lv-Hypp_3001 |
| f4 | SPU_008177 | Lv-Nfkb |
| f4 | SPU_008210 | none |
| f4 | SPU_008262 | Lv-Endrvt66 |
| f4 | SPU_008327 | Lv-PolppL_70 |
| f4 | SPU_008349 | Lv-Amt |
| f4 | SPU_008358 | Lv-Z229 |
| f4 | SPU_008377 | Lv-Z105 |
| f4 | SPU_008403 | Lv-Dhtkd1 |
| f4 | SPU_008405 | Lv-Lztr1 |
| f4 | SPU_008411 | Lv-Wbscr1L |
| f4 | SPU_008423 | Lv-PolppL_72 |
| f4 | SPU_008446 | Lv-Hypp_1784 |
| f4 | SPU_008471 | Lv-Dlc3 |
| f4 | SPU_008529 | Lv-Hypp_552 |

| | | |
|---|---|---|
| f4 | SPU_008534 | Lv-Derl2_1 |
| f4 | SPU_008535 | Lv-Serca_1 |
| f4 | SPU_008567 | Lv-FormL2iB |
| f4 | SPU_008571 | Lv-PolypL_939 |
| f4 | SPU_008624 | NA |
| f4 | SPU_008635 | Lv-Rt31 |
| f4 | SPU_008642 | Lv-Srcr71 |
| f4 | SPU_008643 | Lv-Hypp_34 |
| f4 | SPU_008674 | none |
| f4 | SPU_008810 | Lv-Hypp_1807 |
| f4 | SPU_009121 | Lv-Hypp_570 |
| f4 | SPU_009141 | Lv-Mrpl24 |
| f4 | SPU_009269 | Lv-Uqcrfs1_1 |
| f4 | SPU_009359 | Lv-Tmlhe |
| f4 | SPU_009493 | Lv-C4bpaL |
| f4 | SPU_009536 | Lv-Hypp_50 |
| f4 | SPU_009574 | Lv-PolppL_84 |
| f4 | SPU_009575 | Lv-PolppL_85 |
| f4 | SPU_009598 | none |
| f4 | SPU_009601 | Lv-Cts4 |
| f4 | SPU_009681 | Lv-Pkn |
| f4 | SPU_009863 | none |
| f4 | SPU_009876 | Lv-FoxP |
| f4 | SPU_009945 | none |
| f4 | SPU_010173 | Lv-Endrvt73 |
| f4 | SPU_010211 | Lv-Onpep |
| f4 | SPU_010324 | Lv-Xrcc1 |
| f4 | SPU_010402 | Lv-Nfs1 |
| f4 | SPU_010463 | Lv-Sf3B1 |
| f4 | SPU_010488 | Lv-Hip1_1 |
| f4 | SPU_010546 | Lv-Med7 |
| f4 | SPU_010617 | Lv-Slco4a |
| f4 | SPU_010635 | Lv-Gatae |
| f4 | SPU_010703 | Lv-SemaB |
| f4 | SPU_010749 | Lv-Hypp_1938 |
| f4 | SPU_010760 | none |
| f4 | SPU_010771 | Lv-Endrvt21 |
| f4 | SPU_010871 | Lv-MthL |

| | | |
|---|---|---|
| f4 | SPU_010904 | Lv-PolppL_90 |
| f4 | SPU_010923 | Lv-Mrpl37 |
| f4 | SPU_010933 | Lv-Naglu_1 |
| f4 | SPU_010962 | Lv-Rabac1 |
| f4 | SPU_011010 | none |
| f4 | SPU_011098 | Lv-Svep1L |
| f4 | SPU_011181 | Lv-Hypp_643 |
| f4 | SPU_011223 | Lv-LhH1b |
| f4 | SPU_011255 | Lv-Sult1a |
| f4 | SPU_011260 | Lv-Z181 |
| f4 | SPU_011353 | Lv-Kif27L |
| f4 | SPU_011377 | Lv-Hypp_1977 |
| f4 | SPU_011504 | Lv-Ure |
| f4 | SPU_011656 | Lv-Sc4mol_1 |
| f4 | SPU_011667 | NA |
| f4 | SPU_011690 | Lv-Pias2/3 |
| f4 | SPU_011702 | Lv-RtL_28 |
| f4 | SPU_011853 | Lv-Endrvt24 |
| f4 | SPU_011915 | Lv-Hypp_2007 |
| f4 | SPU_011920 | none |
| f4 | SPU_011973 | Lv-Hgs |
| f4 | SPU_011982 | Lv-Kctd1L_3 |
| f4 | SPU_012112 | Lv-Rab5l |
| f4 | SPU_012177 | none |
| f4 | SPU_012197 | Lv-Hypp_675 |
| f4 | SPU_012242 | Lv-Kmo |
| f4 | SPU_012449 | Lv-Ttc12 |
| f4 | SPU_012551 | none |
| f4 | SPU_012559 | Lv-Ap2s1 |
| f4 | SPU_012592 | Lv-PolypL_942 |
| f4 | SPU_012655 | Lv-Egfi |
| f4 | SPU_012672 | Lv-Ralgps1 |
| f4 | SPU_012694 | Lv-Abcd1 |
| f4 | SPU_012857 | Lv-Mon1a |
| f4 | SPU_012940 | none |
| f4 | SPU_013044 | Lv-Endrvt_928 |
| f4 | SPU_013055 | none |
| f4 | SPU_013065 | Lv-RtL |

| | | |
|---|---|---|
| f4 | SPU_013173 | none |
| f4 | SPU_013176 | Lv-Hypp_2074 |
| f4 | SPU_013198 | Lv-Plekha8 |
| f4 | SPU_013369 | Lv-RgL1 |
| f4 | SPU_013381 | Lv-Trappc2 |
| f4 | SPU_013402 | Lv-Hypp_715 |
| f4 | SPU_013407 | Lv-Z175 |
| f4 | SPU_013409 | Lv-HsF |
| f4 | SPU_013456 | Lv-Endrvt86 |
| f4 | SPU_013501 | Lv-Clcn3 |
| f4 | SPU_013579 | Lv-Moka_1 |
| f4 | SPU_013688 | Lv-Unc50 |
| f4 | SPU_013799 | Lv-Timm9 |
| f4 | SPU_013826 | Lv-Hypp_729 |
| f4 | SPU_013838 | Lv-Etfa |
| f4 | SPU_013858 | none |
| f4 | SPU_013929 | Lv-Rab10 |
| f4 | SPU_013944 | Lv-Slc7a6_2 |
| f4 | SPU_013997 | Lv-Harbi1L |
| f4 | SPU_014005 | Lv-Mrpl1l |
| f4 | SPU_014098 | Lv-Hypp_2128 |
| f4 | SPU_014179 | Lv-Hars |
| f4 | SPU_014244 | Lv-Sptlc2_1 |
| f4 | SPU_014280 | Lv-Pfas_1 |
| f4 | SPU_014284 | Lv-Hypp_747 |
| f4 | SPU_014321 | Lv-Hypp_749 |
| f4 | SPU_014434 | Lv-Hypp_2147 |
| f4 | SPU_014446 | Lv-Med21 |
| f4 | SPU_014469 | NA |
| f4 | SPU_014481 | Lv-Rabif |
| f4 | SPU_014518 | Lv-Trappc10_1 |
| f4 | SPU_014552 | none |
| f4 | SPU_014639 | Lv-Leng8 |
| f4 | SPU_014654 | Lv-Vps33A |
| f4 | SPU_014697 | Lv-Aars_2 |
| f4 | SPU_014815 | none |
| f4 | SPU_014839 | Lv-Lvn1.3 |
| f4 | SPU_014898 | Lv-Rt35 |

| | | |
|---|---|---|
| f4 | SPU_015082 | none |
| f4 | SPU_015086 | none |
| f4 | SPU_015089 | Lv-Auh |
| f4 | SPU_015136 | Lv-Rag1l4 |
| f4 | SPU_015229 | Lv-Vps26hb |
| f4 | SPU_015235 | Lv-Tax1bp1 |
| f4 | SPU_015256 | Lv-Cyp2L29 |
| f4 | SPU_015267 | Lv-Wdrc1 |
| f4 | SPU_015313 | NA |
| f4 | SPU_015343 | Lv-Ugt1L_11 |
| f4 | SPU_015364 | Lv-Whsc2 |
| f4 | SPU_015425 | Lv-Sin3a |
| f4 | SPU_015433 | Lv-Hypp_2212 |
| f4 | SPU_015446 | Lv-Snapc3 |
| f4 | SPU_015467 | Lv-Mcfd2 |
| f4 | SPU_015550 | Lv-PolE3 |
| f4 | SPU_015555 | none |
| f4 | SPU_015635 | Lv-Hypp_795 |
| f4 | SPU_015833 | Lv-Bre |
| f4 | SPU_015849 | Lv-Tropmyh |
| f4 | SPU_015890 | Lv-Hypp_2236 |
| f4 | SPU_016055 | Lv-Nidogen |
| f4 | SPU_016145 | Lv-Endrvt121 |
| f4 | SPU_016173 | Lv-Abi2 |
| f4 | SPU_016221 | Lv-Rpgrip1L_2 |
| f4 | SPU_016233 | none |
| f4 | SPU_016275 | none |
| f4 | SPU_016323 | Lv-Rt7 |
| f4 | SPU_016325 | Lv-RtL_42 |
| f4 | SPU_016418 | Lv-Herc4 |
| f4 | SPU_016436 | Lv-Endrvt91 |
| f4 | SPU_016448 | Lv-Hypp_129 |
| f4 | SPU_016462 | Lv-Rt8 |
| f4 | SPU_016472 | Lv-Hypp_3051 |
| f4 | SPU_016487 | none |
| f4 | SPU_016524 | Lv-Ngly1 |
| f4 | SPU_016537 | Lv-Btbd5 |
| f4 | SPU_016539 | Lv-Rt9 |

| | | |
|---|---|---|
| f4 | SPU_016637 | Lv-Pex11b |
| f4 | SPU_016660 | Lv-Pold2 |
| f4 | SPU_016720 | Lv-Sirt2 |
| f4 | SPU_016728 | Lv-PolypL_91 |
| f4 | SPU_016741 | Lv-PppL_111 |
| f4 | SPU_016790 | Lv-Rac |
| f4 | SPU_016795 | Lv-Hypp_3045 |
| f4 | SPU_016818 | none |
| f4 | SPU_016918 | Lv-Rvt/Exn/End/Pplp_1 |
| f4 | SPU_016929 | none |
| f4 | SPU_017066 | Lv-Hypp_859 |
| f4 | SPU_017072 | Lv-Rab8 |
| f4 | SPU_017074 | Lv-EgfiL_55 |
| f4 | SPU_017174 | Lv-Abhd10 |
| f4 | SPU_017225 | none |
| f4 | SPU_017230 | Lv-Hypp_2313 |
| f4 | SPU_017355 | Lv-Mrpl46 |
| f4 | SPU_017376 | Lv-PppL_115 |
| f4 | SPU_017403 | Lv-Aldh8a1_1 |
| f4 | SPU_017411 | Lv-Sfrs17a |
| f4 | SPU_017463 | Lv-Ergic1 |
| f4 | SPU_017466 | Lv-Mcts1 |
| f4 | SPU_017479 | Lv-Rt10 |
| f4 | SPU_017607 | Lv-Wdr61_1 |
| f4 | SPU_017677 | Lv-PolypL_950 |
| f4 | SPU_017682 | Lv-Z203 |
| f4 | SPU_017683 | none |
| f4 | SPU_017709 | Lv-Endrvt34 |
| f4 | SPU_017723 | Lv-Gtf2A2 |
| f4 | SPU_017817 | NA |
| f4 | SPU_017835 | NA |
| f4 | SPU_017844 | none |
| f4 | SPU_017956 | Lv-Zswim6 |
| f4 | SPU_018066 | Lv-PppL_119 |
| f4 | SPU_018148 | none |
| f4 | SPU_018184 | Lv-Parp1 |
| f4 | SPU_018270 | Lv-Acss1 |
| f4 | SPU_018400 | Lv-Nnt |

| | | |
|---|---|---|
| f4 | SPU_018451 | Lv-PolypL_79 |
| f4 | SPU_018482 | none |
| f4 | SPU_018495 | none |
| f4 | SPU_018496 | none |
| f4 | SPU_018512 | Lv-Anapc2 |
| f4 | SPU_018533 | Lv-Kif26 |
| f4 | SPU_018597 | Lv-Psmd6 |
| f4 | SPU_018686 | Lv-Nfkbi2L |
| f4 | SPU_018716 | Lv-Nenf |
| f4 | SPU_018720 | Lv-Son |
| f4 | SPU_018724 | Lv-Ysk |
| f4 | SPU_018745 | Lv-RtL_97 |
| f4 | SPU_018833 | Lv-Sumo |
| f4 | SPU_018851 | none |
| f4 | SPU_018862 | Lv-FbsL |
| f4 | SPU_018976 | NA |
| f4 | SPU_019021 | Lv-Nek1 |
| f4 | SPU_019031 | Lv-Epc1 |
| f4 | SPU_019046 | Lv-Pccb |
| f4 | SPU_019083 | Lv-Hypp_146 |
| f4 | SPU_019136 | none |
| f4 | SPU_019178 | Lv-Hypp_2422 |
| f4 | SPU_019194 | Lv-Hypp_930 |
| f4 | SPU_019296 | Lv-Golga5 |
| f4 | SPU_019312 | Lv-Gss_1 |
| f4 | SPU_019324 | Lv-Hypp_936 |
| f4 | SPU_019367 | Lv-Endrvt_4 |
| f4 | SPU_019390 | NA |
| f4 | SPU_019426 | Lv-Vps4b |
| f4 | SPU_019519 | Lv-Tada1L |
| f4 | SPU_019525 | Lv-Hydin_2 |
| f4 | SPU_019551 | Lv-Acap2 |
| f4 | SPU_019582 | Lv-Imp3 |
| f4 | SPU_019622 | Lv-Ap3m1_1 |
| f4 | SPU_019643 | Lv-Kctd1L_5 |
| f4 | SPU_019712 | Lv-Acox3_1 |
| f4 | SPU_019732 | Lv-Med27 |
| f4 | SPU_019940 | Lv-Osbpl10 |

| | | |
|---|---|---|
| f4 | SPU_020126 | Lv-Zcchc3 |
| f4 | SPU_020143 | Lv-Cyb5r3 |
| f4 | SPU_020144 | Lv-Rheb |
| f4 | SPU_020165 | none |
| f4 | SPU_020214 | Lv-Wdr3 |
| f4 | SPU_020283 | Lv-Fbxo3 |
| f4 | SPU_020295 | Lv-Surreal/Gpcr113 |
| f4 | SPU_020351 | none |
| f4 | SPU_020401 | none |
| f4 | SPU_020403 | none |
| f4 | SPU_020405 | Lv-Armc6 |
| f4 | SPU_020498 | Lv-Rtcd1 |
| f4 | SPU_020508 | Lv-Ccdc108_2 |
| f4 | SPU_020523 | none |
| f4 | SPU_020534 | Lv-Surf1 |
| f4 | SPU_020558 | Lv-EnRTL2 |
| f4 | SPU_020559 | Lv-Rt39 |
| f4 | SPU_020586 | Lv-Hypp_3035 |
| f4 | SPU_020599 | none |
| f4 | SPU_020638 | Lv-Hypp_2517 |
| f4 | SPU_020707 | Lv-Impk |
| f4 | SPU_020764 | Lv-Shprh |
| f4 | SPU_020833 | Lv-Nutf2 |
| f4 | SPU_020870 | Lv-Gps1 |
| f4 | SPU_020877 | Lv-Hypp_1001 |
| f4 | SPU_020878 | NA |
| f4 | SPU_020917 | Lv-Exoc8 |
| f4 | SPU_020929 | Lv-Bcdin3 |
| f4 | SPU_021024 | Lv-Kiaa0368L |
| f4 | SPU_021045 | Lv-Endrvt7 |
| f4 | SPU_021121 | Lv-Cul3 |
| f4 | SPU_021145 | Lv-Hypp_1015 |
| f4 | SPU_021179 | Lv-PnpLa2 |
| f4 | SPU_021337 | none |
| f4 | SPU_021400 | Lv-RtL_53 |
| f4 | SPU_021483 | Lv-ParP2l |
| f4 | SPU_021591 | Lv-Hypp_2564 |
| f4 | SPU_021678 | Lv-PppL_130 |

| | | |
|---|---|---|
| f4 | SPU_021732 | Lv-Gpn2 |
| f4 | SPU_021774 | Lv-Snf8 |
| f4 | SPU_021823 | Lv-Rt16 |
| f4 | SPU_021830 | Lv-Hypp_2578 |
| f4 | SPU_021877 | none |
| f4 | SPU_021881 | none |
| f4 | SPU_021915 | Lv-Endrvt104 |
| f4 | SPU_021956 | Lv-Rt17 |
| f4 | SPU_022126 | Lv-Kctd1_3 |
| f4 | SPU_022242 | Lv-Smad_ip |
| f4 | SPU_022448 | none |
| f4 | SPU_022449 | Lv-Hypp_184 |
| f4 | SPU_022457 | NA |
| f4 | SPU_022526 | none |
| f4 | SPU_022547 | none |
| f4 | SPU_022553 | Lv-Unc44_161 |
| f4 | SPU_022594 | Lv-Pcnp |
| f4 | SPU_022601 | Lv-Tpase8 |
| f4 | SPU_022767 | Lv-Bbs5_1 |
| f4 | SPU_022828 | Lv-Endrvt44 |
| f4 | SPU_022831 | Lv-PppL_134 |
| f4 | SPU_022855 | Lv-Sfrs16_1 |
| f4 | SPU_022863 | NA |
| f4 | SPU_022888 | Lv-Uqcrc2 |
| f4 | SPU_022899 | Lv-Abhd7_1 |
| f4 | SPU_022916 | Lv-Frizz5/8 |
| f4 | SPU_023006 | Lv-Hypp_2643 |
| f4 | SPU_023081 | Lv-Hypp_1070 |
| f4 | SPU_023188 | Lv-Rvt/Exn/End/Pplp |
| f4 | SPU_023267 | Lv-Hypp_200 |
| f4 | SPU_023342 | Lv-Wdr51a |
| f4 | SPU_023363 | Lv-Slco4a1_14 |
| f4 | SPU_023394 | none |
| f4 | SPU_023450 | Lv-Usp30 |
| f4 | SPU_023476 | Lv-C7orf28b |
| f4 | SPU_023529 | Lv-Luc7l2L |
| f4 | SPU_023560 | Lv-Kif1BL1 |
| f4 | SPU_023562 | Lv-Hypp_2677 |

| | | |
|---|---|---|
| f4 | SPU_023656 | Lv-Uqcrc2_1 |
| f4 | SPU_023661 | NA |
| f4 | SPU_023665 | Lv-Hypp_205 |
| f4 | SPU_023684 | none |
| f4 | SPU_023689 | Lv-Slc24a4 |
| f4 | SPU_023717 | none |
| f4 | SPU_023787 | none |
| f4 | SPU_023827 | Lv-Ccdc24 |
| f4 | SPU_023836 | Lv-Xrcc3 |
| f4 | SPU_023844 | Lv-Notch1-19 |
| f4 | SPU_023873 | Lv-Als2cr4L |
| f4 | SPU_023963 | Lv-Srd5a1_1 |
| f4 | SPU_024100 | Lv-Sf3B1_1 |
| f4 | SPU_024101 | Lv-SF3B1 |
| f4 | SPU_024105 | Lv-Hypp_2709 |
| f4 | SPU_024282 | Lv-PppL_138 |
| f4 | SPU_024283 | none |
| f4 | SPU_024592 | Lv-Endrvt9 |
| f4 | SPU_024670 | Lv-Psmc2 |
| f4 | SPU_024694 | NA |
| f4 | SPU_024725 | Lv-Exoc5 |
| f4 | SPU_024855 | Lv-Endrvt110 |
| f4 | SPU_024917 | Lv-Hypp_1130 |
| f4 | SPU_024920 | Lv-Mic1 |
| f4 | SPU_025054 | none |
| f4 | SPU_025338 | none |
| f4 | SPU_025357 | Lv-Kiaa1772L_1 |
| f4 | SPU_025497 | none |
| f4 | SPU_025579 | Lv-Grb10 |
| f4 | SPU_025619 | none |
| f4 | SPU_025627 | Lv-Ube3a-3 |
| f4 | SPU_025648 | Lv-Harbi1_3 |
| f4 | SPU_025723 | Lv-NippbL |
| f4 | SPU_025730 | Lv-Arg1 |
| f4 | SPU_025786 | Lv-Ercc3 |
| f4 | SPU_025852 | NA |
| f4 | SPU_025907 | none |
| f4 | SPU_025933 | Lv-Hypp_1161 |

| | | |
|---|---|---|
| f4 | SPU_025970 | Lv-Hypp_1162 |
| f4 | SPU_026085 | Lv-Hypp_1167 |
| f4 | SPU_026086 | Lv-Endrvt48 |
| f4 | SPU_026137 | Lv-Actn |
| f4 | SPU_026227 | Lv-Wdr92 |
| f4 | SPU_026256 | Lv-Wasf2_1 |
| f4 | SPU_026357 | Lv-Cog4 |
| f4 | SPU_026484 | Lv-Rt18 |
| f4 | SPU_026600 | none |
| f4 | SPU_026635 | Lv-Gas2 |
| f4 | SPU_026676 | Lv-Fn3f_51 |
| f4 | SPU_026695 | Lv-Rprd1b |
| f4 | SPU_026699 | Lv-Hypp_2861 |
| f4 | SPU_026708 | none |
| f4 | SPU_026728 | Lv-PolypL_85 |
| f4 | SPU_026803 | Lv-PppL_149 |
| f4 | SPU_026852 | NA |
| f4 | SPU_026951 | Lv-Hypp_2877 |
| f4 | SPU_026981 | Lv-Endrvt51 |
| f4 | SPU_027090 | Lv-FibropellinL3 |
| f4 | SPU_027102 | Lv-Hypp_1197 |
| f4 | SPU_027402 | Lv-Btbd33 |
| f4 | SPU_027429 | Lv-Hypp_2903 |
| f4 | SPU_027453 | Lv-WrnL_2 |
| f4 | SPU_027482 | Lv-Dr1 |
| f4 | SPU_027509 | Lv-Hypp_278 |
| f4 | SPU_027536 | Lv-Herc4L_4 |
| f4 | SPU_027600 | Lv-Rag1l |
| f4 | SPU_027628 | Lv-Arhgap28 |
| f4 | SPU_027718 | none |
| f4 | SPU_027785 | Lv-Kat5 |
| f4 | SPU_027791 | Lv-Ftcd_3 |
| f4 | SPU_027829 | Lv-Hypp_276 |
| f4 | SPU_027879 | NA |
| f4 | SPU_027909 | Lv-Ikke_1 |
| f4 | SPU_027959 | Lv-Hypp_9787 |
| f4 | SPU_027985 | Lv-Dhcr24 |
| f4 | SPU_028115 | NA |

| | | |
|---|---|---|
| f4 | SPU_028157 | Lv-Sep15 |
| f4 | SPU_028170 | none |
| f4 | SPU_028180 | Lv-Rad17 |
| f4 | SPU_028225 | none |
| f4 | SPU_028255 | Lv-Nr1x |
| f4 | SPU_028260 | Lv-Hypp_7282 |
| f4 | SPU_028522 | none |
| f4 | SPU_028625 | Lv-Unk_67 |
| f4 | SPU_028640 | Lv-Hypp_289 |
| f4 | SPU_028660 | none |
| f4 | SPU_028768 | Lv-PppL_167 |
| f4 | SPU_028818 | NA |
| f4 | SPU_028847 | none |
| f4 | SPU_028910 | none |
| f4 | SPU_028918 | Lv-C9orf5L |
| f5 | SPU_000035 | Lv-Efhdc2 |
| f5 | SPU_000041 | Lv-Vwd_mucL |
| f5 | SPU_000159 | Lv-Bdnr |
| f5 | SPU_000285 | Lv-Copz1 |
| f5 | SPU_000321 | Lv-PompL |
| f5 | SPU_000513 | Lv-Do1(Udx1) |
| f5 | SPU_000562 | none |
| f5 | SPU_000803 | Lv-Hypp_3117 |
| f5 | SPU_000811 | Lv-Rab3 |
| f5 | SPU_000979 | Lv-Hs3st3a1 |
| f5 | SPU_000992 | Lv-AnthrtrL_1 |
| f5 | SPU_001142 | Lv-Sqle_1 |
| f5 | SPU_001205 | Lv-Unc44L_9 |
| f5 | SPU_001247 | Lv-Psmd12 |
| f5 | SPU_001250 | Lv-Sept2 |
| f5 | SPU_001314 | none |
| f5 | SPU_001616 | none |
| f5 | SPU_001678 | Lv-Galr2L_4 |
| f5 | SPU_001783 | Lv-Hypp_1346 |
| f5 | SPU_001860 | Lv-Atp6v0a1 |
| f5 | SPU_001874 | Lv-Advph |
| f5 | SPU_001875 | NA |
| f5 | SPU_001928 | Lv-Hyalin1 |

| | | |
|---|---|---|
| f5 | SPU_001929 | Lv-Exostl |
| f5 | SPU_002051 | Lv-Atp6v0a1_1 |
| f5 | SPU_002209 | Lv-Acsl6 |
| f5 | SPU_002375 | Lv-Bhmt |
| f5 | SPU_002447 | Lv-C10orf119L |
| f5 | SPU_002528 | Lv-Spast |
| f5 | SPU_002574 | Lv-Hypp_359 |
| f5 | SPU_002579 | Lv-Atg9aL |
| f5 | SPU_002710 | Lv-Hypp_1407 |
| f5 | SPU_002837 | none |
| f5 | SPU_002934 | Lv-Hypp_1421 |
| f5 | SPU_003004 | Lv-Adra1bL_4 |
| f5 | SPU_003108 | none |
| f5 | SPU_003217 | none |
| f5 | SPU_003297 | Lv-Wscd1 |
| f5 | SPU_003336 | Lv-Mel1aL |
| f5 | SPU_003339 | Lv-C17orf85L |
| f5 | SPU_003476 | Lv-MtL |
| f5 | SPU_003569 | none |
| f5 | SPU_003612 | Lv-Astacin1 |
| f5 | SPU_003671 | Lv-Lrp12 |
| f5 | SPU_004404 | Lv-Slc16a12 |
| f5 | SPU_004429 | Lv-Rtn4ip1-3 |
| f5 | SPU_004887 | Lv-Calr |
| f5 | SPU_004896 | none |
| f5 | SPU_004938 | none |
| f5 | SPU_005036 | none |
| f5 | SPU_005086 | Lv-Hrh3L_2 |
| f5 | SPU_005093 | Lv-Wscd2 |
| f5 | SPU_005279 | Lv-Hypp_1574 |
| f5 | SPU_005385 | Lv-Mt5/MmpL3 |
| f5 | SPU_005423 | Lv-Hao1_4 |
| f5 | SPU_005441 | Lv-Apaf1_1 |
| f5 | SPU_005540 | Lv-Slc7a4 |
| f5 | SPU_005579 | Lv-Hypp_7 |
| f5 | SPU_005713 | Lv-Hspbp1 |
| f5 | SPU_005872 | Lv-Hypp_466 |
| f5 | SPU_006056 | Lv-Plcg_1 |

| | | |
|---|---|---|
| f5 | SPU_006096 | Lv-Mcm2_1 |
| f5 | SPU_006386 | Lv-Cyp2L68 |
| f5 | SPU_006452 | Lv-Slc25a1 |
| f5 | SPU_006705 | Lv-Cyc1 |
| f5 | SPU_007080 | none |
| f5 | SPU_007178 | NA |
| f5 | SPU_007249 | Lv-Slc10a2_2 |
| f5 | SPU_007274 | Lv-Surreal/Gpcr106 |
| f5 | SPU_007337 | Lv-Xpo5 |
| f5 | SPU_007384 | none |
| f5 | SPU_007653 | NA |
| f5 | SPU_007682 | Lv-CbpdEL |
| f5 | SPU_007688 | Lv-Hypp_523 |
| f5 | SPU_007775 | Lv-Gamt |
| f5 | SPU_007885 | Lv-Hypp_1751 |
| f5 | SPU_007990 | NA |
| f5 | SPU_008020 | Lv-Endrvt16 |
| f5 | SPU_008022 | Lv-Frizz4 |
| f5 | SPU_008089 | Lv-Vps8h |
| f5 | SPU_008249 | Lv-Elovl6_6 |
| f5 | SPU_008447 | none |
| f5 | SPU_008504 | Lv-Srcr68 |
| f5 | SPU_008552 | Lv-Ism |
| f5 | SPU_008554 | Lv-Htpalpha |
| f5 | SPU_008560 | Lv-Hsp702A |
| f5 | SPU_008588 | NA |
| f5 | SPU_008740 | none |
| f5 | SPU_008773 | Lv-Mt_2 |
| f5 | SPU_008863 | Lv-Timp3 |
| f5 | SPU_008952 | Lv-Hypp_3004 |
| f5 | SPU_008969 | Lv-Gga3_1 |
| f5 | SPU_009154 | Lv-Hypp_42 |
| f5 | SPU_009356 | Lv-Glyt_3 |
| f5 | SPU_009422 | Lv-Cap1_1 |
| f5 | SPU_009773 | Lv-B4galt5_1 |
| f5 | SPU_009849 | Lv-Sept7 |
| f5 | SPU_009907 | Lv-Galr2L_15 |
| f5 | SPU_009960 | Lv-Hypp_597 |

| | | |
|---|---|---|
| f5 | SPU_009982 | Lv-C4st |
| f5 | SPU_010040 | Lv-Golf |
| f5 | SPU_010218 | Lv-Hypp_1904 |
| f5 | SPU_010236 | Lv-Hypp_612 |
| f5 | SPU_010288 | Lv-Tmem26_3 |
| f5 | SPU_010345 | NA |
| f5 | SPU_010554 | Lv-Sarl/CoAds |
| f5 | SPU_010564 | Lv-Slit(N-terminus) |
| f5 | SPU_010971 | Lv-Kdelc1 |
| f5 | SPU_011085 | none |
| f5 | SPU_011100 | Lv-HdLhd3 |
| f5 | SPU_011114 | none |
| f5 | SPU_011127 | Lv-Ppap2b |
| f5 | SPU_011293 | Lv-Alpl |
| f5 | SPU_011671 | Lv-Phb |
| f5 | SPU_011714 | Lv-Rsk1 |
| f5 | SPU_011755 | Lv-Wnt9 |
| f5 | SPU_011913 | Lv-Adam/Ts2/3L2_1 |
| f5 | SPU_012005 | Lv-Itih |
| f5 | SPU_012083 | Lv-Z235 |
| f5 | SPU_012191 | NA |
| f5 | SPU_012345 | none |
| f5 | SPU_012374 | Lv-Scube |
| f5 | SPU_012468 | Lv-Galnac4s-6stL_4 |
| f5 | SPU_012497 | Lv-Mett11L_1 |
| f5 | SPU_012700 | Lv-Fam57aL |
| f5 | SPU_012760 | none |
| f5 | SPU_012772 | Lv-Klf7 |
| f5 | SPU_012951 | Lv-Pan/Clect |
| f5 | SPU_013046 | Lv-Aadat_1 |
| f5 | SPU_013053 | none |
| f5 | SPU_013502 | Lv-Etfdh_1 |
| f5 | SPU_013694 | none |
| f5 | SPU_013932 | Lv-Egf2l |
| f5 | SPU_014015 | Lv-Hypp_2121 |
| f5 | SPU_014521 | NA |
| f5 | SPU_014702 | Lv-Dync1h1_2 |
| f5 | SPU_014792 | none |

| | | |
|---|---|---|
| f5 | SPU_014802 | Lv-Hlx |
| f5 | SPU_014977 | Lv-Glyt_1 |
| f5 | SPU_015127 | Lv-Sarmr5 |
| f5 | SPU_015163 | Lv-Ccp/Hyr/Lnb/7Tm/Gpcr |
| f5 | SPU_015464 | Lv-Lrr/Gpcr_11 |
| f5 | SPU_015548 | Lv-Srcr127 |
| f5 | SPU_015707 | Lv-Chdh-3 |
| f5 | SPU_015715 | Lv-Hypp_2227 |
| f5 | SPU_015748 | none |
| f5 | SPU_015773 | Lv-Dalrd3 |
| f5 | SPU_015848 | none |
| f5 | SPU_015921 | Lv-Adi |
| f5 | SPU_015952 | Lv-Map215prh |
| f5 | SPU_016004 | Lv-Slc25a46_1 |
| f5 | SPU_016100 | Lv-Ris |
| f5 | SPU_016181 | Lv-Lrp4-3 |
| f5 | SPU_016298 | NA |
| f5 | SPU_016377 | Lv-Fbp1 |
| f5 | SPU_016486 | Lv-Aldoaa |
| f5 | SPU_016664 | Lv-Kif21a |
| f5 | SPU_016754 | Lv-SmG1 |
| f5 | SPU_017150 | none |
| f5 | SPU_017258 | Lv-Slsp_1 |
| f5 | SPU_017406 | Lv-Mco1-2 |
| f5 | SPU_017462 | Lv-Got1 |
| f5 | SPU_017502 | Lv-Arp2/3cs2 |
| f5 | SPU_017675 | Lv-Lbhcor1 |
| f5 | SPU_017698 | Lv-PppL_269 |
| f5 | SPU_017855 | Lv-Hypp_896 |
| f5 | SPU_017903 | Lv-Rnpc1L_1 |
| f5 | SPU_017940 | Lv-Tensin_1 |
| f5 | SPU_018017 | Lv-C10orf59L |
| f5 | SPU_018139 | Lv-Psmb1 |
| f5 | SPU_018181 | none |
| f5 | SPU_018369 | Lv-Arp2/3cs3 |
| f5 | SPU_018444 | Lv-Kcnj12 |
| f5 | SPU_018475 | Lv-Fn3/Igf_5 |
| f5 | SPU_018485 | Lv-Chst11 |

| | | |
|---|---|---|
| f5 | SPU_018518 | Lv-Nmral1 |
| f5 | SPU_018583 | NA |
| f5 | SPU_018617 | Lv-Nploc4 |
| f5 | SPU_018635 | Lv-Alox5_7 |
| f5 | SPU_018659 | Lv-St8sia_1 |
| f5 | SPU_018664 | none |
| f5 | SPU_018964 | Lv-Fk506bp2 |
| f5 | SPU_019034 | Lv-Hypp_925 |
| f5 | SPU_019126 | none |
| f5 | SPU_019233 | Lv-Psmd14 |
| f5 | SPU_019337 | none |
| f5 | SPU_019422 | Lv-Tecp2 |
| f5 | SPU_019450 | Lv-Itih_2 |
| f5 | SPU_019483 | Lv-Lass1 |
| f5 | SPU_019631 | Lv-Hyal1 |
| f5 | SPU_019632 | Lv-Hyalin2 |
| f5 | SPU_019683 | Lv-Slc20a1_2 |
| f5 | SPU_019829 | none |
| f5 | SPU_019892 | Lv-Unc44_243 |
| f5 | SPU_019930 | none |
| f5 | SPU_020379 | Lv-SponL |
| f5 | SPU_020421 | Lv-Gp5L |
| f5 | SPU_020546 | none |
| f5 | SPU_020635 | Lv-Siat6_2 |
| f5 | SPU_020661 | Lv-Hypp_983 |
| f5 | SPU_020777 | Lv-C20orf54_2 |
| f5 | SPU_020826 | Lv-Dnah3_4 |
| f5 | SPU_020902 | Lv-Fbn3L |
| f5 | SPU_021051 | Lv-Prei3l |
| f5 | SPU_021067 | Lv-Fut_2 |
| f5 | SPU_021109 | Lv-Letm1 |
| f5 | SPU_021122 | Lv-Hypp_1011 |
| f5 | SPU_021150 | Lv-Hypp_2540 |
| f5 | SPU_021376 | none |
| f5 | SPU_021586 | Lv-Mpg |
| f5 | SPU_021647 | Lv-Psmd10 |
| f5 | SPU_021824 | none |
| f5 | SPU_021939 | Lv-Hs3st3b1_91 |

| | | |
|---|---|---|
| f5 | SPU_022022 | none |
| f5 | SPU_022046 | Lv-Chst14L_1 |
| f5 | SPU_022049 | Lv-Msx |
| f5 | SPU_022057 | Lv-SemaA |
| f5 | SPU_022066 | Lv-Shmt2_1 |
| f5 | SPU_022073 | Lv-C3orf21_1 |
| f5 | SPU_022074 | Lv-Psmb6 |
| f5 | SPU_022099 | Lv-Rars2 |
| f5 | SPU_022165 | Lv-Hypp_1040 |
| f5 | SPU_022240 | Lv-Sc5dl |
| f5 | SPU_022263 | Lv-Ech1 |
| f5 | SPU_022308 | NA |
| f5 | SPU_022424 | Lv-Srcr192 |
| f5 | SPU_022538 | NA |
| f5 | SPU_022631 | Lv-Hypp_2620 |
| f5 | SPU_022824 | none |
| f5 | SPU_023008 | none |
| f5 | SPU_023052 | none |
| f5 | SPU_023157 | Lv-Hypp_1075 |
| f5 | SPU_023325 | Lv-Slc25a22 |
| f5 | SPU_023536 | NA |
| f5 | SPU_023589 | Lv-Ugt2b34_3 |
| f5 | SPU_023639 | Lv-Hypp_2679 |
| f5 | SPU_023757 | Lv-Ephrin |
| f5 | SPU_023946 | NA |
| f5 | SPU_024042 | none |
| f5 | SPU_024273 | Lv-Lv-NiempC1A |
| f5 | SPU_024308 | Lv-Rfc4 |
| f5 | SPU_024577 | none |
| f5 | SPU_024639 | Lv-Alpl_3 |
| f5 | SPU_024790 | Lv-Slc20a1 |
| f5 | SPU_024946 | Lv-WntA |
| f5 | SPU_025085 | Lv-Slc16a7L_2 |
| f5 | SPU_025294 | Lv-C16orf58 |
| f5 | SPU_025362 | Lv-Coro2b |
| f5 | SPU_025711 | none |
| f5 | SPU_025762 | Lv-Vps16 |
| f5 | SPU_025772 | Lv-Calseq |

| | | |
|---|---|---|
| f5 | SPU_025799 | Lv-Ttc39b_1 |
| f5 | SPU_025910 | Lv-Kiaa0789L |
| f5 | SPU_025917 | Lv-Acs16 |
| f5 | SPU_025976 | none |
| f5 | SPU_026051 | none |
| f5 | SPU_026104 | NA |
| f5 | SPU_026164 | none |
| f5 | SPU_026374 | none |
| f5 | SPU_026430 | Lv-Psma2_1 |
| f5 | SPU_026619 | none |
| f5 | SPU_026711 | Lv-Psma3 |
| f5 | SPU_026917 | Lv-Hypp_1191 |
| f5 | SPU_027046 | Lv-Kirrel2L_4 |
| f5 | SPU_027135 | Lv-Hypp_2889 |
| f5 | SPU_027433 | Lv-Chrna7 |
| f5 | SPU_027487 | Lv-Tll_1 |
| f5 | SPU_027492 | Lv-Cog7 |
| f5 | SPU_027498 | Lv-Galnac4s6st |
| f5 | SPU_027527 | Lv-Hsp901 |
| f5 | SPU_028196 | Lv-Hypp_789 |
| f5 | SPU_028223 | Lv-C6orf167L |
| f5 | SPU_028432 | Lv-Clect_56 |
| f5 | SPU_028433 | Lv-Nlr129 |
| f5 | SPU_028450 | Lv-Hypp_2956 |
| f5 | SPU_028470 | Lv-KmanL |
| f5 | SPU_028608 | none |
| f5 | SPU_028698 | Lv-FoxO_1 |
| f5 | SPU_028735 | Lv-Cct4 |
| f5 | SPU_028748 | Lv-Mt1-4/MmpL7 |
| f5 | SPU_028773 | Lv-Dus4l |
| f5 | SPU_028780 | Lv-Slc23a1_3 |
| f5 | SPU_028834 | none |
| f5 | SPU_028934 | Lv-Cyp2L67 |
| f5 | SPU_030236 | Lv-Cytpd1hc |

| Table S7. k-means clusters for TF-encoding transcripts | | |
|---|---|---|
| **CLUSTER** | **SPU ID** | **NAME** |
| a1 | SPU_017249 | Lv-Arxl |
| a1 | SPU_026905 | Lv-Atf2 |
| a1 | SPU_023867 | Lv-Couptf1 |
| a1 | SPU_002874 | Lv-Ets1/2 |
| a1 | SPU_010168 | Lv-Fog |
| a1 | SPU_011635 | Lv-FoxJ2_1 |
| a1 | SPU_017352 | Lv-Hox1 |
| a1 | SPU_027568 | Lv-Hox3 |
| a1 | SPU_003704 | Lv-Lef1 |
| a1 | SPU_018845 | Lv-Nr1m4 |
| a1 | SPU_016650 | Lv-Par1 |
| a1 | SPU_023739 | Lv-Pbx_1 |
| a1 | SPU_006159 | Lv-Pitx3_1 |
| a1 | SPU_012122 | Lv-Pknox |
| a1 | SPU_014405 | Lv-Pnr |
| a1 | SPU_010438 | Lv-Pou6 |
| a1 | SPU_023090 | Lv-Pric |
| a1 | SPU_018366 | Lv-Rara |
| a1 | SPU_017492 | Lv-Reverb |
| a1 | SPU_027147 | Lv-Riz |
| a1 | SPU_022678 | Lv-Rora |
| a1 | SPU_018392 | Lv-Tbx20_1 |
| a1 | SPU_027218 | Lv-Trx |
| a1 | SPU_015421 | Lv-Trx2 |
| a1 | SPU_003378 | Lv-Z108 |
| a1 | SPU_012949 | Lv-Z122 |
| a1 | SPU_003490 | Lv-Z131 |
| a1 | SPU_006497 | Lv-Z136 |
| a1 | SPU_022841 | Lv-Z142 |
| a1 | SPU_007046 | Lv-Z162 |
| a1 | SPU_019365 | Lv-Z23 |
| a1 | SPU_012307 | Lv-Z322 |
| a1 | SPU_012911 | Lv-Z33 |
| a1 | SPU_009474 | Lv-Z364 |

| | | |
|---|---|---|
| a1 | SPU_019384 | Lv-Z367 |
| a1 | SPU_018909 | Lv-Z39 |
| a1 | SPU_018888 | Lv-Z40 |
| a1 | SPU_014256 | Lv-Z411 |
| a1 | SPU_009198 | Lv-Z479 |
| a1 | SPU_012948 | Lv-Z50 |
| a2 | SPU_006808 | Lv-Af9 |
| a2 | SPU_005358 | Lv-Creb |
| a2 | SPU_012546 | Lv-Ctcf |
| a2 | SPU_017404 | Lv-Nr1h6 |
| a2 | SPU_007611 | Lv-Rfx3 |
| a2 | SPU_021566 | Lv-Shrls |
| a2 | SPU_020722 | Lv-Smad1/5/8_1 |
| a2 | SPU_017642 | Lv-Smad2/3 |
| a2 | SPU_022820 | Lv-Soxb1 |
| a2 | SPU_014249 | Lv-Trh |
| a2 | SPU_000016 | Lv-Unk_48 |
| a2 | SPU_006865 | Lv-Unk_50 |
| a2 | SPU_022487 | Lv-Z106 |
| a2 | SPU_012198 | Lv-Z124 |
| a2 | SPU_013307 | Lv-Z124_1 |
| a2 | SPU_027709 | Lv-Z125 |
| a2 | SPU_006427 | Lv-Z126 |
| a2 | SPU_007360 | Lv-Z129 |
| a2 | SPU_007361 | Lv-Z144 |
| a2 | SPU_028222 | Lv-Z146 |
| a2 | SPU_015688 | Lv-Z154 |
| a2 | SPU_018075 | Lv-Z160 |
| a2 | SPU_012645 | Lv-Z166 |
| a2 | SPU_019652 | Lv-Z189 |
| a2 | SPU_013611 | Lv-Z192 |
| a2 | SPU_016149 | Lv-Z265 |
| a2 | SPU_018373 | Lv-Z308 |
| a2 | SPU_020481 | Lv-Z310 |
| a2 | SPU_009053 | Lv-Z316 |
| a2 | SPU_018960 | Lv-Z317 |
| a2 | SPU_018680 | Lv-Z338 |
| a2 | SPU_014045 | Lv-Z35 |

| | | |
|---|---|---|
| a2 | SPU_005559 | Lv-Z356 |
| a2 | SPU_027912 | Lv-Z365 |
| a2 | SPU_016518 | Lv-Z376 |
| a2 | SPU_003773 | Lv-Z392 |
| a2 | SPU_017811 | Lv-Z401 |
| a2 | SPU_025849 | Lv-Z408_1 |
| a2 | SPU_011226 | Lv-Z415 |
| a2 | SPU_008563 | Lv-Z42 |
| a2 | SPU_012955 | Lv-Z422 |
| a2 | SPU_014817 | Lv-Z423 |
| a2 | SPU_019824 | Lv-Z425 |
| a2 | SPU_026385 | Lv-Z427 |
| a2 | SPU_021851 | Lv-Z434 |
| a2 | SPU_011502 | Lv-Z437 |
| a2 | SPU_005229 | Lv-Z440 |
| a2 | SPU_010458 | Lv-Z443 |
| a2 | SPU_026209 | Lv-Z445 |
| a2 | SPU_010922 | Lv-Z458 |
| a2 | SPU_020726 | Lv-Z472 |
| a2 | SPU_002140 | Lv-Z475 |
| a2 | SPU_014686 | Lv-Z478_1 |
| a2 | SPU_013904 | Lv-Z484 |
| a2 | SPU_002988 | Lv-Z49 |
| a2 | SPU_014683 | Lv-Z492 |
| a2 | SPU_012111 | Lv-Z53 |
| a2 | SPU_001483 | Lv-Z59 |
| a2 | SPU_007108 | Lv-Z67 |
| a2 | SPU_016250 | Lv-Z70 |
| a2 | SPU_026510 | Lv-Z74 |
| a2 | SPU_002554 | Lv-Z75 |
| a2 | SPU_014685 | Lv-Z80 |
| a2 | SPU_012633 | none |
| a2 | SPU_021164 | none |
| a3 | SPU_019129 | Lv-Eyg |
| a3 | SPU_000388 | Lv-Hox11/13c |
| a3 | SPU_008752 | Lv-Nfe2_1 |
| a3 | SPU_017380 | Lv-Six4 |
| a3 | SPU_011761 | Lv-Z295 |

| | | |
|---|---|---|
| a3 | SPU_021205 | Lv-Z357 |
| a3 | SPU_002708 | Lv-Z382 |
| a3 | SPU_006058 | Lv-Z464 |
| b | SPU_006312 | Lv-Dp1 |
| b | SPU_024903 | Lv-Ese |
| b | SPU_012253 | Lv-Eve |
| b | SPU_025590 | Lv-FoxM |
| b | SPU_019002 | Lv-FoxQ2_1 |
| b | SPU_013305 | Lv-Grf |
| b | SPU_002631 | Lv-Hox11/13b |
| b | SPU_004148 | Lv-Krl |
| b | SPU_006991 | Lv-Lim1 |
| b | SPU_012448 | Lv-Ovo |
| b | SPU_008112 | Lv-Phb1 |
| b | SPU_008117 | Lv-Shr2 |
| b | SPU_009427 | Lv-Z225 |
| b | SPU_021947 | Lv-Z474 |
| b | SPU_020110 | Lv-Z474_1 |
| b | SPU_012913 | Lv-Z92 |
| b | SPU_028583 | Lv-Zic |
| b | SPU_006421 | none |
| c1 | SPU_022816 | Lv-Alx4 |
| c1 | SPU_003681 | Lv-Atol2 |
| c1 | SPU_003548 | Lv-E78a |
| c1 | SPU_018797 | Lv-Evi1 |
| c1 | SPU_014177 | Lv-Lbx |
| c1 | SPU_006232 | Lv-MyoD2 |
| c1 | SPU_022573 | Lv-Nk7 |
| c1 | SPU_018951 | Lv-Prx1 |
| c2 | SPU_022554 | Lv-Ac/Sc3 |
| c2 | SPU_000990 | Lv-Atol1 |
| c2 | SPU_020975 | Lv-Engrailed |
| c2 | SPU_021309 | Lv-Hox8 |
| c2 | SPU_007852 | Lv-Runt2 |
| c2 | SPU_013962 | Lv-Sim_1 |
| d1 | SPU_016685 | Lv-Ap2 |
| d1 | SPU_024715 | Lv-Cdx |
| d1 | SPU_019656 | Lv-Cdxl |

| | | |
|---|---|---|
| d1 | SPU_027648 | Lv-FoxD_1 |
| d1 | SPU_023894 | Lv-FoxI |
| d1 | SPU_015719 | Lv-FoxL1 |
| d1 | SPU_019345 | Lv-FoxQ1 |
| d1 | SPU_007599 | Lv-Glass2 |
| d1 | SPU_002816 | Lv-Hb9 |
| d1 | SPU_002632 | Lv-Hox11/13a |
| d1 | SPU_011202 | Lv-Meis |
| d1 | SPU_027623 | Lv-Mist |
| d1 | SPU_012008 | Lv-Myor2 |
| d1 | SPU_009231 | Lv-Ncsl |
| d1 | SPU_007147 | Lv-Ngn |
| d1 | SPU_012699 | Lv-Nkx6-1 |
| d1 | SPU_014401 | Lv-NtL |
| d1 | SPU_019290 | Lv-Otp |
| d1 | SPU_000276 | Lv-PaxC |
| d1 | SPU_013464 | Lv-Phox2 |
| d1 | SPU_004599 | Lv-Pitx2 |
| d1 | SPU_025632 | Lv-Pou4f2 |
| d1 | SPU_016786 | Lv-Rhox3 |
| d1 | SPU_002448 | Lv-Sage |
| d1 | SPU_018393 | Lv-Scratch |
| d1 | SPU_011280 | Lv-Sp8 |
| d1 | SPU_020346 | Lv-Tbx6 |
| d1 | SPU_023522 | Lv-Z66 |
| d2 | SPU_014164 | Lv-Barh1 |
| d2 | SPU_024486 | Lv-Dsf |
| d2 | SPU_021173 | Lv-Fos |
| d2 | SPU_018744 | Lv-GlisB |
| d2 | SPU_002630 | Lv-Hox8_1 |
| d2 | SPU_010404 | Lv-Irf |
| d2 | SPU_011246 | Lv-IrxB |
| d2 | SPU_003102 | Lv-Jun |
| d2 | SPU_017375 | Lv-Nr2e6 |
| d2 | SPU_007242 | Lv-Rough |
| d2 | SPU_016881 | Lv-SoxE |
| d2 | SPU_014170 | Lv-SoxF |
| d2 | SPU_006150 | Lv-Tbx1 |

| | | |
|---|---|---|
| d2 | SPU_008703 | Lv-Xbp1 |
| d2 | SPU_010634 | Lv-Z447 |
| e01 | SPU_021491 | Lv-Arxl2 |
| e01 | SPU_027935 | Lv-Bmal |
| e01 | SPU_028061 | Lv-Dac |
| e01 | SPU_023366 | Lv-Enz1 |
| e01 | SPU_027446 | Lv-Enz2 |
| e01 | SPU_021123 | Lv-L3mbt |
| e01 | SPU_013689 | Lv-L3mbt_1 |
| e01 | SPU_013569 | Lv-Lmo2t |
| e01 | SPU_016168 | Lv-Mef2 |
| e01 | SPU_000861 | Lv-Myb |
| e01 | SPU_009262 | Lv-Oct1-2 |
| e01 | SPU_004217 | Lv-SoxD1 |
| e01 | SPU_011080 | Lv-SoxH |
| e01 | SPU_009831 | Lv-Sp2 |
| e01 | SPU_025584 | Lv-Tbr |
| e01 | SPU_028479 | Lv-Tel |
| e01 | SPU_014332 | Lv-Usf |
| e01 | SPU_019407 | Lv-Z101 |
| e01 | SPU_010426 | Lv-Z128 |
| e01 | SPU_011189 | Lv-Z141 |
| e01 | SPU_018673 | Lv-Z17 |
| e01 | SPU_001937 | Lv-Z19 |
| e01 | SPU_004806 | Lv-Z197 |
| e01 | SPU_009646 | Lv-Z223 |
| e01 | SPU_004802 | Lv-Z232 |
| e01 | SPU_007440 | Lv-Z282 |
| e01 | SPU_004807 | Lv-Z32 |
| e01 | SPU_005035 | Lv-Z339 |
| e01 | SPU_012583 | Lv-Z383 |
| e01 | SPU_020563 | Lv-Z395 |
| e01 | SPU_015071 | Lv-Z398 |
| e01 | SPU_006871 | Lv-Z45 |
| e01 | SPU_012914 | Lv-Z62 |
| e01 | SPU_006063 | Lv-Z69 |
| e01 | SPU_011720 | Lv-Z90 |
| e02 | SPU_004702 | Lv-Ebf3 |

| | | |
|---|---|---|
| e02 | SPU_011348 | Lv-Fxr |
| e02 | SPU_018692 | Lv-Groucho |
| e02 | SPU_013436 | Lv-Gsx |
| e02 | SPU_017470 | Lv-Klf3/8/12 |
| e02 | SPU_000948 | Lv-Lass6 |
| e02 | SPU_005787 | Lv-Mlx |
| e02 | SPU_006683 | Lv-Pax1-9 |
| e02 | SPU_019332 | Lv-Ppar1 |
| e02 | SPU_010383 | Lv-Z137 |
| e02 | SPU_020846 | Lv-Z233 |
| e02 | SPU_022800 | Lv-Z236 |
| e02 | SPU_015682 | Lv-Z417_1 |
| e02 | SPU_011583 | Lv-Z421 |
| e02 | SPU_003685 | Lv-Z450 |
| e02 | SPU_014197 | Lv-Z55 |
| e03 | SPU_011315 | Lv-Ato |
| e03 | SPU_006803 | Lv-Creb3l3 |
| e03 | SPU_027969 | Lv-FoxJ1 |
| e03 | SPU_014528 | Lv-Giant |
| e03 | SPU_017627 | Lv-GliA |
| e03 | SPU_001975 | Lv-Lhx3-4 |
| e03 | SPU_025486 | Lv-Mox |
| e03 | SPU_024918 | Lv-NeuroD1 |
| e03 | SPU_023339 | Lv-Nfia |
| e03 | SPU_024307 | Lv-NfIL3 |
| e03 | SPU_005472 | Lv-Nkx2.5 |
| e03 | SPU_015456 | Lv-Nr1h6b |
| e03 | SPU_017491 | Lv-Nr1m1 |
| e03 | SPU_013178 | Lv-Nr1m3 |
| e03 | SPU_021289 | Lv-Ppar2 |
| e03 | SPU_027362 | Lv-Top1L |
| e03 | SPU_017656 | Lv-Z10 |
| e03 | SPU_000437 | Lv-Z211 |
| e03 | SPU_015495 | Lv-Z224 |
| e03 | SPU_026110 | Lv-Z262 |
| e04 | SPU_001519 | Lv-Barx |
| e04 | SPU_017287 | Lv-Hand |
| e05 | SPU_025302 | Lv-Alx1 |

| | | |
|---|---|---|
| e05 | SPU_023037 | Lv-Bbx |
| e05 | SPU_026633 | Lv-Dmtf |
| e05 | SPU_015358 | Lv-Egr |
| e05 | SPU_017636 | Lv-Hypp_886 |
| e05 | SPU_004844 | Lv-Lztf1_1 |
| e05 | SPU_022163 | Lv-Max |
| e05 | SPU_026465 | Lv-Mll3 |
| e05 | SPU_026205 | Lv-Mnt |
| e05 | SPU_013134 | Lv-Nr2C |
| e05 | SPU_009413 | Lv-Nxf |
| e05 | SPU_010424 | Lv-Otx |
| e05 | SPU_017725 | Lv-P3a2 |
| e05 | SPU_018351 | Lv-PaxB |
| e05 | SPU_025113 | Lv-Soxb2 |
| e05 | SPU_014793 | Lv-Spalt |
| e05 | SPU_027774 | Lv-Srf |
| e05 | SPU_021210 | Lv-Tead4 |
| e05 | SPU_010295 | Lv-Yyi |
| e05 | SPU_008377 | Lv-Z105 |
| e05 | SPU_028045 | Lv-Z110 |
| e05 | SPU_019435 | Lv-Z114 |
| e05 | SPU_011611 | Lv-Z134 |
| e05 | SPU_021382 | Lv-Z202 |
| e05 | SPU_012724 | Lv-Z218 |
| e05 | SPU_008358 | Lv-Z229 |
| e05 | SPU_011687 | Lv-Z258 |
| e05 | SPU_009553 | Lv-Z263 |
| e05 | SPU_017846 | Lv-Z313 |
| e05 | SPU_025959 | Lv-Z314 |
| e05 | SPU_021757 | Lv-Z347 |
| e05 | SPU_028753 | Lv-Z359 |
| e05 | SPU_014643 | Lv-Z372 |
| e05 | SPU_022024 | Lv-Z410 |
| e05 | SPU_012632 | Lv-Z417 |
| e05 | SPU_024745 | Lv-Z52 |
| e05 | SPU_021383 | Lv-Z56 |
| e05 | SPU_022892 | Lv-Z61 |
| e05 | SPU_006064 | Lv-Z79 |

| | | |
|---|---|---|
| e05 | SPU_013576 | Lv-Z96 |
| e06 | SPU_007749 | Lv-Atf6b |
| e06 | SPU_027235 | Lv-Blimp1 |
| e06 | SPU_020451 | Lv-Bra_1 |
| e06 | SPU_012838 | Lv-Creb3l1 |
| e06 | SPU_006753 | Lv-E2f3 |
| e06 | SPU_028827 | Lv-E2f4 |
| e06 | SPU_002592 | Lv-Emx |
| e06 | SPU_018483 | Lv-Erg |
| e06 | SPU_015243 | Lv-FoxN2/3 |
| e06 | SPU_006462 | Lv-Gcm |
| e06 | SPU_023177 | Lv-Hbn |
| e06 | SPU_021608 | Lv-HesC |
| e06 | SPU_015374 | Lv-Id |
| e06 | SPU_020311 | Lv-Klf2/4 |
| e06 | SPU_003705 | Lv-Mta1_1 |
| e06 | SPU_003166 | Lv-Myc |
| e06 | SPU_000756 | Lv-Nk2-2 |
| e06 | SPU_002129 | Lv-Not |
| e06 | SPU_026763 | Lv-Scml1 |
| e06 | SPU_001998 | Lv-Smad6/7 |
| e06 | SPU_002603 | Lv-SoxC |
| e06 | SPU_023386 | Lv-Tbx2/3 |
| e06 | SPU_000440 | Lv-Z115 |
| e06 | SPU_017427 | Lv-Z155 |
| e06 | SPU_013861 | Lv-Z190 |
| e06 | SPU_015800 | Lv-Z246 |
| e06 | SPU_017848 | Lv-Z313_2 |
| e06 | SPU_027708 | Lv-Z491 |
| e07 | SPU_018954 | Lv-Awh |
| e07 | SPU_014809 | Lv-FoxL2 |
| e07 | SPU_013843 | Lv-FtzF |
| e07 | SPU_005169 | Lv-Hox5 |
| e07 | SPU_015983 | Lv-MyoD3 |
| e07 | SPU_030060 | Lv-Pu1 |
| e07 | SPU_013119 | Lv-Sage_1 |
| e07 | SPU_015767 | Lv-Z57 |
| e07 | SPU_018850 | Lv-Z63 |

| | | |
|---|---|---|
| e07 | SPU_021843 | Lv-Z64 |
| e08 | SPU_022817 | Lv-Alx1_1 |
| e08 | SPU_012586 | Lv-Fax1 |
| e08 | SPU_027015 | Lv-GataC |
| e08 | SPU_016445 | Lv-Myor4 |
| e08 | SPU_028093 | Lv-Scl |
| e08 | SPU_019268 | Lv-Shox |
| e08 | SPU_017947 | Lv-Z116 |
| e08 | SPU_007636 | Lv-Z306 |
| e09 | SPU_017348 | Lv-Atbf1 |
| e09 | SPU_028698 | Lv-FoxO_1 |
| e09 | SPU_010403 | Lv-FoxY |
| e09 | SPU_026292 | Lv-Glis1_1 |
| e09 | SPU_014802 | Lv-Hlx |
| e09 | SPU_016449 | Lv-Hnf6 |
| e09 | SPU_002634 | Lv-Hox7 |
| e09 | SPU_026877 | Lv-Irf4 |
| e09 | SPU_010351 | Lv-IrxA |
| e09 | SPU_022049 | Lv-Msx |
| e09 | SPU_014539 | Lv-Pax2-5-8 |
| e09 | SPU_002677 | Lv-Ptf1a |
| e09 | SPU_028422 | Lv-Rxr |
| e09 | SPU_017379 | Lv-Six1/2 |
| e09 | SPU_001113 | Lv-Sna |
| e09 | SPU_018126 | Lv-Tgif |
| e09 | SPU_027487 | Lv-Tll_1 |
| e09 | SPU_001739 | Lv-Unc4-1 |
| e09 | SPU_015362 | Lv-Z167 |
| e10 | SPU_003920 | Lv-Barx_1 |
| e10 | SPU_000485 | Lv-Chx10 |
| e10 | SPU_021192 | Lv-Hnf4 |
| e10 | SPU_021119 | Lv-MyoD |
| e10 | SPU_014461 | Lv-Pitx1 |
| e10 | SPU_015640 | Lv-ScratchX |
| e10 | SPU_027753 | Lv-Z347_1 |
| e11 | SPU_019338 | Lv-Alx |
| e11 | SPU_016443 | Lv-Brn1-2-4 |
| e11 | SPU_024139 | Lv-FoxC |

| | | |
|---|---|---|
| e11 | SPU_000975 | Lv-FoxF |
| e11 | SPU_015712 | Lv-Hey4 |
| e11 | SPU_023730 | Lv-Isl |
| e11 | SPU_026418 | Lv-Klf11 |
| e11 | SPU_015772 | Lv-Klf15 |
| e11 | SPU_012772 | Lv-Klf7 |
| e11 | SPU_014157 | Lv-Lmx1 |
| e11 | SPU_026099 | Lv-Lox |
| e11 | SPU_006583 | Lv-Mad |
| e11 | SPU_011297 | Lv-Mbx1 |
| e11 | SPU_020565 | Lv-Msxl |
| e11 | SPU_024877 | Lv-Osr |
| e11 | SPU_016523 | Lv-Rar |
| e11 | SPU_007253 | Lv-Srebp |
| e11 | SPU_012050 | Lv-Z11 |
| e11 | SPU_006637 | Lv-Z193 |
| e11 | SPU_012083 | Lv-Z235 |
| f1 | SPU_000129 | Lv-Arnt |
| f1 | SPU_021689 | Lv-Eif2Ba |
| f1 | SPU_004723 | Lv-Err |
| f1 | SPU_025010 | Lv-FoxK |
| f1 | SPU_010635 | Lv-Gatae |
| f1 | SPU_023727 | Lv-Klf13 |
| f1 | SPU_025126 | Lv-Large_1 |
| f1 | SPU_019586 | Lv-Lmo4 |
| f1 | SPU_007981 | Lv-Lmpt |
| f1 | SPU_005667 | Lv-Mcm10 |
| f1 | SPU_011576 | Lv-Nr1m2 |
| f1 | SPU_014576 | Lv-Pea |
| f1 | SPU_015984 | Lv-Prox1 |
| f1 | SPU_016617 | Lv-Tulp4L |
| f1 | SPU_009225 | Lv-Unk_49 |
| f1 | SPU_028301 | Lv-Z163 |
| f1 | SPU_013407 | Lv-Z175 |
| f1 | SPU_011260 | Lv-Z181 |
| f1 | SPU_017682 | Lv-Z203 |
| f1 | SPU_007169 | Lv-Z219 |
| f1 | SPU_010914 | Lv-Z25_1 |

| | | |
|---|---|---|
| f1 | SPU_003848 | Lv-Z298 |
| f1 | SPU_019653 | Lv-Z309 |
| f1 | SPU_024953 | Lv-Z34 |
| f1 | SPU_001648 | Lv-Z355 |
| f1 | SPU_007957 | Lv-Z370 |
| f1 | SPU_003239 | Lv-Z373 |
| f1 | SPU_026390 | Lv-Z426 |
| f1 | SPU_028679 | Lv-Z481 |
| f1 | SPU_014684 | Lv-Z94 |
| f2 | SPU_019915 | Lv-Apa |
| f2 | SPU_018423 | Lv-Ash2 |
| f2 | SPU_001262 | Lv-Birc6 |
| f2 | SPU_017407 | Lv-Clock |
| f2 | SPU_014836 | Lv-Cp2 |
| f2 | SPU_003595 | Lv-Cutl |
| f2 | SPU_016343 | Lv-E12 |
| f2 | SPU_014628 | Lv-Erf |
| f2 | SPU_017579 | Lv-GlisC |
| f2 | SPU_006813 | Lv-Hairy |
| f2 | SPU_026962 | Lv-Ldb2 |
| f2 | SPU_008845 | Lv-Mlx/Ip |
| f2 | SPU_009642 | Lv-Rreb1 |
| f2 | SPU_004287 | Lv-Smad4 |
| f2 | SPU_003470 | Lv-Spz12 |
| f2 | SPU_018861 | Lv-Thrb |
| f2 | SPU_013406 | Lv-Z118 |
| f2 | SPU_019959 | Lv-Z14 |
| f2 | SPU_012912 | Lv-Z150 |
| f2 | SPU_009847 | Lv-Z156 |
| f2 | SPU_017750 | Lv-Z159 |
| f2 | SPU_007190 | Lv-Z185 |
| f2 | SPU_012315 | Lv-Z215 |
| f2 | SPU_015751 | Lv-Z220 |
| f2 | SPU_009685 | Lv-Z227 |
| f2 | SPU_011001 | Lv-Z257 |
| f2 | SPU_024748 | Lv-Z276 |
| f2 | SPU_014923 | Lv-Z28 |
| f2 | SPU_027918 | Lv-Z288 |

| | | |
|---|---|---|
| f2 | SPU_000017 | Lv-Z307 |
| f2 | SPU_015683 | Lv-Z326 |
| f2 | SPU_013415 | Lv-Z328 |
| f2 | SPU_027919 | Lv-Z344 |
| f2 | SPU_007011 | Lv-Z346 |
| f2 | SPU_002586 | Lv-Z366 |
| f2 | SPU_011380 | Lv-Z368 |
| f2 | SPU_019508 | Lv-Z371 |
| f2 | SPU_013902 | Lv-Z386 |
| f2 | SPU_002707 | Lv-Z388 |
| f2 | SPU_019651 | Lv-Z391 |
| f2 | SPU_025850 | Lv-Z408 |
| f2 | SPU_011879 | Lv-Z412 |
| f2 | SPU_010924 | Lv-Z416 |
| f2 | SPU_023510 | Lv-Z444 |
| f2 | SPU_000578 | Lv-Z460 |
| f2 | SPU_002008 | Lv-Z480 |
| f2 | SPU_002009 | Lv-Z480_1 |
| f2 | SPU_014687 | Lv-Z494 |
| f2 | SPU_017441 | Lv-Z65 |
| f2 | SPU_020480 | Lv-Z77 |
| f2 | SPU_005071 | Lv-Z87 |
| f2 | SPU_000540 | Lv-Z98 |
| f3 | SPU_026767 | Lv-Abcg10 |
| f3 | SPU_025292 | Lv-Cic |
| f3 | SPU_020123 | Lv-ElfB |
| f3 | SPU_021172 | Lv-Fra2 |
| f3 | SPU_025492 | Lv-Gbx |
| f3 | SPU_009465 | Lv-Hey |
| f3 | SPU_027215 | Lv-Hhex |
| f3 | SPU_004414 | Lv-Hlf |
| f3 | SPU_005171 | Lv-Hox6 |
| f3 | SPU_008177 | Lv-Nfkb |
| f3 | SPU_000757 | Lv-Nkx2.1 |
| f3 | SPU_013047 | Lv-Nkx3-2 |
| f3 | SPU_004526 | Lv-Nr1h6c |
| f3 | SPU_028255 | Lv-Nr1x |
| f3 | SPU_027334 | Lv-PaxA |

| | | |
|---|---|---|
| f3 | SPU_024093 | Lv-Phb2 |
| f3 | SPU_018908 | Lv-Six3 |
| f3 | SPU_022242 | Lv-Smad_ip |
| f3 | SPU_024189 | Lv-Sp5 |
| f3 | SPU_002674 | Lv-TfIIIa |
| f3 | SPU_023320 | Lv-Trpc5_2 |
| f3 | SPU_016490 | Lv-Z145 |
| f3 | SPU_025125 | Lv-Z147 |
| f3 | SPU_017849 | Lv-Z313_3 |
| f3 | SPU_002090 | Lv-Z342 |
| f4 | SPU_028148 | Lv-Ac/Sc |
| f4 | SPU_025482 | Lv-Ash1 |
| f4 | SPU_002815 | Lv-Dlx |
| f4 | SPU_006676 | Lv-FoxA |
| f4 | SPU_004551 | Lv-FoxB |
| f4 | SPU_009771 | Lv-FoxG |
| f4 | SPU_015982 | Lv-Gsc |
| f4 | SPU_006814 | Lv-Hairy2/4 |
| f4 | SPU_002633 | Lv-Hox9/10 |
| f4 | SPU_027472 | Lv-Ia1 |
| f4 | SPU_004021 | Lv-Limc1 |
| f4 | SPU_025888 | Lv-Maf |
| f4 | SPU_008175 | Lv-Mitf |
| f4 | SPU_011174 | Lv-Nfe2 |
| f4 | SPU_012491 | Lv-Nk1 |
| f4 | SPU_000255 | Lv-Nurr1 |
| f4 | SPU_006917 | Lv-Runt1 |
| f4 | SPU_018246 | Lv-Smad6/7_1 |
| f4 | SPU_019089 | Lv-Z133 |
| f4 | SPU_000603 | Lv-Z135 |
| f4 | SPU_006484 | Lv-Z204 |
